# Feedback regulation of RNase E during UV-stress response in the cyanobacterium *Synechocystis* sp. PCC 6803

**DOI:** 10.1101/2022.02.07.478427

**Authors:** Satoru Watanabe, Damir Stazic, Jens Georg, Shota Ohtake, Megumi Numakura, Munehiko Asayama, Taku Chibazakura, Annegret Wilde, Claudia Steglich, Wolfgang R. Hess

## Abstract

Endoribonucleases govern the maturation and degradation of RNA and are indispensable in the posttranscriptional regulation of gene expression. A key endoribonuclease in many bacteria is RNase E. To ensure an appropriate supply of RNase E, some bacteria, such as *E. coli,* have evolved tightly functioning feedback regulation of RNase E that is mediated in *cis* by the *rne* 5′-untranslated region (5′ UTR); however, the mechanisms involved in the control of RNase E in other bacteria largely remain unknown. Cyanobacteria rely on solar light as an energy source for photosynthesis, despite the inherent ultraviolet (UV) irradiation. Here, we investigated the global gene expression response in the cyanobacterium *Synechocystis* sp. PCC 6803 after exposure to UV light and discovered a unique response of RNase E: a rapidly increasing enzymatic activity, although the stability of the protein was decreased. In parallel, we observed an increased accumulation of full-length *rne* mRNA that was caused by the stabilization of its 5′ UTR and suppression of premature transcriptional termination but not by an increased transcription rate. Mapping of RNA 3′ ends and *in vitro* cleavage assays revealed that RNase E cleaves within a stretch of six consecutive uridine residues within the *rne* 5′ UTR, indicating autoregulation *via* its own 5′ UTR. These observations imply that RNase E in cyanobacteria contributes substantially to reshaping the transcriptome during the UV stress response and that its required activity level is maintained despite enhanced turnover of the protein by posttranscriptional feedback regulation.

## Introduction

Messenger RNA (mRNA) degradation plays a key and universal role in the posttranscriptional control of gene expression. In both prokaryotic and eukaryotic organisms, mRNA lifetimes can vary by up to two orders of magnitude, with proportionate effects on protein production (1). In *E. coli,* mRNA decay mechanisms involving the sequential action of endonucleases and 3′ exonucleases have been well studied (2, 3). The endonuclease that is most important for mRNA turnover in *E. coli* is endoribonuclease (RNase) E. In addition to its function in the degradation of most mRNAs, RNase E also participates in rRNA and tRNA maturation (4). *E. coli* RNase E cuts RNA within single-stranded regions that are AU-rich, although the presence of a guanosine residue two nucleotides upstream of the cleavage site increases reactivity (5–8). The recognized core motif of RNase E of *Salmonella typhimurium*, a close relative of *E. coli*, has been specified as ‘‘RN↓WUU’’; the enzyme shows a marked preference for uridine at position +2 after the cleavage site (indicated by ↓) (9). RNase E is also the key enzyme in the interactions between bacterial regulatory small RNAs (sRNAs) and their targets, to which it can be recruited upon sRNA binding (10) or excluded from accessing possible cleavage sites (11, 12). In view of its many crucial biological functions, it is not surprising that RNase E is an essential enzyme in *E. coli* and that imbalanced production of RNase E can impede cell growth (13–15). To ensure a steady supply of RNase E, *E. coli* and related bacteria have evolved a homeostatic mechanism for tightly regulating its synthesis in which the level and rate of decay of *rne* mRNA are modulated in response to changes in cellular RNase E activity (16). The feedback regulation of RNase E is mediated in *cis* by the *rne* 5′ UTR (16, 17). Compared to the 5′ UTRs of other genes, the *E. coli rne* 5′ UTR is with a length of 361 nucleotides very long (18). Through its cleavage by RNase E and its expediting of cleavage elsewhere within the *rne* transcript, this long 5′ UTR is critically involved in the control of RNase E synthesis (19).

Cyanobacteria are the only bacteria that perform oxygenic photosynthesis similar to the photosynthesis that occurs in plant and algal chloroplasts. Cyanobacterial RNase E proteins are smaller than their homologs in most other bacteria: the RNase E of the cyanobacterium *Synechocystis* sp. PCC 6803 (hereafter *Synechocystis* 6803) contains 674 amino acid residues, whereas RNase E of *E. coli* and *Pseudomonas* spp. contain more than 1,000 residues. Nevertheless, the compact form of RNase E is highly conserved among cyanobacteria and occurs in plant and algal chloroplasts, where import of the enzyme is mediated via an additional N-terminal targeting sequence (20). In the cyanobacterium *Synechocystis* 6803, complete genetic disruption of the *rne* gene (gene *slr1129*) failed to segregate completely into a homozygous mutant line (21, 22), and partial disruption of RNase E led to severe growth inhibition and affected the expression of a large number of genes (22), indicating that RNase E is essential. There is circumstantial evidence for feedback regulation of RNase E in the cyanobacteria *Synechocystis* 6803 and *Prochlorococcus marinus* MED4 (22, 23). This feedback regulation appears to be mediated via the *rne* 5′ UTR. Comprehensive transcriptome analysis revealed the transcriptional start sites (TSSs) of *rne* and indicated that, compared to *E. coli* (361 nt), even longer 5′ UTRs (458-622 nt) are typically associated with the *rne* gene in several different cyanobacteria, such as *Synechocystis* sp., *Anabaena* sp. and *Synechococcus elongatus* (24–27) (Supplementary Figure S1A). However, the molecular details of such putative regulation remain largely unknown.

RNase E in *Synechocystis* 6803 participates in the posttranscriptional regulation of *psbA2*, which encodes the photosystem (PS) II reaction center D1 protein. During darkness, when *psbA2* expression is not required, RNase E cleaves at two tightly spaced sites, the AU box and within the ribosome binding site, both of which are located in the 5′ UTR of the *psbA2* transcript (28, 29). However, these sites are not cleaved when the cells are cultivated in the light and *psbA2* expression is high (28, 29). PsbA2R and PsbA3R, two cis-encoded antisense RNAs (asRNAs), are involved in the stabilization of *psbA2* and *psbA3* transcripts in the light, and this protective effect has physiological relevance (30). Another function relevant to the proper functioning of the photosynthetic apparatus is the recruitment of RNase E upon binding of the sRNA PsrR1 to a cleavage site located 4 nucleotides downstream of the start codon within the *psaLI* dicistronic mRNA encoding two PS I proteins (31).

RNase E has also been shown to be involved in the processing of polycistronic transcripts. In *Synechocystis* 6803, the DEAD-box RNA helicase CrhR responds to cold stress (32, 33). The *crhR* gene forms an operon with *rimO*, which encodes a methylthiotransferase. *In vitro* cleavage experiments suggested that RNase E cleaves the polycistronic *rimO-crhR* transcript and that it is required for the autoregulation of CrhR expression (34). Another critical role of RNase E in *Synechocystis* 6803 is in the clustered regularly interspaced short palindromic repeat (CRISPR)-Cas defense mechanism, where it is involved in the maturation of CRISPR-derived RNAs (crRNAs) (35).

RNase E thus appears to play a pivotal role in cyanobacteria. Indeed, the recent mapping of RNase E-dependent cleavage sites in *Synechocystis* 6803 after transient inactivation of RNase E by temperature shift (TIER-seq) yielded 1,472 such sites (36). The dominating cleavage signature was found to consist of an adenine at the -3 position and a uridine at the +2 position within a single-stranded segment of the RNA (36).

As an energy source for their photosynthetic lifestyle, cyanobacteria rely on solar energy. Under natural conditions, light intensity varies frequently and substantially, as does the inherent fraction of ultraviolet (UV) light. Therefore, cyanobacteria must employ specific mechanisms to cope with UV light-induced damage to biomolecules. Since nucleic acid molecules (not only DNA but also RNA) are primary targets of UV radiation (37) and damaged RNA may perturb cellular gene expression (38), it is reasonable that RNase E becomes activated after UV treatment. In *Synechocystis* 6803, *rne* transcript levels increased approximately two-to-threefold after UV treatment (39), but this also occurred following sulfur starvation (40) or redox stress (41). However, neither the functional relevance of the enhanced expression of *rne* under these conditions nor the mechanisms underlying it have been elucidated.

Here, we demonstrate that the UV stress response in *Synechocystis* 6803 involves dynamic changes in the transcriptome and triggers feedback regulation of RNase E. After UV irradiation, full-length forms of *rne* mRNA significantly accumulated; this was caused by selective stabilization of its 5′ UTR and suppressed premature termination of *rne* transcription, while an increased transcription rate was not involved. In parallel, the activity of RNase E increased while the amount of RNase E protein remained constant, although RNase E protein stability decreased. Mapping of RNA 3′ ends and *in vitro* cleavage assays indicated that *Synechocystis* 6803 RNase E cleaves close to and within a U-rich region in the 5′ UTR of *rne* mRNA. Our findings suggest that RNase E is required for reshaping of the transcriptome during the UV stress response in cyanobacteria, that its required level of activity is ensured despite enhanced turnover of the protein, and that the mechanism underlying this involves a feedback mechanism acting on a U-rich element within the *rne* 5′ UTR.

## Materials and Methods

### Bacterial strains and growth conditions

*Synechocystis* sp. PCC 6803 PCC-M strain (42) was grown photoautotrophically (40 µmol photons·m^-2^·s^-1^) at 30 °C in BG-11 medium (43) supplemented with 20 mM HEPES-NaOH (pH 7.5).

### UV irradiation and viability assay

For viability testing, exponentially growing cells at a density of 2 × 10^7^ cells ml^-1^ were transferred to plastic dishes without lids and irradiated with UV-C (254 nm) using UV lamps (UVP Inc., Upland, CA, USA) at a dose of 400–16,000 J/m^2^. The cells were harvested and then spread on solid medium before or after UV irradiation. Surviving colonies were counted after 7 days of growth. Colony formation assays conducted after irradiation with UV-C at 400 J/m^2^ showed 80% viability of *Synechocystis* cells (Supplementary Figure S2), consistent with previous reports (44). This intensity of irradiation was therefore used in all assays.

### RNA extraction and northern blot analysis

After UV-C irradiation at 400 J/m^2^ or mock treatment, *Synechocystis* 6803 cells were harvested by rapid filtration through hydrophilic polyethersulfone filters (Supor 800 filter, 0.8 μm, Pall, New York, NY, USA). The filter covered with cells was immediately immersed in 1 ml of PGTX solution (45) and frozen in liquid nitrogen. Total RNA was extracted as described previously (46). To eliminate contaminating genomic DNA, 5 μg of each RNA sample was incubated twice with 2 units of TURBO DNase (TURBO DNA-free Kit, Ambion, Austin, TX, USA) for 30 min at 37 °C. DNase was inactivated according to the manufacturer’s instructions. RNA was re-extracted in phenol/chloroform and purified by ethanol precipitation. The total RNA samples were analyzed by electrophoretic separation of 3 μg of RNA, and northern hybridization experiments were performed as previously described (46) using single-stranded transcript probes generated by PCR and *in vitro* transcription. The PCR-generated probe templates were obtained using the primers slr1129-5UTR-f and slr1129-5UTR-rT7 (*rne* 5′ UTR probe) or slr1129orf-f and slr1129orf-rT7 (*rne* ORF probe); for the primer sequences, see Supplementary Table S1 and Figure S4A.

### Microarray analysis

The microarray design, hybridization and data analysis have been described previously (47). The Agilent microarrays contain oligonucleotide probes representing all annotated mRNAs as well as most other expressed transcripts, allowing precise determination of individual transcripts with respect to both DNA strand and genomic location. Total RNA (5 µg) was extracted from *Synechocystis* 6803 cells collected 1 h or 2 h after UV-C irradiation at 400 J/m^2^ or following mock treatment and directly labeled with Cy5 (without cDNA synthesis) using Kreatech’s ULS labeling kit (Kreatech Diagnostics, B.V., Netherlands) according to the protocol provided by the manufacturer. RNA fragmentation and hybridization for Agilent one-color microarrays were performed according to the manufacturer’s instructions using 1.65 µg of each labeled RNA. Array analysis was performed using biological duplicates. The raw data were quantile normalized. The differences in the transcriptomes of cells subjected to UV-C and mock conditions were determined for each time point. A transcript was considered differentially expressed when it met the significance criteria (log_2_FC ≥│1│, adj. p value ≤ 0.05). P values were adjusted for multiple testing using the Benjamini–Hochberg method. The comparative microarray data are shown in Supplementary Data 1, and the raw data have been deposited in the GEO database under the accession number GSE186330.

### Estimation of transcript half-lives by quantitative real-time reverse transcription PCR (RT–qPCR)

We designed specific primer sets for use in analyzing two segments within the *rne* 5′ UTR (5′ UTR-1 and 5′ UTR-2) and two segments within the coding region of the RNA (ORF-1 and ORF-2) by RT–qPCR (Supplementary Figures S4B and S4C). To estimate the half-lives of the *rne* 5′ UTR and coding region segments, 200 µg/ml rifampicin, a transcription inhibitor (48), was added 2 hours after UV-C treatment of the cells at 400 J/m^2^. RNA samples were prepared from cells collected 0, 5, 10 and 30 minutes after rifampicin addition by rapid filtration of the cells onto Supor 800 membranes as described above, and relative amounts of the RNA were quantified by RT–qPCR. For this, cDNA was prepared from 2 µg of each RNA sample using the PrimeScript II 1^st^ strand cDNA synthesis kit (TaKaRa, Shiga, Japan) with 40 units of RNasin Plus RNase inhibitor (Promega, Madison, WI, USA) according to the manufacturer’s instructions. RT–qPCR was performed using a StepOnePlus real-time PCR System (Applied Biosystems, Foster City, CA, USA) in standard mode (10 min at 95 °C followed by 40 cycles of 15 s at 95 °C, 15 s at 55 °C, and 60 s at 72 °C). Each 20 μl reaction contained 10 μl of Power SYBR Green Mix (Applied Biosystems), 2 μl of cDNA, and 0.4 μl each of the forward and reverse primers (final conc. 200 nM). The primers (Supplementary Table S1) were synthesized by Eurofins MWG Operon, Ebersberg, Germany. All reactions were conducted in triplicate, and 16S rRNA was amplified as a reference. Melting curves for the amplifications showed only single products. The data were analyzed using the StepOnePlus system SDS software (Applied Biosystems) with manual Ct and automatic baseline settings. Relative transcript quantities were calculated using the ΔΔCt method. The RT–qPCR data were used to calculate transcript half-lives by fitting the decay time-course abundance curves to an exponential decay function. The degradation constants and half-lives were calculated by fitting the data to an exponential decay curve with the R nls function.

### Calculation of synthesis rates

Assuming steady-state expression of a given transcript, the transcript levels (*Int*) are defined by the ratio of its synthesis rate to its transcription rate (*α*) and the degradation constant (*λ*), as follows:

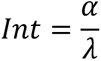

The fold change (*FC*) in the expression of two transcripts (*i,j*) is

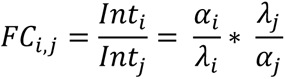

With the use of the microarray intensity data and the fits for the degradation constants, we can calculate the ratio of synthesis rates (FC_synt).

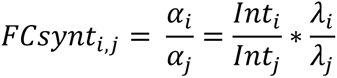

Assuming a single promoter for *rne* transcription, a synthesis ratio *FCsynt_UTR,coding_* > 1 indicates a termination after the 5′ UTR. Comparing the synthesis ratios of the UTR under mock and UV stress conditions, *FCsynt_UV,mock_* < 1 indicates a reduced transcription rate under UV stress conditions.

### Fluorogenic cleavage assay

A fluorogenic cleavage assay was performed as described previously (23) with minor modifications in the method used to prepare the cell extract. After UV irradiation at a dose of 400 J/m^2^, the cells were collected immediately or after cultivation at room temperature for 2–4 hours and were then stored at -20 °C. The cell pellets were resuspended in 300 µl of reaction buffer (25 mM Tris-HCl (pH 8.0), 60 mM KCl) in a vessel containing 300 mg of glass beads and broken in a Mini-BeadBeater-16 (BioSpec, Bartlesville, OK, USA). The resulting cell extract was centrifuged at 400 × *g* for 5 min at 4 °C. The supernatant containing the soluble and membrane proteins was transferred to a new tube and used in the assay. The cleavage reaction was monitored for 70 min at one-minute intervals by fluorometry using a Victor^TM^X^3^ multilabel plate reader (PerkinElmer, U.S.; excitation at 480 nm, emission at 520 nm).

### Overexpression and purification of Synechocystis 6803 RNase E and preparation of polyclonal antisera

Expression of recombinant RNase E and affinity purification under native conditions were performed as described (29). The preparation of rabbit antiserum against the purified RNase E protein was based on previous methods (49) (Protein Purify Co. Ltd, Japan). The recombinant RNase E was subjected to SDS–PAGE and recovered from the gel. The gel slices were crushed, mixed with adjuvant and injected beneath the skin of a rabbit in the area of the back as an antigen. Each antigen was injected five times over a period of three months, and the resulting antibody titers were measured by ELISA. Whole blood was collected, and whole antiserum was obtained by centrifugation. The antiserum was stabilized by the addition of NaN_3_ at a final concentration of 0.1% (w/v) and stored at - 80 °C until use.

### Western blot analysis and determination of RNA half-life

After UV-C irradiation at 400 J/m^2^ or mock treatment, *Synechocystis* 6803 cells were harvested by centrifugation at 6,000 *g* for 10 min at 4 °C. To analyze the cellular localization of RNase E protein, cytosolic and membrane fractions were separated by centrifugation according to a previously described procedure (50). Crude extracts of the cells were prepared using 10% trichloroacetic acid as previously described (51). To avoid degradation of RNase E protein, the crude extracts were prepared immediately after cell harvesting. Twenty micrograms of each sample was analyzed by western blotting using primary antisera against RNase E and RbcL (Agrisera) at dilutions of 1:3,000 and 1:8,000, respectively; HRP-conjugated anti-rabbit IgG (GE Healthcare) was used as the secondary antibody. The images and quantities of each protein signal were obtained using a ChemiDoc XRS + system with Image Lab software (Bio–Rad laboratories).

To estimate the half-life of RNase E protein, 250 µg/ml of chloramphenicol, a translation inhibitor, was added 2 hours after UV-C treatment of the cells at 400 J/m^2^. Crude extracts were prepared from the cells as described above and electrophoretically separated by SDS–PAGE followed by western blot analysis.

### Rapid Amplification of 3′ ends

Rapid amplification of 3′ ends (3′ RACE) was performed according to Argaman et al. (52). Total RNA was prepared from a *Synechocystis* 6803 culture grown under standard conditions. After ligation of an RNA adapter, the RNA was reverse-transcribed and PCR-amplified using primers that anneal to the 5′ UTR of *rne* or to the adapter sequence. An electrophoresis gel image of PCR products is shown in Supplementary Figure S7. DNA fragments were ligated into the pGEM-T vector and transferred into *E. coli*. Single colonies were picked, and the inserts were sequenced by Sanger sequencing. The sequences of the RNA adapters and DNA primers used in this study are listed in Supplementary Table S1.

### In vitro cleavage assay

The *rne* 5′ UTR was transcribed *in vitro* from a PCR-generated template using the oligonucleotides rne5UTR-T7-fw and rneATG-rev. For *in vitro* transcription of the variants of the *rne* 5′ UTR (mutation 1 and mutation 2, Figure 6E), template DNA was synthesized in a fusion-PCR approach. Briefly, *rne* 5′ UTR fragments 1, 2 and 3 were amplified using the oligonucleotide pairs rne5UTR-T7-fw/frag1-rev, frag2-fw/rneATG-rev and frag3-fw/rneATG-rev, respectively. Next, fragments 1 and 2 (fusion product 1) and fragments 1 and 3 (fusion product 2) were combined and used as template DNA for fusion PCR, each in combination with oligonucleotides rne5UTR-T7-fw and rneATG-rev. Fusion products 1 and 2 were used as template DNA for *in vitro* transcription of *rne* 5′ UTR mutation 1 and mutation 2 variants, respectively. Residual template DNA was depleted as described (23), and the full-length RNAs generated *in vitro* were purified from polyacrylamide (PAA) gels as described (31). *In vitro* RNase E cleavage assays were performed as described previously (23) with the following modifications: 0.8 pmol of RNA was incubated with 7 pmol of *Synechocystis* 6803 recombinant RNase E for 30 min at 30 °C; after separation of the mixture on 7 M urea-6% PAA gels, RNA was transferred to a Hybond-N nylon membrane (Amersham) and subjected to northern blot hybridization. RNA gel blot hybridizations, 5′-radiolabeling and purification of oligonucleotide probes were performed as described previously (23). The oligonucleotides used as probes are listed in Supplementary Table S1.

### RNase E protection assay

Reaction mixtures (3 µl each) containing 0.5 pmol of *in vitro* transcribed RNA (*rne* 5′ UTR, *rne* 5′ UTR mutation 1, and *rne* 5′ UTR mutation 2) and 2 pmol of the oligonucleotides as-rne 210/228 and as-rne 200/234 were incubated for 5 min at 85 °C and then briefly chilled on ice. The reaction mixture was then supplemented with 1 µl 5× RNase reaction buffer (125 mM Tris–HCl (pH 8.0), 300 mM KCl, 25 mM MgCl_2_, 500 mM NH_4_Cl, 0.5 mM DTT) and incubated at room temperature for 15 min. Recombinant RNase E (7 pmol) was added to increase the total reaction volume to 5 µl, and incubation was continued at 30 °C for 15 min. RNase E activity was quenched by the addition of 1 µl 0.5 M EDTA and 1× volume loading buffer. Following heating at 95 °C for 3–5 min, cleavage products were separated on 7 M urea–6% polyacrylamide gels and subjected to northern blot hybridization as described above.

## Results

### Microarray analysis and induction of rne transcripts after UV irradiation

To study the regulation of *rne* expression in cyanobacteria, we focused on the UV stress response because UV is an inevitable fraction of solar irradiation and because the *rne* gene has been reported to be induced in *Synechocystis* 6803 under conditions of UV stress (39). We first conducted a UV-C irradiation assay by exposing cells from exponentially growing cultures to 400–16,000 J/m^2^ UV irradiation. Colony formation assays indicated that 80% survival of the cells was obtained after irradiation at 400 J/m^2^ (Supplementary Figure S2); thus, this intensity of irradiation was used in all subsequent experiments.

A microarray analysis in which cells subjected to mock and UV stress conditions were revealed dynamic transcriptome changes during the UV stress response. After 1 h of UV irradiation, we observed 275 upregulated and 306 downregulated genes, and at 2 h after the initiation of UV stress we observed 189 upregulated and 218 downregulated genes (log_2_FC ≥│1│, adj. p values ≤0.01; Supplementary Data 1). Because the microarray also contained probes that detect UTRs and noncoding transcripts, the term “gene” here includes not only protein-coding genes but also sRNAs and separate UTRs. A genome-wide graphical overview of probe localization and signal intensities is shown in Supplementary Data 2. A volcano plot indicating log-transformed fold changes (FCs) at 2 hours after UV treatment (compared to cells that received mock treatment) is shown in Figure 1A. Several genes classified in the Cyanobase (53, 54) GO categories “translation” and “photosynthesis and respiration”, including gene clusters encoding ribosomal proteins, ATPase subunits and RubisCo subunits, were dramatically downregulated 1 hour after UV irradiation (Supplementary Figure S3A–C, Supplementary Data 1). In contrast, *slr1639,* which encodes SmpB, a protein that binds to tmRNA (*ssrA* RNA) and works in concert with it to rescue stalled ribosomes (55), was upregulated, as were some specific ribosomal genes (Supplementary Figure S3D, Supplementary Data 1). These results suggest that translation arrest and reconstruction of the ribosome occurred during this period. Several sRNAs were also upregulated at these time points. Among them, PsrR1, a negative posttranscriptional regulator of multiple PSI genes in response to high light stress (31), was induced; furthermore, HLIP genes (*hliA*: *ssl2542*; *hliB*: *ssr2595*; *hliC*: *ssl1633*) (Supplementary Figure S3E-H), the products of which quench absorbed light energy and assist chlorophyll biosynthesis and PS II assembly (56). In addition, the sRNA *ncl0380*, which corresponds to the 5′ UTR of *sll1799*, was upregulated at both 1 h and 2 h after UV treatment, while the downstream-located ribosomal gene cluster was strongly downregulated at 1 h after UV treatment and then showed a gradient of differential partial recovery over time (Supplementary Figure S3A); this recovery was most pronounced for the first genes in the cluster and strongly decreased toward the end of the operon. Interestingly, we did not observe differences in the transcript levels of *lexA*, consistent with previous observations that the *Synechocystis lexA* gene is not induced by DNA damage (57). Regarding the regulation of genes that encode ribonucleases, we observed marked upregulation of *rne* 2 h after UV irradiation (Figure 1B), consistent with a previous report (39). Likewise, the *rnj* gene (*slr0551*) encoding RNase J (Supplementary Figure S3I) was also upregulated at this time point. RNase E and RNase J are key enzymes in RNA metabolism. Therefore, their enhanced transcript accumulation likely indicates their involvement in transcriptome remodeling following UV-induced damage. We also noted a unique UV stress response of the *rimO-crhR* operon, one of the targets of RNase E (34). The *rimO* transcript level was significantly upregulated after UV treatment, whereas *crhR* mRNA levels responded in an inverse fashion by transiently decreasing one hour after UV treatment (Supplementary Figure S3J). This is consistent with a previous observation of posttranscriptional operon discoordination in the UTR between *rimO* and *crhR* (34). The fact that the expression pattern of *crhR* was similar to the pattern of expression of ribosome and ATPase gene clusters suggests that CrhR is involved in their regulation.

**Figure 1.**
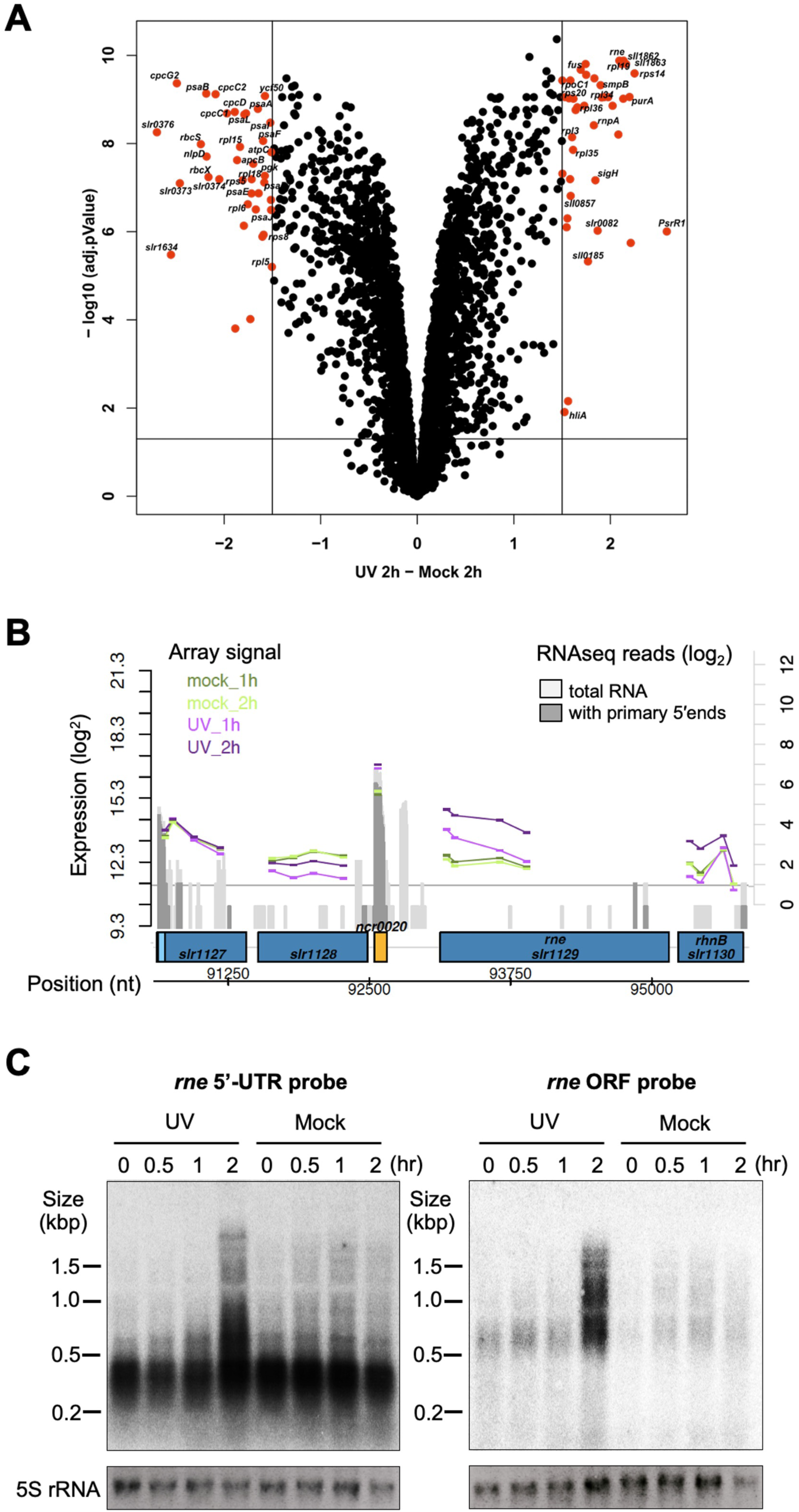
Transcriptomic response to UV treatment and increased accumulation of the *rne* transcript. (A) Volcano plot: log-transformed fold changes (FCs) between samples taken 2 hours after UV irradiation and after mock treatment (x-axis, difference in log_2_ expression values) and −log10 (adjusted p value, y-axis). The lines indicate the adjusted p value threshold of 0.05 and the FC thresholds of 1 and −1. The entire dataset is shown in the genome-wide expression plot (Supplementary Data 2), and numeric values are presented in Supplementary Data 1. (B) Detailed view of the *rne* locus with array probes indicated by vertical bars connected by colored lines. The *rne* gene, which is transcribed together with the *rnh* gene encoding RNase H, has a long 5′ UTR from which separate shorter transcripts can also accumulate, annotated as *ncr0020*. The signal intensities are given as log_2_ values. The graphs shown in gray represent RNA sequencing data given as log_2_ read numbers; these were extracted from the previous genome-wide mapping of TSSs (82). (C) Northern blot analysis of *rne* expression after UV treatment using single-stranded RNA probes that hybridize either to the *rne* 5′ UTR or to the coding region (Supplementary Figure S4A and Supplementary Table S1). 5S rRNA accumulation is shown for the control.

We next investigated the UV stress response of the *rne* gene in more detail. Northern blot analyses using probes specific for the *rne* 5′ UTR or the coding sequence 5′ portion (Supplementary Figure S4A) revealed that transcripts in the 200–500 nt range originating from the 5′ UTR were abundant under all test conditions (Figure 1C, left). In contrast, the *rne* full-length transcript became detectable 2 hours after UV treatment (Figure 1C, right, UV), whereas the *rne* transcript steady-state level remained low in the mock condition without UV treatment (Figure 1C, right, mock). The largest distinct mRNA, with a length of ∼3.2 kb, was seen 2 hours after UV treatment (Figure 1C); this mRNA originates from the dicistronic transcriptional unit that consists of *rne* and the downstream located *rnh* gene encoding RNase H (58). Consistent with previous reports (39), these results indicate that *rne* expression is upregulated after UV-C irradiation and that the *rne* 5′ UTR accumulates as an abundant and separate transcript, consistent with previous transcriptome data in *Synechocystis* 6803 (Figure 1B and Supplementary Figure S1B) (58) and *Synechocystis* sp. PCC 6714 (Supplementary Figure S1B) (59), a closely related strain (58).

### Elevated RNase E activity after UV-C irradiation

To determine whether the observed change in *rne* gene expression resulted in higher RNase E enzyme activity, we used a previously established activity assay (23). This assay is based on a fluorogenic RNA oligonucleotide that consists of a FAM tag, a BHQ-1 quenching tag and a previously reported *Synechocystis* 6803 RNase E recognition site. In this assay, RNase E activity is monitored via fluorescence from the cleaved FAM oligonucleotide fragment. Using this system, we followed the RNase E activity in *Synechocystis* cells before and after UV-C treatment. Samples were collected immediately and at 2, 3 and 4 h after irradiation of the cells with UV (400 J/m^2^), and cell extracts were prepared. RNase E cleavage was measured at one-minute intervals over a period of 70 min using equal amounts of cell lysate protein per sample (Figure 2A). In all incubations except the buffer control, the initial steep increase in fluorescence was followed by a plateau. The reaction efficiency of RNase E obtained by mixing the crude extract with the substrate and incubating for 15 min was compared (Figure 2B). RNase E activity was higher at all measured time points following UV irradiation than in the nonirradiated controls. Fluorescence increased with time after UV treatment (Figure 2), suggesting that stimulation of RNase E enzymatic activity is a time-dependent process. This result is consistent with the northern blot result, in which substantial accumulation of full-length *rne* mRNA was observed at 2 h but not earlier (Figure 1C). A similar increase in RNase E activity in protein extracts from UV-irradiated cultures was consistently observed, even when the experiment was repeated using smaller amounts of protein (Supplementary Figure S5A). The activity was constant in the mock treatment experiment (Supplementary Figure S5B).

**Figure 2.**
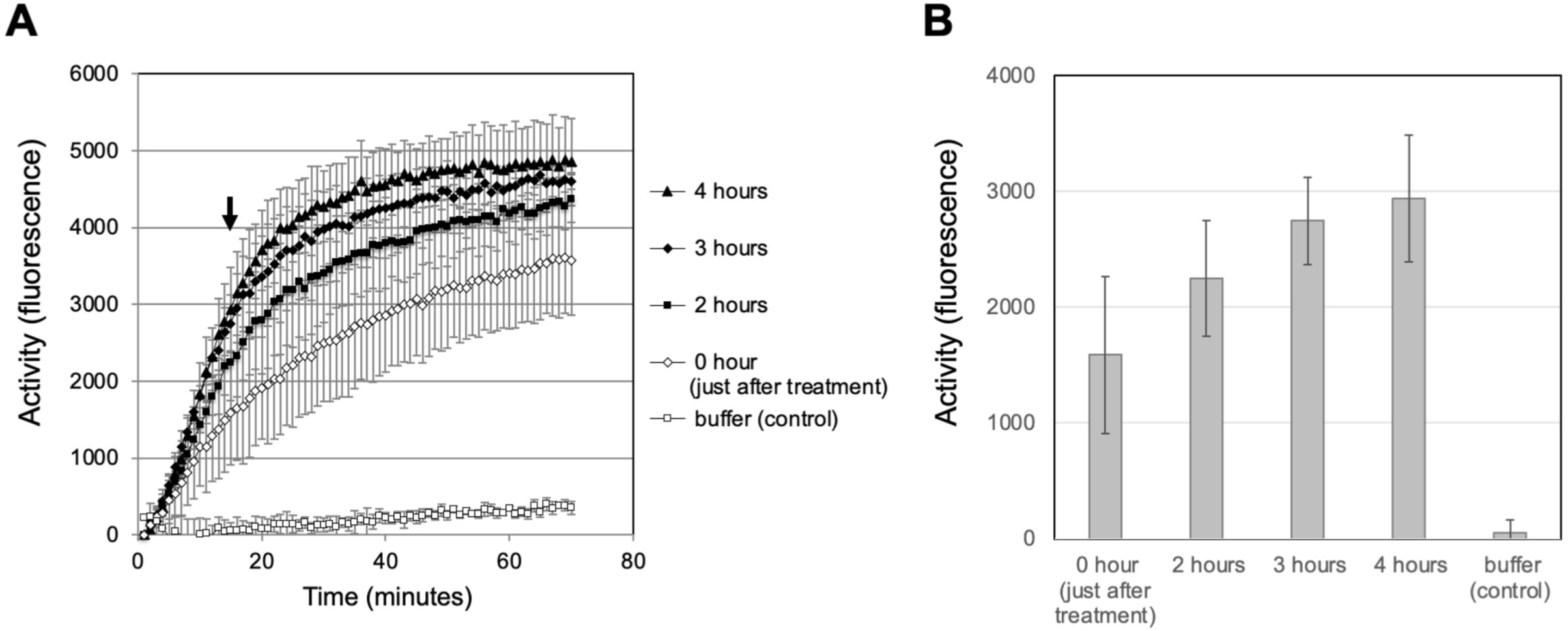
Induction of RNase E activity after UV-C treatment. (A) RNase E activity in crude extracts prepared at various times after UV-C treatment (open diamonds: just after treatment; closed squares: after 2 h; closed diamonds: after 3 h; closed triangles: after 4 h; open squares: buffer control) measured in an *in vivo* fluorescence-based assay with a duration of 70 min. The arrow indicates the high reaction efficiency achieved after 15 minutes of incubation. (B) Comparison of RNase E activities at 15 minutes incubation. The standard deviations of the values obtained for three biological replicates are shown.

### Turnover of RNase E protein after UV irradiation

The observed enhancement of RNase E activity in cell lysates (Figure 2) after UV irradiation could result from the presence of more RNase E protein, higher specific activity of the enzyme, or both. To distinguish these possibilities and to permit the direct detection of RNase E, an antiserum against recombinant *Synechocystis* 6803 RNase E was generated. When used in western blots, this antiserum showed specific signals for RNase E protein at ∼100 kDa, higher than the predicted molecular mass of RNase E, together with likely nonspecific signals at ∼65 and 55 kDa (Figure 3). This is consistent with previous results in *E. coli,* in which RNase E of a larger size than predicted was detected (60). The fact that RNase E was clearly observed at the protein level (Figure 3A) indicates that the low amounts of its mRNA that are present in the nonstress condition (Figure 1) are sufficient for expression of RNase E protein. For comparison, we used an antiserum against the large subunit of RubisCo and normalized the RNase E signal detected on the immunoblot to the RubisCo signal. This comparison showed that the protein levels of RNase E at 1 and 2 hours after UV irradiation were almost the same as those present in the mock condition (Figure 3A), in contrast to the increased accumulation of *rne* full-length transcripts and upregulated RNase E activity observed 2 hours after UV irradiation (Figure 1 and Figure 2). This suggests either that the additional mRNA copies were not efficiently translated or that the stability of the protein was decreased. Thus, we compared the stability of RNase E in cells that had or had not received UV treatment. Chloramphenicol, a translation inhibitor, was added after UV irradiation, and the cells were harvested at the indicated time points. Western blot analysis revealed that the protein levels of RNase E in UV-treated cells greatly decreased 30 and 60 min after the addition of chloramphenicol, while the RbcL signal and the intensity of the nonspecific bands remained constant (Figure 3B). In cells that were not UV-treated, the intensity of the RNase E signal remained constant despite the addition of chloramphenicol (Figure 3B, mock). We conclude that the enhancement of RNase E activity we observed in the earlier experiment was not caused by an increase in the amount of RNase E. We further conclude that active translation of mRNA was required to keep the cellular amount of the enzyme constant following UV treatment and that the decreased stability of RNase E we observed was likely linked to the degradation of UV-damaged RNase E. These results suggest that the turnover of RNase E protein was accelerated by activation of protein degradation after UV irradiation and concerted initiation of resynthesis of the enzyme from the increased amount of *rne* full-length mRNA.

**Figure 3.**
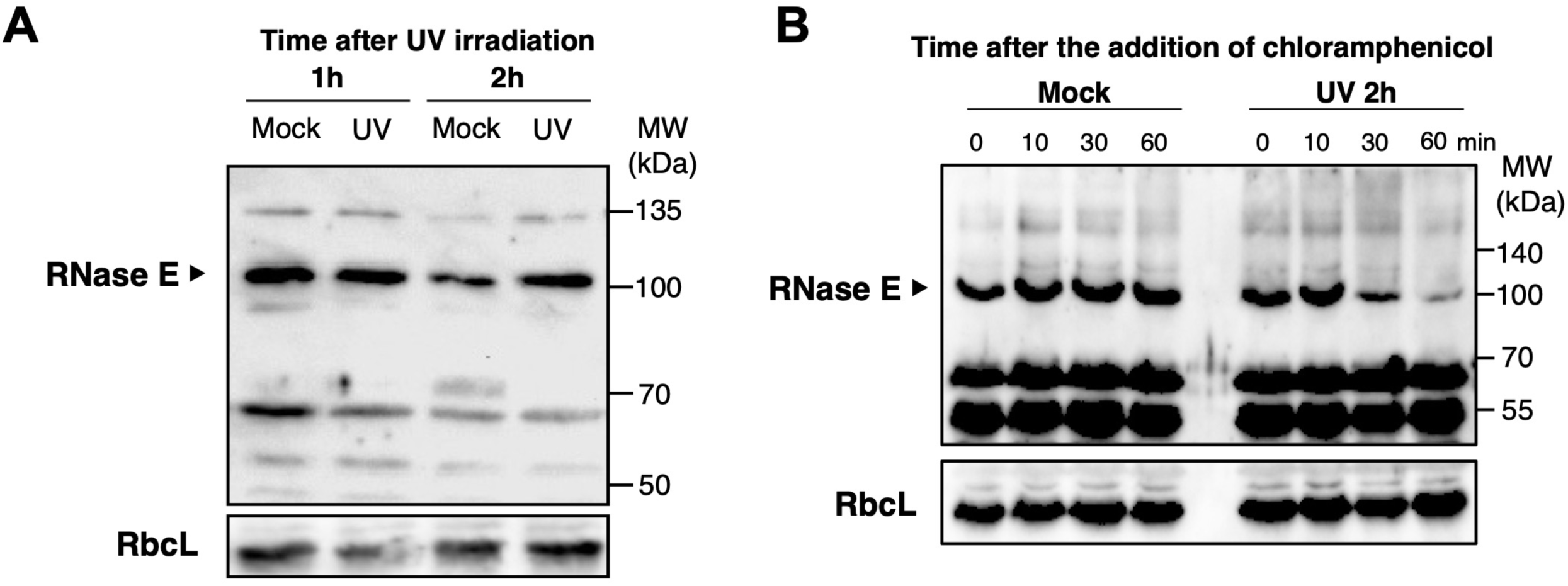
Turnover of RNase E protein after UV-C treatment. (A) Comparison of RNase E expression levels. Twenty micrograms of total protein were prepared from bacterial cultures one and two hours after UV-C treatment, loaded on an SDS-PAA gel and subjected to western blot analysis using antibodies against RNase E (arrowhead) or RbcL used as an internal control. (B) Comparison of the stability of RNase E. The translation inhibitor chloramphenicol was added to the cultures 2 hours after UV-C treatment, and cells were harvested at the indicated time points for western blot analysis.

### Stabilization of rne transcripts during the UV stress response

To identify the mechanisms that underlie the increased amount of *rne* full-length mRNA and hence permit the enhanced turnover of RNase E during the UV stress response, transcript half-lives were determined. After UV irradiation, rifampicin, which inhibits the initiation of transcription by binding to the β subunit of RNA polymerase (48), was added to the cultures. For more robust results, the following calculations were based on technical triplicates and biological replicates of the respective two 5′ UTR and ORF segments. The relative amounts of 5′ UTR-1, 5′ UTR-2, ORF-1 and ORF-2 present at the indicated time points were determined by RT–qPCR. In parallel, a mock treatment was performed in which UV irradiation was omitted.

Under the mock conditions, signal intensities for the probed segments within the 5′ UTR and those within the coding region decreased rapidly (Figure 4). The calculated half-life of the *rne* 5′ UTR segments was 3.1 min, while that of the ORF was 7.2 min (Table 1), suggesting low stability of the *rne* transcript, similar to *E. coli* (16). Compared to the mock condition, substantial stabilization of the 5′ UTR segments but not of the coding region segments was observed after UV stress treatment. The calculated half-life increased to 17.1 min for the *rne* 5′ UTR, while a half-life of 7.5 min was determined for the coding sequence (ORF, Figure 4 and Table 1).

**Figure 4.**
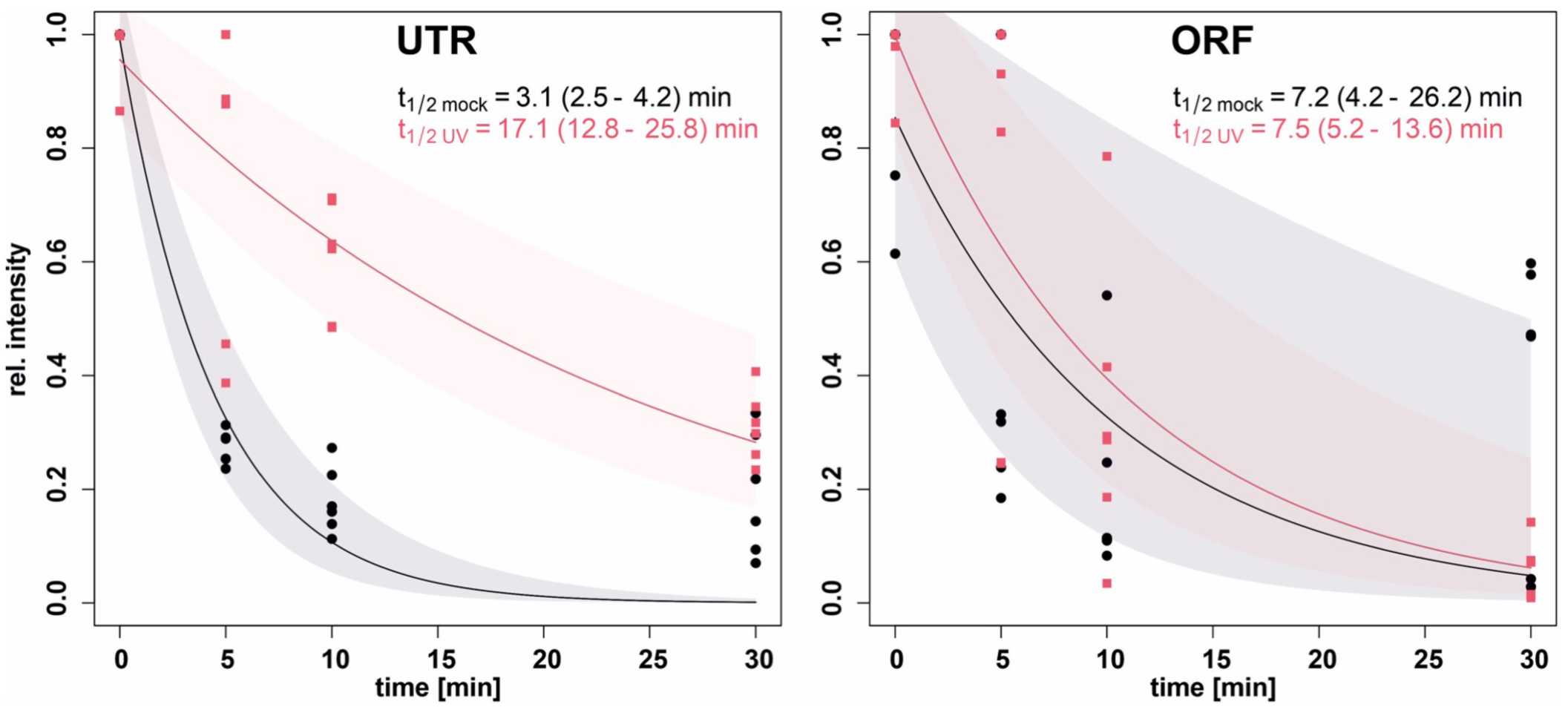
Upregulation of *rne* full-length transcripts after UV-C treatment. The ratios of the amounts of *rne* 5′ UTR and coding region transcripts are shown, together with the half-lives of each transcript. The transcription inhibitor rifampicin was added to the cultures 2 hours after UV-C treatment, and cells were harvested at the indicated time points for RT–qPCR analysis using primer sets that anneal to two different regions of the *rne* 5′ UTR (5′ UTR-1 and 5′ UTR-2) or the coding region (ORF-1 and ORF-2) (Supplementary Figure S4B, C and Supplementary Table S1). The data from tests conducted in triplicates were normalized to the amount of 16S rRNA, and the ratios were calculated. Half-life and decay were calculated based on two independent regions in the UTR and the ORF each with three biological replicates for each region. The fitting curves for the mock treatment and UV-stress are given in black and red, respectively. The 95% confidence interval areas are shaded accordingly.

**Table 1.** Intensities and stabilities of the 5′ UTR and ORF in the rne transcript based on microarray signals and half-lives measured by RT–qPCR. The intensity data are based on two biological replicates. Half-life and decay were calculated based on two separately probed regions in the 5′ UTR and ORF, each with three biological replicates for each region. For the intensity the standard deviation is indicated and for half-life and decay constant the 95% confidence interval is given.

|  | Mock |  | UV 2h |  |
| --- | --- | --- | --- | --- |
|  | 5'UTR | ORF | 5'UTR | ORF |
| Intensity (microarray) [AU] | $5 \cdot 10^4 \pm 2.1 \cdot 10^3$ | $4.5 \cdot 10^3 \pm 6.1 \cdot 10^1$ | $1.9 \cdot 10^4 \pm 3 \cdot 10^2$ | $1.3 \cdot 10^5 \pm 4.7 \cdot 10^2$ |
| Half life of transcript [min] | 3.1<br>(2.5-4.2) | 7.2<br>(4.2 -26.2) | 17.1<br>(12.8-25.8) | 7.5<br>(5.2-13.6) |
| Decay constant<br>[1/min] | 0.22<br>(0.17-0.28) | 0.1<br>(0.03-0.17) | 0.04<br>(0.03-0.05) | 0.09<br>(0.05-0.13) |

To distinguish between transcriptional and posttranscriptional effects on the stability of *rne* transcripts in *Synechocystis* 6803 exposed to UV stress, we used the experimentally observed array signal intensities and the estimated half-lives of the transcripts to calculate the expected changes in the rate of synthesis of the protein, as summarized in Tables 1 and 2 and Supplementary Figure S6. Under mock conditions, the stability of the coding region was 2.2 times higher than that of the UTR (Table 2). Because there is only a single promoter in front of the *rne* gene (26), the observed difference in expression of the 5′ UTR and the coding region can only be explained by premature transcription termination after the 5′ UTR; this is in good agreement with our calculations: ∼95.9% of the polymerases terminate within the 5′ UTR (Supplementary Figure S6). Under UV stress conditions, the stability of the coding region was roughly unchanged (Table 1), while the increased intensity levels indicated a reduction in transcription termination to ∼65.3% (Table 2 and Supplementary Figure S6). Interestingly, the stability of the 5′ UTR under UV stress conditions increased 5.5-fold (in other words, the decay constant decreased by a factor of 0.18) compared to its stability under mock conditions, while the intensity level increased by only a factor of 2.52, indicating that the rate of transcription of *rne* was reduced by a factor of ∼2.2 under UV. Together, the data show that the increased *rne* mRNA levels observed in cells under UV stress are due to posttranscriptional stabilization and reduced premature termination (Figure 7A).

**Table 2.** Change in synthesis rates for the 5′ UTR and ORF regions of the rne transcript under mock and UV stress conditions. FC, Fold change.

|  | mock | UV | 5'UTR |
| --- | --- | --- | --- |
|  | 5'UTR/orf | 5'UTR/orf | UV/mock |
| FC intensity | 11 | 6.5 | 2.52 |
| FC decay<br>constant | 2.2 | 0.4 | 0.18 |
| FC synthesis-<br>rate | 24.2 | 2.9 | 0.46 |
| Termination<br>after UTR | 95.9% | 65.3% | NA |

### Detailed analysis of the 5′ UTR in Synechocystis 6803

We observed a particularly large difference in the calculated half-life of the 5′ UTR in UV-treated versus untreated cells (Figure 4) and an abundant accumulation of separate, 5′ UTR-derived transcripts (Figure 1). Both of these observations suggest that the 5′ UTR plays a pivotal role in the control of transcript stability and the determination of whether full-length mRNA will accumulate. These findings led us to hypothesize that RNase E expression may be regulated via its extremely long 5′ UTR. Recent mapping of RNase E-dependent cleavage sites in *Synechocystis* 6803 by TIER-seq, which also points at sites within the *rne* 5′ UTR, supports this idea (36). However, in the TIER-seq analysis, mainly 5′ ends were mapped, whereas the 3′ ends remain largely unexplored.

Therefore, we conducted 3′ RACE analysis to map possible 3′ ends within the *rne* 5′ UTR. The cDNA inserts of 17 clones were sequenced, and the mapped 3′ ends were positioned within the *rne* 5′ UTR. These 3′ ends primarily mapped to two regions that are located 78-90 nt (shorter 3′ ends) and 214-229 nt (longer 3′ ends) downstream of the TSS (Figure 5A). The most prominent RNase E cleavage sites within the *rne* 5′ UTR mapped by TIER-seq (36) also lie within the AU-rich region (nt positions 92762 and 92763) or very close by in the 3′ direction (positions 92771 and 92864) (Figure 5B). Moreover, compared to the shorter 3′ ends, the sequence surrounding the longer 3′ ends of the *rne* 5′ UTR is well conserved in *Synechocystis* 6803 and *Synechocystis* 6714 (Supplementary Figure S8 and Figure 5A). These observations further support the hypothesis that the expression of RNase E is self-regulated by cleavage within the 5′ UTR, with the AU-rich region as a key target site.

**Figure 5.**
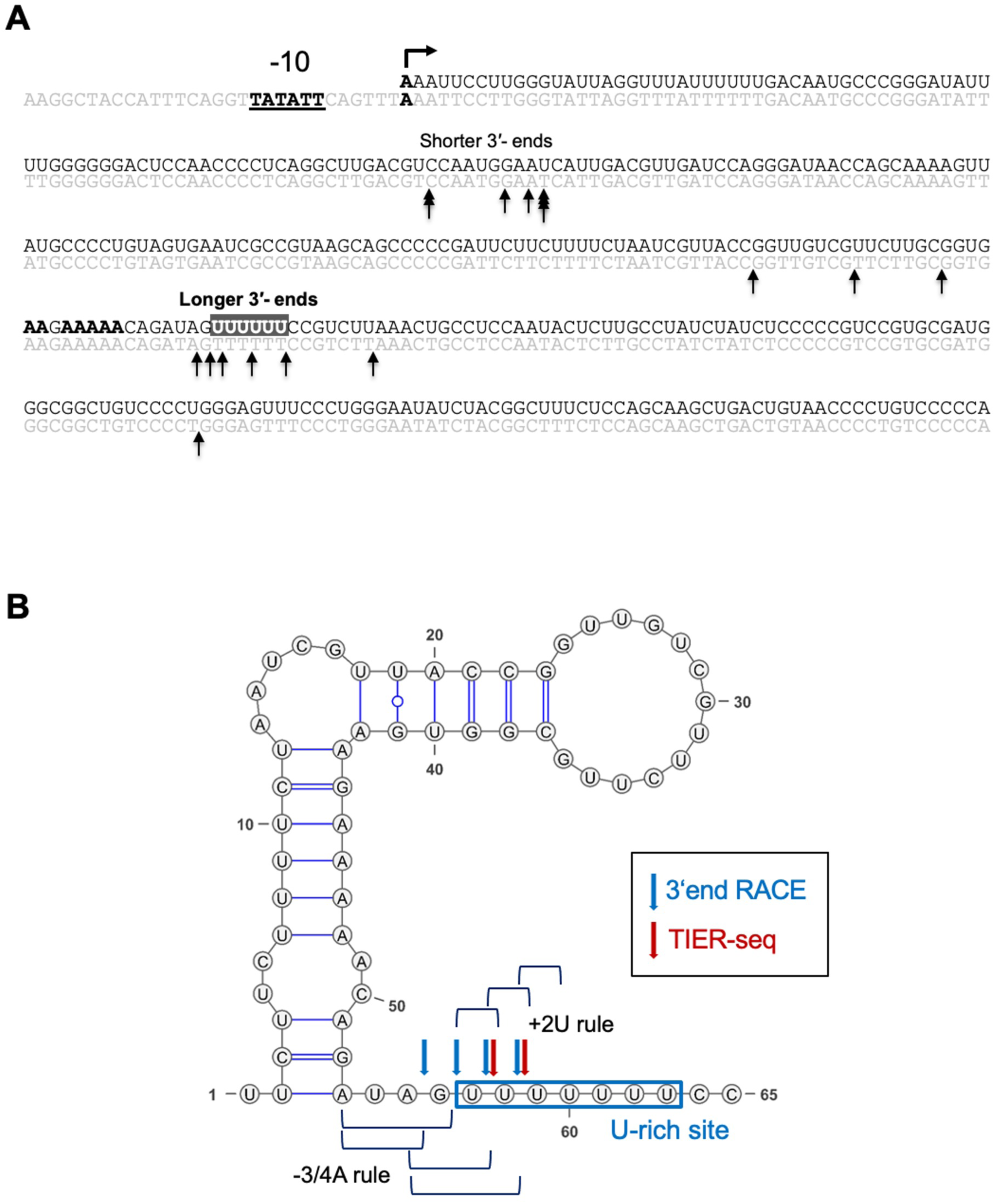
Analysis of 3′ ends within the *rne* 5′ UTR. (A) The 3′ ends mapped by the 3′ RACE assay within the first 355 nt of the *rne* 5′ UTR are indicated by arrows. The promoter (-10), TSS, and A-rich sites are represented by boldface letters. The U-rich site is indicated by the gray box and white characters. The 3′ ends mapped primarily to two regions that were located 78-90 nt (shorter 3′ ends) and 214-229 nt (longer 3′ ends) downstream from the TSS of *rne*. (B) Secondary structure of the U-rich site that forms part of the *rne* 5′ UTR predicted by RNAfold on the ViennaRNA website (84) with default settings and visualized using VARNA version 3.93 (85). RNA 3′ ends mapped by 3′ RACE and 5′ ends mapped by TIER-seq (36) are indicated by blue and red arrows, respectively. The RNase E consensus sequence suggested by TIER-seq (+2U rule: uridine at 2 nt downstream of the cleavage site; -3/4A rule: adenine at 3 to 4 nt upstream) is also shown.

In *Synechocystis,* not only RNase E but also RNase J, which can function as an endoribonuclease, are active (22). To discriminate between these two enzymes and to gain further insight into the location of cleavage sites within the 5′ UTR, *in vitro* assays of RNase E cleavage were performed. The complete *rne* 5′ UTR RNA (583 nt, Figure 6A) was transcribed *in vitro* and incubated with purified recombinant RNase E, and the cleavage products were separated by polyacrylamide gel electrophoresis. After treatment with RNase E, several distinct RNA fragments were observed, confirming the presence of multiple RNase E sites within the *rne* 5′ UTR (Figure 6B). Next, we performed northern blot analysis to identify the major fragments generated by cleavage within the *rne* 5′ UTR (Figure 6C). We used probes that recognize either the 5′ or the 3′ parts of the *rne* 5′ UTR with respect to the cleavage site (Supplementary Figure S4D). A particular fragment (labeled by the asterisks in the figure) resulting from the RNase E cleavage was noted; as this fragment was observed with probes 1 and 2 but not with probes 3 or 4, it must be derived from the first part of the 5′ UTR (Figure 6C). The length of this fragment corresponds well to that of an RNA that could extend from the TSS to the AU-rich site. To unambiguously demonstrate that the observed cleavage was performed by RNase E, we focused on the AU-rich region, which is conserved in *Synechocystis* 6803 and *Synechocystis* 6714, together with the mapped 3′ ends of *rne* 5′ UTR subfragments (Supplementary Figure S8). We used the fragment to test for the effects of point mutations in the AU-rich region (Figure 6D). Although substitution of A to C at the A-rich site had no effect, the cleavage product was no longer observed when nucleotides were substituted at both the A-rich and the U-rich sites (Figure 6E).

**Figure 6.**
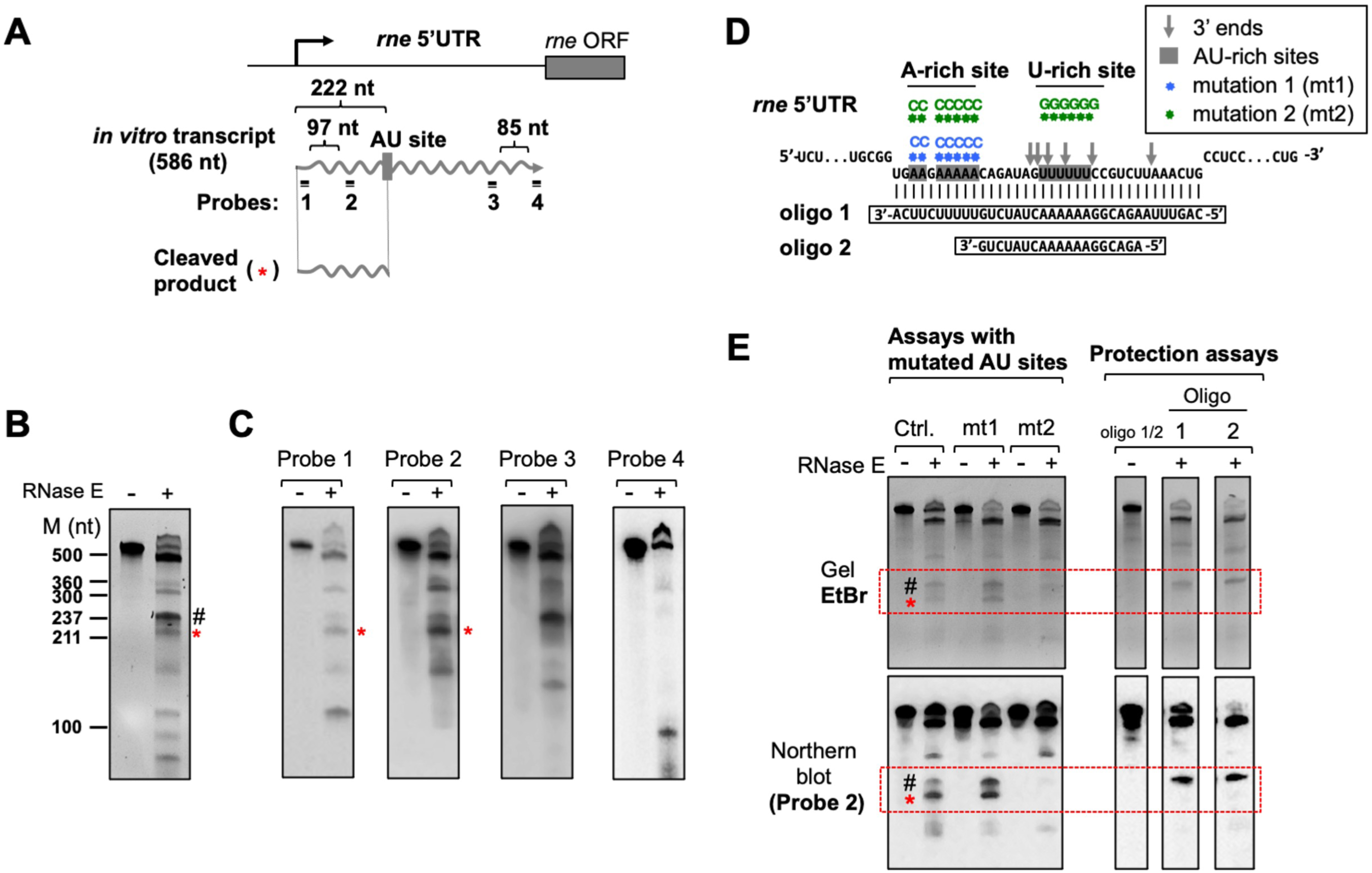
*In vitro* RNase E cleavage assay. (A) Summary of the cleavage of the *rne* 5′ UTR transcript by RNase E recombinant protein. The *rne* 5′ UTR transcript and the major products detected in the assay are shown, together with the locations at which probes that recognize different regions within the *rne* 5′ UTR transcript hybridize (Supplementary Figure S4D and Supplementary Table S1). (B) *In vitro* transcripts of the *rne* 5′ UTR were incubated with (+) and without (-) recombinant *Synechocystis* RNase E, and the resulting RNA cleavage patterns were visualized by ethidium bromide staining. Fragment sizes were estimated using NEB ssRNA markers. The major bands generated by RNase E digestion are marked by red asterisks and hash signs. The red asterisks mark a longer fragment that is similar in length to a fragment that could extend from the TSS to the two major 3′ ends as determined by 3′ RACE (Figure 5). (C) Northern blot analysis of RNase E-digested *rne* 5′ UTR transcripts. *In vitro* transcripts of the *rne* 5′ UTR were incubated with (+) and without (-) recombinant *Synechocystis* RNase E. After separation of the digestion products on polyacrylamide gels, the gels were subjected to northern blot analysis using specific probes (Supplementary Figure S4D and Table S1). (D, E) *In vitro* RNase E cleavage assays using mutant *rne* 5′ UTR transcripts and protection of the transcripts from RNase E attack. (D) Scheme of point mutations within the *rne* 5′ UTR transcript and the sequences of oligo-RNAs used in the protection assay. The 3′ ends of the *rne* 5′ UTR, mapped by 3′ RACE, are indicated by gray arrows. (E) RNase E cleavage assay (left) and protection assay (right). The RNA cleavage patterns were visualized on ethidium bromide-stained 7 M urea-6% polyacrylamide gels (upper image) and by northern blot analysis using probe 2 (lower image). The specific bands generated by RNase E digestion are marked by the same asterisk and hash symbols.

To further verify this result, protection assays were performed using oligo RNAs that cover either only the U-rich site or both the A-rich and the U-rich sites. The cleavage product in question disappeared after the addition of either of the two tested oligo RNAs, both of which cover the U-rich site, whereas the missing complementarity to the A-rich site in the case of oligo 2 was not relevant (Figure 6E). These results indicate that RNase E cleaves the U-rich site rather than the A-rich site in the *rne* 5′ UTR *in vitro* and are consistent with the results of our analysis of RNA ends *in vivo* by 3′ RACE mapping (Figure 5B) and with the results of 5′ end mapping according to the TIER-seq dataset (36); those experiments also showed that the U-rich site is a preferred site for RNase E cleavage within the *rne* 5′ UTR (Figure 5B).

## Discussion

Here, we demonstrate that in *Synechocystis* 6803, comprehensive transcriptome remodeling occurs and RNase E turnover is accelerated during the UV stress response and identify a posttranscriptional regulatory mechanism that is involved in this process. The results of *in vitro* RNase E assays in the presence and absence of protecting oligonucleotides and with RNAs containing specific point mutations (Figure 6) are consistent with the results of 3′ end RACE mapping and 5′ end mapping by TIER-seq, confirming that RNase E cleaves within the 5′ UTR of its own mRNA in the U-rich site (Figure 5). Moreover, the TIER-seq data indicate the possible presence of additional sites in the region approximately 10 to 100 nt downstream of the U-rich site (36). These facts strongly suggest that RNase E regulates its level of expression by targeting the 5′ UTR of its own transcript, superficially resembling the autoregulation of RNase E levels in *E. coli* (16, 19). However, the observed increase in the half-life of the full-length *rne* transcript after UV stress cannot be fully explained by stability control of the *rne* 5′ UTR. Our data indicate that autoregulation also depends on the premature termination of *rne* mRNA transcripts and does not involve an increased transcription rate (Figure 7A and Supplementary Figure S6). Cleavage at the 5′ UTR site by RNase E might also lead to termination of transcription. Hence, termination would cease if the cleavage was prevented due to UV inactivation of RNase E.

**Figure 7.**
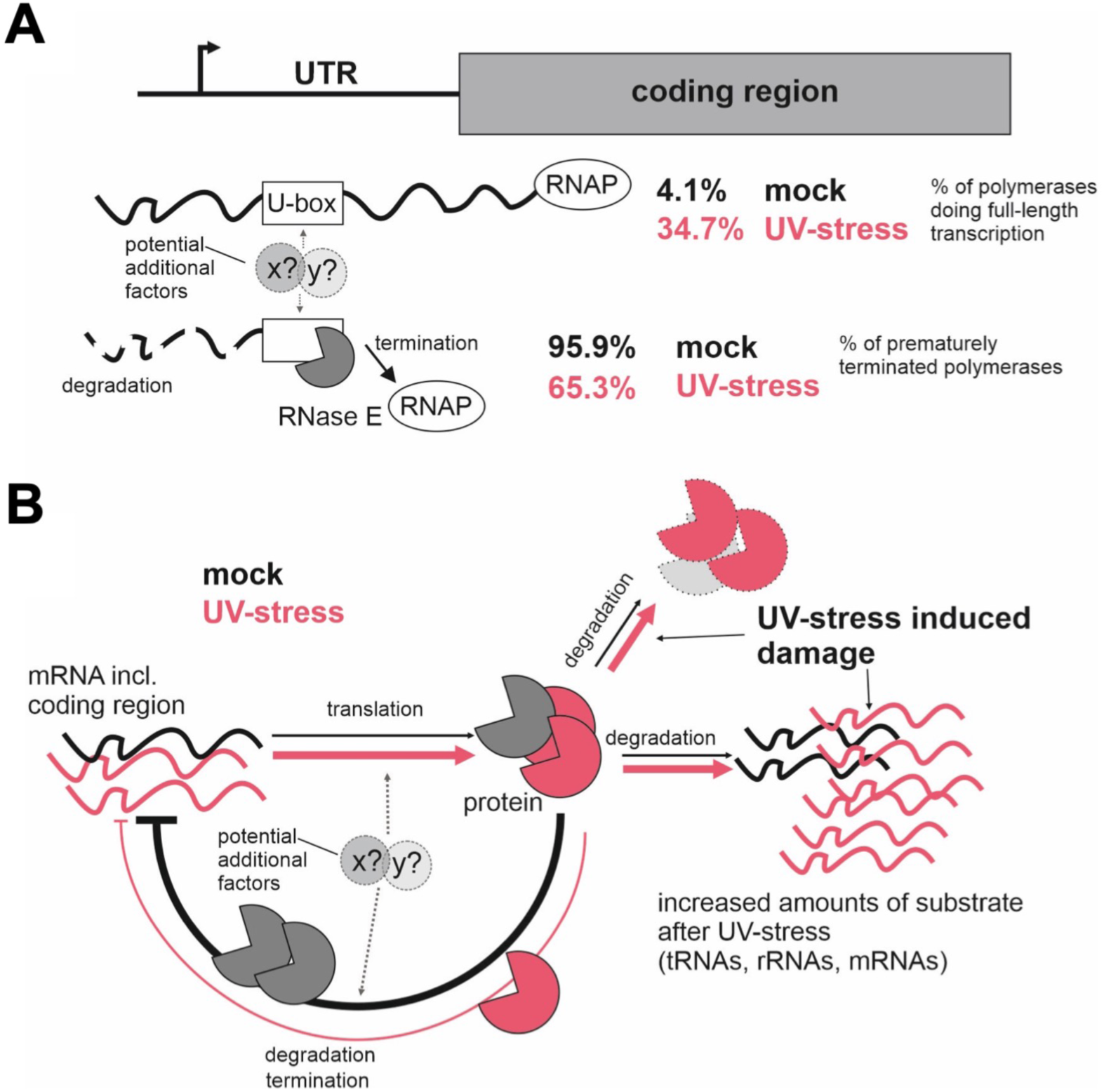
Hypothetical model of the regulation of RNase E expression in *Synechocystis* 6803. (A) The 5′ UTR of the *rne* message contains an RNase E-sensitive U-box. The (initial) endonucleolytic cleavage by RNase E destabilizes the UTR and might also lead to termination of transcription. Regardless of the exact molecular mechanism, termination is the main factor that accounts for differences in expression of the 5′ UTR and the coding region. Additional factors might be involved in these processes. In the absence of UV stress, 95.9% of all RNA polymerase molecules terminate at a point prior to the coding region, and only 4.1% transcribe the full-length message. Under UV stress conditions, the percentage of prematurely terminating polymerases decreases to 65.3%. (B) Hypothetical model of the autoregulatory negative feedback loop. Under normal growth conditions, the number of *rne* transcripts is maintained at a low level by a combination of RNA cleavage and transcription termination. When UV stress occurs, the number of alternative RNase E targets increases, and the stability of the RNase E protein is reduced. As a result, less RNase E activity is allocated to its own UTR, the autoregulation is relieved, and the system reaches a new equilibrium in which the concentrations of RNase E mRNA and protein are higher. For further details, see text.

Our proposed model for the feedback regulation of *Synechocystis* RNase E is shown in Figure 7B. Under nonstress conditions, the number of *rne* transcripts remains at a low level. This appears to be regulated by cleavage within the *rne* 5′ UTR U-rich region as well as by termination of transcript elongation. Following UV irradiation, degradation of UV-damaged RNase E protein is selectively accelerated, a process to which inactivation of RNase E by UV-induced irreversible crosslinking of RNase E and RNA may contribute. In parallel, we assume that the presence of UV-damaged RNAs drastically increases the substrate pool for RNase E. Consequently, less RNase E activity is allocated to its own mRNA, and this leads to relief of the feedback inhibition of *rne* full-length transcripts caused by stabilization of the *rne* 5′ UTR and reduced transcription termination prior to the ORF. As a result, more active RNase E is synthesized.

RNase E has a variety of functions, including degradation of damaged RNA and processing of newly synthesized RNA, and is considered to be particularly important in UV-damaged cells (61). The selective upregulation of *rne* transcripts that occurs during the UV stress response may be achieved by a comprehensive and cooperative system in the cell.

A similar control mechanism might also exist in other cyanobacteria. Like *Synechocystis* sp., *Anabaena* sp., *Synechococcus elongatus*, and even *Prochlorococcus* sp. have extremely long *rne* 5′ UTRs (Supplementary Figure S1A). An additional regulatory mechanism was reported in the marine cyanobacterium *Prochlorococcus* sp. MED4, in which RNase E levels were found to increase during lytic infection by the cyanophage P-SSP7; the increase in RNase E levels may support phage replication by generating a source of nucleotides from stimulated RNA degradation (62, 63). During phage infection, *rne* transcription proceeds from an alternative TSS, resulting in a shorter mRNA variant that lacks the regulatory 5′ UTR (63). These observations suggest that there is additional diversity in the control mechanisms of RNase E in cyanobacteria. While the mechanism described here ensures the presence of a certain level of RNase E, it does not explain the increased enzymatic activity we observed 2, 3 and 4 h after UV irradiation. Because there was no concomitant increase in the amount of RNase E protein, either the specific activity of the enzyme is enhanced by a regulatory factor or by protein modification or the fraction of active enzyme is higher among freshly synthesized enzymes than among the existing pool.

Our microarray analysis revealed changes in transcriptome composition after UV treatment over time, and these changes showed similarities and differences to those observed in other bacteria. When DNA is damaged by UV light, an SOS response is triggered, leading to DNA repair and the rebuilding of cellular components; this is accomplished by pausing cell division and energy production. Consistent with this widely conserved mechanism, we observed transient repression of genes involved in translation and energy metabolism one hour after UV irradiation of *Synechocystis* 6803. On the other hand, no significant induction of SOS gene homologs, such as *lexA*, *recA*, *uvrA*, *uvrB*, and *uvrC,* occurred; instead, a marked induction of *rne* and *rnj* was observed (Figure 1 and Supplementary Figure S3I). In *E. coli*, SOS-responsive genes were not induced in mutants of *rne* and *rng* (the latter encodes the RNase G homolog of *rne*), suggesting that RNase E is involved in the control of the SOS response (61). While the cellular response to UV has been studied in *Synechocystis* 6803 at the protein level (64), the regulatory mechanisms involved, including those that control the SOS response, are not clear in cyanobacteria. The functions of homologs of the *E. coli*-type SOS response regulator LexA are not conserved in several species of cyanobacteria (65). LexA has been reported to be unrelated to the regulation of the SOS response in *Synechocystis* 6803 (44, 66), in which it instead functions as a transcriptional regulator of fatty acid metabolism (67) and salt stress response (68). In addition, transcript levels of the *recA* gene, which plays a major role in DNA repair and recombination, are negatively regulated by UV light and oxidative stress at the posttranscriptional level (44) through a mechanism in which both RNase E and RNase J are thought to be involved. Further study of the posttranscriptional regulatory mechanisms involved in the cyanobacterial SOS response, including those related to RNase E, is needed.

In *Synechocystis* 6803, it has been reported that RNase E binds to a short hairpin RNA structure and then cleaves the RNA in an AU-rich region immediately downstream of the hairpin. Such sites have been described within the 5′ UTR of *psbA*, upstream of *crhR* (29, 34) and in maturation of crRNAs from long precursor RNA in a CRISPR–Cas subtype III-Bv system (35). Systematic mapping of RNase E sites, moreover, points to the frequent presence of a uridine 2 nt downstream of the cleavage site (+2U rule) and an adenine 3 or 4 nt upstream (-3/4A rule); together, these residues are capable of forming an “AU clamp” (36). We observed a very similar architecture of the U-rich site determined here (Figure 5B).

Further studies targeting proteins that may bind to the long *rne* 5′ UTR or modulate RNase E activity will be necessary to elucidate the detailed mechanism of the selective posttranscriptional upregulation of *rne* transcripts during the UV stress response. The widely distributed RNA chaperone Hfq modulates regulatory sRNA and target RNA structures and their interactions with each other (69); however, there is no evidence that Hfq binds RNA in *Synechocystis* 6803 (70, 71). RNA-binding proteins (Rbp) containing a single RNA recognition motif (RRM) have been identified in a number of cyanobacteria (72–75) and are also conserved among plant chloroplasts (76). In *Synechocystis* 6803, Rbp2 and Rbp3 are involved in the regulation of photosynthetic gene expression and thylakoid membrane targeting through transcript binding (77). In our study, the expression of these *rbp* genes was found to be induced at the same time as the expression of *rne* in response to UV stress (Supplementary Figure S3K-M), suggesting the involvement of Rbps in some aspects of transcriptome reshaping during the UV stress response.

In *E. coli*, RNA degradation and processing by RNase E is known to be modulated by RNase E-binding proteins such as RraA, RraB, and RapZ (20,78–80); however, no homologs of RraA or RraB have been characterized in cyanobacteria. Homologs of RapZ are found in certain cyanobacteria (*Synechococcales*, *Leptolyngbya*, *Gloeobacter*, and others) but are not conserved in *Synechocystis* species, and their function is still unknown.

In the cyanobacterium *Synechococcus elongatus* PCC 7942, the two heat shock proteins DnaK2 and DnaJ2 inhibit RNase E activity in an ATP-dependent manner, suggesting that both are involved in RNA degradation through interaction with RNase E (81).

The identification and functional characterization of proteins that interact with RNase E or its 5′ UTR is a promising topic for future research and should lead to a better understanding of the feedback regulation of RNase E and regulatory circuits within the RNA metabolism of cyanobacteria.

## Acknowledgments

We thank Matthias Kopf for helpful discussions and assistance with data analysis. This work was supported by Grants-in-Aid 25850056, S2306, and 20K05793 from the Ministry of Education, Culture, Sports, Science and Technology of Japan, the Advanced Low Carbon Technology Research and Development Program (ALCA) of the Japan Science and Technology Agency (JST) (to S.W.), the German Ministry of Education and Research (BMBF), grant no. 031L0164B “RNAProNet” (to W.R.H.) and the German Science Foundation (DFG) research training group MeInBio 322977937/GRK2344 (to A.W. and W.R.H.) and grant STE 1119/4-2 (to C.S.).

## Supplementary Figures

**Supplementary Figure S1.**
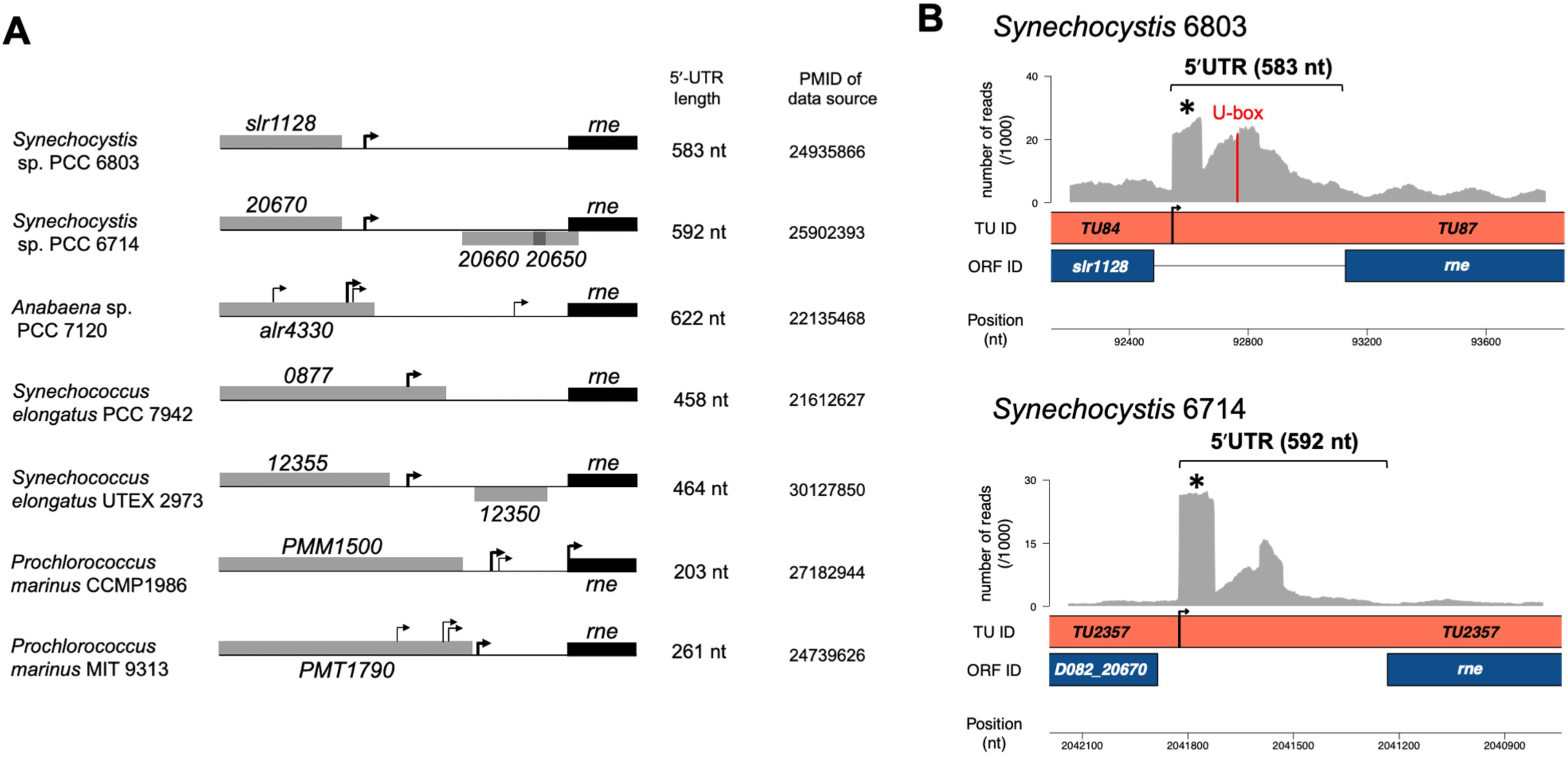
Long 5′ UTRs of the *rne* gene in cyanobacteria. (A) Comparison of *rne* 5′ UTRs in different cyanobacteria. Transcription start sites (TSSs, arrows) and upstream regions of the *rne* in several species are illustrated, as is the length of the *rne* 5′ UTR. Information on the TSS in each organism was obtained from comprehensive experimental data, including TSS-seq analysis. The bold arrows indicate major TSSs. (B) The *rne* genomic locus in *Synechocystis* 6803 and *Synechocystis* 6714 with coverage by RNA-seq data extracted from previous analyses (58). Protein-encoding genes are shown in blue, and transcription units (TUs) are shown in red. The graphs shown in gray represent the coverage by RNA-seq reads. The y-axes of the graphs indicate square-root-scaled coverage values, and the x-axes show the chromosomal position in bp. A red bar indicates the location of the U-box in the *rne* 5′ UTR. Asterisks indicate the highly expressed *rne* 5′ UTRs in *Synechocystis* 6803, annotated as sRNA *ncr0020* (Figure 1B).

**Supplementary Figure S2.**
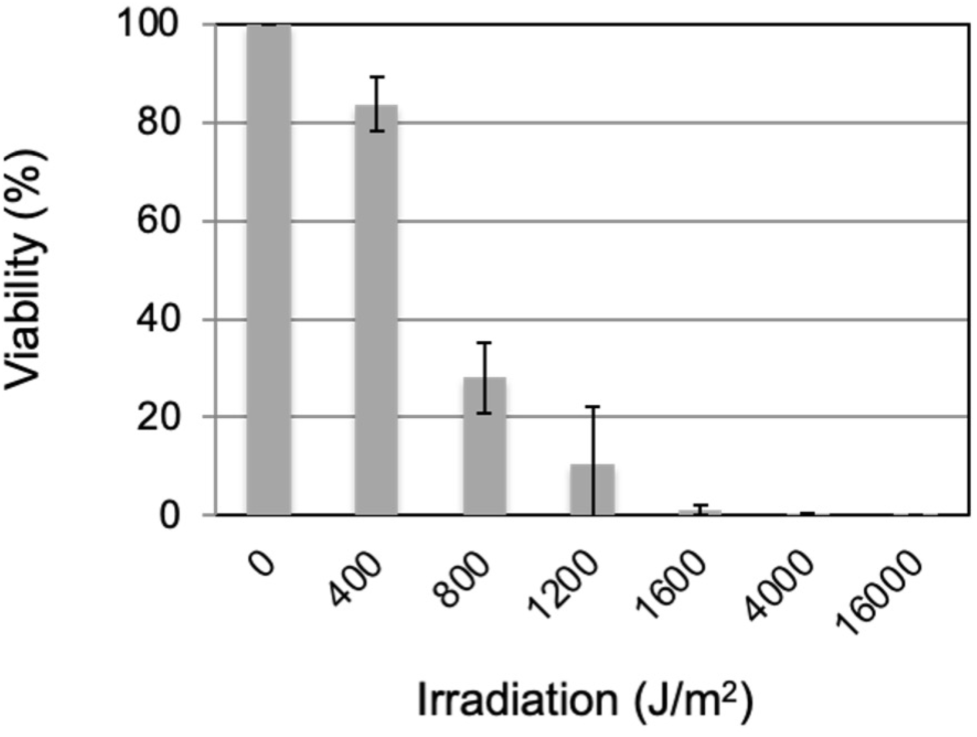
Survival of *Synechocystis* following UV-C irradiation. One hundred milliliters of exponentially growing cells (O.D._750_ = 0.8) were transferred to a petri dish, and the cells were irradiated with UV-C (254 nm) at a dose of 200 µW/cm^2^, leading to the indicated amounts of total irradiation. Cells were harvested at various times and tested in the viability assay. The cells were spread on solid medium before and after UV-C irradiation. Surviving colonies were counted after 7 days. The data shown are the mean± SD of values obtained in triplicate experiments.

**Supplementary Figure S3.**
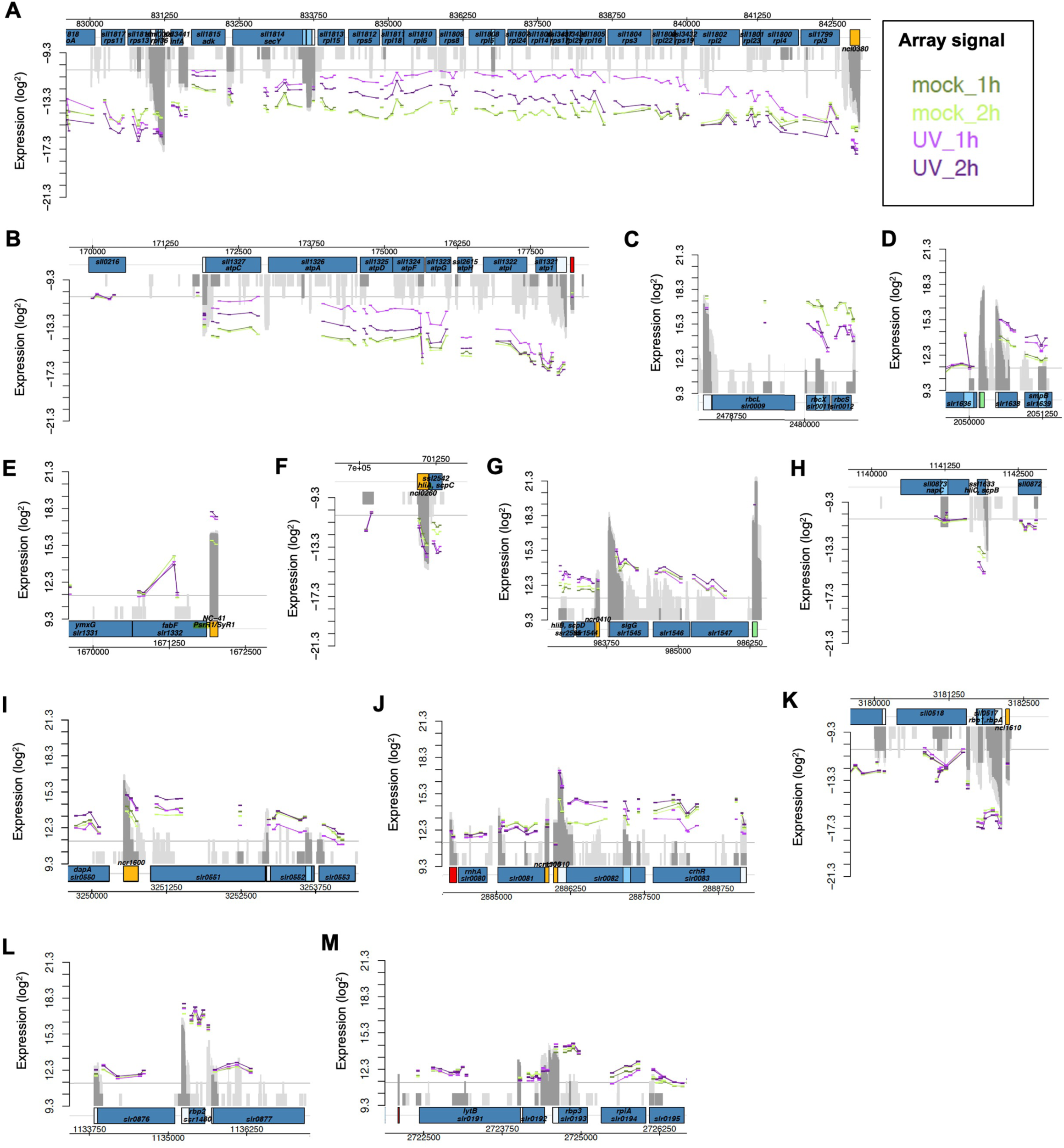
UV stress response in *Synechocystis* 6803. Genomic loci of significantly up- or downregulated genes were selected and are shown with the respective mapped array probes (vertical tabs). All data are available in Supplementary Data 2. The signal intensities are given as log_2_ values. (A) Operon of genes encoding ribosomal proteins (*sll1799*-*sll1817*) and *ncl0380*; (B) *atp* operon (*sll1321*-*sll1327*); (C) *rbc* operon (*slr0009*-*slr0012*); (D) *smpB* (*slr1639*); (E) *psrR1*; (F) *hliA* (*ssl2542*); (G) *hliB* (*ssr2595*); (H) *hliC* (*ssl1633*); (I) *rnj* (*slr0551*); (J) *rimO* (*slr0082*)-*crhR* operon; (K) *rbp1* (*sll0517*); (L) *rbp2* (*ssr1480*); and (M) *rbp3* (*slr0193*).

**Supplementary Figure S4.**
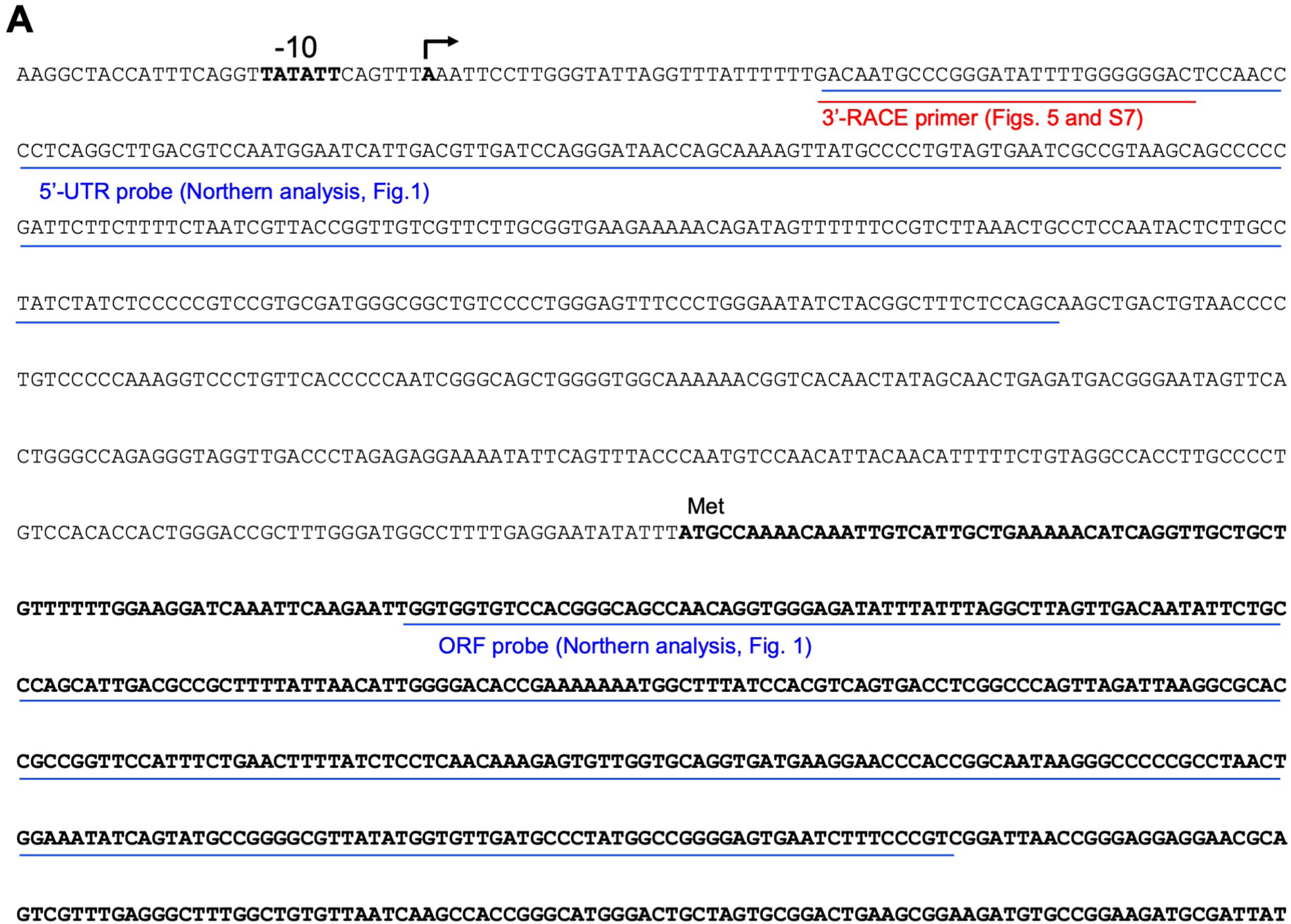
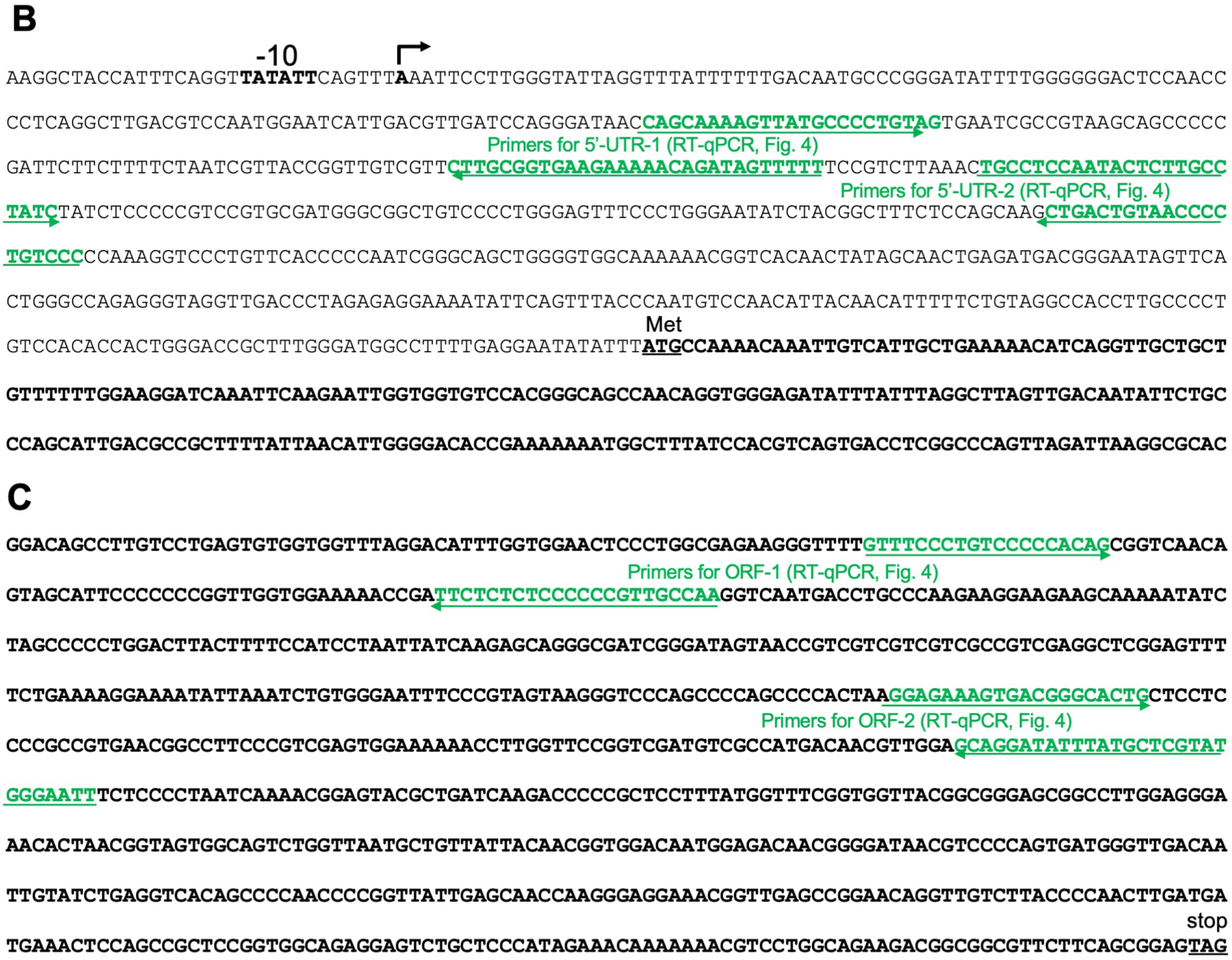
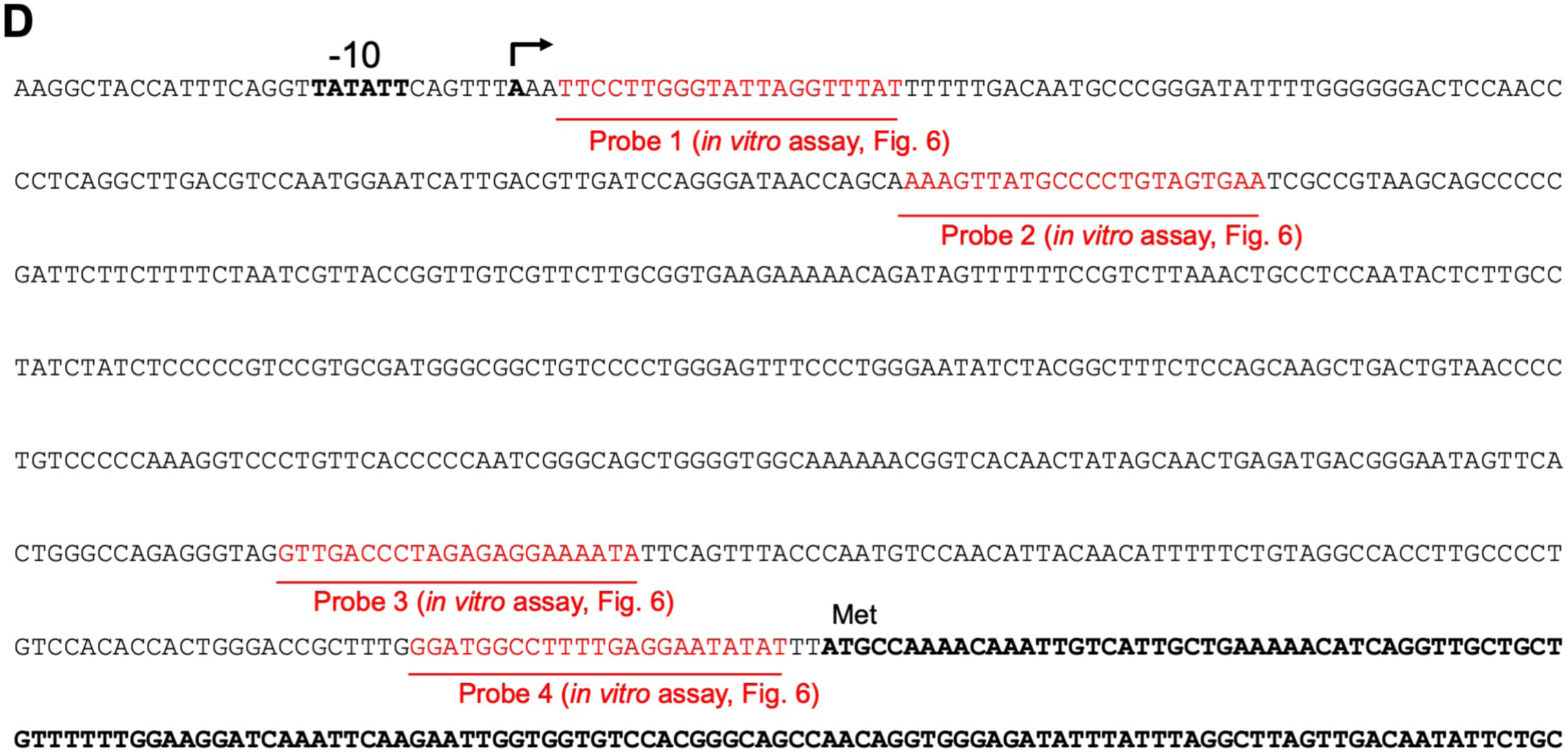
Locations of the probes and primers used in this study. (A) The positions of the probes (5′ UTR and ORF, underlined in blue) used in the northern analysis (Figure 1C) and the primer used in 3′ RACE (underlined in red, Figure 5 and Supplementary Figure S7) within the sequence of the *rne* (*slr1129*) ORF and its upstream region are shown. The promoter (-10), TSS, and ORF region (1-515 bp) of *rne* are shown in bold type. (B, C) The positions of the primers used in RT–qPCR (Figure 4) are shown as green arrows along the sequence of the *rne* upstream region (B) and its ORF (C). (D) The positions of probes used in the *in vitro* cleavage assay (probes 1-4, Figure 6) are underlined in red.

**Supplementary Figure S5.**
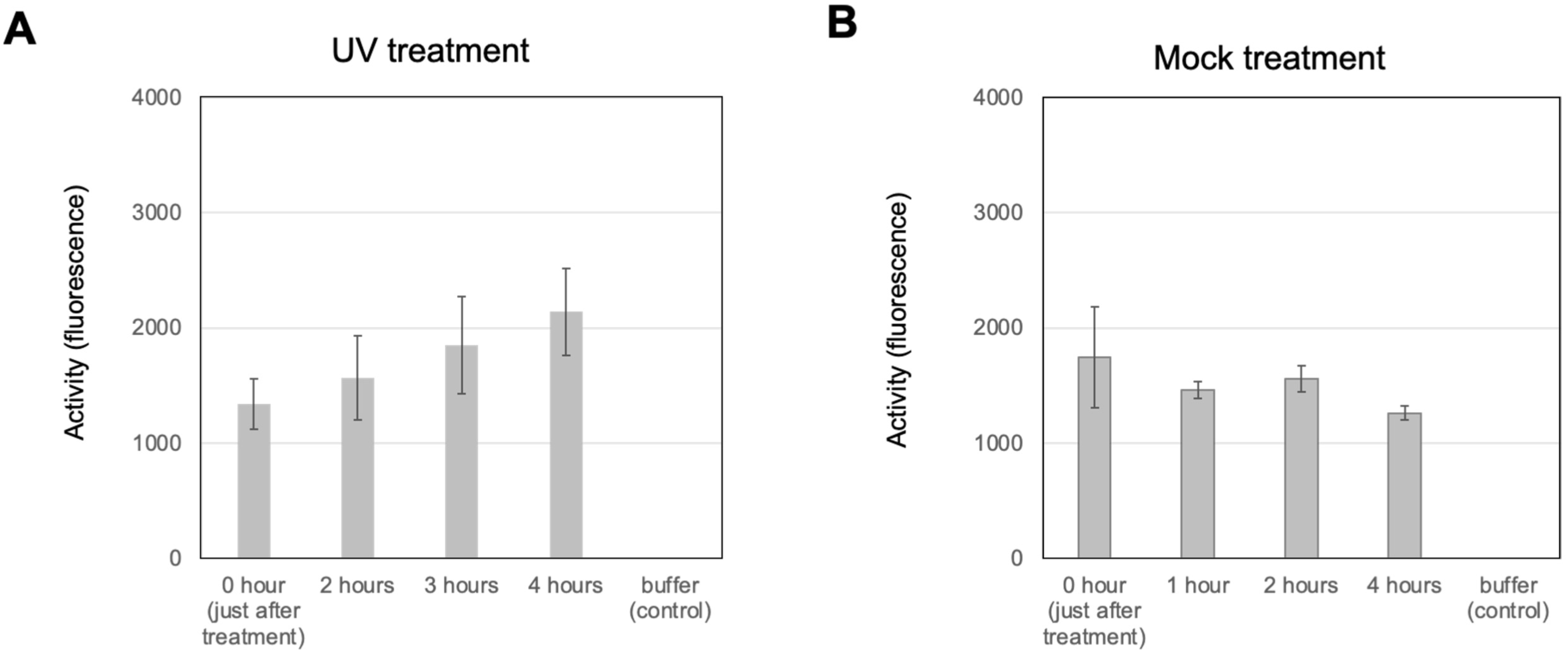
Comparison of RNase E activity values after 15 minutes of incubation. Crude extracts containing 15 µg of protein were prepared from cultures after UV (A) or mock treatment (B). The standard deviations in (A) were calculated from 3 biological replicates, while those in (B) were derived from technical triplicates.

**Supplementary Figure S6.**
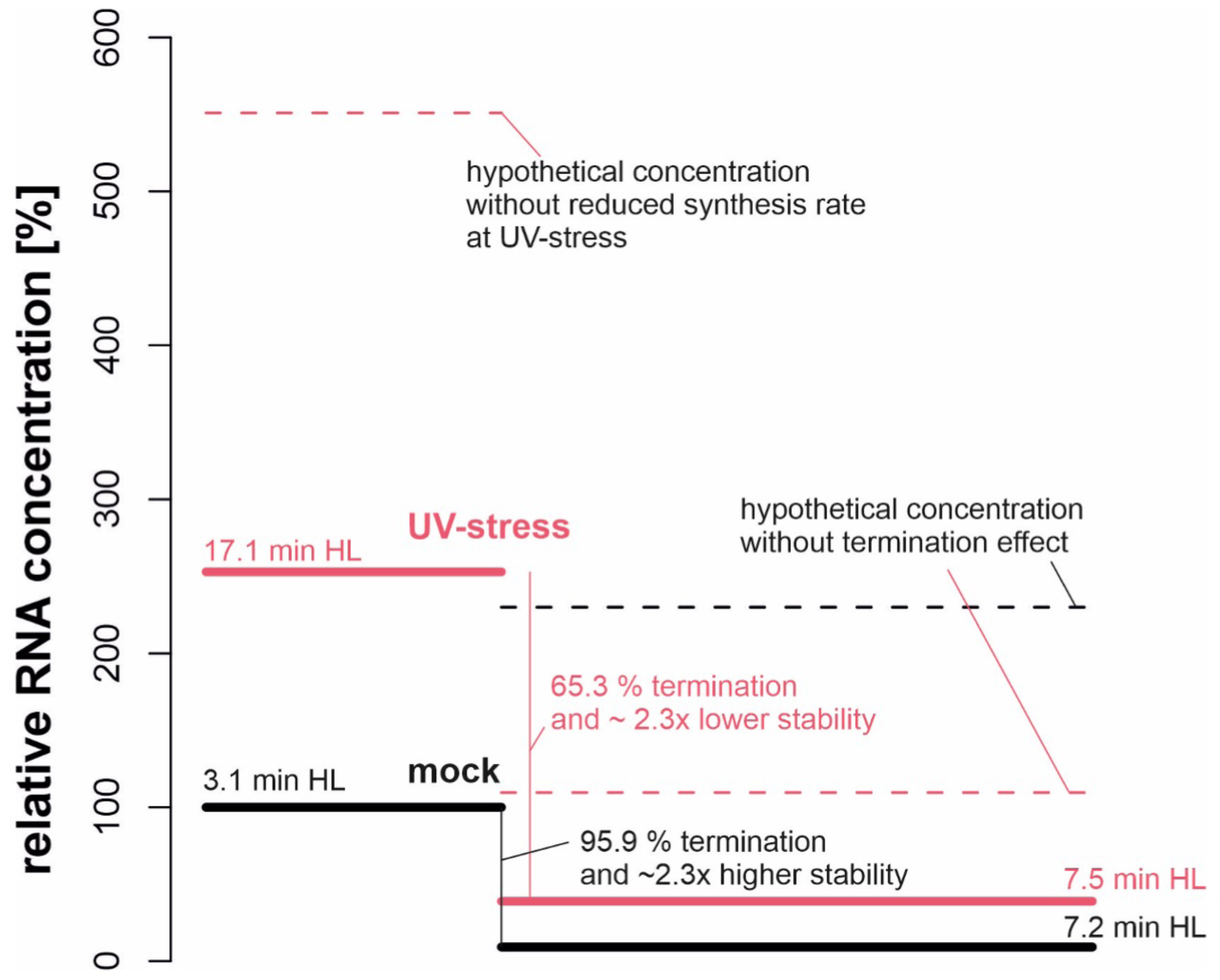
Schematic representation of the *rne* 5′ UTR and coding region transcript concentrations under mock and UV stress conditions, together with their respective half-lives (HL). The concentration of the 5′ UTR under mock conditions was set to 100%. The broken line in the 5′ UTR indicates the hypothetical RNA concentration under UV stress conditions if the synthesis rate remained unchanged and only the stability of the protein increased. The broken lines in the coding region indicate the hypothetical transcript concentrations based on differences in stability without premature termination.

**Supplementary Figure S7.**
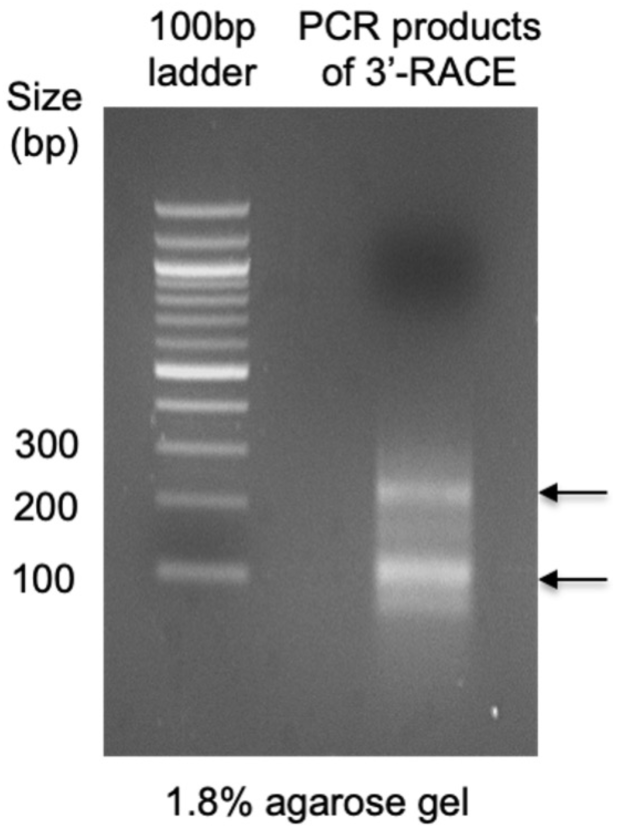
Electrophoresis gel image of the PCR products used in the 3′ RACE assay. RNA samples prepared from *Synechocystis* cultures were ligated to 3′- adaptor RNAs and reverse-transcribed using a primer specific for the 3′ adaptor. Using the resulting cDNA products as a template, DNA fragments were PCR-amplified using the primer set slr1129-5UTR-f and 3RACE_Tm55 (Supplementary Figure S4A and Table S1) and subjected to electrophoretic separation on an agarose gel.

**Supplementary Figure S8.**
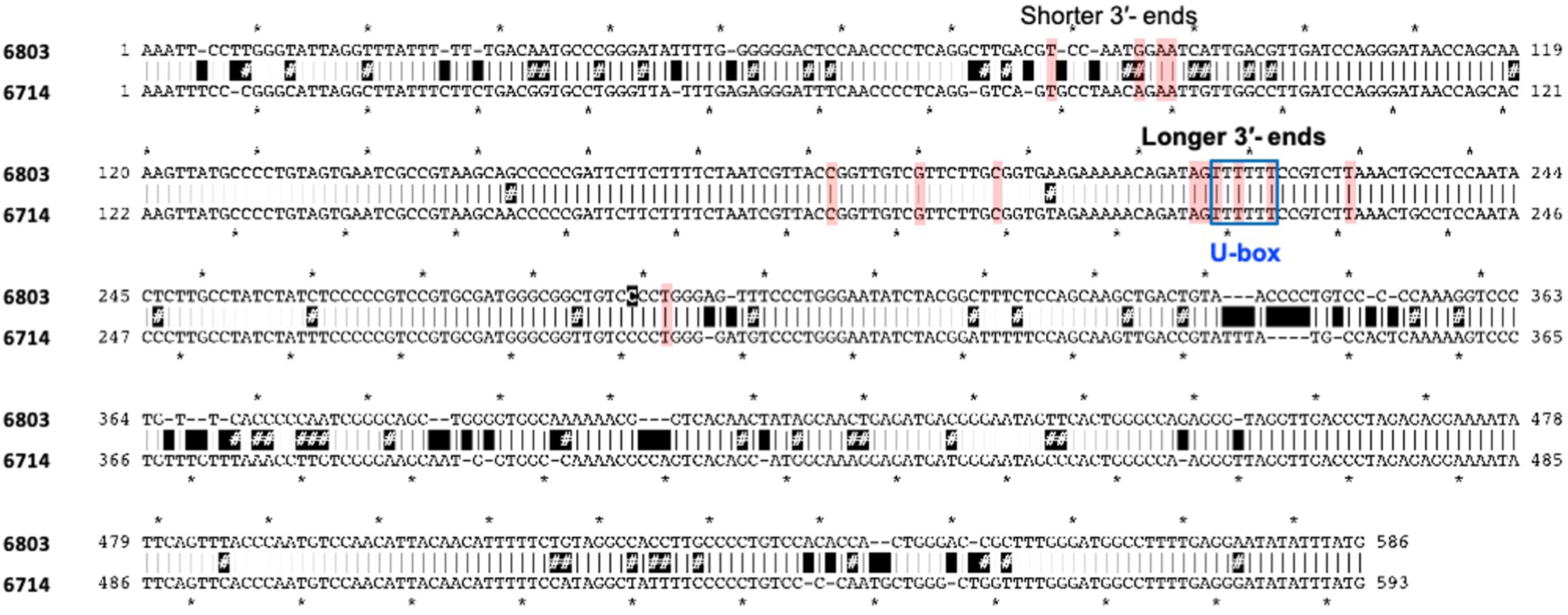
Alignment of *rne* 5′ UTRs in *Synechocystis* 6803 (6803) with those in *Synechocystis* 6714 (6714). The 3′ ends of the 5′ UTR transcripts identified by 3′ RACE are indicated by the red boxes. The U-rich region is boxed in blue. Mismatches between the two sequences are indicated by black boxes with hashtags.

**Supplementary Data 1.** Comparative microarray data for the UV stress response compared to mock conditions in *Synechocystis* 6803 cells. Transcripts are categorized into mRNAs (labeled with their respective gene IDs), antisense RNAs (labeled “as”), potentially trans-encoded sRNAs, 5′ UTRs and transcripts derived from gene-internal segments (labeled “int”). The table displays log_2_ FCs in transcript abundance under the compared conditions (UV_1 h versus Mock 1 h and UV_2 h versus Mock 2 h).

**Supplementary Data 2.** Transcriptomic response to UV treatment. Detailed genomic view with array probes indicated by vertical bars connected by colored lines. The signal intensities are given as log_2_ values. The graphs shown in gray represent RNA sequencing data given as log_2_ read numbers; these were extracted from the previous genome-wide mapping of TSSs (82).

## Supplementary Table

**Supplementary Table S1.**
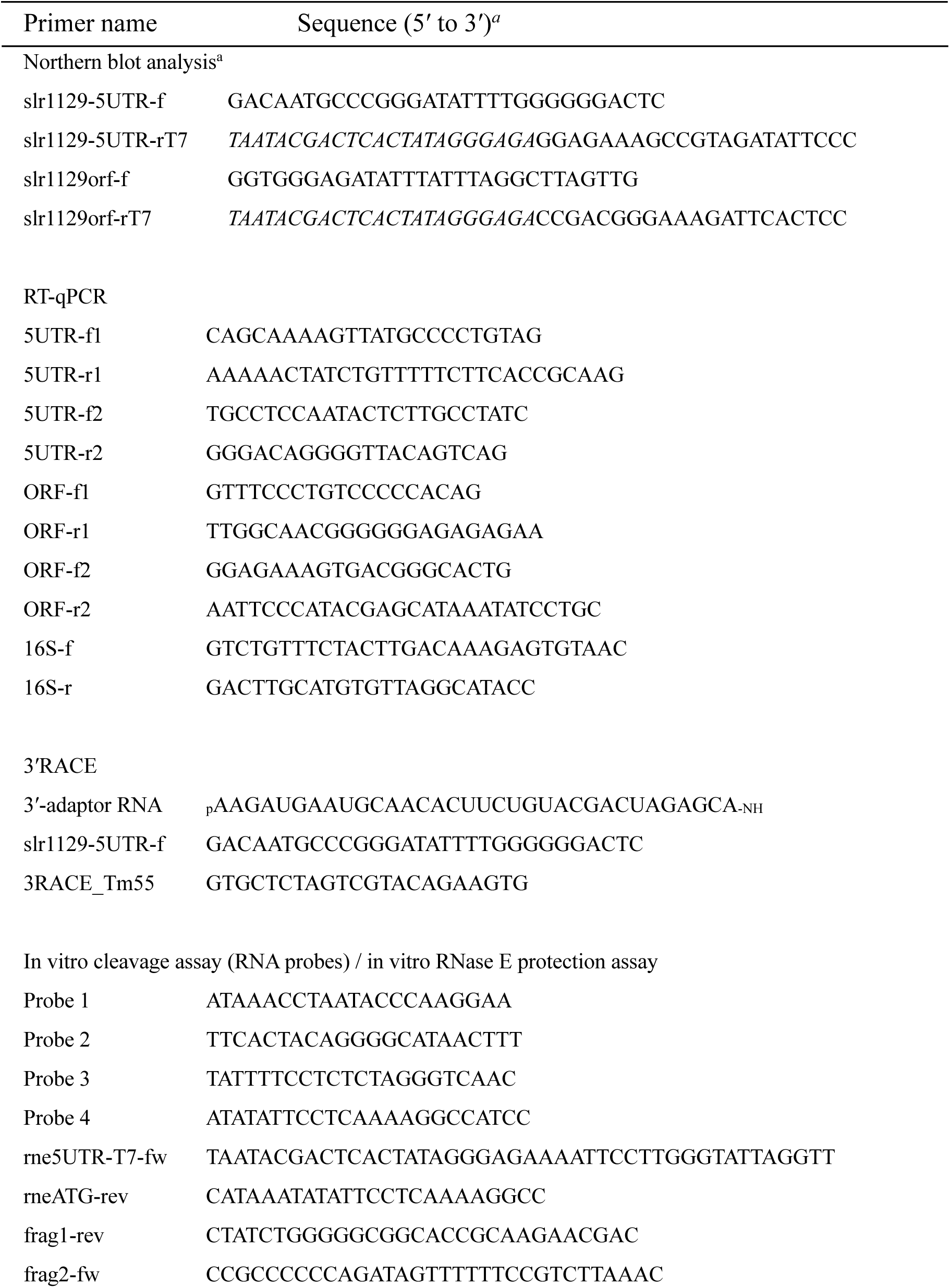
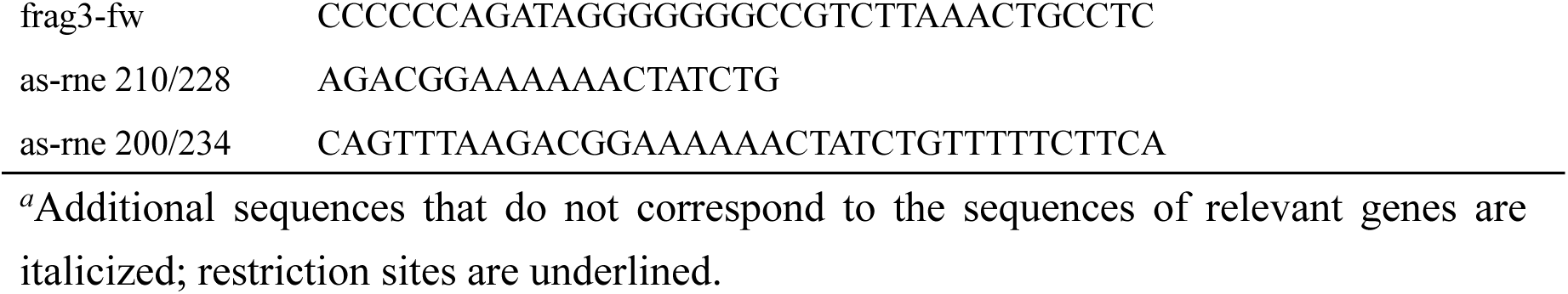
Oligonucleotide primers used in this study. Primer name Sequence (5′ to 3′)*^a^*

