## Supplementary material for "Feedback regulation of RNase E during UV-stress response in the cyanobacterium *Synechocystis* sp. PCC 6803": Supplemental DataSet 2.pdf

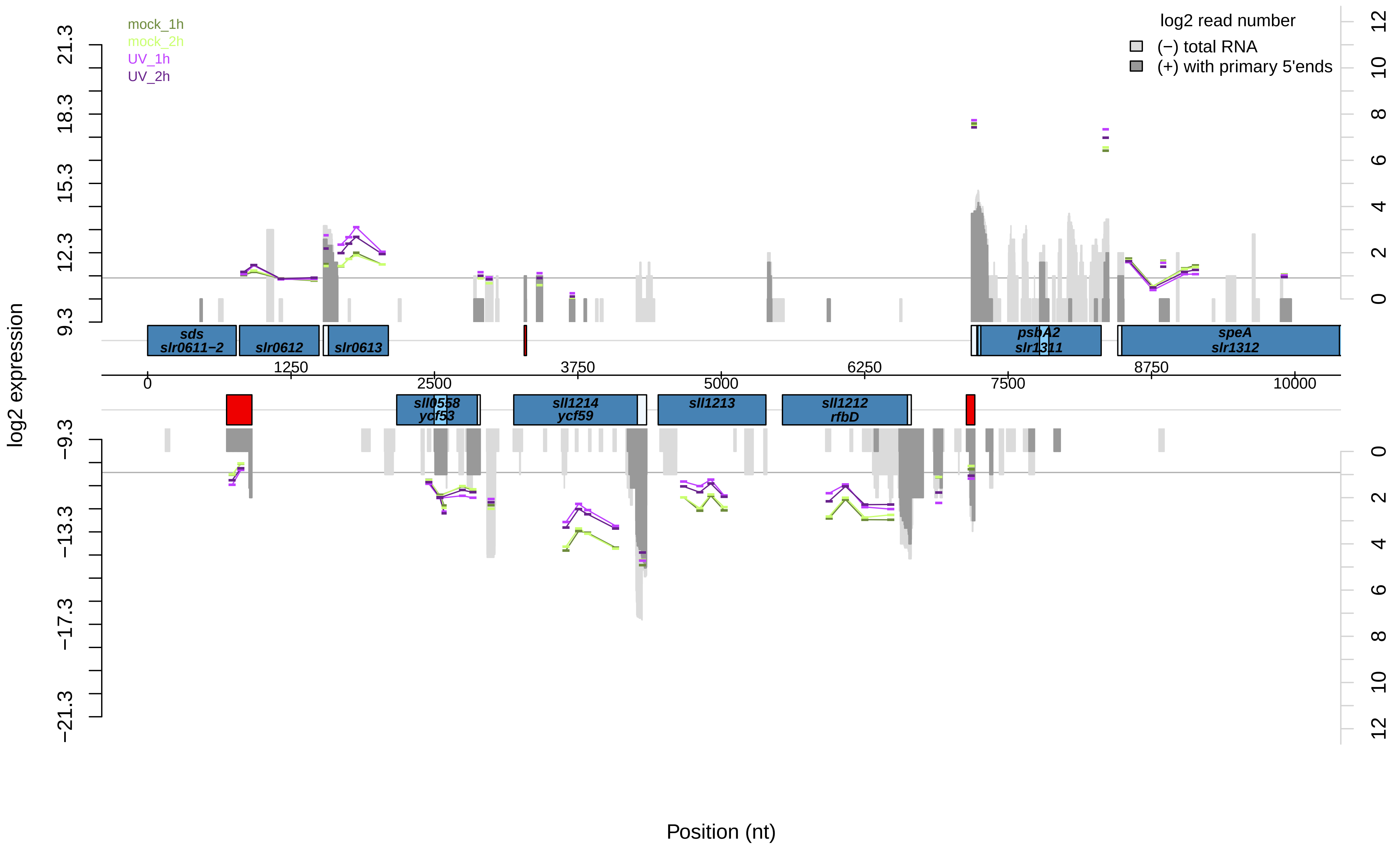

mock_1h
mock_2h
UV_1h
UV_2h
log2 read number
(−) total RNA
(+) with primary 5'ends
log2 expression
21.3
18.3
15.3
12.3
9.3
−9.3
−13.3
−17.3
−21.3
12
10
8
6
4
2
0
sds
slr0611−2
slr0612
slr0613
psbA2
slr1311
speA
slr1312
0
1250
2500
3750
5000
6250
7500
8750
10000
sll0558
ycf53
sll1214
ycf59
sll1213
sll1212
rfbD
Position (nt)

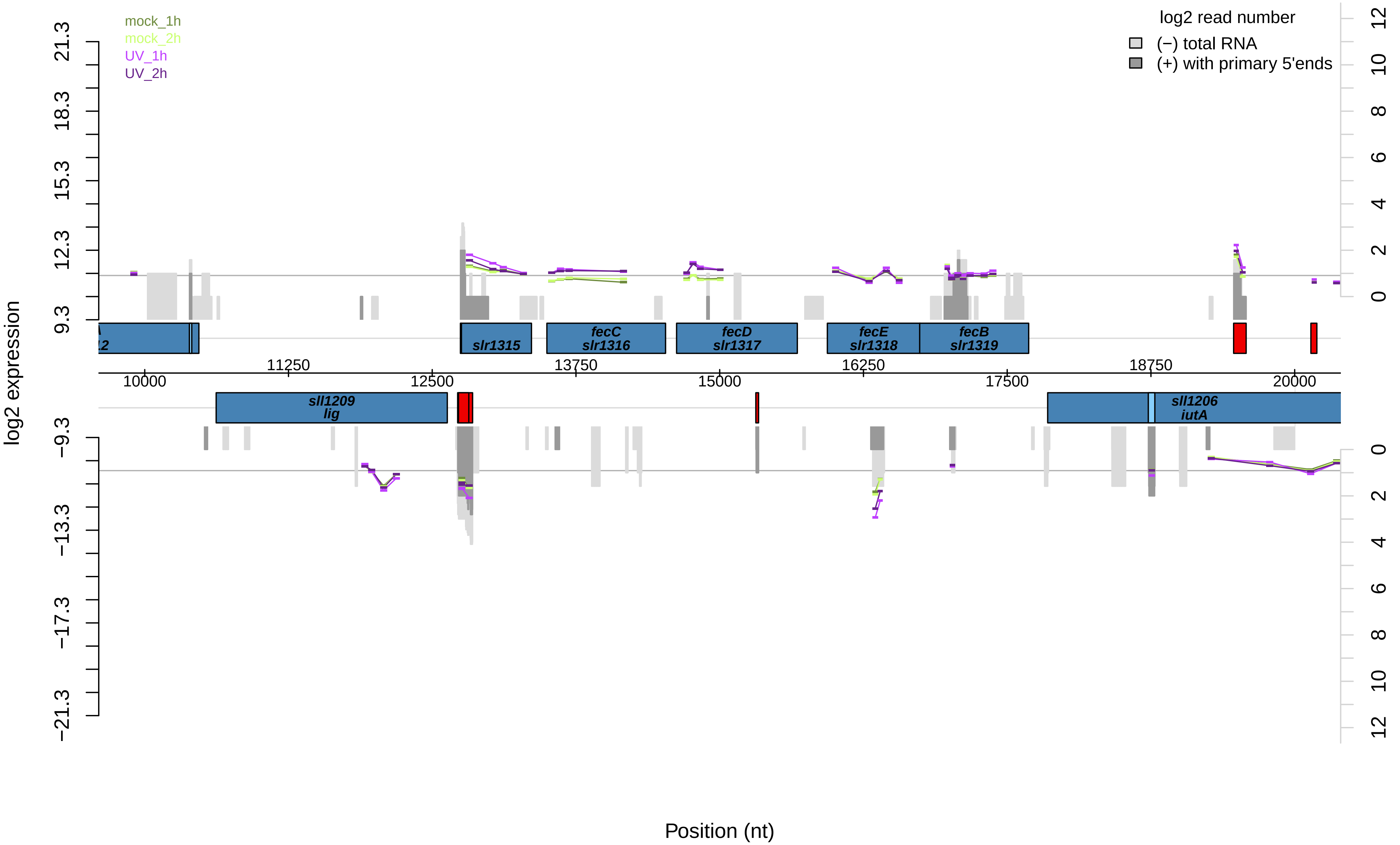

mock_1h
mock_2h
UV_1h
UV_2h
log2 read number
(−) total RNA
(+) with primary 5'ends
log2 expression
21.3
18.3
15.3
12.3
9.3
−9.3
−13.3
−17.3
−21.3
12
10
8
6
4
2
0
slr1315
fecC
slr1316
fecD
slr1317
fecE
slr1318
fecB
slr1319
sll1209
lig
sll1206
iutA
11250
13750
16250
18750
10000
12500
15000
17500
20000
Position (nt)

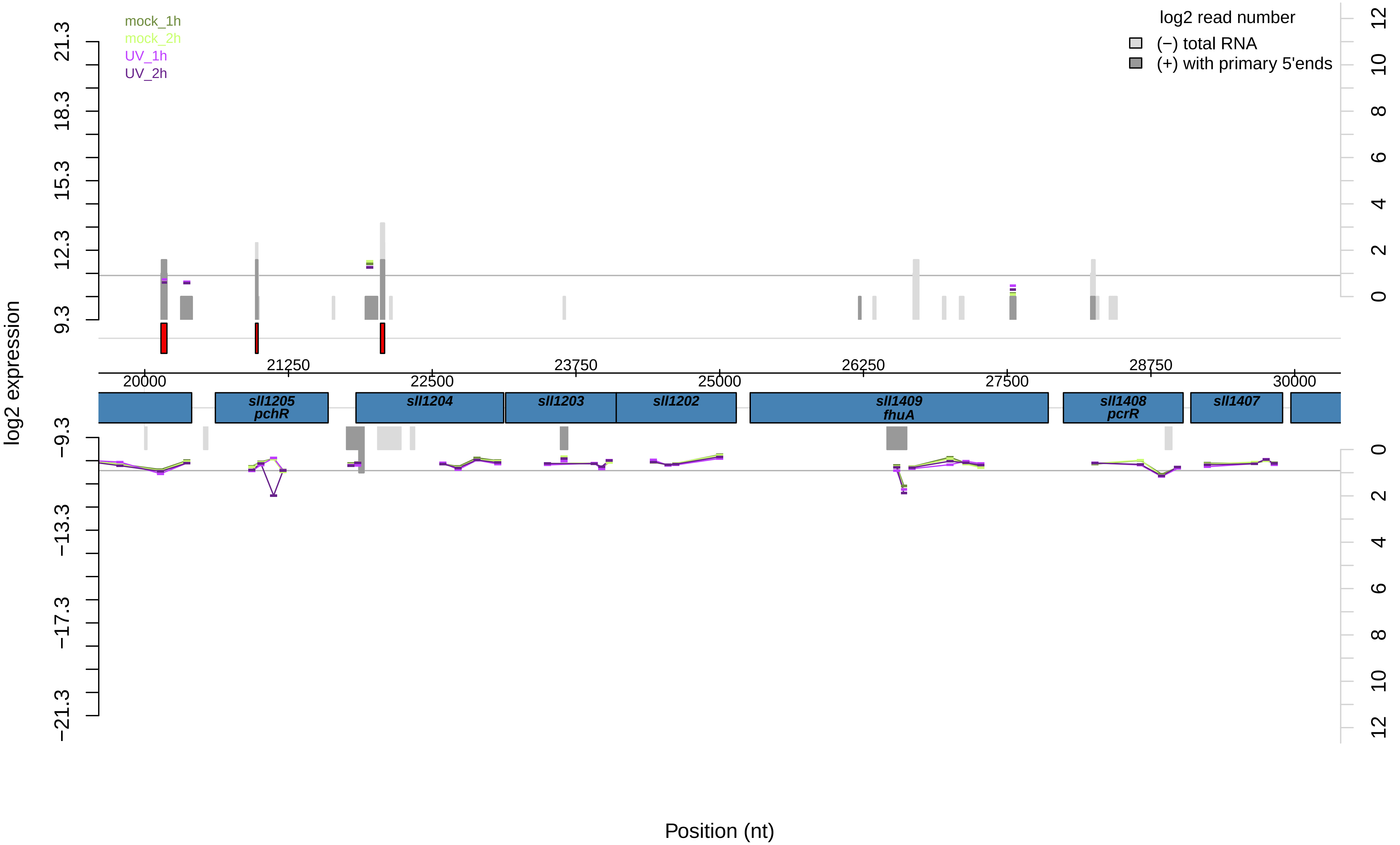

mock_1h
mock_2h
UV_1h
UV_2h
log2 read number
(−) total RNA
(+) with primary 5'ends
log2 expression
21.3
18.3
15.3
12.3
9.3
−9.3
−13.3
−17.3
−21.3
12
10
8
6
4
2
0
20000
21250
22500
23750
25000
26250
27500
28750
30000
sll1205
pchR
sll1204
sll1203
sll1202
sll1409
fhuA
sll1408
pcrR
sll1407
Position (nt)

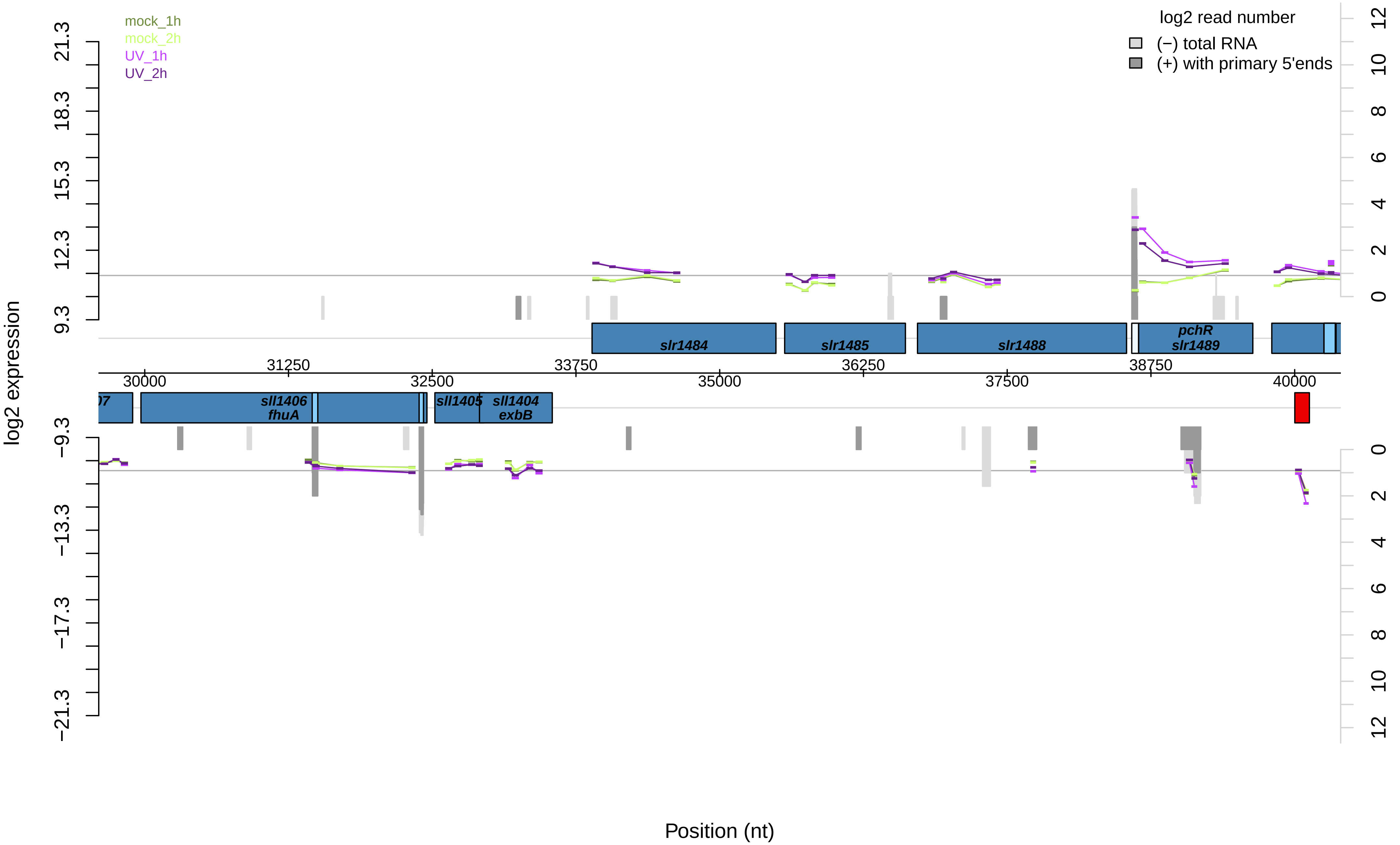

mock_1h
mock_2h
UV_1h
UV_2h
log2 read number
(−) total RNA
(+) with primary 5'ends
log2 expression
21.3
18.3
15.3
12.3
9.3
−9.3
−13.3
−17.3
−21.3
12
10
8
6
4
2
0
slr1484
slr1485
slr1488
pchR
slr1489
31250
33750
36250
38750
30000
32500
35000
37500
40000
07
sll1406
fhuA
sll1405
sll1404
exbB
Position (nt)

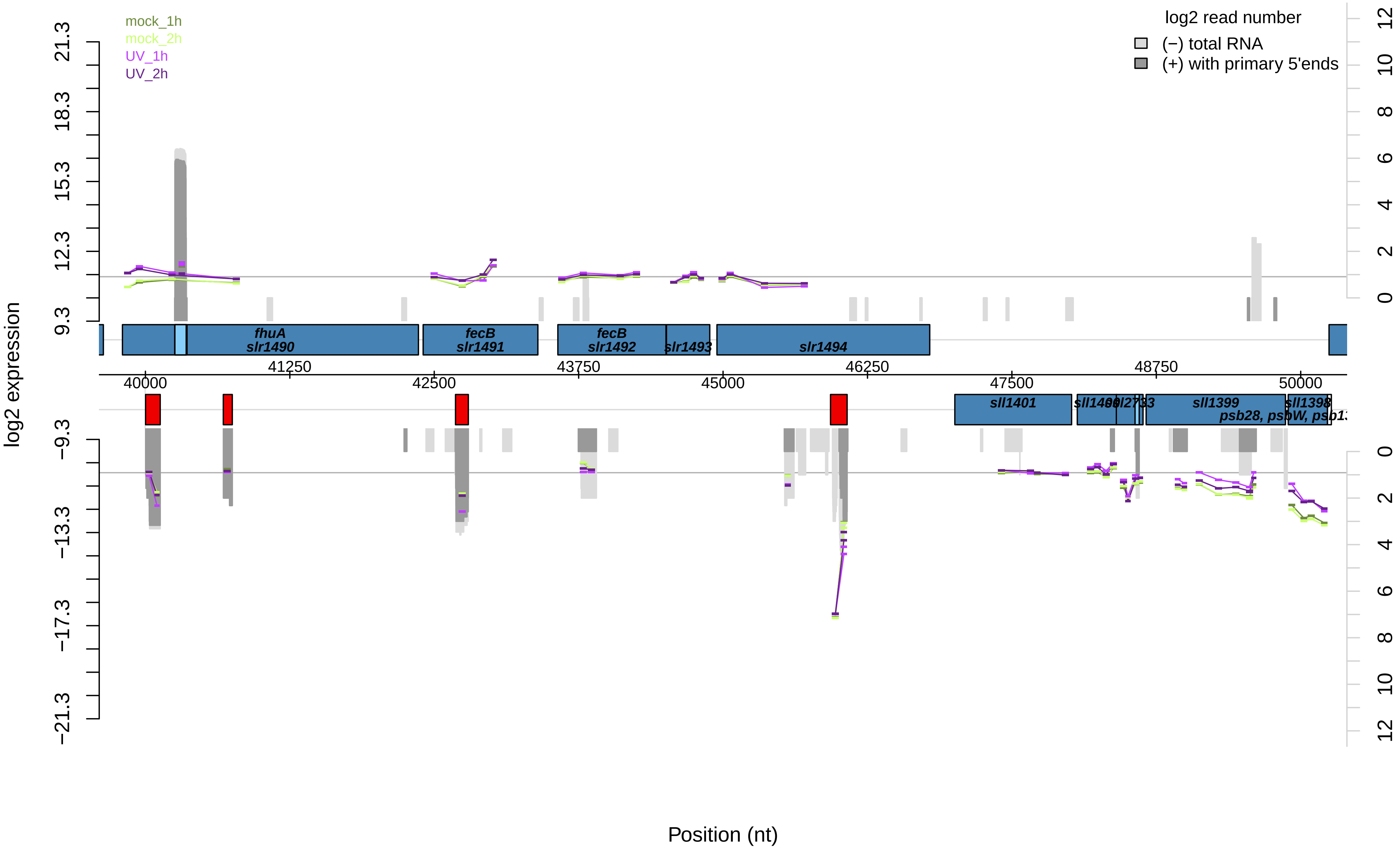

mock_1h
mock_2h
UV_1h
UV_2h
log2 read number
(−) total RNA
(+) with primary 5'ends
log2 expression
21.3
18.3
15.3
12.3
9.3
−9.3
−13.3
−17.3
−21.3
fhuA
slr1490
fecB
slr1491
fecB
slr1492
slr1493
slr1494
41250
43750
46250
48750
40000
42500
45000
47500
50000
sll1401
sll1399
psb28, psbW, psb1
sll1398
Position (nt)

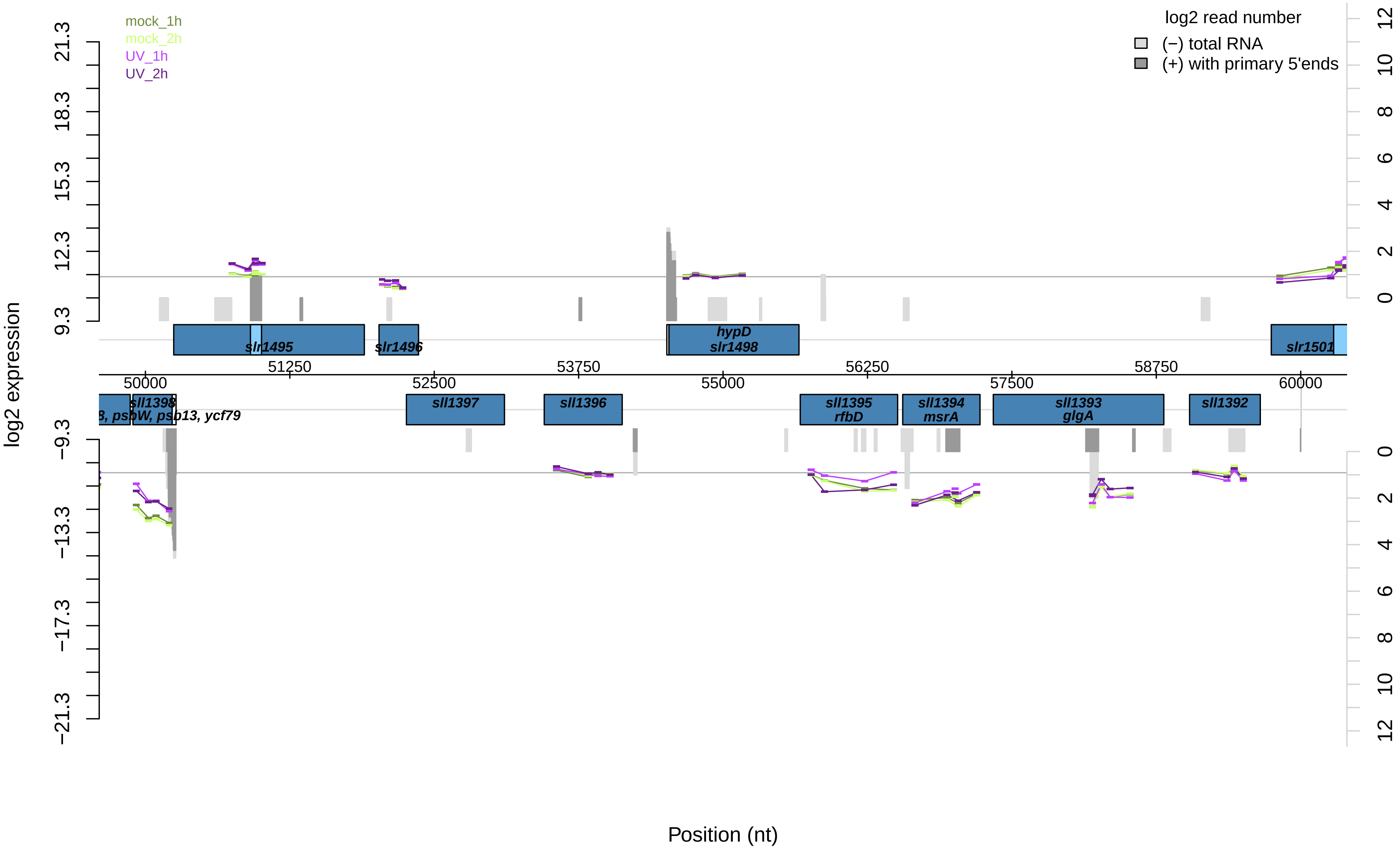

mock_1h
mock_2h
UV_1h
UV_2h
log2 read number
(−) total RNA
(+) with primary 5'ends
log2 expression
21.3
18.3
15.3
12.3
9.3
−9.3
−13.3
−17.3
−21.3
12
10
8
6
4
2
0
slr1495
slr1496
hypD
slr1498
slr1501
51250
53750
56250
58750
50000
52500
55000
57500
60000
sll1398
8, psbW, psb13, ycf79
sll1397
sll1396
sll1395
rfbD
sll1394
msrA
sll1393
glgA
sll1392
Position (nt)

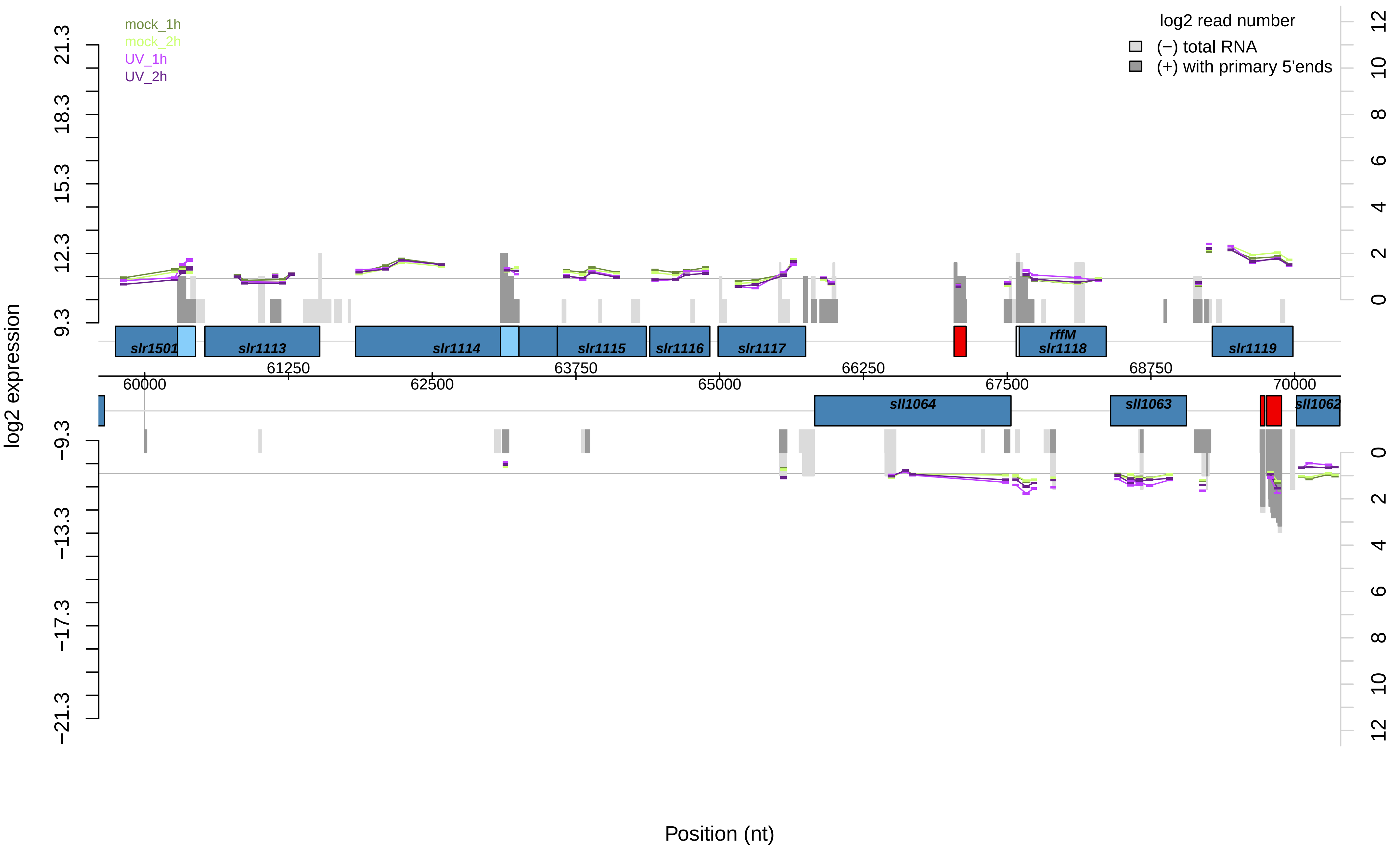

mock_1h
mock_2h
UV_1h
UV_2h
log2 read number
(−) total RNA
(+) with primary 5'ends
log2 expression
21.3
18.3
15.3
12.3
9.3
−9.3
−13.3
−17.3
−21.3
12
10
8
6
4
2
0
slr1501
slr1113
slr1114
slr1115
slr1116
slr1117
rffM
slr1118
slr1119
61250
63750
66250
68750
60000
62500
65000
67500
70000
sll1064
sll1063
sll1062
Position (nt)

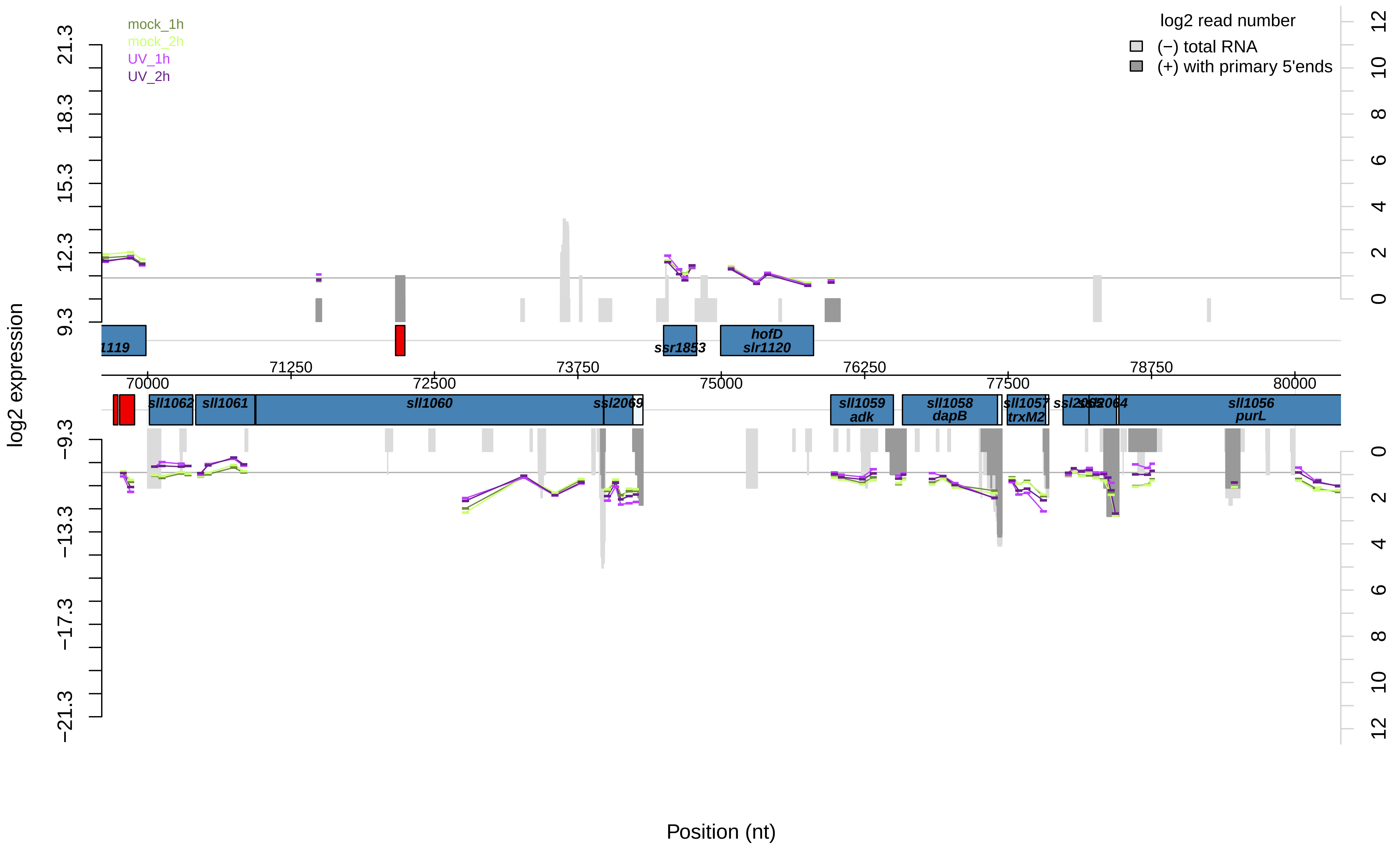
mock_1h
mock_2h
UV_1h
UV_2h
log2 read number
(−) total RNA
(+) with primary 5'ends
log2 expression
21.3
18.3
15.3
12.3
9.3
−9.3
−13.3
−17.3
−21.3
12
10
8
6
4
2
0
1119
ssr1853
hofD
slr1120
71250
73750
76250
78750
70000
72500
75000
77500
80000
sll1062
sll1061
sll1060
ssl2069
sll1059
adk
sll1058
dapB
sll1057
trxM2
sll2064
sll1056
purL
Position (nt)

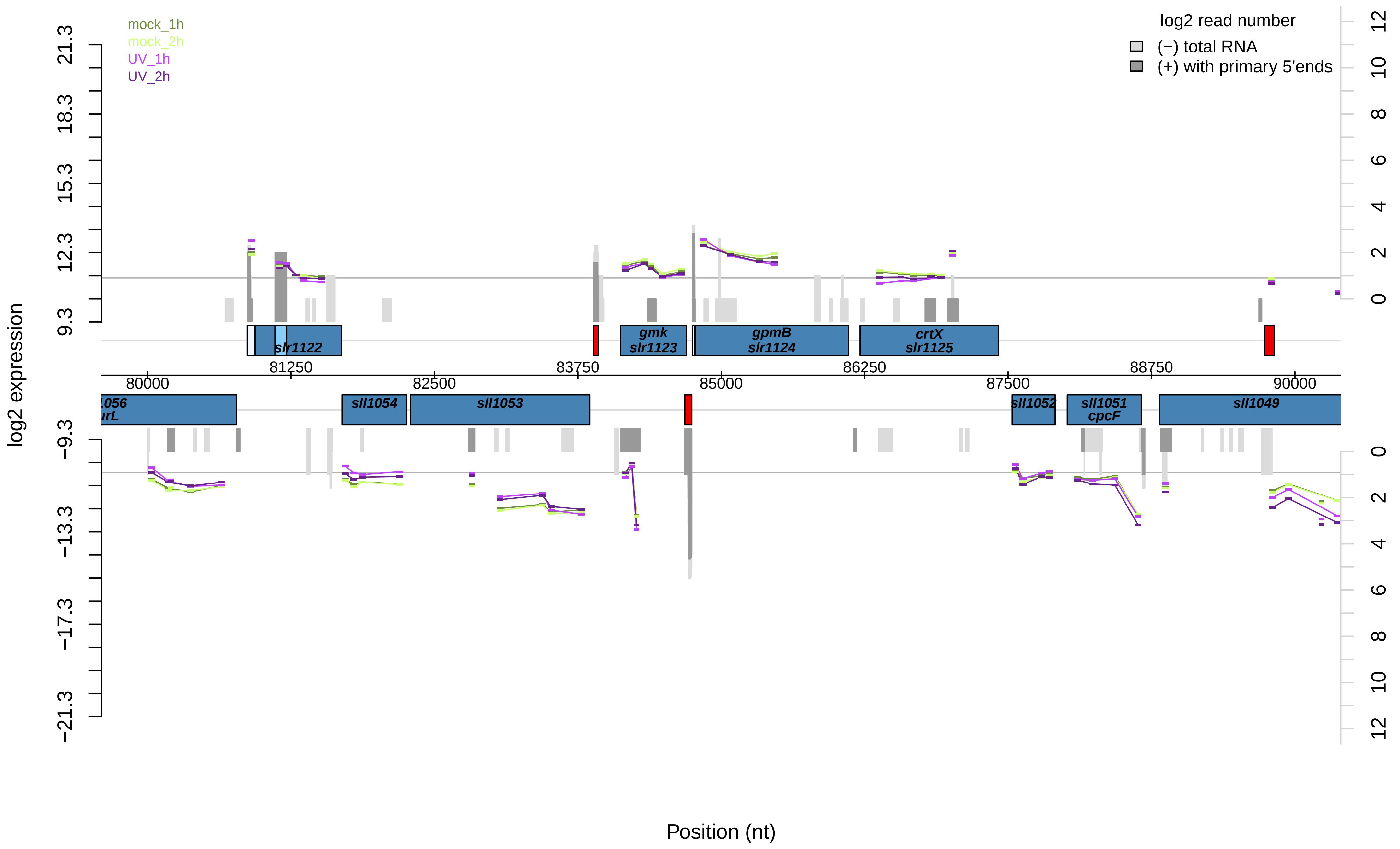

mock_1h
mock_2h
UV_1h
UV_2h
log2 read number
(−) total RNA
(+) with primary 5'ends
log2 expression
21.3
18.3
15.3
12.3
9.3
−9.3
−13.3
−17.3
−21.3
12
10
8
6
4
2
0
slr1122
gmk
slr1123
gpmB
slr1124
crtX
slr1125
81250
83750
86250
88750
80000
82500
85000
87500
90000
sll1054
sll1053
sll1052
sll1051
cpcF
sll1049
Position (nt)

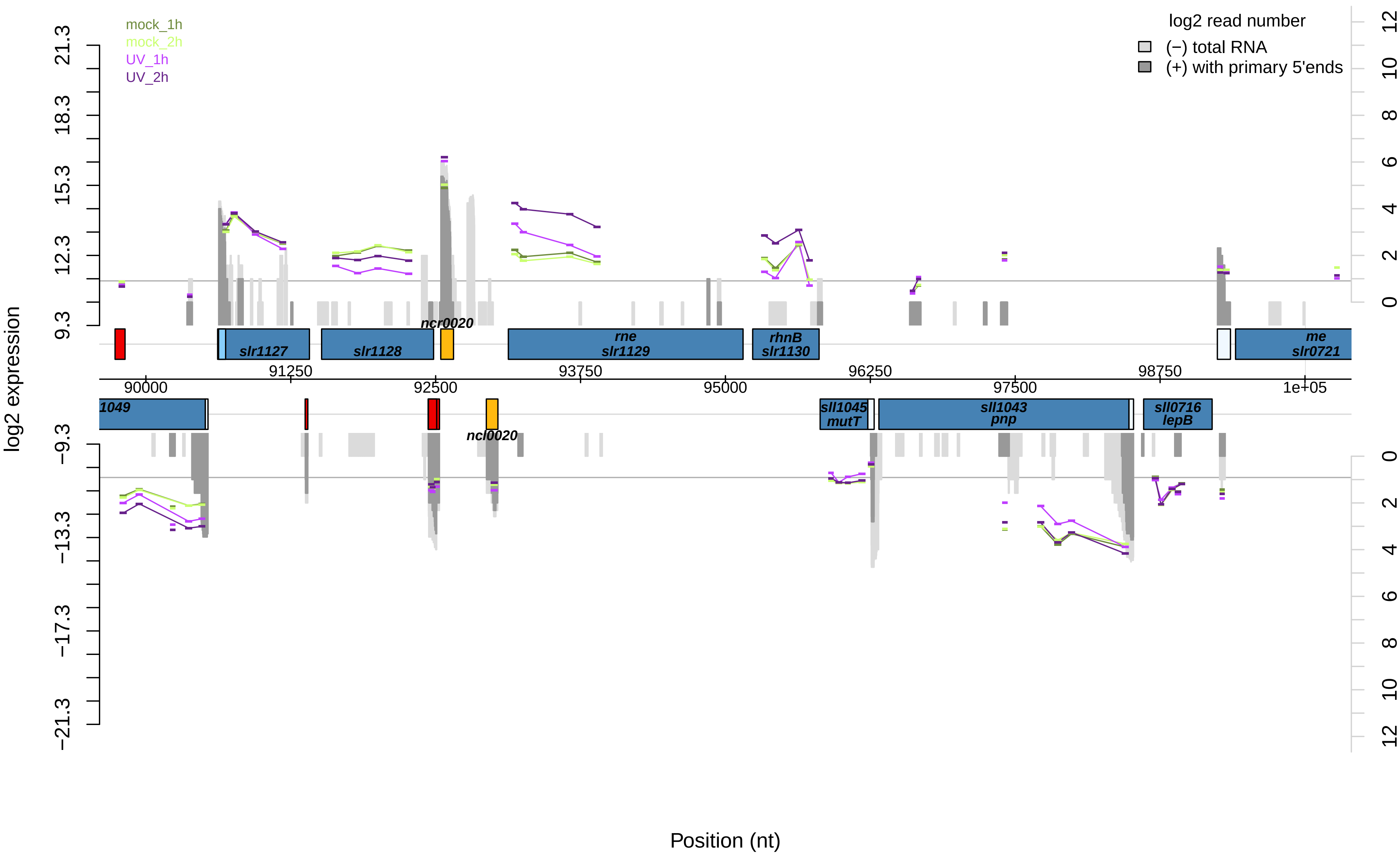

mock_1h
mock_2h
UV_1h
UV_2h
log2 read number
(−) total RNA
(+) with primary 5'ends
log2 expression
21.3
18.3
15.3
12.3
9.3
−9.3
−13.3
−17.3
−21.3
12
10
8
6
4
2
0
slr1127
slr1128
ncr0020
rne
slr1129
rhnB
slr1130
me
slr0721
1049
ncl0020
sll1045
mutT
sll1043
pnp
sll0716
lepB
90000
91250
92500
93750
95000
96250
97500
98750
1e+05
Position (nt)

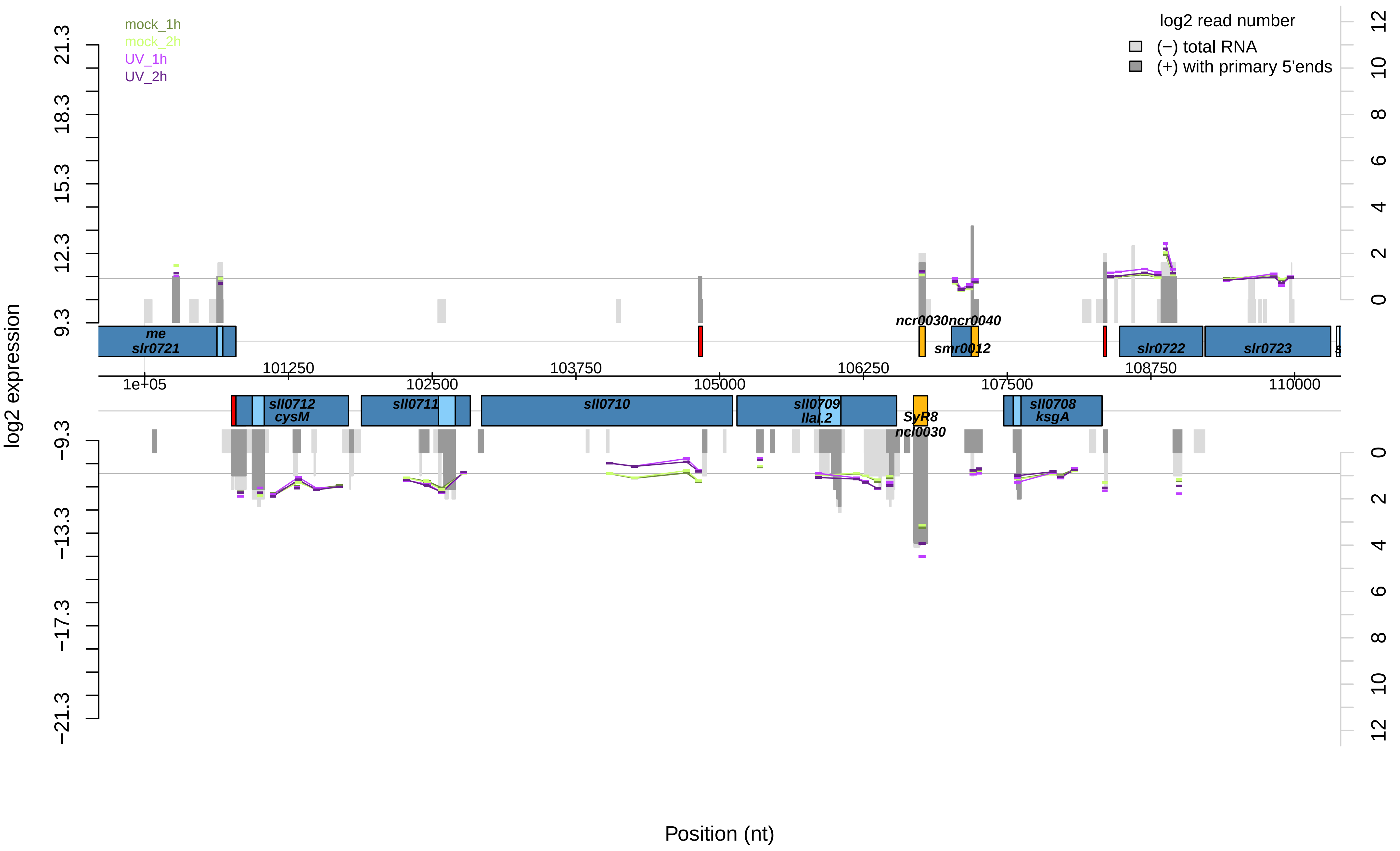

mock_1h
mock_2h
UV_1h
UV_2h
log2 read number
(−) total RNA
(+) with primary 5'ends
log2 expression
21.3
18.3
15.3
12.3
9.3
−9.3
−13.3
−17.3
−21.3
me
slr0721
ncr0030ncr0040
smr0012
slr0722
slr0723
101250
102500
103750
105000
106250
107500
108750
110000
1e+05
sll0712
cysM
sll0711
sll0710
sll0709
llaL.2
SyR8
ncl0030
sll0708
ksgA
Position (nt)

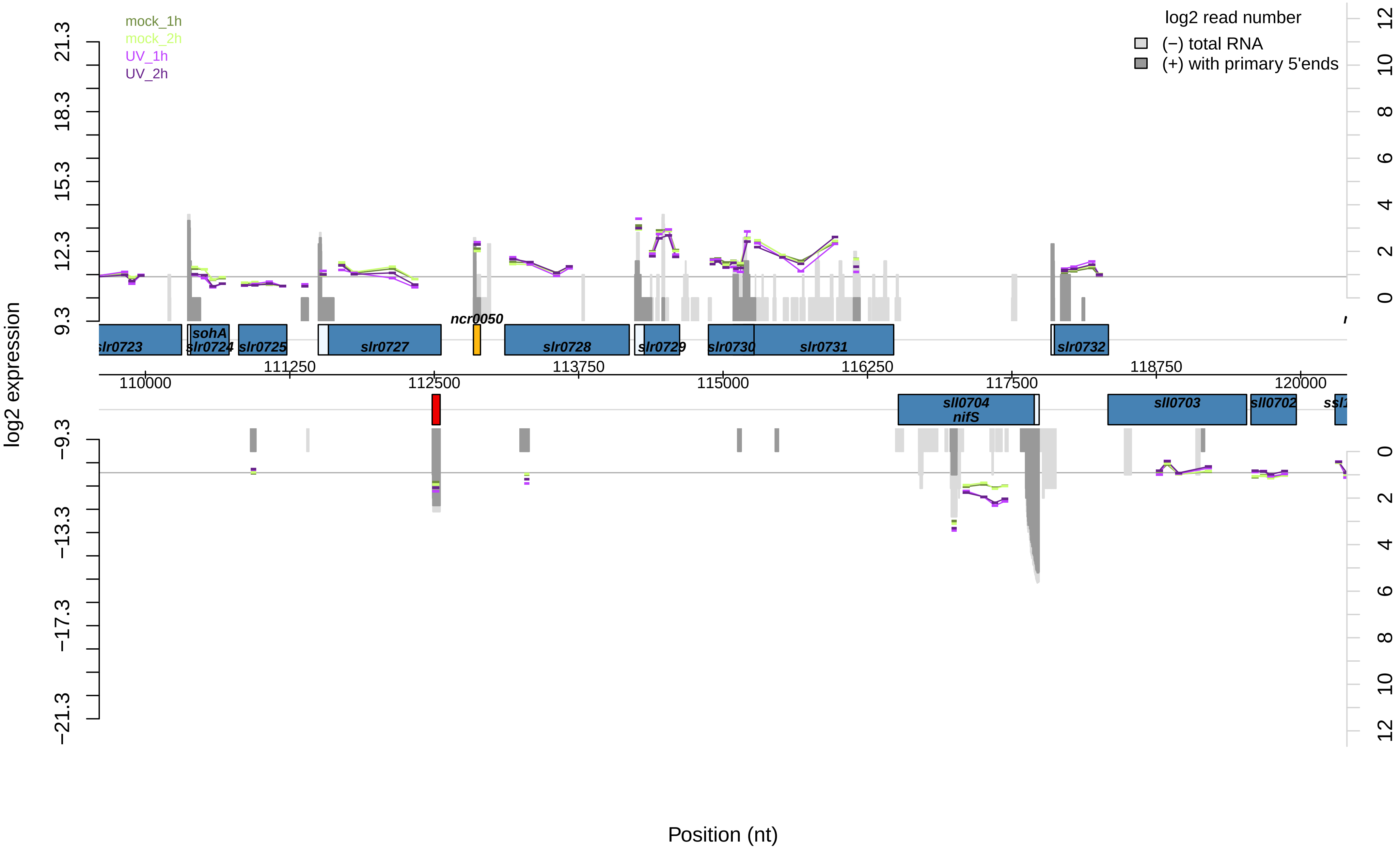
mock_1h
mock_2h
UV_1h
UV_2h
log2 read number
(−) total RNA
(+) with primary 5'ends
log2 expression
21.3
18.3
15.3
12.3
9.3
−9.3
−13.3
−17.3
−21.3
12
10
8
6
4
2
0
slr0723
sohA
slr0724
slr0725
slr0727
ncr0050
slr0728
slr0729
slr0730
slr0731
slr0732
111250
113750
116250
118750
110000
112500
115000
117500
120000
sll0704
nifS
sll0703
sll0702
Position (nt)

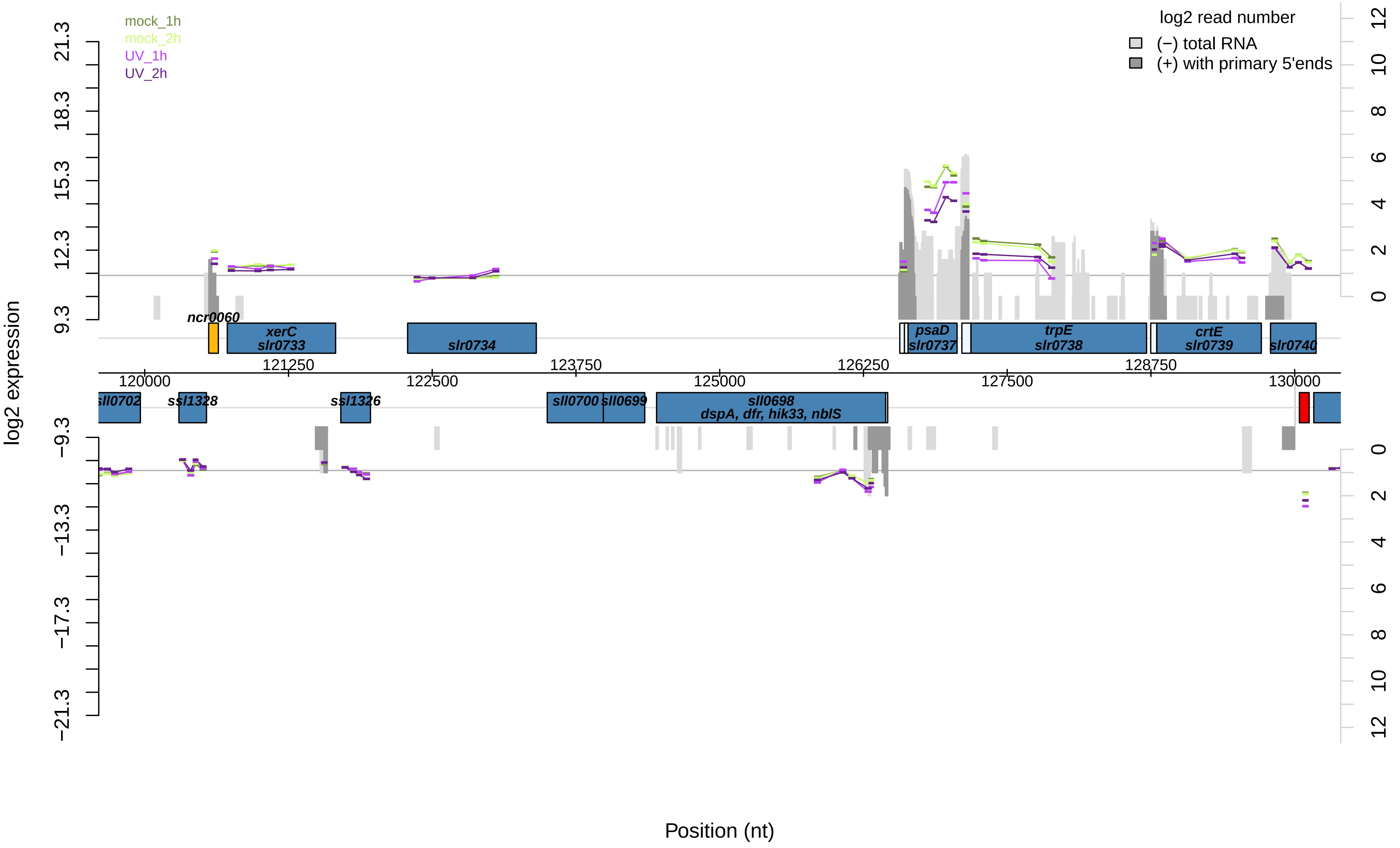

mock_1h
mock_2h
UV_1h
UV_2h
log2 read number
(−) total RNA
(+) with primary 5'ends
log2 expression
21.3
18.3
15.3
12.3
9.3
−9.3
−13.3
−17.3
−21.3
12
10
8
6
4
2
0
ncr0060
xerC
slr0733
slr0734
psaD
slr0737
trpE
slr0738
crtE
slr0739
slr0740
121250
123750
126250
128750
120000
122500
125000
127500
130000
sll0702
ssl1328
ssl1326
sll0700
sll0699
sll0698
dspA, dfr, hik33, nblS
Position (nt)

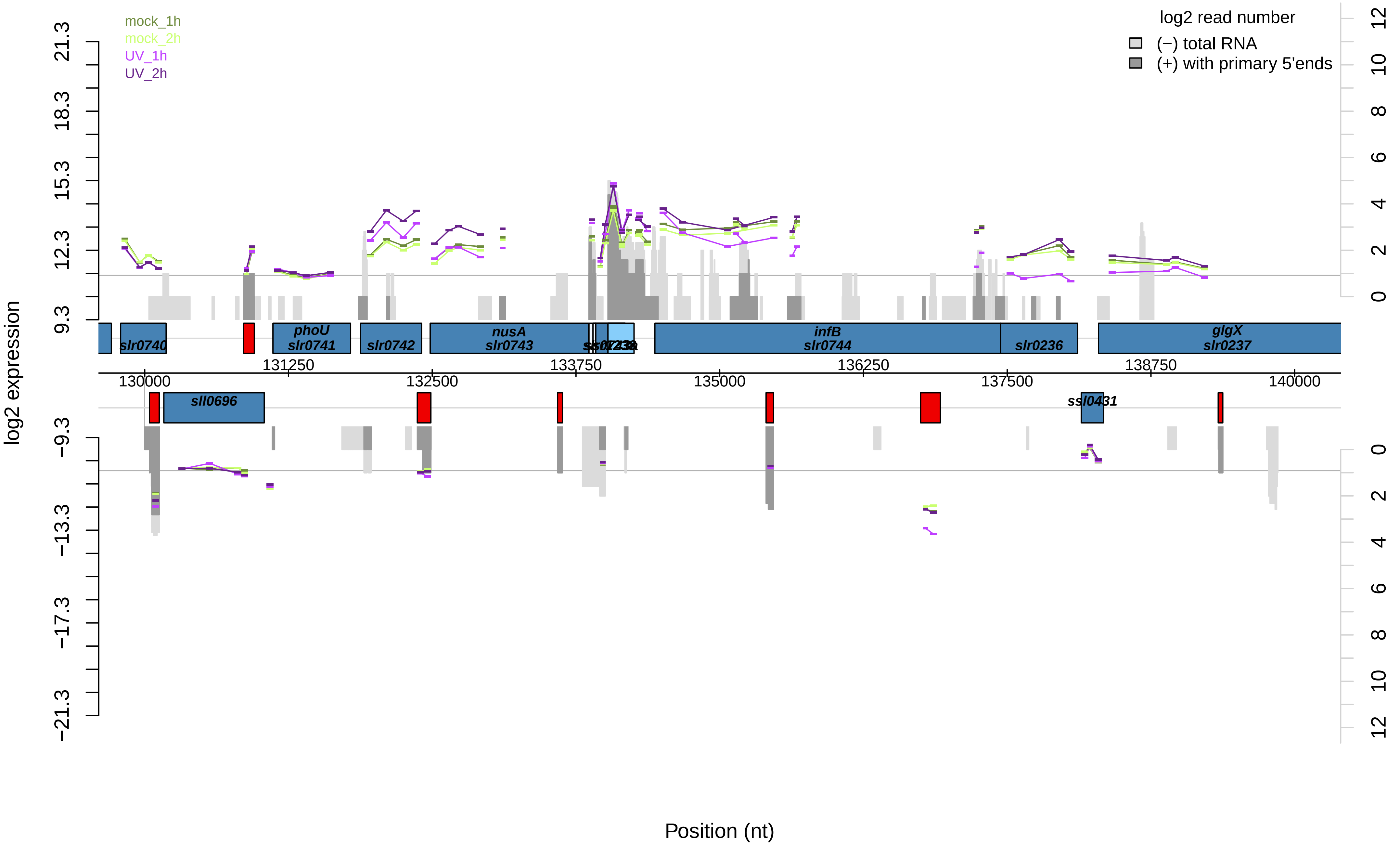

mock_1h
mock_2h
UV_1h
UV_2h
log2 read number
(−) total RNA
(+) with primary 5'ends
log2 expression
Position (nt)
phoU
slr0741
slr0740
slr0742
nusA
slr0743
infB
slr0744
slr0236
glgX
slr0237
sll0696
ssl0431
130000
131250
132500
133750
135000
136250
137500
138750
140000

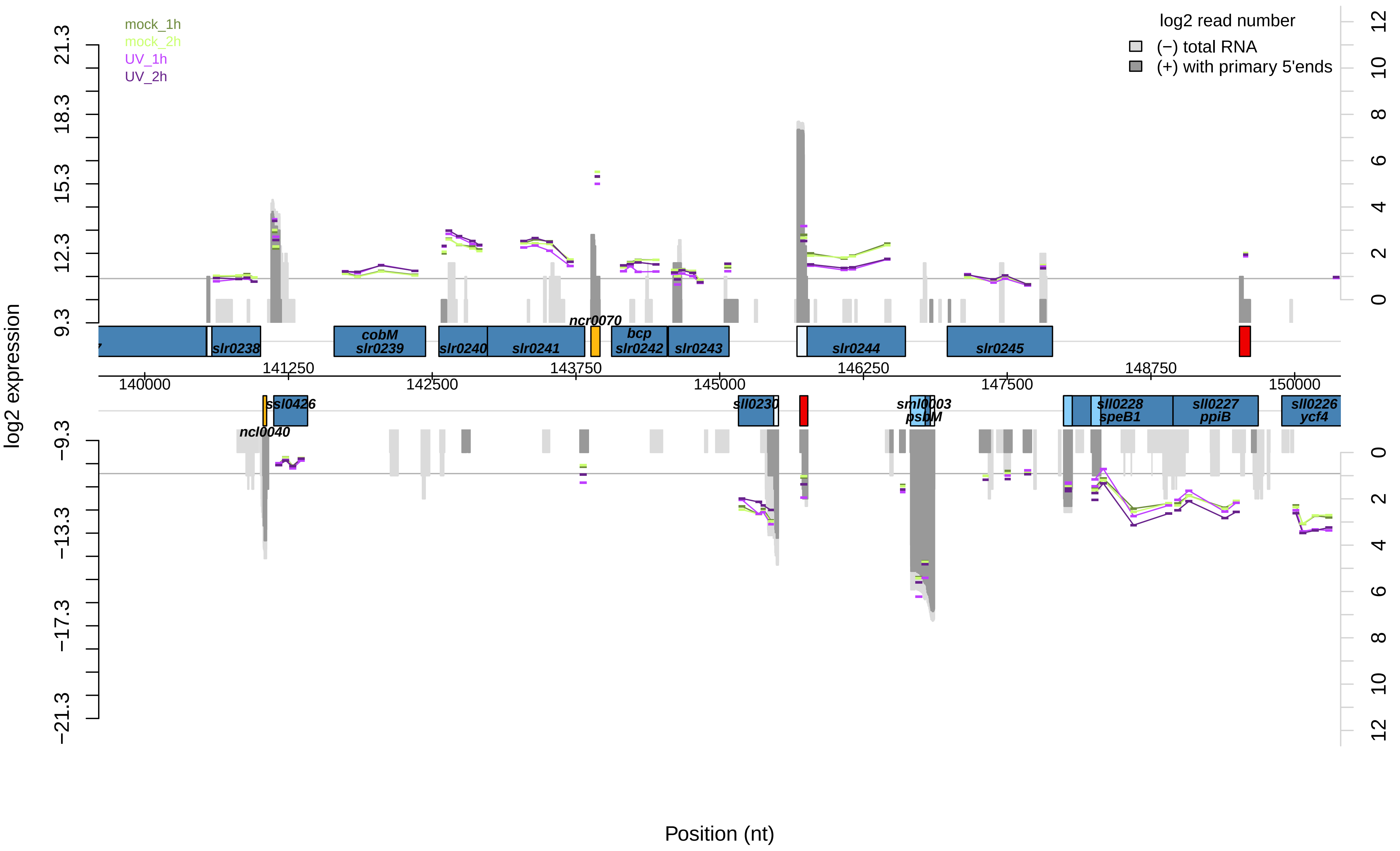
mock_1h
mock_2h
UV_1h
UV_2h
log2 read number
(−) total RNA
(+) with primary 5'ends
log2 expression
21.3
18.3
15.3
12.3
9.3
−9.3
−13.3
−17.3
−21.3
12
10
8
6
4
2
0
slr0238
cobM
slr0239
slr0240
slr0241
ncr0070
bcp
slr0242
slr0243
slr0244
slr0245
141250
143750
146250
148750
140000
142500
145000
147500
150000
ssl0426
ncl0040
sll0230
sml0003
psbM
sll0228
speB1
sll0227
ppiB
sll0226
ycf4
Position (nt)

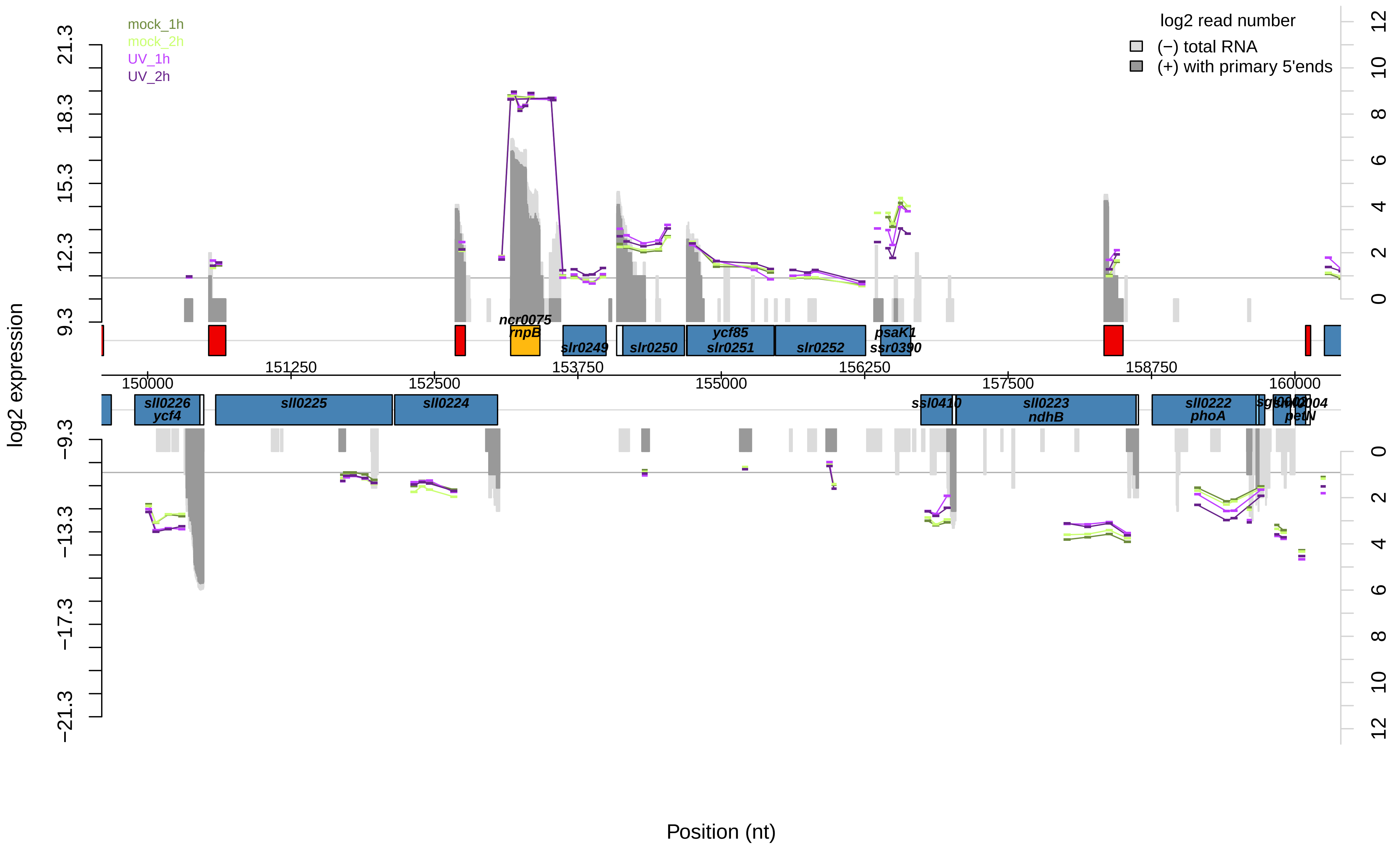
mock_1h
mock_2h
UV_1h
UV_2h
log2 read number
(−) total RNA
(+) with primary 5'ends
log2 expression
Position (nt)
21.3
18.3
15.3
12.3
9.3
−9.3
−13.3
−17.3
−21.3
150000
151250
152500
153750
155000
156250
157500
158750
160000
ncr0075
rnpB
slr0249
slr0250
ycf85
slr0251
slr0252
psaK1
ssr0390
sll0226
ycf4
sll0225
sll0224
ssl0410
sll0223
ndhB
sll0222
phoA
petN

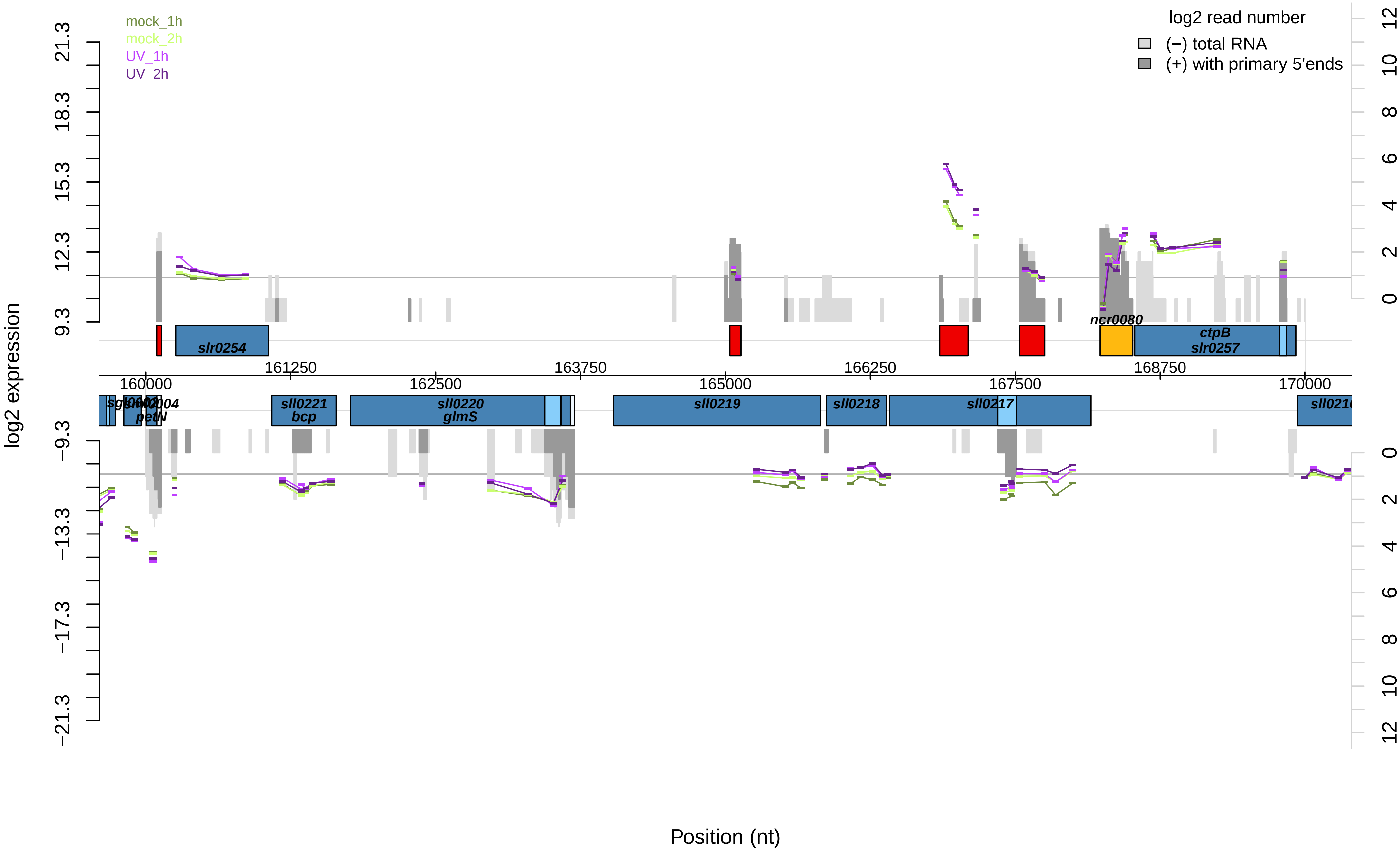

mock_1h
mock_2h
UV_1h
UV_2h
log2 read number
(−) total RNA
(+) with primary 5'ends
log2 expression
21.3
18.3
15.3
12.3
9.3
−9.3
−13.3
−17.3
−21.3
12
10
8
6
4
2
0
ncr0080
slr0254
ctpB
slr0257
160000
161250
162500
163750
165000
166250
167500
168750
170000
petN
sll0221
bcp
sll0220
glmS
sll0219
sll0218
sll0217
Position (nt)

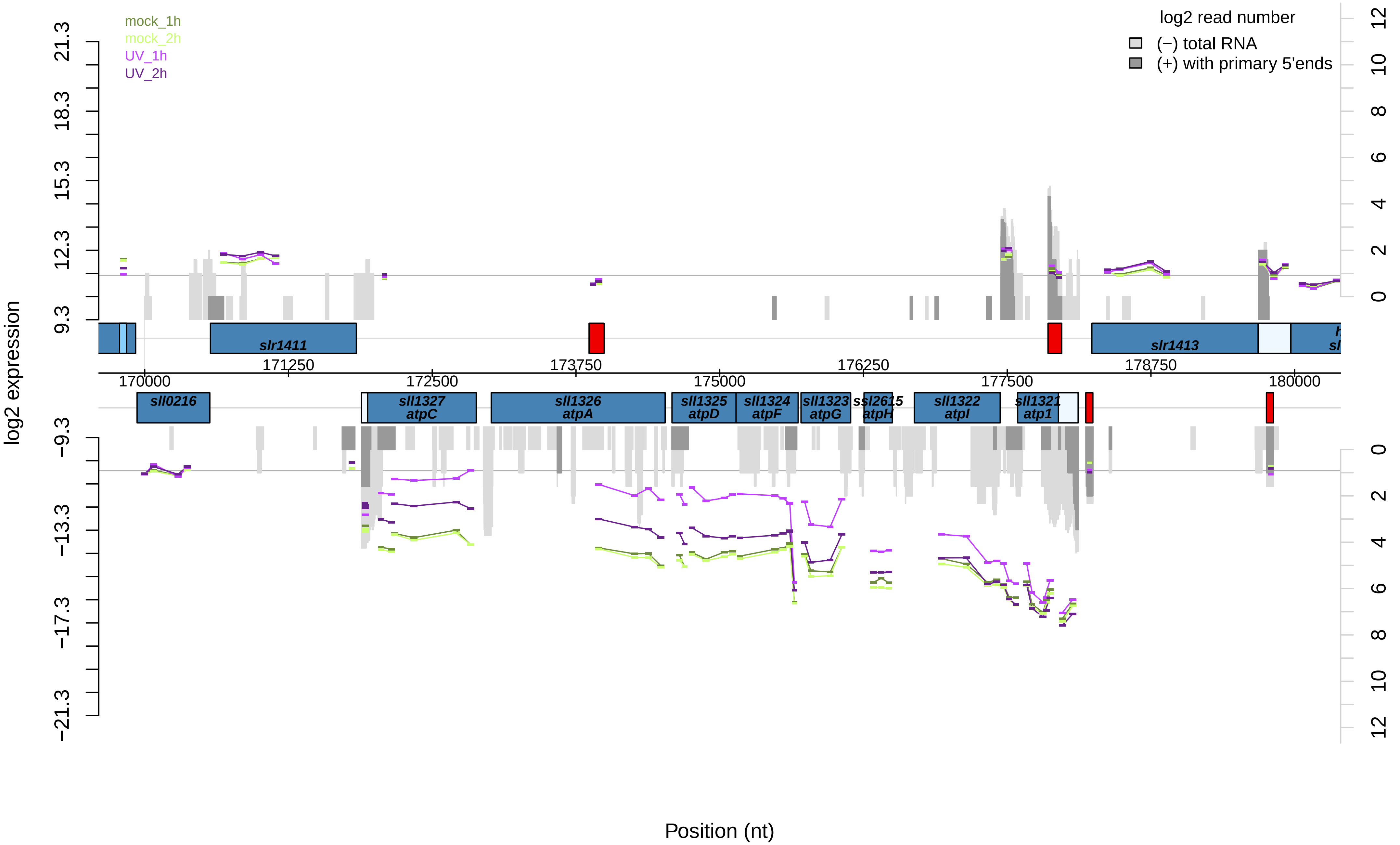

mock_1h
mock_2h
UV_1h
UV_2h
log2 read number
(−) total RNA
(+) with primary 5'ends
log2 expression
21.3
18.3
15.3
12.3
9.3
−9.3
−13.3
−17.3
−21.3
12
10
8
6
4
2
0
slr1411
slr1413
171250
173750
176250
178750
170000
172500
175000
177500
180000
sll0216
sll1327
atpC
sll1326
atpA
sll1325
atpD
sll1324
atpF
sll1323
atpG
ssl2615
atpH
sll1322
atpI
sll1321
atp1
Position (nt)

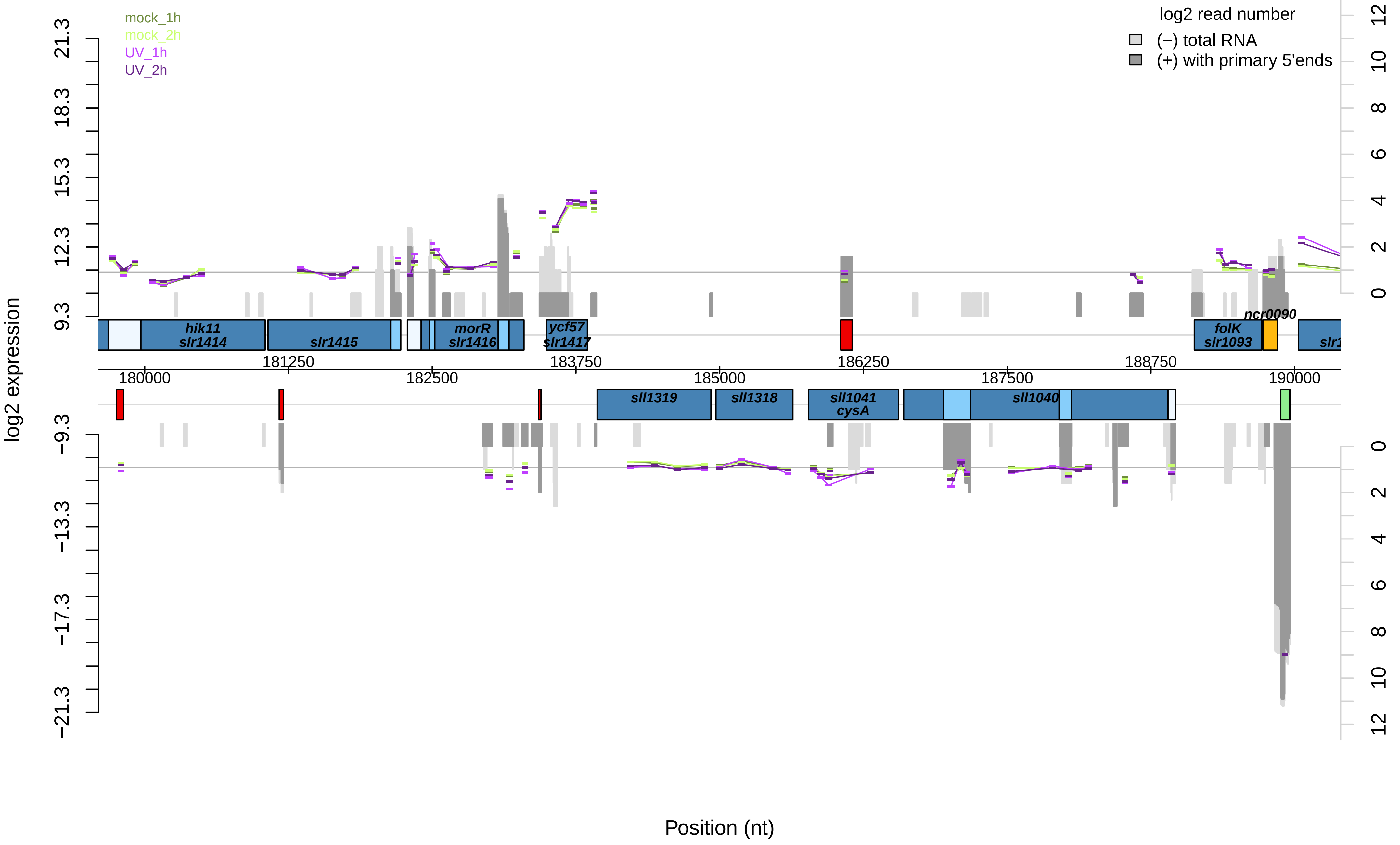

mock_1h
mock_2h
UV_1h
UV_2h
log2 read number
(−) total RNA
(+) with primary 5'ends
log2 expression
21.3
18.3
15.3
12.3
9.3
−9.3
−13.3
−17.3
−21.3
12
10
8
6
4
2
0
hik11
slr1414
slr1415
morR
slr1416
ycf57
slr1417
folK
slr1093
ncr0090
181250
183750
186250
188750
180000
182500
185000
187500
190000
sll1319
sll1318
sll1041
cysA
sll1040
Position (nt)

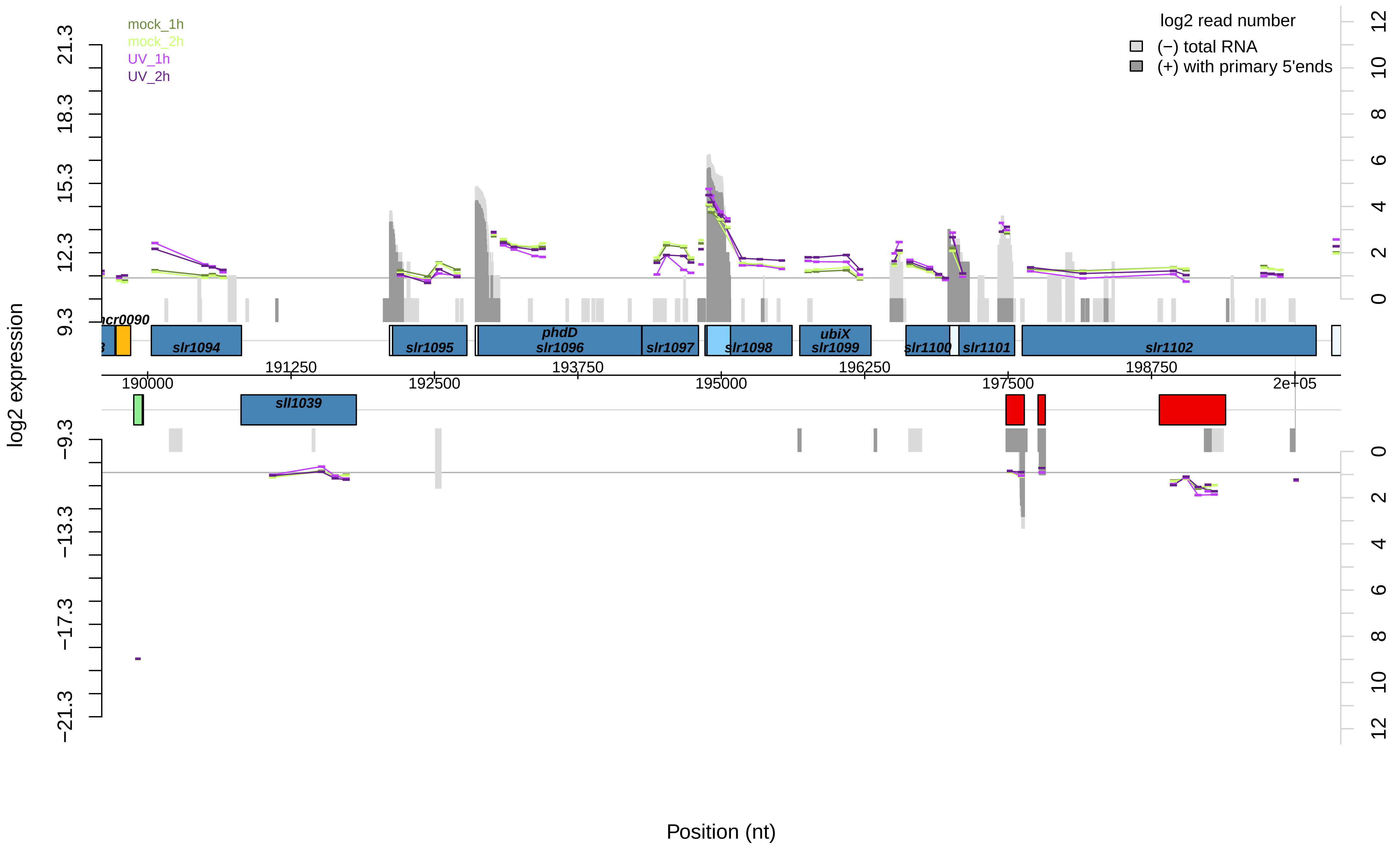
mock_1h
mock_2h
UV_1h
UV_2h
log2 read number
(−) total RNA
(+) with primary 5'ends
log2 expression
21.3
18.3
15.3
12.3
9.3
−9.3
−13.3
−17.3
−21.3
12
10
8
6
4
2
0
ncr0090
slr1094
slr1095
phdD
slr1096
slr1097
slr1098
ubiX
slr1099
slr1100
slr1101
slr1102
191250
193750
196250
198750
190000
192500
195000
197500
2e+05
sll1039
Position (nt)

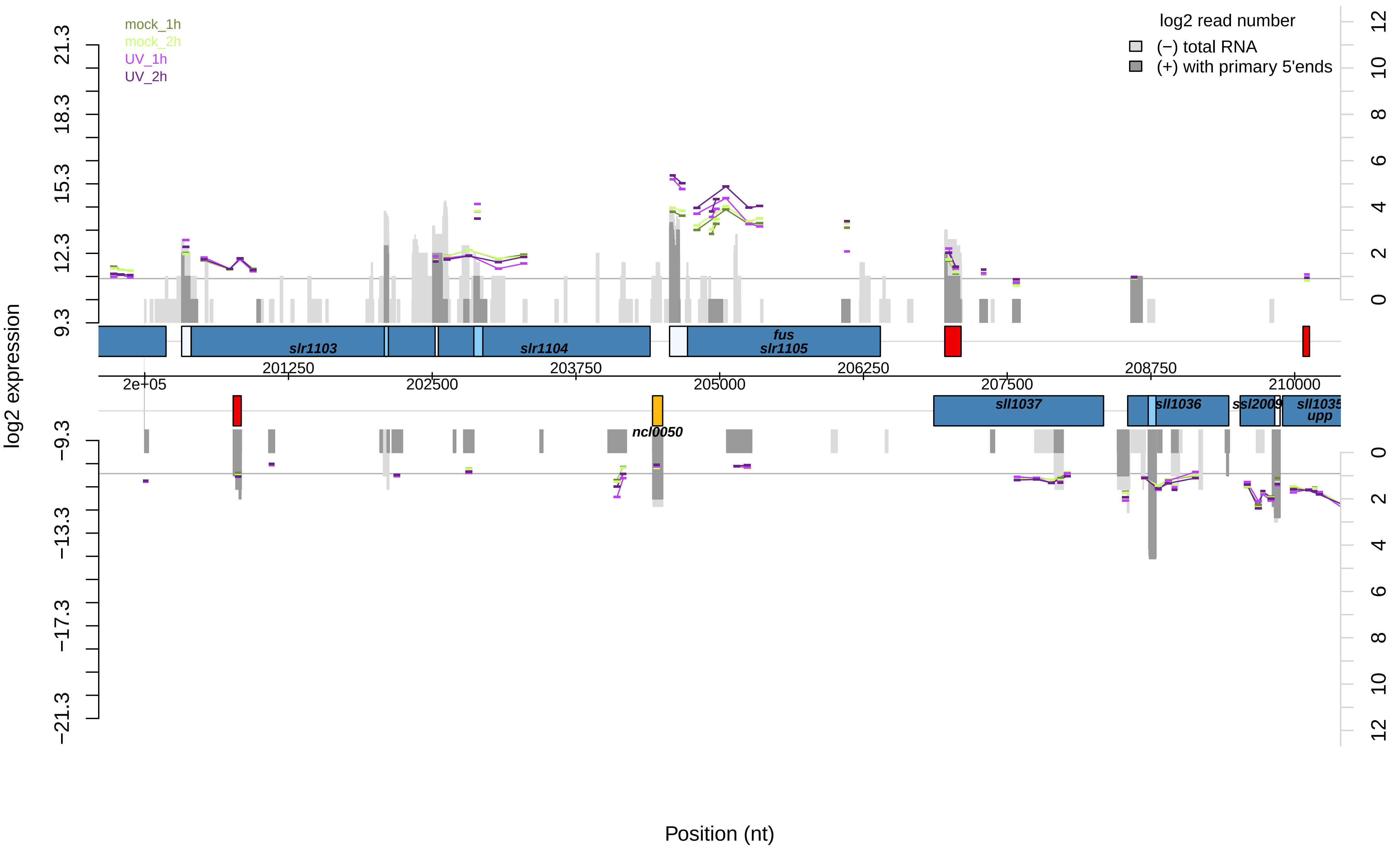
mock_1h
mock_2h
UV_1h
UV_2h
log2 read number
(−) total RNA
(+) with primary 5'ends
log2 expression
21.3
18.3
15.3
12.3
9.3
−9.3
−13.3
−17.3
−21.3
12
10
8
6
4
2
0
slr1103
slr1104
fus
slr1105
201250
203750
206250
208750
2e+05
202500
205000
207500
210000
sll1037
sll1036
ssl2009
sll1035
upp
ncl0050
Position (nt)

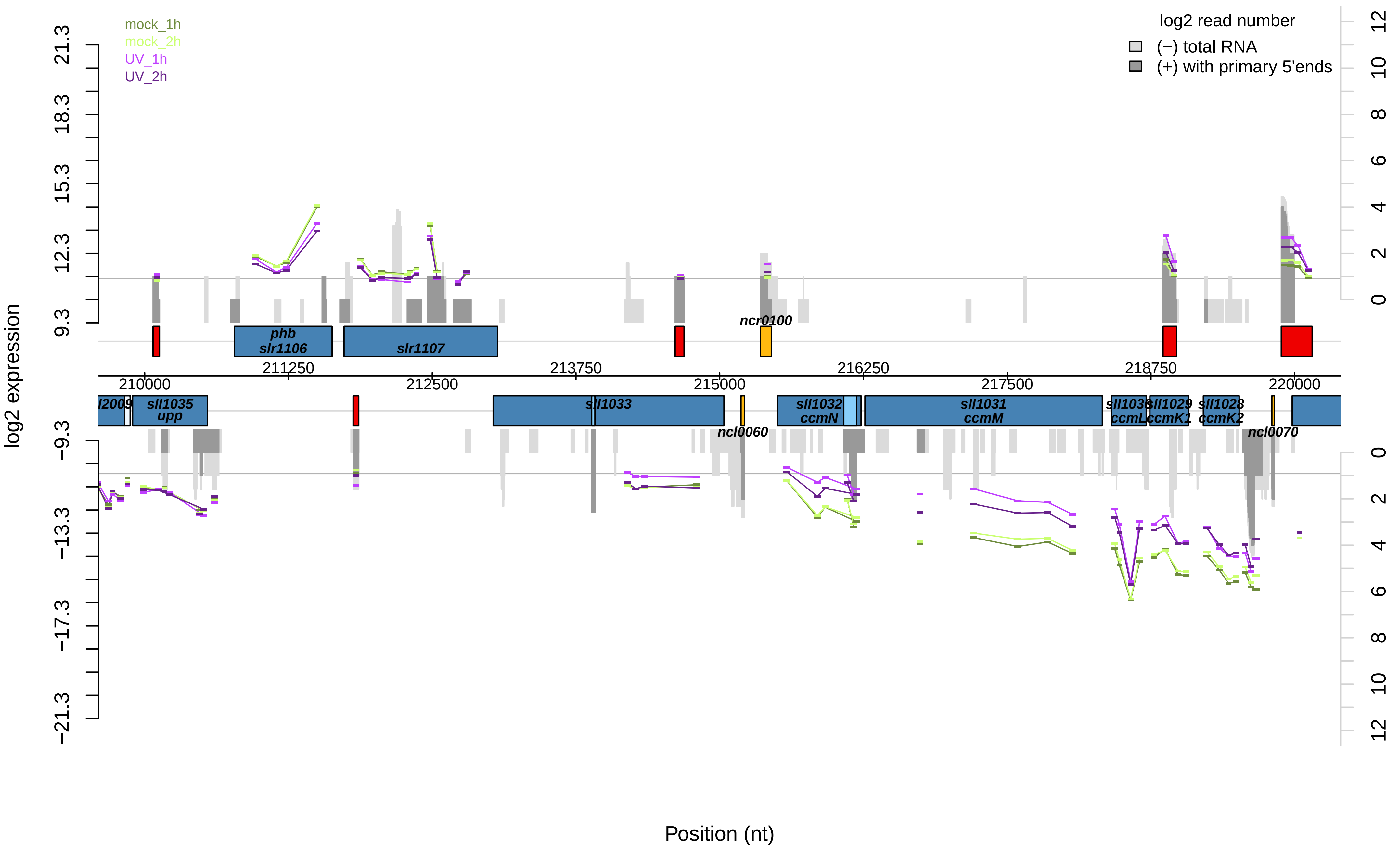

mock_1h
mock_2h
UV_1h
UV_2h
log2 read number
(−) total RNA
(+) with primary 5'ends
log2 expression
Position (nt)
21.3
18.3
15.3
12.3
9.3
−9.3
−13.3
−17.3
−21.3
12
10
8
6
4
2
0
phb
slr1106
slr1107
ncr0100
211250
213750
216250
218750
210000
212500
215000
217500
220000
sll1035
upp
sll1033
ncl0060
sll1032
ccmN
sll1031
ccmM
sll1030
ccmL
sll1029
ccmK1
sll1028
ccmK2
ncl0070

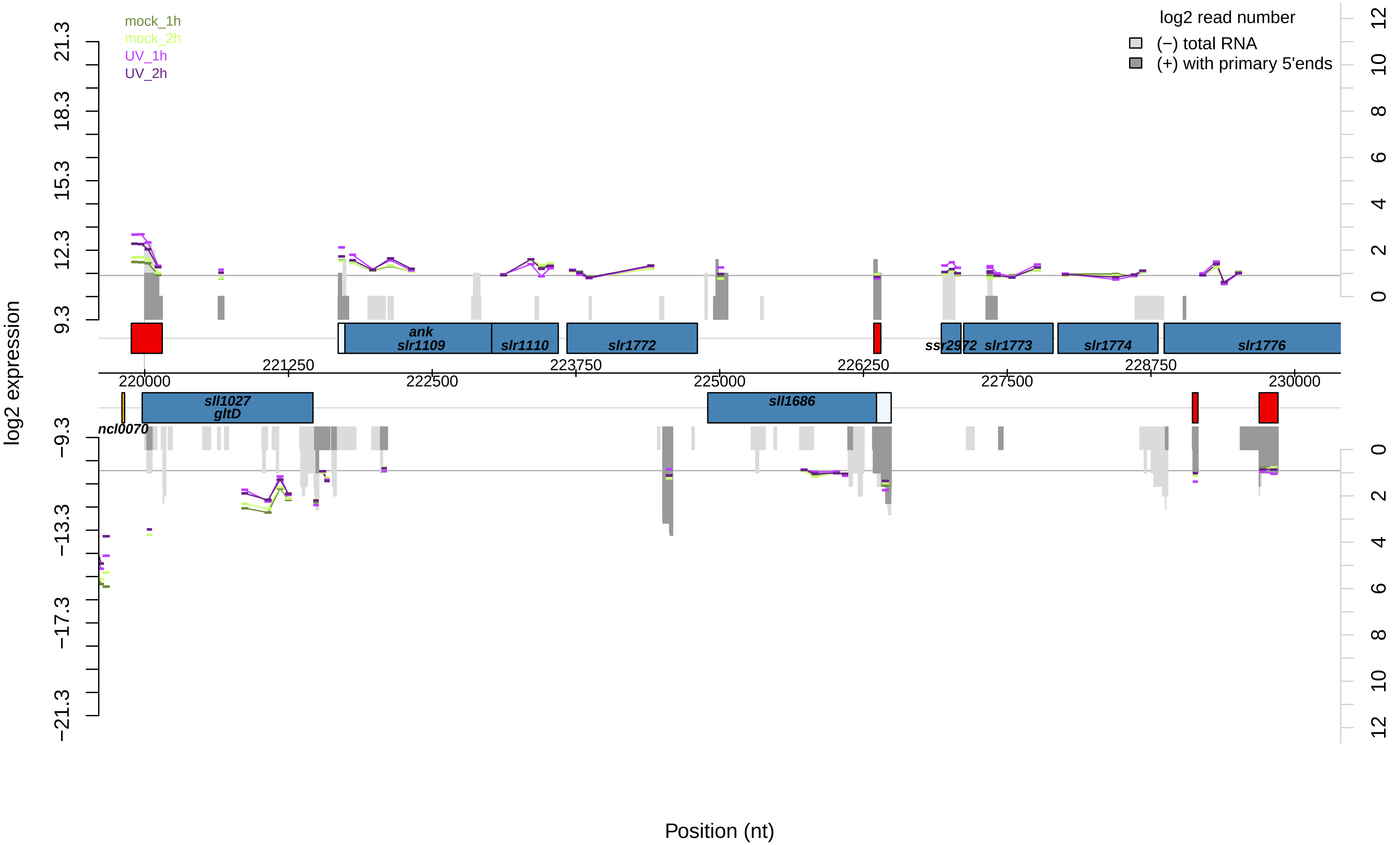

mock_1h
mock_2h
UV_1h
UV_2h
log2 read number
(−) total RNA
(+) with primary 5'ends
log2 expression
21.3
18.3
15.3
12.3
9.3
−9.3
−13.3
−17.3
−21.3
12
10
8
6
4
2
0
ank
slr1109
slr1110
slr1772
ssr2972
slr1773
slr1774
slr1776
221250
223750
226250
228750
220000
222500
225000
227500
230000
sll1027
gltD
sll1686
ncl0070
Position (nt)

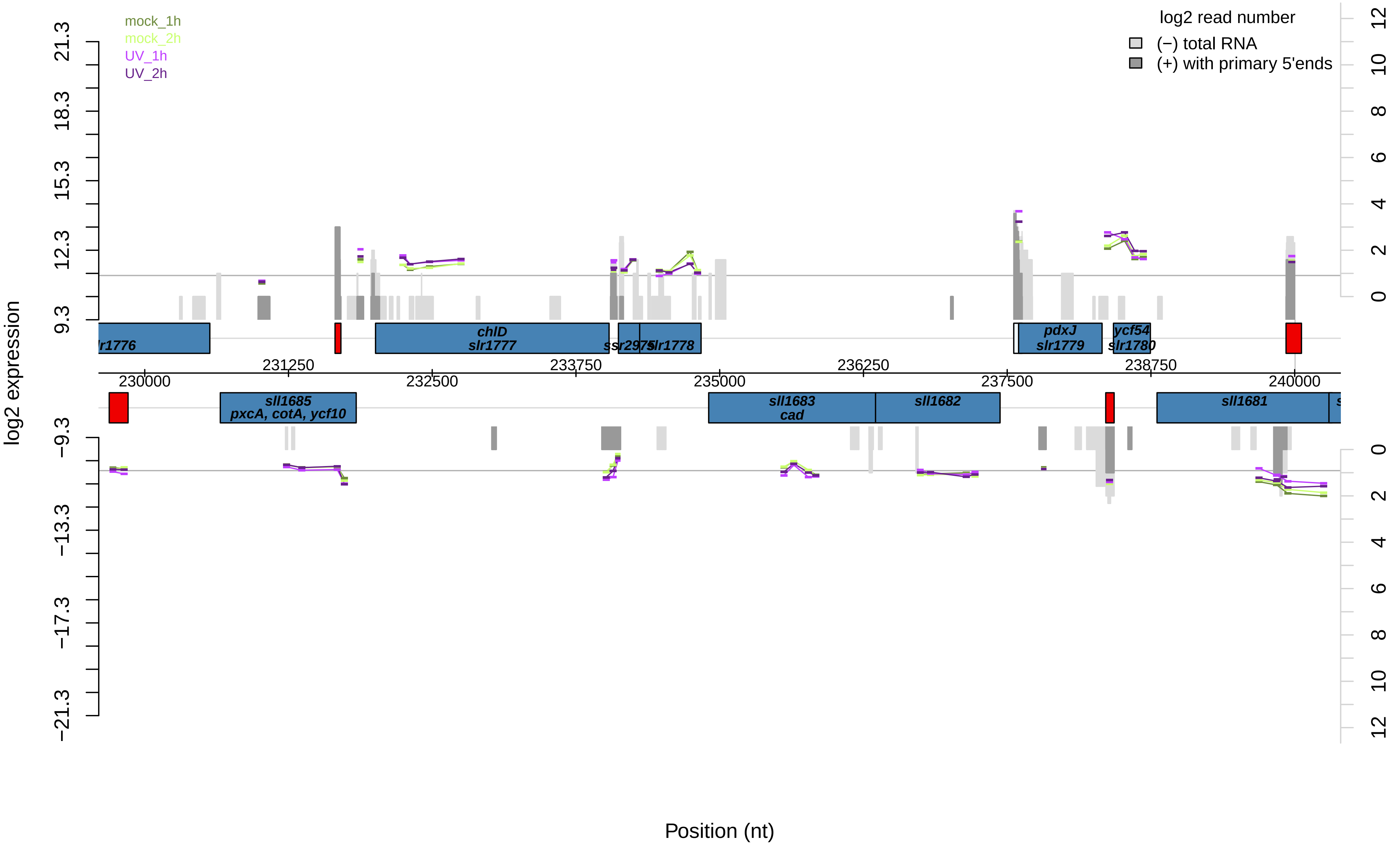

mock_1h
mock_2h
UV_1h
UV_2h
log2 read number
(−) total RNA
(+) with primary 5'ends
log2 expression
21.3
18.3
15.3
12.3
9.3
−9.3
−13.3
−17.3
−21.3
12
10
8
6
4
2
0
lr1776
chlD
slr1777
ssr2975
slr1778
pdxJ
slr1779
ycf54
slr1780
231250
233750
236250
238750
230000
232500
235000
237500
240000
sll1685
pxcA, cotA, ycf10
sll1683
cad
sll1682
sll1681
Position (nt)

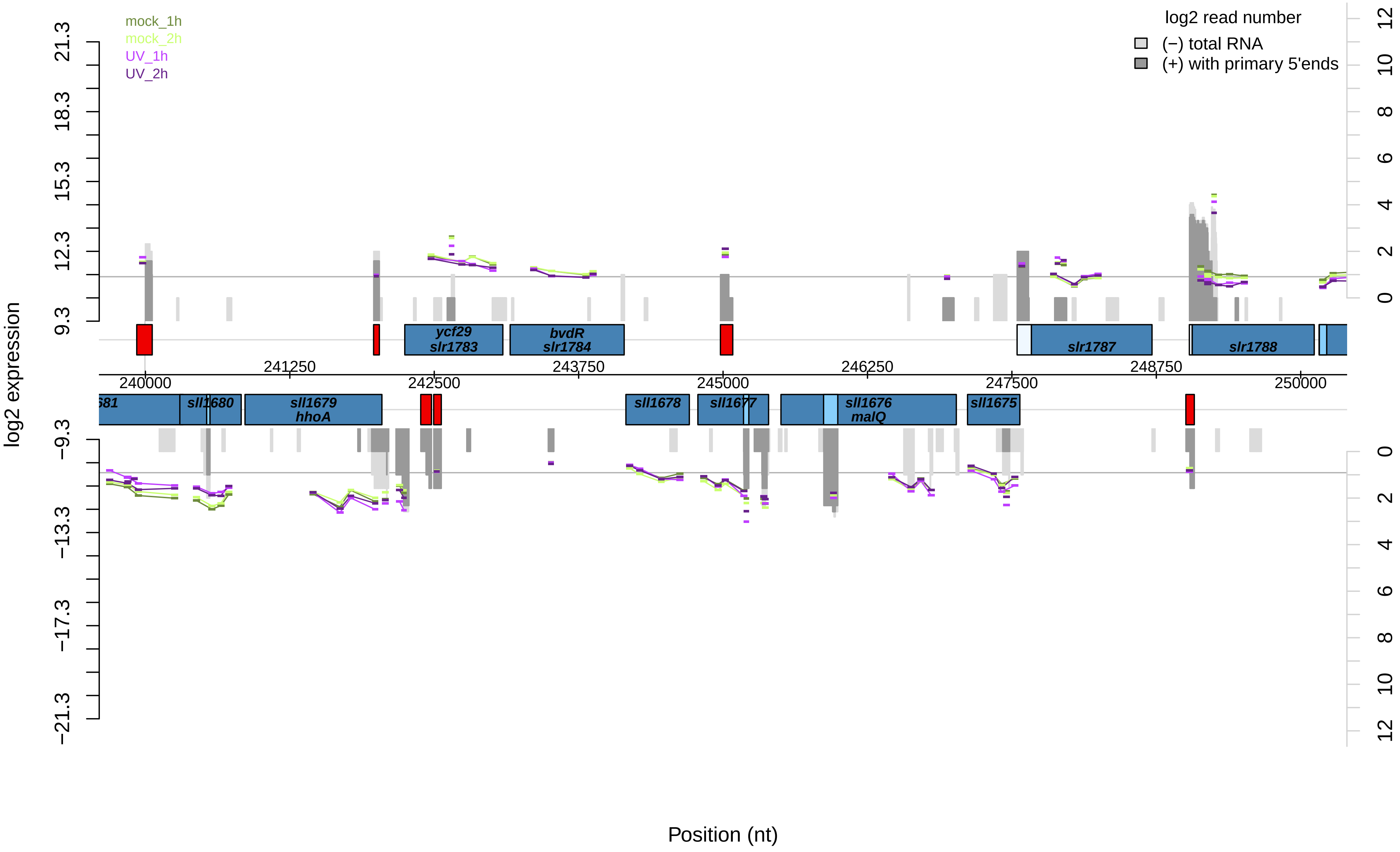

mock_1h
mock_2h
UV_1h
UV_2h
log2 read number
(−) total RNA
(+) with primary 5'ends
log2 expression
21.3
18.3
15.3
12.3
9.3
−9.3
−13.3
−17.3
−21.3
12
10
8
6
4
2
0
ycf29
slr1783
bvdR
slr1784
slr1787
slr1788
241250
242500
243750
245000
246250
247500
248750
240000
250000
sll1680
sll1679
hhoA
sll1678
sll1677
sll1676
malQ
sll1675
Position (nt)

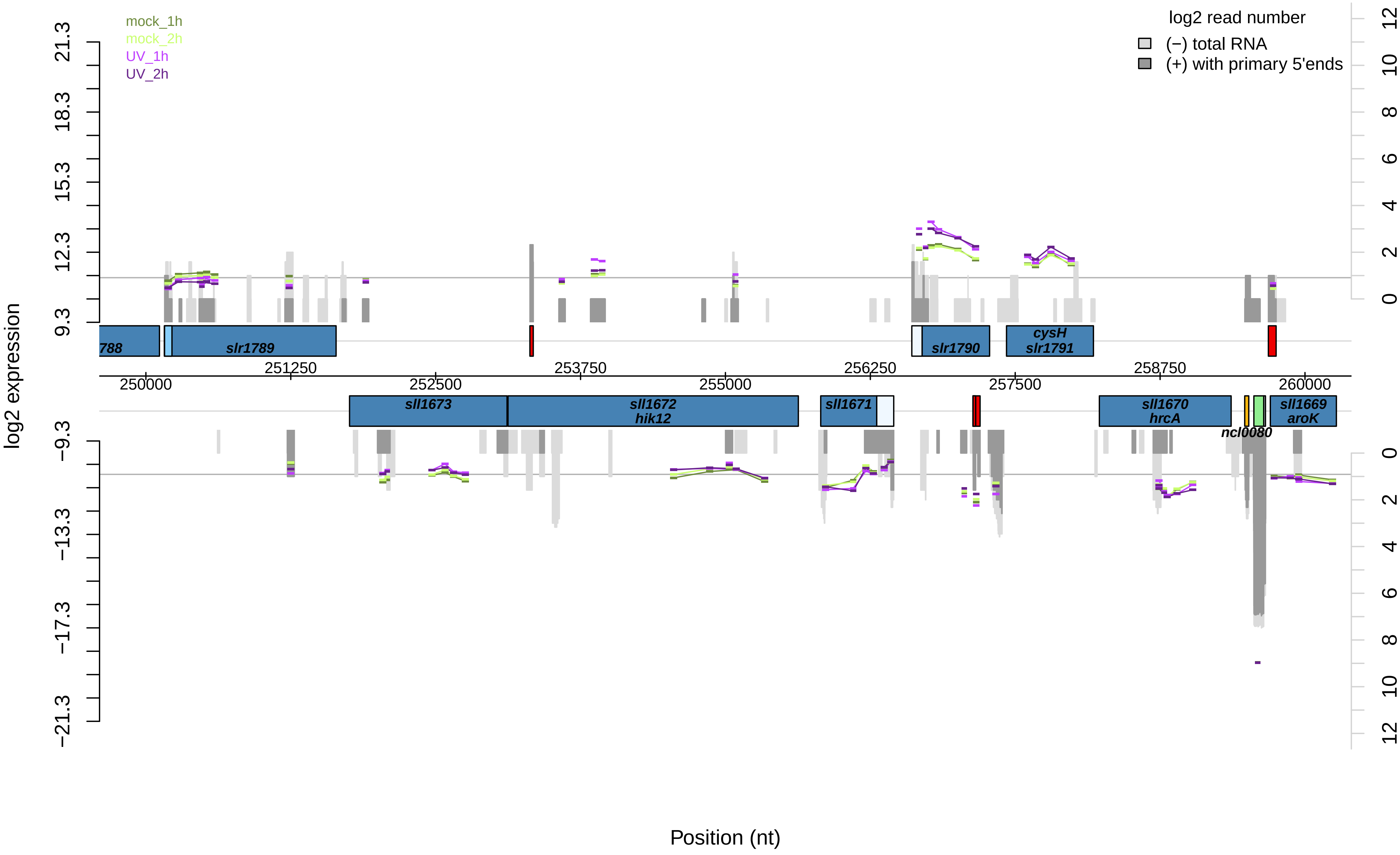

mock_1h
mock_2h
UV_1h
UV_2h
log2 read number
(−) total RNA
(+) with primary 5'ends
log2 expression
21.3
18.3
15.3
12.3
9.3
−9.3
−13.3
−17.3
−21.3
788
slr1789
slr1790
cysH
slr1791
sll1673
sll1672
hik12
sll1671
sll1670
hrcA
ncl0080
sll1669
aroK
250000
251250
252500
253750
255000
256250
257500
258750
260000
Position (nt)

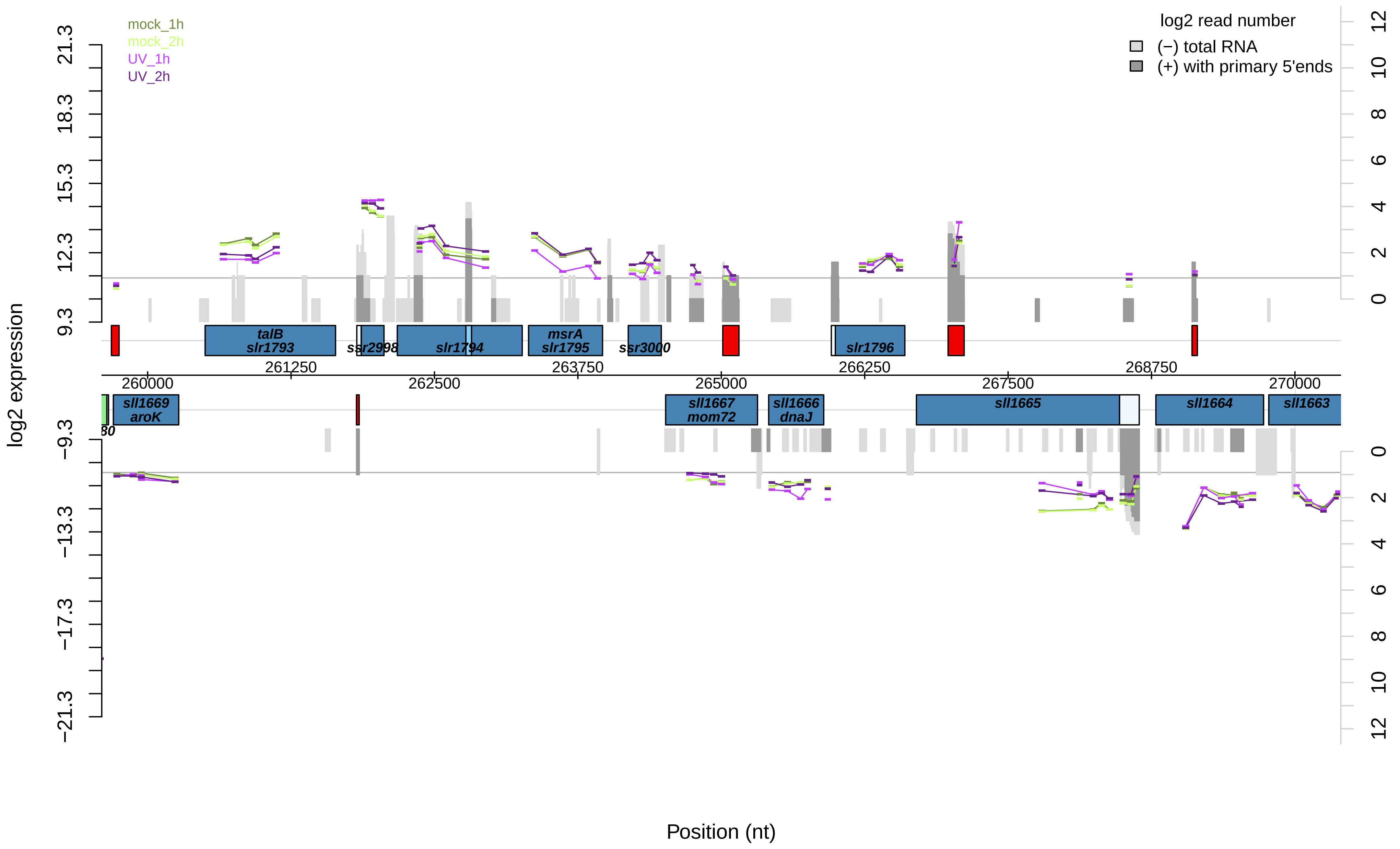

mock_1h
mock_2h
UV_1h
UV_2h
log2 read number
(−) total RNA
(+) with primary 5'ends
log2 expression
21.3
18.3
15.3
12.3
9.3
−9.3
−13.3
−17.3
−21.3
12
10
8
6
4
2
0
talB
slr1793
ssr2998
slr1794
msrA
slr1795
ssr3000
slr1796
261250
263750
266250
268750
260000
262500
265000
267500
270000
sll1669
aroK
sll1667
mom72
sll1666
dnaJ
sll1665
sll1664
sll1663
Position (nt)

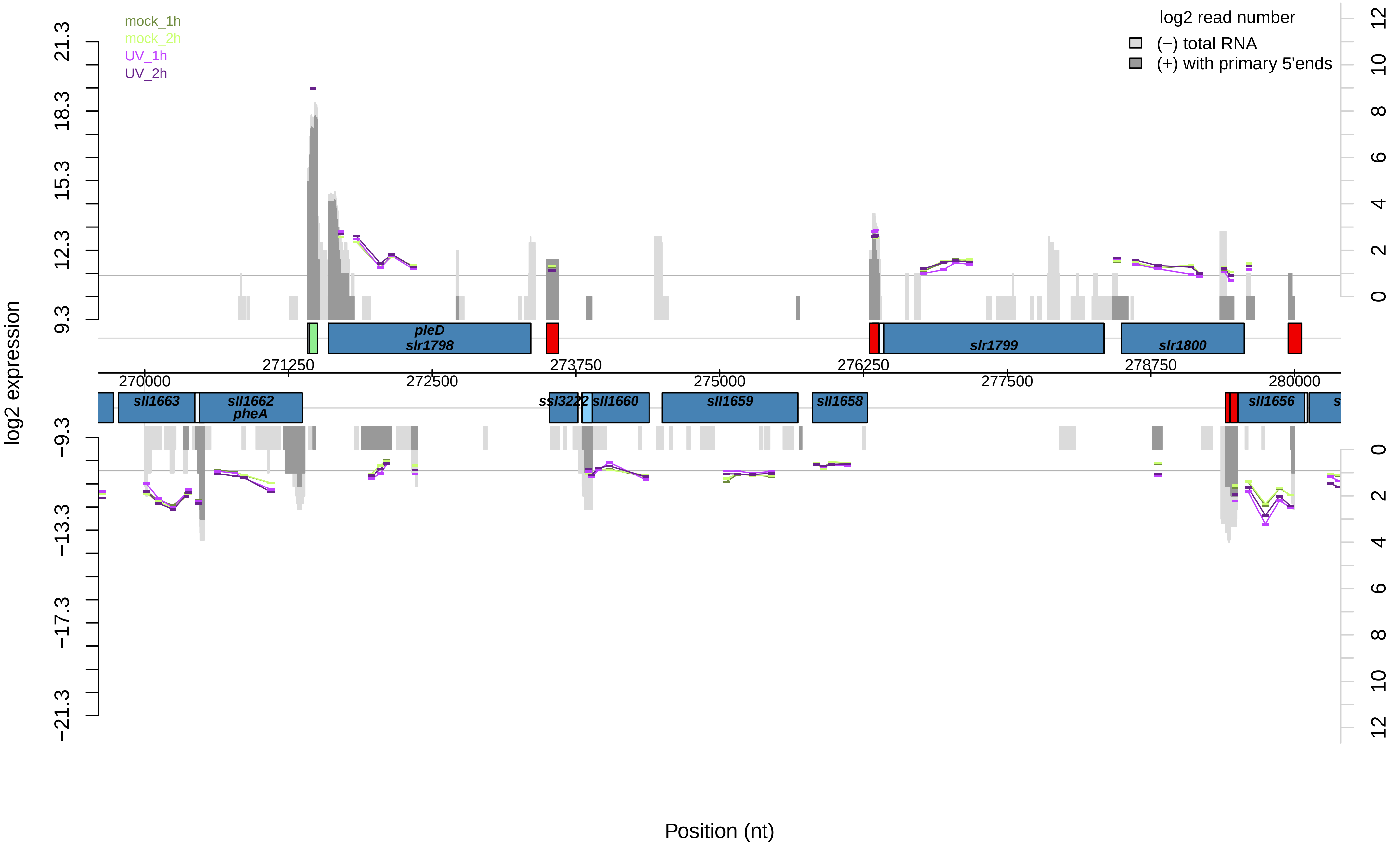
mock_1h
mock_2h
UV_1h
UV_2h
log2 read number
(−) total RNA
(+) with primary 5'ends
log2 expression
21.3
18.3
15.3
12.3
9.3
−9.3
−13.3
−17.3
−21.3
12
10
8
6
4
2
0
pleD
slr1798
slr1799
slr1800
271250
273750
276250
278750
270000
272500
275000
277500
280000
sll1663
sll1662
pheA
ssl3222
sll1660
sll1659
sll1658
sll1656
Position (nt)

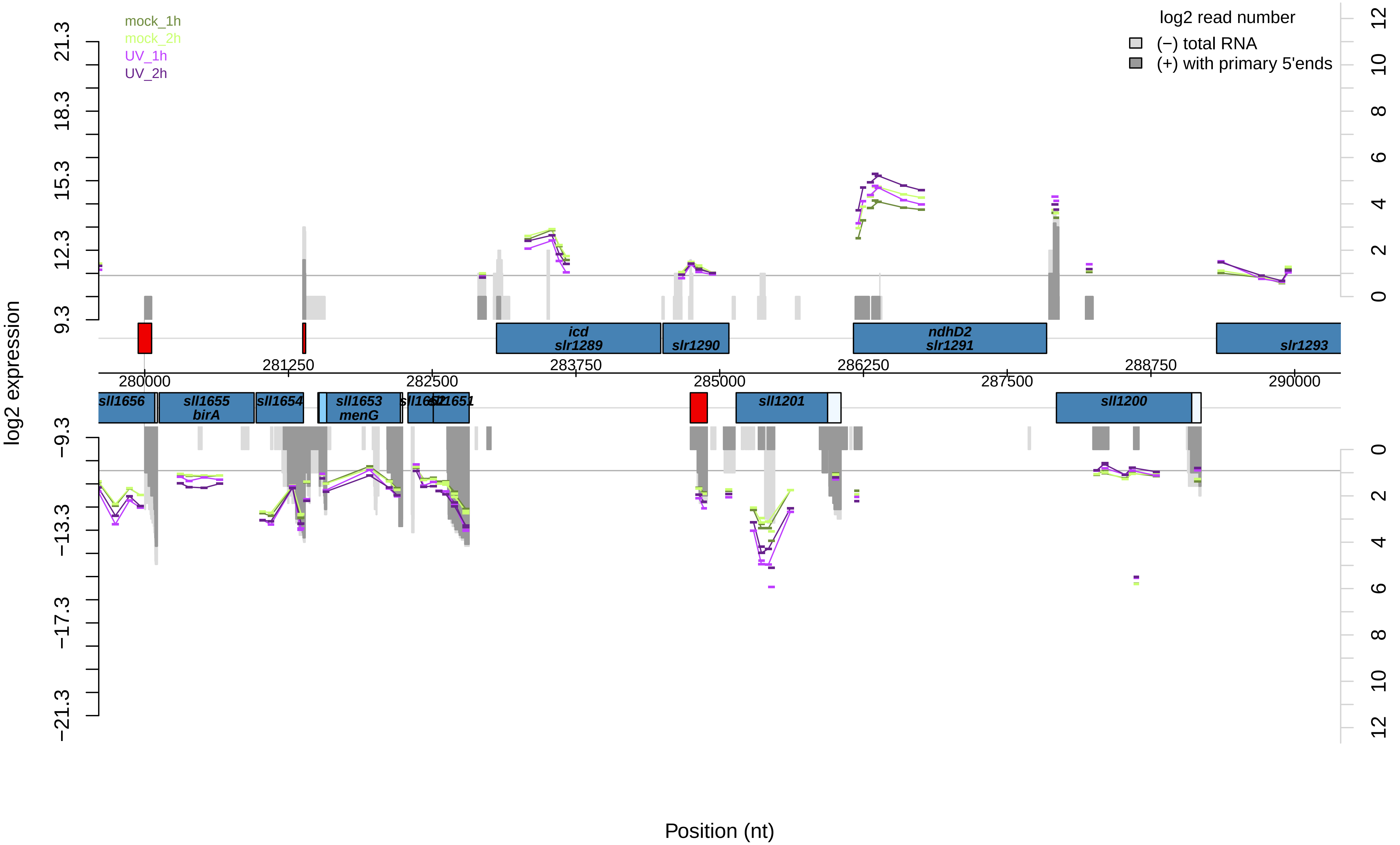

mock_1h
mock_2h
UV_1h
UV_2h
log2 read number
(−) total RNA
(+) with primary 5'ends
log2 expression
Position (nt)
icd
slr1289
slr1290
ndhD2
slr1291
slr1293
sll1656
sll1655
birA
sll1654
sll1653
menG
sll1652
sll1651
sll1201
sll1200
280000
281250
282500
283750
285000
286250
287500
288750
290000
21.3
18.3
15.3
12.3
9.3
−9.3
−13.3
−17.3
−21.3

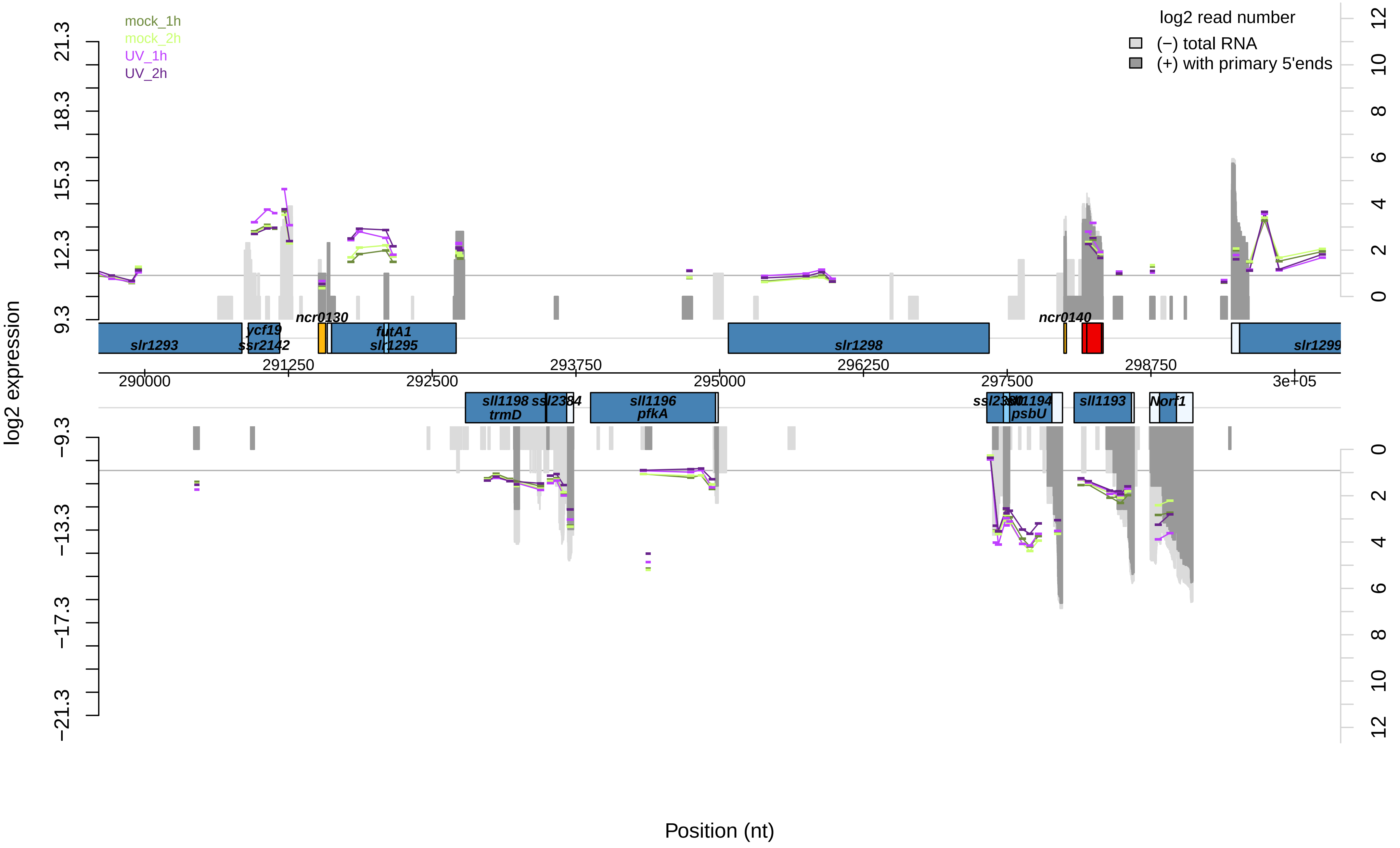

mock_1h
mock_2h
UV_1h
UV_2h
log2 read number
(−) total RNA
(+) with primary 5'ends
log2 expression
21.3
18.3
15.3
12.3
9.3
−9.3
−13.3
−17.3
−21.3
12
10
8
6
4
2
0
slr1293
ycf19
ssr2142
ncr0130
futA1
slr1295
slr1298
ncr0140
slr1299
291250
293750
296250
298750
290000
292500
295000
297500
3e+05
sll1198
trmD
ssl2384
sll1196
pfkA
sll1194
psbU
sll1193
Norf1
Position (nt)

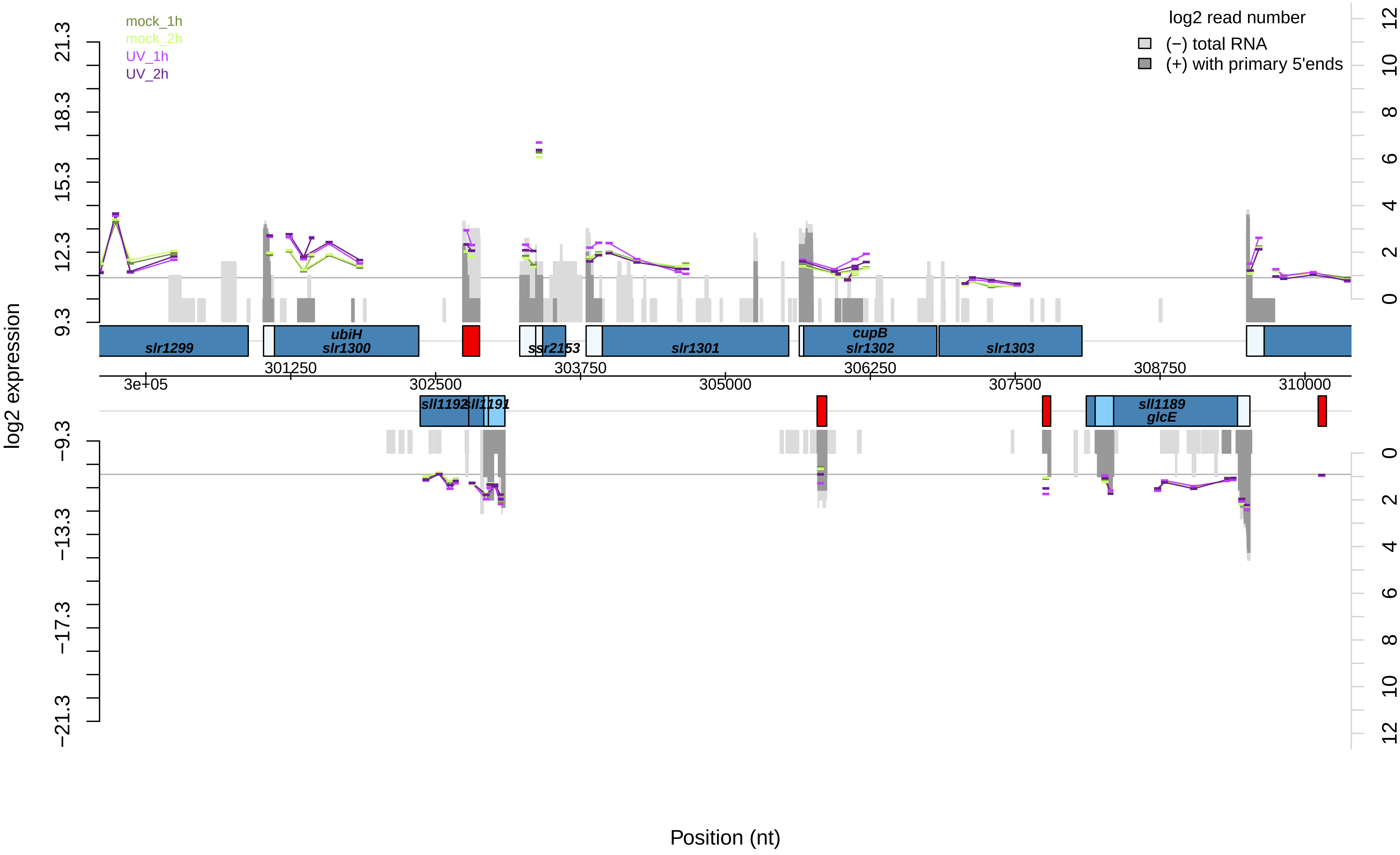

mock_1h
mock_2h
UV_1h
UV_2h
log2 read number
(−) total RNA
(+) with primary 5'ends
log2 expression
21.3
18.3
15.3
12.3
9.3
−9.3
−13.3
−17.3
−21.3
12
10
8
6
4
2
0
slr1299
ubiH
slr1300
ssr2153
slr1301
cupB
slr1302
slr1303
301250
303750
306250
308750
3e+05
302500
305000
307500
310000
sll1192
sll1191
sll1189
glcE
Position (nt)

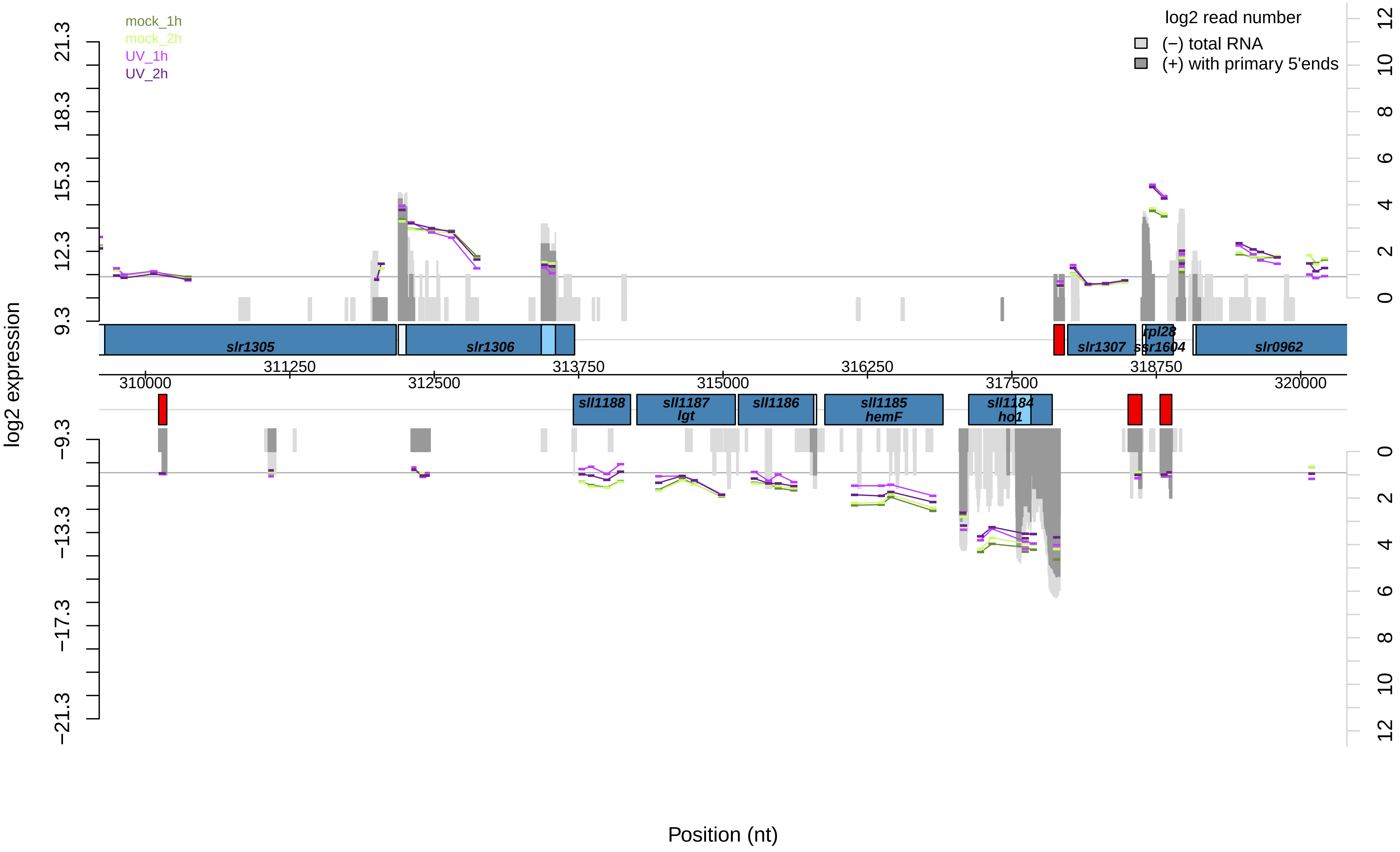

mock_1h
mock_2h
UV_1h
UV_2h
log2 read number
(−) total RNA
(+) with primary 5'ends
log2 expression
Position (nt)
slr1305
slr1306
slr1307
rpl28
ssr1604
slr0962
sll1188
sll1187
lgt
sll1186
sll1185
hemF
sll1184
ho1
310000
311250
312500
313750
315000
316250
317500
318750
320000
21.3
18.3
15.3
12.3
9.3
−9.3
−13.3
−17.3
−21.3

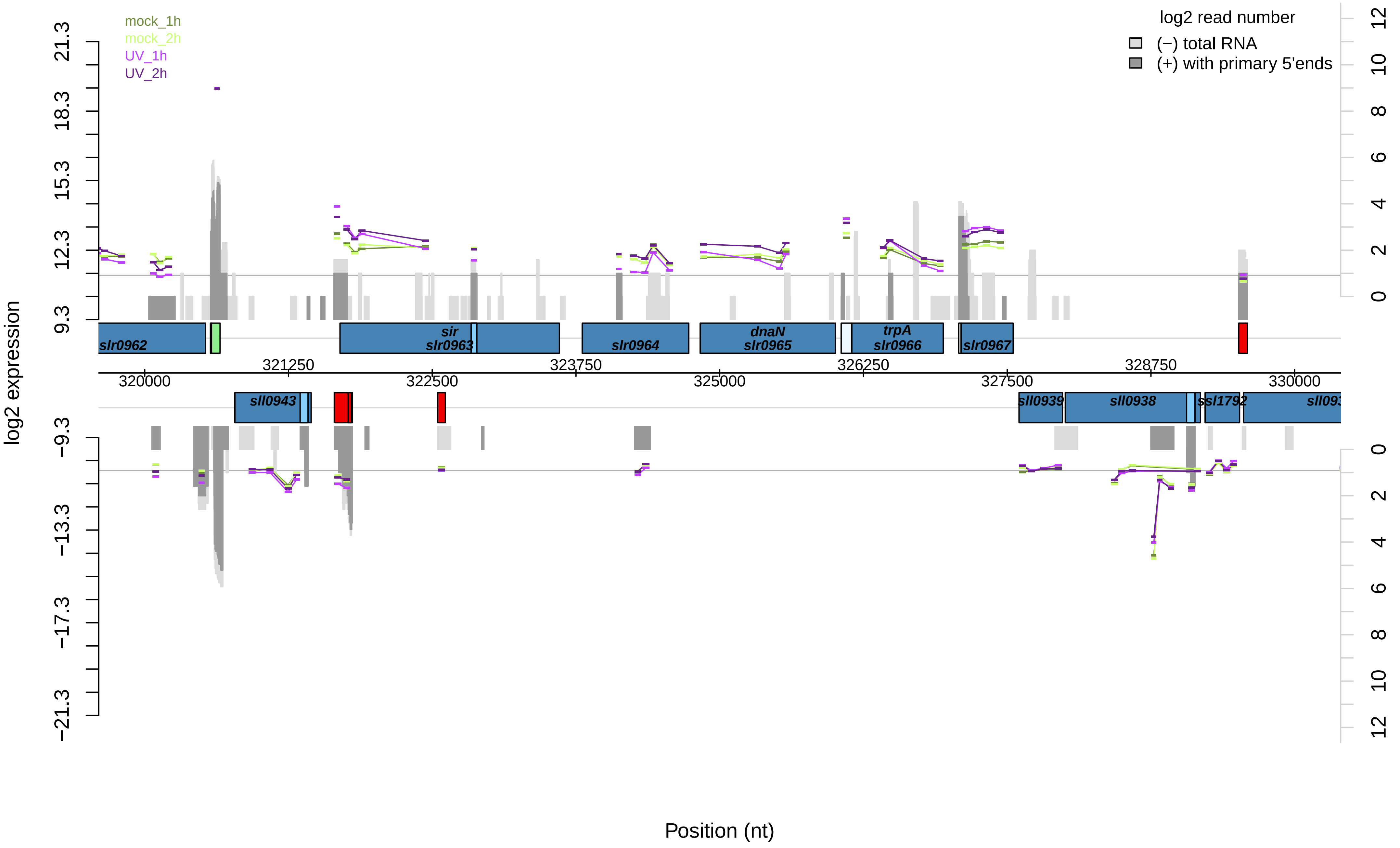

mock_1h
mock_2h
UV_1h
UV_2h
log2 read number
(−) total RNA
(+) with primary 5'ends
log2 expression
21.3
18.3
15.3
12.3
9.3
−9.3
−13.3
−17.3
−21.3
12
10
8
6
4
2
0
slr0962
sir
slr0963
slr0964
dnaN
slr0965
trpA
slr0966
slr0967
sll0943
sll0939
sll0938
ssl1792
320000
321250
322500
323750
325000
326250
327500
328750
330000
Position (nt)

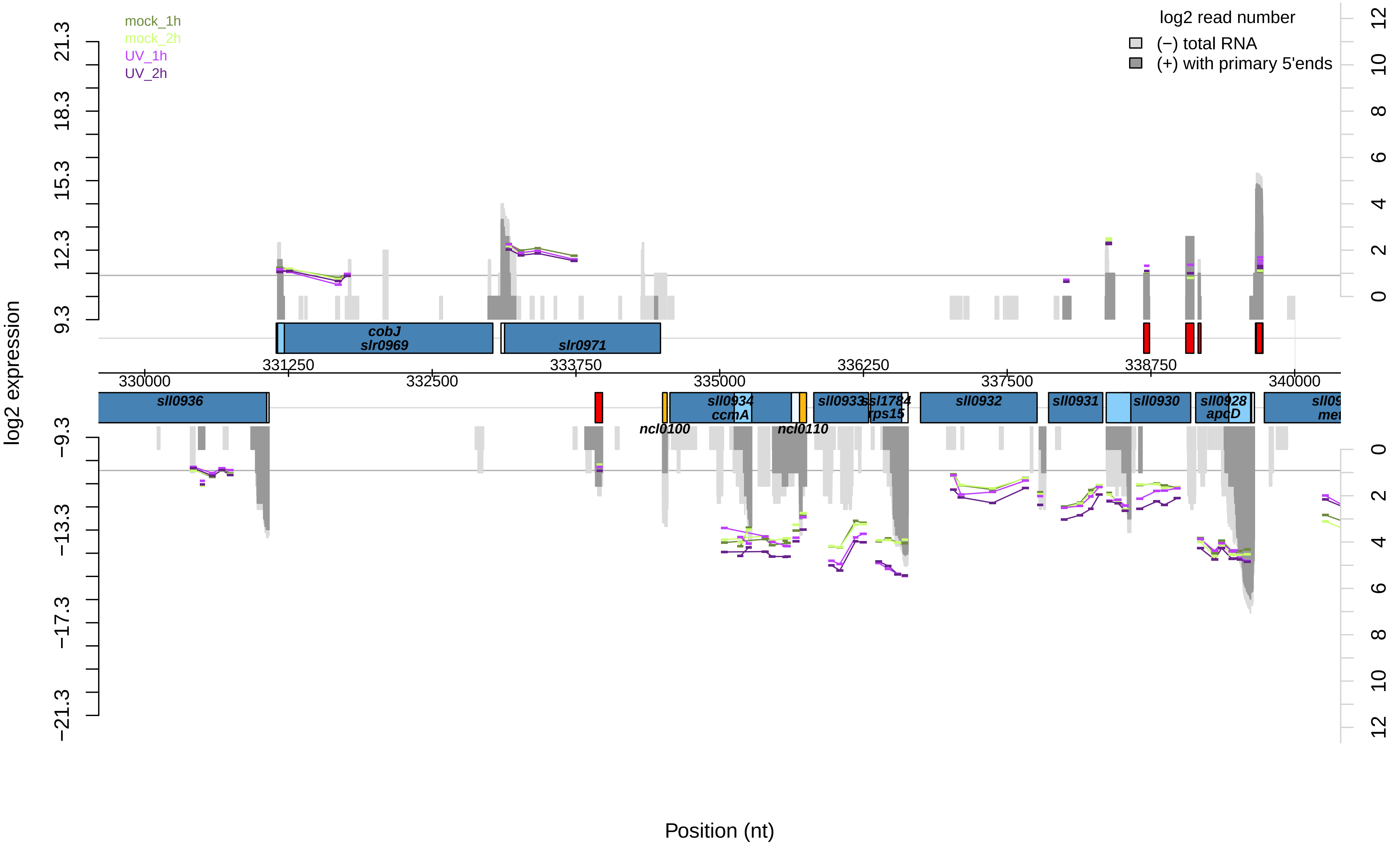

mock_1h
mock_2h
UV_1h
UV_2h
log2 read number
(−) total RNA
(+) with primary 5'ends
log2 expression
21.3
18.3
15.3
12.3
9.3
−9.3
−13.3
−17.3
−21.3
12
10
8
6
4
2
0
cobJ
slr0969
slr0971
331250
333750
336250
338750
330000
332500
335000
337500
340000
sll0936
sll0934
ccmA
ncl0100
ncl0110
sll0933
ssl1784
rps15
sll0932
sll0931
sll0930
sll0928
apcD
Position (nt)

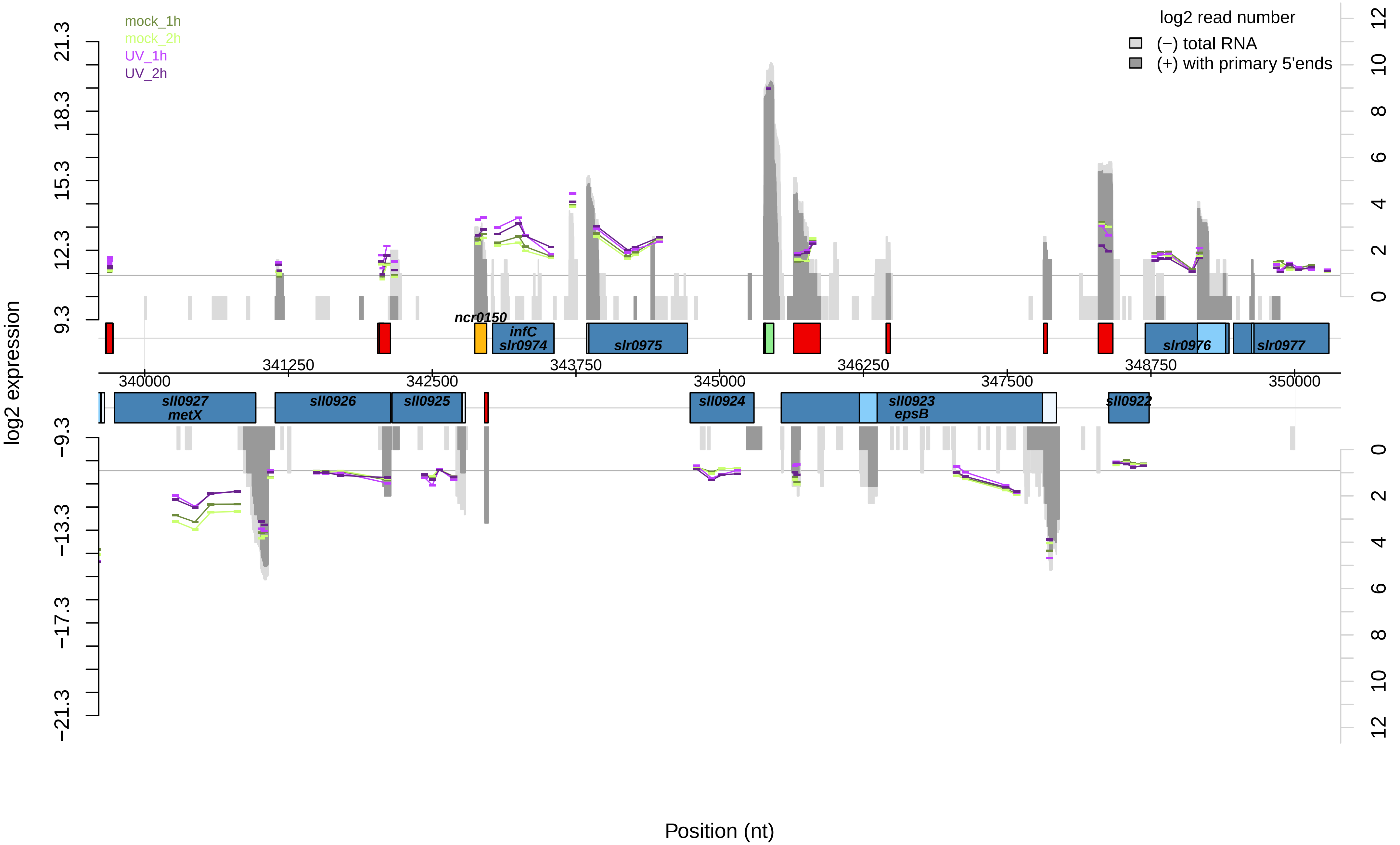

mock_1h
mock_2h
UV_1h
UV_2h
log2 read number
(−) total RNA
(+) with primary 5'ends
log2 expression
21.3
18.3
15.3
12.3
9.3
−9.3
−13.3
−17.3
−21.3
12
10
8
6
4
2
0
ncr0150
infC
slr0974
slr0975
slr0976
slr0977
341250
343750
346250
348750
340000
342500
345000
347500
350000
sll0927
metX
sll0926
sll0925
sll0924
sll0923
epsB
sll0922
Position (nt)

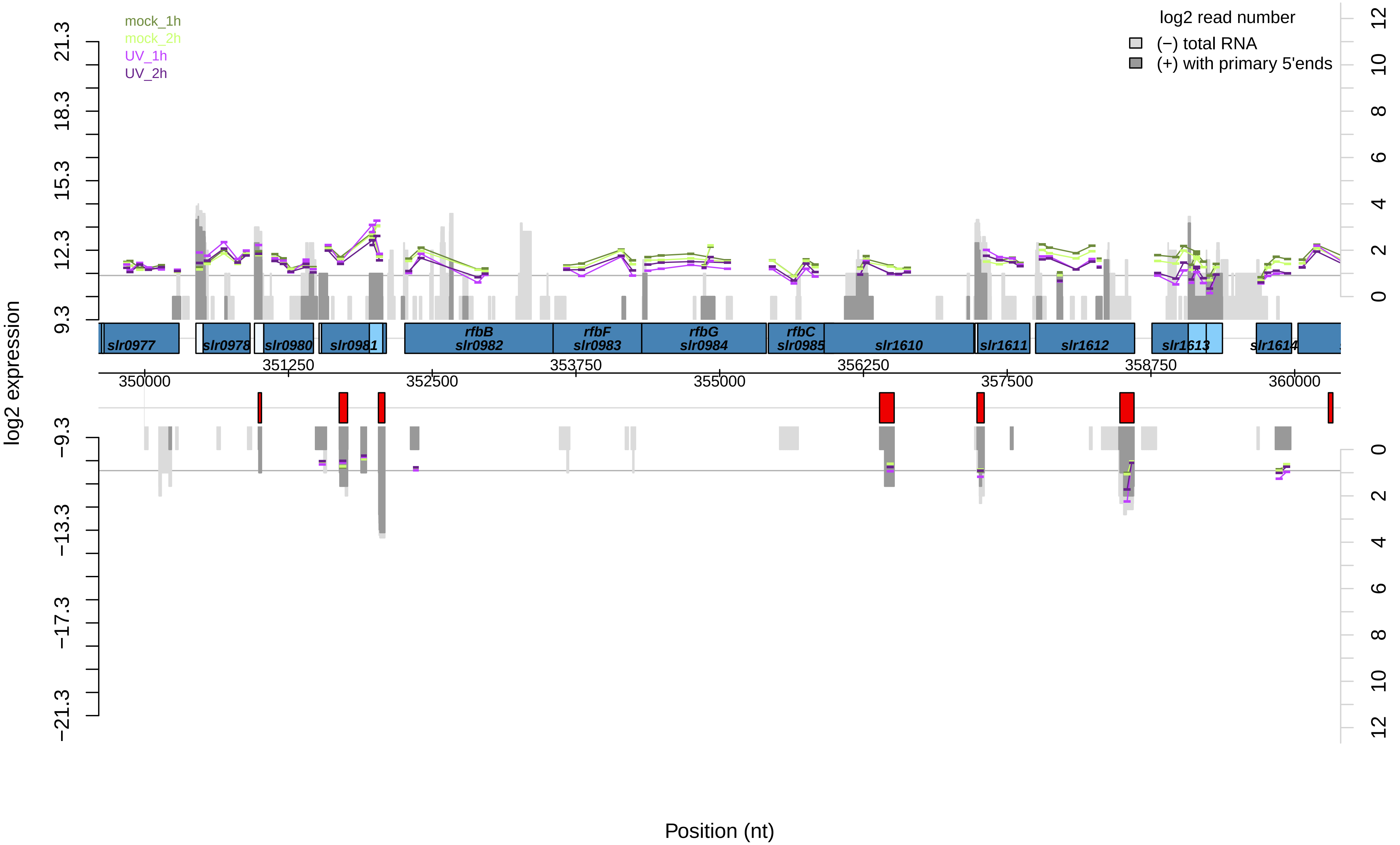

mock_1h
mock_2h
UV_1h
UV_2h
log2 read number
(−) total RNA
(+) with primary 5'ends
log2 expression
21.3
18.3
15.3
12.3
9.3
−9.3
−13.3
−17.3
−21.3
12
10
8
6
4
2
0
slr0977
slr0978
slr0980
slr0981
rfbB
slr0982
rfbF
slr0983
rfbG
slr0984
rfbC
slr0985
slr1610
slr1611
slr1612
slr1613
slr1614
350000
351250
352500
353750
355000
356250
357500
358750
360000
Position (nt)

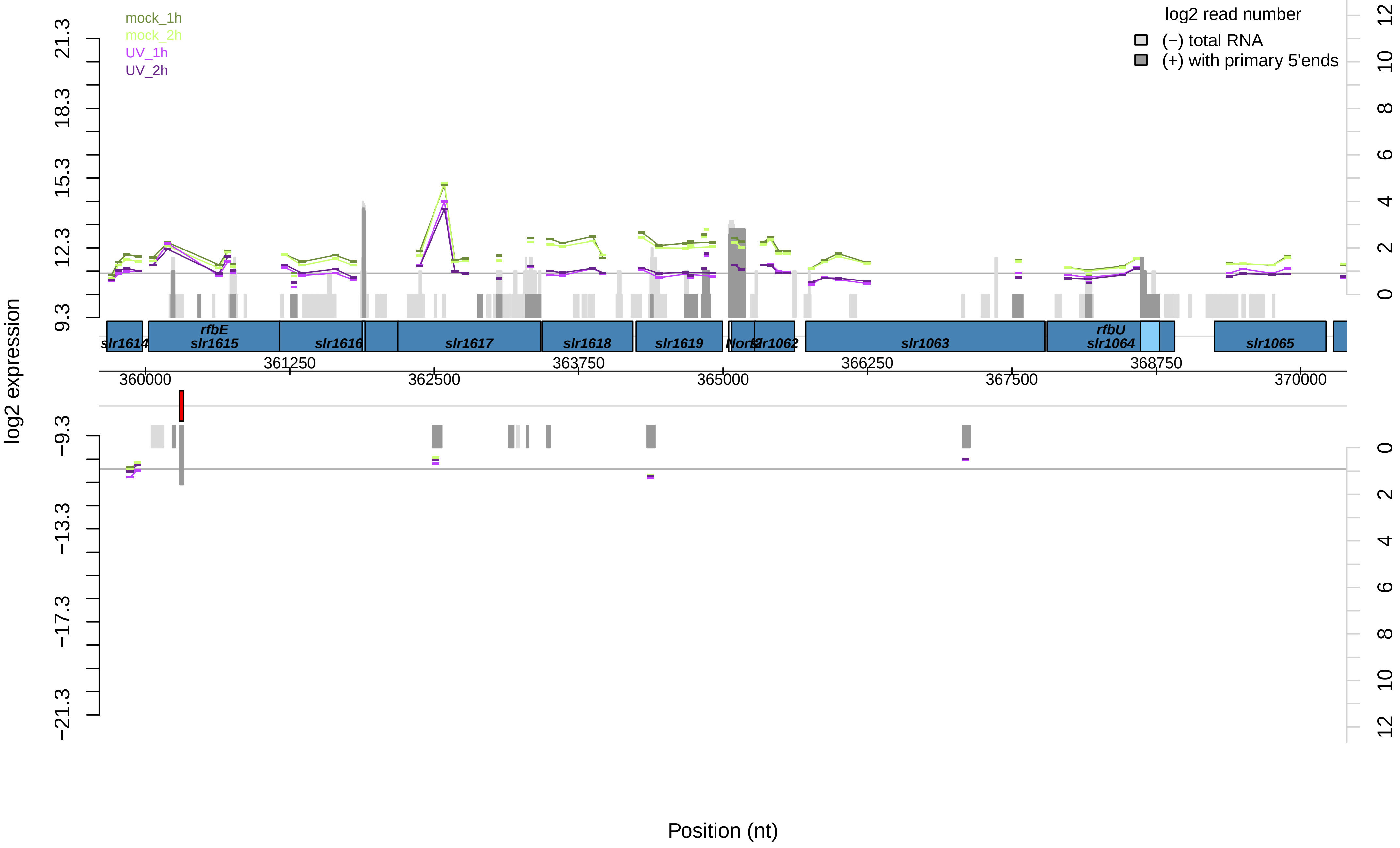

mock_1h
mock_2h
UV_1h
UV_2h
log2 read number
(−) total RNA
(+) with primary 5'ends
log2 expression
21.3
18.3
15.3
12.3
9.3
−9.3
−13.3
−17.3
−21.3
12
10
8
6
4
2
0
slr1614
rfbE
slr1615
slr1616
slr1617
slr1618
slr1619
slr1062
slr1063
rfbU
slr1064
slr1065
361250
363750
366250
368750
360000
362500
365000
367500
370000
Position (nt)

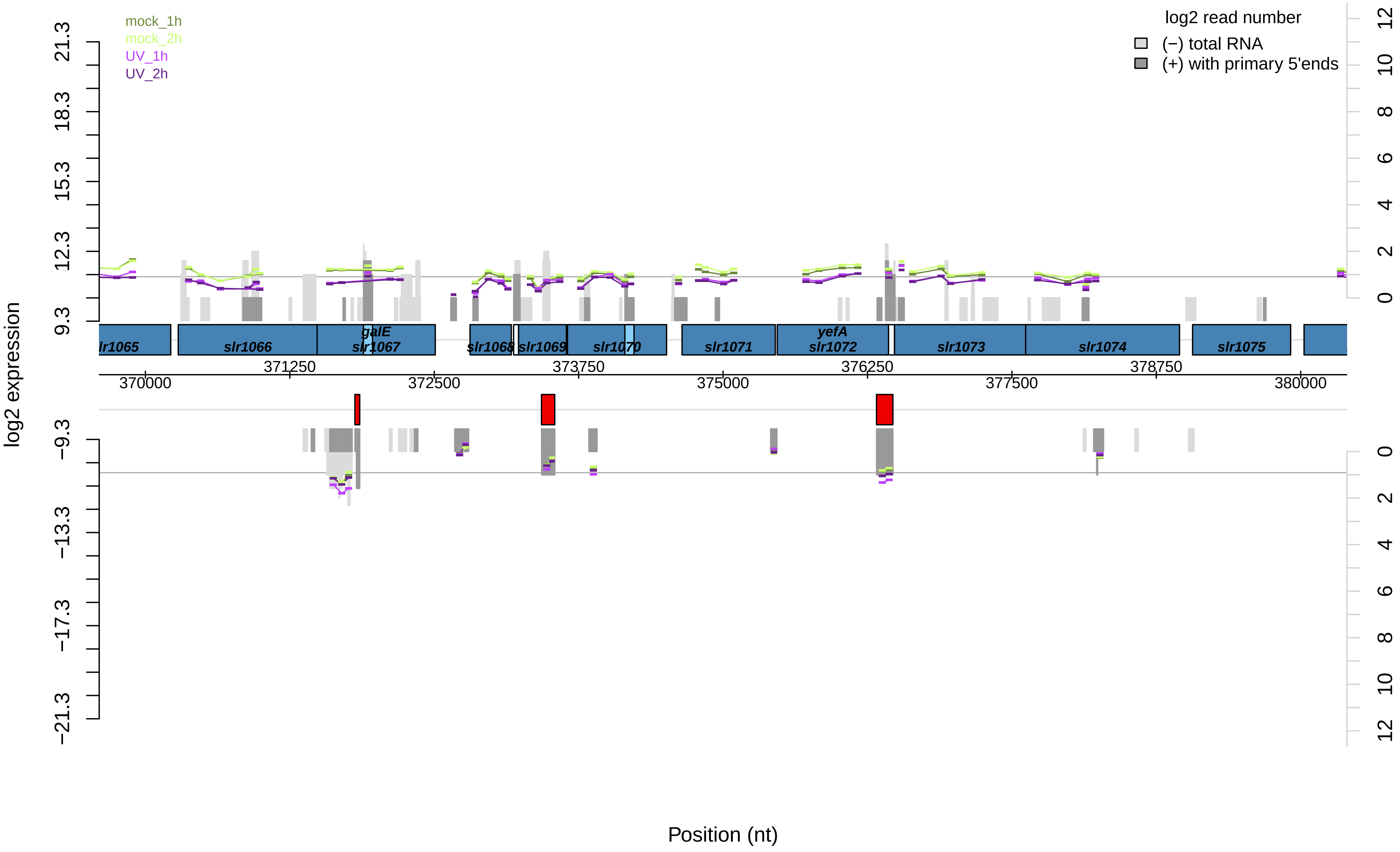

mock_1h
mock_2h
UV_1h
UV_2h
log2 read number
(−) total RNA
(+) with primary 5'ends
log2 expression
21.3
18.3
15.3
12.3
9.3
−9.3
−13.3
−17.3
−21.3
12
10
8
6
4
2
0
lr1065
slr1066
galE
slr1067
slr1068
slr1069
slr1070
slr1071
yefA
slr1072
slr1073
slr1074
slr1075
371250
373750
376250
378750
370000
372500
375000
377500
380000
Position (nt)

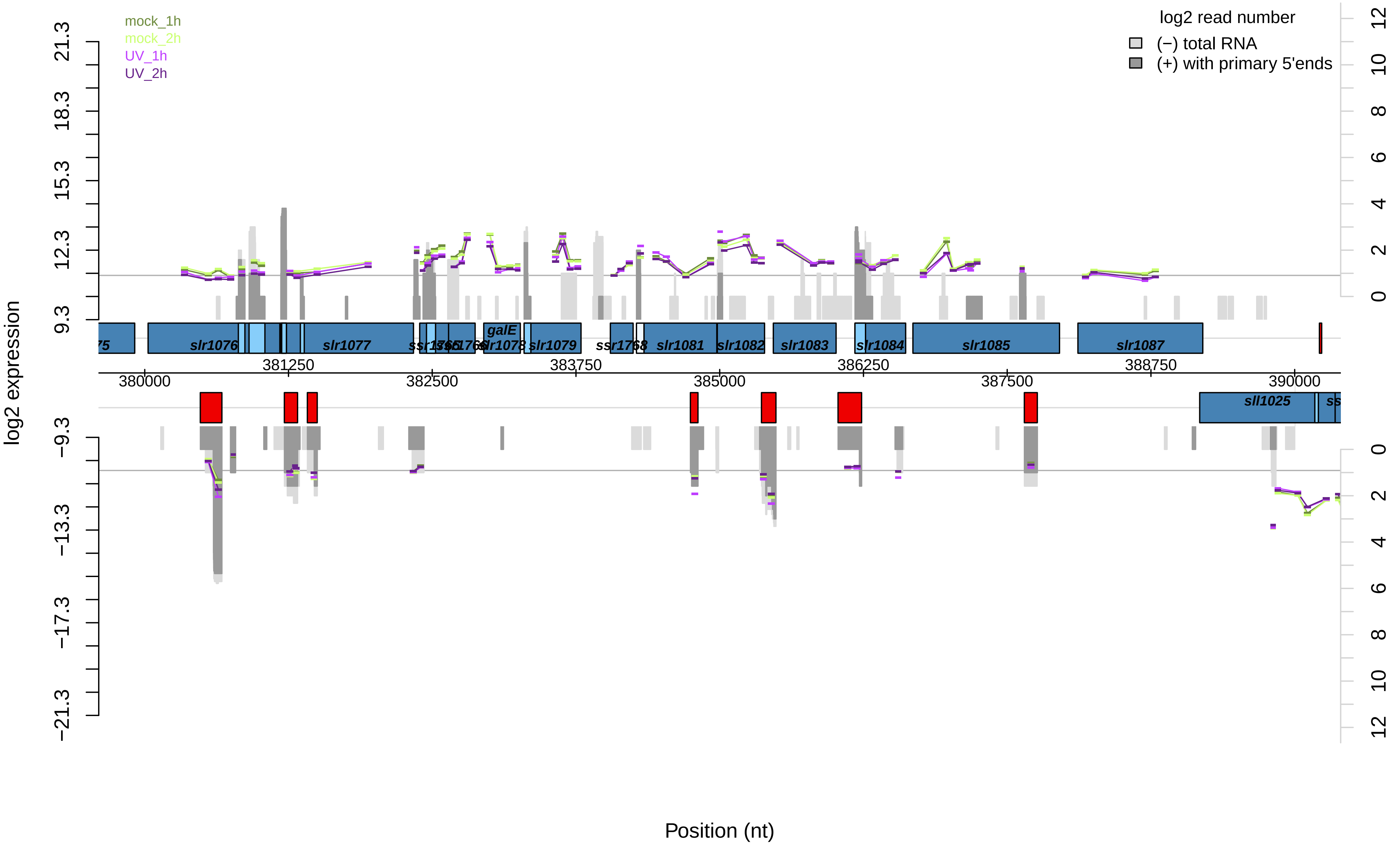

mock_1h
mock_2h
UV_1h
UV_2h
log2 read number
(−) total RNA
(+) with primary 5'ends
log2 expression
21.3
18.3
15.3
12.3
9.3
−9.3
−13.3
−17.3
−21.3
12
10
8
6
4
2
0
slr1076
slr1077
galE
slr1078
slr1079
ssr1768
slr1081
slr1082
slr1083
slr1084
slr1085
slr1087
sll1025
380000
381250
382500
383750
385000
386250
387500
388750
390000
Position (nt)

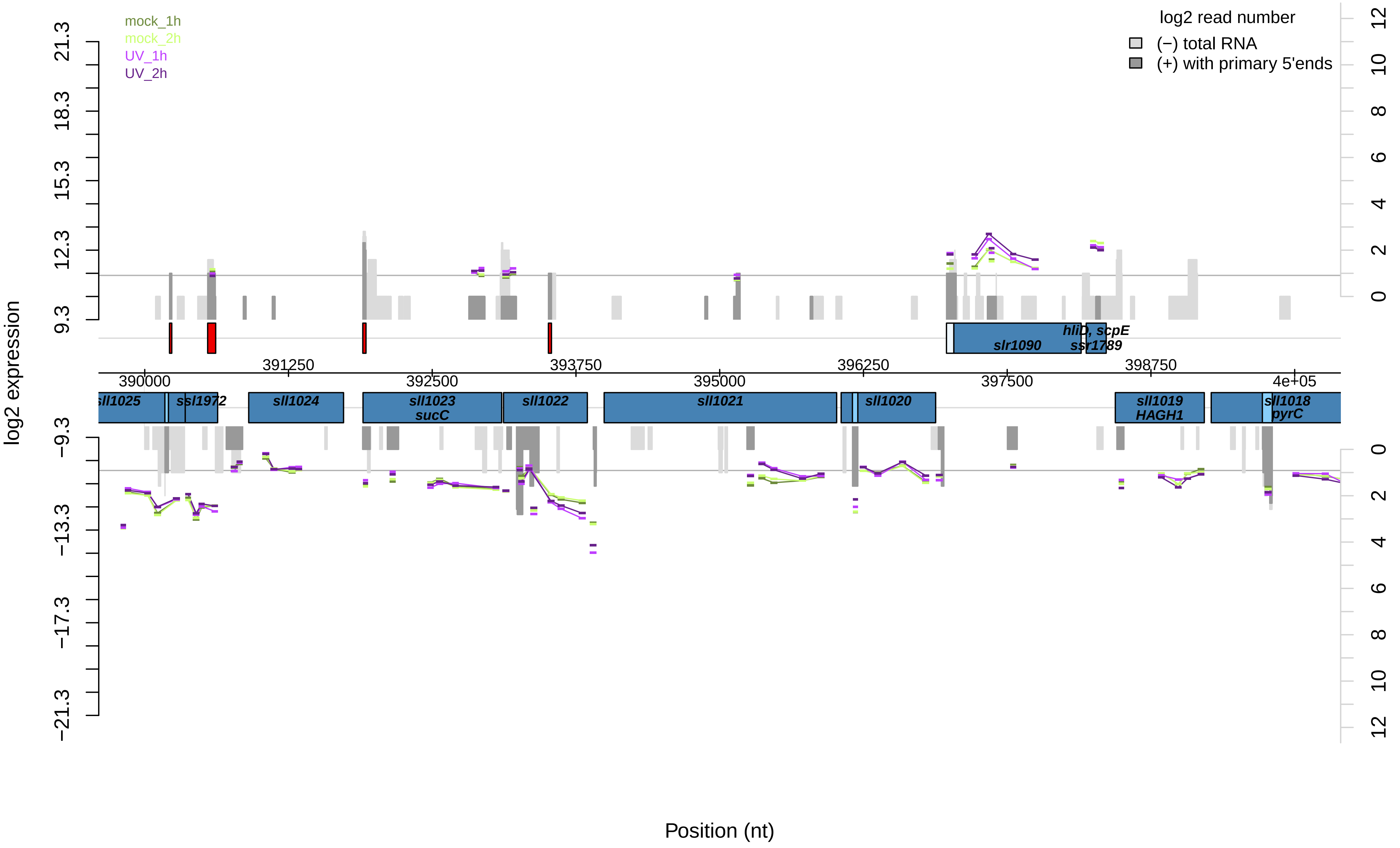
mock_1h
mock_2h
UV_1h
UV_2h
log2 read number
(−) total RNA
(+) with primary 5'ends
log2 expression
21.3
18.3
15.3
12.3
9.3
−9.3
−13.3
−17.3
−21.3
12
10
8
6
4
2
0
hliD, scpE
slr1090
ssr1789
391250
393750
396250
398750
390000
392500
395000
397500
4e+05
sll1025
ssl1972
sll1024
sll1023
succ
sll1022
sll1021
sll1020
sll1019
HAGH1
sll1018
pyrC
Position (nt)

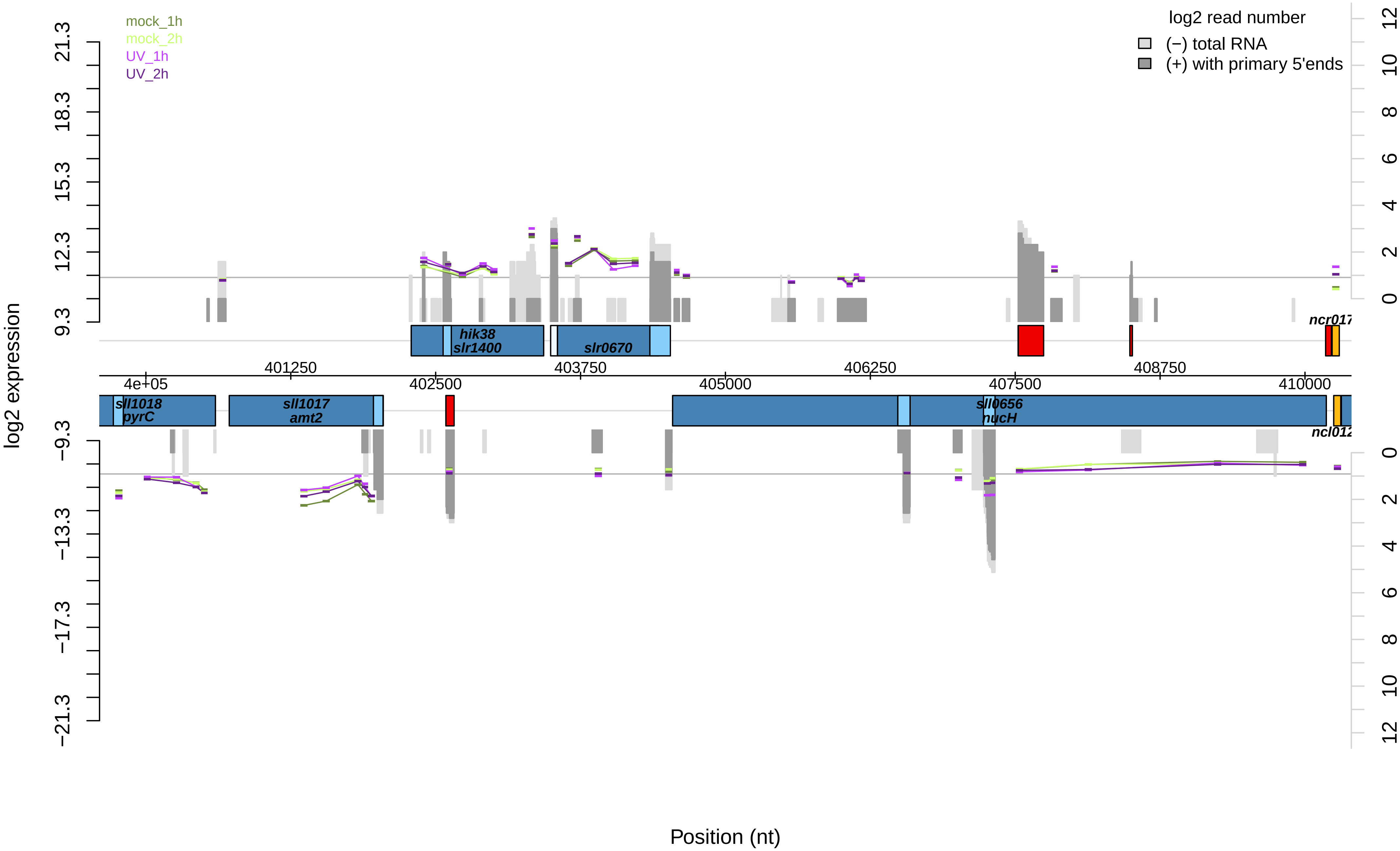

mock_1h
mock_2h
UV_1h
UV_2h
log2 read number
(−) total RNA
(+) with primary 5'ends
log2 expression
21.3
18.3
15.3
12.3
9.3
−9.3
−13.3
−17.3
−21.3
0
2
4
6
8
10
12
hik38
slr1400
slr0670
ncr017
401250
403750
406250
408750
4e+05
402500
405000
407500
410000
sll1018
pyrC
sll1017
amt2
sll0656
nucH
ncl012
Position (nt)

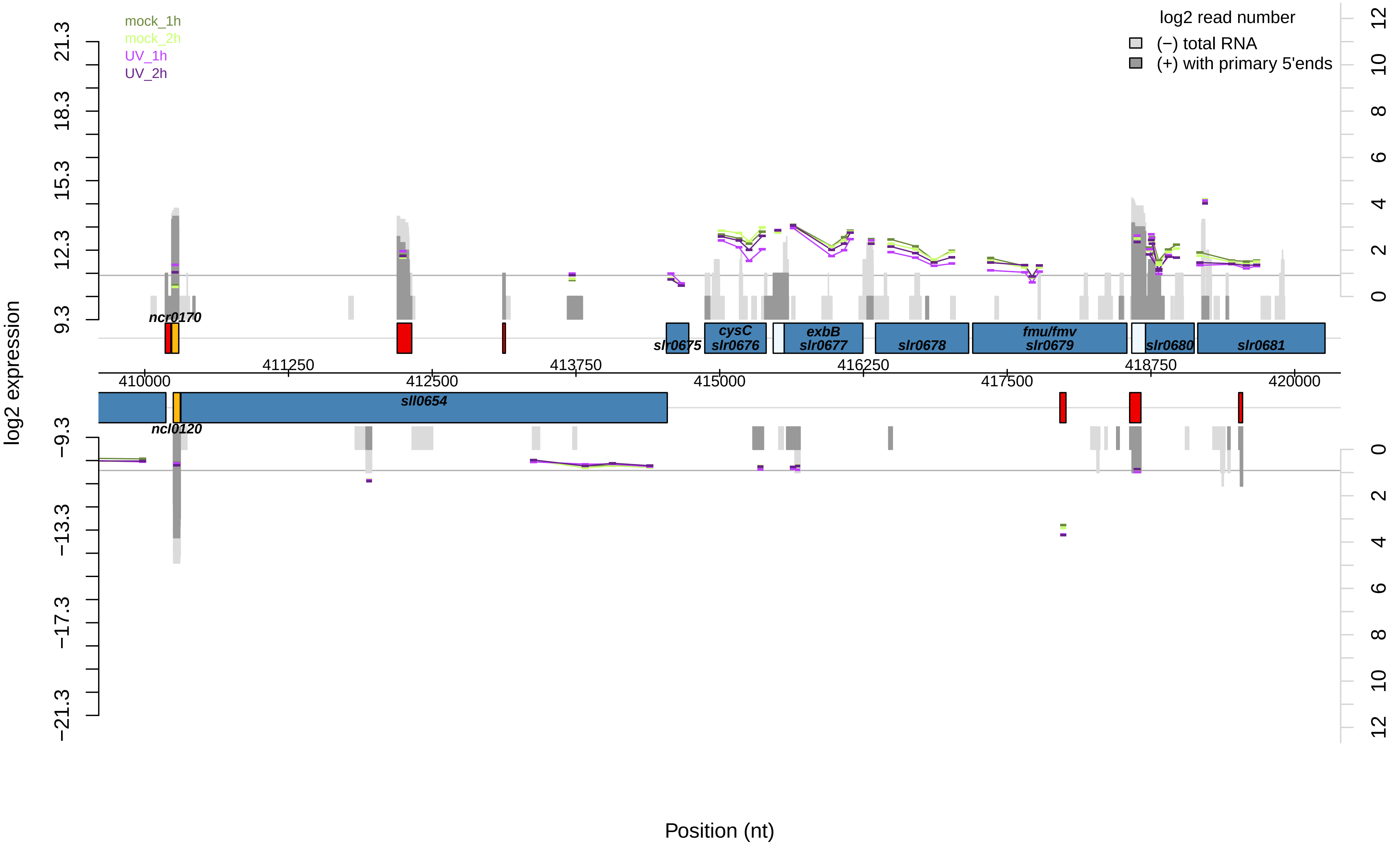

mock_1h
mock_2h
UV_1h
UV_2h
log2 read number
(−) total RNA
(+) with primary 5'ends
log2 expression
21.3
18.3
15.3
12.3
9.3
−9.3
−13.3
−17.3
−21.3
12
10
8
6
4
2
0
ncr0170
ncl0120
slr0675
cysC
slr0676
exbB
slr0677
slr0678
fmu/fmv
slr0679
slr0680
slr0681
sll0654
410000
411250
412500
413750
415000
416250
417500
418750
420000
Position (nt)

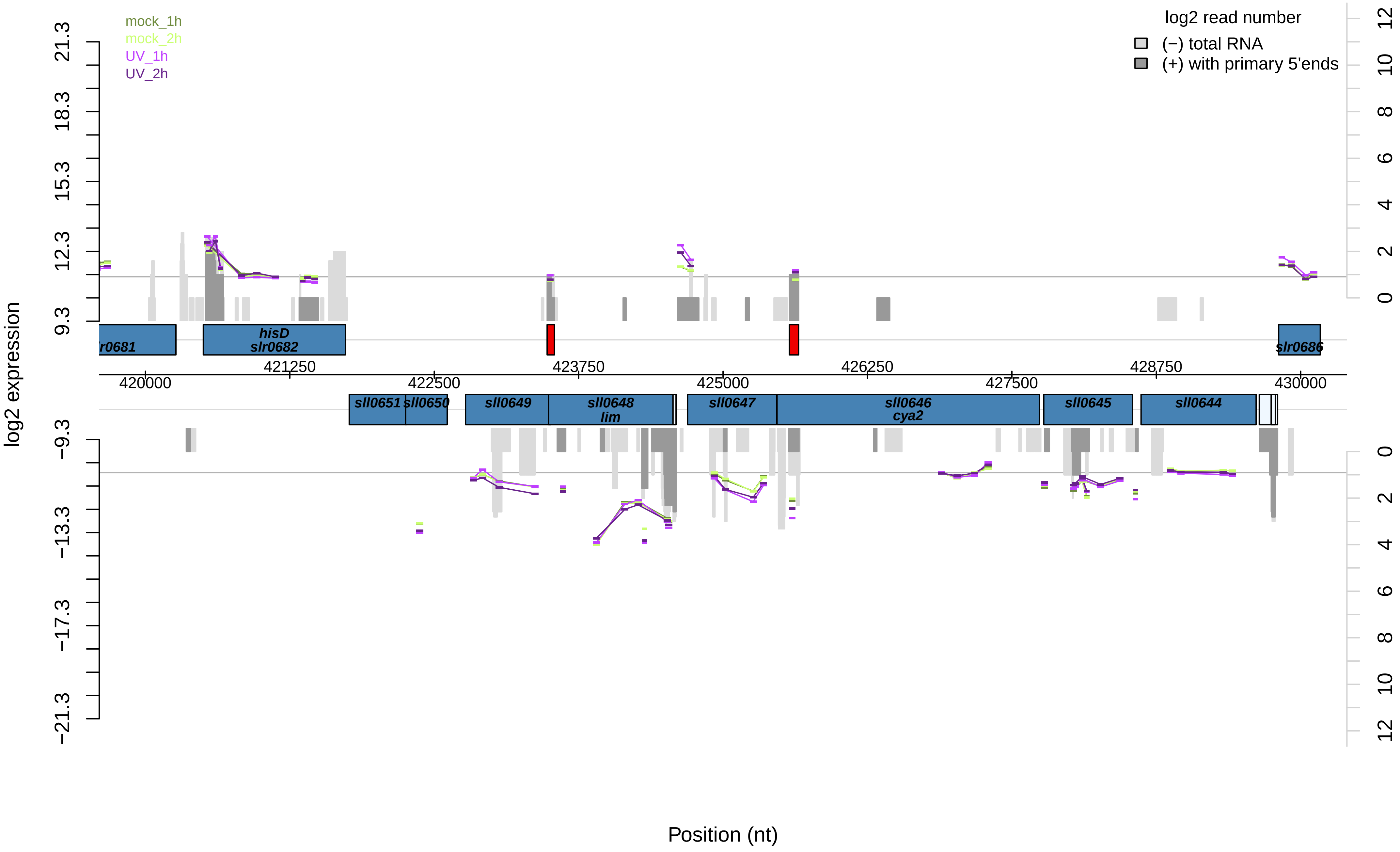
mock_1h
mock_2h
UV_1h
UV_2h
log2 read number
(−) total RNA
(+) with primary 5'ends
log2 expression
21.3
18.3
15.3
12.3
9.3
−9.3
−13.3
−17.3
−21.3
12
10
8
6
4
2
0
r0681
hisD
slr0682
slr0686
421250
423750
426250
428750
420000
422500
425000
427500
430000
sll0651
sll0650
sll0649
sll0648
lim
sll0647
sll0646
cya2
sll0645
sll0644
Position (nt)

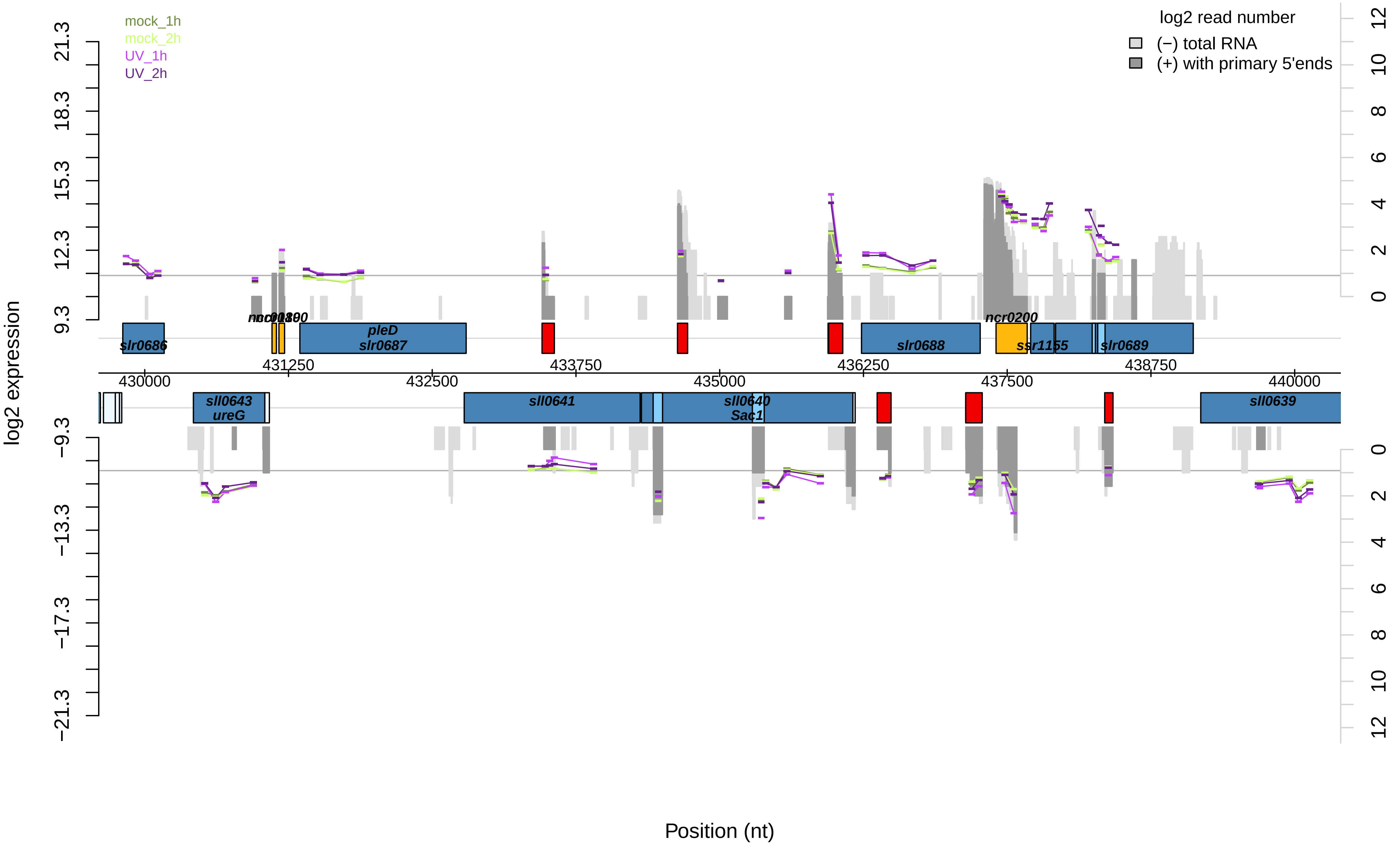

mock_1h
mock_2h
UV_1h
UV_2h
log2 read number
(−) total RNA
(+) with primary 5'ends
log2 expression
21.3
18.3
15.3
12.3
9.3
−9.3
−13.3
−17.3
−21.3
12
10
8
6
4
2
0
slr0686
ncr0180
ncr0190
pleD
slr0687
slr0688
ncr0200
ssr1155
slr0689
431250
433750
436250
438750
430000
432500
435000
437500
440000
sll0643
ureG
sll0641
sll0640
Sac1
sll0639
Position (nt)

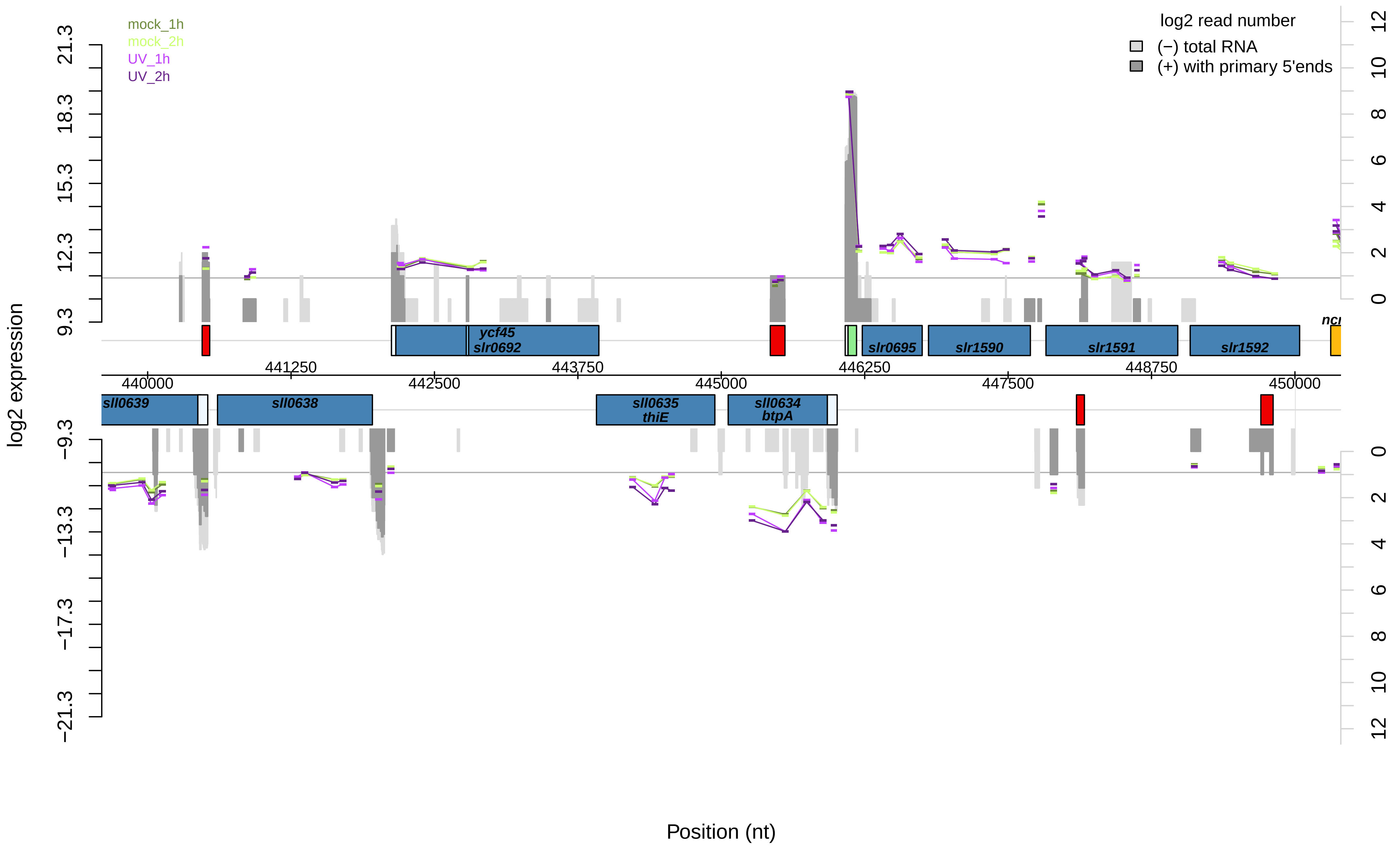

mock_1h
mock_2h
UV_1h
UV_2h
log2 read number
(−) total RNA
(+) with primary 5'ends
log2 expression
21.3
18.3
15.3
12.3
9.3
−9.3
−13.3
−17.3
−21.3
12
10
8
6
4
2
0
ycf45
slr0692
slr0695
slr1590
slr1591
slr1592
nci
441250
443750
446250
448750
440000
442500
445000
447500
450000
sll0639
sll0638
sll0635
thiE
sll0634
btpA
Position (nt)

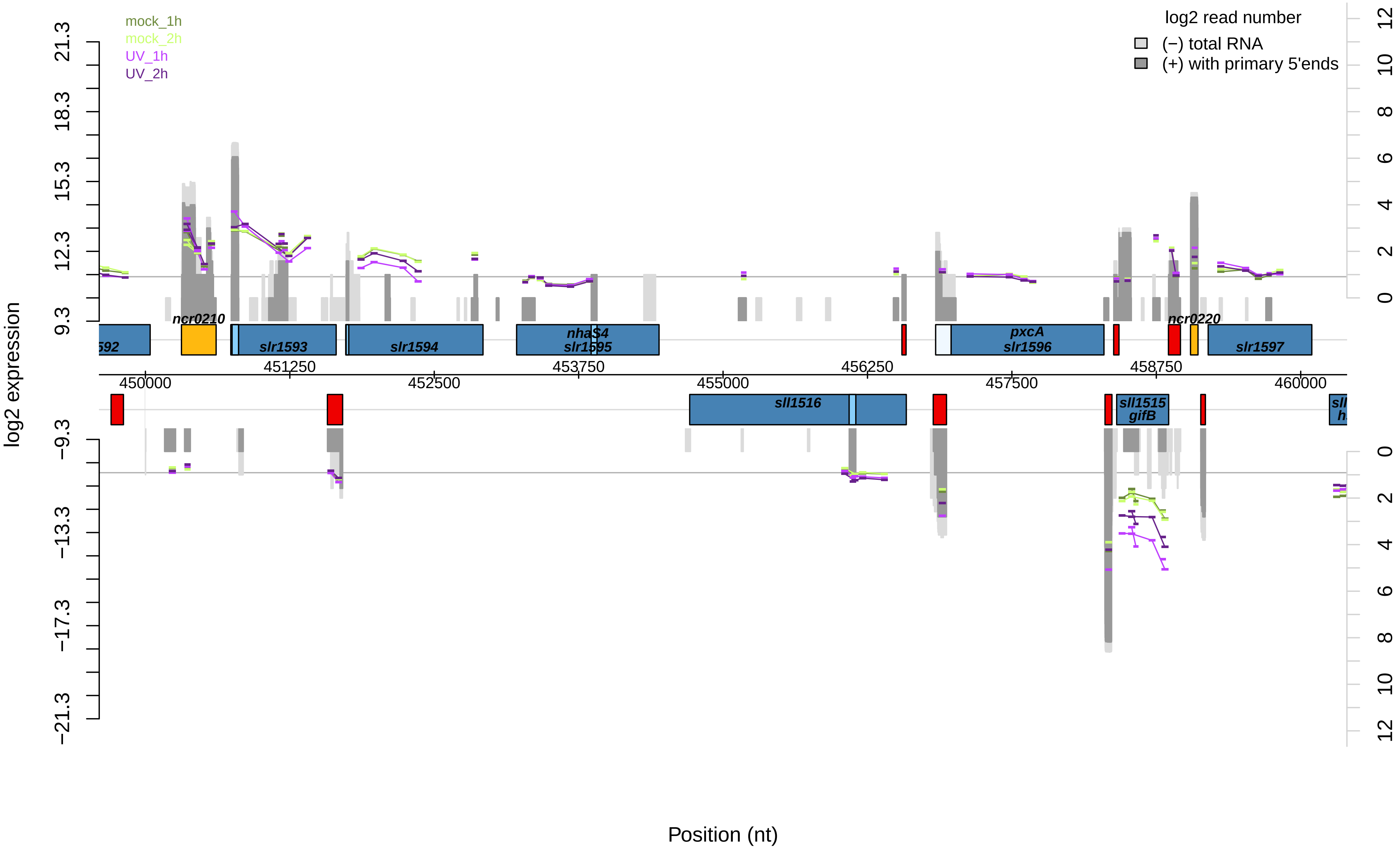

mock_1h
mock_2h
UV_1h
UV_2h
log2 read number
(−) total RNA
(+) with primary 5'ends
log2 expression
21.3
18.3
15.3
12.3
9.3
−9.3
−13.3
−17.3
−21.3
12
10
8
6
4
2
0
ncr0210
slr1593
slr1594
nhaS4
slr1595
pxcA
slr1596
ncr0220
slr1597
sll1516
sll1515
gifB
450000
451250
452500
453750
455000
456250
457500
458750
460000
Position (nt)

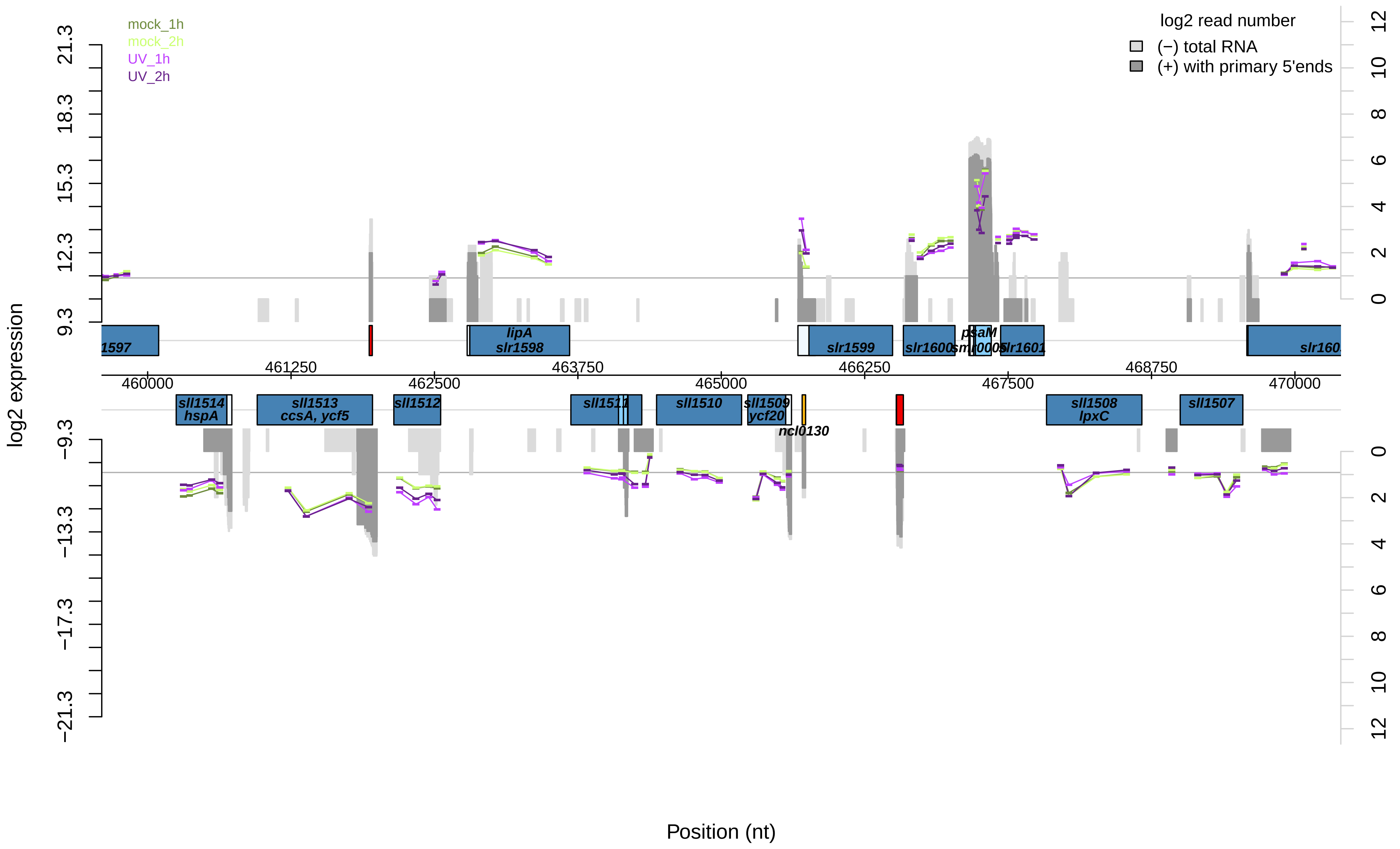

mock_1h
mock_2h
UV_1h
UV_2h
log2 read number
(−) total RNA
(+) with primary 5'ends
log2 expression
Position (nt)
sll1514 hspA
sll1513 ccsA, ycf5
sll1512
lipA slr1598
sll1511
sll1510
sll1509 ycf20
ncl0130
slr1599
slr1600
psaM
slr1601
sll1508 lpxC
sll1507
460000
461250
462500
463750
465000
466250
467500
468750
470000

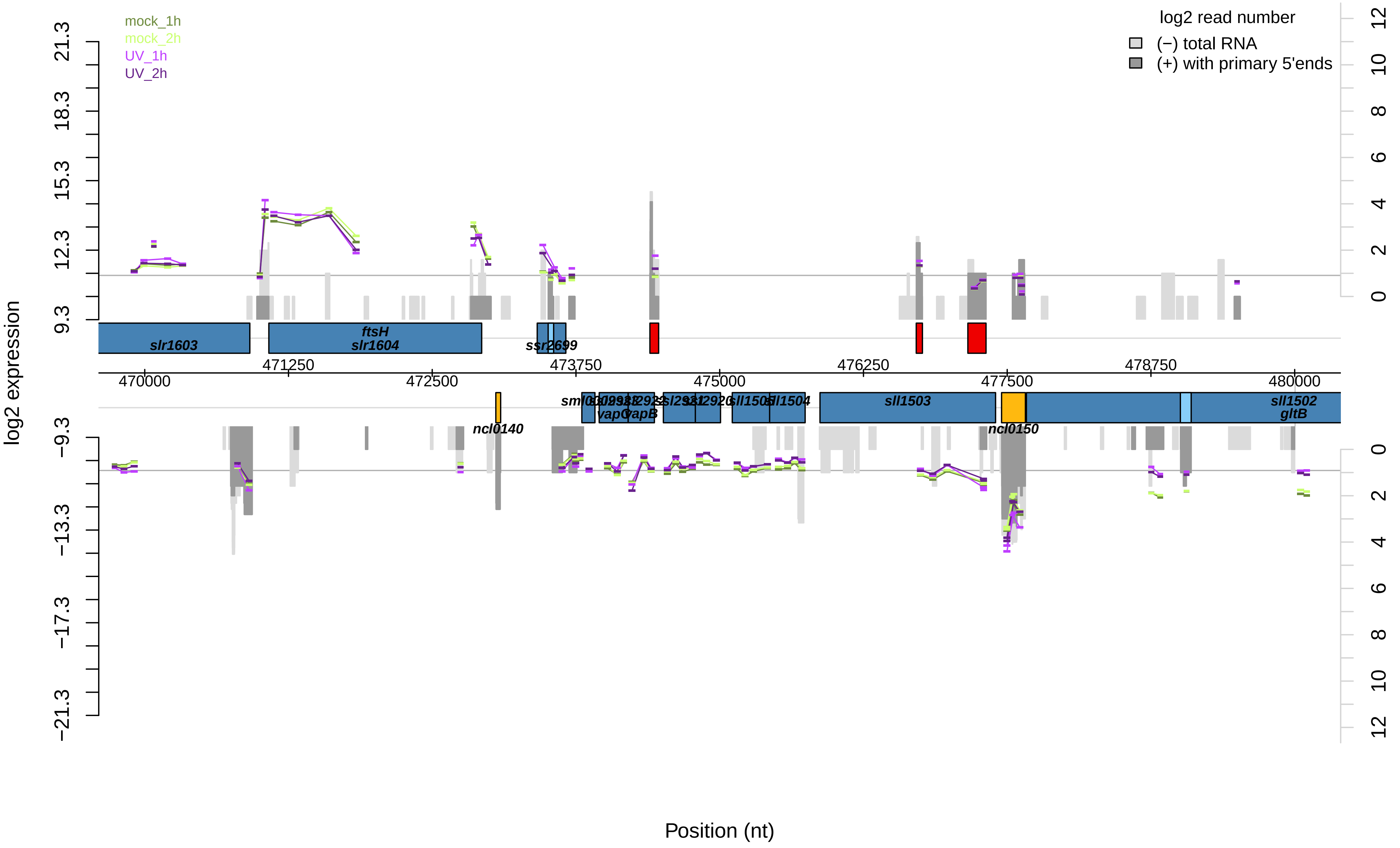

mock_1h
mock_2h
UV_1h
UV_2h
log2 read number
(−) total RNA
(+) with primary 5'ends
log2 expression
21.3
18.3
15.3
12.3
9.3
−9.3
−13.3
−17.3
−21.3
12
10
8
6
4
2
0
slr1603
ftsH
slr1604
ssr2699
ncl0140
ncl0150
sll1503
sll1502
gltB
sll1504
470000
471250
472500
473750
475000
476250
477500
478750
480000
Position (nt)

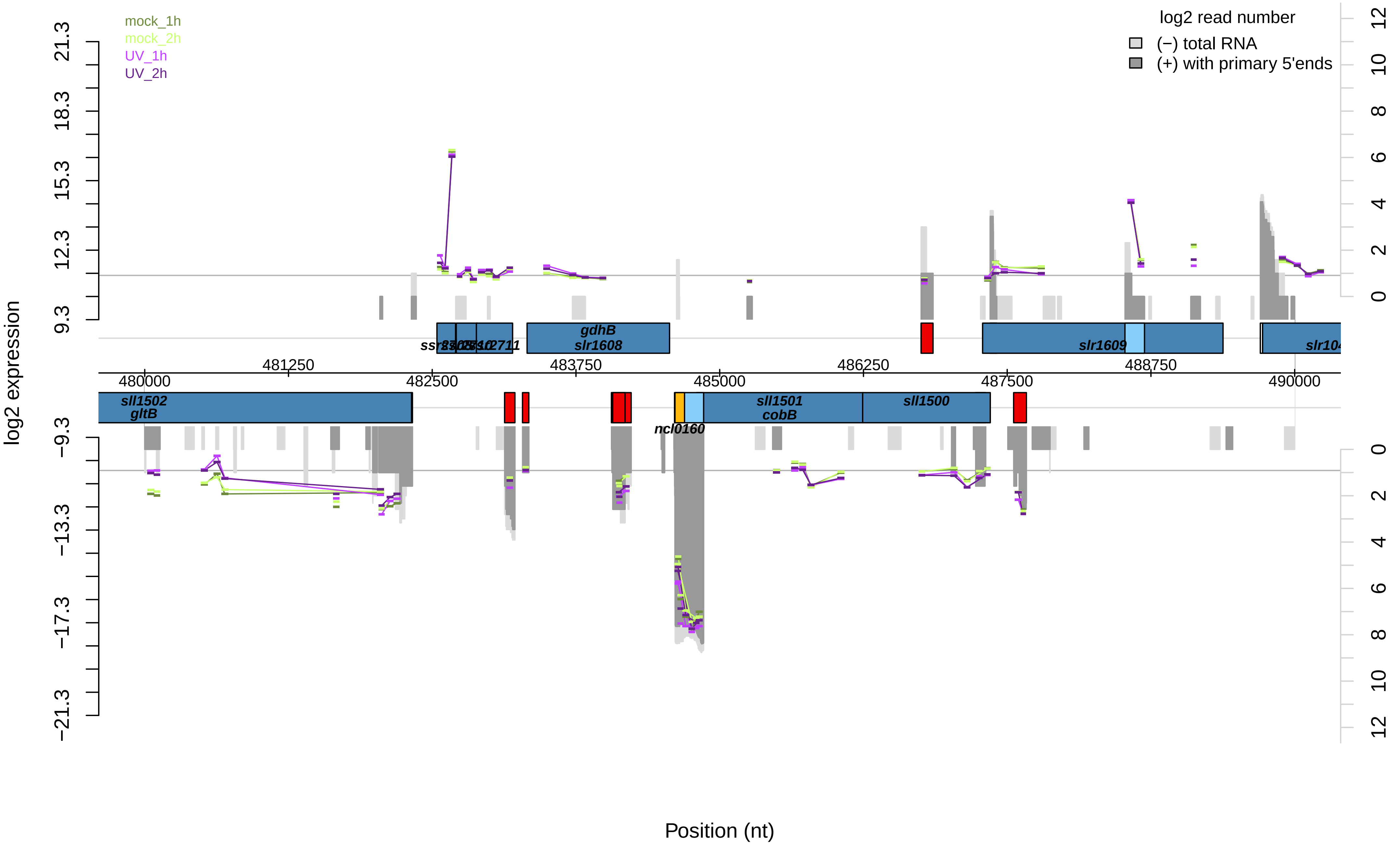

mock_1h
mock_2h
UV_1h
UV_2h
log2 read number
(−) total RNA
(+) with primary 5'ends
log2 expression
21.3
18.3
15.3
12.3
9.3
−9.3
−13.3
−17.3
−21.3
12
10
8
6
4
2
0
gdhB
slr1608
slr1609
480000
481250
482500
483750
485000
486250
487500
488750
490000
sll1502
gltB
sll1501
cobB
sll1500
ncl0160
Position (nt)

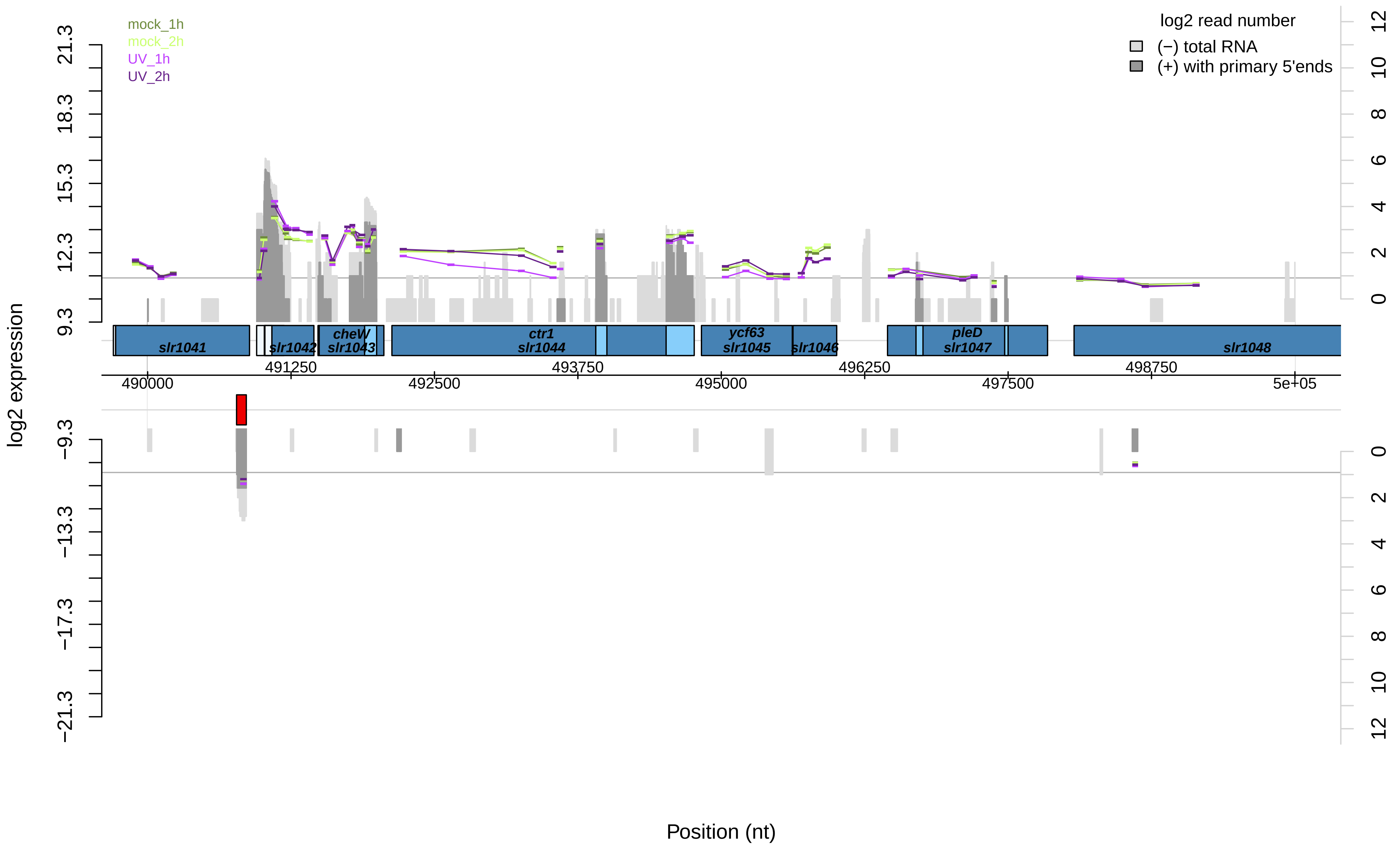

mock_1h
mock_2h
UV_1h
UV_2h
log2 read number
(−) total RNA
(+) with primary 5'ends
log2 expression
21.3
18.3
15.3
12.3
9.3
−9.3
−13.3
−17.3
−21.3
12
10
8
6
4
2
0
slr1041
slr1042
cheW
slr1043
ctr1
slr1044
ycf63
slr1045
slr1046
pleD
slr1047
slr1048
490000
491250
492500
493750
495000
496250
497500
498750
5e+05
Position (nt)

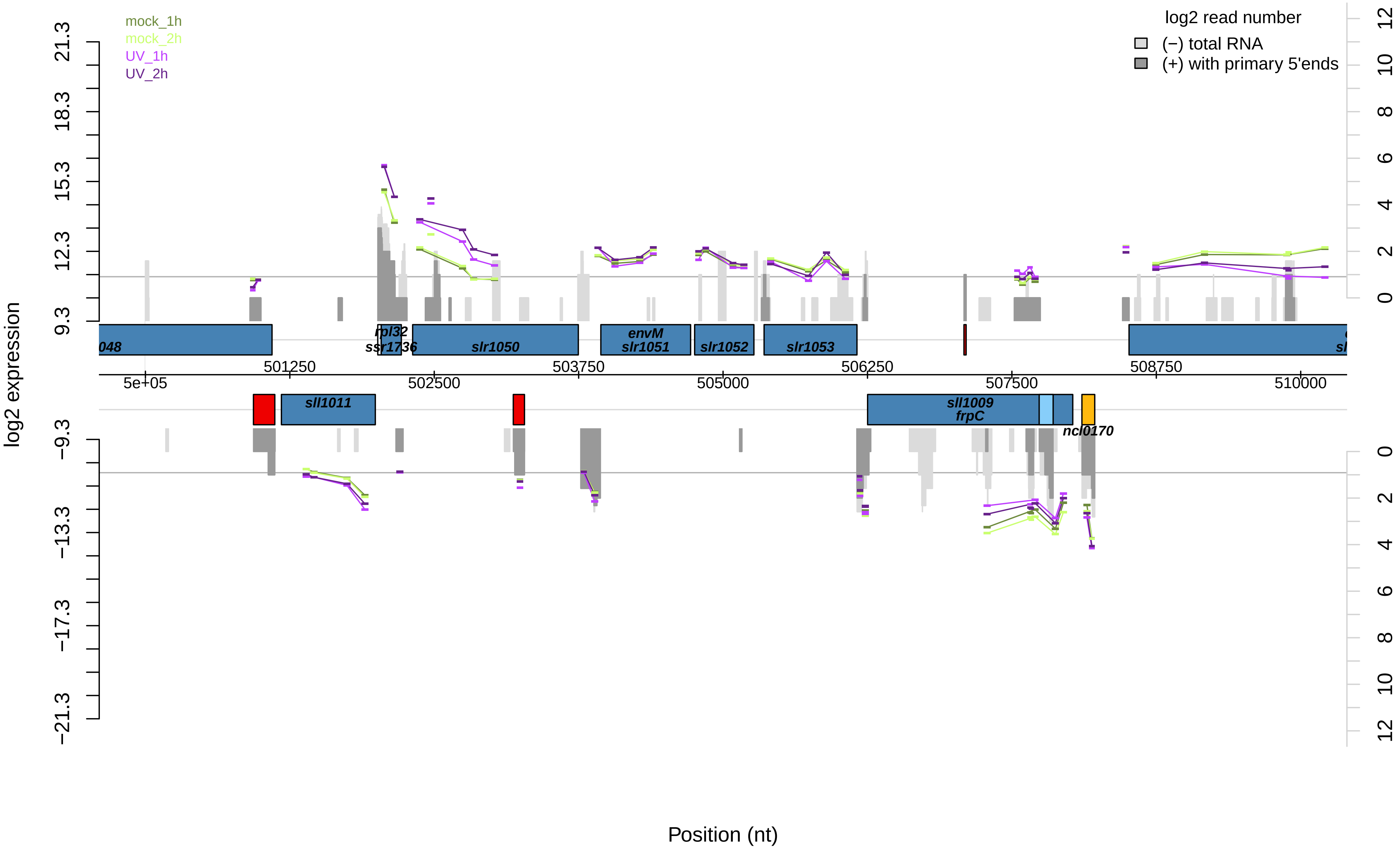

mock_1h
mock_2h
UV_1h
UV_2h
log2 read number
(−) total RNA
(+) with primary 5'ends
log2 expression
Position (nt)
rpl32
ssr1736
slr1050
envM
slr1051
slr1052
slr1053
sll1011
sll1009
frpC
ncl0170
048
5e+05
501250
502500
503750
505000
506250
507500
508750
510000
21.3
18.3
15.3
12.3
9.3
−9.3
−13.3
−17.3
−21.3

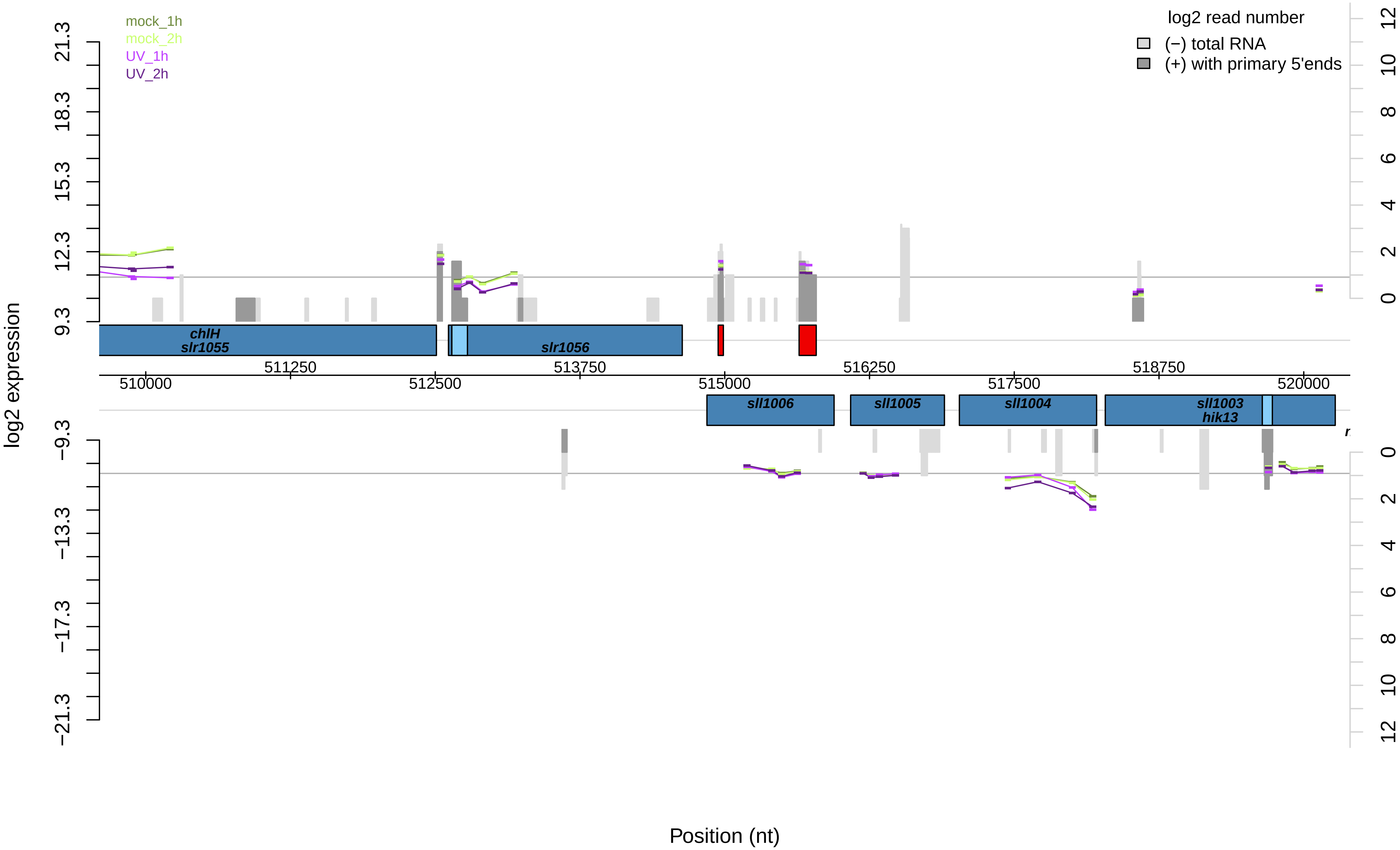

mock_1h
mock_2h
UV_1h
UV_2h
log2 read number
(−) total RNA
(+) with primary 5'ends
log2 expression
21.3
18.3
15.3
12.3
9.3
−9.3
−13.3
−17.3
−21.3
12
10
8
6
4
2
0
chlH
slr1055
slr1056
511250
513750
516250
518750
510000
512500
515000
517500
520000
sll1006
sll1005
sll1004
sll1003
hik13
Position (nt)

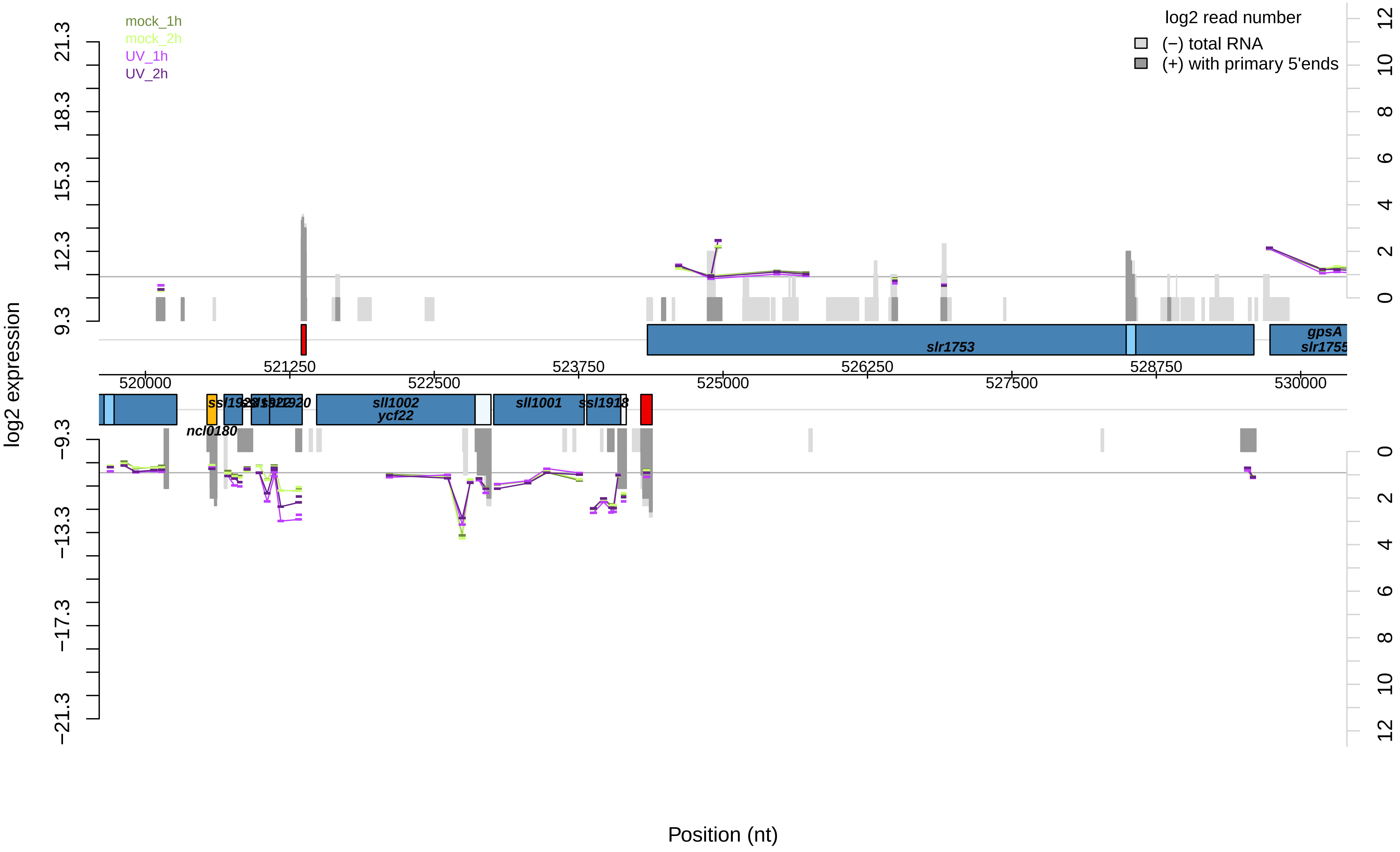
mock_1h
mock_2h
UV_1h
UV_2h
log2 read number
(−) total RNA
(+) with primary 5'ends
log2 expression
21.3
18.3
15.3
12.3
9.3
−9.3
−13.3
−17.3
−21.3
12
10
8
6
4
2
0
slr1753
gpsA
slr1755
521250
523750
526250
528750
520000
522500
525000
527500
530000
ssl1922
ssl1920
sll1002
ycf22
sll1001
ssl1918
ncl0180
Position (nt)

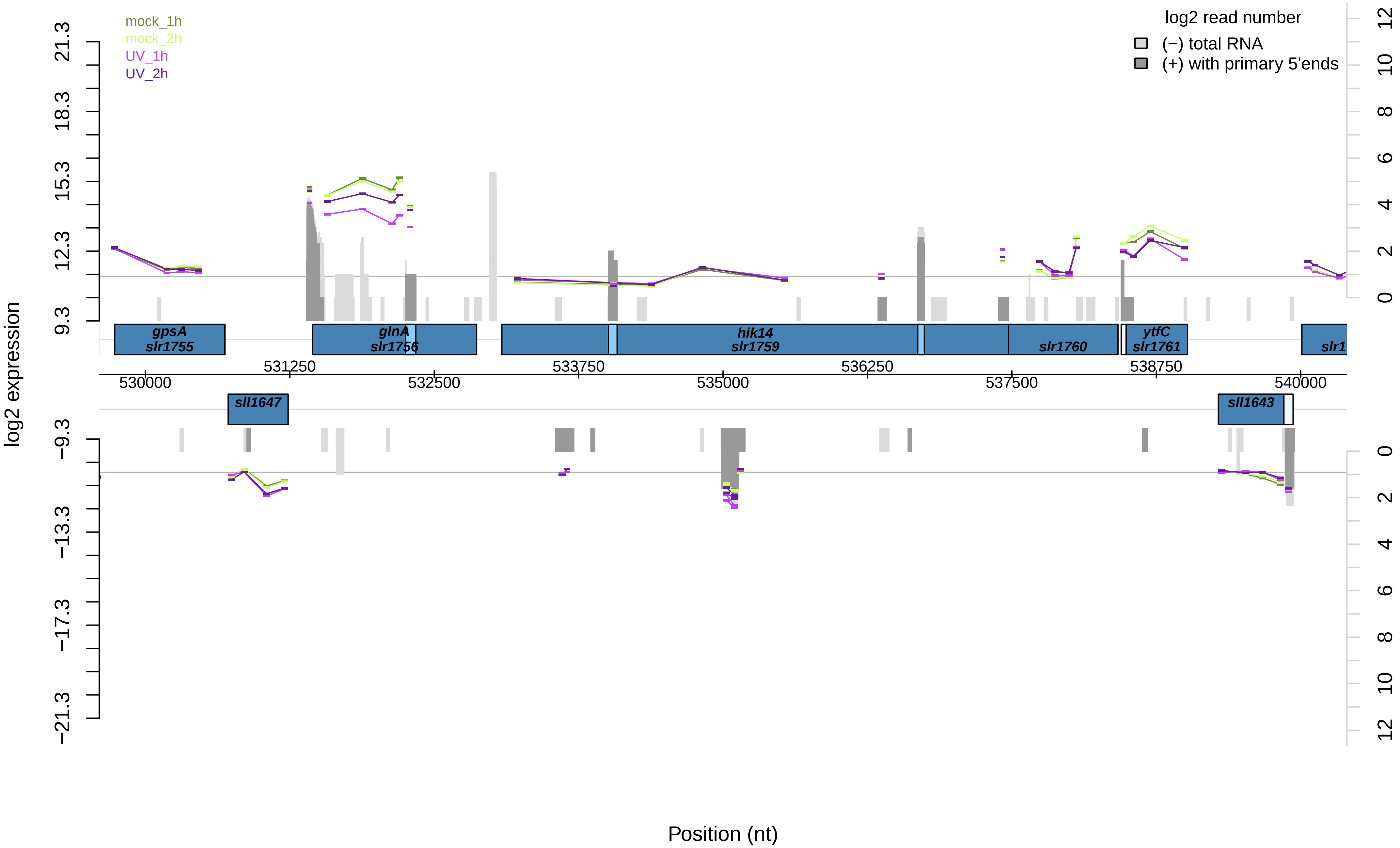
mock_1h
mock_2h
UV_1h
UV_2h
log2 read number
(−) total RNA
(+) with primary 5'ends
log2 expression
gpsA
slr1755
glnA
slr1756
hik14
slr1759
slr1760
ytfC
slr1761
slr1
sll1647
sll1643
530000
531250
532500
533750
535000
536250
537500
538750
540000
Position (nt)

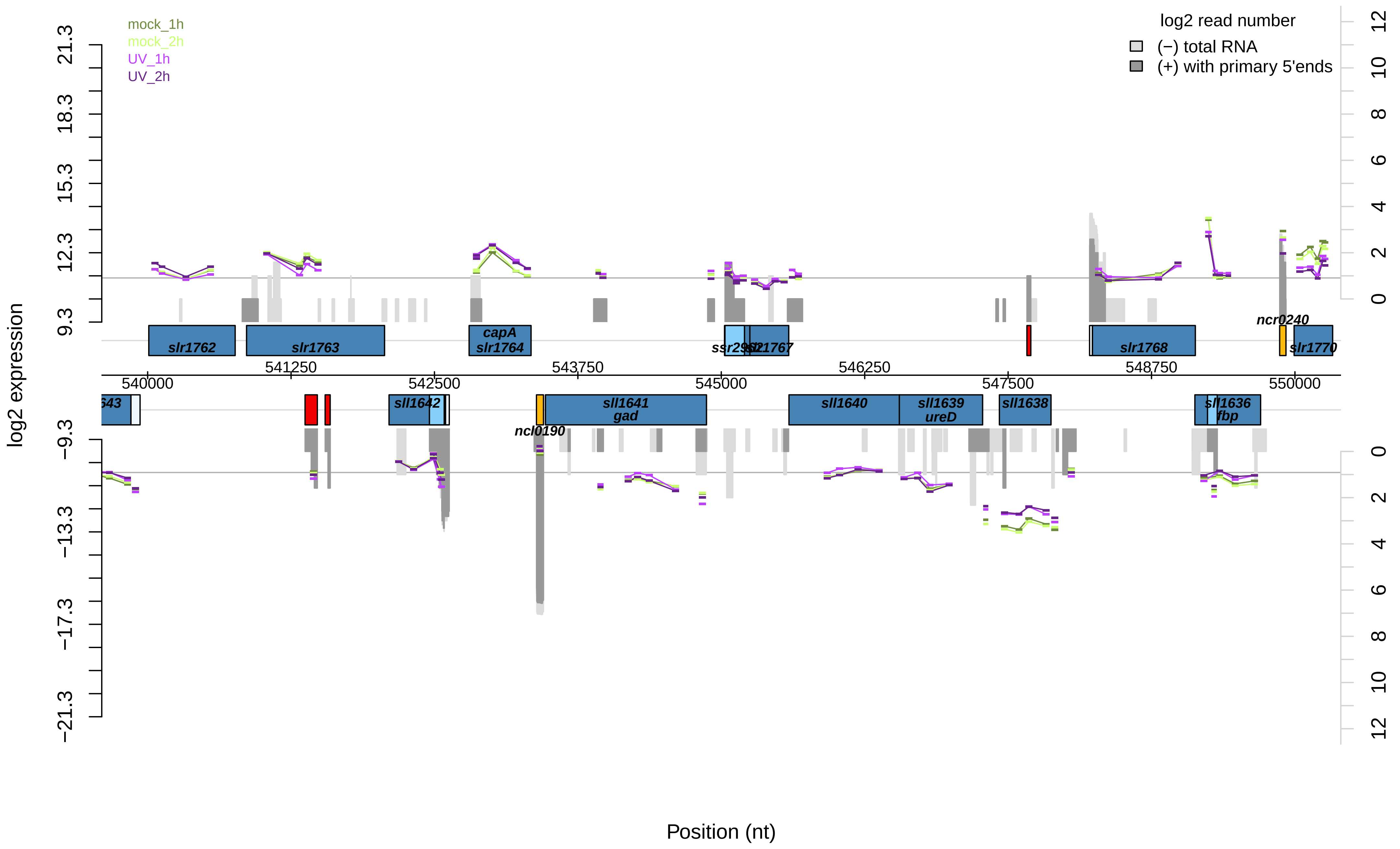

mock_1h
mock_2h
UV_1h
UV_2h
log2 read number
(−) total RNA
(+) with primary 5'ends
log2 expression
Position (nt)
slr1762
slr1763
capA
slr1764
ssr2988
slr1767
slr1768
ncr0240
slr1770
sll1642
ncl0190
sll1641
gad
sll1640
sll1639
ureD
sll1638
sll1636
fbp

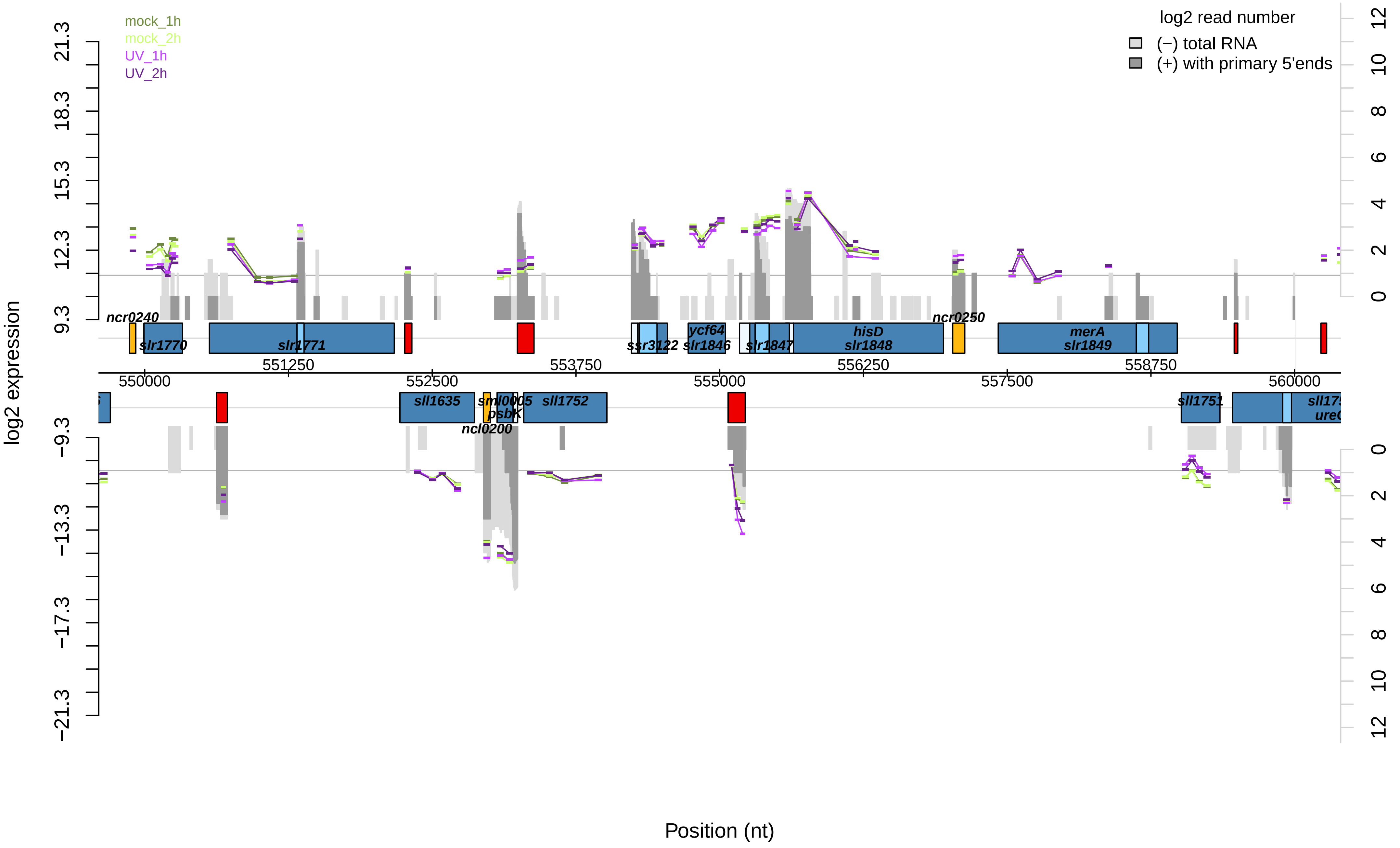

mock_1h
mock_2h
UV_1h
UV_2h
log2 read number
(−) total RNA
(+) with primary 5'ends
log2 expression
21.3
18.3
15.3
12.3
9.3
−9.3
−13.3
−17.3
−21.3
ncr0240
slr1770
slr1771
ssr3122
ycf64
slr1846
slr1847
hisD
slr1848
ncr0250
merA
slr1849
551250
553750
556250
558750
550000
552500
555000
557500
560000
sll1635
sml0005
psbK
ncl0200
sll1752
sll1751
Position (nt)

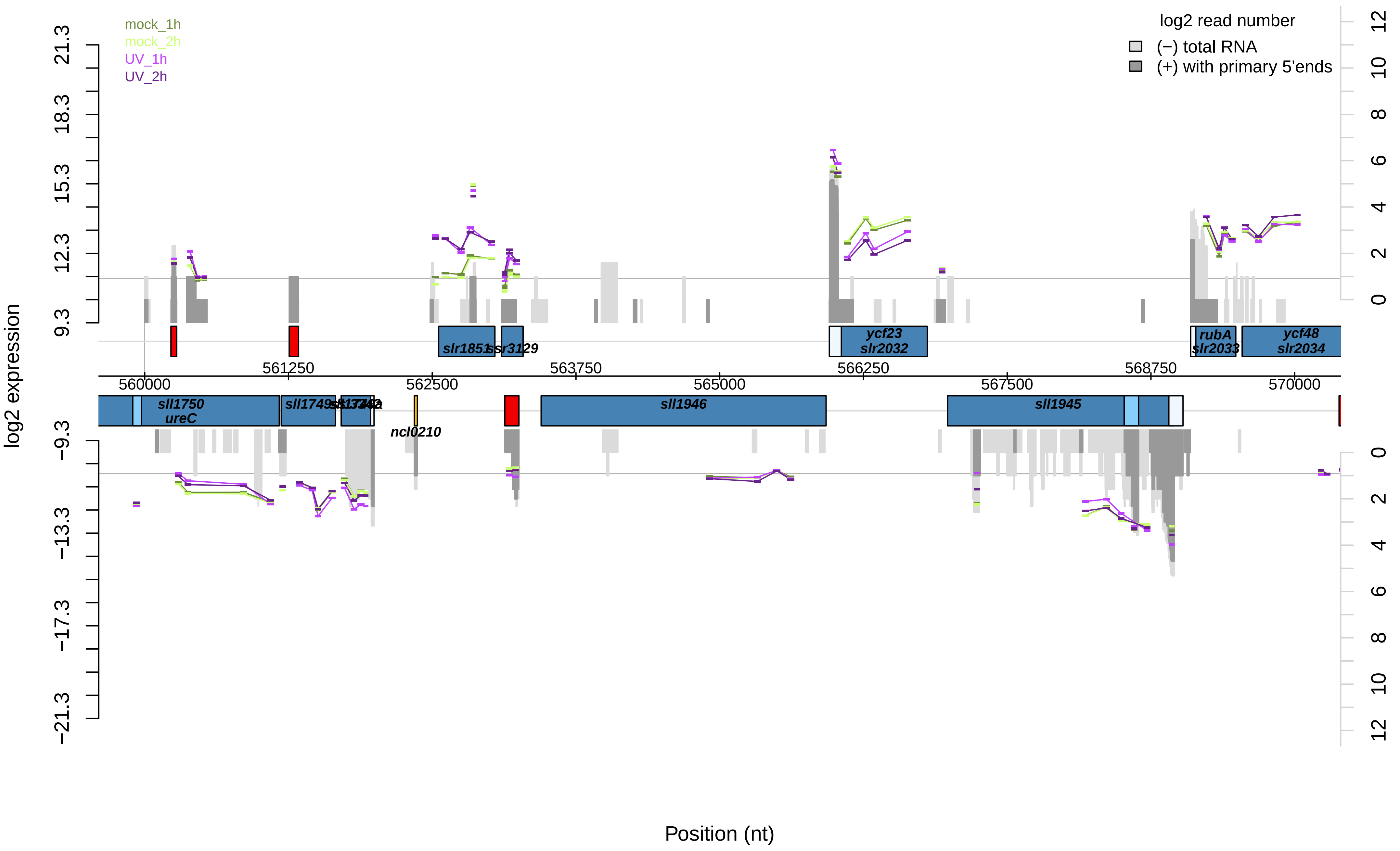

mock_1h
mock_2h
UV_1h
UV_2h
log2 read number
(−) total RNA
(+) with primary 5'ends
log2 expression
21.3
18.3
15.3
12.3
9.3
−9.3
−13.3
−17.3
−21.3
12
10
8
6
4
2
0
ycf23
slr2032
rubA
slr2033
ycf48
slr2034
slr1851
ssr3129
561250
563750
566250
568750
560000
562500
565000
567500
570000
sll1750
ureC
sll1946
sll1945
ncl0210
Position (nt)

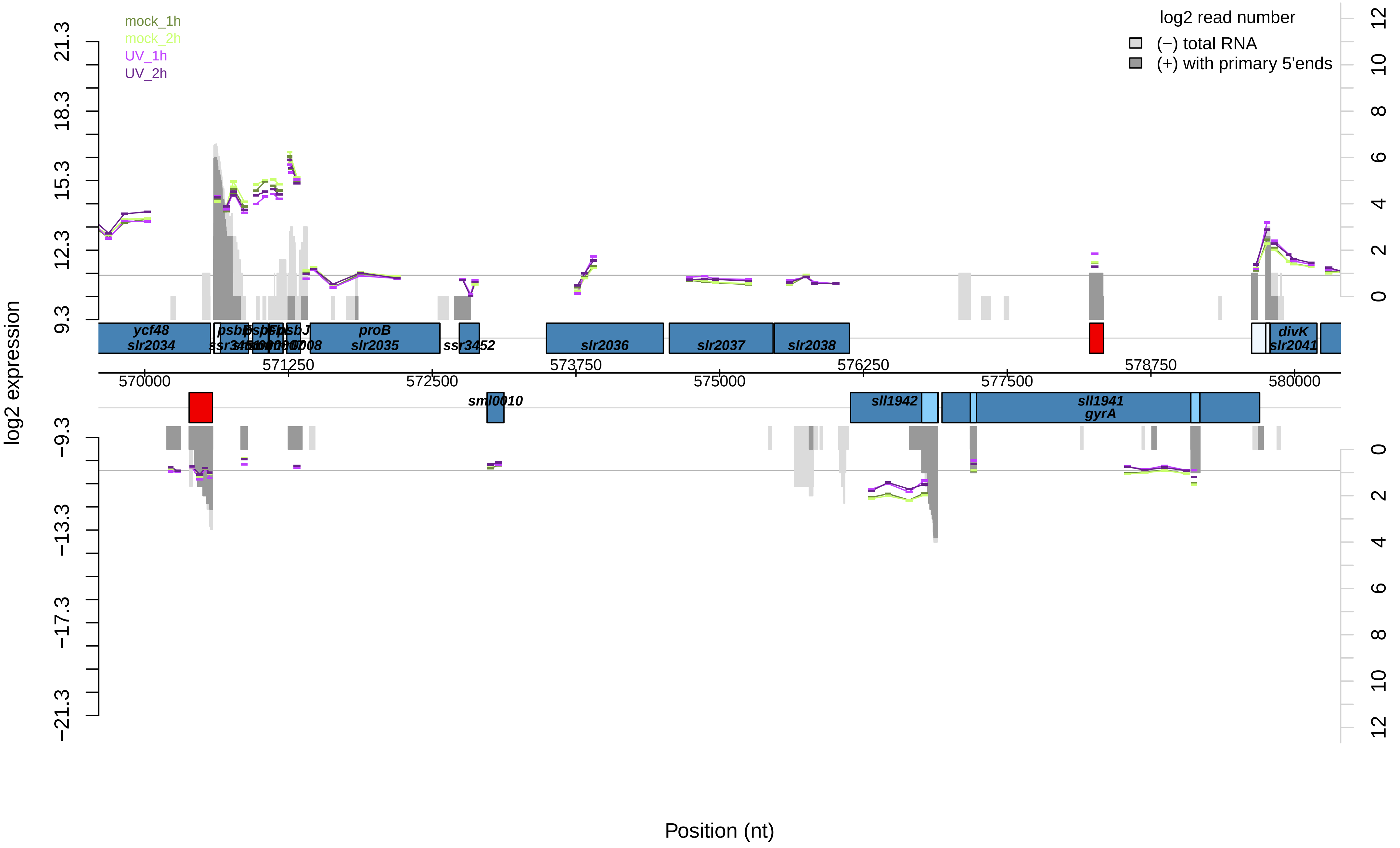

mock_1h
mock_2h
UV_1h
UV_2h
log2 read number
(−) total RNA
(+) with primary 5'ends
log2 expression
21.3
18.3
15.3
12.3
9.3
−9.3
−13.3
−17.3
−21.3
12
10
8
6
4
2
0
ycf48
slr2034
proB
slr2035
ssr3452
slr2036
slr2037
slr2038
divK
slr2041
sml0010
sll1942
sll1941
gyrA
570000
571250
572500
573750
575000
576250
577500
578750
580000
Position (nt)

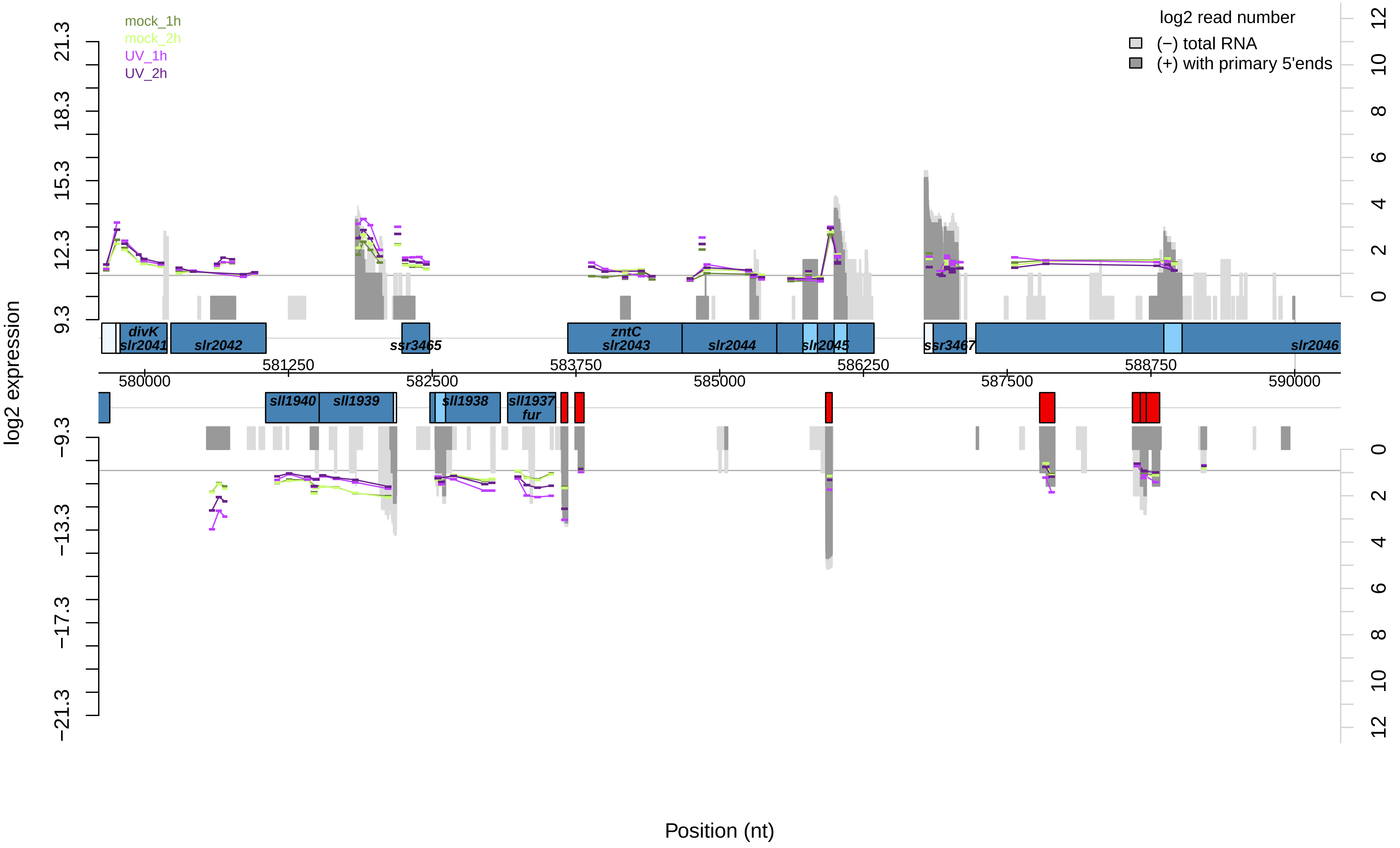

mock_1h
mock_2h
UV_1h
UV_2h
log2 read number
(−) total RNA
(+) with primary 5'ends
log2 expression
21.3
18.3
15.3
12.3
9.3
−9.3
−13.3
−17.3
−21.3
12
10
8
6
4
2
0
divK
slr2041
slr2042
ssr3465
zntC
slr2043
slr2044
slr2045
ssr3467
slr2046
581250
583750
586250
588750
580000
582500
585000
587500
590000
sll1940
sll1939
sll1938
sll1937
fur
Position (nt)

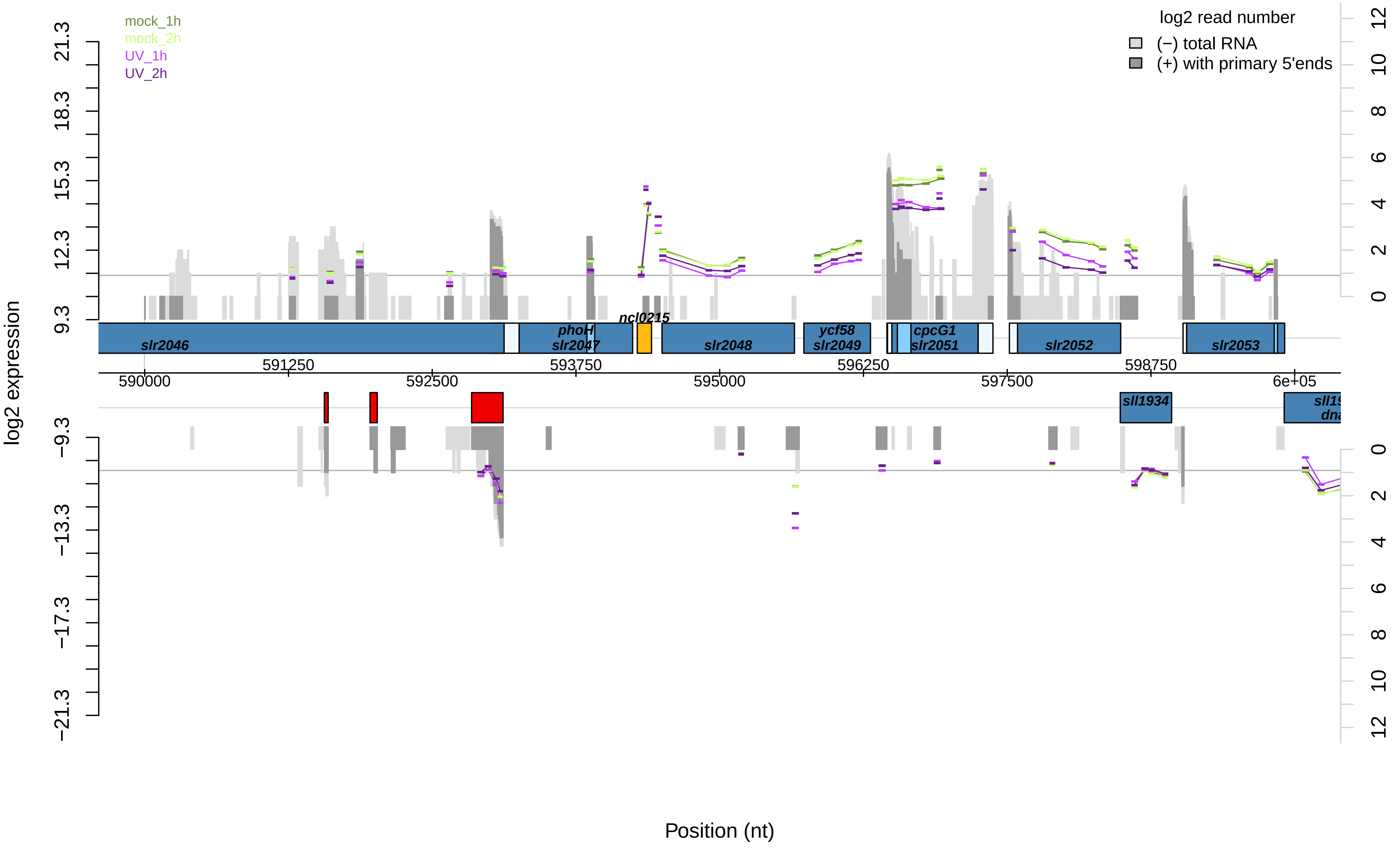

mock_1h
mock_2h
UV_1h
UV_2h
log2 read number
(−) total RNA
(+) with primary 5'ends
log2 expression
slr2046
phoH
slr2047
ncl0215
slr2048
ycf58
slr2049
cpcG1
slr2051
slr2052
slr2053
sll1934
590000
591250
592500
593750
595000
596250
597500
598750
6e+05
Position (nt)

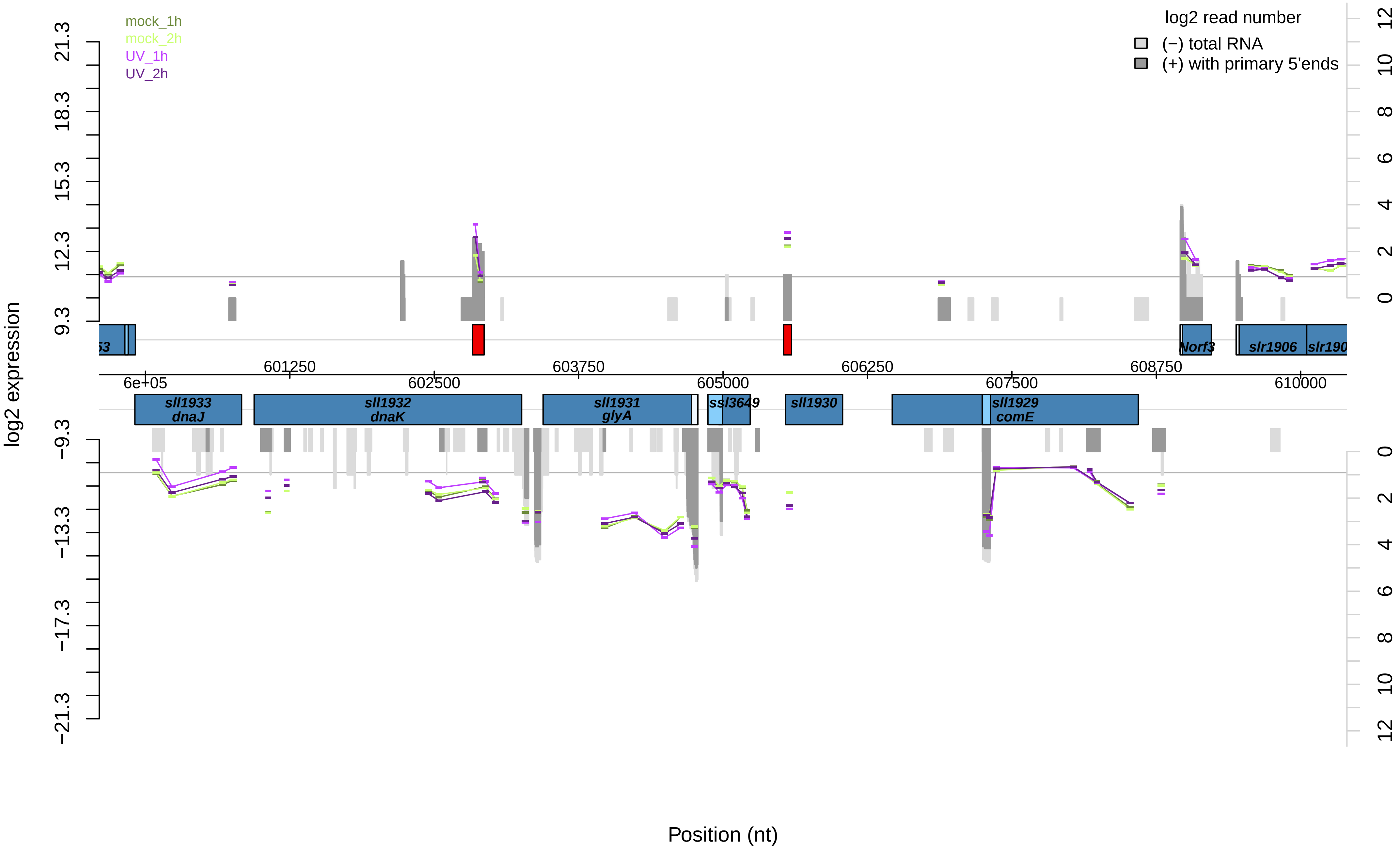

mock_1h
mock_2h
UV_1h
UV_2h
log2 read number
(−) total RNA
(+) with primary 5'ends
log2 expression
21.3
18.3
15.3
12.3
9.3
−9.3
−13.3
−17.3
−21.3
12
10
8
6
4
2
0
601250
603750
606250
608750
6e+05
602500
605000
607500
610000
sll1933
dnaJ
sll1932
dnaK
sll1931
glyA
ssl3649
sll1930
sll1929
comE
Norf3
slr1906
Position (nt)

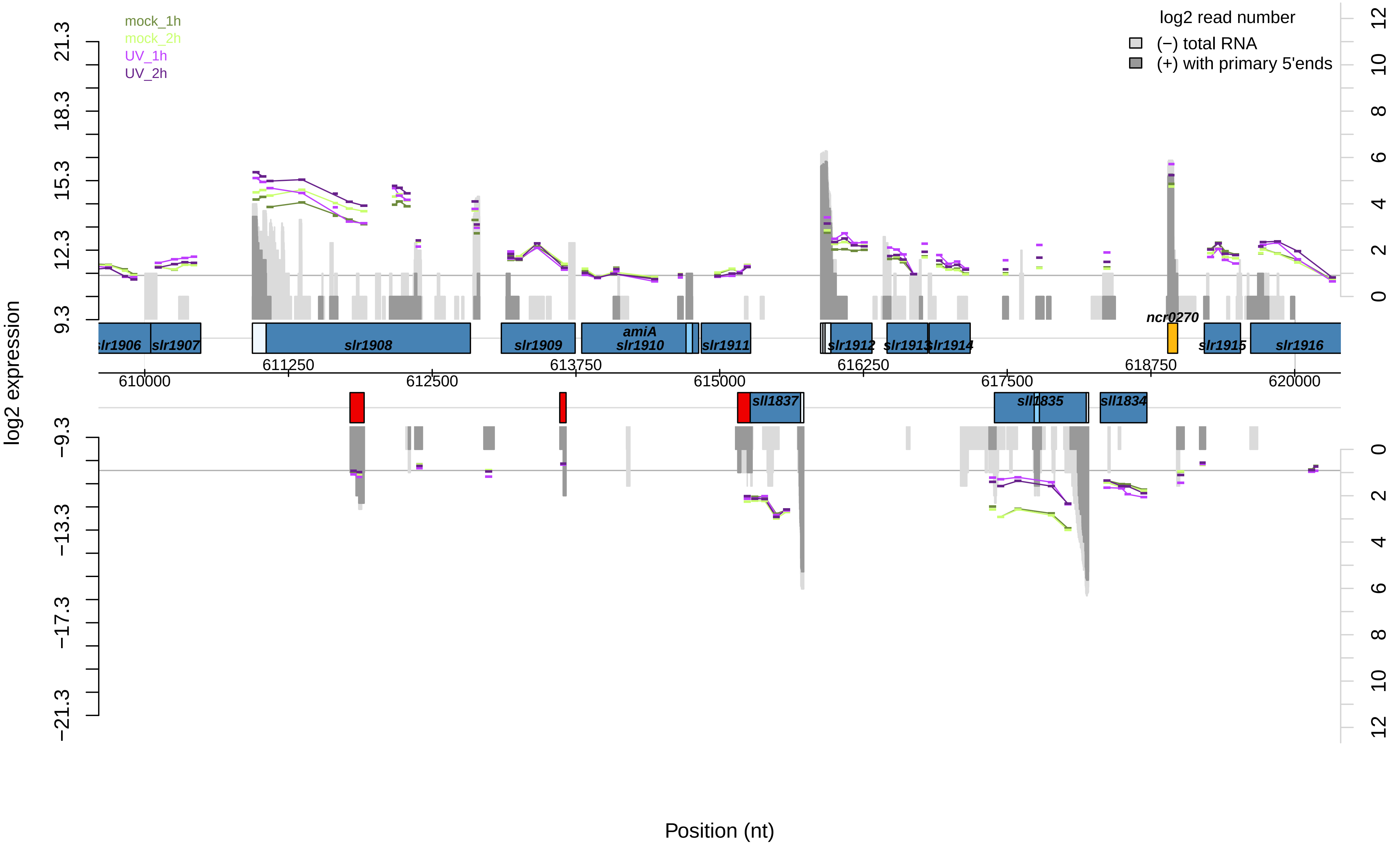

mock_1h
mock_2h
UV_1h
UV_2h
log2 read number
(−) total RNA
(+) with primary 5'ends
log2 expression
21.3
18.3
15.3
12.3
9.3
−9.3
−13.3
−17.3
−21.3
12
10
8
6
4
2
0
slr1906
slr1907
slr1908
slr1909
amiA
slr1910
slr1911
slr1912
slr1913
slr1914
ncr0270
slr1915
slr1916
sll1837
sll1835
sll1834
611250
613750
616250
618750
610000
612500
615000
617500
620000
Position (nt)

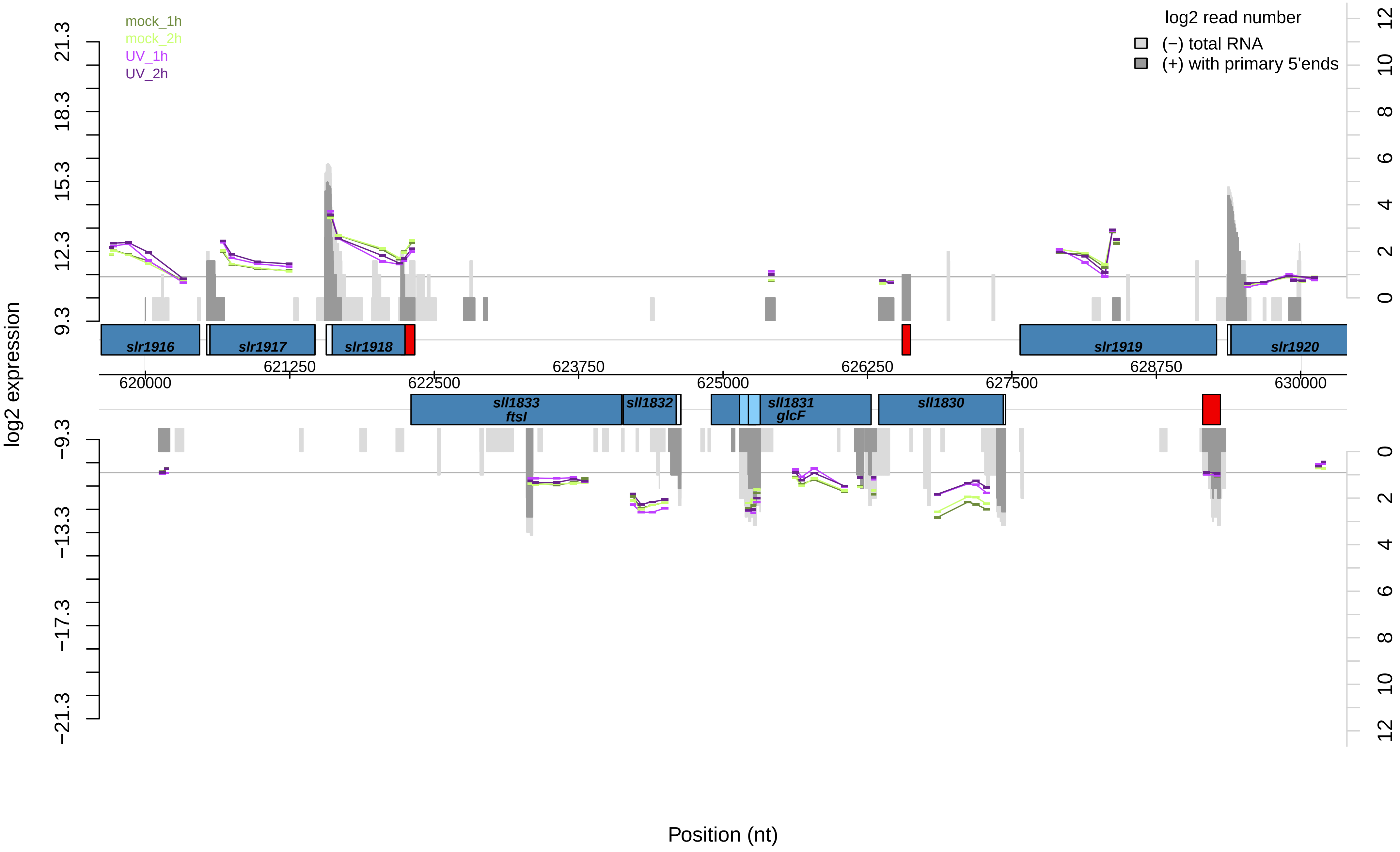
mock_1h
mock_2h
UV_1h
UV_2h
log2 read number
(−) total RNA
(+) with primary 5'ends
log2 expression
21.3
18.3
15.3
12.3
9.3
−9.3
−13.3
−17.3
−21.3
12
10
8
6
4
2
0
slr1916
slr1917
slr1918
slr1919
slr1920
sll1833
ftsI
sll1832
sll1831
glcF
sll1830
620000
621250
622500
623750
625000
626250
627500
628750
630000
Position (nt)

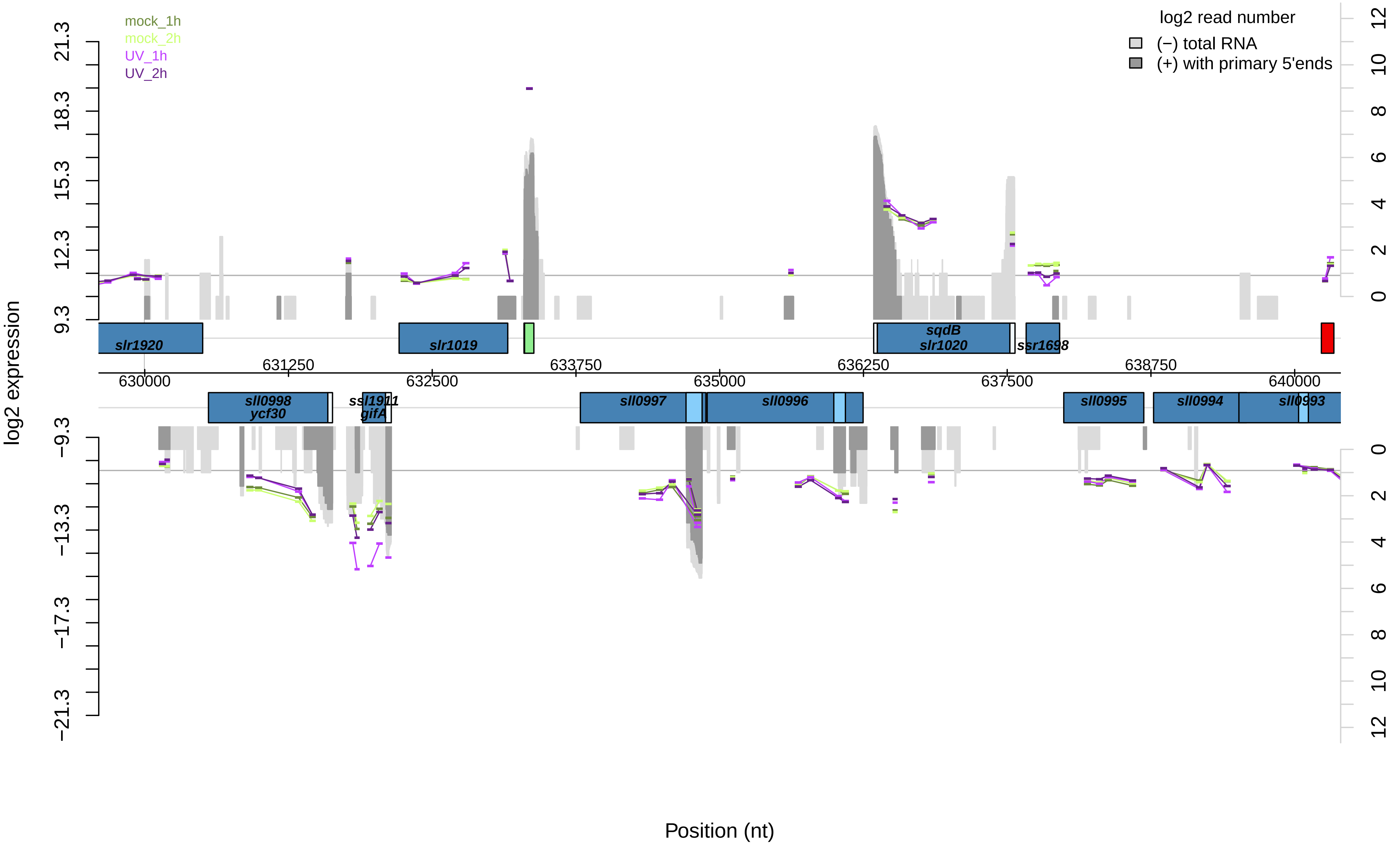

mock_1h
mock_2h
UV_1h
UV_2h
log2 read number
(−) total RNA
(+) with primary 5'ends
log2 expression
21.3
18.3
15.3
12.3
9.3
−9.3
−13.3
−17.3
−21.3
12
10
8
6
4
2
0
slr1920
slr1019
sqdB
slr1020
ssr1698
631250
633750
636250
638750
630000
632500
635000
637500
640000
sll0998
ycf30
ssl1911
gifA
sll0997
sll0996
sll0995
sll0994
sll0993
Position (nt)

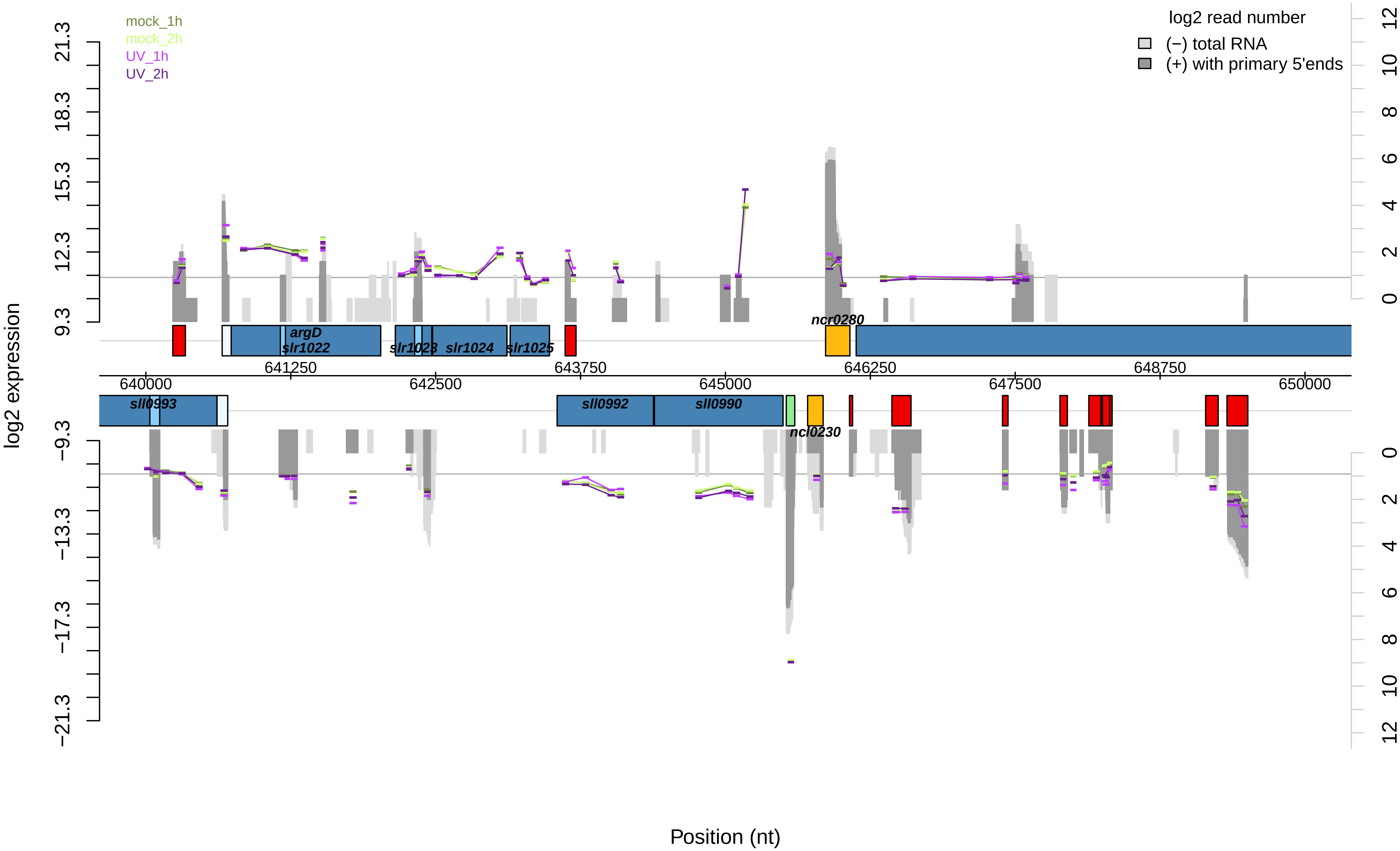

mock_1h
mock_2h
UV_1h
UV_2h
log2 read number
(−) total RNA
(+) with primary 5'ends
21.3
18.3
15.3
12.3
9.3
−9.3
−13.3
−17.3
−21.3
log2 expression
12
10
8
6
4
2
0
argD
slr1022
slr1023
slr1024
slr1025
ncr0280
sll0993
sll0992
sll0990
ncl0230
641250
643750
646250
648750
640000
642500
645000
647500
650000
Position (nt)

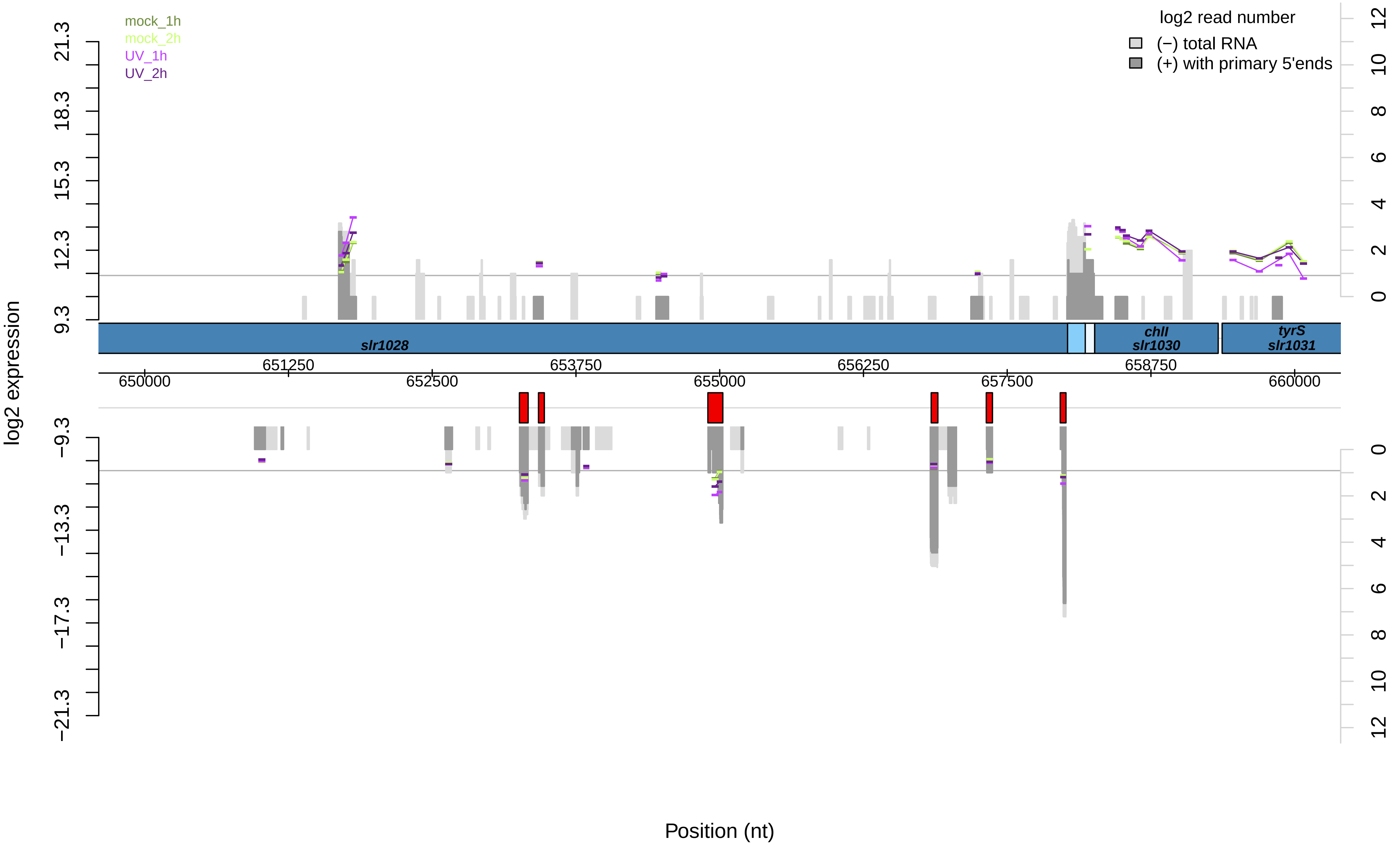
mock_1h
mock_2h
UV_1h
UV_2h
log2 read number
(−) total RNA
(+) with primary 5'ends
log2 expression
21.3
18.3
15.3
12.3
9.3
−9.3
−13.3
−17.3
−21.3
12
10
8
6
4
2
0
slr1028
chlI
slr1030
tyrS
slr1031
651250
653750
656250
658750
650000
652500
655000
657500
660000
Position (nt)

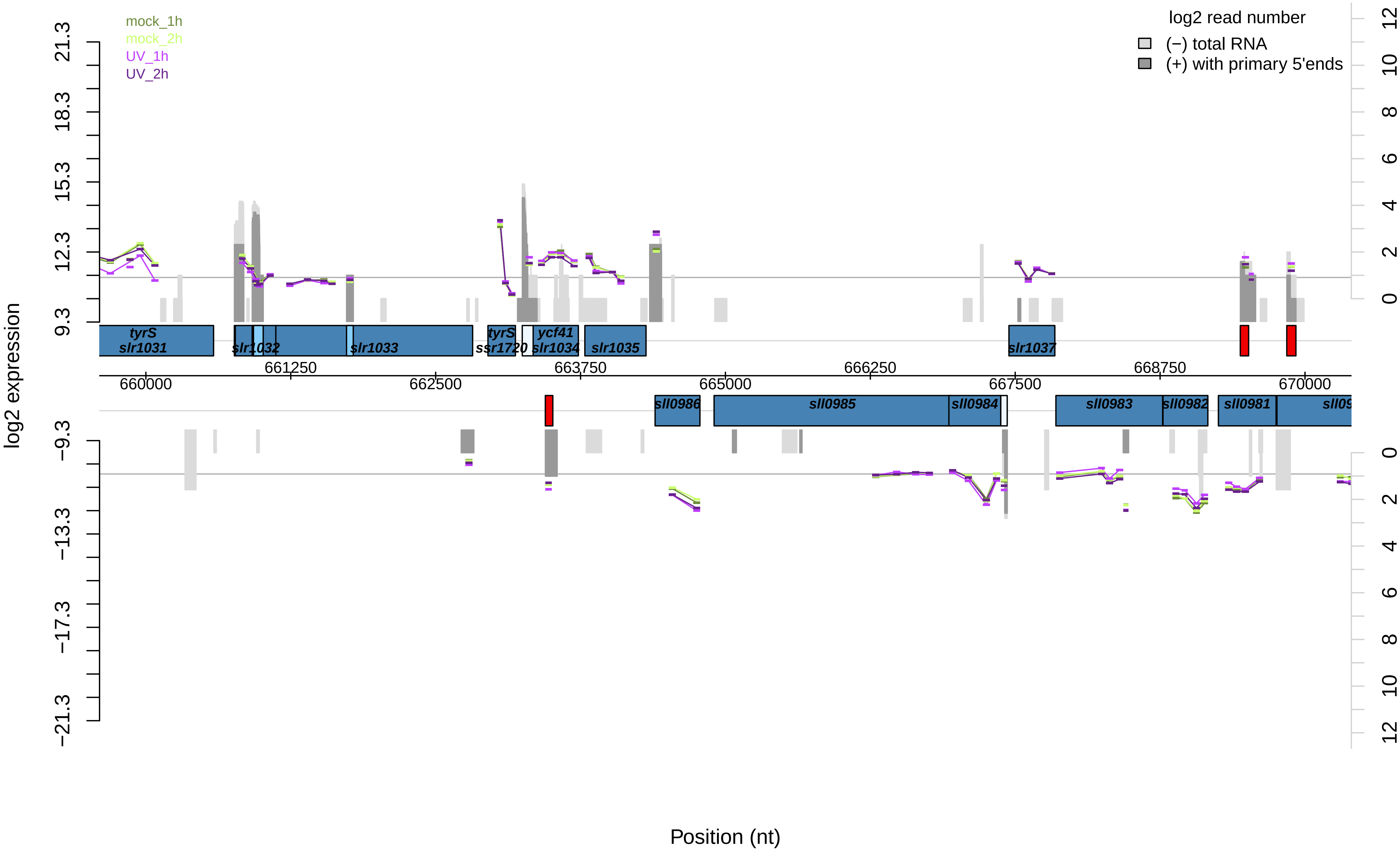

mock_1h
mock_2h
UV_1h
UV_2h
log2 read number
(−) total RNA
(+) with primary 5'ends
log2 expression
Position (nt)
tyrS
slr1031
slr1032
slr1033
tyrS
ssr1720
ycf41
slr1034
slr1035
slr1037
sll0986
sll0985
sll0984
sll0983
sll0982
sll0981
661250
663750
666250
668750
660000
662500
665000
667500
670000
21.3
18.3
15.3
12.3
9.3
−9.3
−13.3
−17.3
−21.3

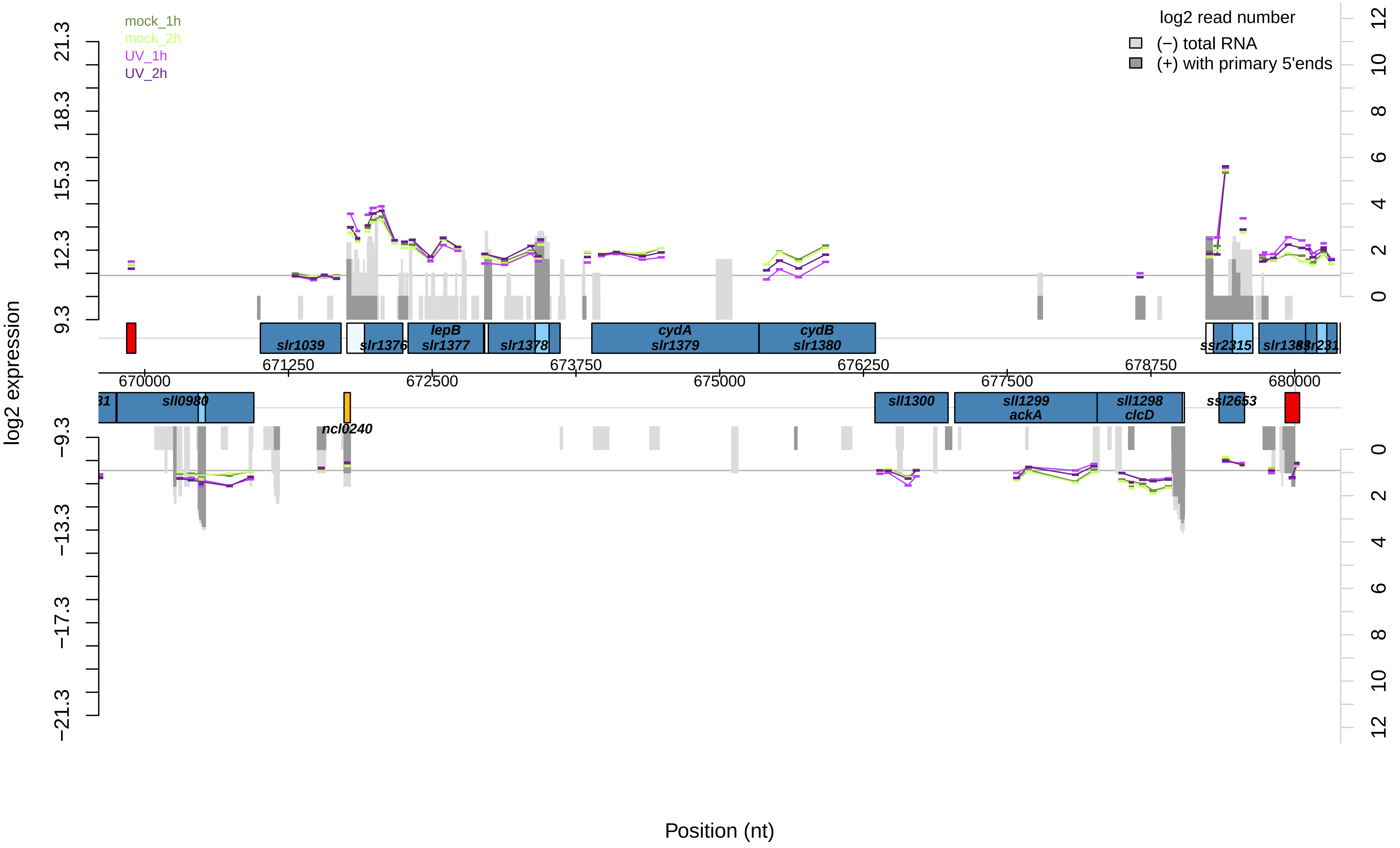

mock_1h
mock_2h
UV_1h
UV_2h
log2 read number
(−) total RNA
(+) with primary 5'ends
log2 expression
21.3
18.3
15.3
12.3
9.3
−9.3
−13.3
−17.3
−21.3
12
10
8
6
4
2
0
slr1039
slr1376
lepB
slr1377
slr1378
cydA
slr1379
cydB
slr1380
ssr2315
slr1383
ssr231
671250
673750
676250
678750
670000
672500
675000
677500
680000
sll0980
ncl0240
sll1300
sll1299
ackA
sll1298
clcD
ssl2653
Position (nt)

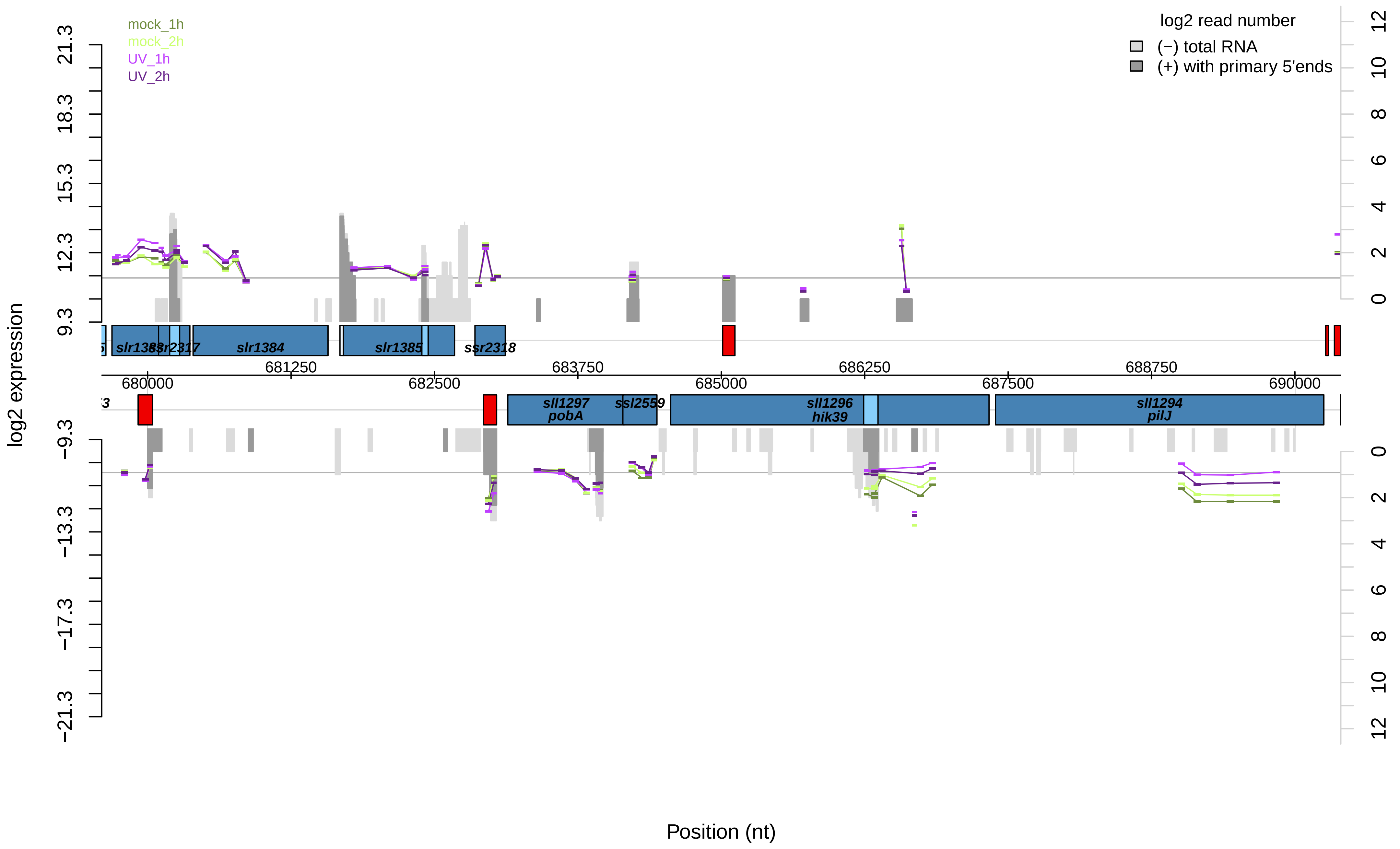

mock_1h
mock_2h
UV_1h
UV_2h
log2 read number
(−) total RNA
(+) with primary 5'ends
log2 expression
21.3
18.3
15.3
12.3
9.3
−9.3
−13.3
−17.3
−21.3
12
10
8
6
4
2
0
slr1383
ssr2317
slr1384
slr1385
ssr2318
681250
683750
686250
688750
680000
682500
685000
687500
690000
sll1297
pobA
ssl2559
sll1296
hik39
sll1294
pilJ
Position (nt)

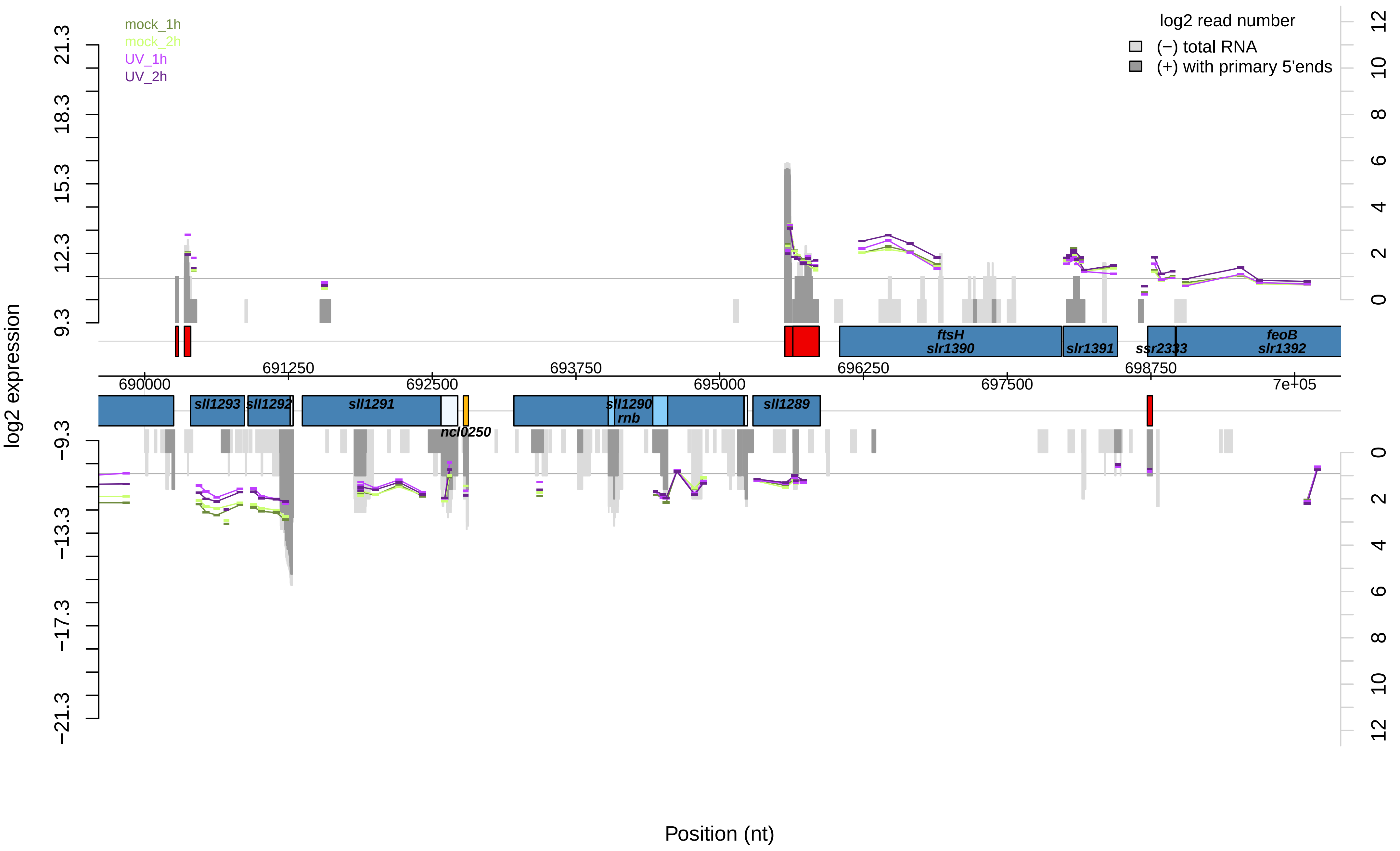

mock_1h
mock_2h
UV_1h
UV_2h
log2 read number
(−) total RNA
(+) with primary 5'ends
log2 expression
21.3
18.3
15.3
12.3
9.3
−9.3
−13.3
−17.3
−21.3
12
10
8
6
4
2
0
ftsH
slr1390
slr1391
ssr2333
feoB
slr1392
691250
693750
696250
698750
690000
692500
695000
697500
7e+05
sll1293
sll1292
sll1291
ncl0250
sll1290
rnb
sll1289
Position (nt)

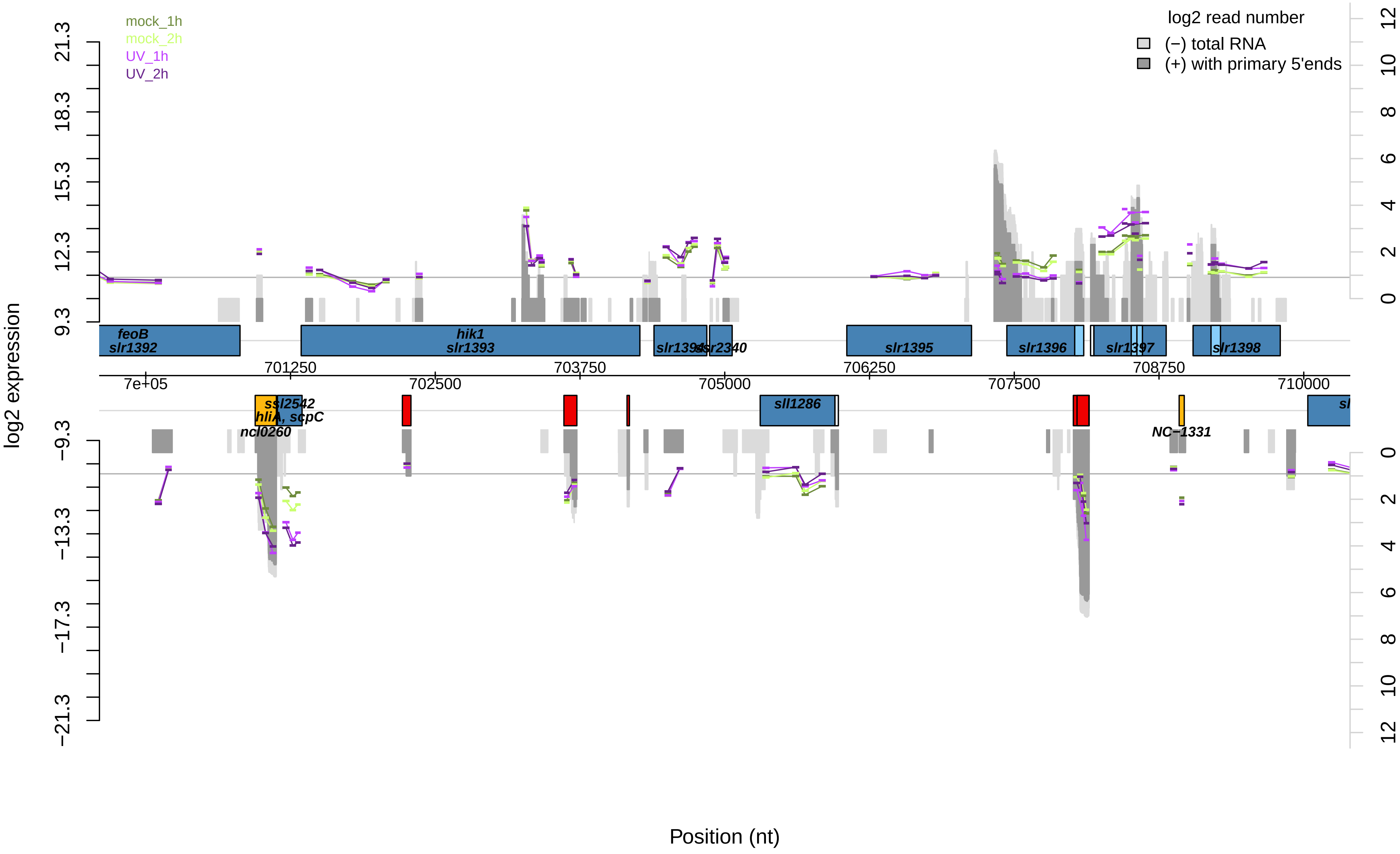
mock_1h
mock_2h
UV_1h
UV_2h
log2 read number
(−) total RNA
(+) with primary 5'ends
log2 expression
21.3
18.3
15.3
12.3
9.3
−9.3
−13.3
−17.3
−21.3
12
10
8
6
4
2
0
feoB
slr1392
hik1
slr1393
slr1394
ssr2340
slr1395
slr1396
slr1397
slr1398
701250
703750
706250
708750
7e+05
702500
705000
707500
710000
ssl2542
hliA, scpC
ncl0260
sll1286
NC−1331
Position (nt)

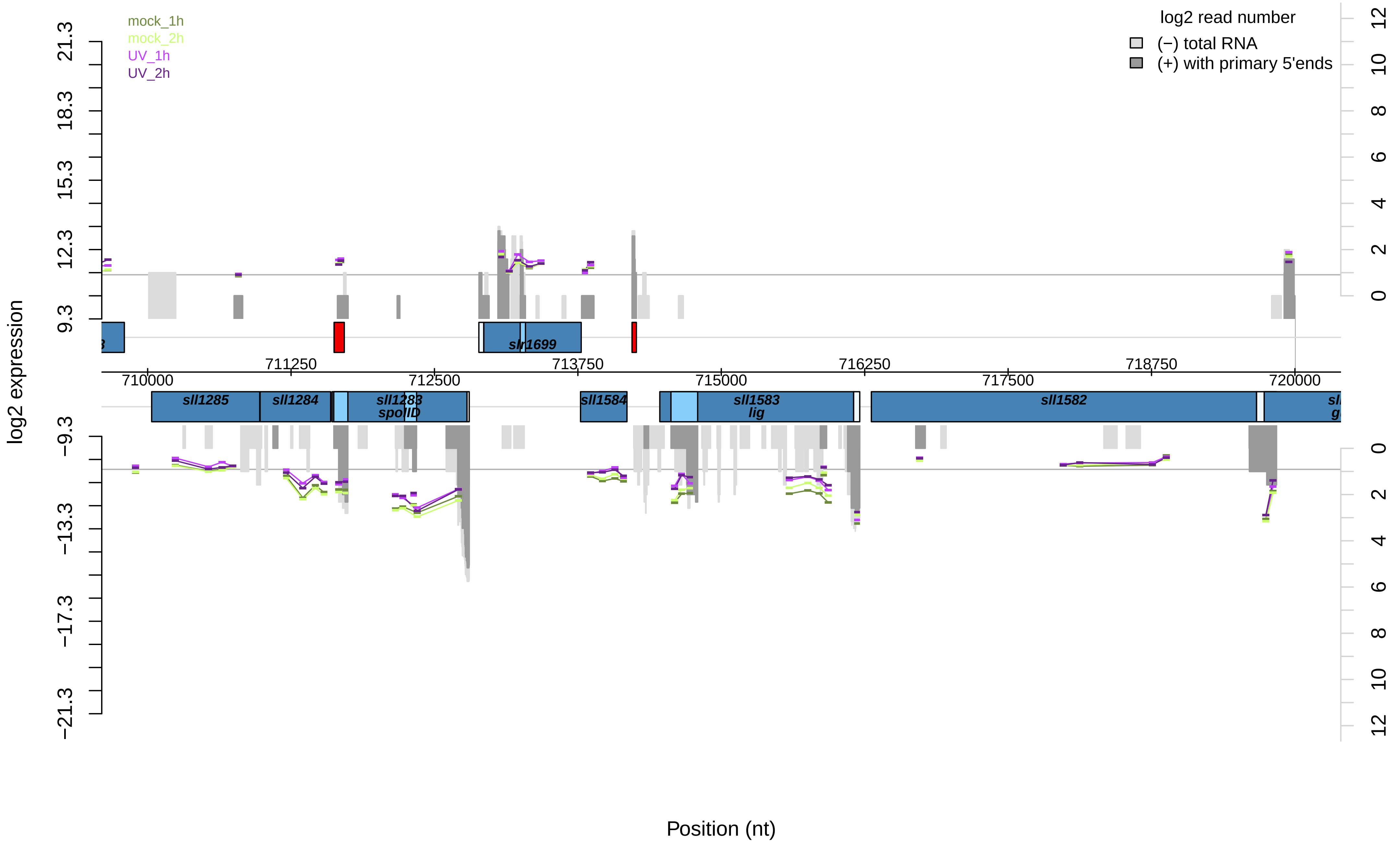

mock_1h
mock_2h
UV_1h
UV_2h
log2 read number
(−) total RNA
(+) with primary 5'ends
log2 expression
21.3
18.3
15.3
12.3
9.3
−9.3
−13.3
−17.3
−21.3
12
10
8
6
4
2
0
sll1285
sll1284
sll1283
spoIID
slr1699
sll1584
sll1583
lig
sll1582
710000
711250
712500
713750
715000
716250
717500
718750
720000
Position (nt)

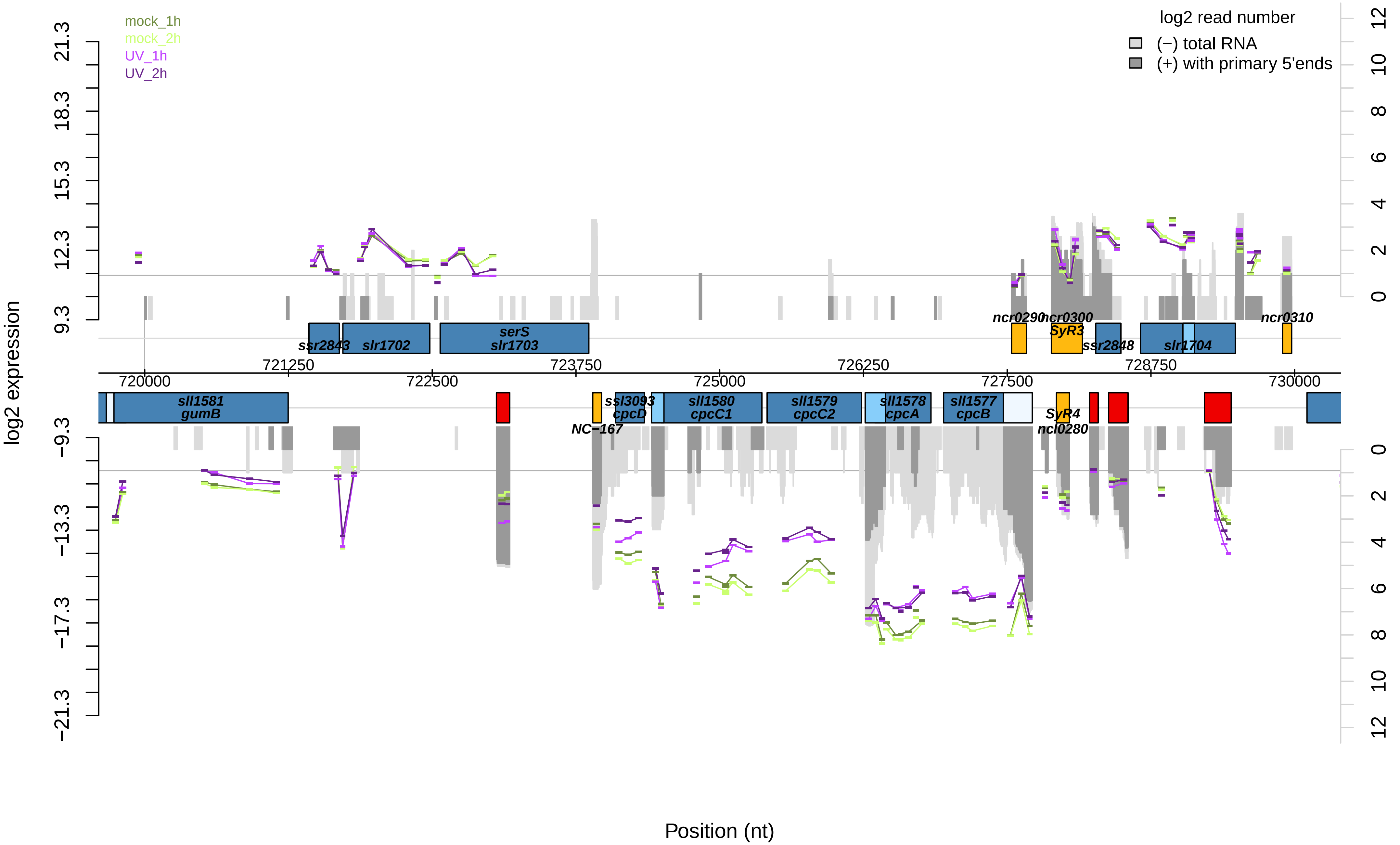

mock_1h
mock_2h
UV_1h
UV_2h
log2 read number
(−) total RNA
(+) with primary 5'ends
log2 expression
21.3
18.3
15.3
12.3
9.3
−9.3
−13.3
−17.3
−21.3
ssr2843
slr1702
serS
slr1703
ncr0290
ncr0300
SyR3
ssr2848
slr1704
ncr0310
721250
723750
726250
728750
720000
722500
725000
727500
730000
sll1581
gumB
ssl3093
cpcD
NC−167
sll1580
cpcC1
sll1579
cpcC2
sll1578
cpcA
sll1577
cpcB
SyR4
ncl0280
Position (nt)

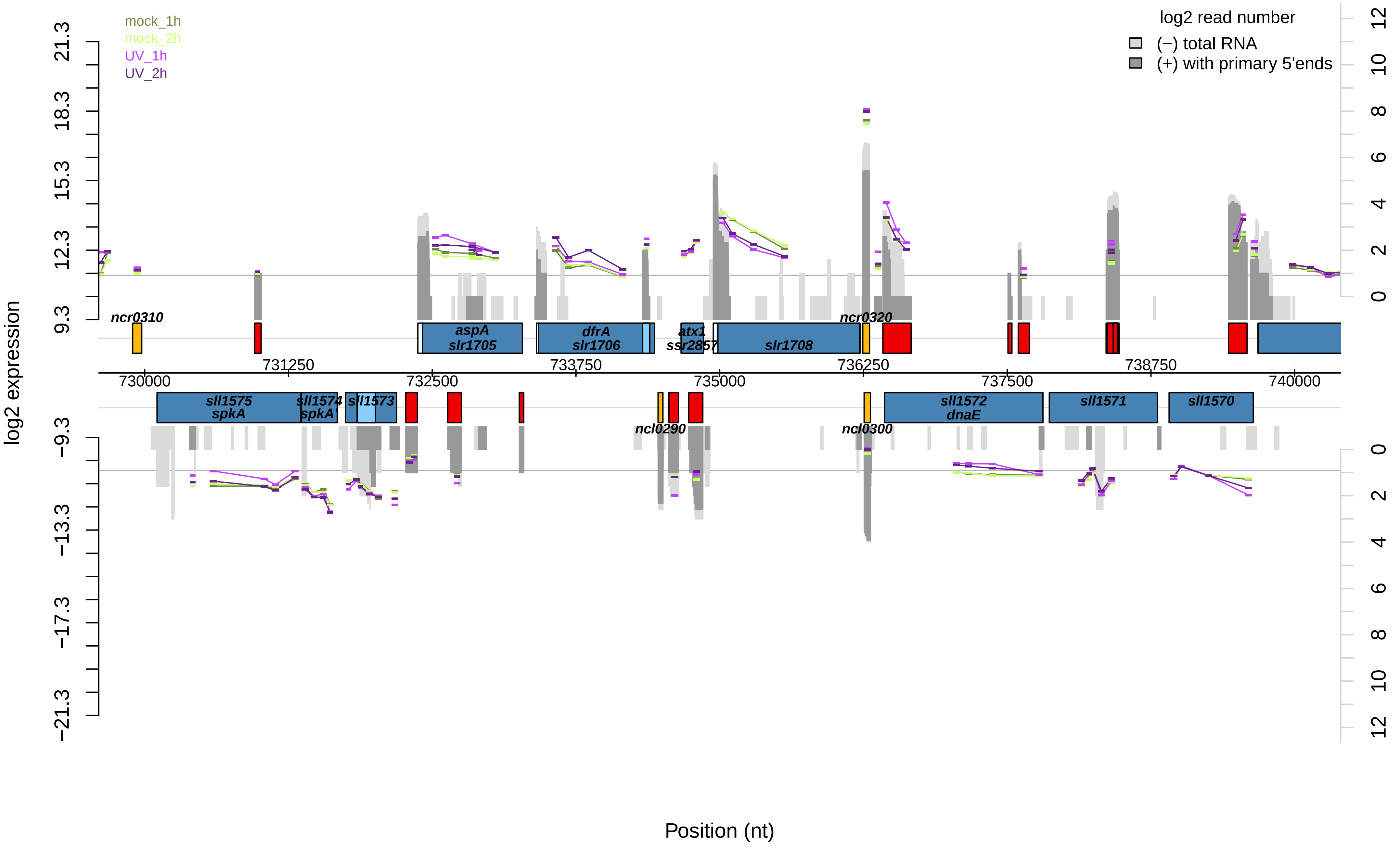

mock_1h
mock_2h
UV_1h
UV_2h
log2 read number
(−) total RNA
(+) with primary 5'ends
log2 expression
21.3
18.3
15.3
12.3
9.3
−9.3
−13.3
−17.3
−21.3
12
10
8
6
4
2
0
ncr0310
aspA
slr1705
dfrA
slr1706
atx1
ssr2857
slr1708
ncr0320
731250
733750
736250
738750
730000
732500
735000
737500
740000
sll1575
spkA
sll1574
spkA
sll1573
sll1572
dnaE
sll1571
sll1570
ncl0290
ncl0300
Position (nt)

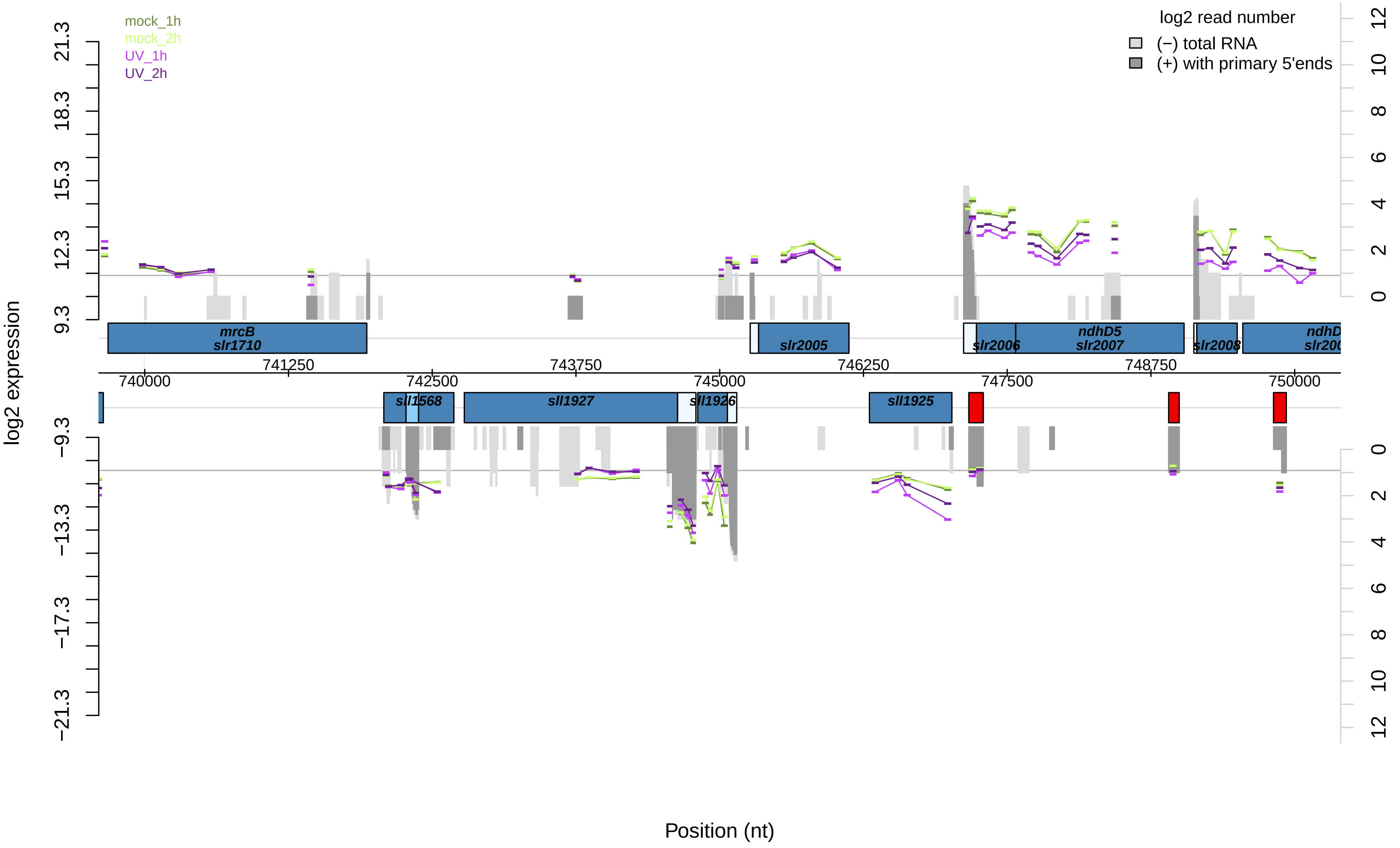

mock_1h
mock_2h
UV_1h
UV_2h
log2 read number
(−) total RNA
(+) with primary 5'ends
log2 expression
21.3
18.3
15.3
12.3
9.3
−9.3
−13.3
−17.3
−21.3
mrcB
slr1710
slr2005
slr2006
ndhD5
slr2007
slr2008
sll1568
sll1927
sll1926
sll1925
741250
743750
746250
748750
740000
742500
745000
747500
750000
Position (nt)

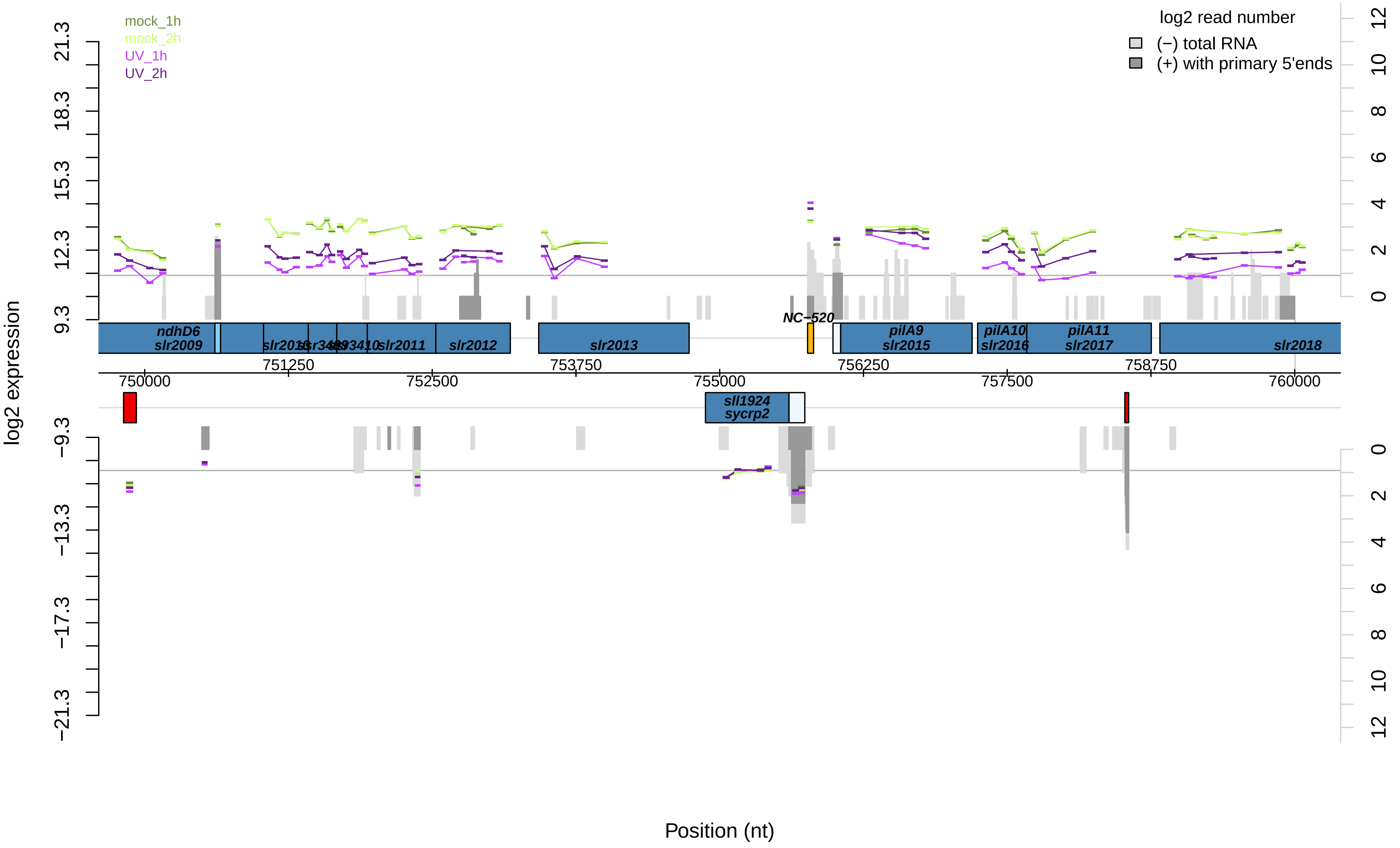
mock_1h
mock_2h
UV_1h
UV_2h
log2 read number
(−) total RNA
(+) with primary 5'ends
log2 expression
21.3
18.3
15.3
12.3
9.3
−9.3
−13.3
−17.3
−21.3
ndhD6
slr2009
slr2010
ssr3409
ssr3410
slr2011
slr2012
slr2013
NC−520
pilA9
slr2015
pilA10
slr2016
pilA11
slr2017
slr2018
751250
753750
756250
758750
750000
752500
755000
757500
760000
sll1924
sycrp2
Position (nt)

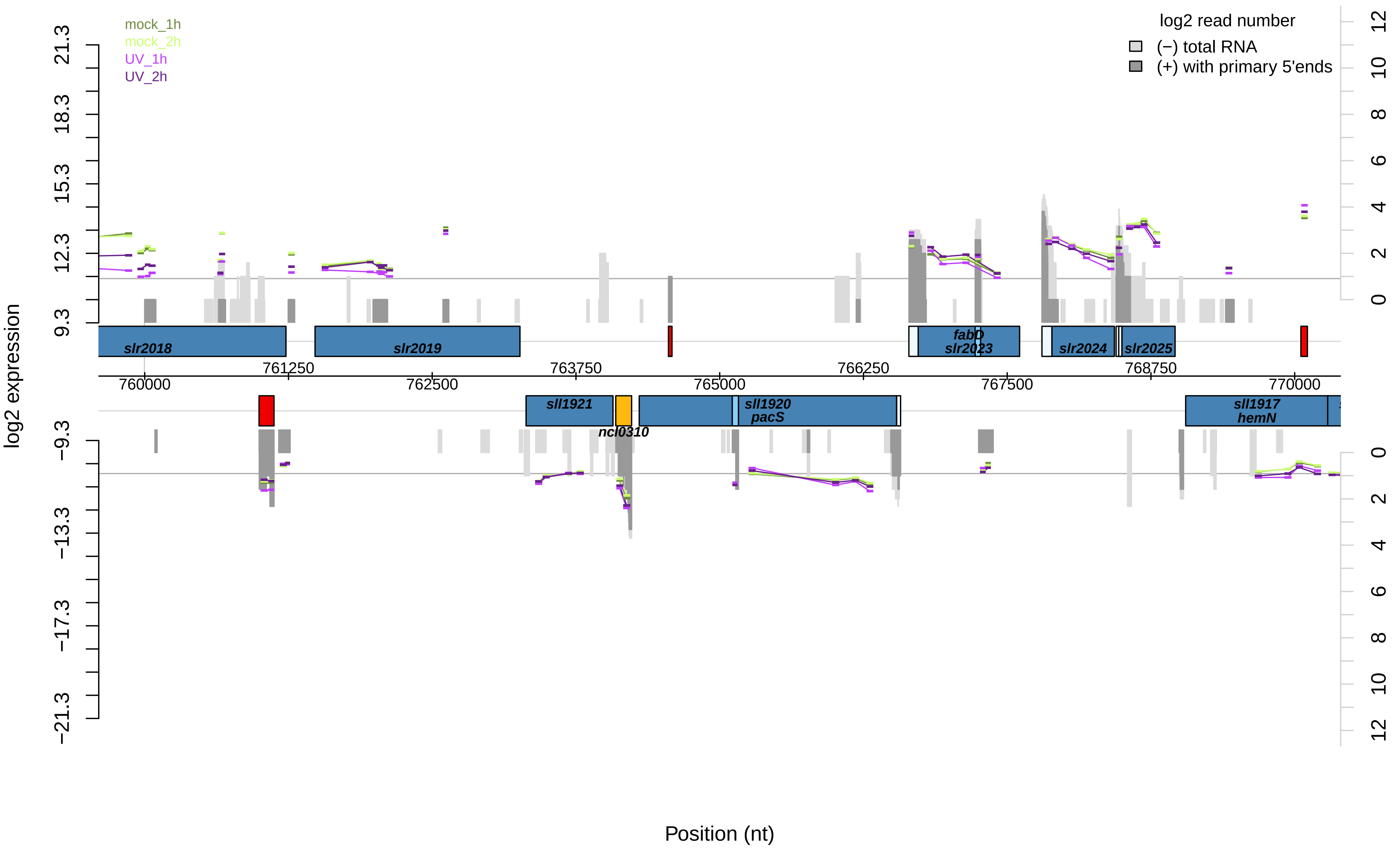

mock_1h
mock_2h
UV_1h
UV_2h
log2 read number
(−) total RNA
(+) with primary 5'ends
log2 expression
21.3
18.3
15.3
12.3
9.3
−9.3
−13.3
−17.3
−21.3
12
10
8
6
4
2
0
slr2018
slr2019
fabD
slr2023
slr2024
slr2025
761250
763750
766250
768750
760000
762500
765000
767500
770000
sll1921
ncl0310
sll1920
pacS
sll1917
hemN
Position (nt)

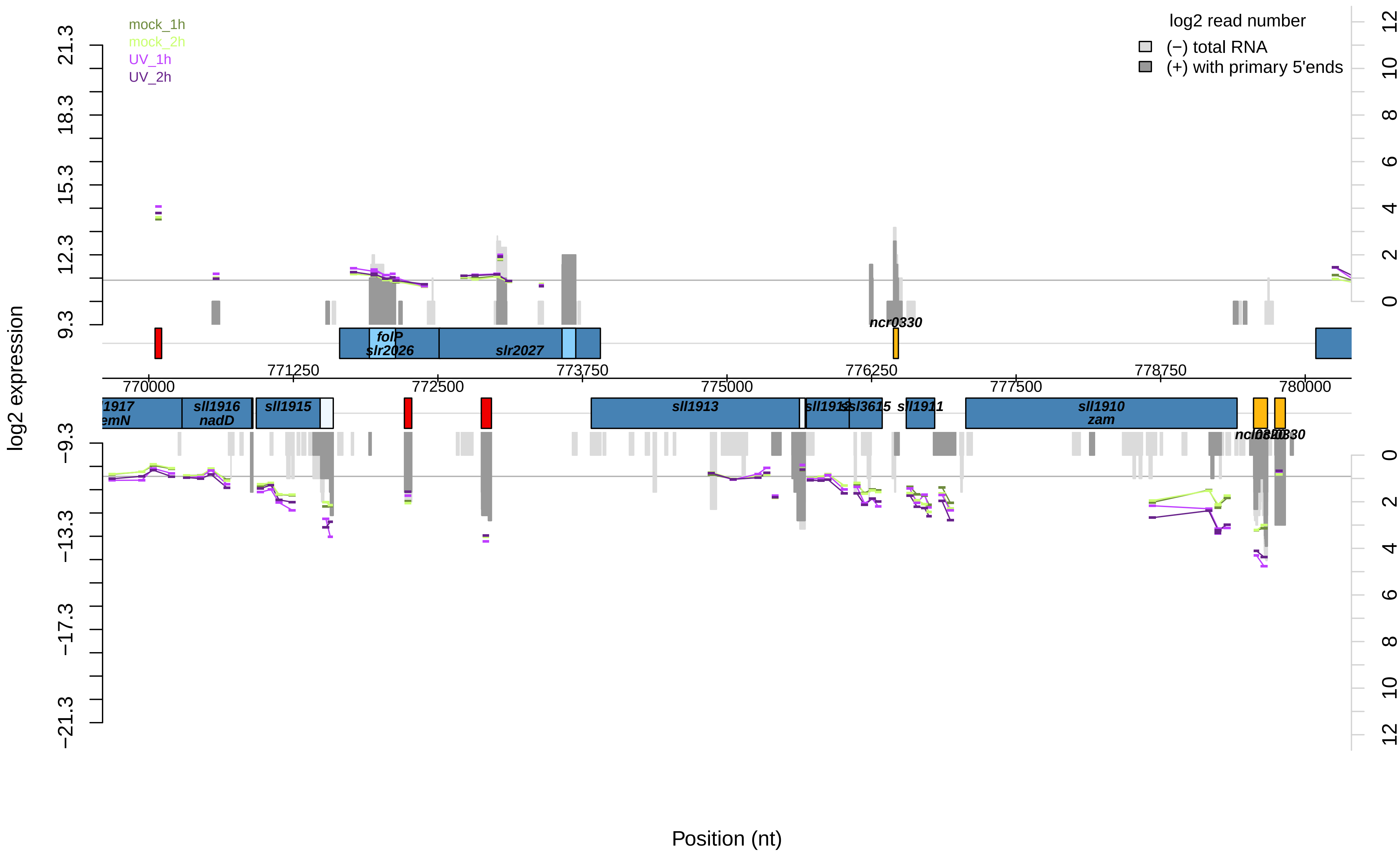

mock_1h
mock_2h
UV_1h
UV_2h
log2 read number
(−) total RNA
(+) with primary 5'ends
log2 expression
21.3
18.3
15.3
12.3
9.3
−9.3
−13.3
−17.3
−21.3
12
10
8
6
4
2
0
folP
slr2026
slr2027
ncr0330
771250
773750
776250
778750
770000
772500
775000
777500
780000
sll1916
nadD
sll1915
sll1913
sll3615
sll1911
sll1910
zam
Position (nt)

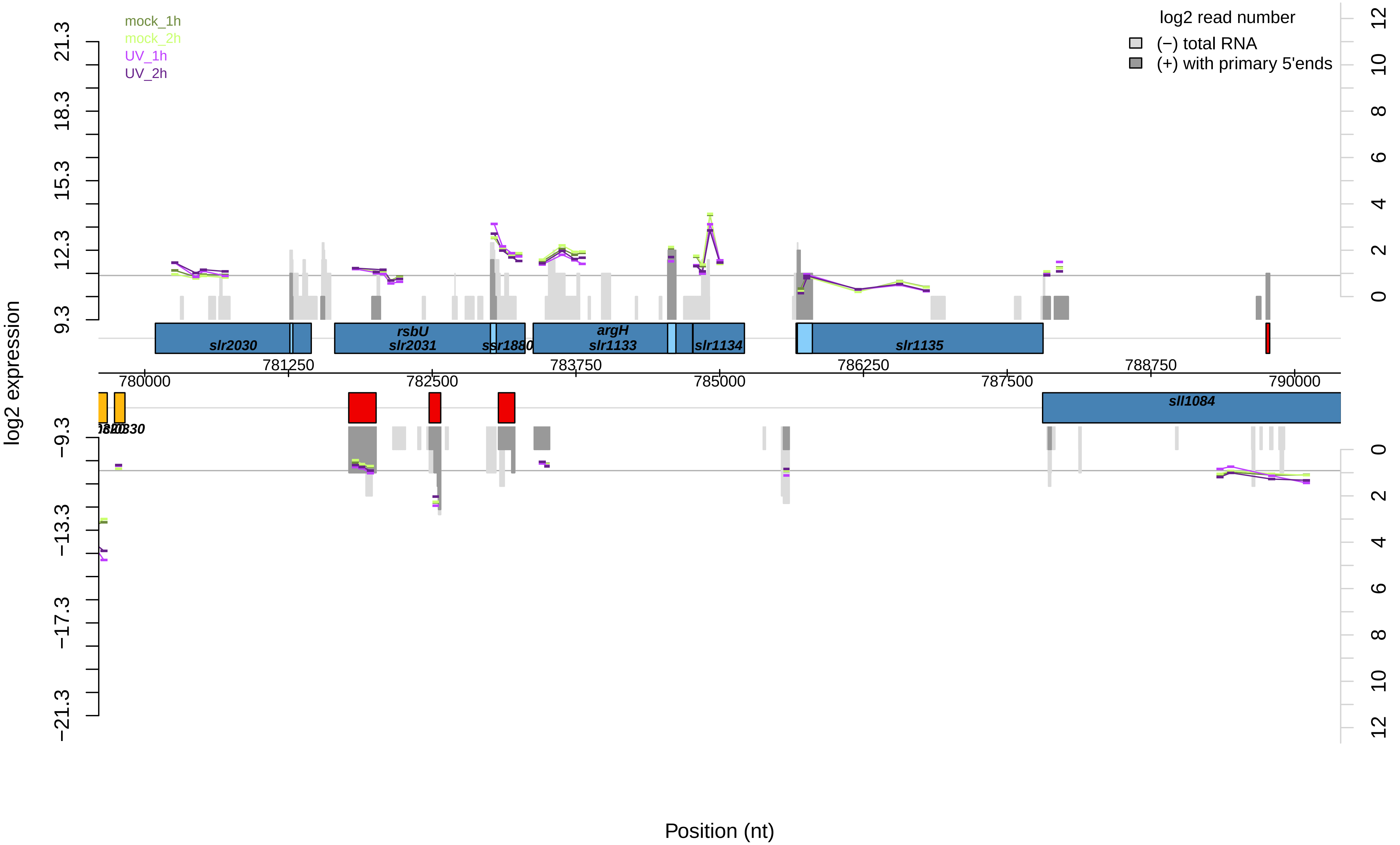

mock_1h
mock_2h
UV_1h
UV_2h
log2 read number
(−) total RNA
(+) with primary 5'ends
log2 expression
21.3
18.3
15.3
12.3
9.3
−9.3
−13.3
−17.3
−21.3
12
10
8
6
4
2
0
slr2030
rsbU
slr2031
ssr1880
argH
slr1133
slr1134
slr1135
sll1084
781250
783750
786250
788750
780000
782500
785000
787500
790000
Position (nt)

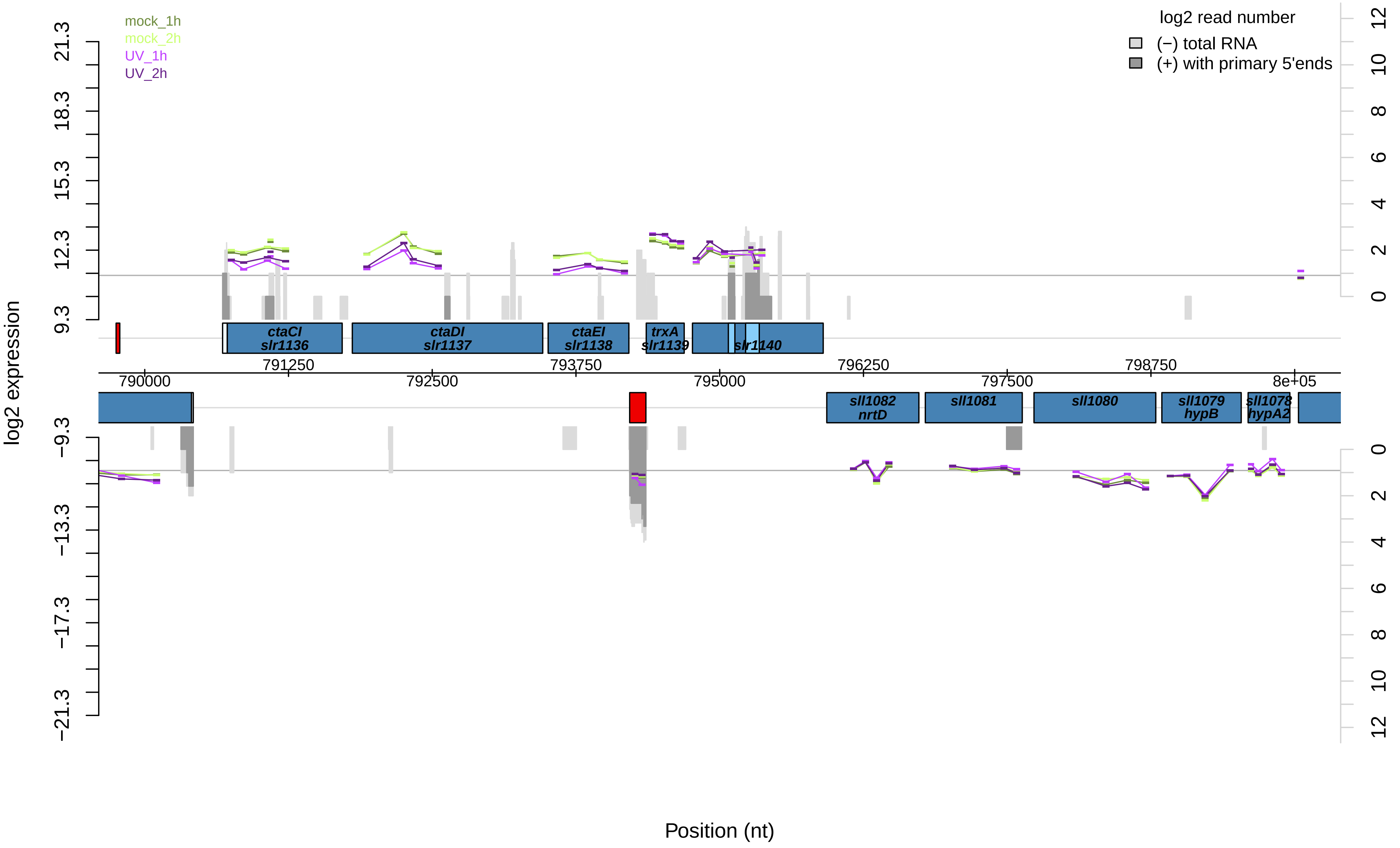

mock_1h
mock_2h
UV_1h
UV_2h
log2 read number
(−) total RNA
(+) with primary 5'ends
log2 expression
21.3
18.3
15.3
12.3
9.3
−9.3
−13.3
−17.3
−21.3
12
10
8
6
4
2
0
ctaCI
slr1136
ctaDI
slr1137
ctaEI
slr1138
trxA
slr1139
slr1140
790000
791250
792500
793750
795000
796250
797500
798750
8e+05
sll1082
nrtD
sll1081
sll1080
sll1079
hypB
sll1078
hypA2
Position (nt)

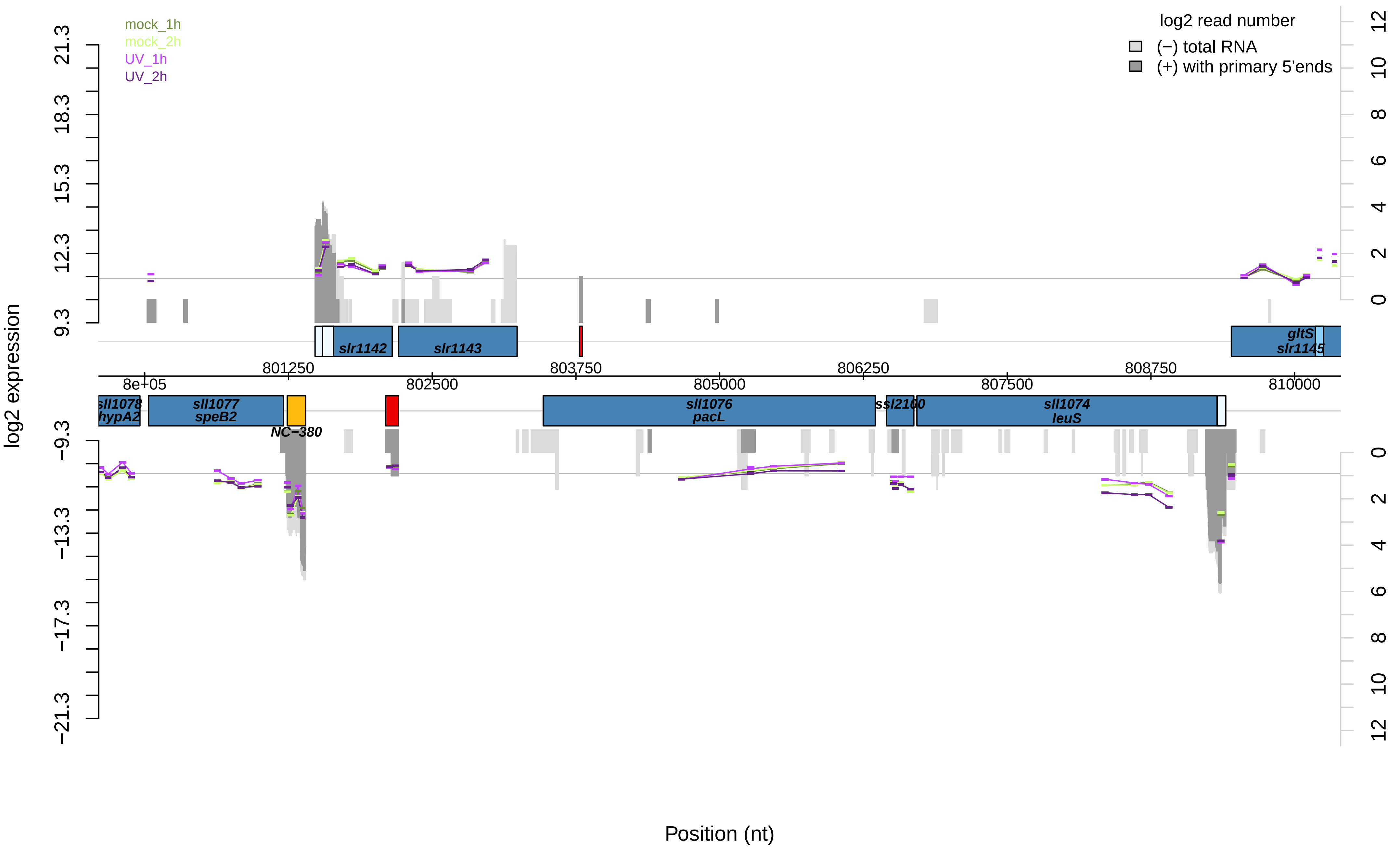

mock_1h
mock_2h
UV_1h
UV_2h
log2 read number
(−) total RNA
(+) with primary 5'ends
log2 expression
21.3
18.3
15.3
12.3
9.3
−9.3
−13.3
−17.3
−21.3
12
10
8
6
4
2
0
slr1142
slr1143
gltS
slr1145
801250
803750
806250
808750
8e+05
802500
805000
807500
810000
sll1078
hypA2
sll1077
speB2
NC−380
sll1076
pacL
ssl2100
sll1074
leuS
Position (nt)

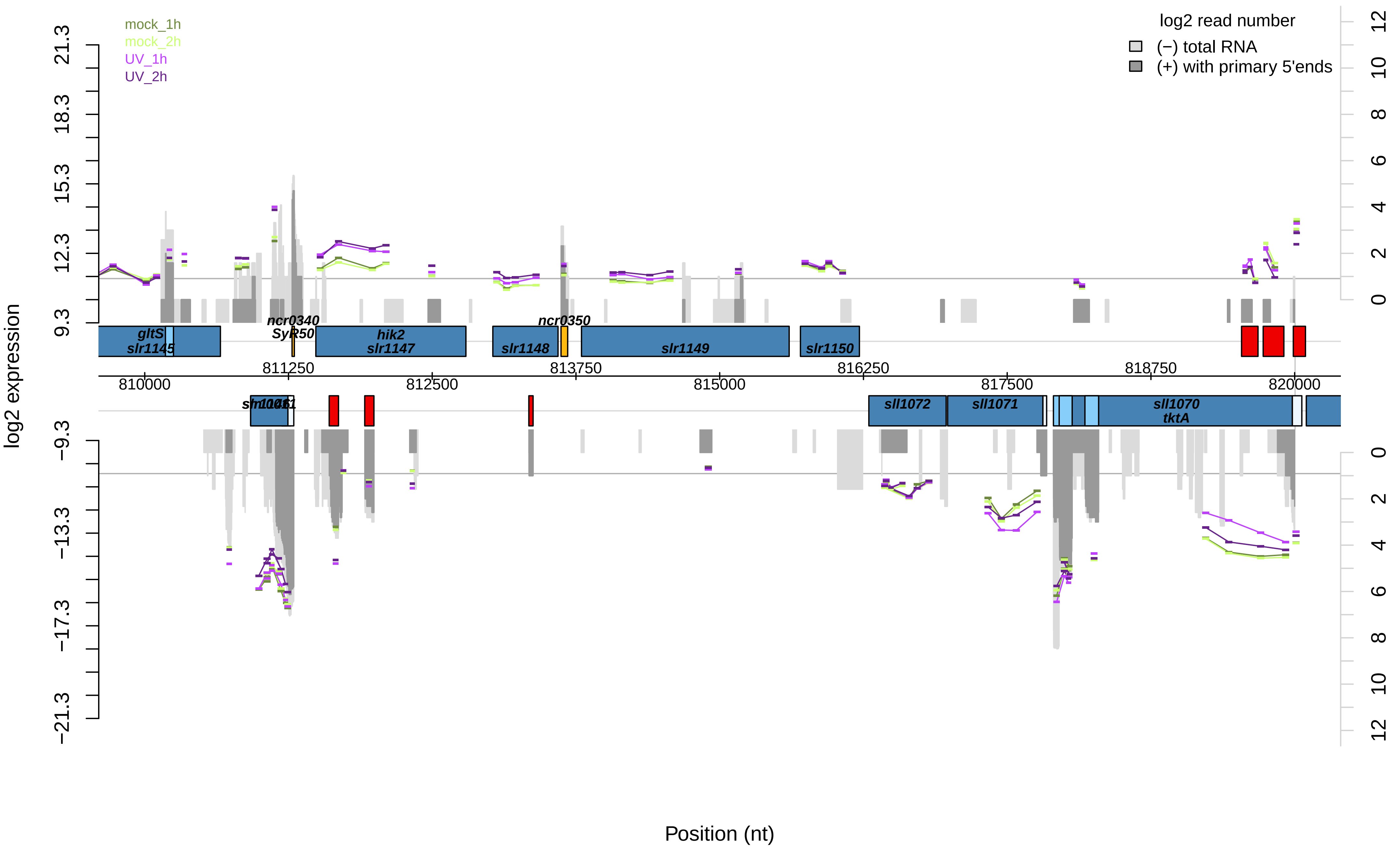

mock_1h
mock_2h
UV_1h
UV_2h
log2 read number
(−) total RNA
(+) with primary 5'ends
log2 expression
Position (nt)
21.3
18.3
15.3
12.3
9.3
−9.3
−13.3
−17.3
−21.3
12
10
8
6
4
2
0
gltS
slr1145
ncr0340
SyR50
hik2
slr1147
slr1148
ncr0350
slr1149
slr1150
810000
811250
812500
813750
815000
816250
817500
818750
820000
sll1072
sll1071
sll1070
tktA

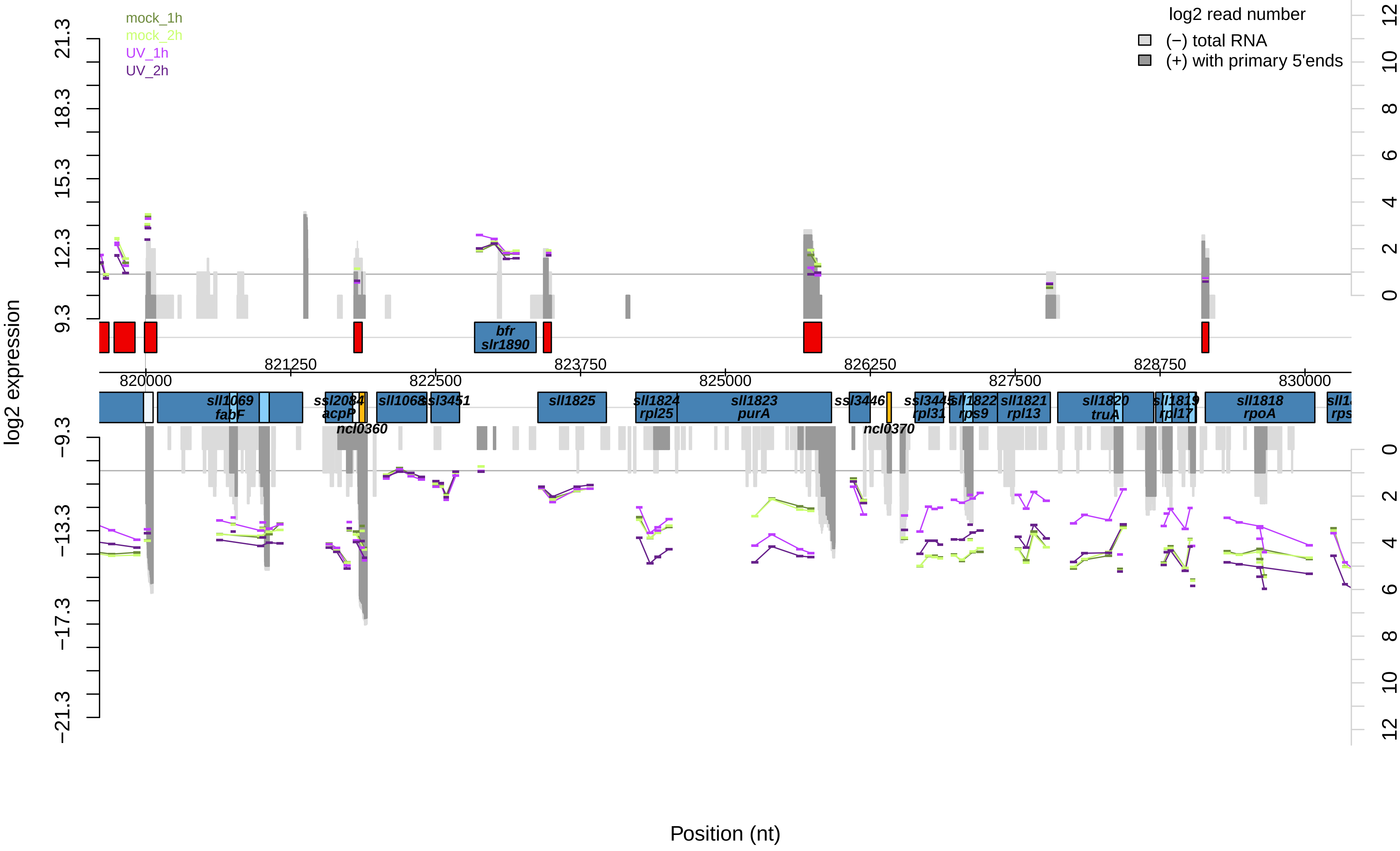

mock_1h
mock_2h
UV_1h
UV_2h
log2 read number
(−) total RNA
(+) with primary 5'ends
log2 expression
21.3
18.3
15.3
12.3
9.3
−9.3
−13.3
−17.3
−21.3
12
10
8
6
4
2
0
bfr
slr1890
821250
822500
823750
825000
826250
827500
828750
830000
820000
sll1069
fabF
ssl2084
acpP
ncl0360
sll1068
ssl3451
sll1825
sll1824
rpl25
sll1823
purA
ssl3446
ncl0370
ssl3445
rpl31
sll1822
rps9
sll1821
rpl13
sll1820
truA
sll1819
rpl17
sll1818
rpoA
Position (nt)

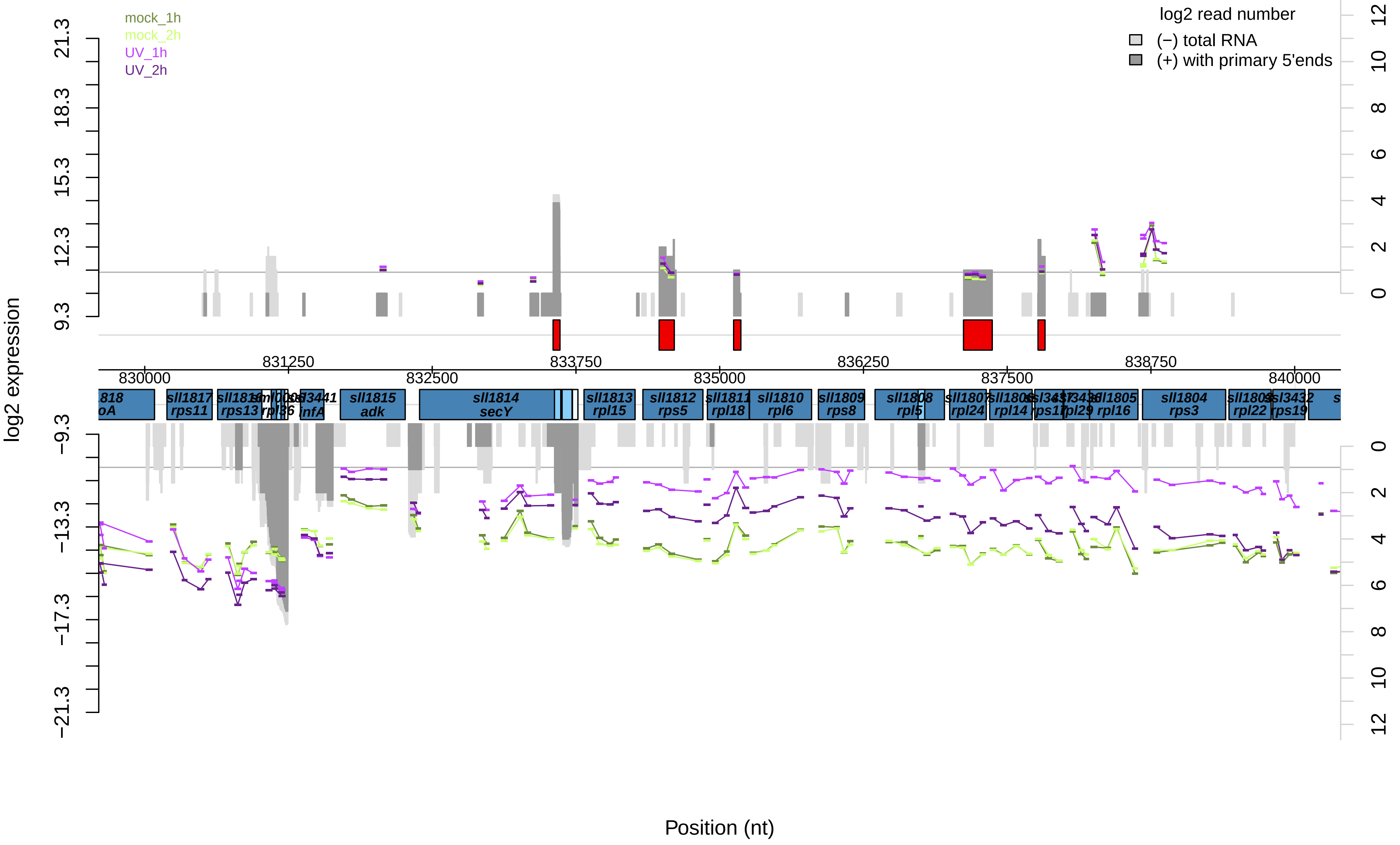

mock_1h
mock_2h
UV_1h
UV_2h
log2 read number
(−) total RNA
(+) with primary 5'ends
log2 expression
Position (nt)
21.3
18.3
15.3
12.3
9.3
−9.3
−13.3
−17.3
−21.3
831250
833750
836250
838750
830000
832500
835000
837500
840000
sll1817 rps11
sll1816 rps13
rpl36
infA
sll1815 adk
sll1814 secY
sll1813 rpl15
sll1812 rps5
sll1811 rpl18
sll1810 rpl6
sll1809 rps8
sll1808 rpl5
rpl24
rpl14
rps17
rpl29
sll1805 rpl16
sll1804 rps3
rpl22
rps19

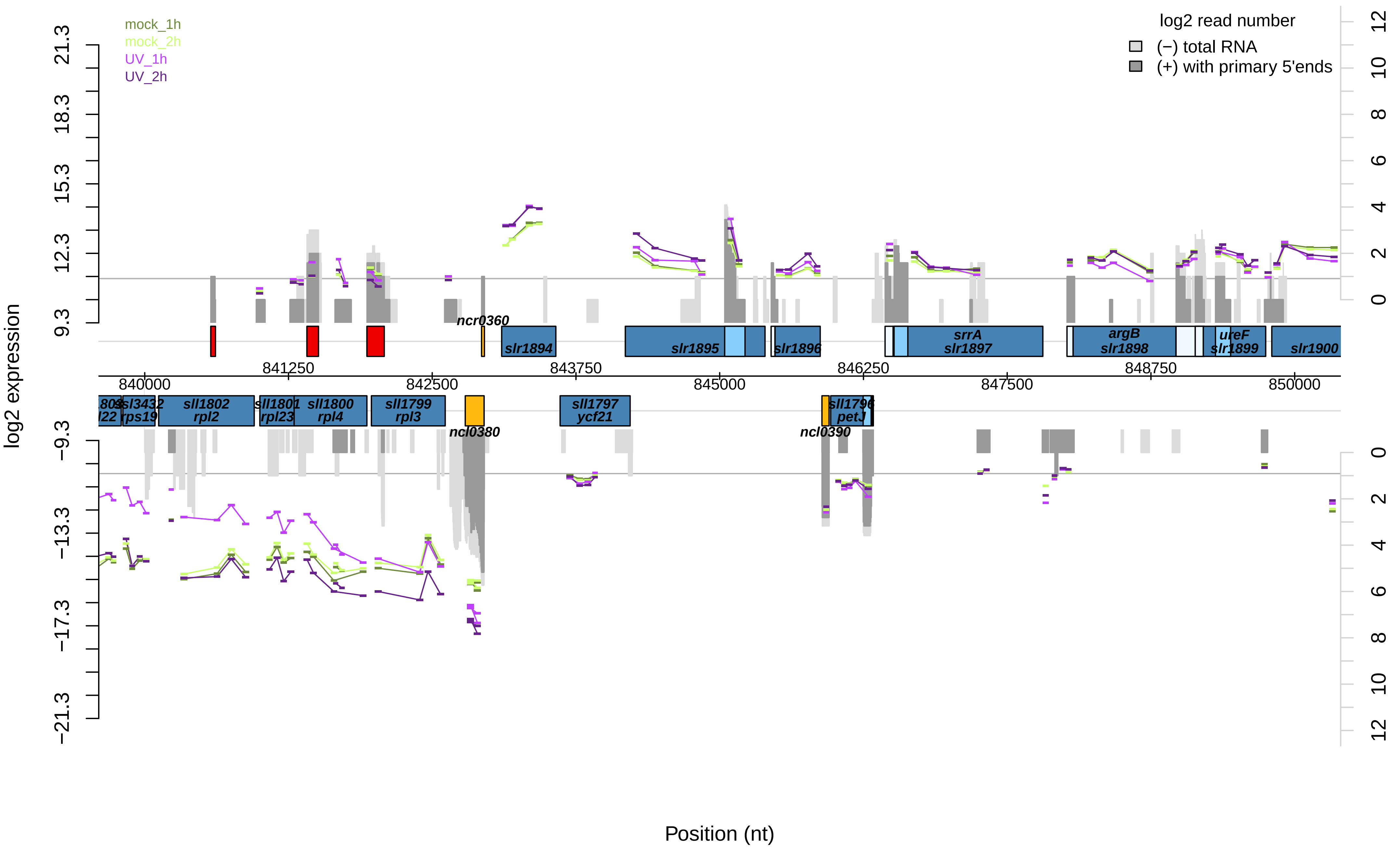

mock_1h
mock_2h
UV_1h
UV_2h
log2 read number
(−) total RNA
(+) with primary 5'ends
log2 expression
Position (nt)
ncr0360
slr1894
slr1895
slr1896
srrA
slr1897
argB
slr1898
ureF
slr1899
slr1900
sll1802
rpl2
sll1801
rpl23
sll1800
rpl4
sll1799
rpl3
ncl0380
sll1797
ycf21
ncl0390
sll1796
petJ
rps19

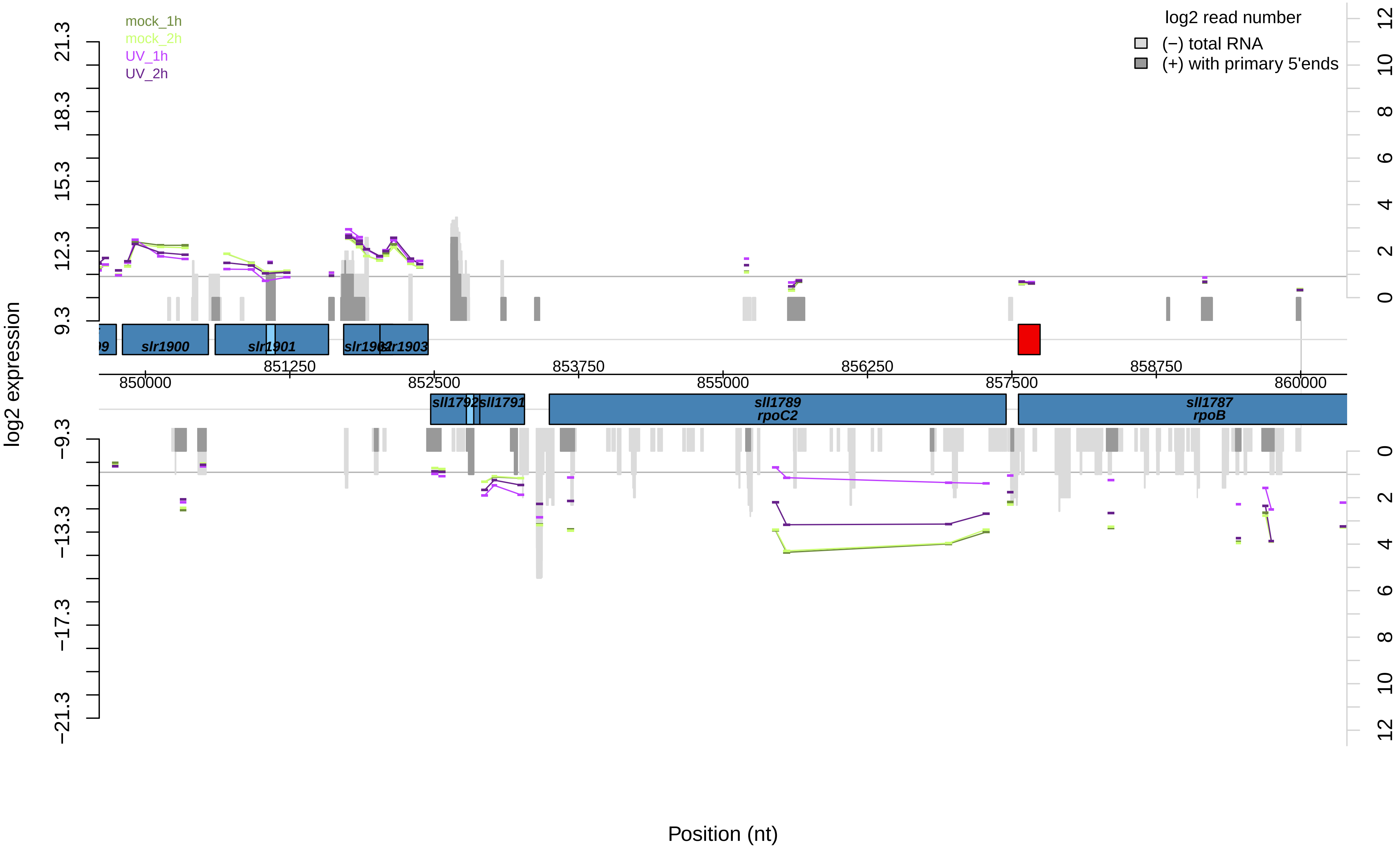
mock_1h
mock_2h
UV_1h
UV_2h
log2 read number
(−) total RNA
(+) with primary 5'ends
log2 expression
21.3
18.3
15.3
12.3
9.3
−9.3
−13.3
−17.3
−21.3
12
10
8
6
4
2
0
slr1900
slr1901
slr1902
slr1903
sll1792
sll1791
sll1789
rpoC2
sll1787
rpoB
850000
851250
852500
853750
855000
856250
857500
858750
860000
Position (nt)

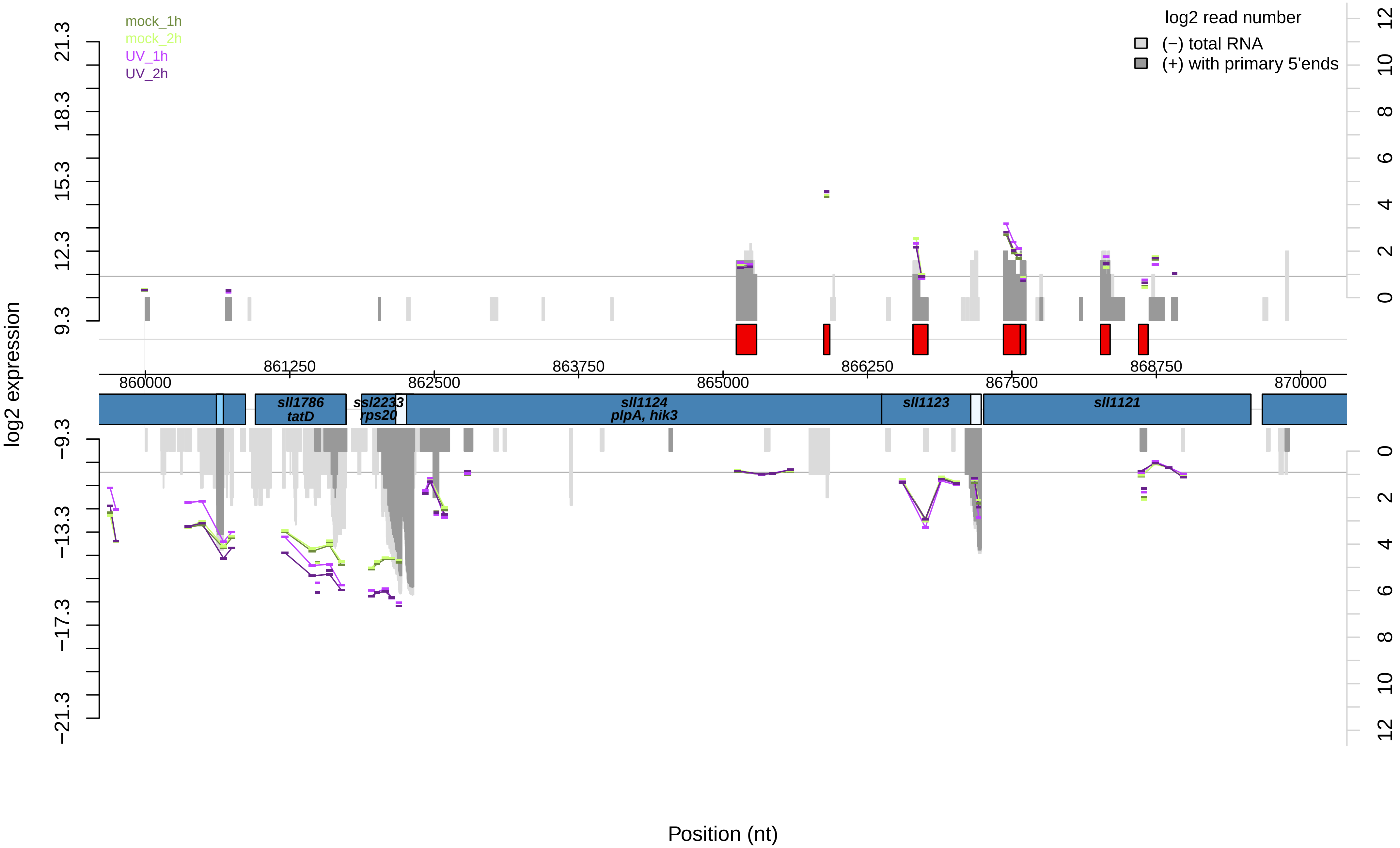

mock_1h
mock_2h
UV_1h
UV_2h
log2 read number
(−) total RNA
(+) with primary 5'ends
log2 expression
21.3
18.3
15.3
12.3
9.3
−9.3
−13.3
−17.3
−21.3
12
10
8
6
4
2
0
860000
861250
862500
863750
865000
866250
867500
868750
870000
sll1786
tatD
ssl2233
rps20
sll1124
plpA, hik3
sll1123
sll1121
Position (nt)

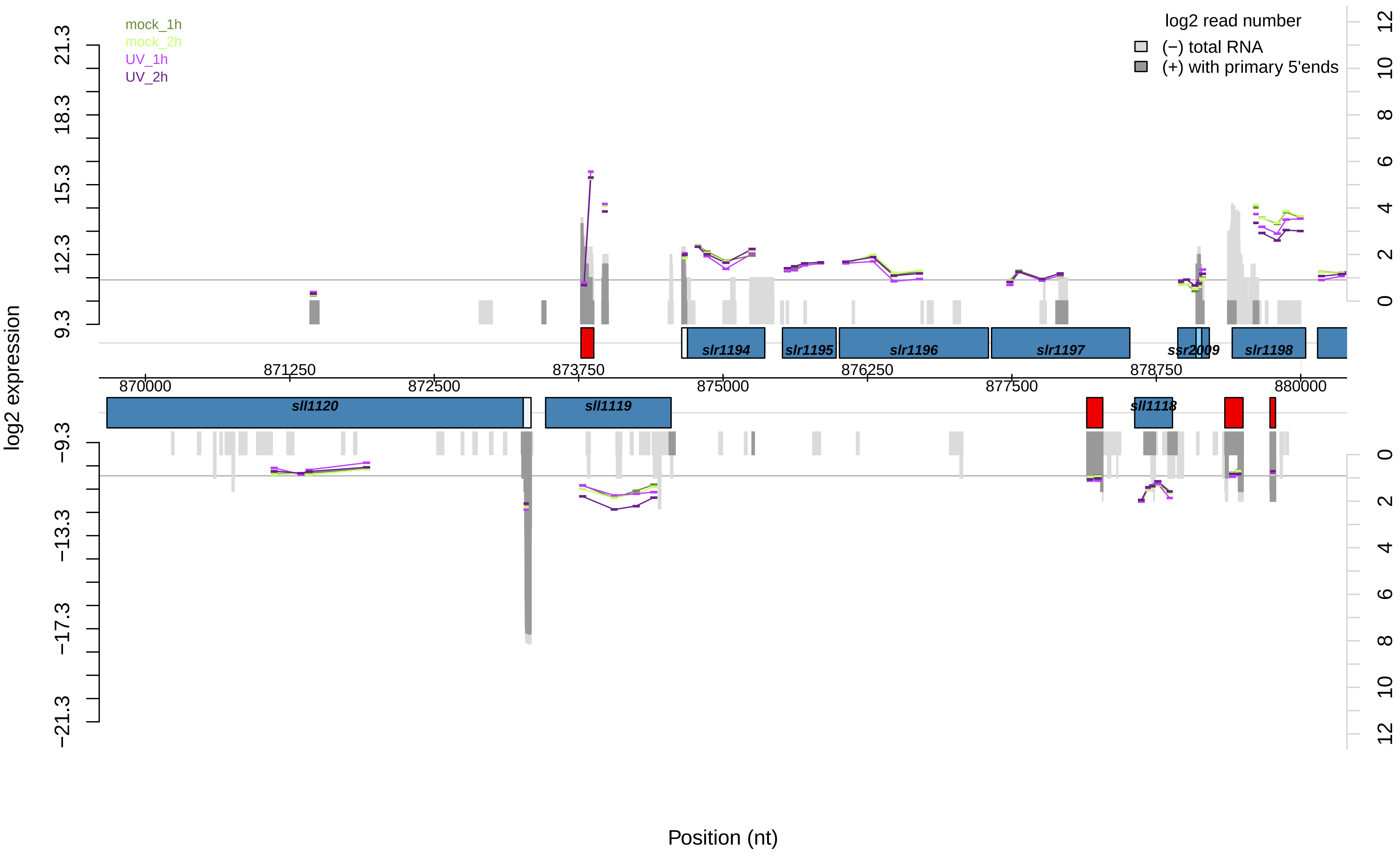
mock_1h
mock_2h
UV_1h
UV_2h
log2 read number
(−) total RNA
(+) with primary 5'ends
log2 expression
21.3
18.3
15.3
12.3
9.3
−9.3
−13.3
−17.3
−21.3
12
10
8
6
4
2
0
slr1194
slr1195
slr1196
slr1197
ssr2009
slr1198
871250
873750
876250
878750
870000
872500
875000
877500
880000
sll1120
sll1119
sll1118
Position (nt)

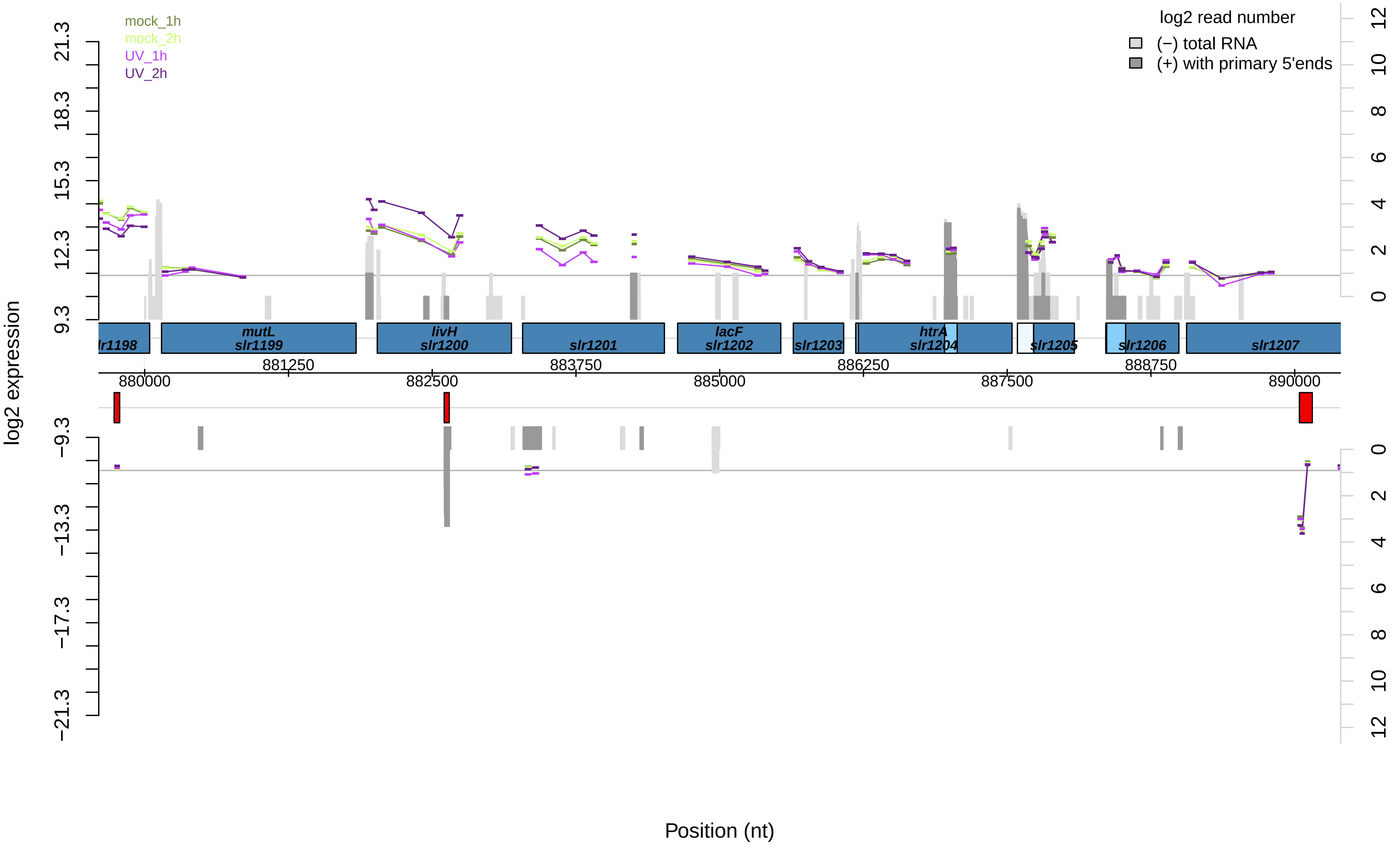
mock_1h
mock_2h
UV_1h
UV_2h
log2 read number
(−) total RNA
(+) with primary 5'ends
log2 expression
21.3
18.3
15.3
12.3
9.3
−9.3
−13.3
−17.3
−21.3
12
10
8
6
4
2
0
mutL
slr1199
livH
slr1200
slr1201
lacF
slr1202
slr1203
htrA
slr1204
slr1205
slr1206
slr1207
881250
883750
886250
888750
880000
882500
885000
887500
890000
Position (nt)

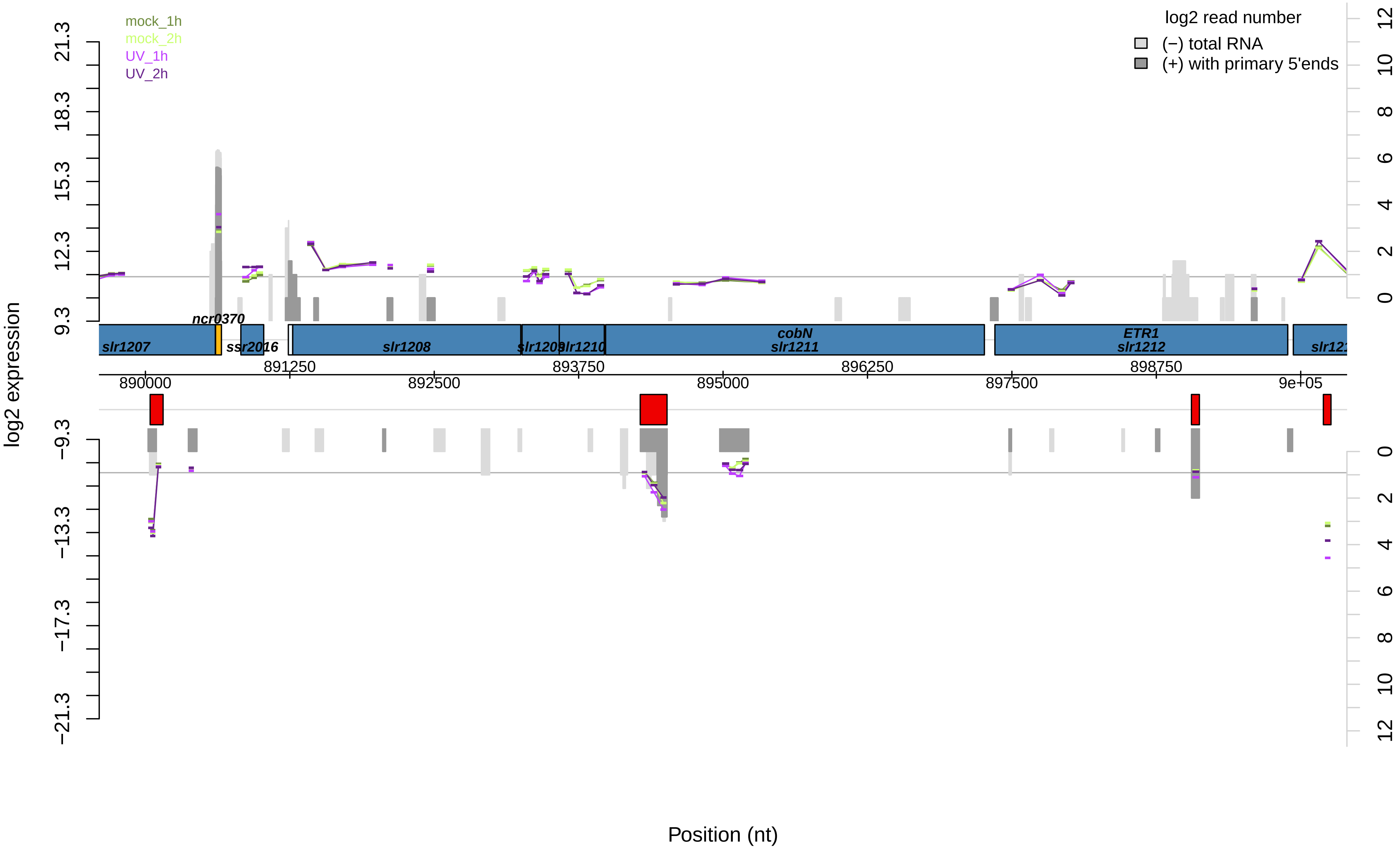

mock_1h
mock_2h
UV_1h
UV_2h
log2 read number
(−) total RNA
(+) with primary 5'ends
log2 expression
21.3
18.3
15.3
12.3
9.3
−9.3
−13.3
−17.3
−21.3
12
10
8
6
4
2
0
ncr0370
slr1207
ssr2016
slr1208
slr1209
slr1210
cobN
slr1211
ETR1
slr1212
891250
893750
896250
898750
890000
892500
895000
897500
9e+05
Position (nt)

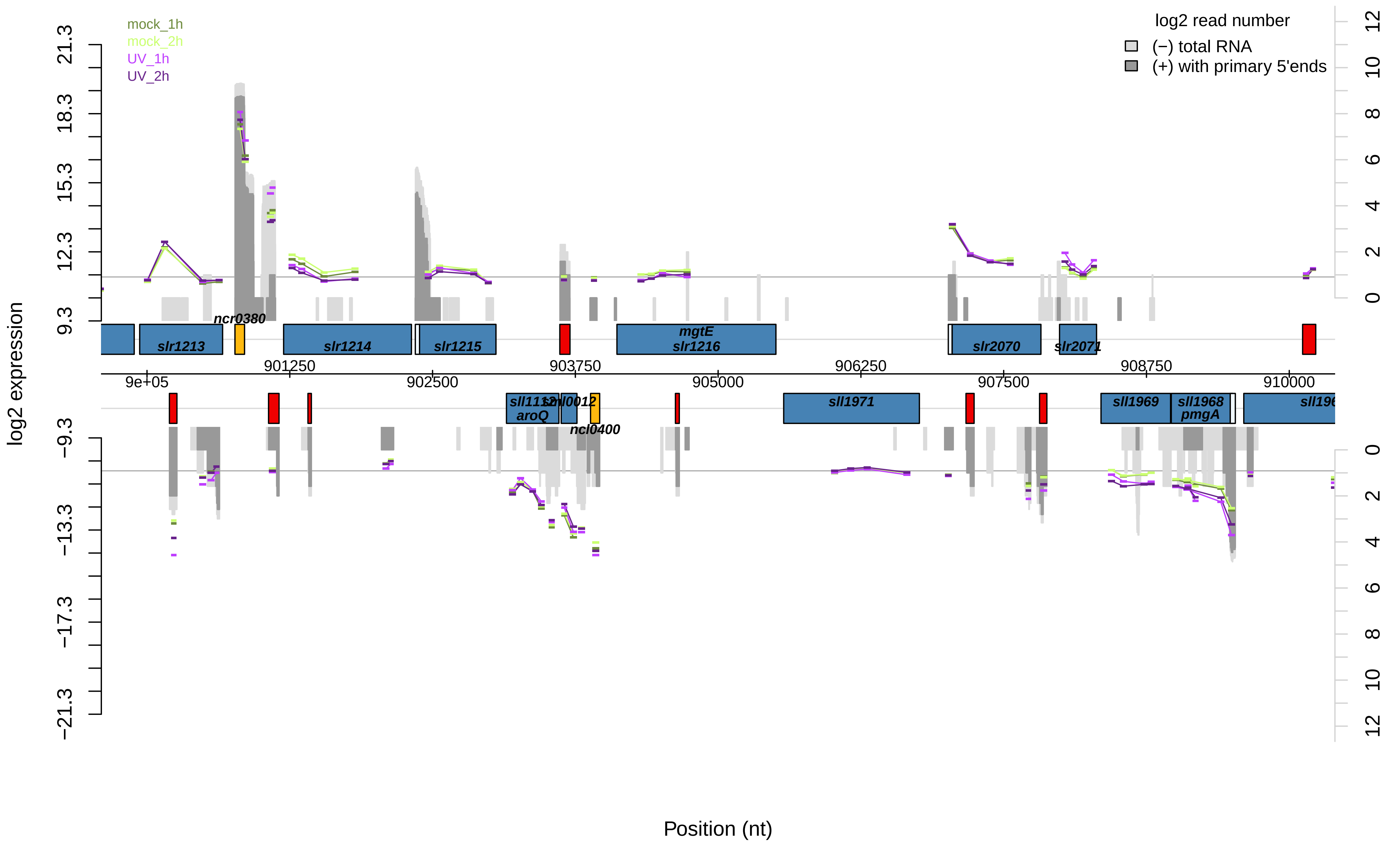
mock_1h
mock_2h
UV_1h
UV_2h
log2 read number
(−) total RNA
(+) with primary 5'ends
log2 expression
21.3
18.3
15.3
12.3
9.3
−9.3
−13.3
−17.3
−21.3
slr1213
ncr0380
slr1214
slr1215
mgtE
slr1216
slr2070
slr2071
901250
903750
906250
908750
9e+05
902500
905000
907500
910000
sll1112
aroQ
ncl0400
sll1971
sll1969
sll1968
pmgA
Position (nt)

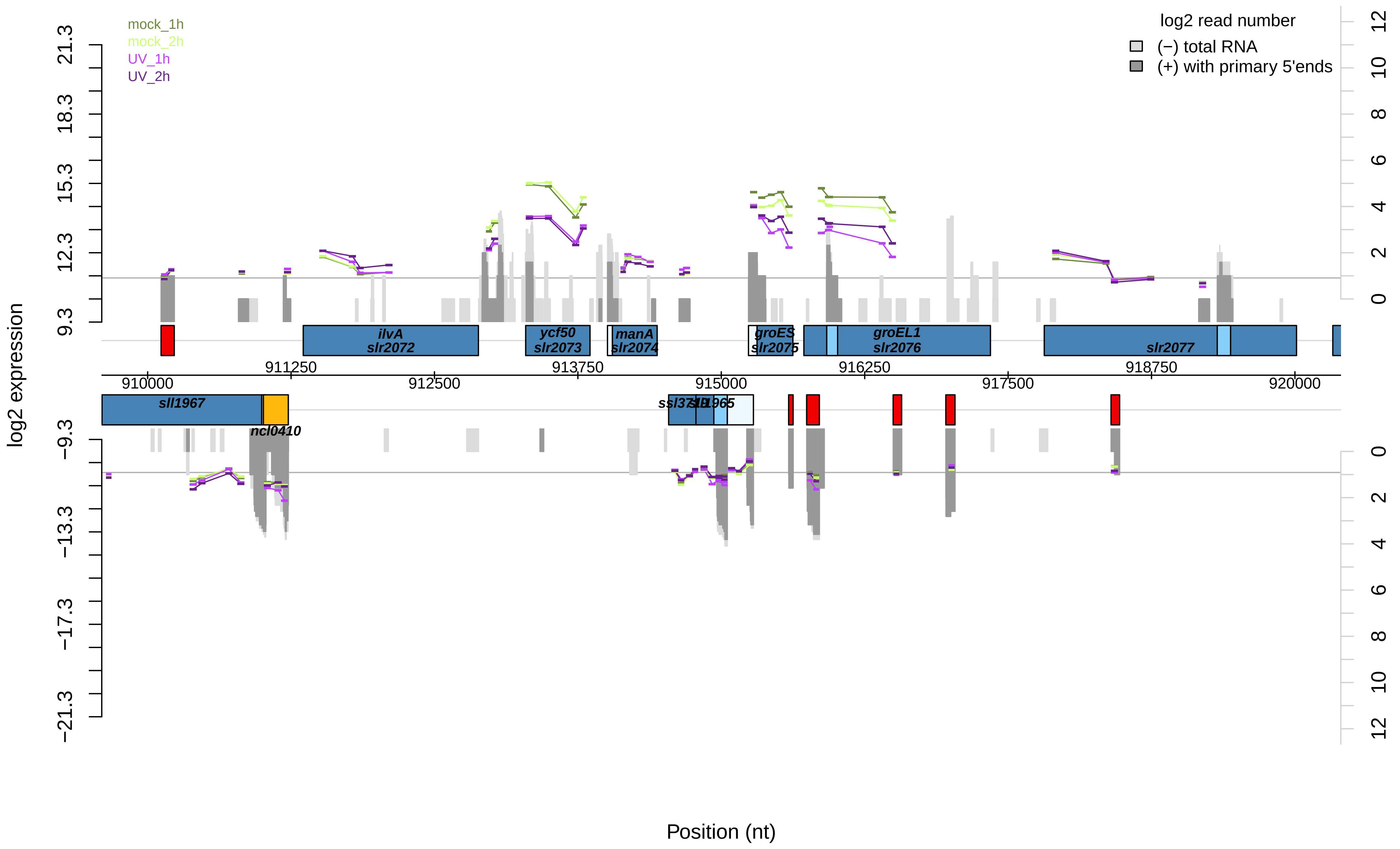

mock_1h
mock_2h
UV_1h
UV_2h
log2 read number
(−) total RNA
(+) with primary 5'ends
log2 expression
21.3
18.3
15.3
12.3
9.3
−9.3
−13.3
−17.3
−21.3
12
10
8
6
4
2
0
ilvA
slr2072
ycf50
slr2073
manA
slr2074
groES
slr2075
groEL1
slr2076
slr2077
sll1967
ncl0410
911250
913750
916250
918750
910000
912500
915000
917500
920000
Position (nt)

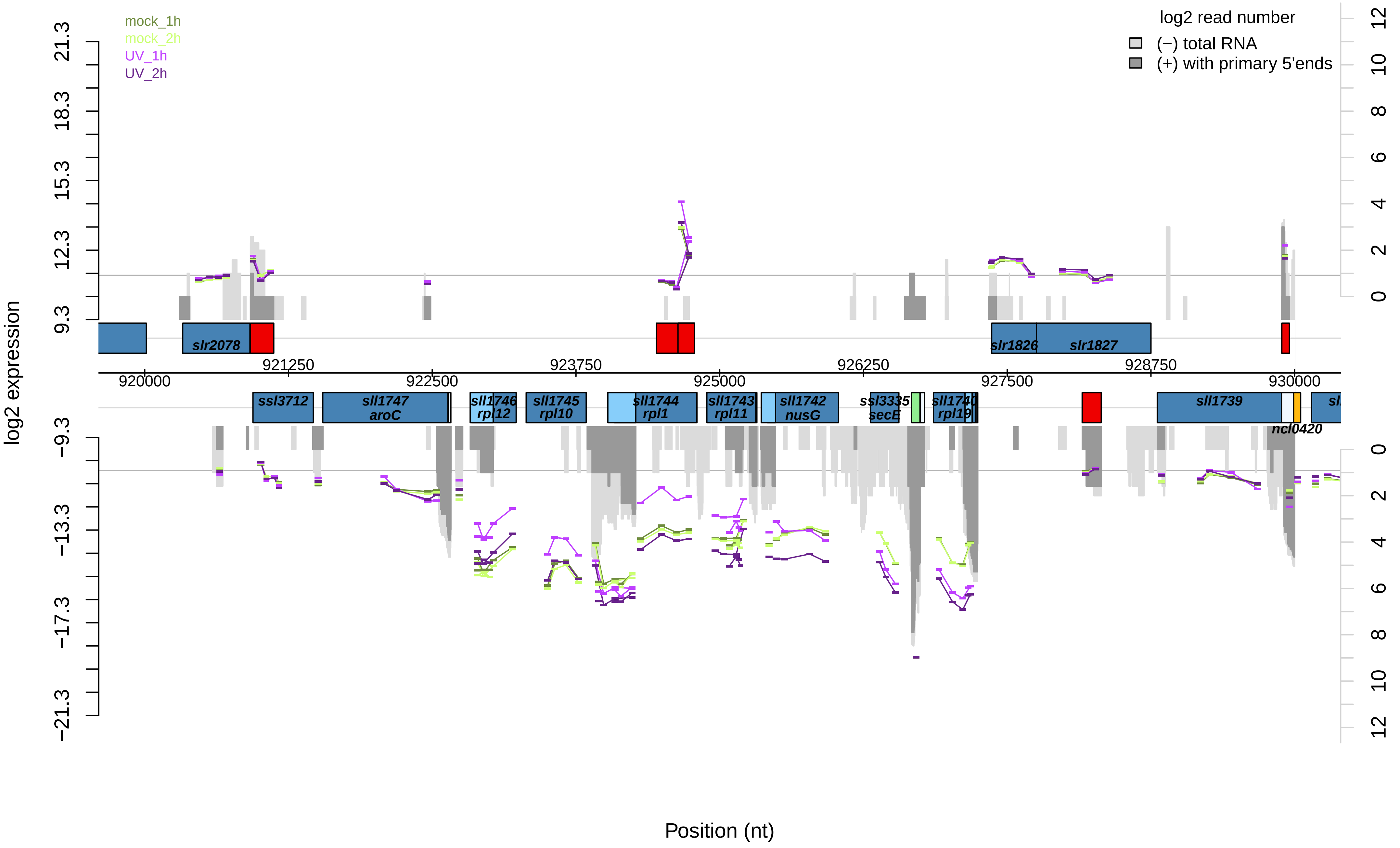

mock_1h
mock_2h
UV_1h
UV_2h
log2 read number
(−) total RNA
(+) with primary 5'ends
log2 expression
Position (nt)
21.3
18.3
15.3
12.3
9.3
−9.3
−13.3
−17.3
−21.3
12
10
8
6
4
2
0
slr2078
slr1826
slr1827
921250
923750
926250
928750
920000
922500
925000
927500
930000
ssl3712
sll1747
aroC
sll1746
rpl12
sll1745
rpl10
sll1744
rpl1
sll1743
rpl11
sll1742
nusG
ssl3335
secE
sll1740
rpl19
sll1739
ncl0420

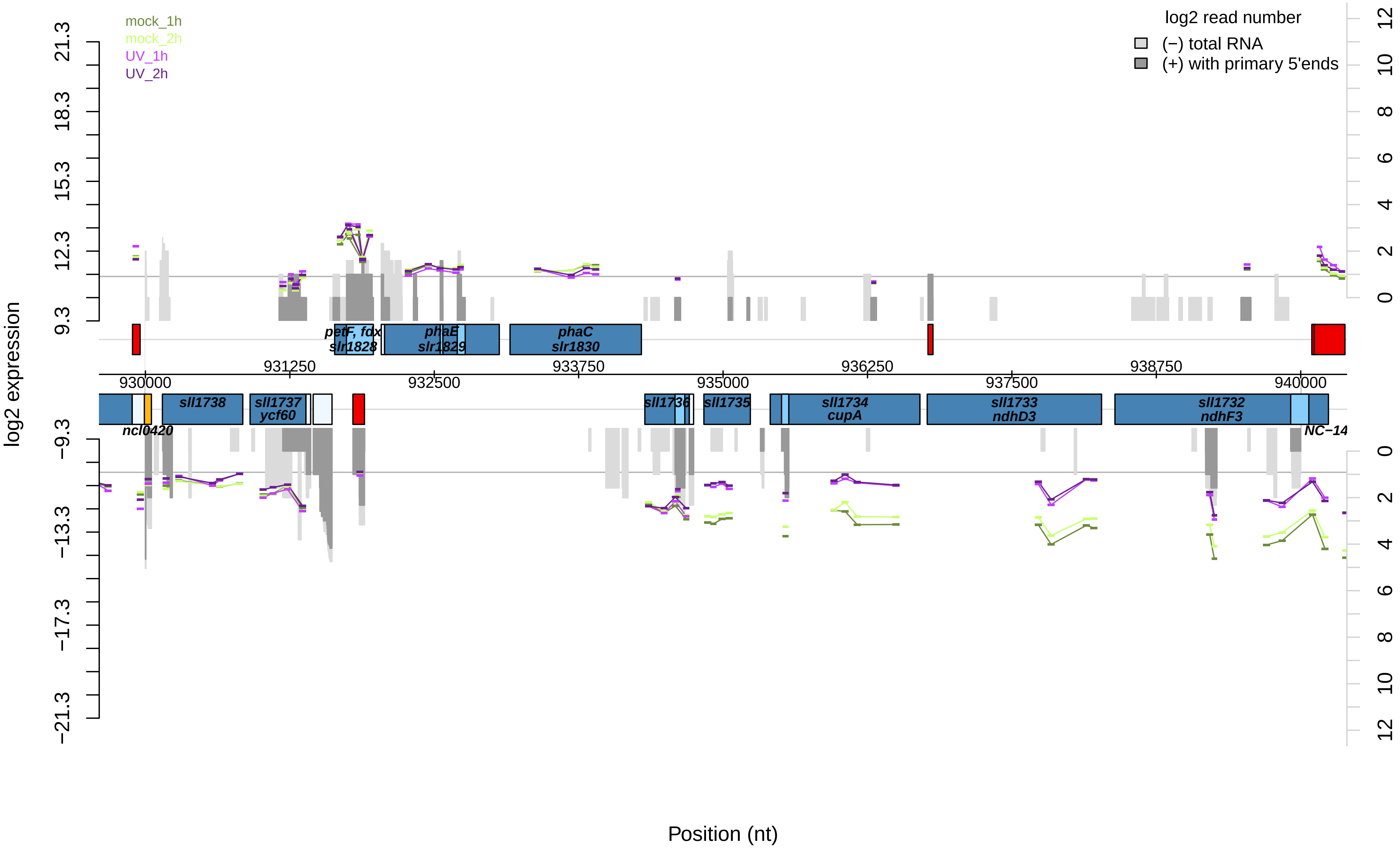

mock_1h
mock_2h
UV_1h
UV_2h
log2 read number
(−) total RNA
(+) with primary 5'ends
log2 expression
Position (nt)
930000
931250
932500
933750
935000
936250
937500
938750
940000
petF, fdx
slr1828
phaE
slr1829
phaC
slr1830
sll1738
sll1737
ycf60
ncl0420
sll1736
sll1735
sll1734
cupA
sll1733
ndhD3
sll1732
ndhF3
NC−14
21.3
18.3
15.3
12.3
9.3
−9.3
−13.3
−17.3
−21.3

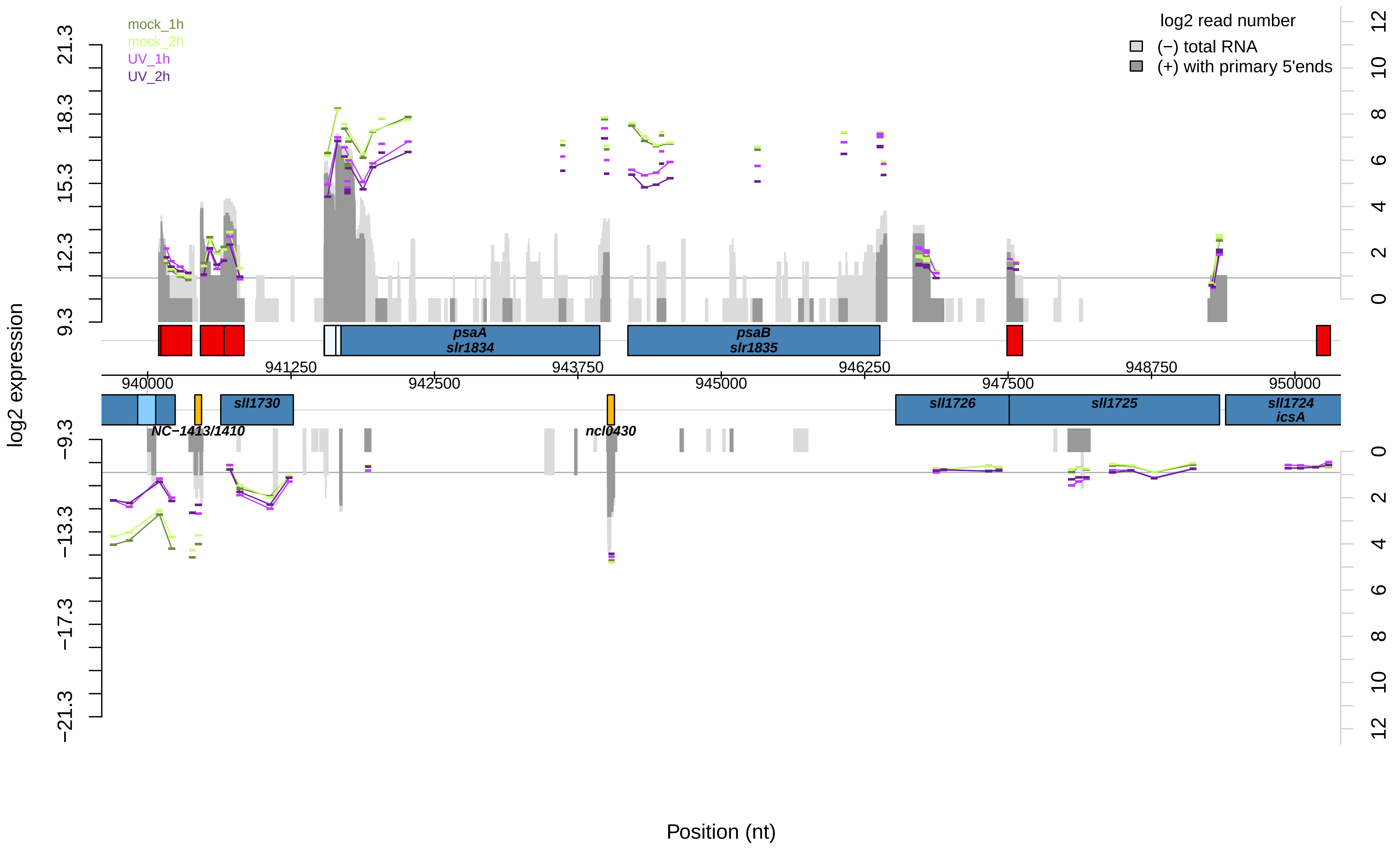
mock_1h
mock_2h
UV_1h
UV_2h
log2 read number
(−) total RNA
(+) with primary 5'ends
log2 expression
psaA
slr1834
psaB
slr1835
sll1730
NC−1413/1410
ncl0430
sll1726
sll1725
sll1724
icsA
941250
943750
946250
948750
940000
942500
945000
947500
950000
21.3
18.3
15.3
12.3
9.3
−9.3
−13.3
−17.3
−21.3
Position (nt)

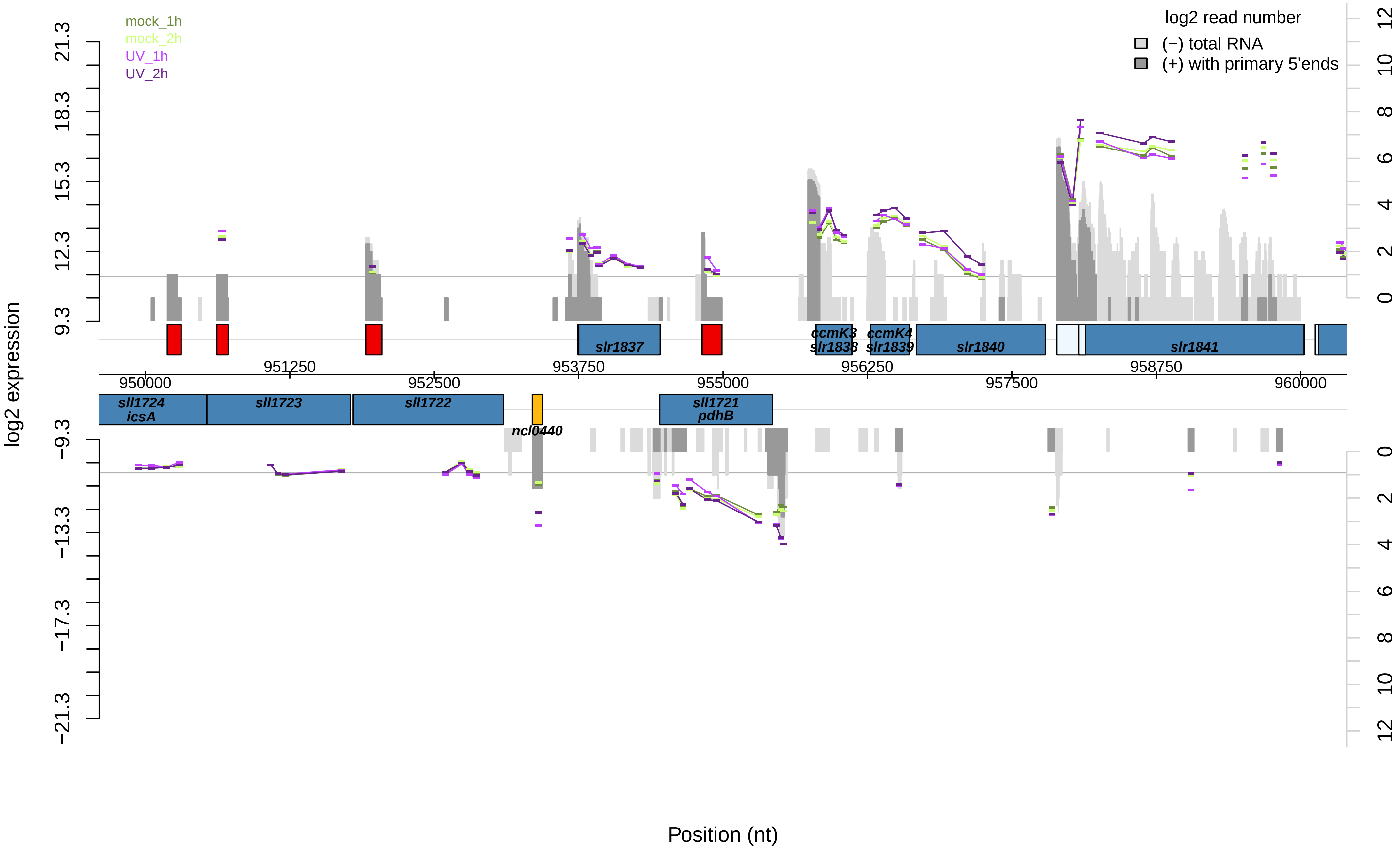
mock_1h
mock_2h
UV_1h
UV_2h
log2 read number
(−) total RNA
(+) with primary 5'ends
log2 expression
Position (nt)
sll1724 icsA
sll1723
sll1722
ncl0440
sll1721 pdhB
slr1837
ccmK3 slr1838
ccmK4 slr1839
slr1840
slr1841
950000
951250
952500
953750
955000
956250
957500
958750
960000

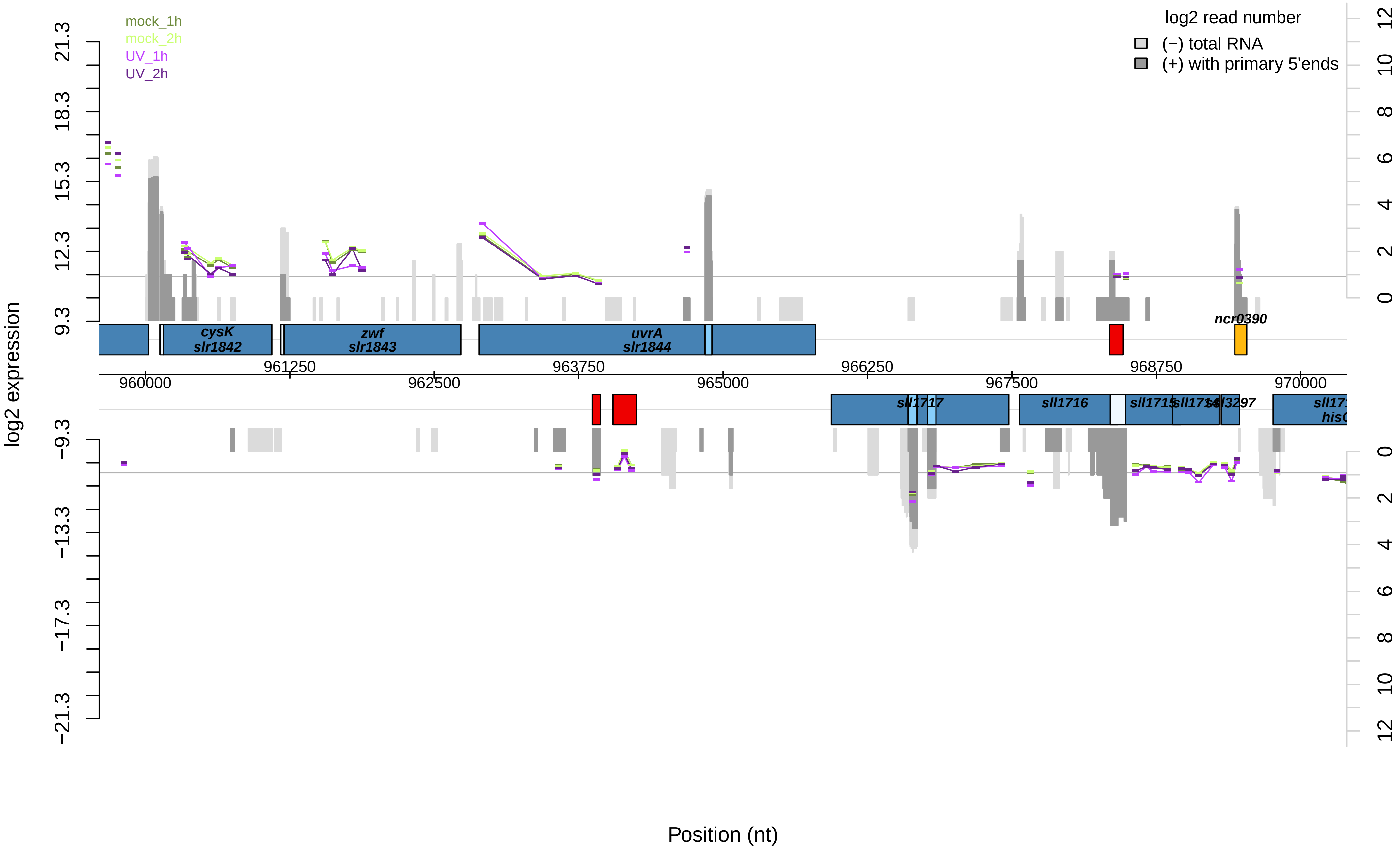

mock_1h
mock_2h
UV_1h
UV_2h
log2 read number
(−) total RNA
(+) with primary 5'ends
log2 expression
21.3
18.3
15.3
12.3
9.3
−9.3
−13.3
−17.3
−21.3
cysK
slr1842
zwf
slr1843
uvrA
slr1844
ncr0390
961250
963750
966250
968750
960000
962500
965000
967500
970000
sll1717
sll1716
sll1715
sll1714
ssl3297
Position (nt)

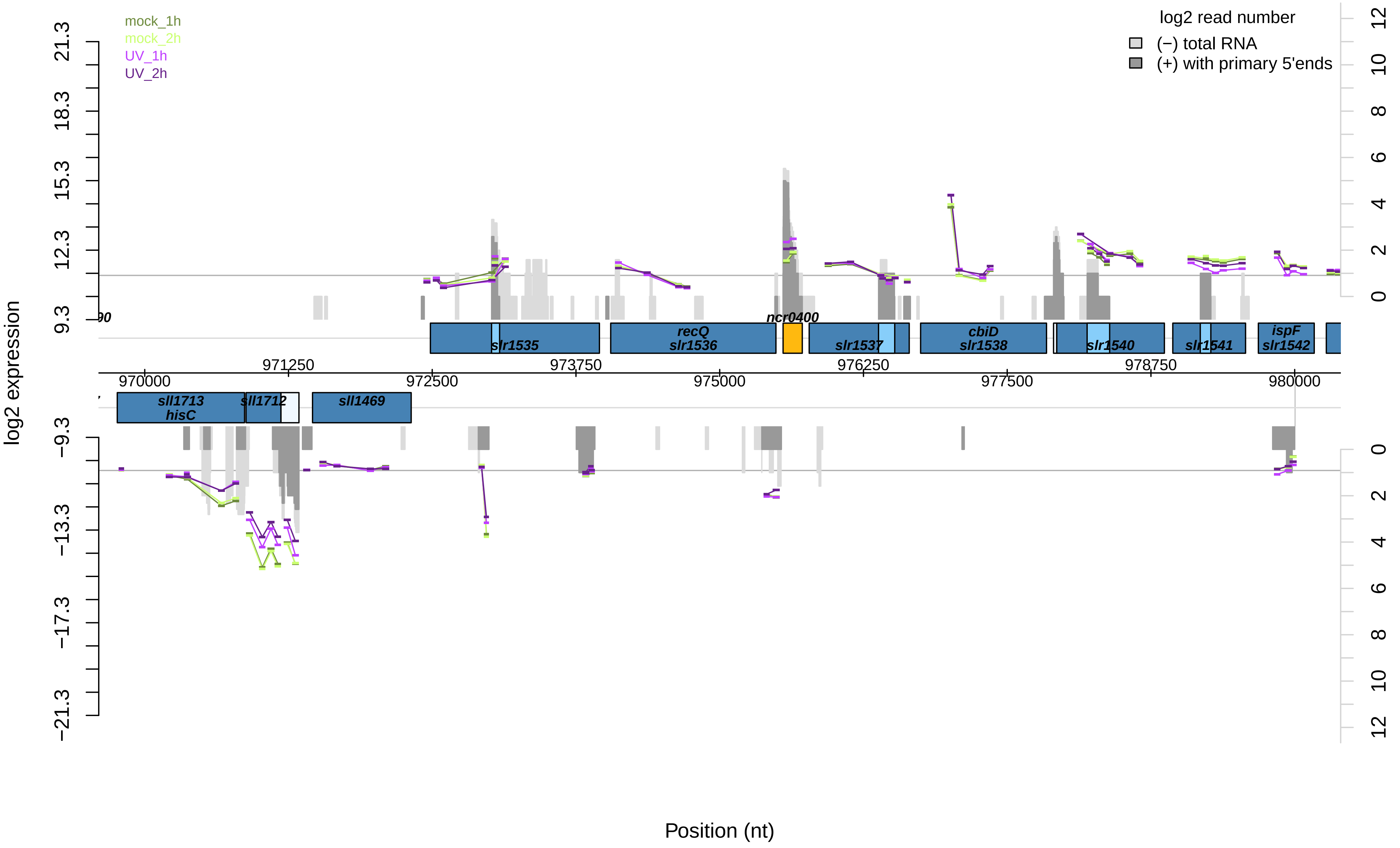
mock_1h
mock_2h
UV_1h
UV_2h
log2 read number
(−) total RNA
(+) with primary 5'ends
log2 expression
21.3
18.3
15.3
12.3
9.3
−9.3
−13.3
−17.3
−21.3
12
10
8
6
4
2
0
slr1535
recQ
slr1536
ncr0400
slr1537
cbiD
slr1538
slr1540
slr1541
ispF
slr1542
971250
973750
976250
978750
970000
972500
975000
977500
980000
sll1713
hisC
sll1712
sll1469
Position (nt)

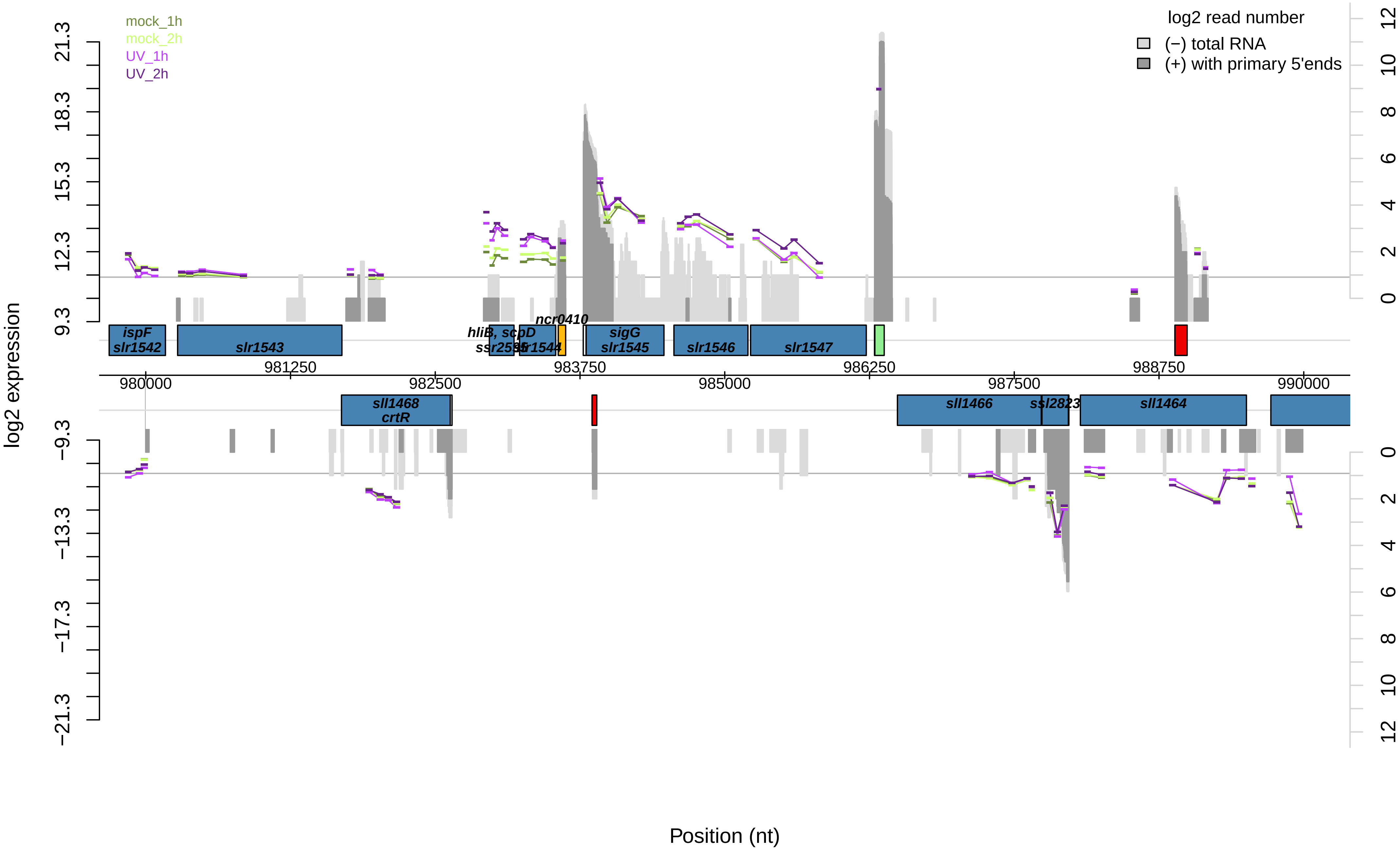

mock_1h
mock_2h
UV_1h
UV_2h
log2 read number
(−) total RNA
(+) with primary 5'ends
log2 expression
Position (nt)
21.3
18.3
15.3
12.3
9.3
−9.3
−13.3
−17.3
−21.3
0
2
4
6
8
10
12
980000
981250
982500
983750
985000
986250
987500
988750
990000
ispF
slr1542
slr1543
hliB, scpD
slr1544
ncr0410
sigG
slr1545
slr1546
slr1547
sll1468
crtR
sll1466
ssl2823
sll1464

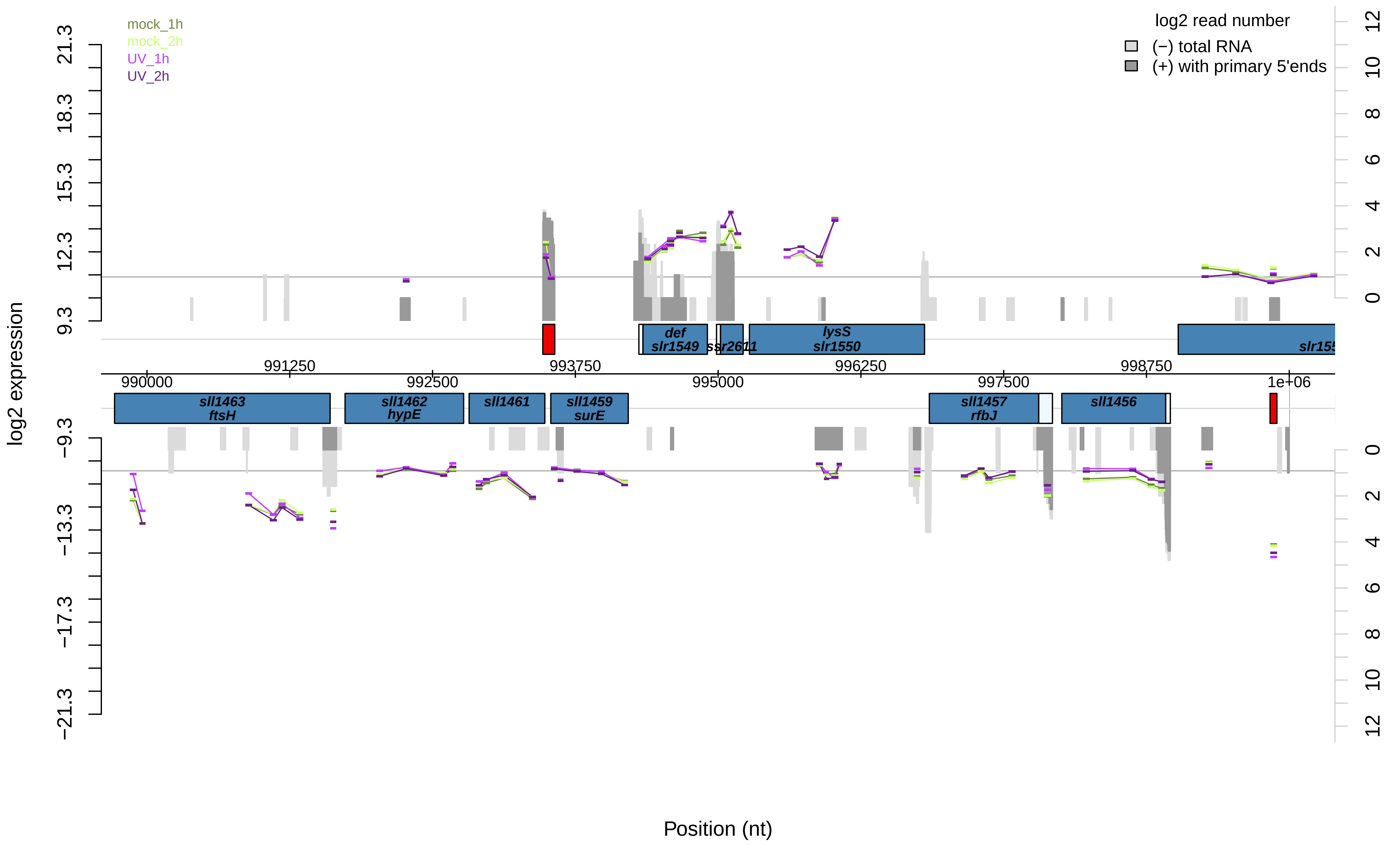

mock_1h
mock_2h
UV_1h
UV_2h
log2 read number
(−) total RNA
(+) with primary 5'ends
log2 expression
Position (nt)
21.3
18.3
15.3
12.3
9.3
−9.3
−13.3
−17.3
−21.3
12
10
8
6
4
2
0
def
slr1549
ssr2611
lysS
slr1550
slr155
991250
993750
996250
998750
990000
992500
995000
997500
1e+06
sll1463
ftsH
sll1462
hypE
sll1461
sll1459
surE
sll1457
rfbJ
sll1456

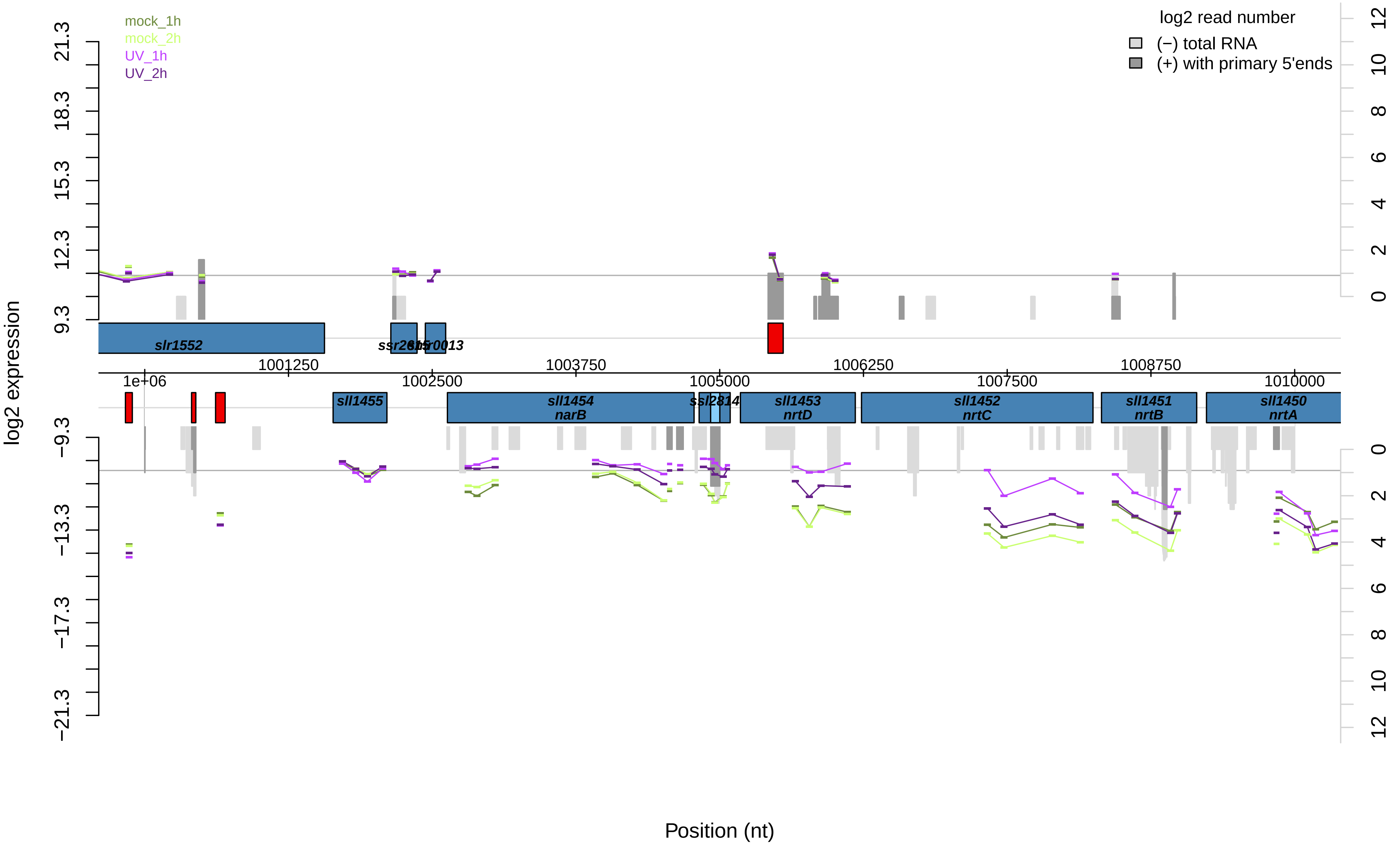

mock_1h
mock_2h
UV_1h
UV_2h
log2 read number
(−) total RNA
(+) with primary 5'ends
log2 expression
21.3
18.3
15.3
12.3
9.3
−9.3
−13.3
−17.3
−21.3
12
10
8
6
4
2
0
slr1552
ssr2615
slr0013
1001250
1003750
1006250
1008750
1e+06
1002500
1005000
1007500
1010000
sll1455
sll1454
narB
ssl2814
sll1453
nrtD
sll1452
nrtC
sll1451
nrtB
sll1450
nrtA
Position (nt)

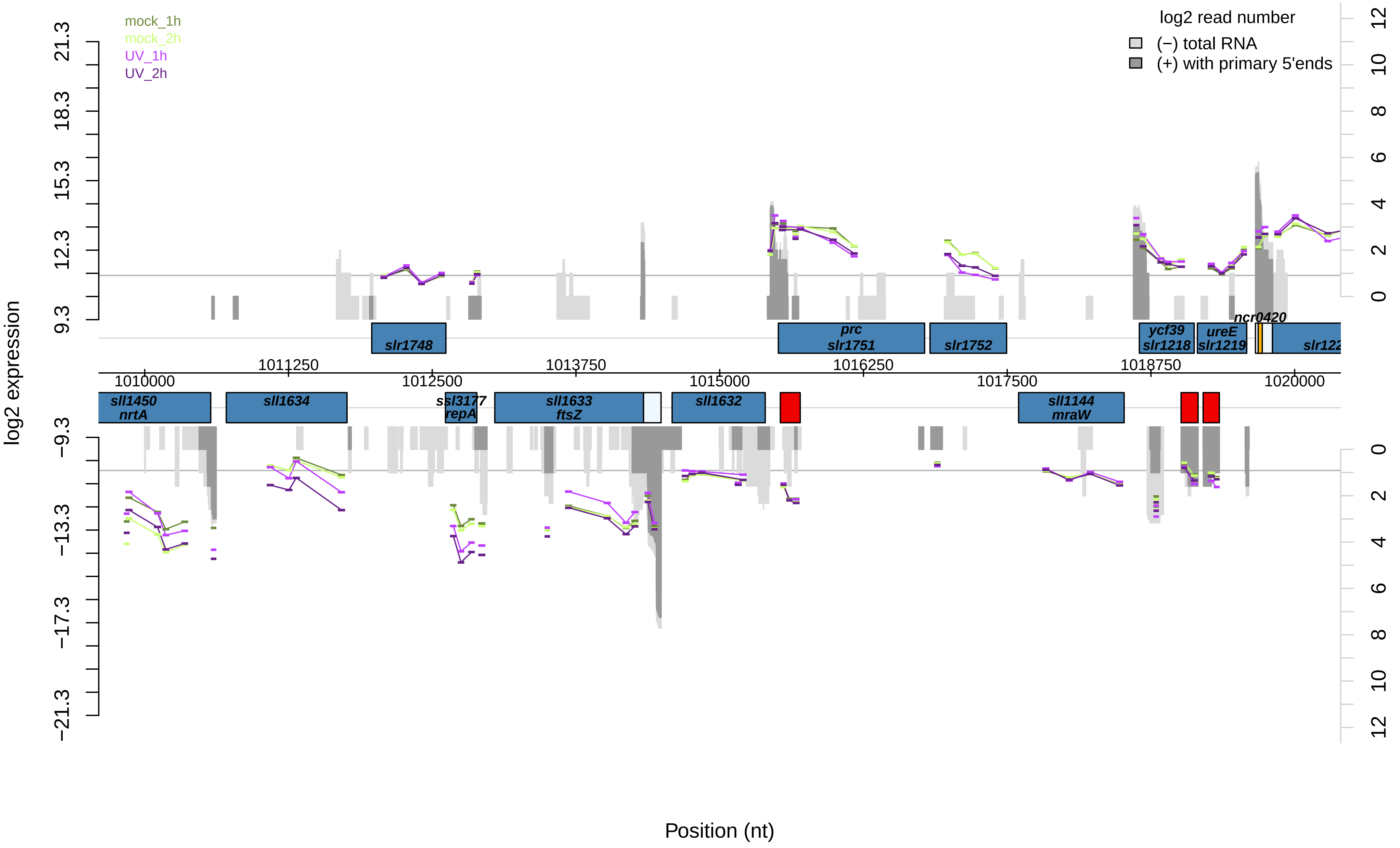
mock_1h
mock_2h
UV_1h
UV_2h
log2 read number
(−) total RNA
(+) with primary 5'ends
log2 expression
21.3
18.3
15.3
12.3
9.3
−9.3
−13.3
−17.3
−21.3
12
10
8
6
4
2
0
slr1748
prc
slr1751
slr1752
ycf39
slr1218
ureE
slr1219
ncr0420
1011250
1013750
1016250
1018750
1010000
1012500
1015000
1017500
1020000
sll1450
nrtA
sll1634
ssl3177
repA
sll1633
ftsZ
sll1632
sll1144
mraW
Position (nt)

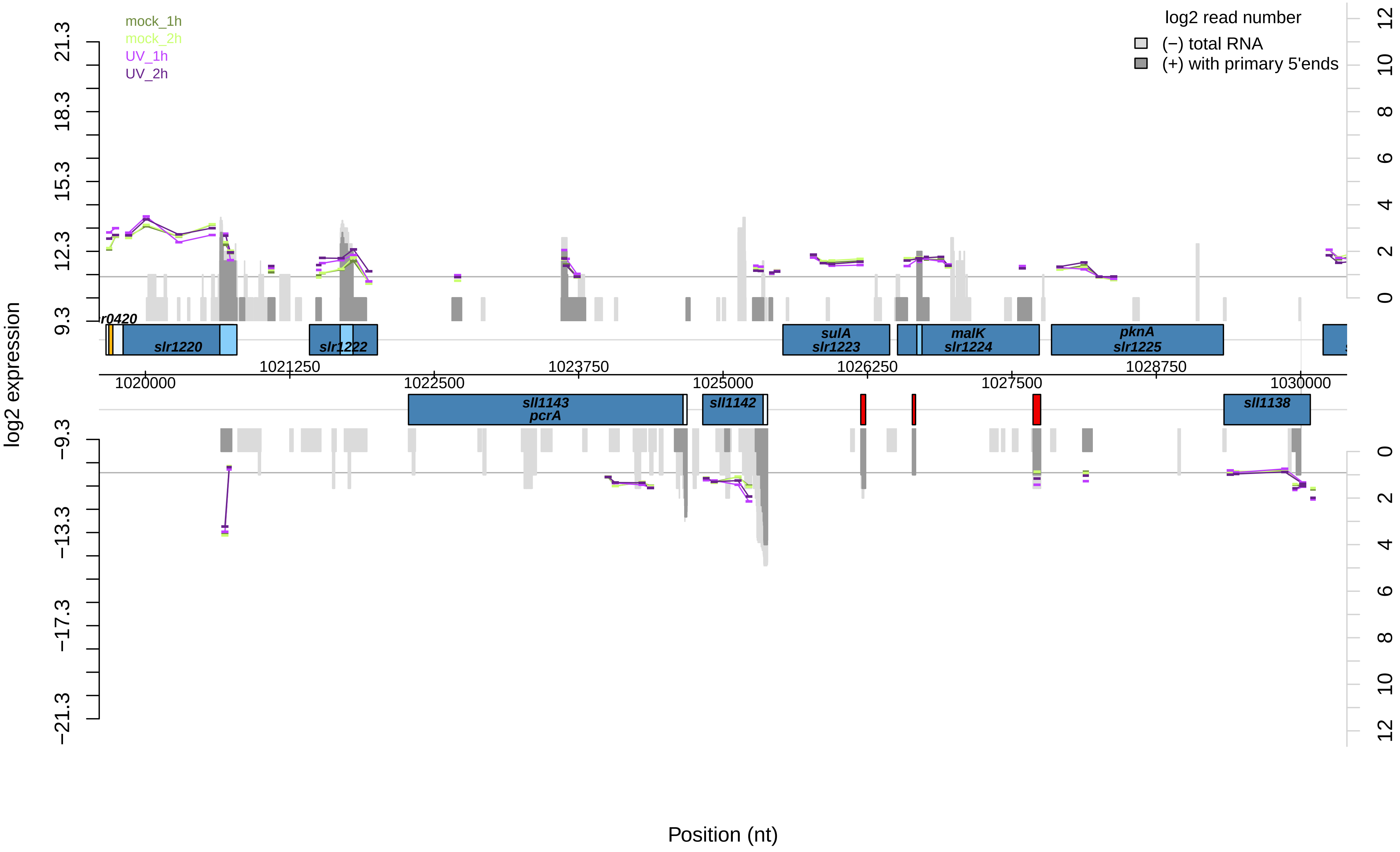

mock_1h
mock_2h
UV_1h
UV_2h
log2 read number
(−) total RNA
(+) with primary 5'ends
log2 expression
21.3
18.3
15.3
12.3
9.3
−9.3
−13.3
−17.3
−21.3
12
10
8
6
4
2
0
r0420
slr1220
slr1222
sulA
slr1223
malK
slr1224
pknA
slr1225
1021250
1023750
1026250
1028750
1020000
1022500
1025000
1027500
1030000
sll1143
pcrA
sll1142
sll1138
Position (nt)

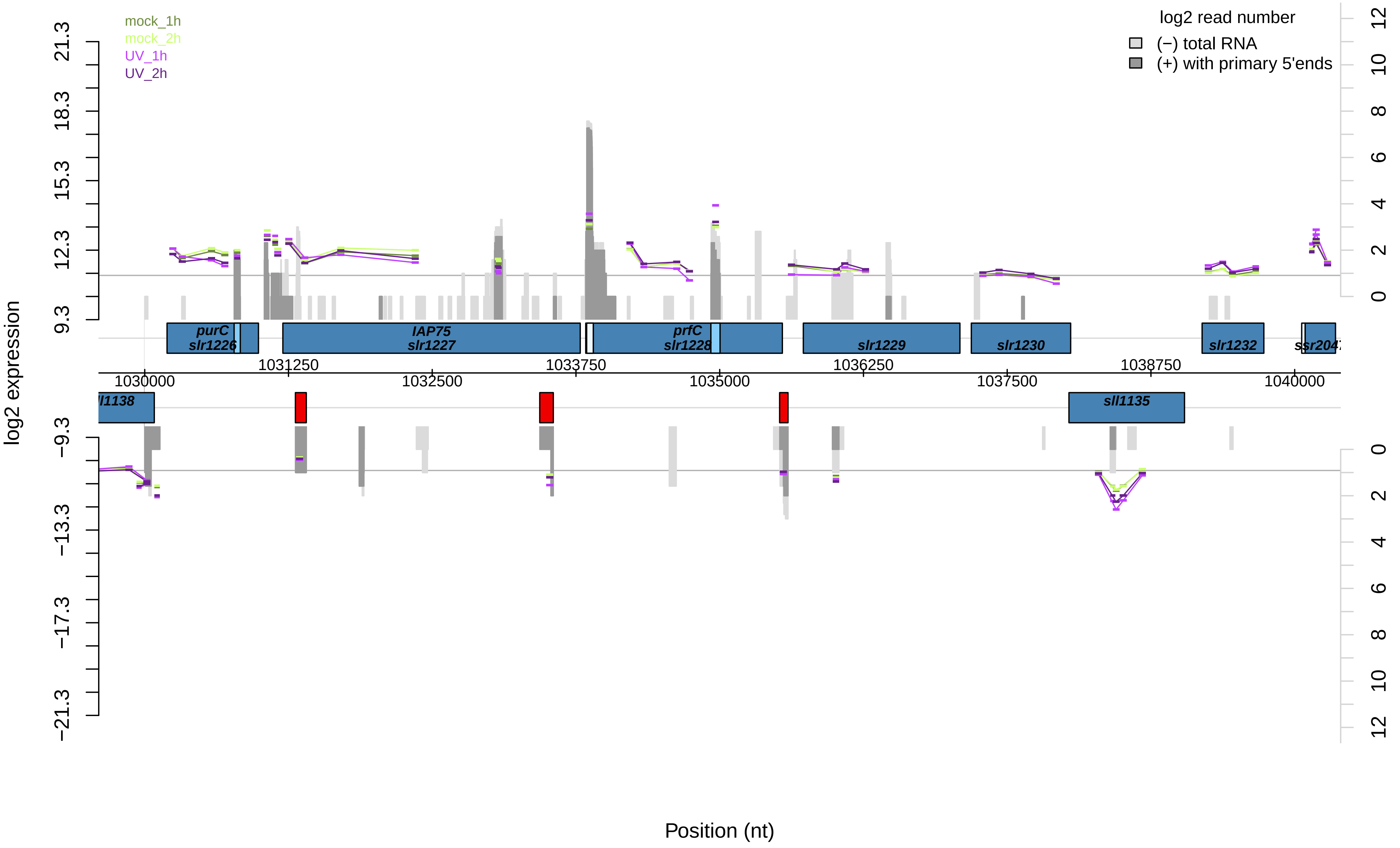

mock_1h
mock_2h
UV_1h
UV_2h
log2 read number
(−) total RNA
(+) with primary 5'ends
log2 expression
21.3
18.3
15.3
12.3
9.3
−9.3
−13.3
−17.3
−21.3
12
10
8
6
4
2
0
purC
slr1226
IAP75
slr1227
prfC
slr1228
slr1229
slr1230
slr1232
ll1138
sll1135
1030000
1031250
1032500
1033750
1035000
1036250
1037500
1038750
1040000
Position (nt)

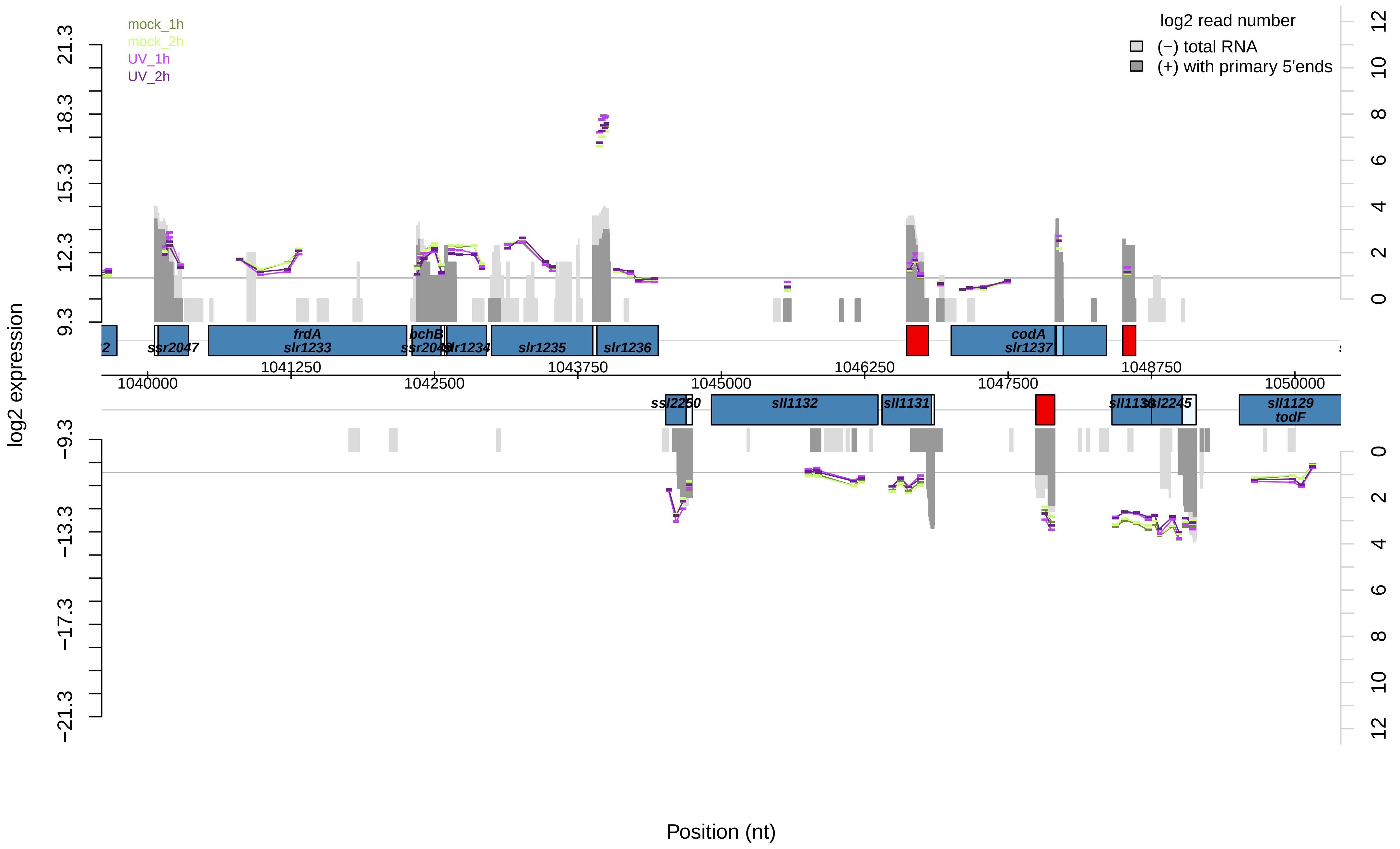
mock_1h
mock_2h
UV_1h
UV_2h
log2 read number
(−) total RNA
(+) with primary 5'ends
log2 expression
21.3
18.3
15.3
12.3
9.3
−9.3
−13.3
−17.3
−21.3
12
10
8
6
4
2
0
frdA
slr1233
ssr2047
bchB
ssr2048
slr1234
slr1235
slr1236
codA
slr1237
1041250
1042500
1043750
1046250
1048750
1040000
1045000
1047500
1050000
ssl2250
sll1132
sll1131
sll1130
ssl2245
sll1129
todF
Position (nt)

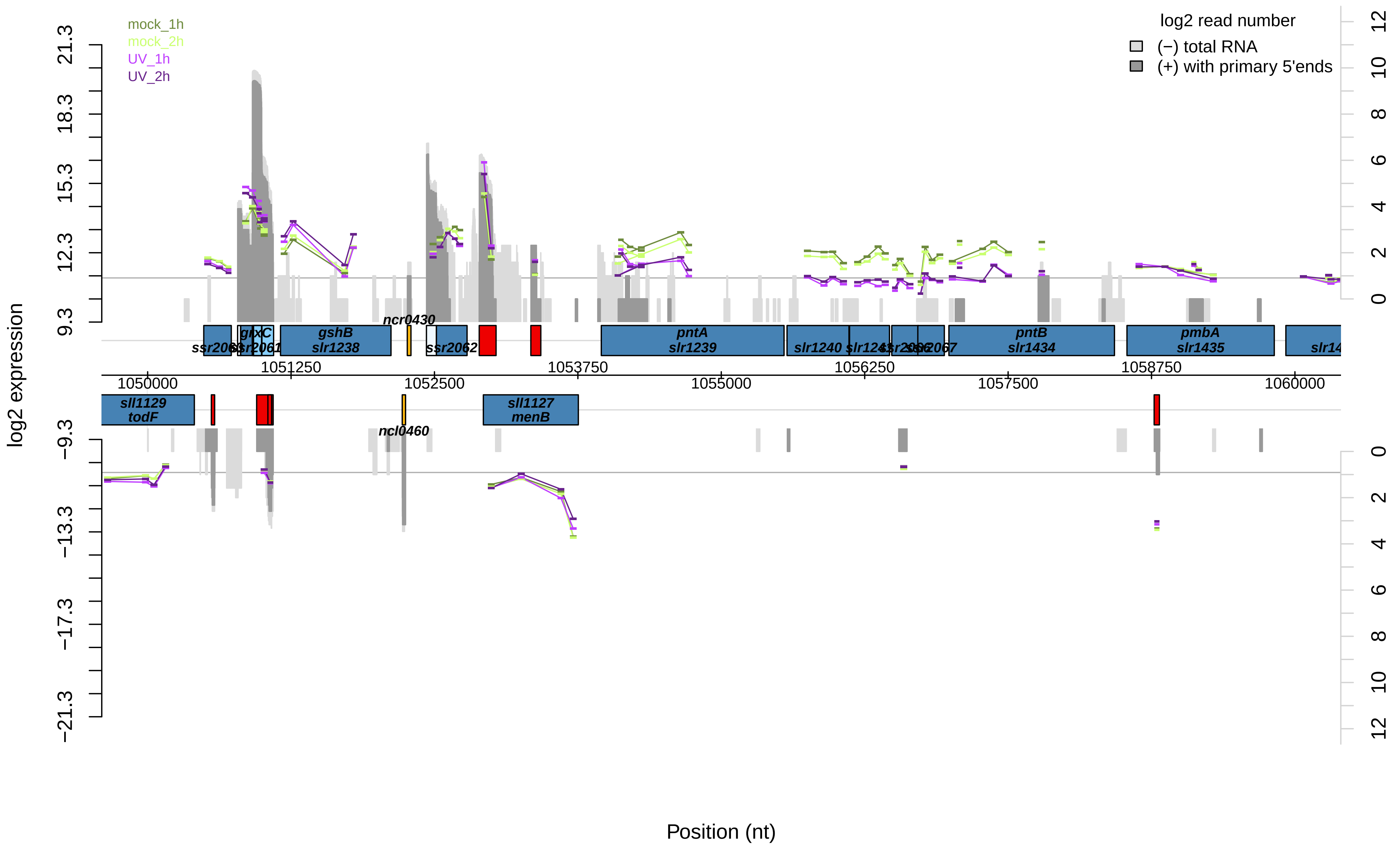

mock_1h
mock_2h
UV_1h
UV_2h
log2 read number
(−) total RNA
(+) with primary 5'ends
log2 expression
Position (nt)
ncr0430
ncl0460
grxC
ssr2061
gshB
slr1238
ssr2062
pntA
slr1239
slr1240
pntB
slr1434
pmbA
slr1435
sll1129
todF
sll1127
menB
1050000
1051250
1052500
1053750
1055000
1056250
1057500
1058750
1060000

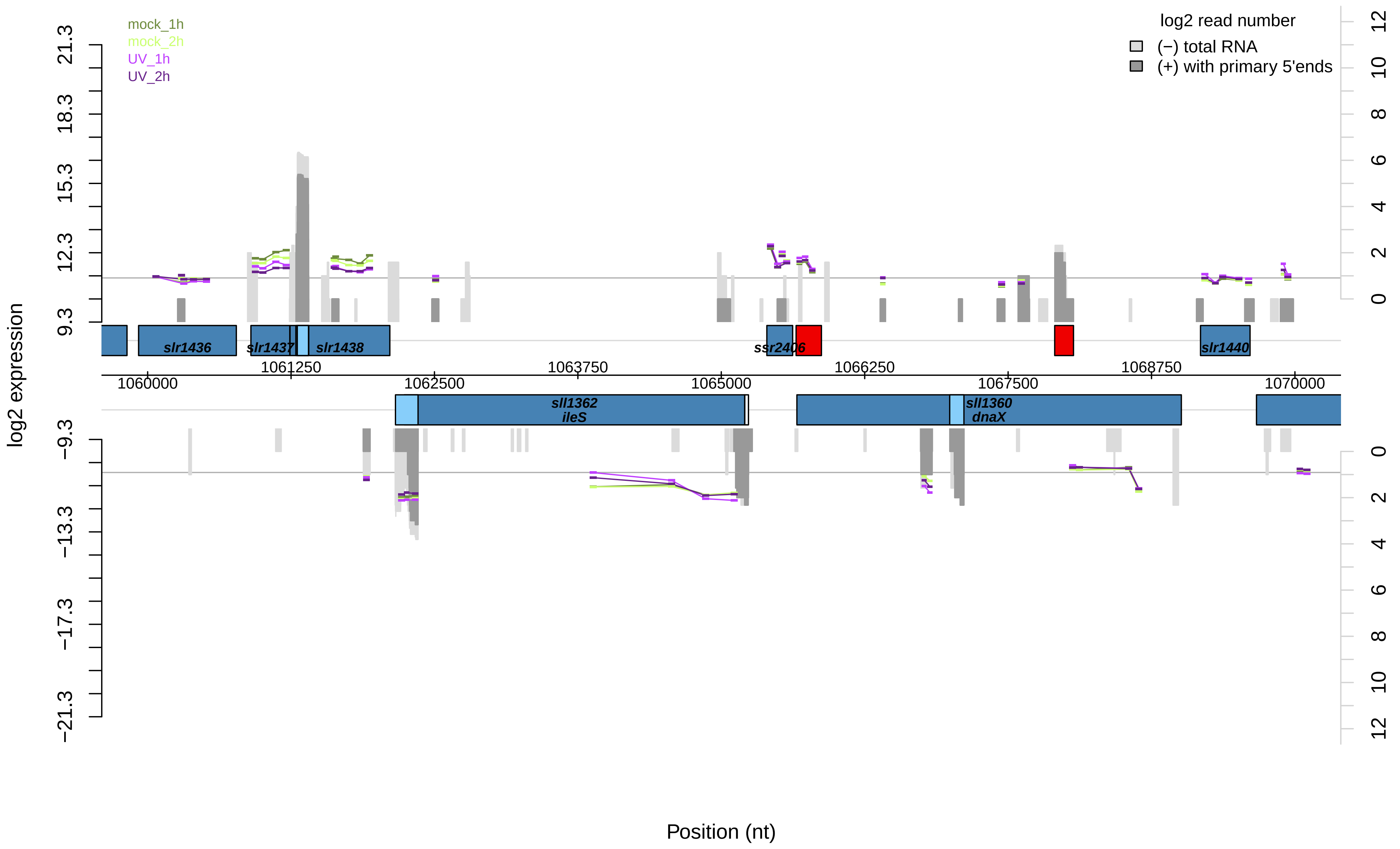

mock_1h
mock_2h
UV_1h
UV_2h
log2 read number
(−) total RNA
(+) with primary 5'ends
log2 expression
21.3
18.3
15.3
12.3
9.3
−9.3
−13.3
−17.3
−21.3
12
10
8
6
4
2
0
slr1436
slr1437
slr1438
ssr2406
slr1440
1060000
1061250
1062500
1063750
1065000
1066250
1067500
1068750
1070000
sll1362
ileS
sll1360
dnaX
Position (nt)

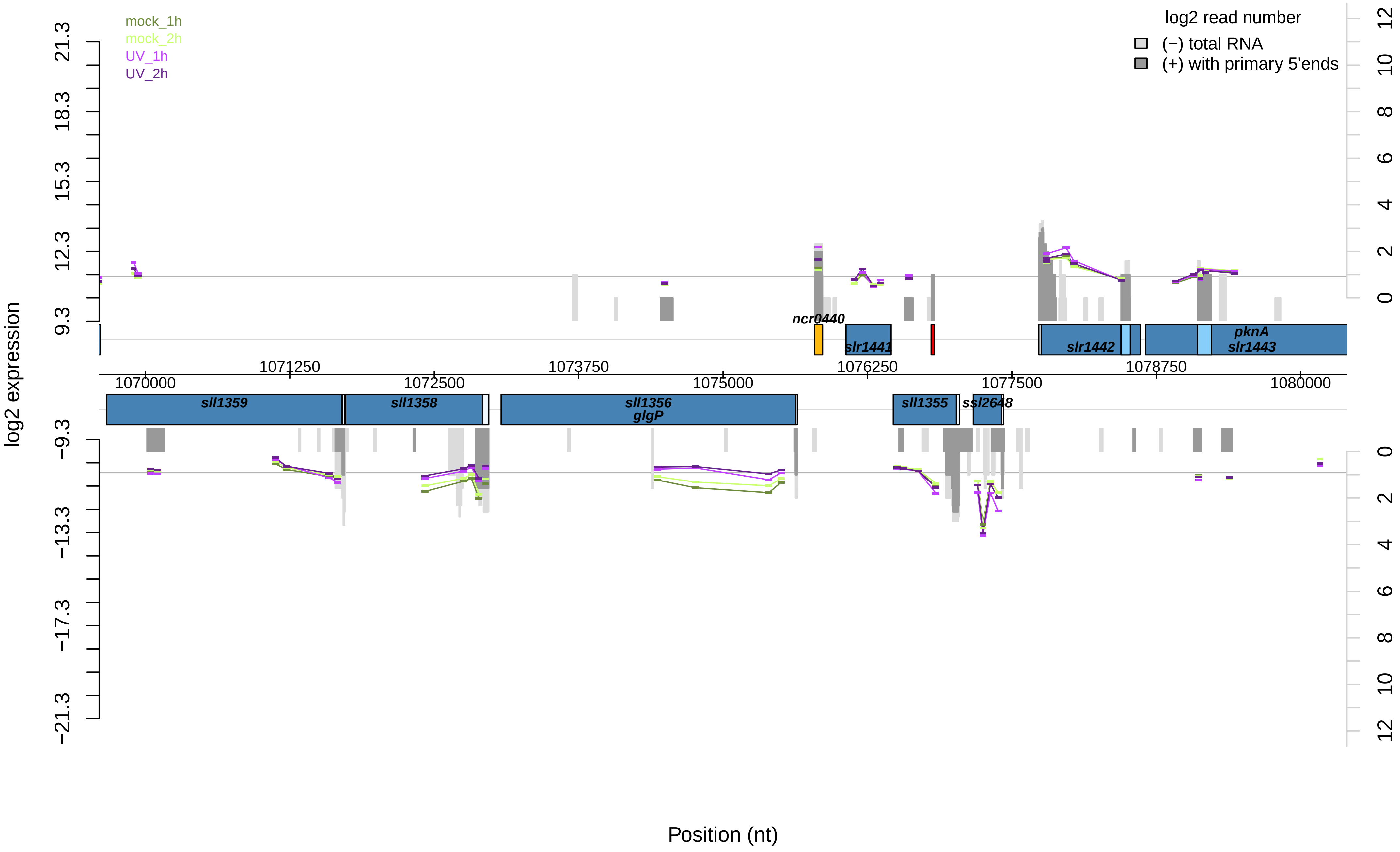

mock_1h
mock_2h
UV_1h
UV_2h
log2 read number
(−) total RNA
(+) with primary 5'ends
log2 expression
21.3
18.3
15.3
12.3
9.3
−9.3
−13.3
−17.3
−21.3
ncr0440
slr1441
slr1442
pknA
slr1443
sll1359
sll1358
sll1356
glgP
sll1355
ssl2648
1070000
1071250
1072500
1073750
1075000
1076250
1077500
1078750
1080000
Position (nt)

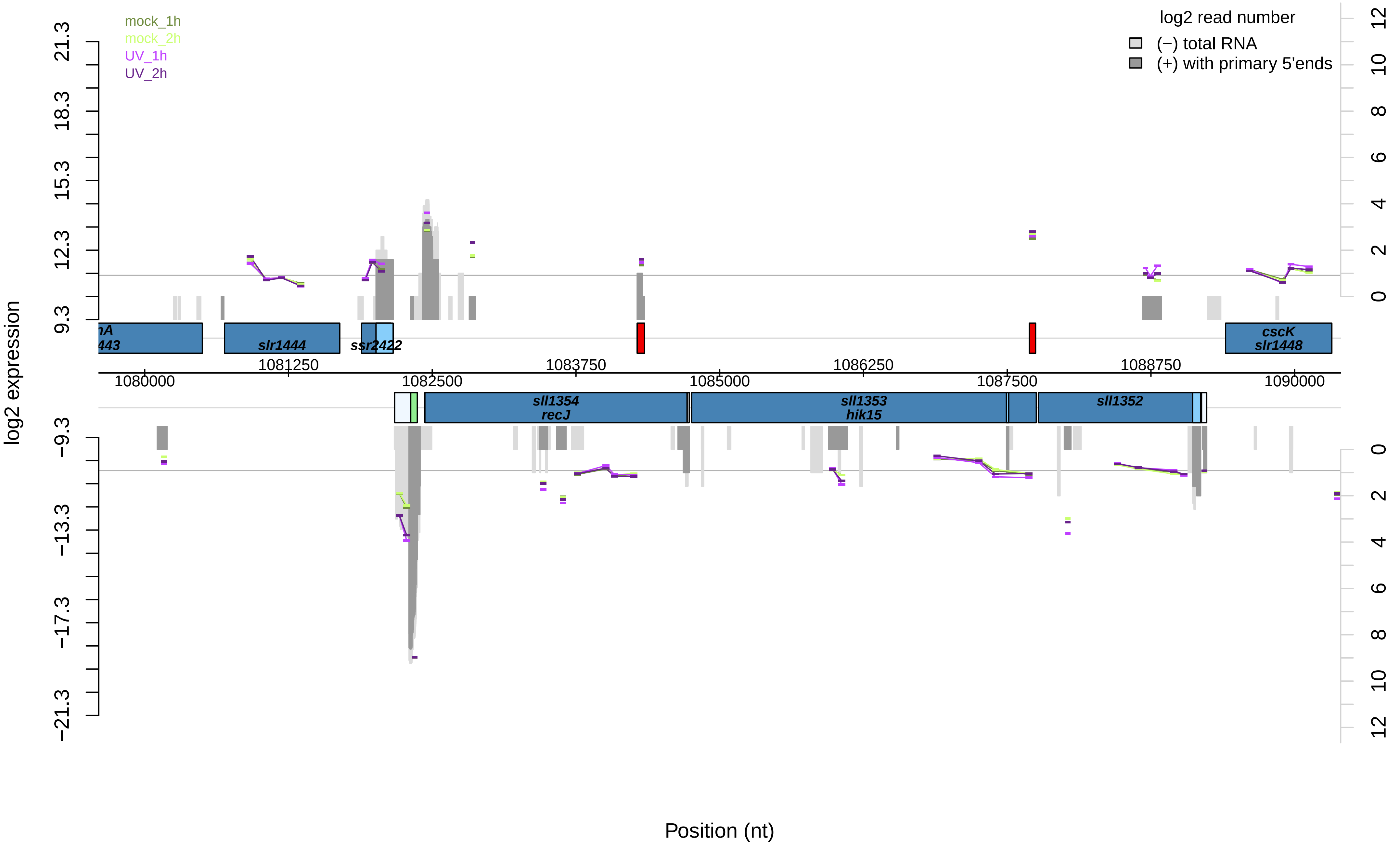

mock_1h
mock_2h
UV_1h
UV_2h
log2 read number
(−) total RNA
(+) with primary 5'ends
log2 expression
Position (nt)
21.3
18.3
15.3
12.3
9.3
−9.3
−13.3
−17.3
−21.3
12
10
8
6
4
2
0
1080000
1081250
1082500
1083750
1085000
1086250
1087500
1088750
1090000
slr1444
ssr2422
cscK
slr1448
sll1354
recJ
sll1353
hik15
sll1352

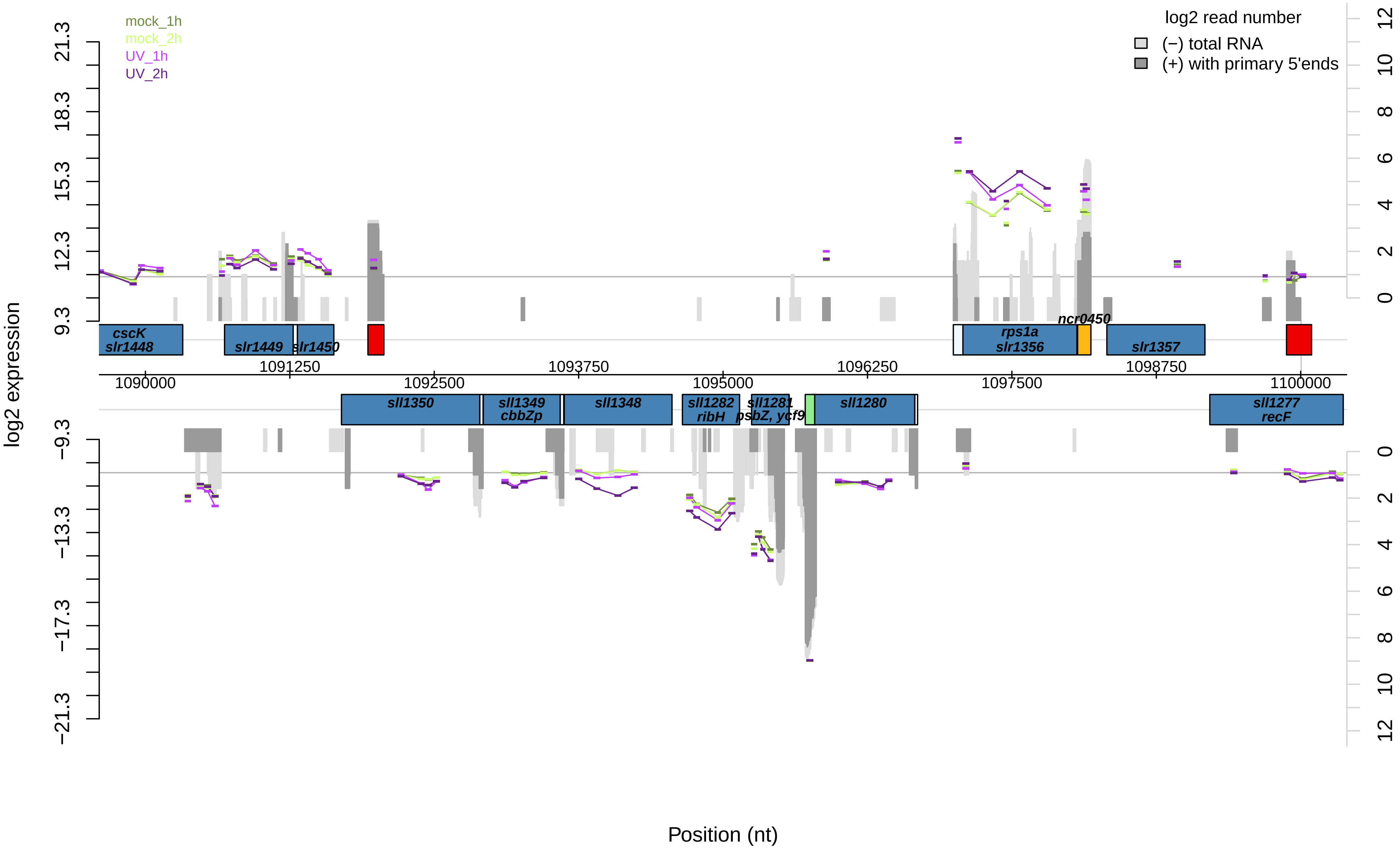

mock_1h
mock_2h
UV_1h
UV_2h
log2 read number
(−) total RNA
(+) with primary 5'ends
log2 expression
21.3
18.3
15.3
12.3
9.3
−9.3
−13.3
−17.3
−21.3
0
2
4
6
8
10
12
cscK
slr1448
slr1449
slr1450
rps1a
slr1356
ncr0450
slr1357
sll1350
sll1349
cbbZp
sll1348
sll1282
ribH
sll1281
psbZ, ycf9
sll1280
sll1277
recF
1090000
1091250
1092500
1093750
1095000
1096250
1097500
1098750
1100000
Position (nt)

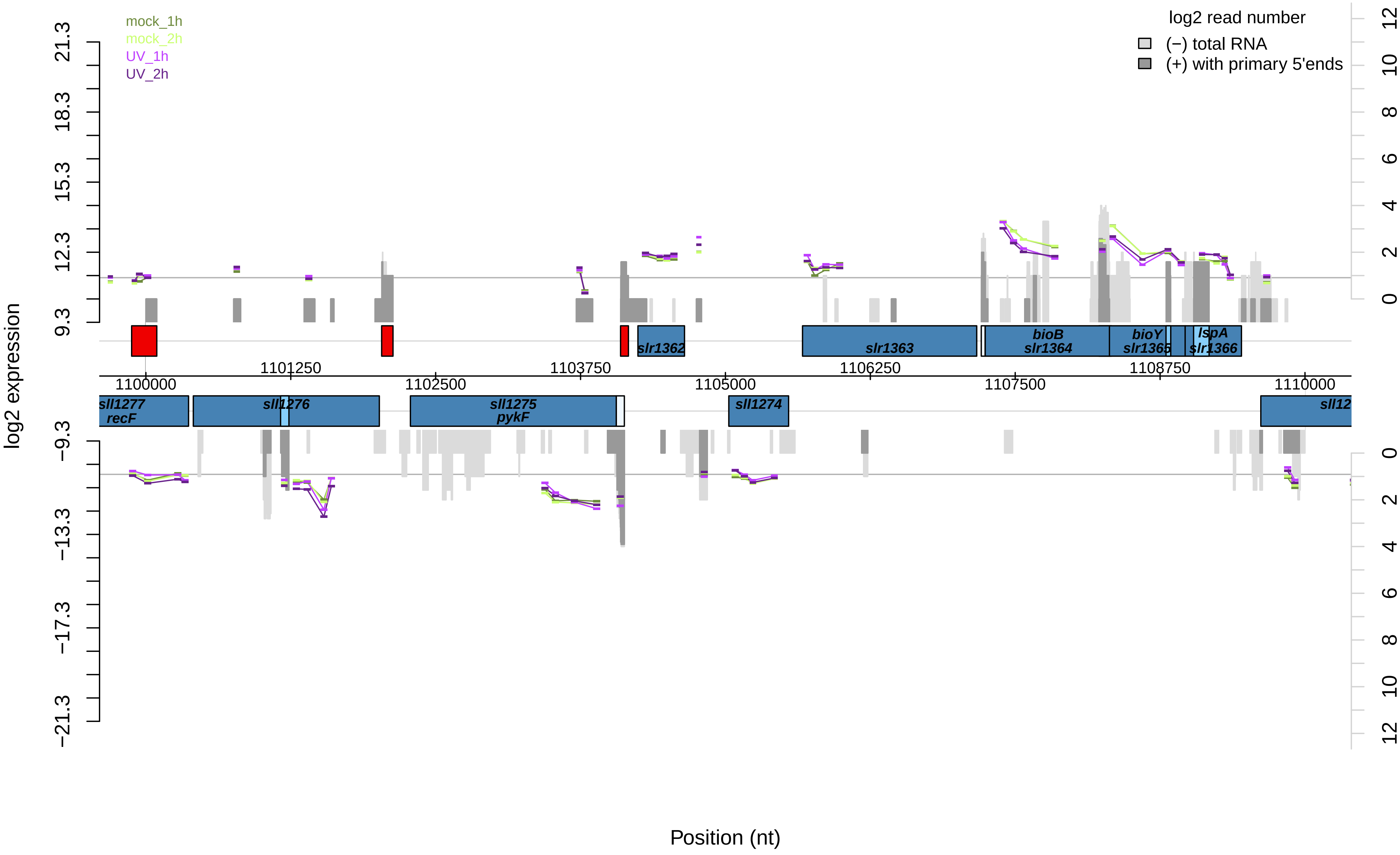

mock_1h
mock_2h
UV_1h
UV_2h
log2 read number
(−) total RNA
(+) with primary 5'ends
log2 expression
21.3
18.3
15.3
12.3
9.3
−9.3
−13.3
−17.3
−21.3
12
10
8
6
4
2
0
recF
sll1277
sll1276
sll1275
pykF
sll1274
sll12
slr1362
slr1363
bioB
slr1364
bioY
slr1365
lspA
slr1366
1100000
1101250
1102500
1103750
1105000
1106250
1107500
1108750
1110000
Position (nt)

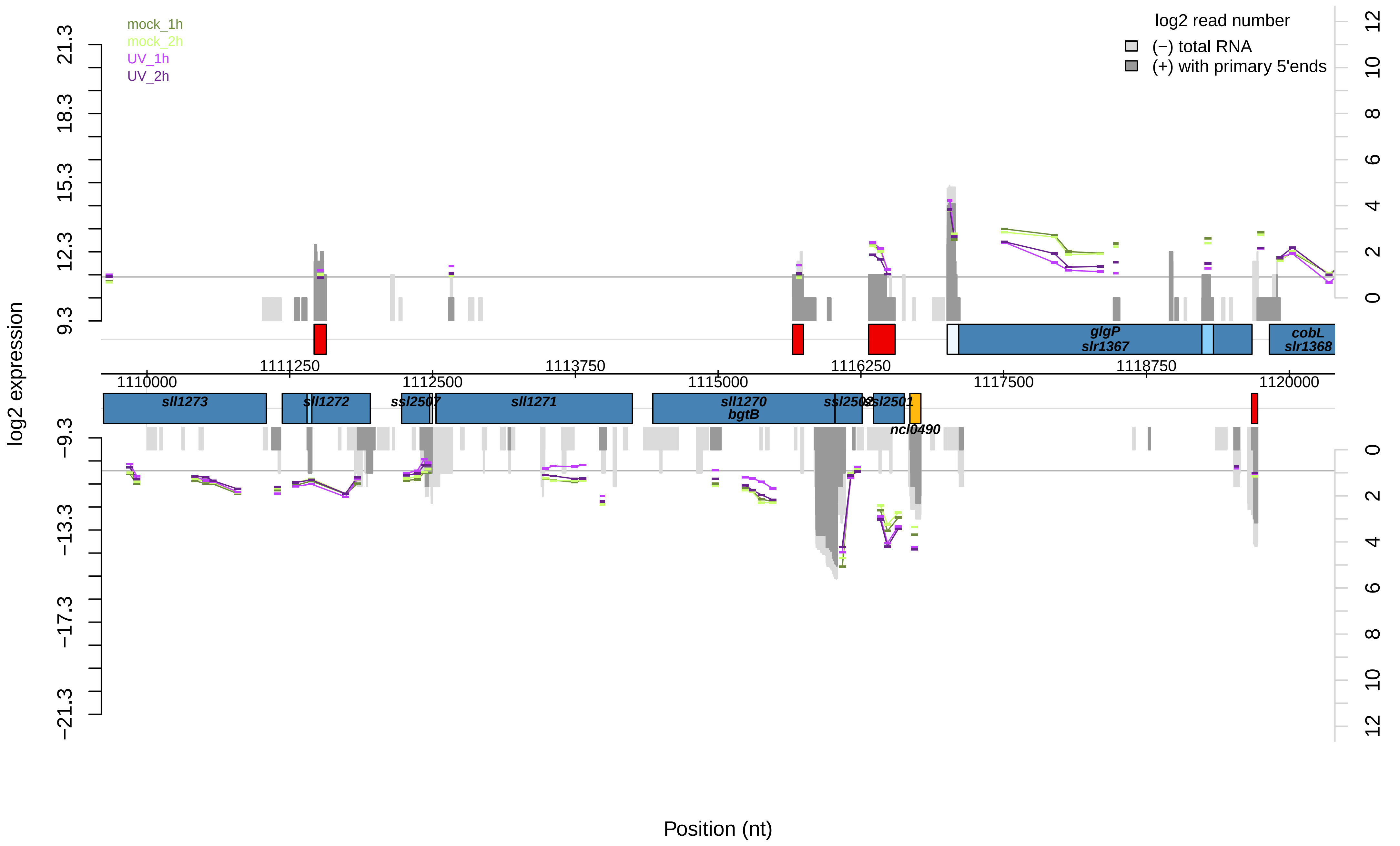

mock_1h
mock_2h
UV_1h
UV_2h
log2 read number
(−) total RNA
(+) with primary 5'ends
log2 expression
21.3
18.3
15.3
12.3
9.3
−9.3
−13.3
−17.3
−21.3
12
10
8
6
4
2
0
glgP
slr1367
cobL
slr1368
1110000
1111250
1112500
1113750
1115000
1116250
1117500
1118750
1120000
sll1273
sll1272
ssl2507
sll1271
sll1270
bgtB
ssl2502
ssl2501
ncl0490
Position (nt)

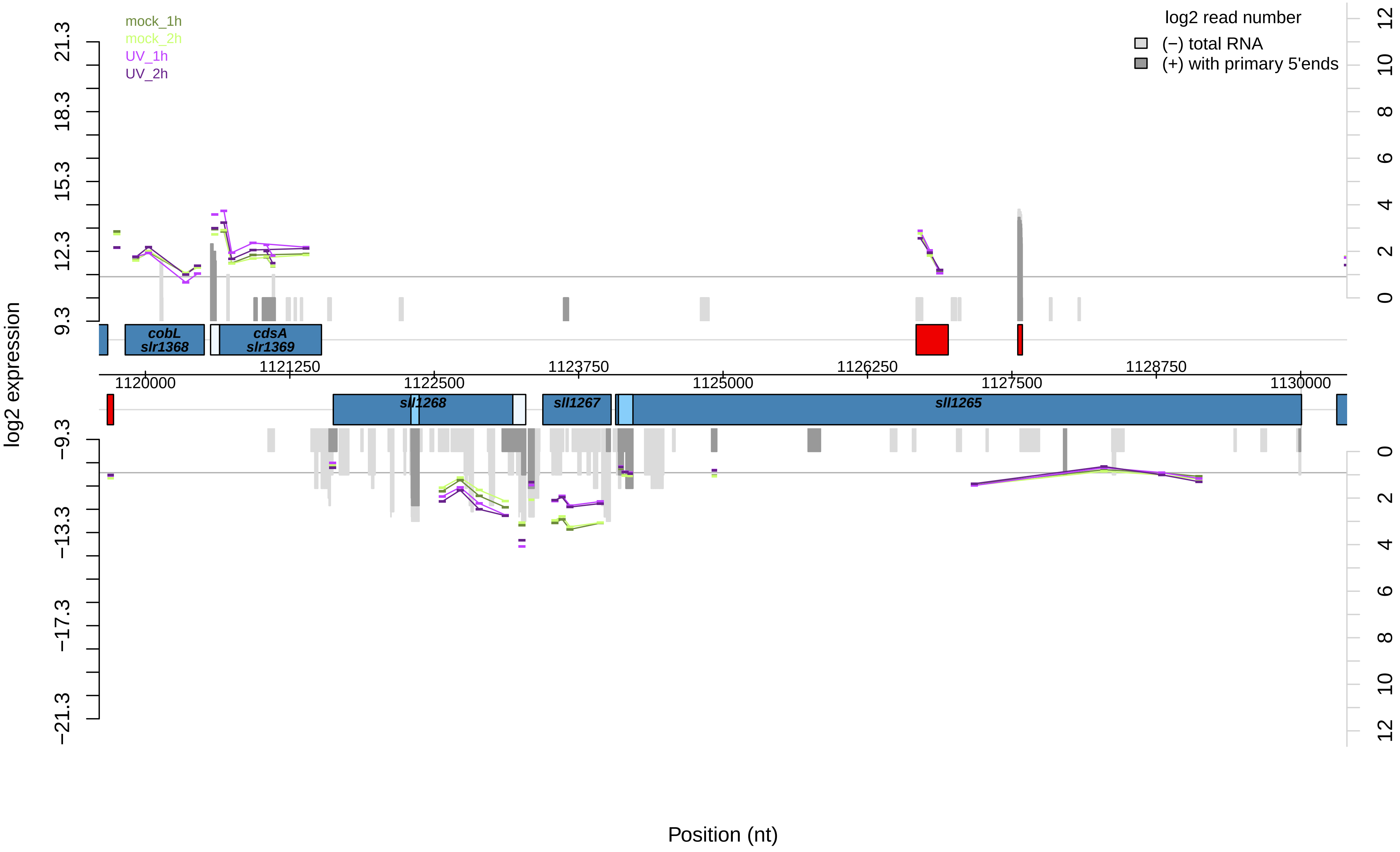

mock_1h
mock_2h
UV_1h
UV_2h
log2 read number
(−) total RNA
(+) with primary 5'ends
log2 expression
21.3
18.3
15.3
12.3
9.3
−9.3
−13.3
−17.3
−21.3
12
10
8
6
4
2
0
cobL
slr1368
cdsA
slr1369
1121250
1123750
1126250
1128750
1120000
1122500
1125000
1127500
1130000
sll1268
sll1267
sll1265
Position (nt)

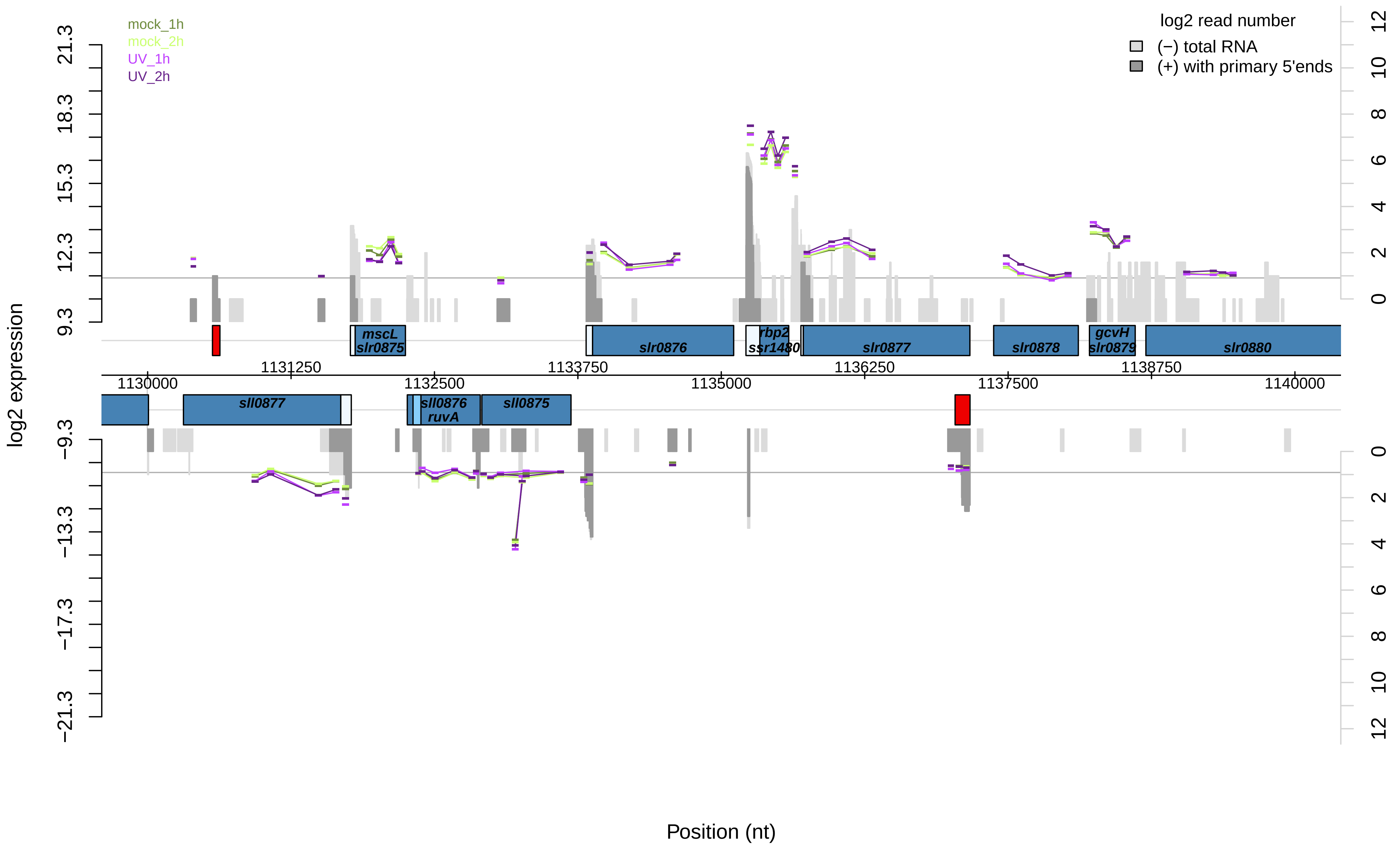

mock_1h
mock_2h
UV_1h
UV_2h
log2 read number
(−) total RNA
(+) with primary 5'ends
log2 expression
21.3
18.3
15.3
12.3
9.3
−9.3
−13.3
−17.3
−21.3
12
10
8
6
4
2
0
mscL
slr0875
slr0876
rbp2
ssr1480
slr0877
slr0878
gcvH
slr0879
slr0880
1131250
1133750
1136250
1138750
1130000
1132500
1135000
1137500
1140000
sll0877
sll0876
ruvA
sll0875
Position (nt)

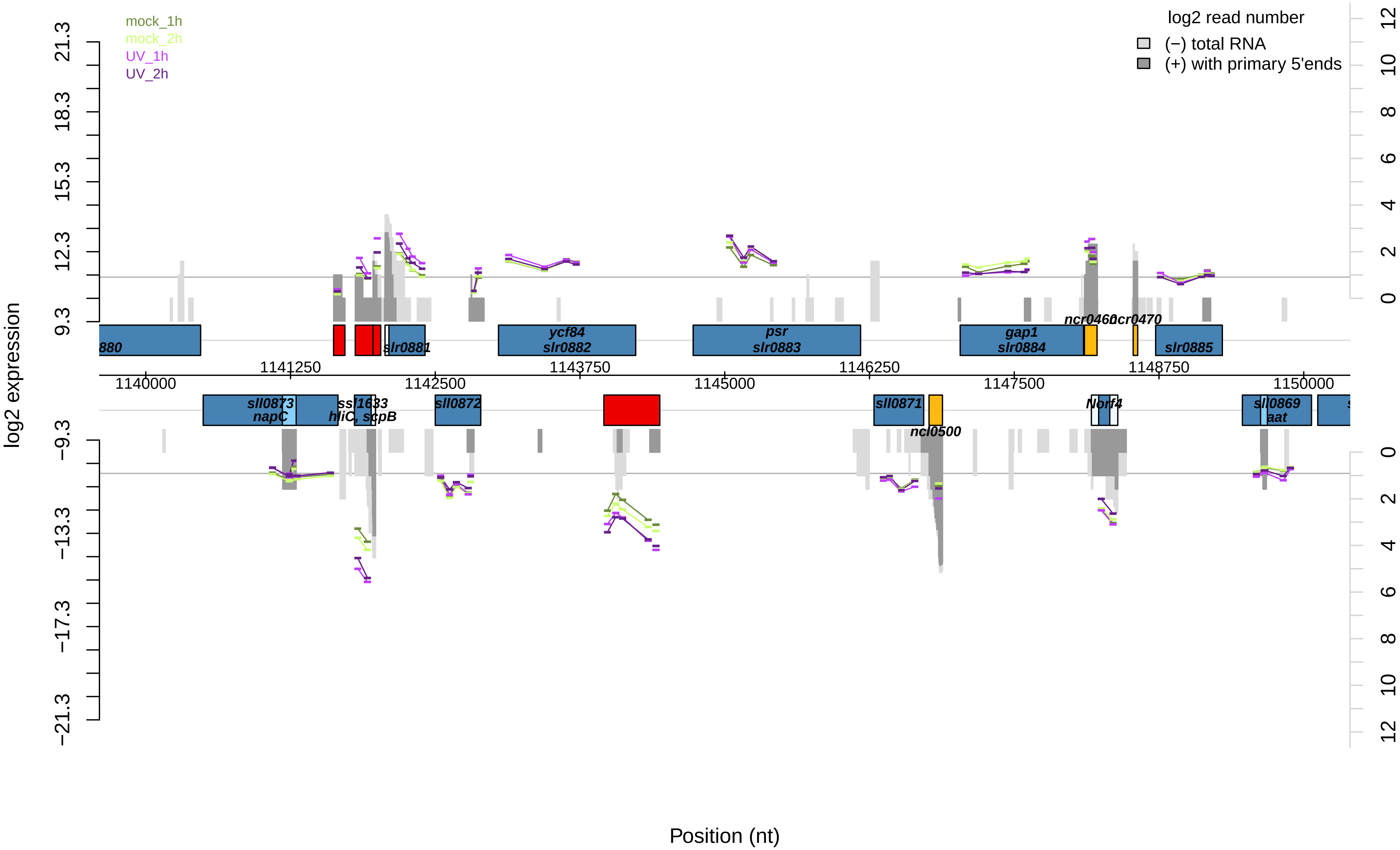

mock_1h
mock_2h
UV_1h
UV_2h
log2 read number
(−) total RNA
(+) with primary 5'ends
log2 expression
21.3
18.3
15.3
12.3
9.3
−9.3
−13.3
−17.3
−21.3
12
10
8
6
4
2
0
880
ycf84
slr0882
psr
slr0883
gap1
slr0884
ncr0460
ncr0470
slr0885
slr0881
1141250
1142500
1143750
1145000
1146250
1147500
1148750
1140000
1150000
sll0873
napC
ssl1633
hliC, scpB
sll0872
sll0871
ncl0500
Norf4
sll0869
aat
Position (nt)

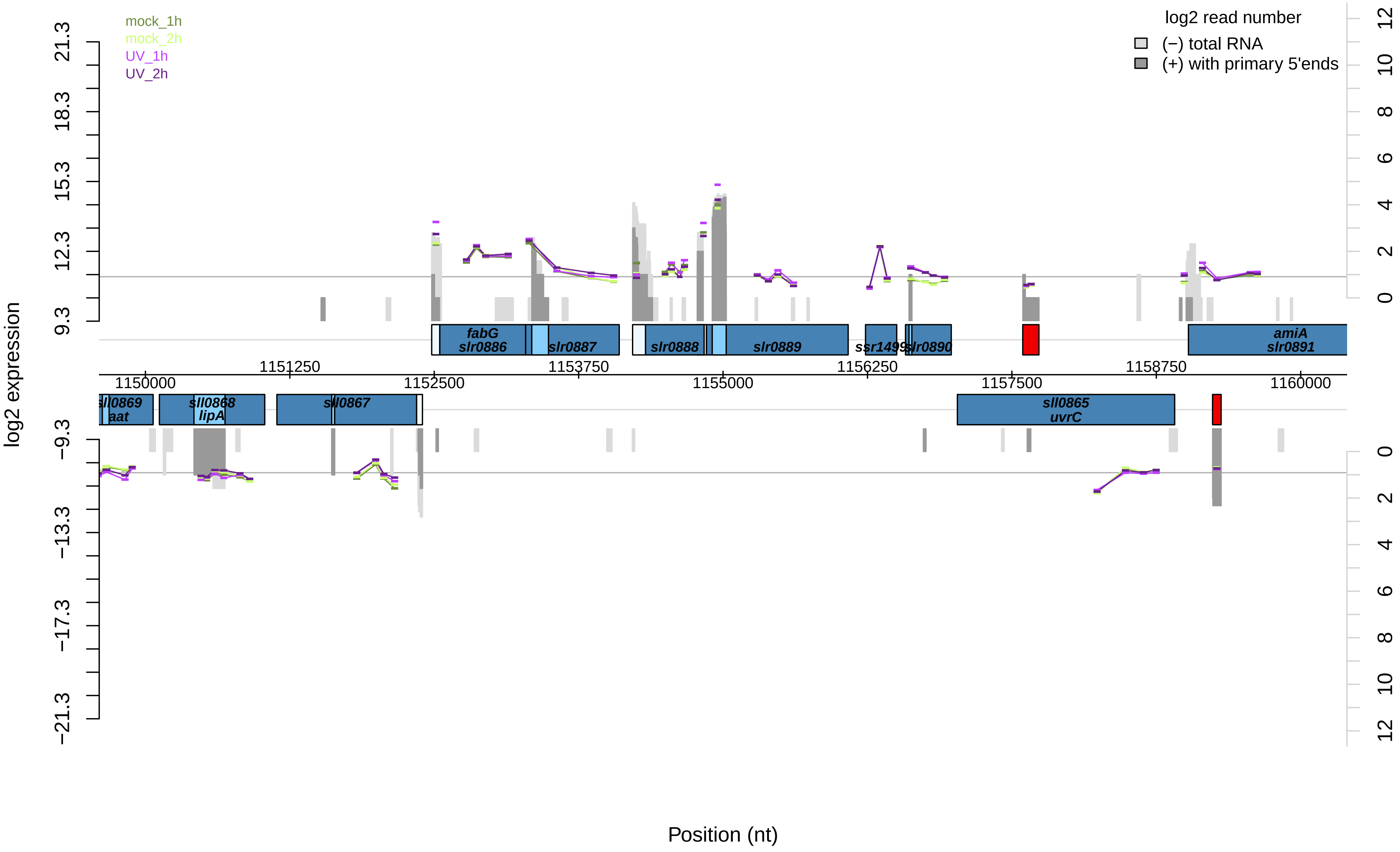
mock_1h
mock_2h
UV_1h
UV_2h
log2 read number
(−) total RNA
(+) with primary 5'ends
log2 expression
21.3
18.3
15.3
12.3
9.3
−9.3
−13.3
−17.3
−21.3
12
10
8
6
4
2
0
fabG
slr0886
slr0887
slr0888
slr0889
ssr1499
slr0890
amiA
slr0891
1151250
1153750
1156250
1158750
1150000
1152500
1155000
1157500
1160000
sll0869
aat
sll0868
lipA
sll0867
sll0865
uvrC
Position (nt)

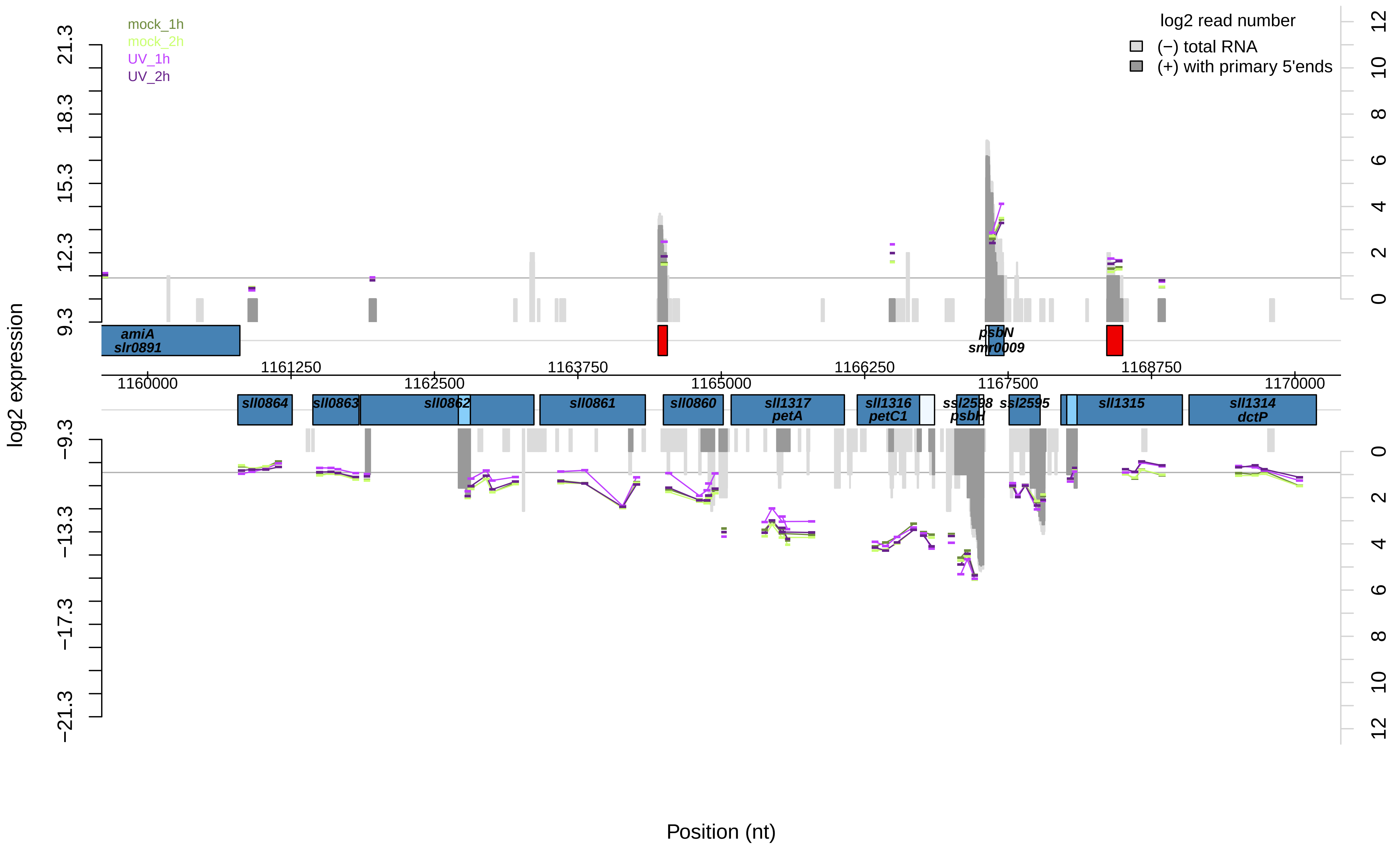

mock_1h
mock_2h
UV_1h
UV_2h
log2 read number
(−) total RNA
(+) with primary 5'ends
log2 expression
21.3
18.3
15.3
12.3
9.3
−9.3
−13.3
−17.3
−21.3
12
10
8
6
4
2
0
amiA
slr0891
psbN
smr0009
1161250
1163750
1166250
1168750
1160000
1162500
1165000
1167500
1170000
sll0864
sll0863
sll0862
sll0861
sll0860
sll1317
petA
sll1316
petC1
ssl2598
psbH
ssl2595
sll1315
sll1314
dctP
Position (nt)

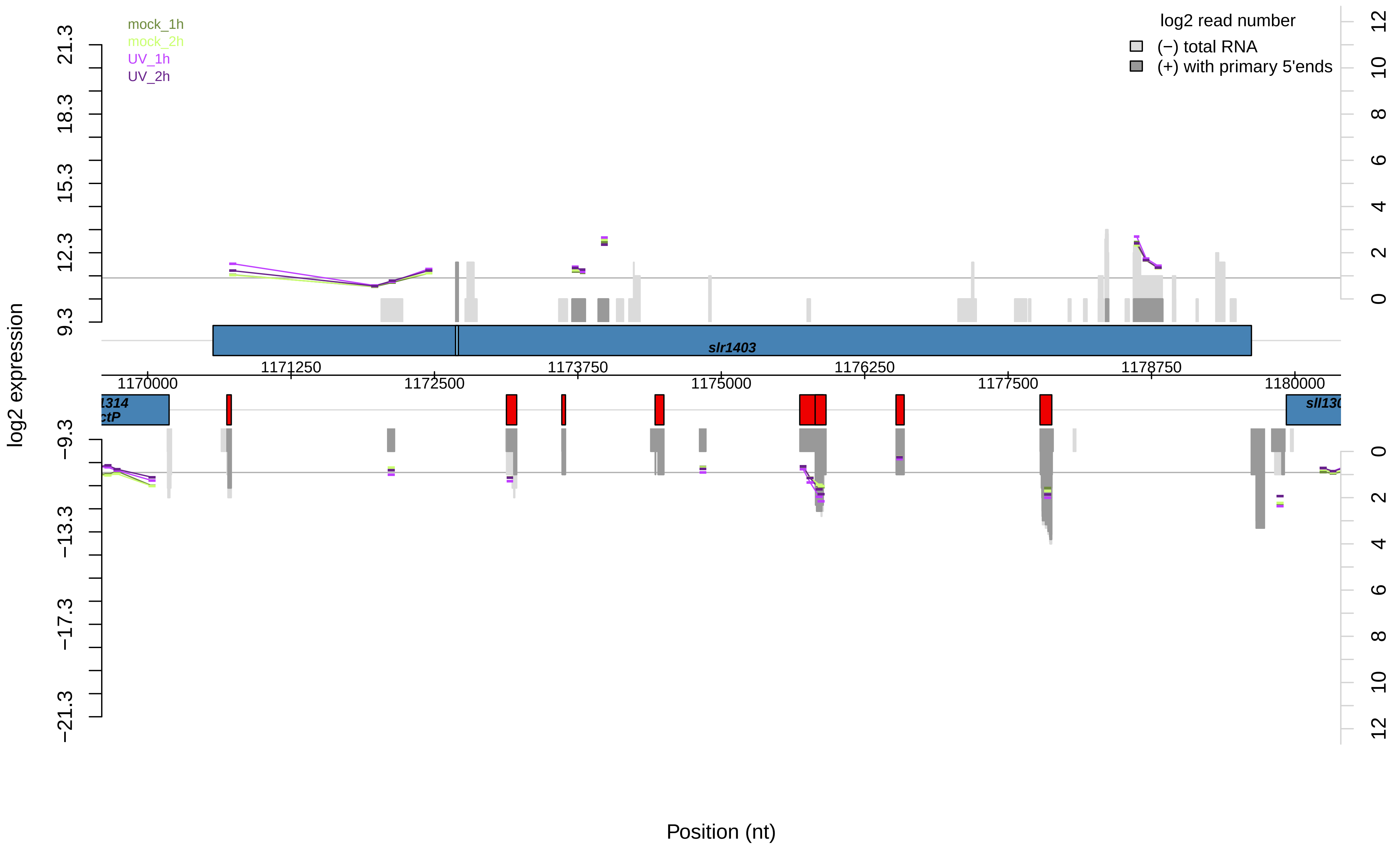

mock_1h
mock_2h
UV_1h
UV_2h
log2 read number
(−) total RNA
(+) with primary 5'ends
log2 expression
21.3
18.3
15.3
12.3
9.3
−9.3
−13.3
−17.3
−21.3
slr1403
1171250
1173750
1176250
1178750
1170000
1172500
1175000
1177500
1180000
ctP
Position (nt)

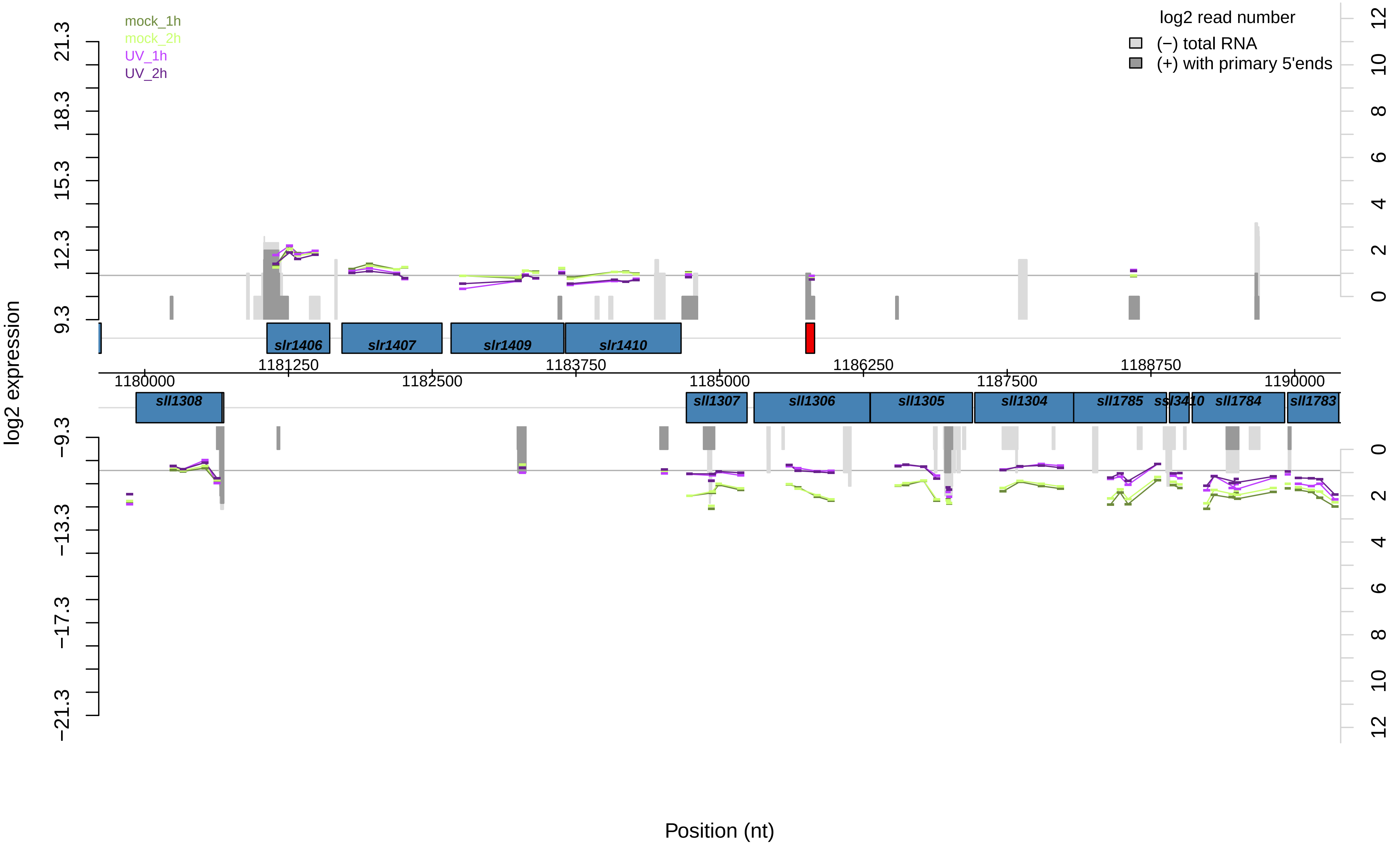
mock_1h
mock_2h
UV_1h
UV_2h
log2 read number
(−) total RNA
(+) with primary 5'ends
log2 expression
21.3
18.3
15.3
12.3
9.3
−9.3
−13.3
−17.3
−21.3
12
10
8
6
4
2
0
slr1406
slr1407
slr1409
slr1410
1181250
1183750
1186250
1188750
1180000
1182500
1185000
1187500
1190000
sll1308
sll1307
sll1306
sll1305
sll1304
sll1785
ssl3410
sll1784
sll1783
Position (nt)

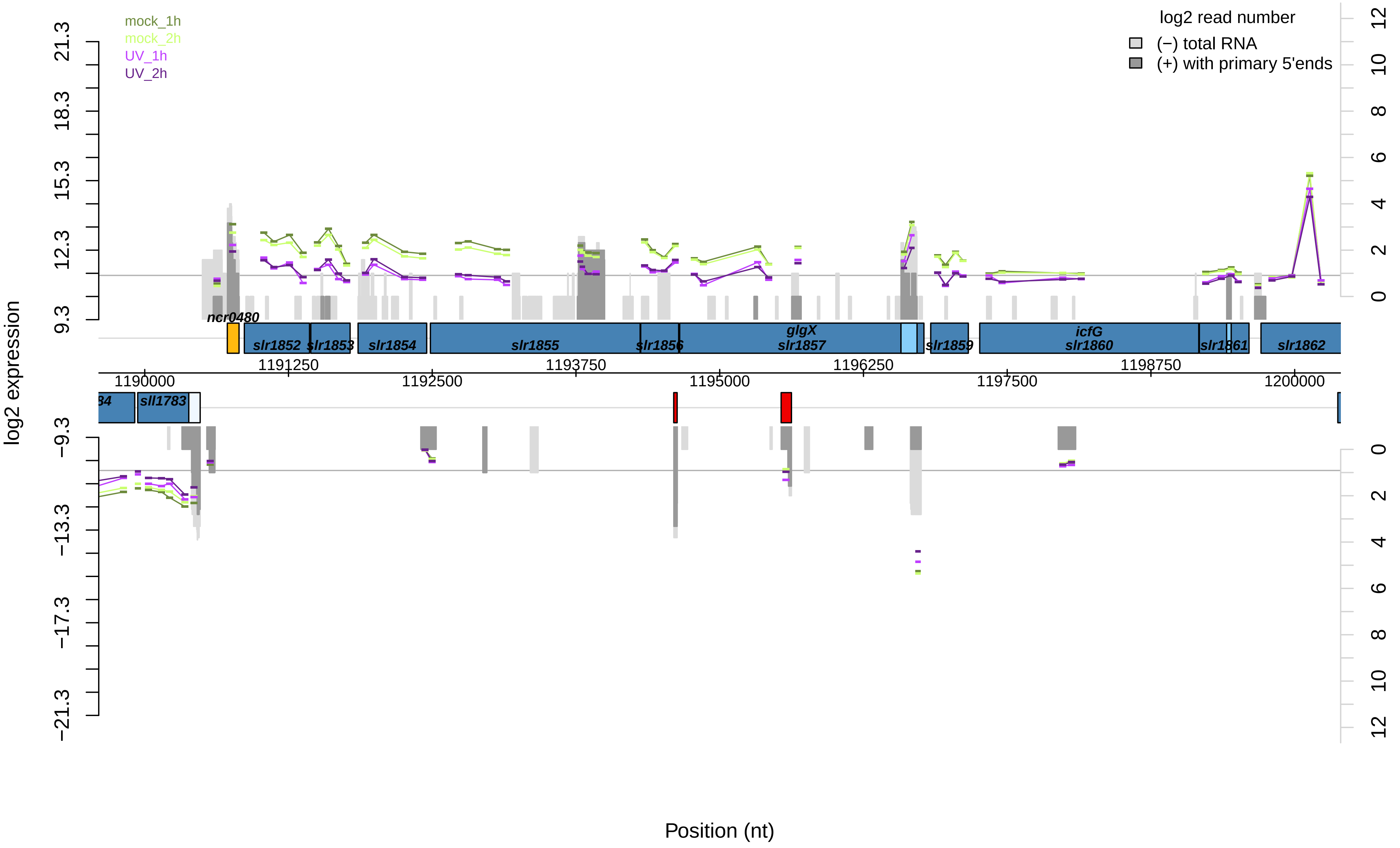

mock_1h
mock_2h
UV_1h
UV_2h
log2 read number
(−) total RNA
(+) with primary 5'ends
log2 expression
21.3
18.3
15.3
12.3
9.3
−9.3
−13.3
−17.3
−21.3
ncr0480
slr1852
slr1853
slr1854
slr1855
slr1856
glgX
slr1857
slr1859
icfG
slr1860
slr1861
slr1862
sll1783
1191250
1193750
1196250
1198750
1190000
1192500
1195000
1197500
1200000
Position (nt)

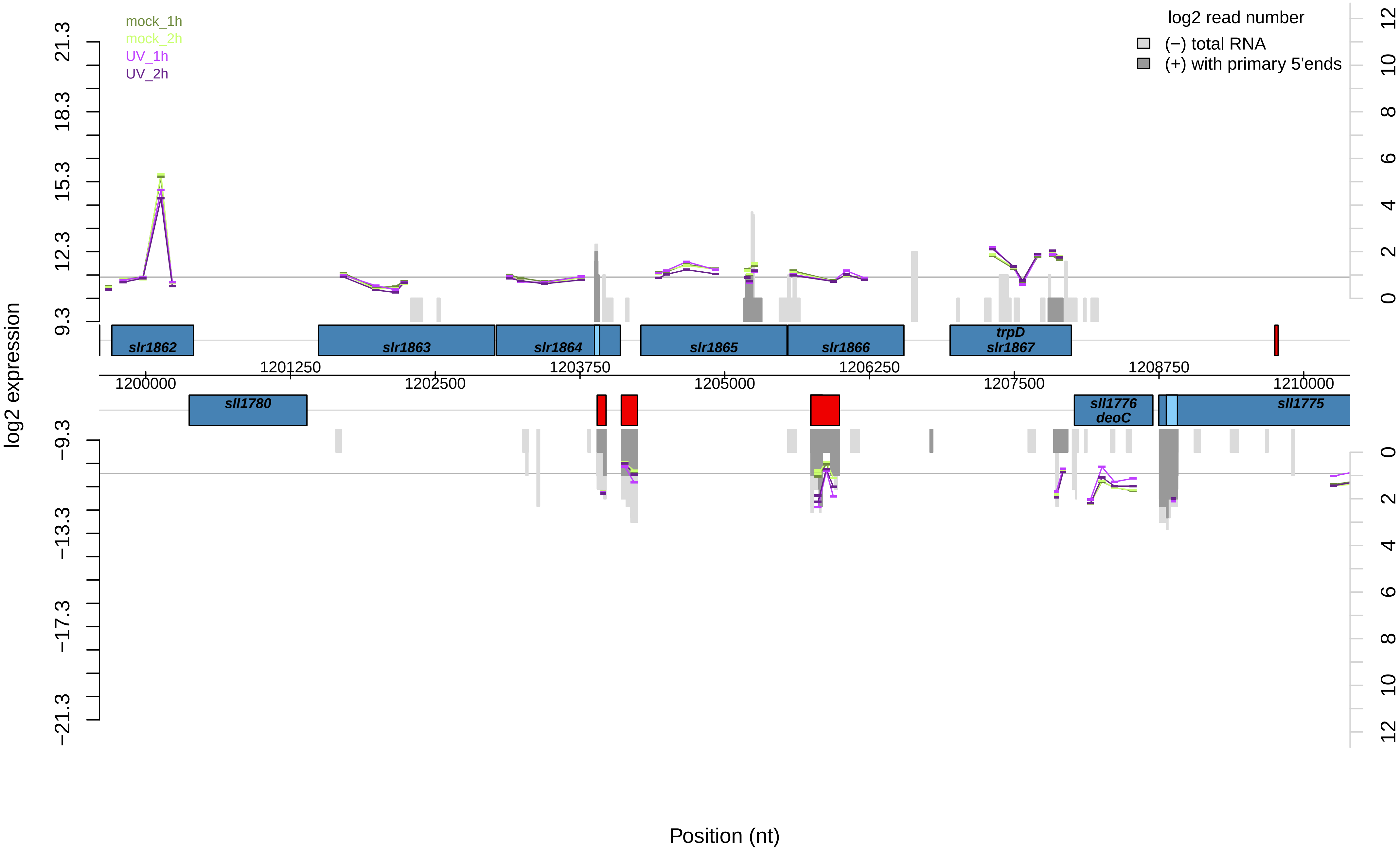

mock_1h
mock_2h
UV_1h
UV_2h
log2 read number
(−) total RNA
(+) with primary 5'ends
log2 expression
21.3
18.3
15.3
12.3
9.3
−9.3
−13.3
−17.3
−21.3
slr1862
slr1863
slr1864
slr1865
slr1866
trpD
slr1867
sll1780
sll1776
deoC
sll1775
1200000
1201250
1202500
1203750
1205000
1206250
1207500
1208750
1210000
Position (nt)

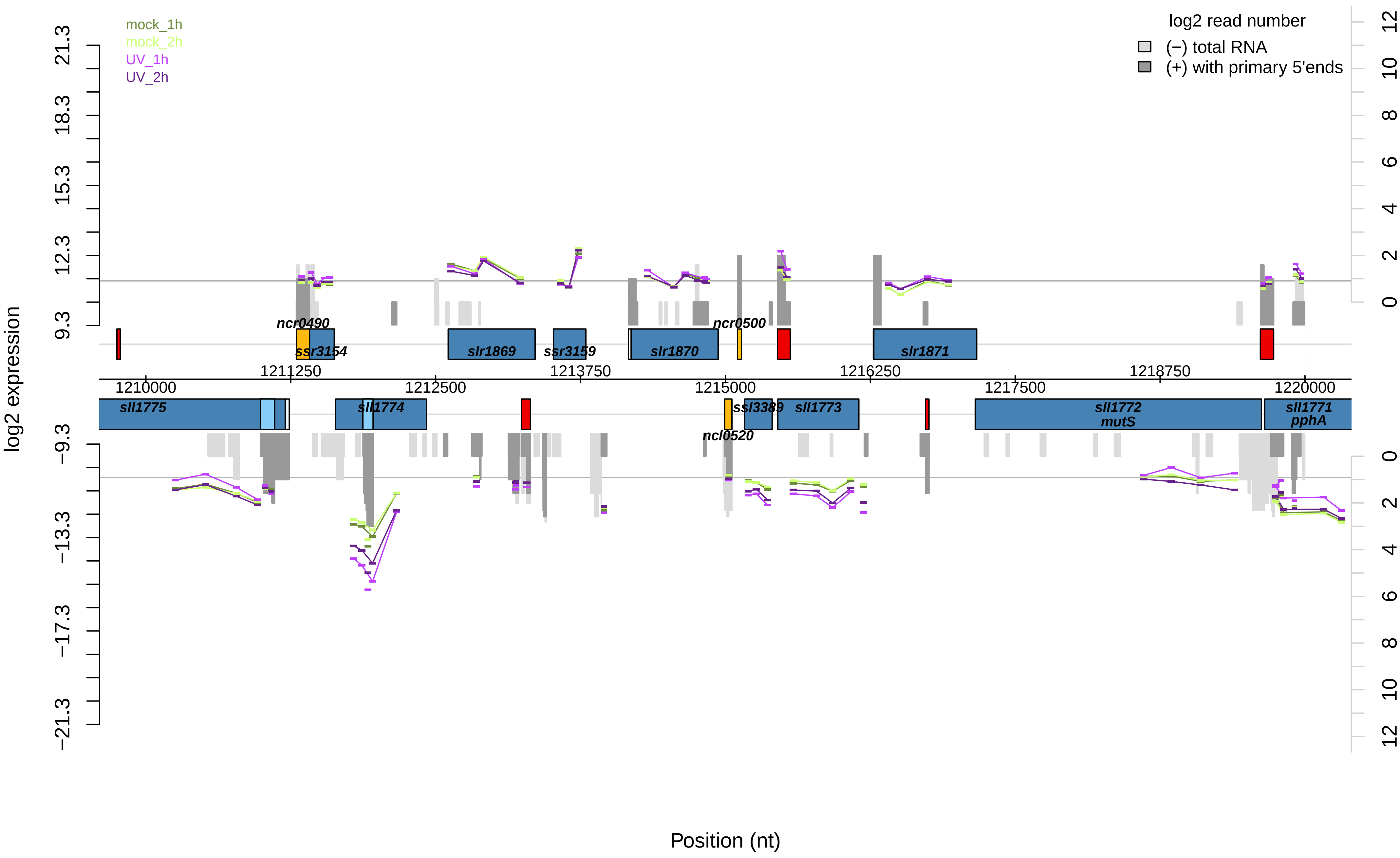

mock_1h
mock_2h
UV_1h
UV_2h
log2 read number
(−) total RNA
(+) with primary 5'ends
log2 expression
21.3
18.3
15.3
12.3
9.3
−9.3
−13.3
−17.3
−21.3
0
2
4
6
8
10
12
ncr0490
ssr3154
slr1869
ssr3159
slr1870
ncr0500
slr1871
1211250
1213750
1216250
1218750
1210000
1212500
1215000
1217500
1220000
sll1775
sll1774
ssl3389
sll1773
sll1772
mutS
sll1771
pphA
ncl0520
Position (nt)

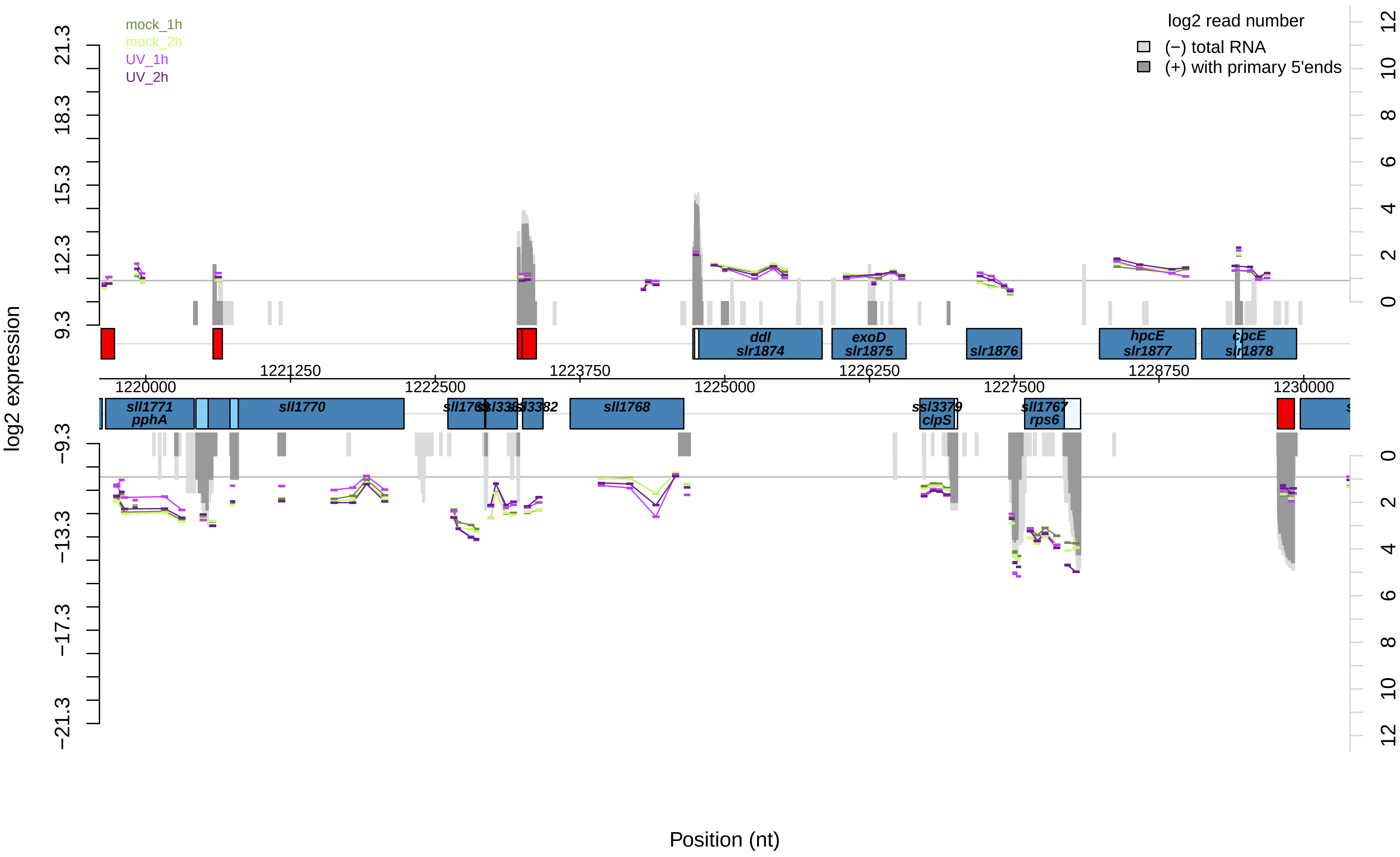

mock_1h
mock_2h
UV_1h
UV_2h
log2 read number
(−) total RNA
(+) with primary 5'ends
log2 expression
21.3
18.3
15.3
12.3
9.3
−9.3
−13.3
−17.3
−21.3
12
10
8
6
4
2
0
ddl
slr1874
exoD
slr1875
slr1876
hpcE
slr1877
cpcE
slr1878
1221250
1222500
1223750
1225000
1226250
1227500
1228750
1230000
1220000
sll1771
pphA
sll1770
sll1768
ssl3379
clpS
sll1767
rps6
Position (nt)

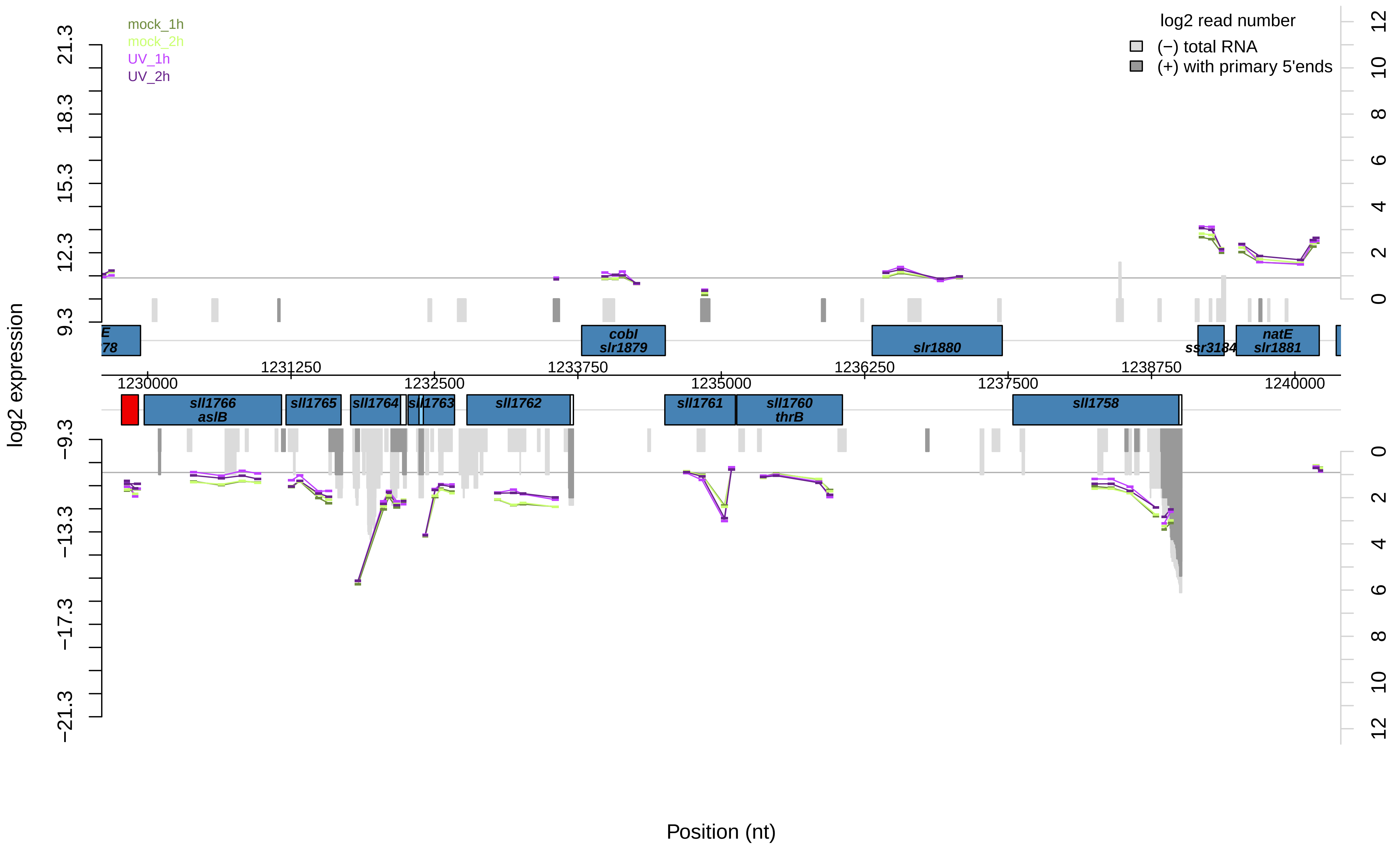

mock_1h
mock_2h
UV_1h
UV_2h
log2 read number
(−) total RNA
(+) with primary 5'ends
log2 expression
21.3
18.3
15.3
12.3
9.3
−9.3
−13.3
−17.3
−21.3
12
10
8
6
4
2
0
cobI
slr1879
slr1880
ssr3184
natE
slr1881
1231250
1233750
1236250
1238750
1230000
1232500
1235000
1237500
1240000
sll1766
aslB
sll1765
sll1764
sll1763
sll1762
sll1761
sll1760
thrB
sll1758
Position (nt)

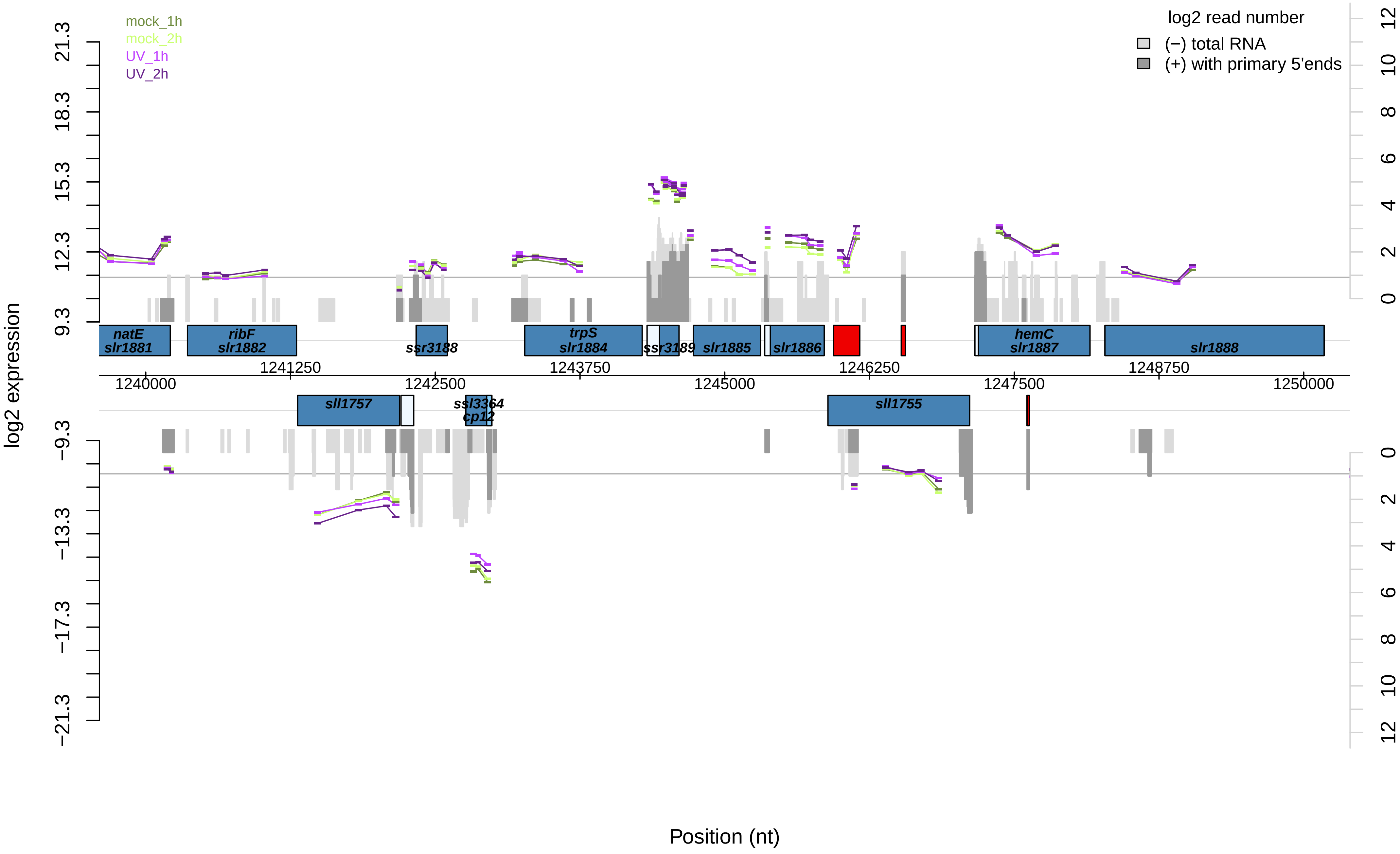

mock_1h
mock_2h
UV_1h
UV_2h
log2 read number
(−) total RNA
(+) with primary 5'ends
log2 expression
21.3
18.3
15.3
12.3
9.3
−9.3
−13.3
−17.3
−21.3
12
10
8
6
4
2
0
natE
slr1881
ribF
slr1882
ssr3188
trpS
slr1884
ssr3189
slr1885
slr1886
hemC
slr1887
slr1888
1240000
1241250
1242500
1243750
1245000
1246250
1247500
1248750
1250000
sll1757
ssl3364
cp12
sll1755
Position (nt)

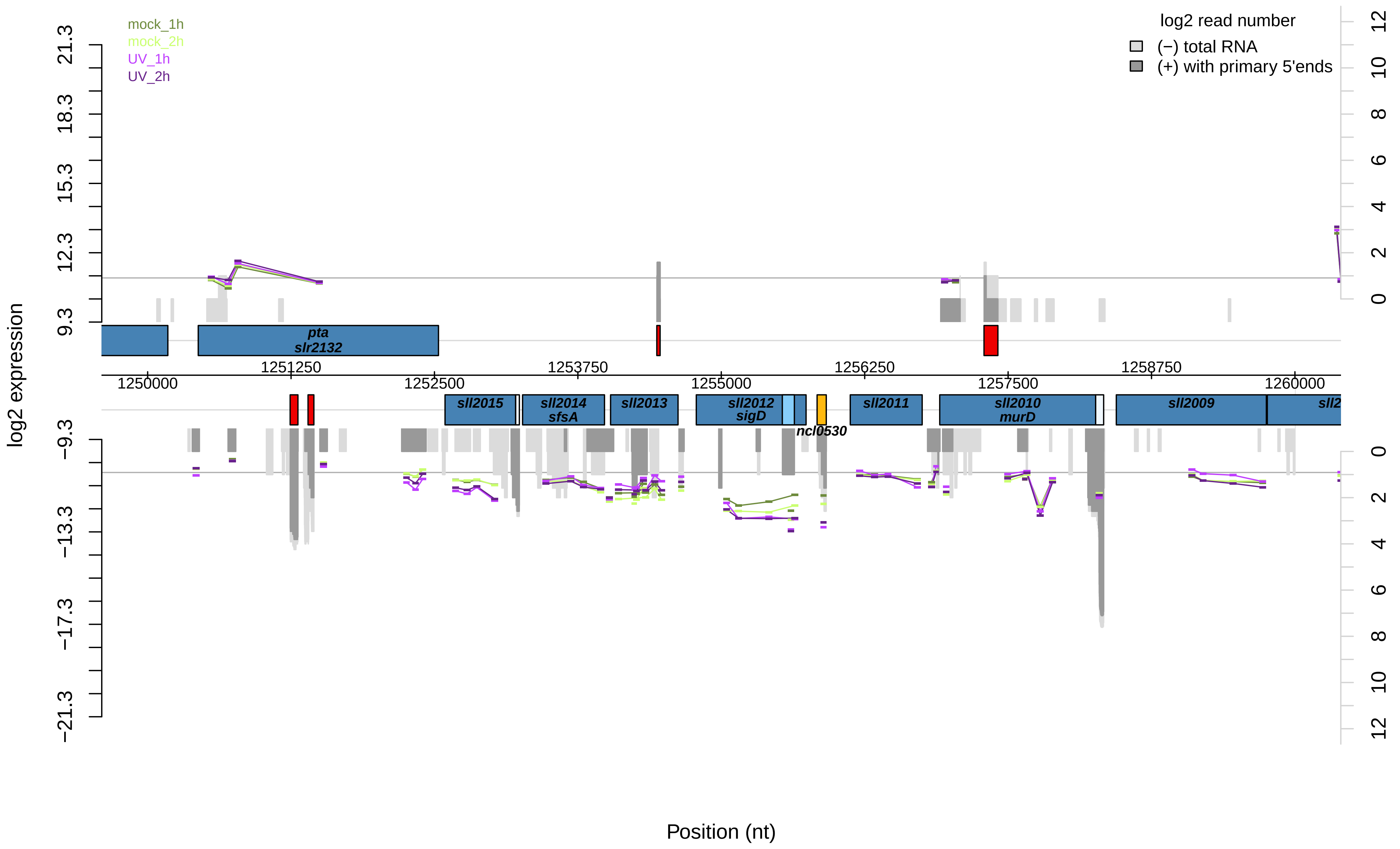
mock_1h
mock_2h
UV_1h
UV_2h
log2 read number
(−) total RNA
(+) with primary 5'ends
log2 expression
21.3
18.3
15.3
12.3
9.3
−9.3
−13.3
−17.3
−21.3
pta
slr2132
1250000
1251250
1252500
1253750
1255000
1256250
1257500
1258750
1260000
sll2015
sll2014
sfsA
sll2013
sll2012
sigD
ncl0530
sll2011
sll2010
murD
sll2009
Position (nt)

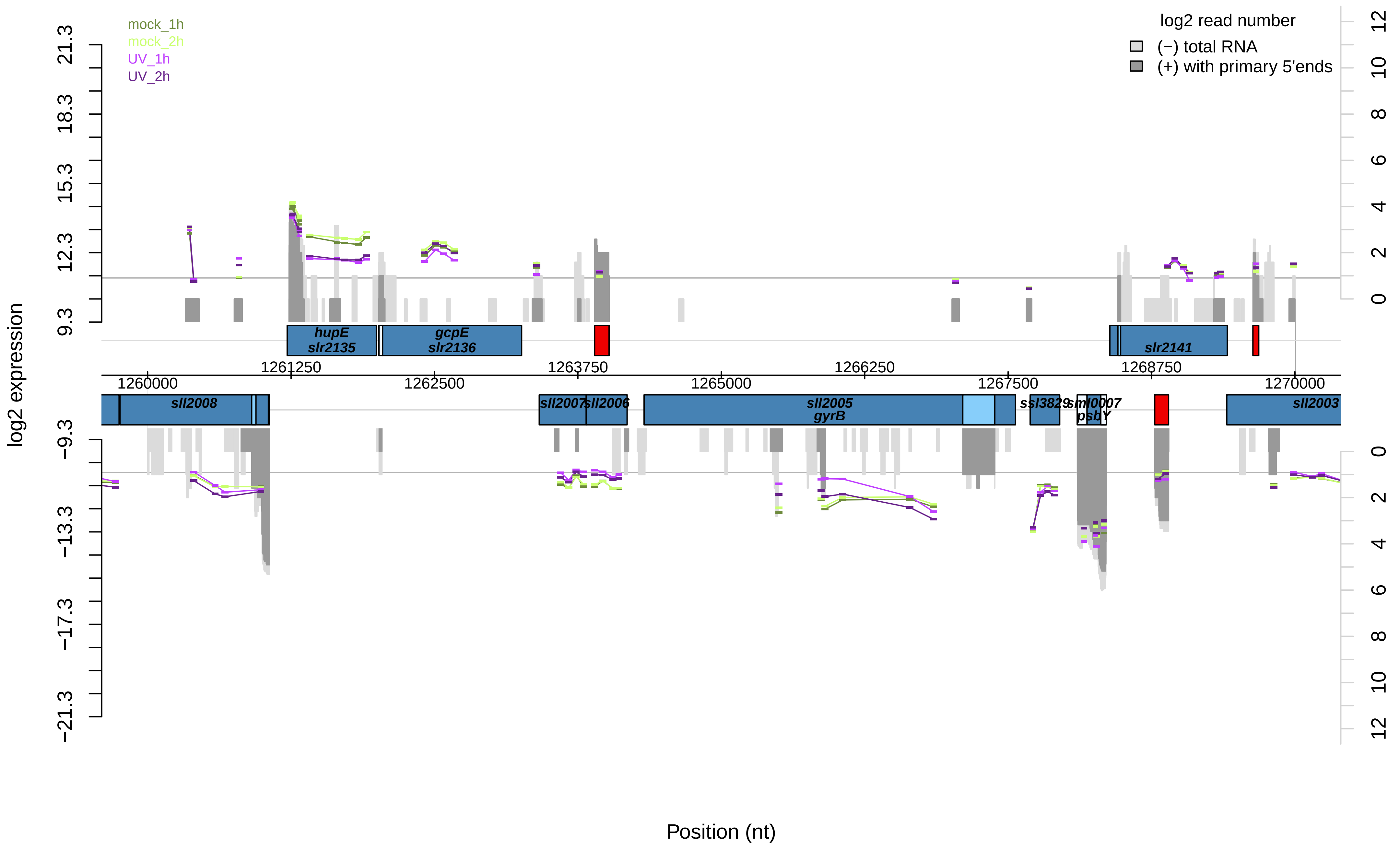

mock_1h
mock_2h
UV_1h
UV_2h
log2 read number
(−) total RNA
(+) with primary 5'ends
log2 expression
21.3
18.3
15.3
12.3
9.3
−9.3
−13.3
−17.3
−21.3
12
10
8
6
4
2
0
hupE
slr2135
gcpE
slr2136
slr2141
1261250
1263750
1266250
1268750
1260000
1262500
1265000
1267500
1270000
sll2008
sll2007
sll2006
sll2005
gyrB
ssl3829
sml0007
psbY
sll2003
Position (nt)

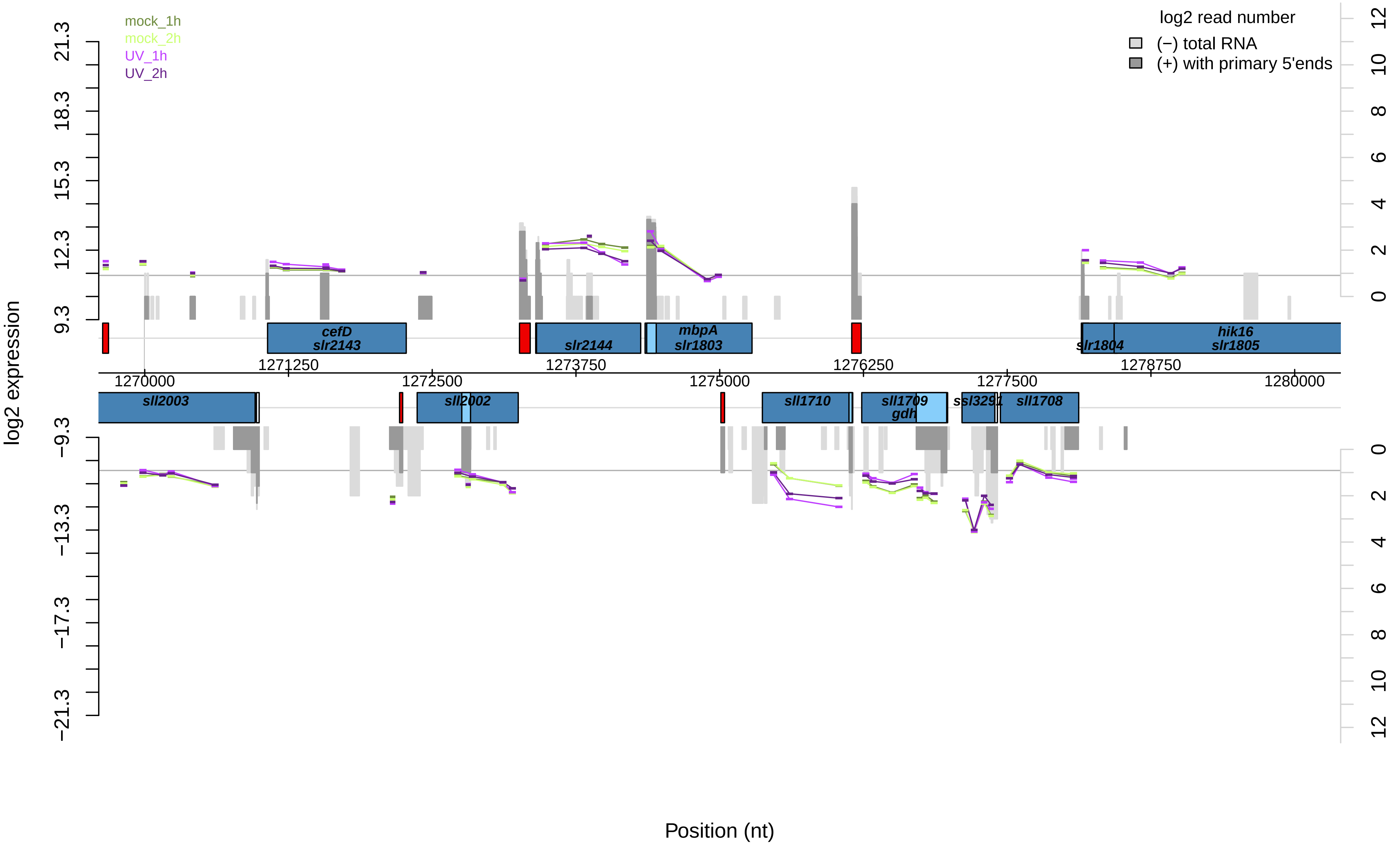

mock_1h
mock_2h
UV_1h
UV_2h
log2 read number
(−) total RNA
(+) with primary 5'ends
log2 expression
21.3
18.3
15.3
12.3
9.3
−9.3
−13.3
−17.3
−21.3
12
10
8
6
4
2
0
cefD
slr2143
slr2144
mbpA
slr1803
slr1804
hik16
slr1805
1271250
1273750
1276250
1278750
1270000
1272500
1275000
1277500
1280000
sll2003
sll2002
sll1710
sll1709
gdh
ssl3291
sll1708
Position (nt)

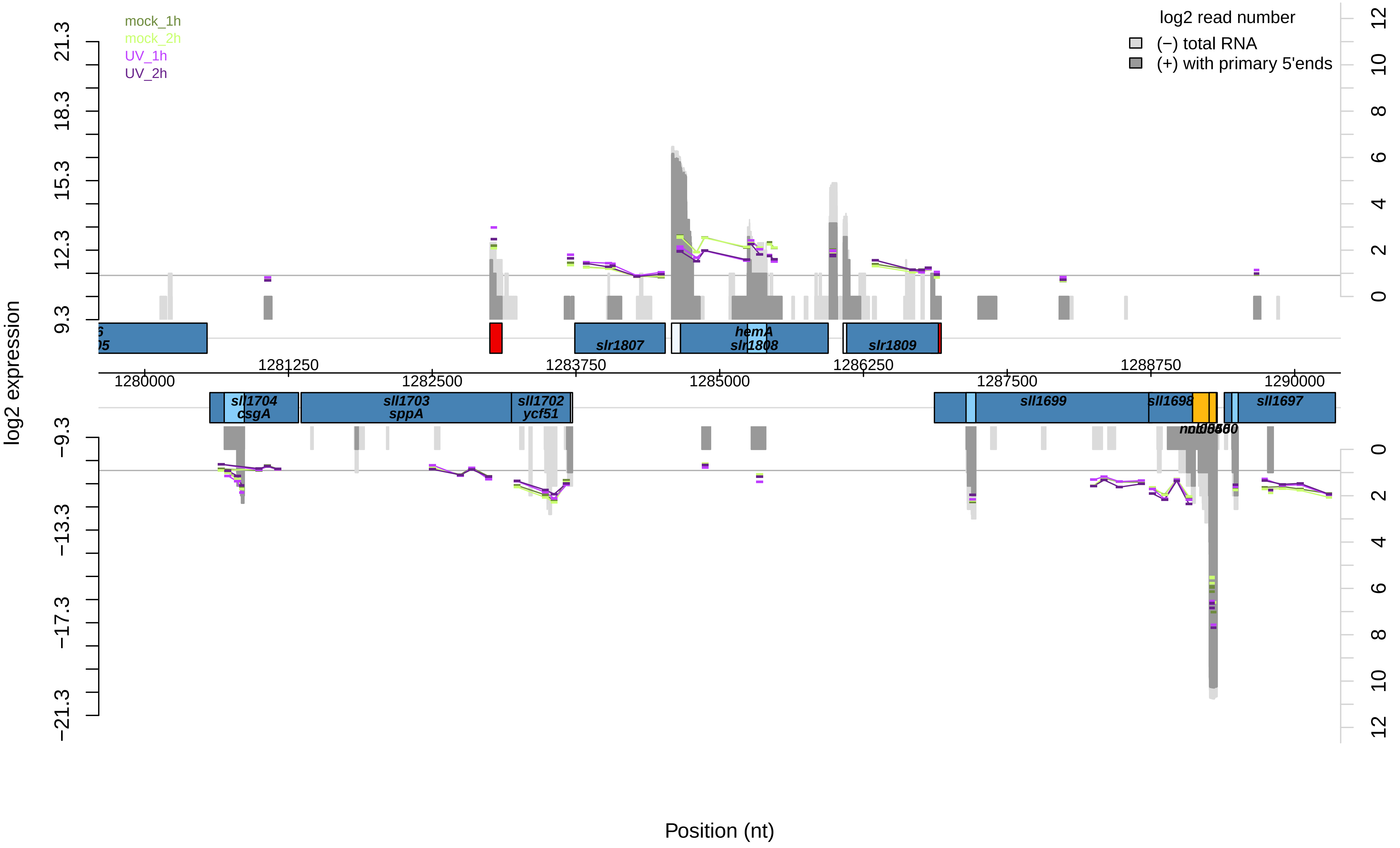

mock_1h
mock_2h
UV_1h
UV_2h
log2 read number
(−) total RNA
(+) with primary 5'ends
log2 expression
21.3
18.3
15.3
12.3
9.3
−9.3
−13.3
−17.3
−21.3
slr1807
hemA
slr1808
slr1809
sll1704
csgA
sll1703
sppA
sll1702
ycf51
sll1699
sll1698
sll1697
1280000
1281250
1282500
1283750
1285000
1286250
1287500
1288750
1290000
Position (nt)

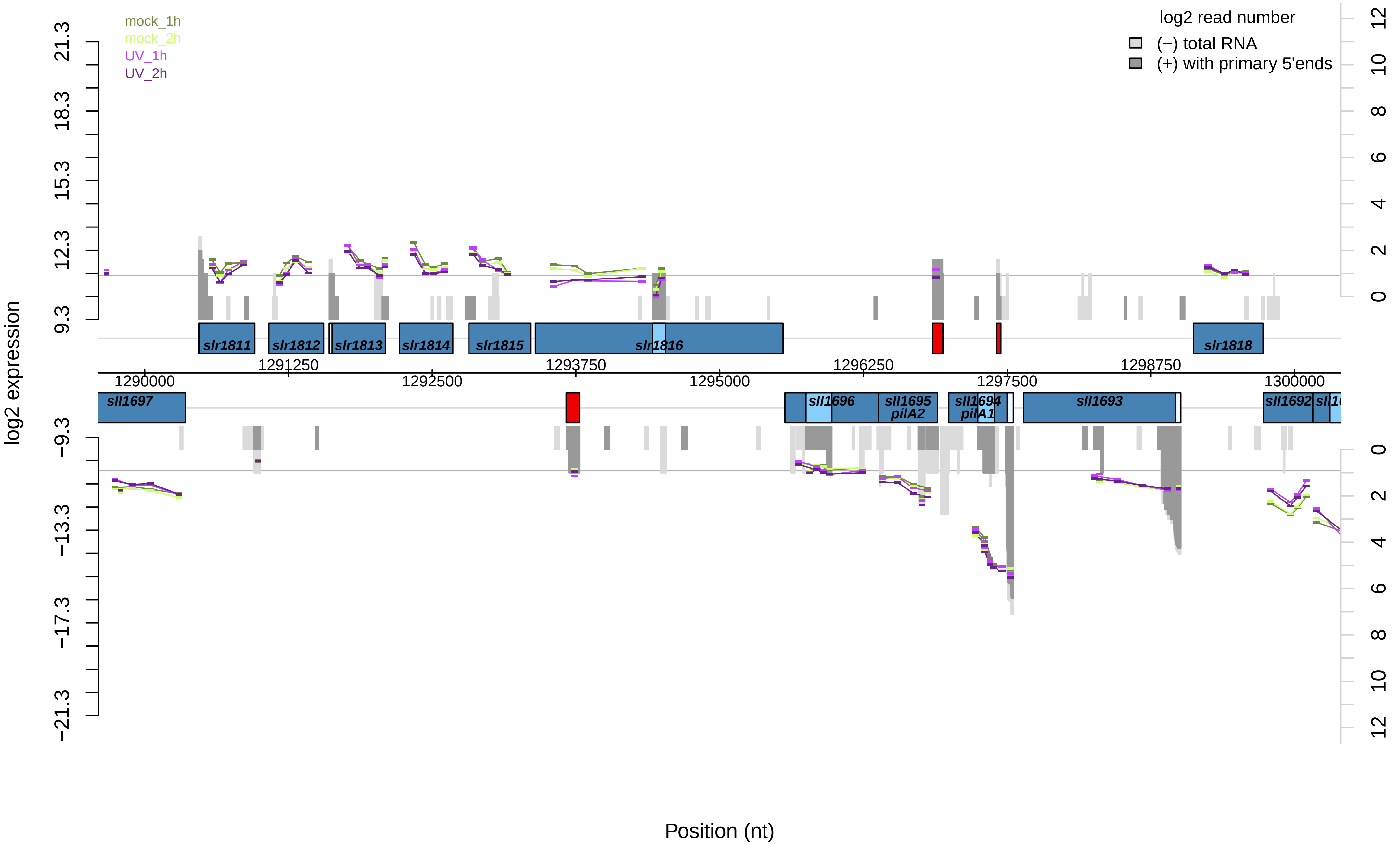

mock_1h
mock_2h
UV_1h
UV_2h
log2 read number
(−) total RNA
(+) with primary 5'ends
log2 expression
21.3
18.3
15.3
12.3
9.3
−9.3
−13.3
−17.3
−21.3
12
10
8
6
4
2
0
slr1811
slr1812
slr1813
slr1814
slr1815
slr1816
slr1818
1291250
1293750
1296250
1298750
1290000
1292500
1295000
1297500
1300000
sll1697
sll1696
sll1695
pilA2
sll1694
pilA1
sll1693
sll1692
Position (nt)

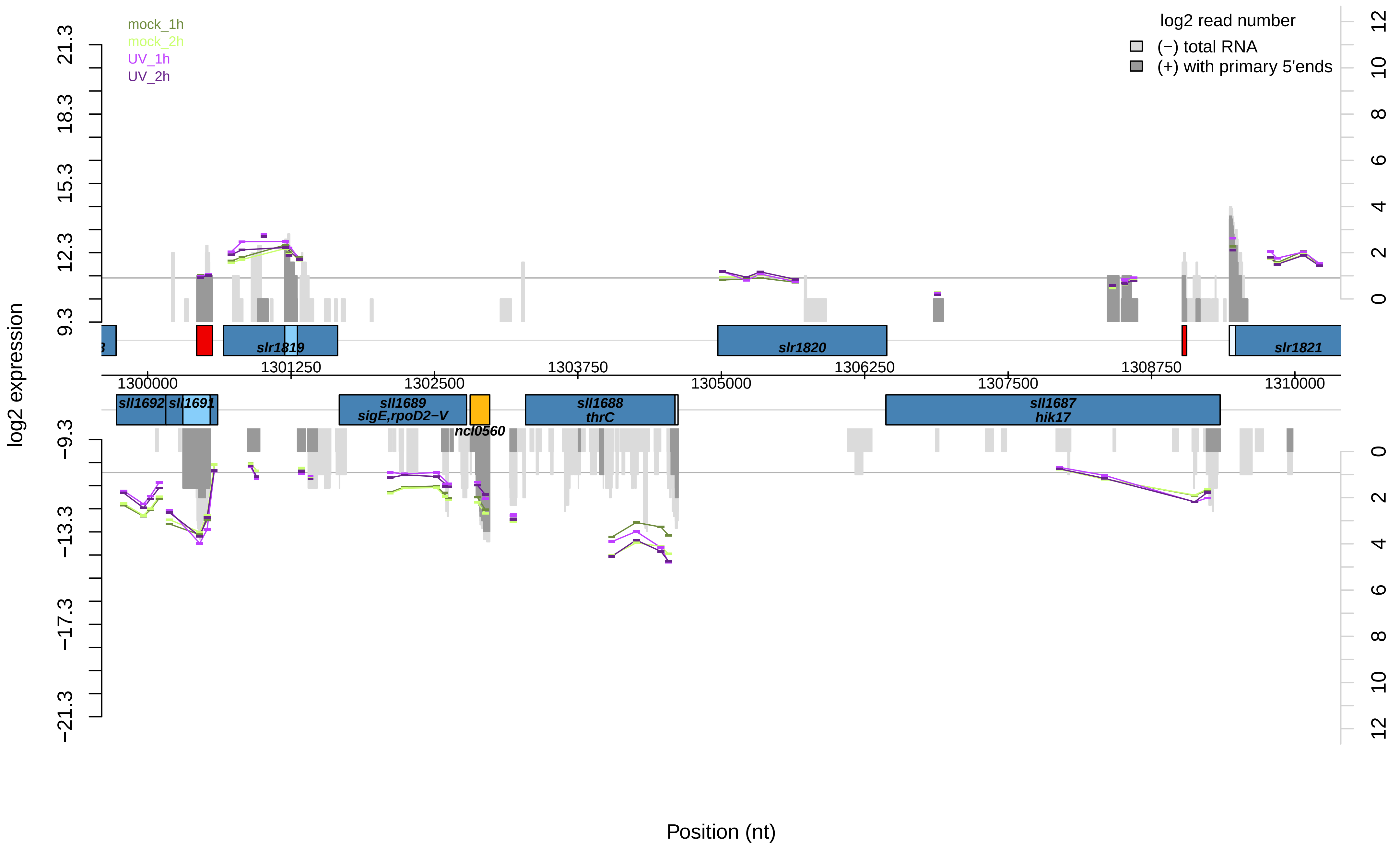

mock_1h
mock_2h
UV_1h
UV_2h
log2 read number
(−) total RNA
(+) with primary 5'ends
log2 expression
21.3
18.3
15.3
12.3
9.3
−9.3
−13.3
−17.3
−21.3
12
10
8
6
4
2
0
slr1819
slr1820
slr1821
1301250
1303750
1306250
1308750
1300000
1302500
1305000
1307500
1310000
sll1692
sll1691
sll1689
sigE,rpoD2−V
ncl0560
sll1688
thrC
sll1687
hik17
Position (nt)

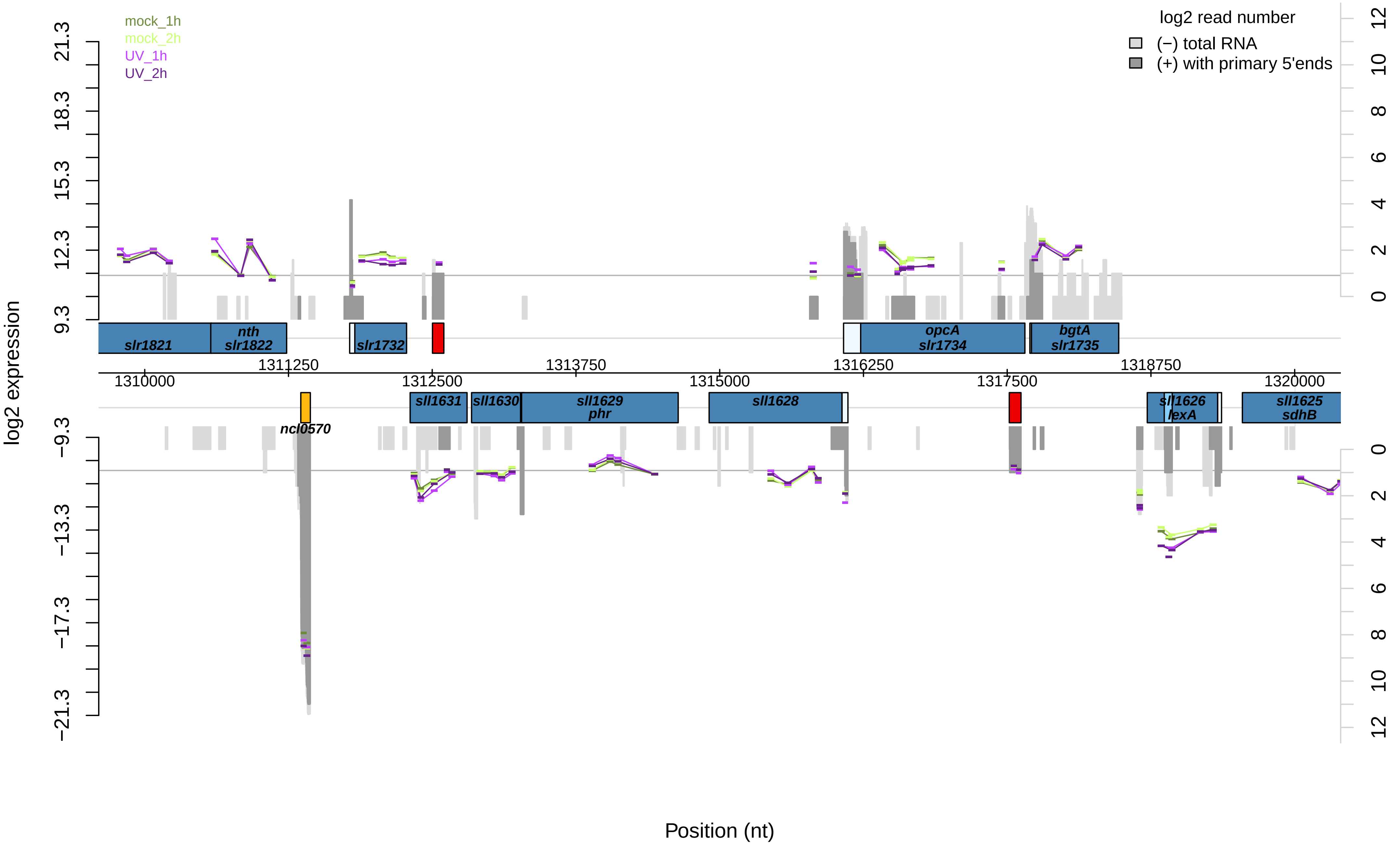
mock_1h
mock_2h
UV_1h
UV_2h
log2 read number
(−) total RNA
(+) with primary 5'ends
log2 expression
21.3
18.3
15.3
12.3
9.3
−9.3
−13.3
−17.3
−21.3
12
10
8
6
4
2
0
slr1821
nth
slr1822
slr1732
opcA
slr1734
bgtA
slr1735
ncl0570
sll1631
sll1630
sll1629
phr
sll1628
sll1626
lexA
sll1625
sdhB
1310000
1311250
1312500
1313750
1315000
1316250
1317500
1318750
1320000
Position (nt)

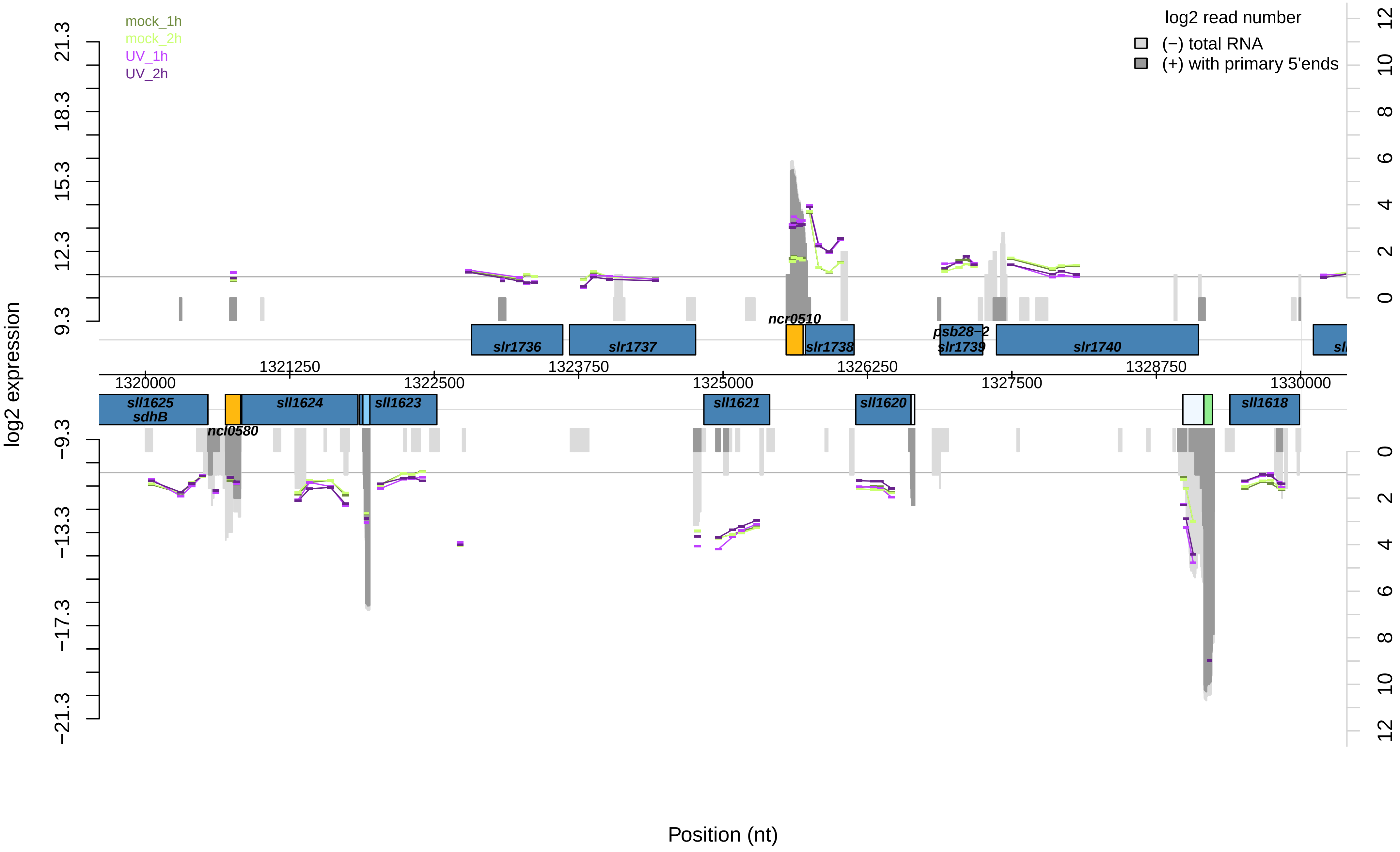

mock_1h
mock_2h
UV_1h
UV_2h
log2 read number
(−) total RNA
(+) with primary 5'ends
log2 expression
21.3
18.3
15.3
12.3
9.3
−9.3
−13.3
−17.3
−21.3
12
10
8
6
4
2
0
ncr0510
slr1736
slr1737
slr1738
psb28−2
slr1739
slr1740
1321250
1322500
1323750
1325000
1326250
1327500
1328750
1320000
1330000
sll1625
sdhB
ncl0580
sll1624
sll1623
sll1621
sll1620
sll1618
Position (nt)

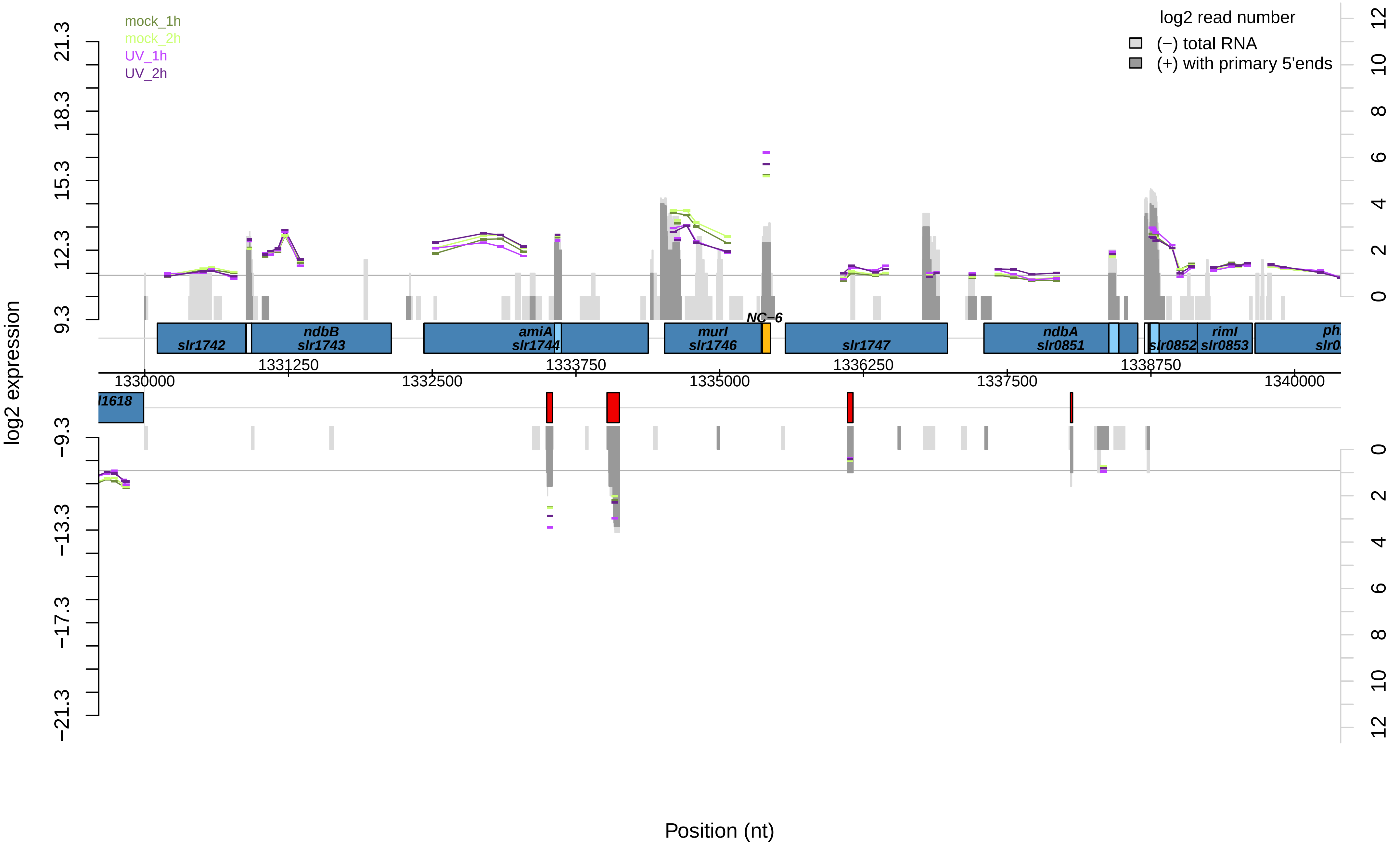
mock_1h
mock_2h
UV_1h
UV_2h
log2 read number
(−) total RNA
(+) with primary 5'ends
log2 expression
Position (nt)
slr1742
ndbB
slr1743
amiA
slr1744
murI
slr1746
NC−6
slr1747
ndbA
slr0851
slr0852
rimI
slr0853
1330000
1331250
1332500
1333750
1335000
1336250
1337500
1338750
1340000
21.3
18.3
15.3
12.3
9.3
−9.3
−13.3
−17.3
−21.3

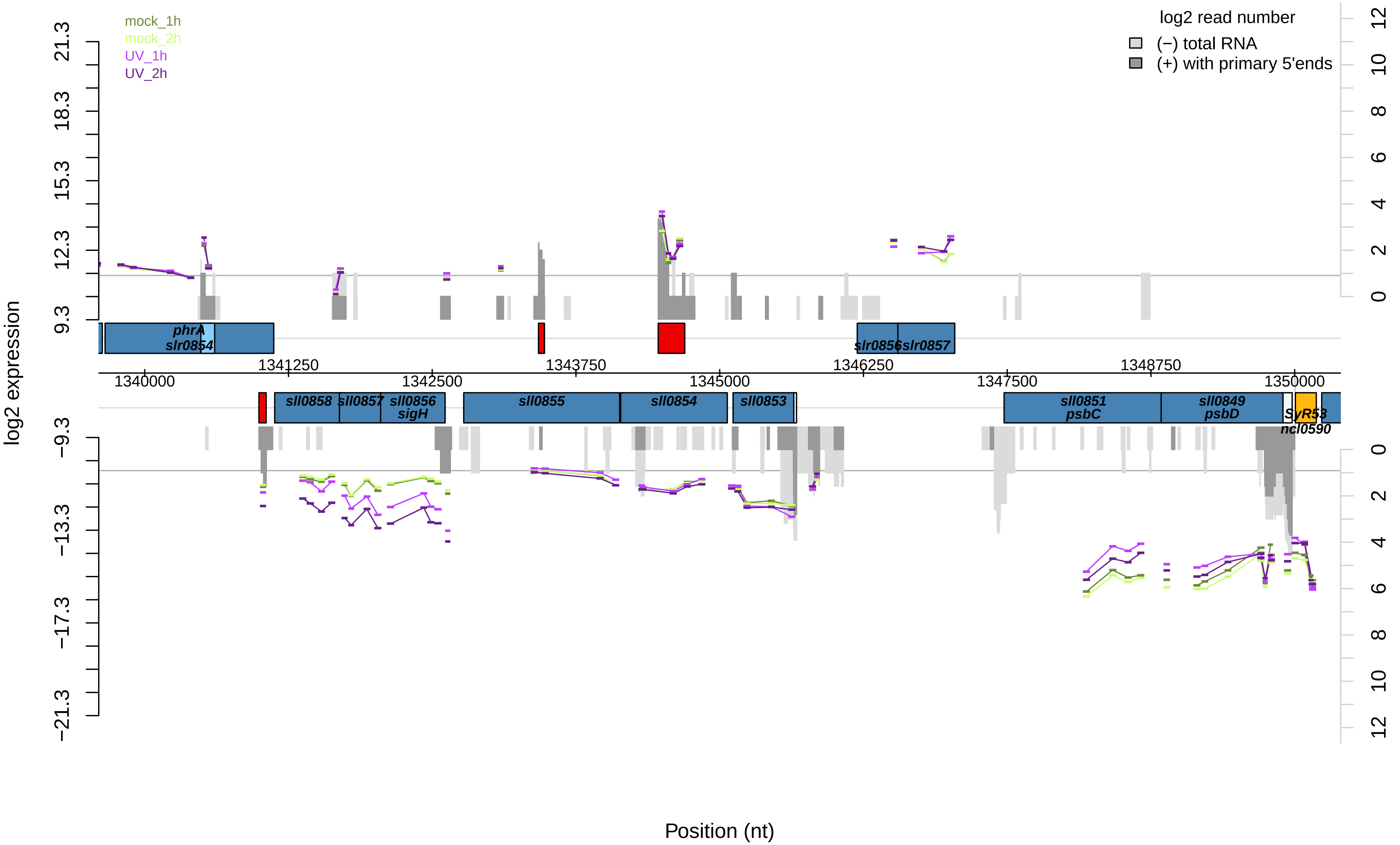

mock_1h
mock_2h
UV_1h
UV_2h
log2 read number
(−) total RNA
(+) with primary 5'ends
log2 expression
21.3
18.3
15.3
12.3
9.3
−9.3
−13.3
−17.3
−21.3
12
10
8
6
4
2
0
phrA
slr0854
slr0856
slr0857
1340000
1341250
1342500
1343750
1345000
1346250
1347500
1348750
1350000
sll0858
sll0857
sll0856
sigH
sll0855
sll0854
sll0853
sll0851
psbC
sll0849
psbD
SyR53
ncl0590
Position (nt)

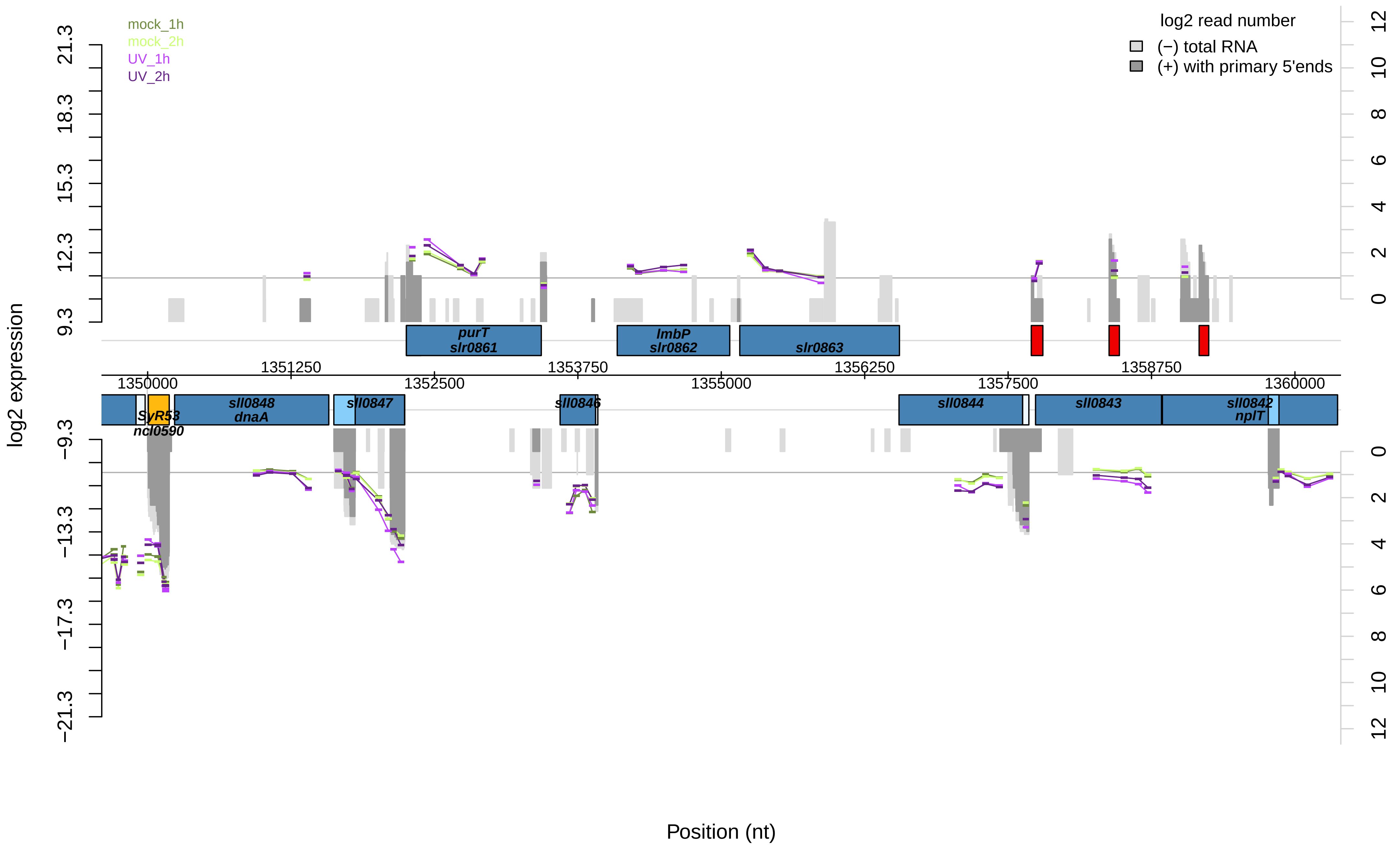

mock_1h
mock_2h
UV_1h
UV_2h
log2 read number
(−) total RNA
(+) with primary 5'ends
log2 expression
21.3
18.3
15.3
12.3
9.3
−9.3
−13.3
−17.3
−21.3
12
10
8
6
4
2
0
purT
slr0861
lmbP
slr0862
slr0863
1351250
1353750
1356250
1358750
1350000
1352500
1355000
1357500
1360000
sll0848
dnaA
SyR53
ncl0590
sll0847
sll0846
sll0844
sll0843
sll0842
nplT
Position (nt)

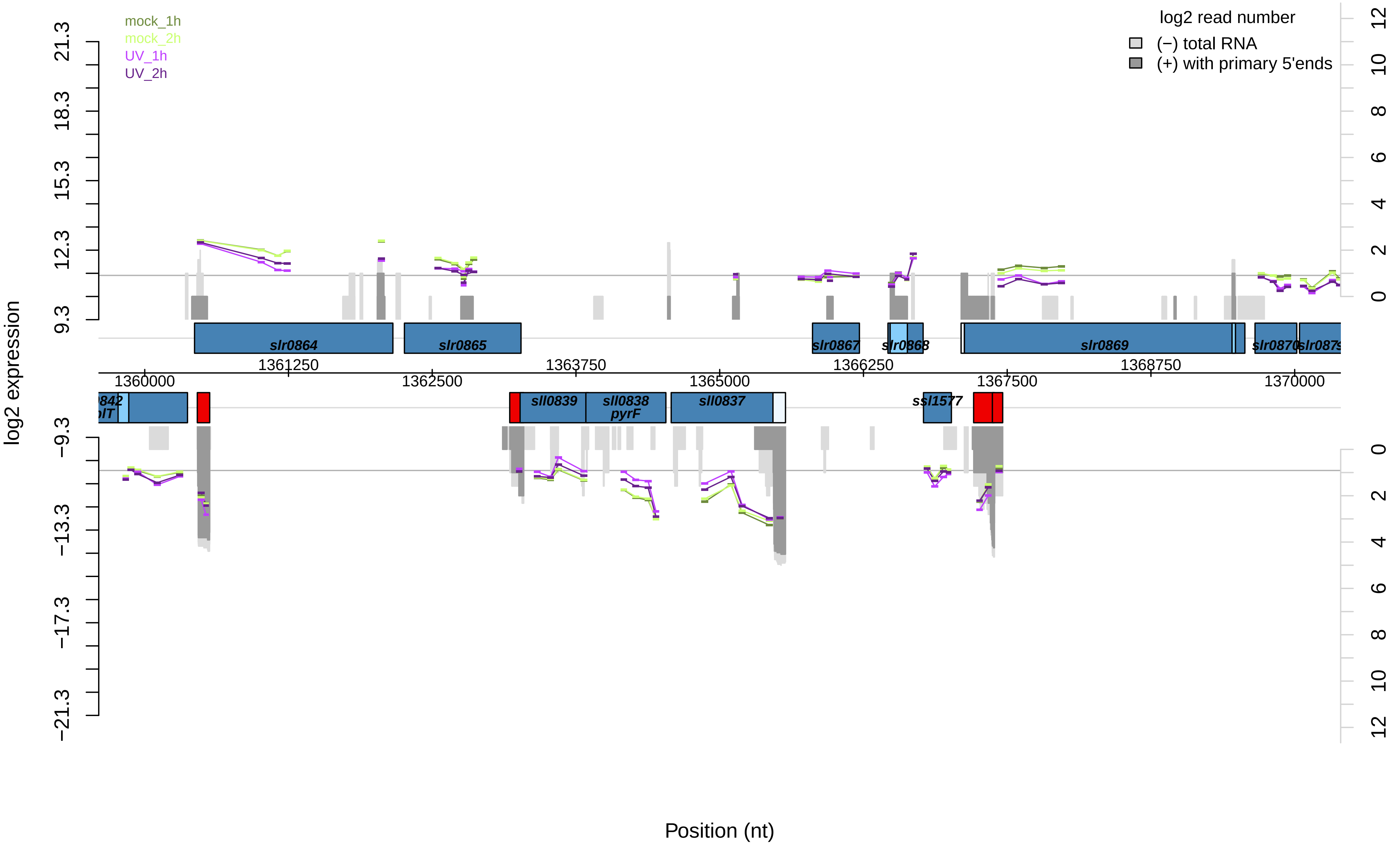

mock_1h
mock_2h
UV_1h
UV_2h
log2 read number
(−) total RNA
(+) with primary 5'ends
log2 expression
Position (nt)
slr0864
slr0865
slr0867
slr0868
slr0869
slr0870
sll0839
sll0838
pyrF
sll0837
ssl1577
1360000
1361250
1362500
1363750
1365000
1366250
1367500
1368750
1370000

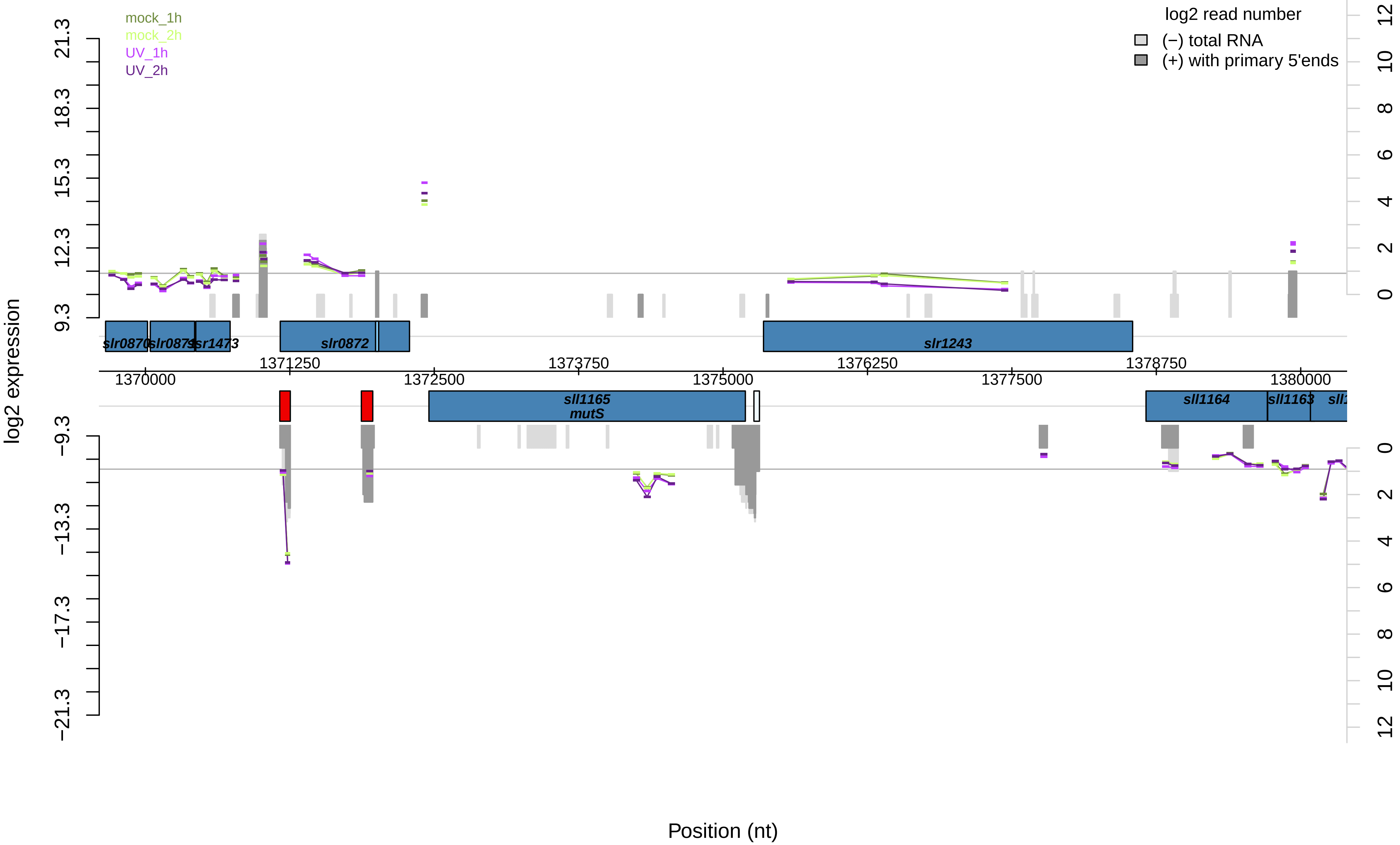

mock_1h
mock_2h
UV_1h
UV_2h
log2 read number
(−) total RNA
(+) with primary 5'ends
log2 expression
21.3
18.3
15.3
12.3
9.3
−9.3
−13.3
−17.3
−21.3
12
10
8
6
4
2
0
slr0870
slr0871
ssr1473
slr0872
slr1243
1371250
1373750
1376250
1378750
1370000
1372500
1375000
1377500
1380000
sll1165
mutS
sll1164
sll1163
Position (nt)

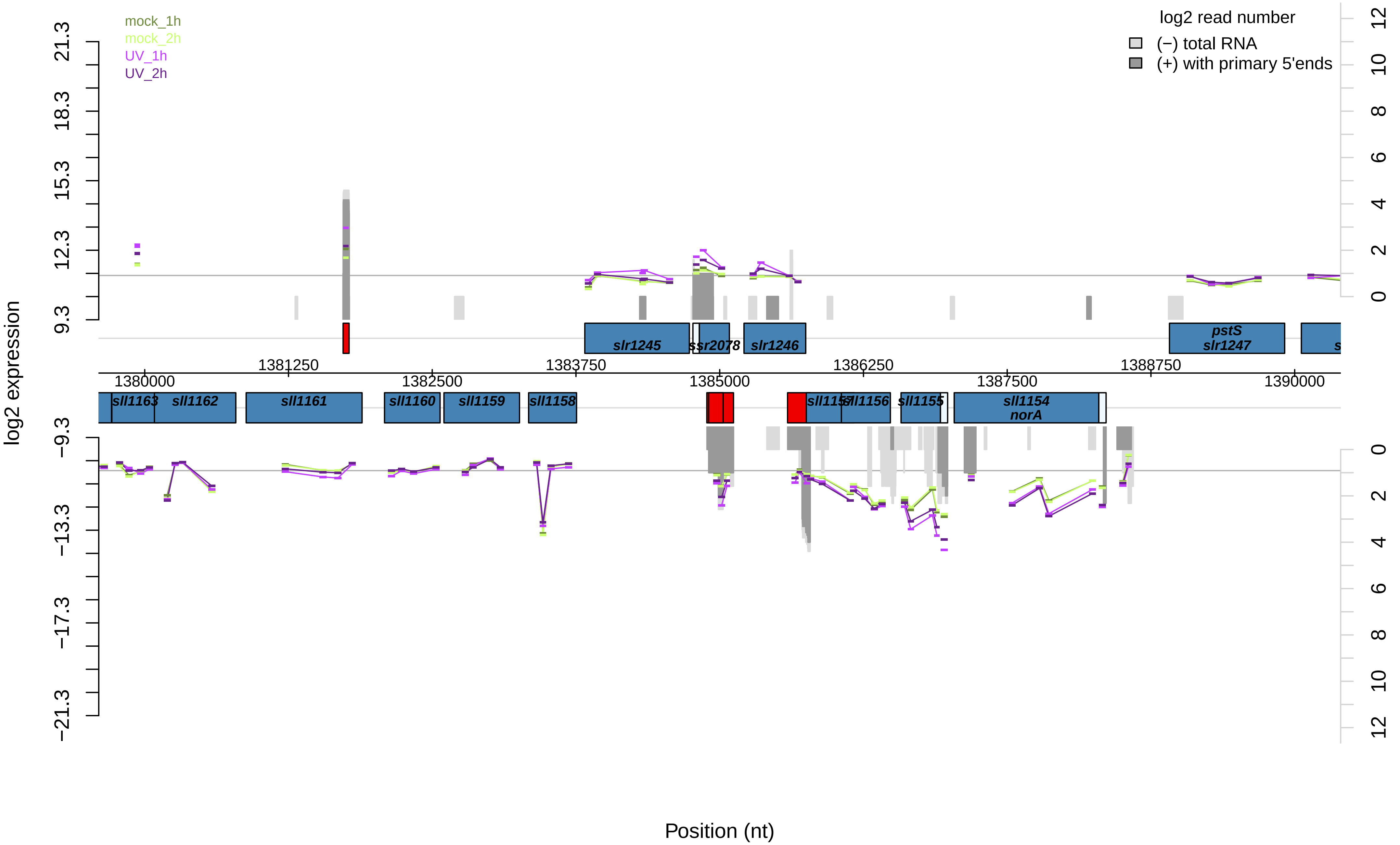

mock_1h
mock_2h
UV_1h
UV_2h
log2 read number
(−) total RNA
(+) with primary 5'ends
log2 expression
21.3
18.3
15.3
12.3
9.3
−9.3
−13.3
−17.3
−21.3
12
10
8
6
4
2
0
slr1245
ssr2078
slr1246
pstS
slr1247
1381250
1383750
1386250
1388750
1380000
1382500
1385000
1387500
1390000
sll1163
sll1162
sll1161
sll1160
sll1159
sll1158
sll1157
sll1156
sll1155
sll1154
norA
Position (nt)

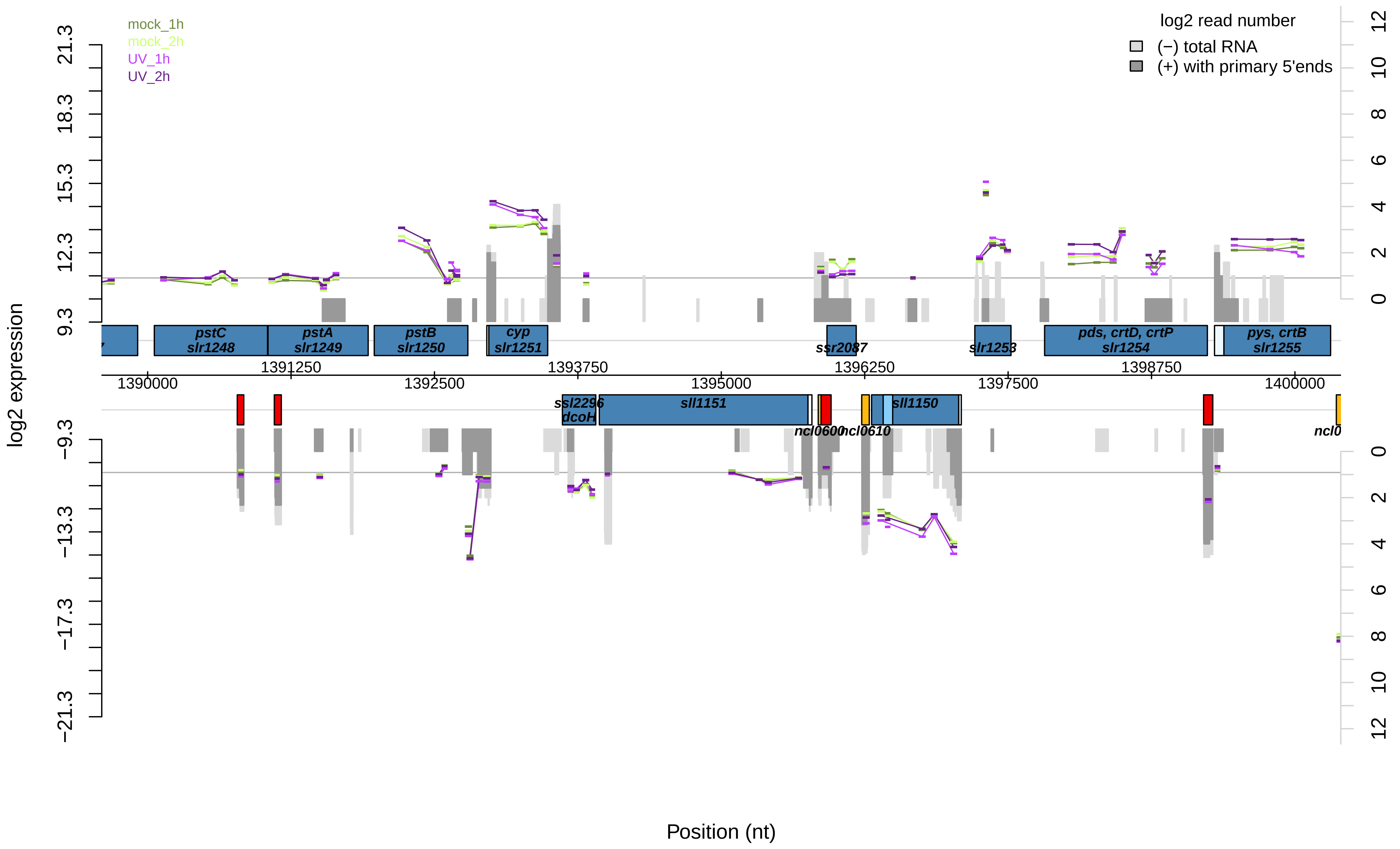

mock_1h
mock_2h
UV_1h
UV_2h
log2 read number
(−) total RNA
(+) with primary 5'ends
log2 expression
21.3
18.3
15.3
12.3
9.3
−9.3
−13.3
−17.3
−21.3
pstC slr1248
pstA slr1249
pstB slr1250
cyp slr1251
ssr2087
slr1253
pds, crtD, crtP slr1254
pys, crtB slr1255
1391250
1393750
1396250
1398750
1390000
1392500
1395000
1397500
1400000
ssl2296 dcoH
sll1151
sll1150
ncl0600
ncl0610
Position (nt)

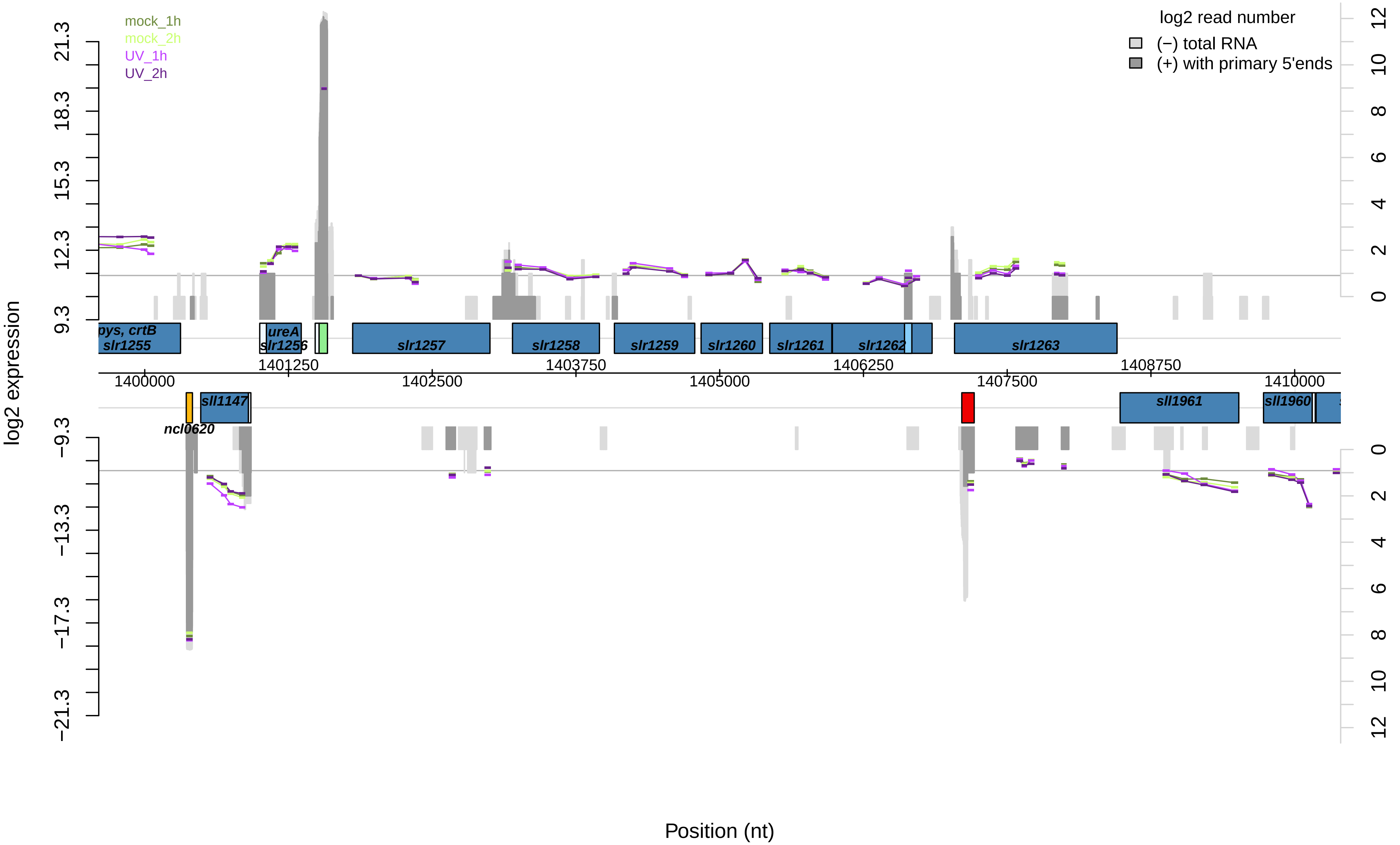
mock_1h
mock_2h
UV_1h
UV_2h
log2 read number
(−) total RNA
(+) with primary 5'ends
log2 expression
21.3
18.3
15.3
12.3
9.3
−9.3
−13.3
−17.3
−21.3
12
10
8
6
4
2
0
pys, crtB
slr1255
ureA
slr1256
slr1257
slr1258
slr1259
slr1260
slr1261
slr1262
slr1263
1401250
1403750
1406250
1408750
1400000
1402500
1405000
1407500
1410000
sll1147
ncl0620
sll1961
sll1960
Position (nt)

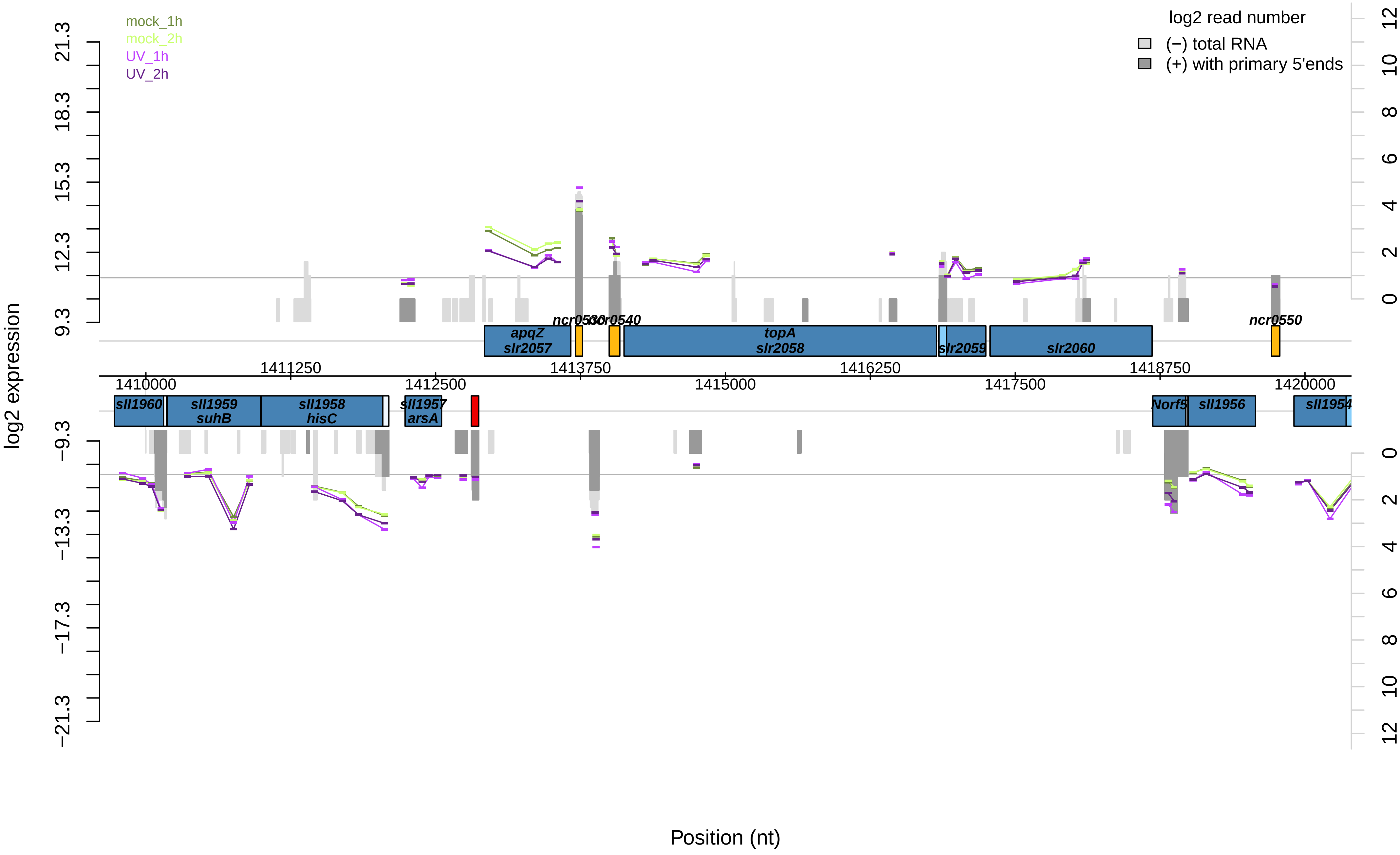

mock_1h
mock_2h
UV_1h
UV_2h
log2 read number
(−) total RNA
(+) with primary 5'ends
log2 expression
21.3
18.3
15.3
12.3
9.3
−9.3
−13.3
−17.3
−21.3
12
10
8
6
4
2
0
ncr0530
ncr0540
ncr0550
apqZ
slr2057
topA
slr2058
slr2059
slr2060
1411250
1413750
1416250
1418750
1410000
1412500
1415000
1417500
1420000
sll1960
sll1959
suhB
sll1958
hisC
sll1957
arsA
Norf5
sll1956
sll1954
Position (nt)

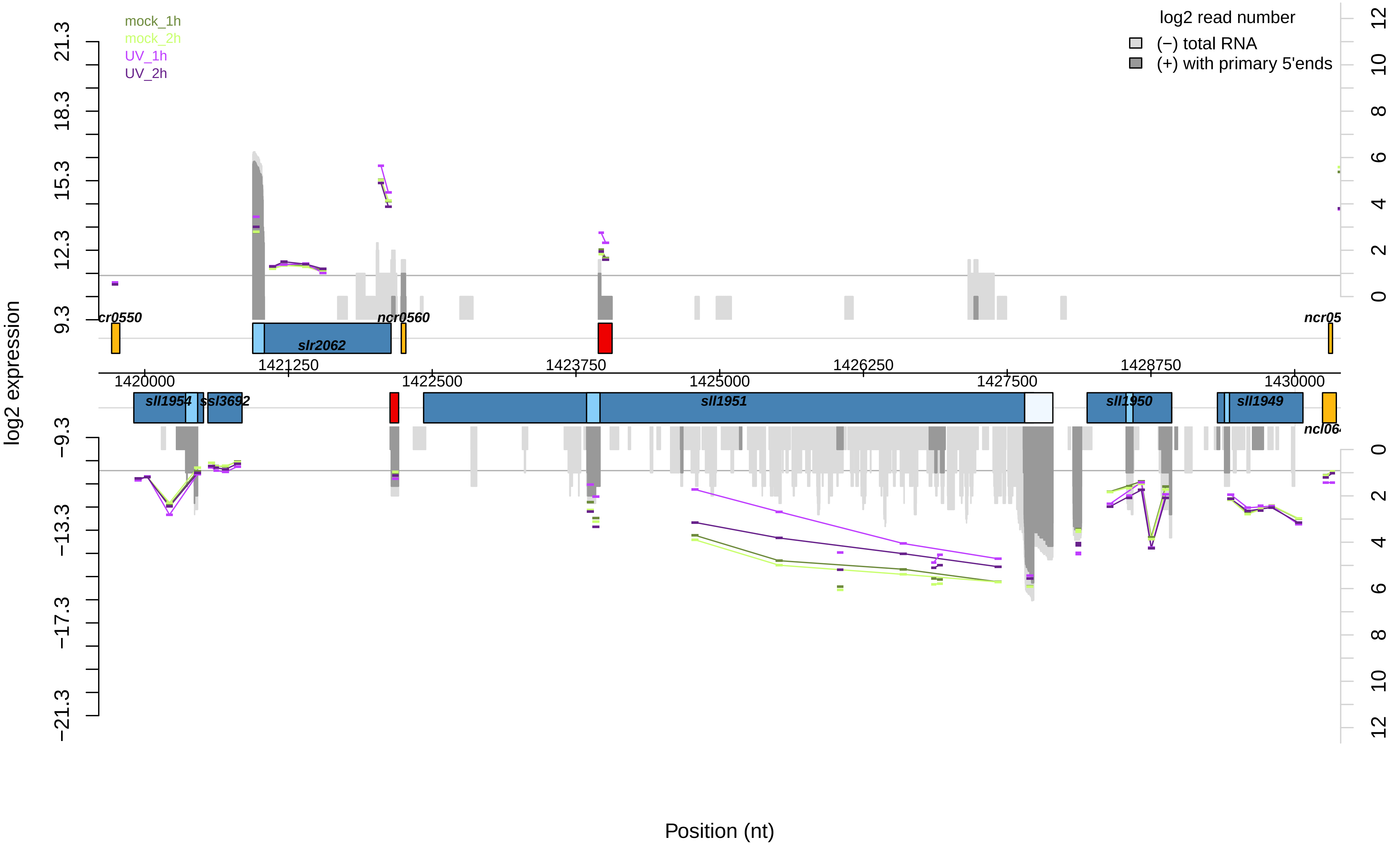

mock_1h
mock_2h
UV_1h
UV_2h
log2 read number
(−) total RNA
(+) with primary 5'ends
log2 expression
Position (nt)
ncr0550
ncr0560
slr2062
ncr05
sll1954
ssl3692
sll1951
sll1950
sll1949
ncl06
1420000
1421250
1422500
1423750
1425000
1426250
1427500
1428750
1430000
21.3
18.3
15.3
12.3
9.3
−9.3
−13.3
−17.3
−21.3

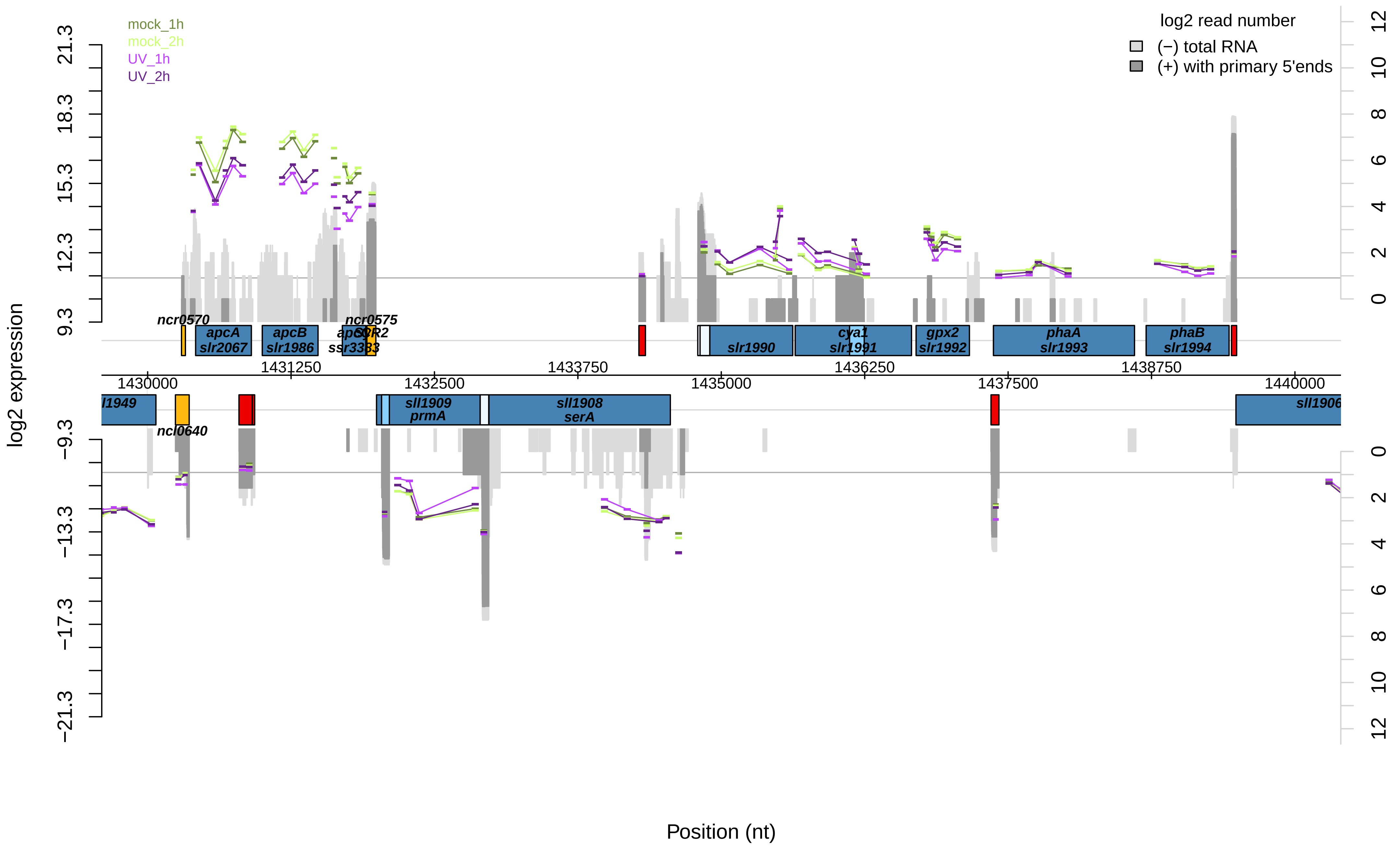

mock_1h
mock_2h
UV_1h
UV_2h
log2 read number
(−) total RNA
(+) with primary 5'ends
log2 expression
21.3
18.3
15.3
12.3
9.3
−9.3
−13.3
−17.3
−21.3
12
10
8
6
4
2
0
ncr0570
apcA
slr2067
apcB
slr1986
apcC
ssr3383
ncr0575
SOR2
slr1990
cya1
slr1991
gpx2
slr1992
phaA
slr1993
phaB
slr1994
1431250
1432500
1433750
1435000
1436250
1437500
1438750
1430000
1440000
l1949
ncl0640
sll1909
prmA
sll1908
serA
sll1906
Position (nt)

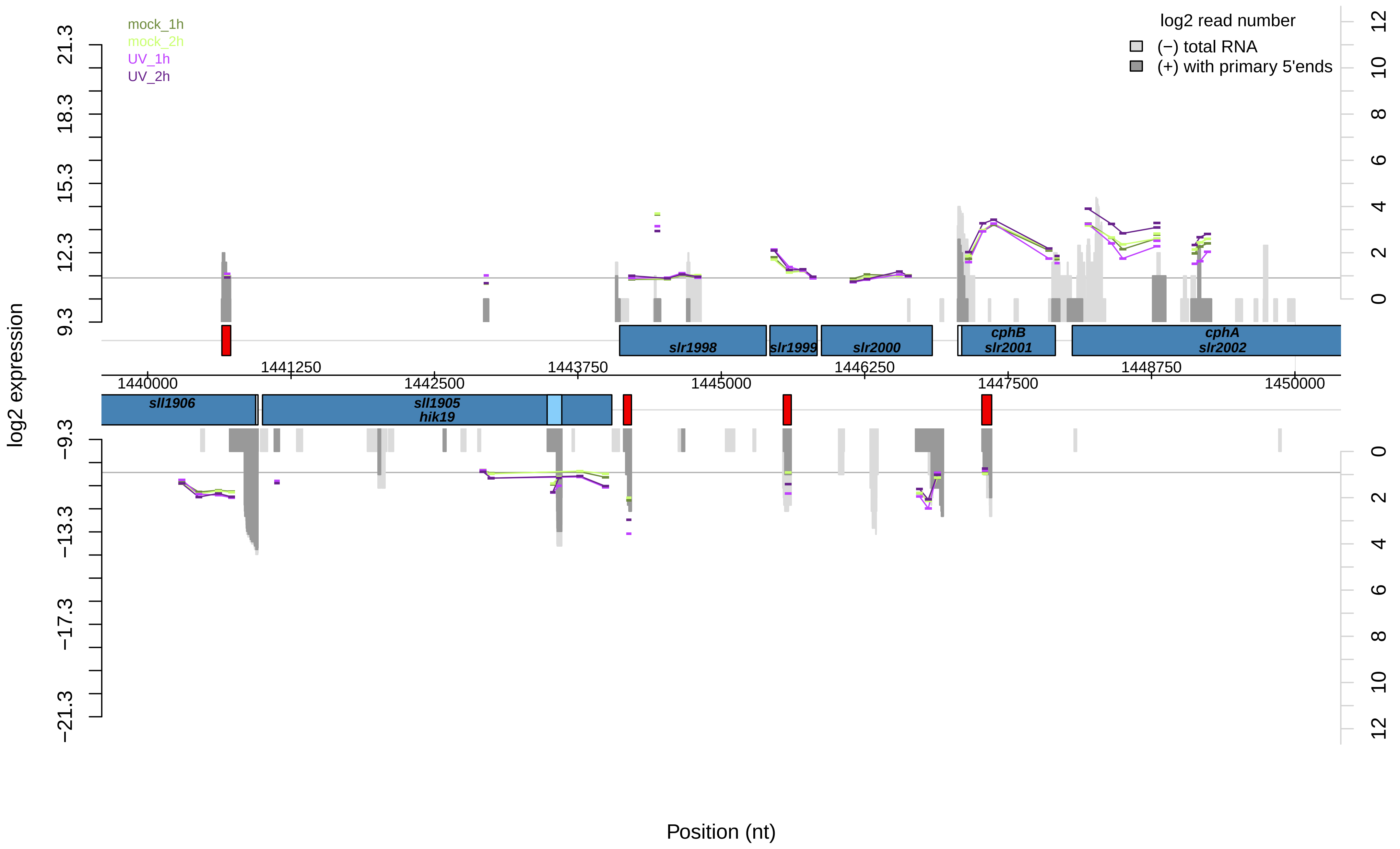

mock_1h
mock_2h
UV_1h
UV_2h
log2 read number
(−) total RNA
(+) with primary 5'ends
log2 expression
21.3
18.3
15.3
12.3
9.3
−9.3
−13.3
−17.3
−21.3
12
10
8
6
4
2
0
slr1998
slr1999
slr2000
cphB
slr2001
cphA
slr2002
1441250
1443750
1446250
1448750
1440000
1442500
1445000
1447500
1450000
sll1906
sll1905
hik19
Position (nt)

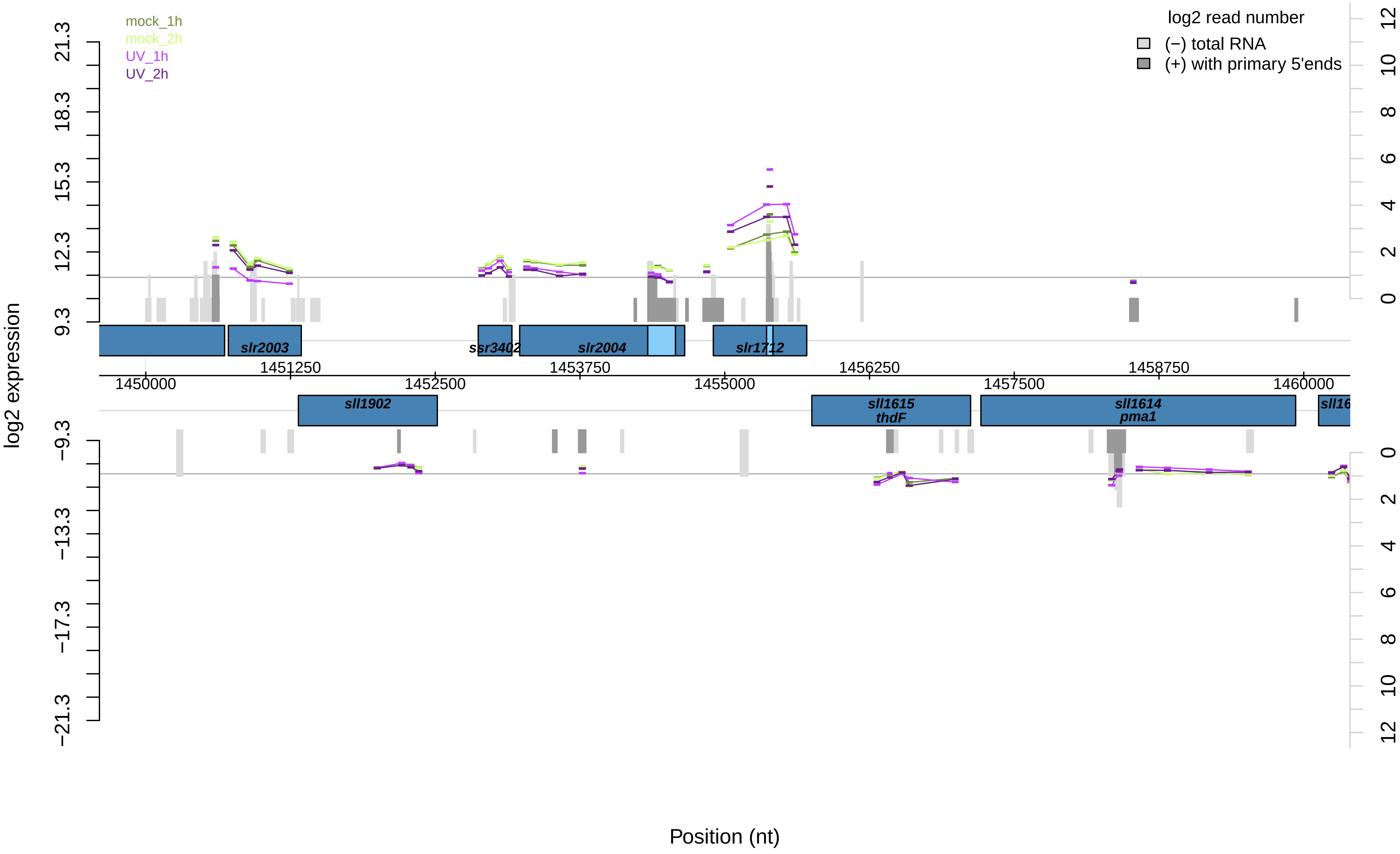

mock_1h
mock_2h
UV_1h
UV_2h
log2 read number
(−) total RNA
(+) with primary 5'ends
log2 expression
21.3
18.3
15.3
12.3
9.3
−9.3
−13.3
−17.3
−21.3
12
10
8
6
4
2
0
slr2003
ssr3402
slr2004
slr1712
1450000
1451250
1452500
1453750
1455000
1456250
1457500
1458750
1460000
sll1902
sll1615
thdF
sll1614
pma1
sll16
Position (nt)

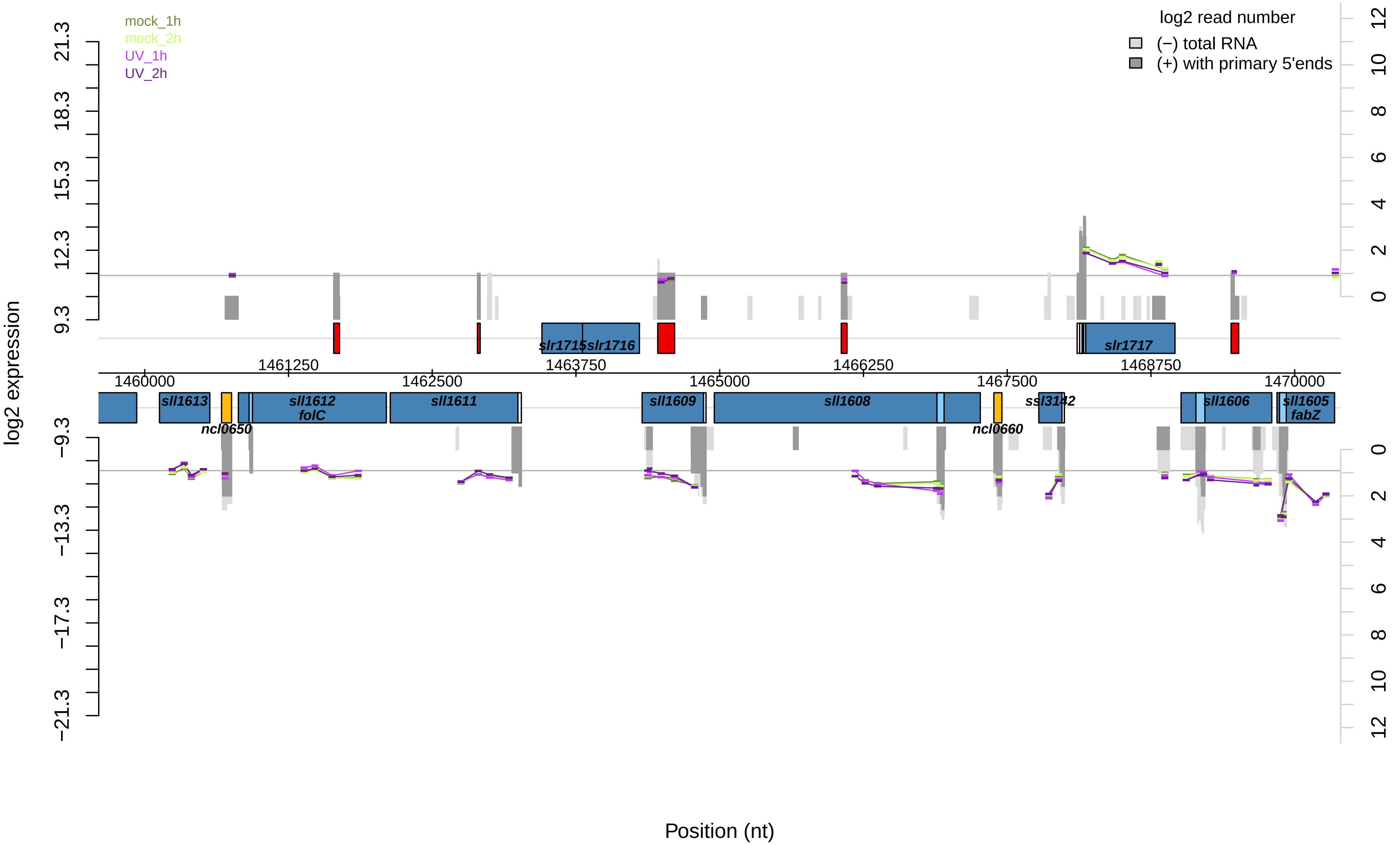

mock_1h
mock_2h
UV_1h
UV_2h
log2 read number
(−) total RNA
(+) with primary 5'ends
log2 expression
21.3
18.3
15.3
12.3
9.3
−9.3
−13.3
−17.3
−21.3
12
10
8
6
4
2
0
slr1715
slr1716
slr1717
1461250
1463750
1466250
1468750
1460000
1462500
1465000
1467500
1470000
sll1613
sll1612
folC
sll1611
sll1609
sll1608
ssl3142
sll1606
sll1605
fabZ
ncl0650
ncl0660
Position (nt)

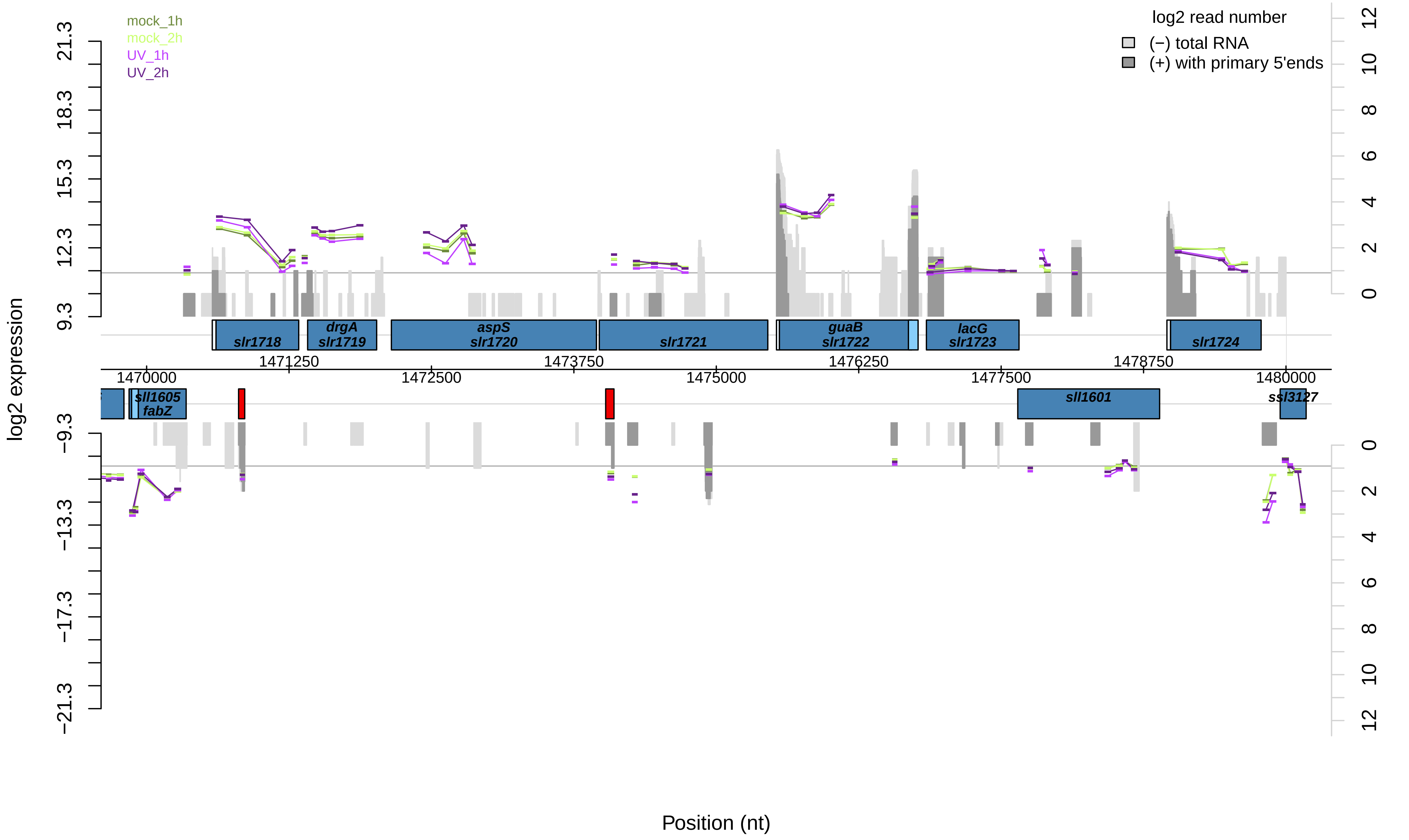

mock_1h
mock_2h
UV_1h
UV_2h
log2 read number
(−) total RNA
(+) with primary 5'ends
log2 expression
21.3
18.3
15.3
12.3
9.3
−9.3
−13.3
−17.3
−21.3
12
10
8
6
4
2
0
slr1718
drgA
slr1719
aspS
slr1720
slr1721
guaB
slr1722
lacG
slr1723
slr1724
1471250
1473750
1476250
1478750
1470000
1472500
1475000
1477500
1480000
sll1605
fabZ
sll1601
ssl3127
Position (nt)

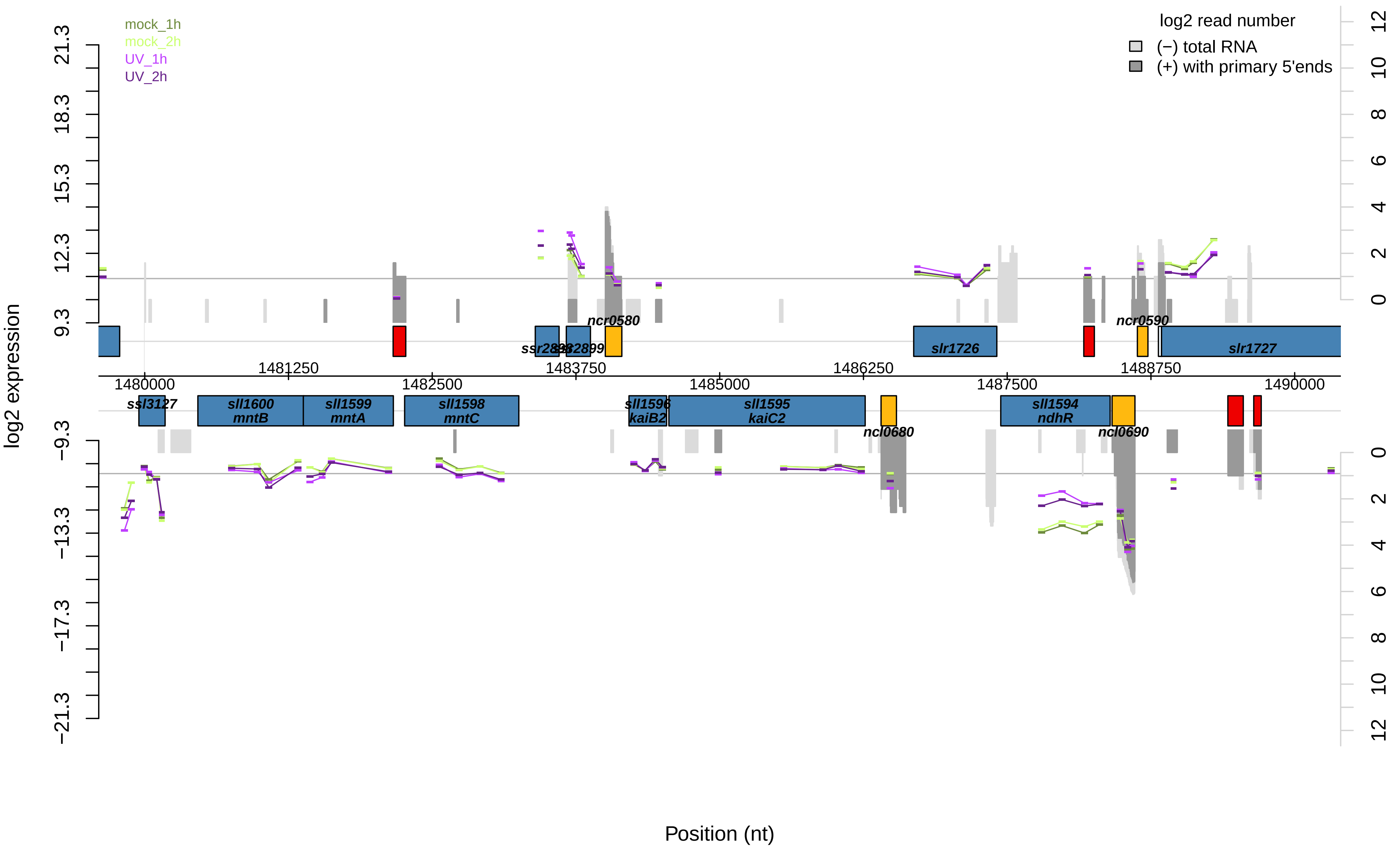

mock_1h
mock_2h
UV_1h
UV_2h
log2 read number
(−) total RNA
(+) with primary 5'ends
log2 expression
21.3
18.3
15.3
12.3
9.3
−9.3
−13.3
−17.3
−21.3
12
10
8
6
4
2
0
ncr0580
ncr0590
ssr2898
ssl2899
slr1726
slr1727
1481250
1483750
1486250
1488750
1480000
1482500
1485000
1487500
1490000
ssl3127
sll1600
mntB
sll1599
mntA
sll1598
mntC
sll1596
kaiB2
sll1595
kaiC2
ncl0680
sll1594
ndhR
ncl0690
Position (nt)

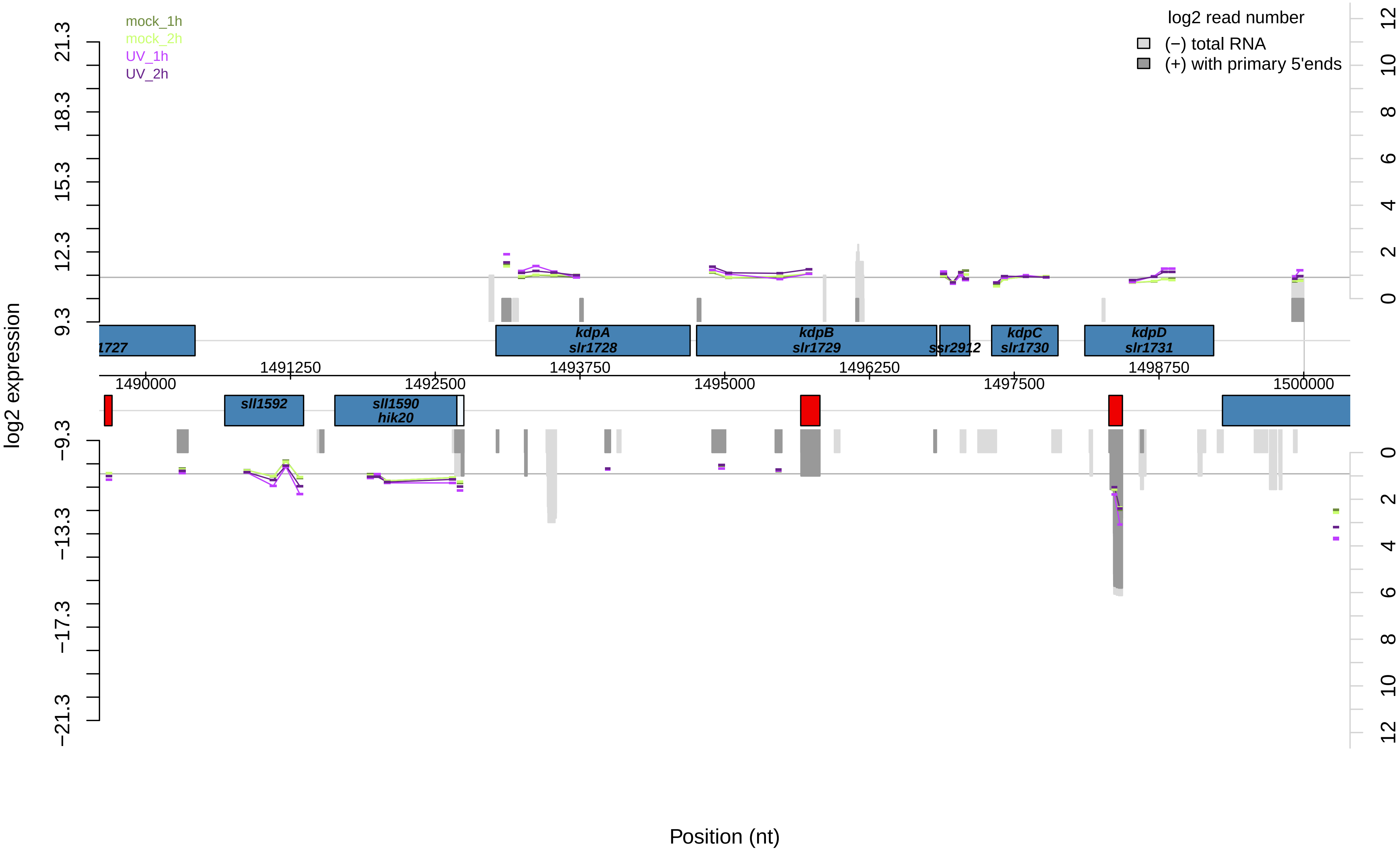

mock_1h
mock_2h
UV_1h
UV_2h
log2 read number
(−) total RNA
(+) with primary 5'ends
log2 expression
21.3
18.3
15.3
12.3
9.3
−9.3
−13.3
−17.3
−21.3
12
10
8
6
4
2
0
1727
kdpA
slr1728
kdpB
slr1729
ssr2912
kdpC
slr1730
kdpD
slr1731
sll1592
sll1590
hik20
1490000
1491250
1492500
1493750
1495000
1496250
1497500
1498750
1500000
Position (nt)

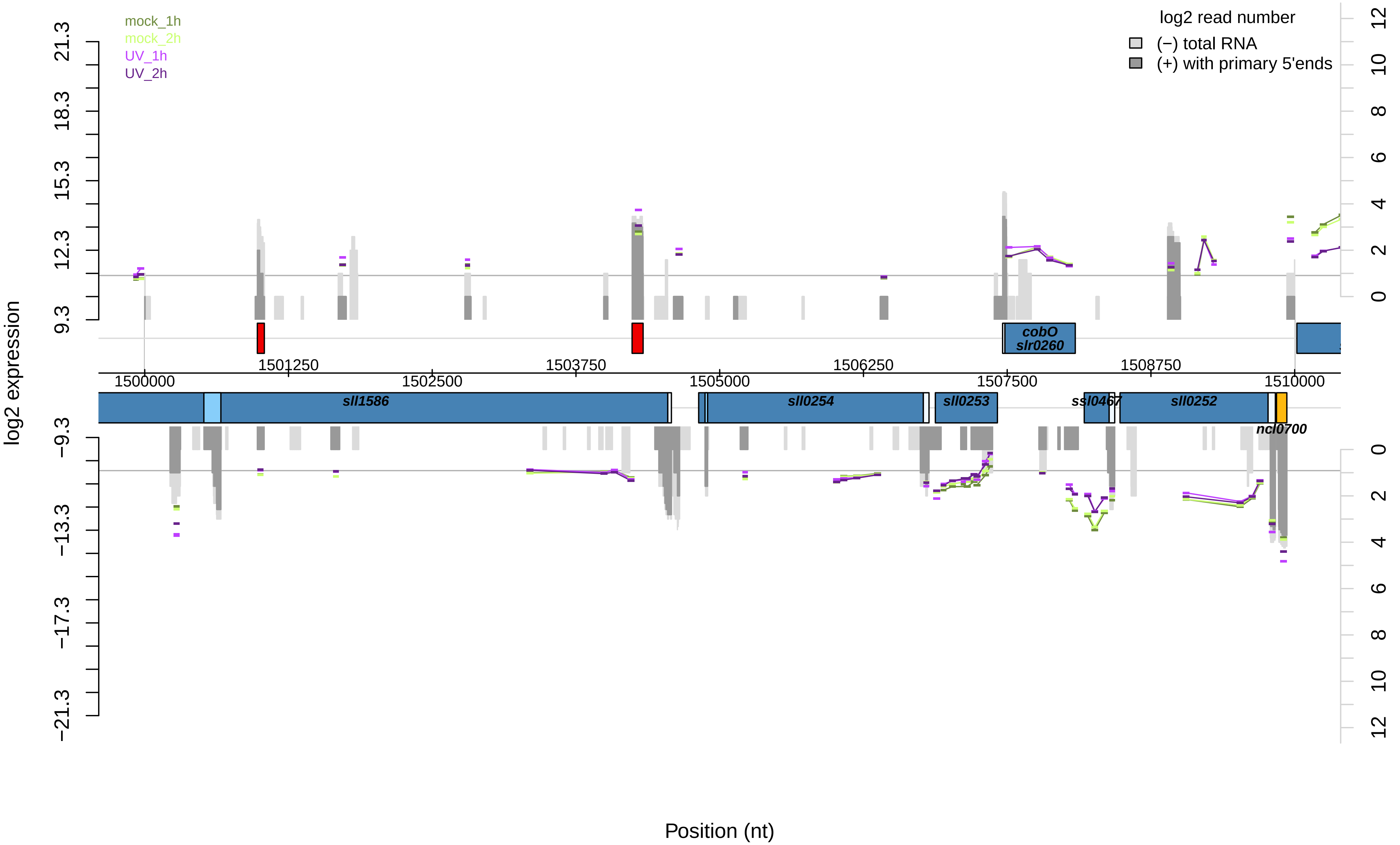
mock_1h
mock_2h
UV_1h
UV_2h
log2 read number
(−) total RNA
(+) with primary 5'ends
log2 expression
21.3
18.3
15.3
12.3
9.3
−9.3
−13.3
−17.3
−21.3
12
10
8
6
4
2
0
cobO
slr0260
sll1586
sll0254
sll0253
ssl0467
sll0252
ncl0700
1500000
1501250
1502500
1503750
1505000
1506250
1507500
1508750
1510000
Position (nt)

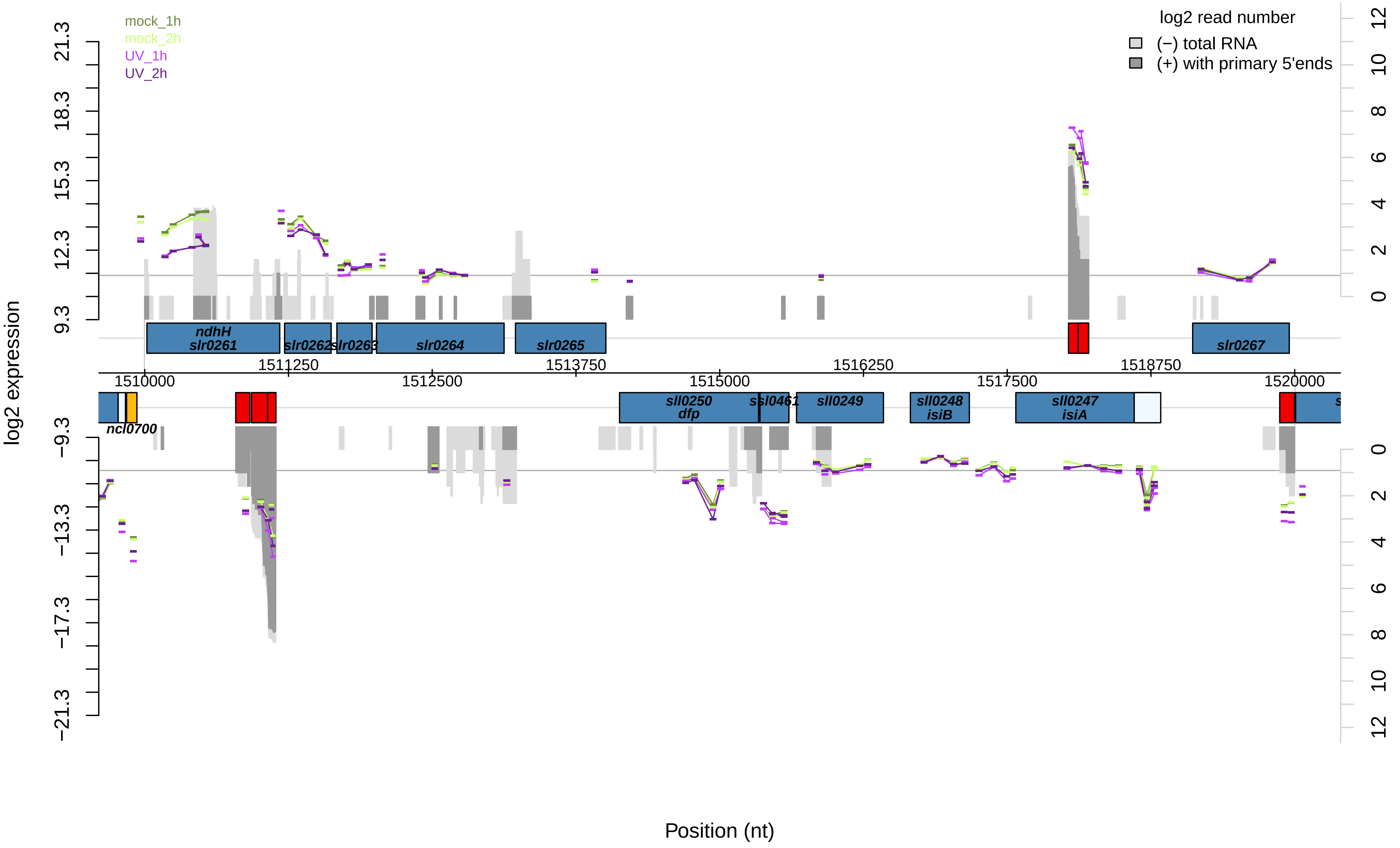

mock_1h
mock_2h
UV_1h
UV_2h
log2 read number
(−) total RNA
(+) with primary 5'ends
log2 expression
21.3
18.3
15.3
12.3
9.3
−9.3
−13.3
−17.3
−21.3
12
10
8
6
4
2
0
ndhH
slr0261
slr0262
slr0263
slr0264
slr0265
slr0267
1511250
1512500
1513750
1515000
1516250
1517500
1518750
1520000
1510000
ncl0700
sll0250
dfp
ssl0461
sll0249
sll0248
isiB
sll0247
isiA
Position (nt)

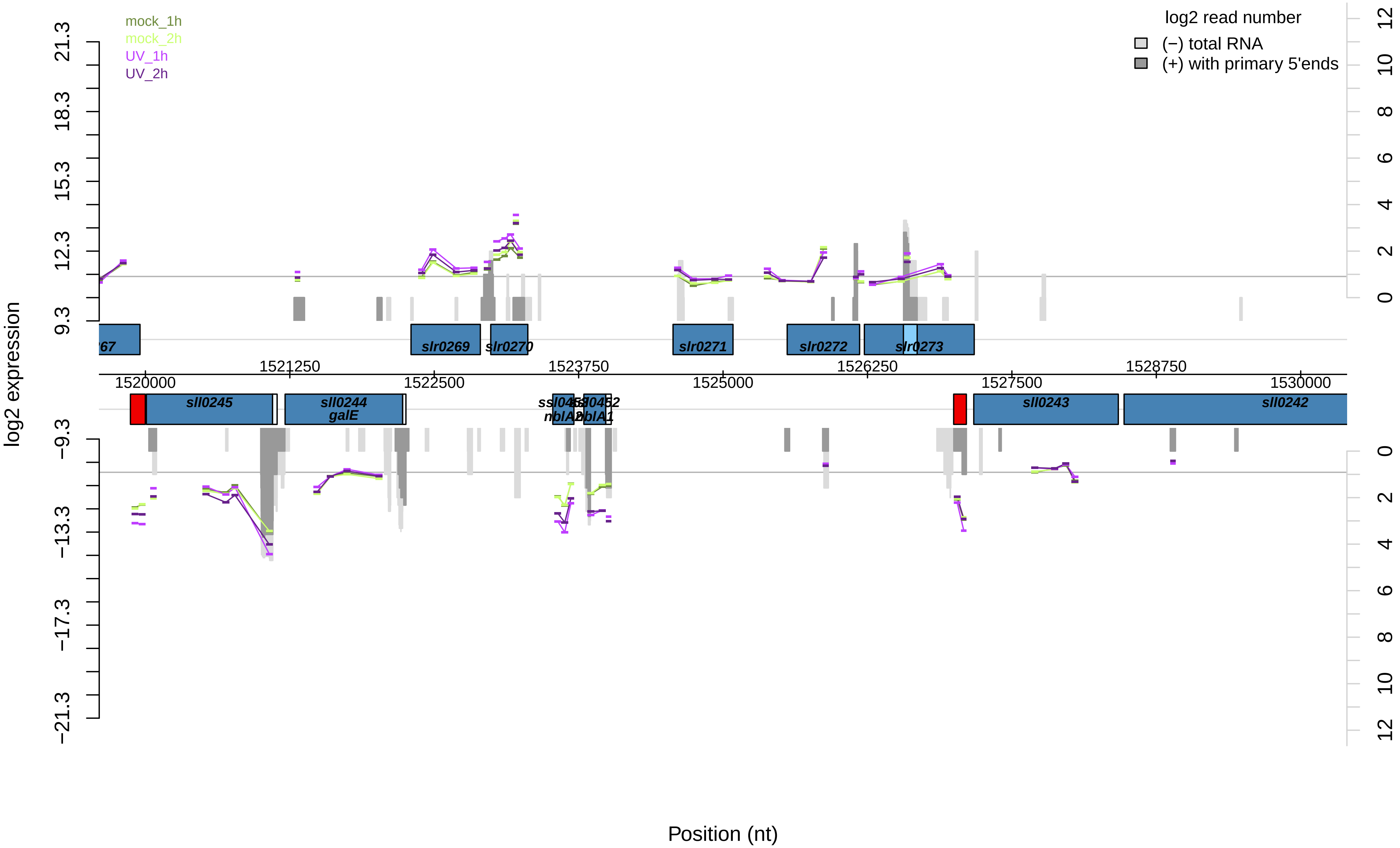
mock_1h
mock_2h
UV_1h
UV_2h
log2 read number
(−) total RNA
(+) with primary 5'ends
21.3
18.3
15.3
12.3
9.3
−9.3
−13.3
−17.3
−21.3
log2 expression
12
10
8
6
4
2
0
slr0269
slr0270
slr0271
slr0272
slr0273
1520000
1521250
1522500
1523750
1525000
1526250
1527500
1528750
1530000
sll0245
sll0244
galE
sll0243
sll0242
Position (nt)

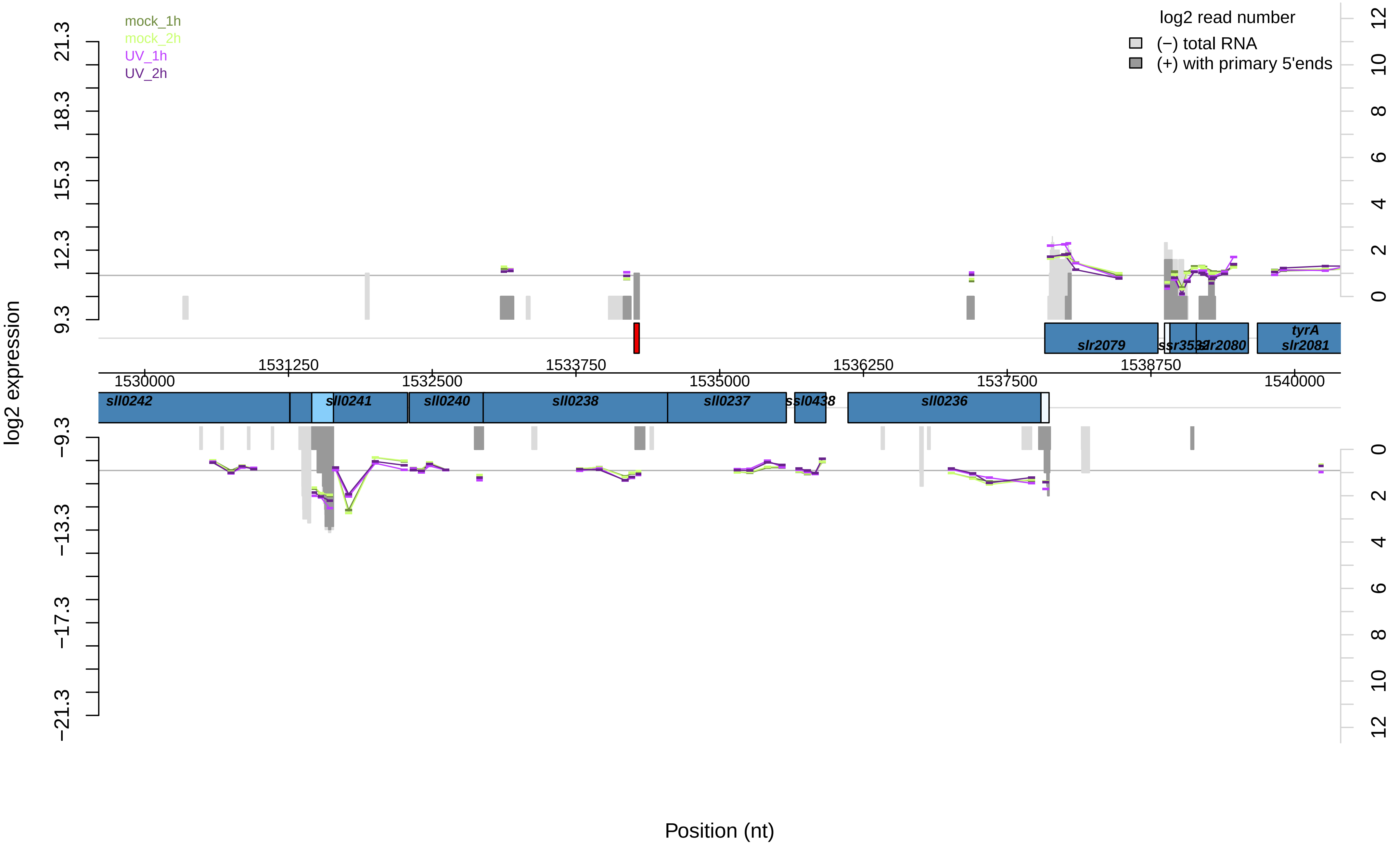
mock_1h
mock_2h
UV_1h
UV_2h
log2 read number
(−) total RNA
(+) with primary 5'ends
log2 expression
21.3
18.3
15.3
12.3
9.3
−9.3
−13.3
−17.3
−21.3
12
10
8
6
4
2
0
slr2079
ssr3532
slr2080
tyrA
slr2081
1530000
1531250
1532500
1533750
1535000
1536250
1537500
1538750
1540000
sll0242
sll0241
sll0240
sll0238
sll0237
ssl0438
sll0236
Position (nt)

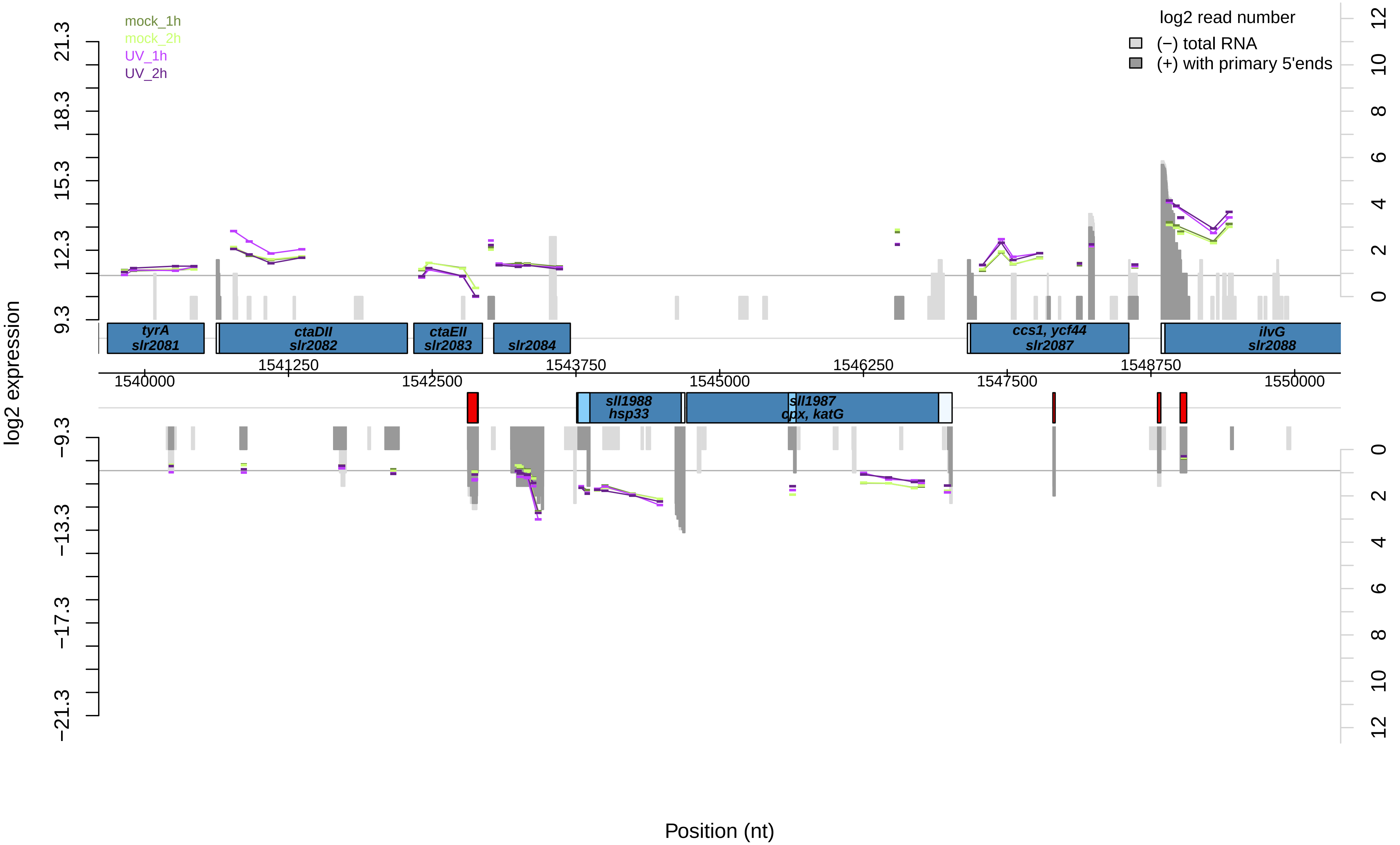

mock_1h
mock_2h
UV_1h
UV_2h
log2 read number
(−) total RNA
(+) with primary 5'ends
log2 expression
21.3
18.3
15.3
12.3
9.3
−9.3
−13.3
−17.3
−21.3
12
10
8
6
4
2
0
tyrA slr2081
ctaDII slr2082
ctaEII slr2083
slr2084
ccs1, ycf44 slr2087
ilvG slr2088
sll1988 hsp33
sll1987 cpx, katG
1541250
1543750
1546250
1548750
1540000
1542500
1545000
1547500
1550000
Position (nt)

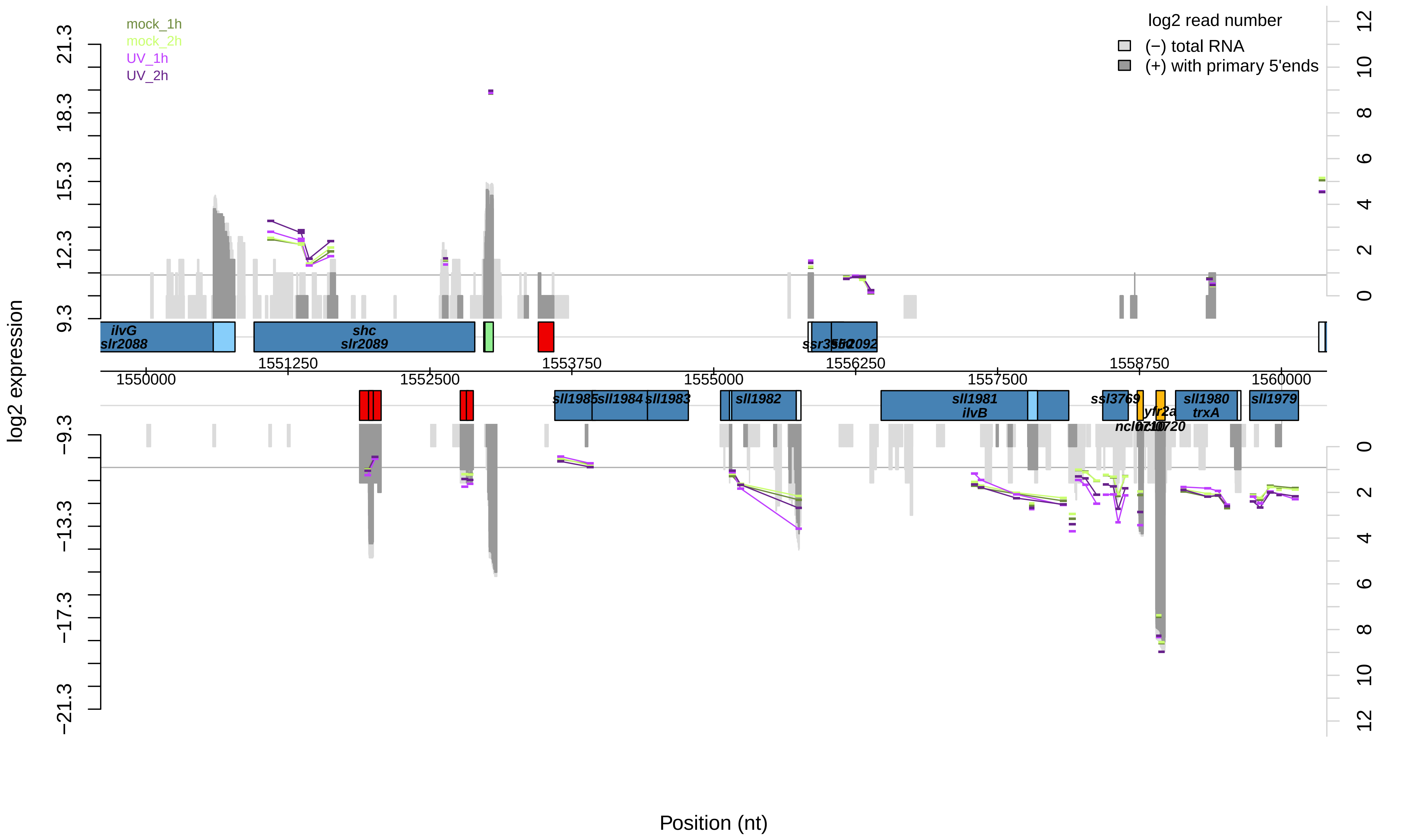

mock_1h
mock_2h
UV_1h
UV_2h
log2 read number
(−) total RNA
(+) with primary 5'ends
log2 expression
Position (nt)
21.3
18.3
15.3
12.3
9.3
−9.3
−13.3
−17.3
−21.3
0
2
4
6
8
10
12
ilvG
slr2088
shc
slr2089
1551250
1553750
1556250
1558750
1550000
1552500
1555000
1557500
1560000
sll1985
sll1984
sll1983
sll1982
sll1981
ilvB
ssl3769
yfr2a
sll1980
trxA
sll1979

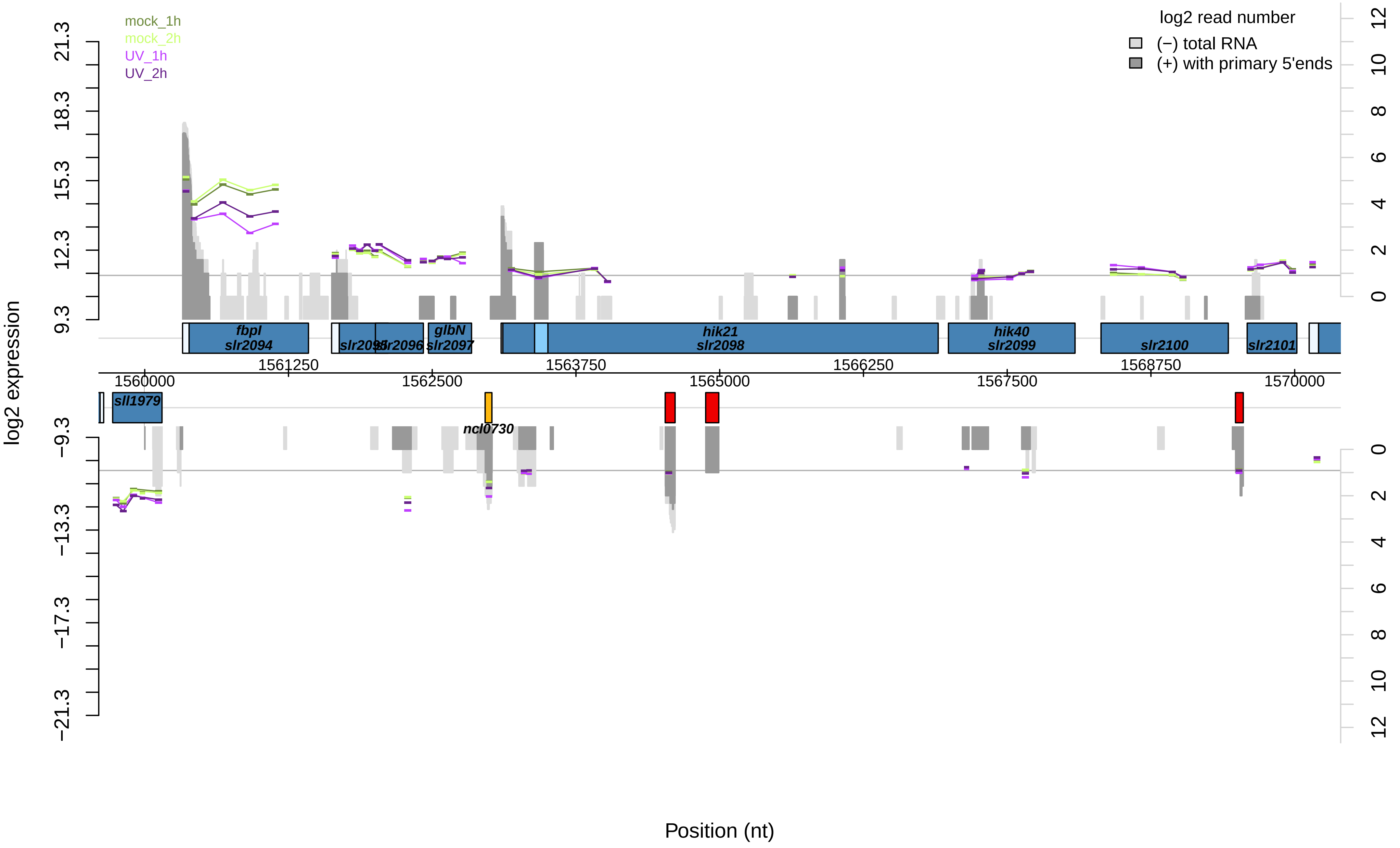

mock_1h
mock_2h
UV_1h
UV_2h
log2 read number
(−) total RNA
(+) with primary 5'ends
log2 expression
21.3
18.3
15.3
12.3
9.3
−9.3
−13.3
−17.3
−21.3
fbpI
slr2094
slr2095
slr2096
glbN
slr2097
hik21
slr2098
hik40
slr2099
slr2100
slr2101
sll1979
ncl0730
1560000
1561250
1562500
1563750
1565000
1566250
1567500
1568750
1570000
Position (nt)

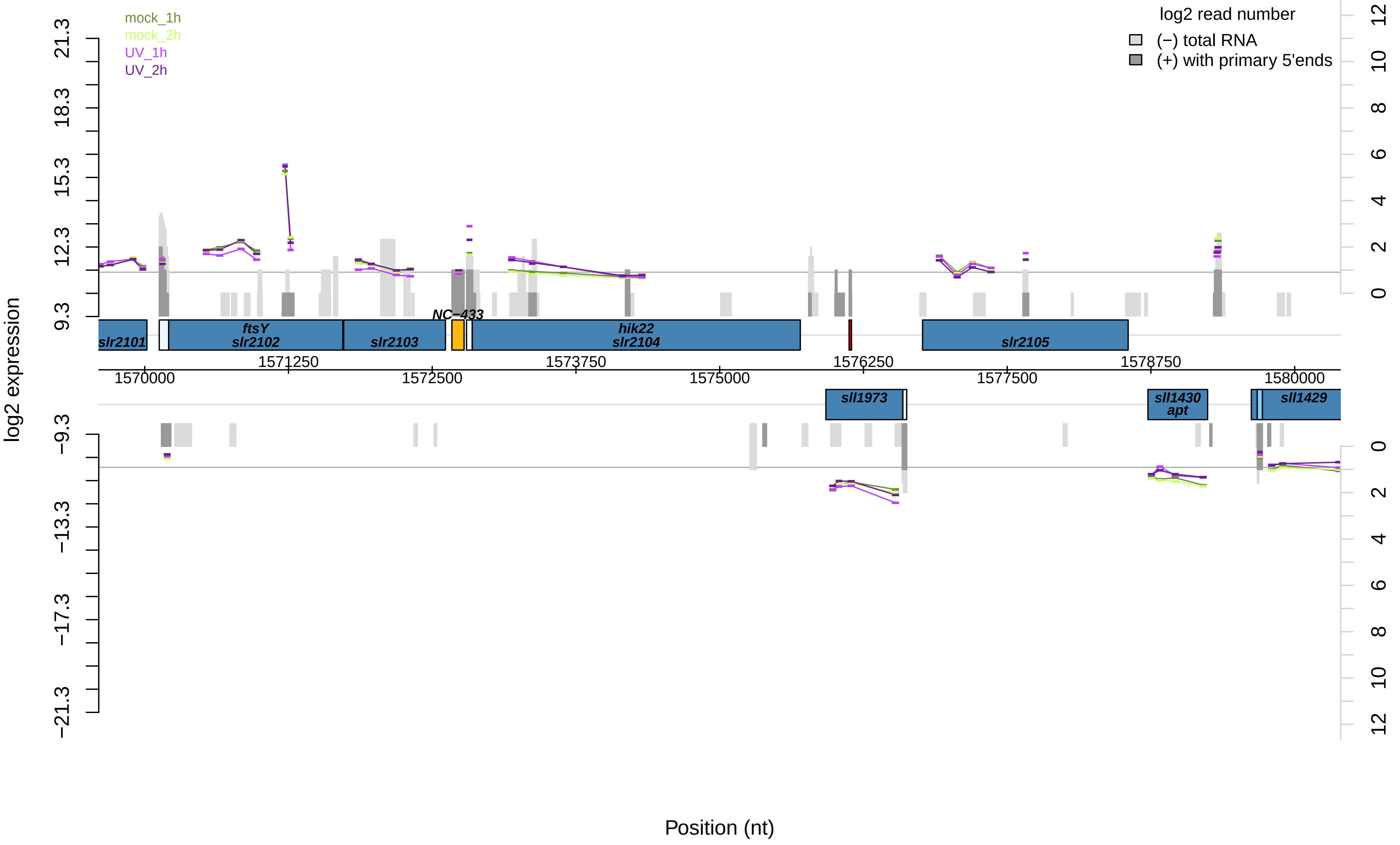

mock_1h
mock_2h
UV_1h
UV_2h
log2 read number
(−) total RNA
(+) with primary 5'ends
log2 expression
21.3
18.3
15.3
12.3
9.3
−9.3
−13.3
−17.3
−21.3
12
10
8
6
4
2
0
slr2101
ftsY
slr2102
slr2103
NC−433
hik22
slr2104
slr2105
sll1973
sll1430
apt
sll1429
1570000
1571250
1572500
1573750
1575000
1576250
1577500
1578750
1580000
Position (nt)

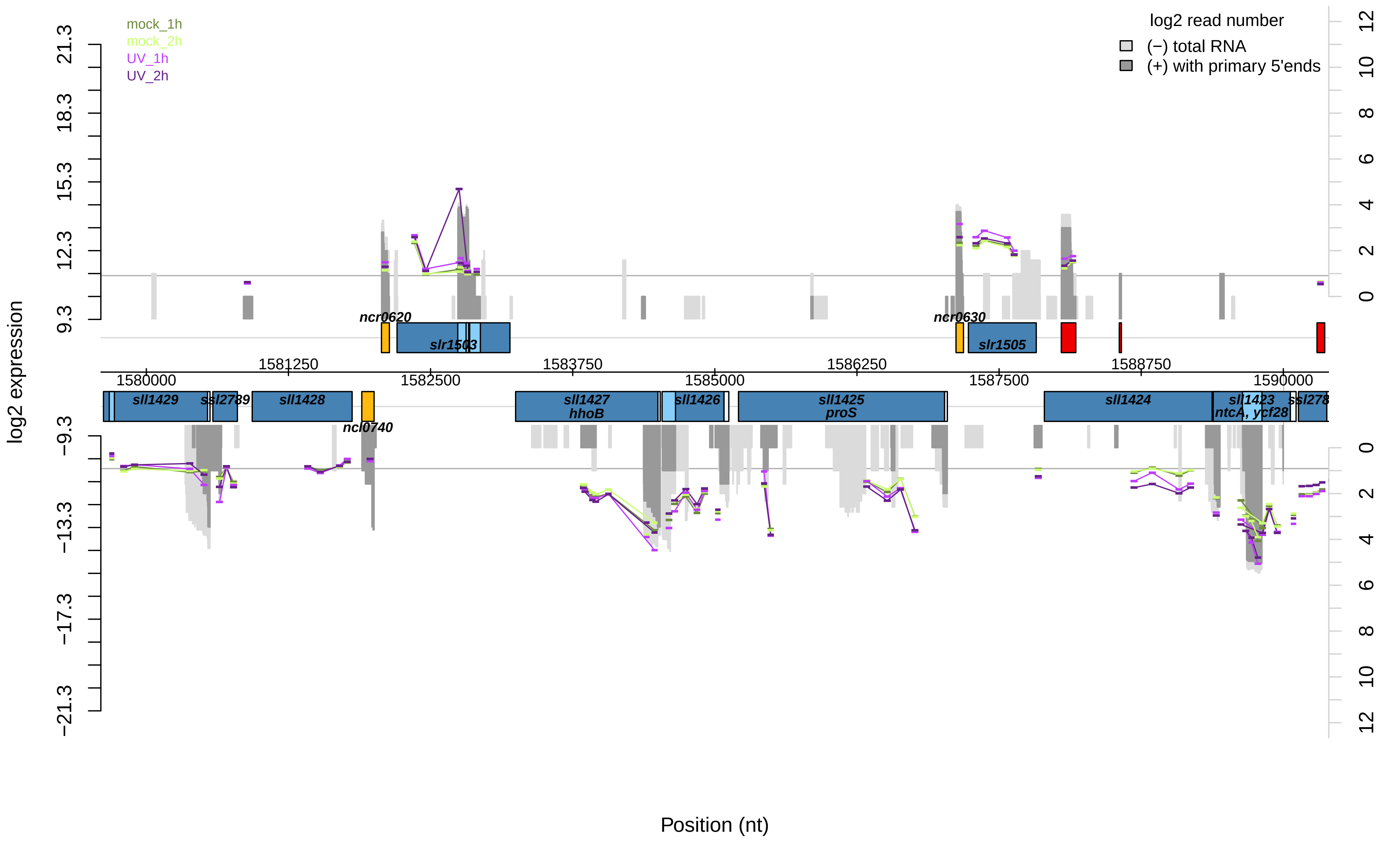

mock_1h
mock_2h
UV_1h
UV_2h
log2 read number
(−) total RNA
(+) with primary 5'ends
log2 expression
21.3
18.3
15.3
12.3
9.3
−9.3
−13.3
−17.3
−21.3
12
10
8
6
4
2
0
ncr0620
slr1503
ncr0630
slr1505
1580000
1581250
1582500
1583750
1585000
1586250
1587500
1588750
1590000
sll1429
ssl2789
sll1428
ncl0740
sll1427
hhoB
sll1426
sll1425
proS
sll1424
sll1423
ntcA, ycf28
ssl278
Position (nt)

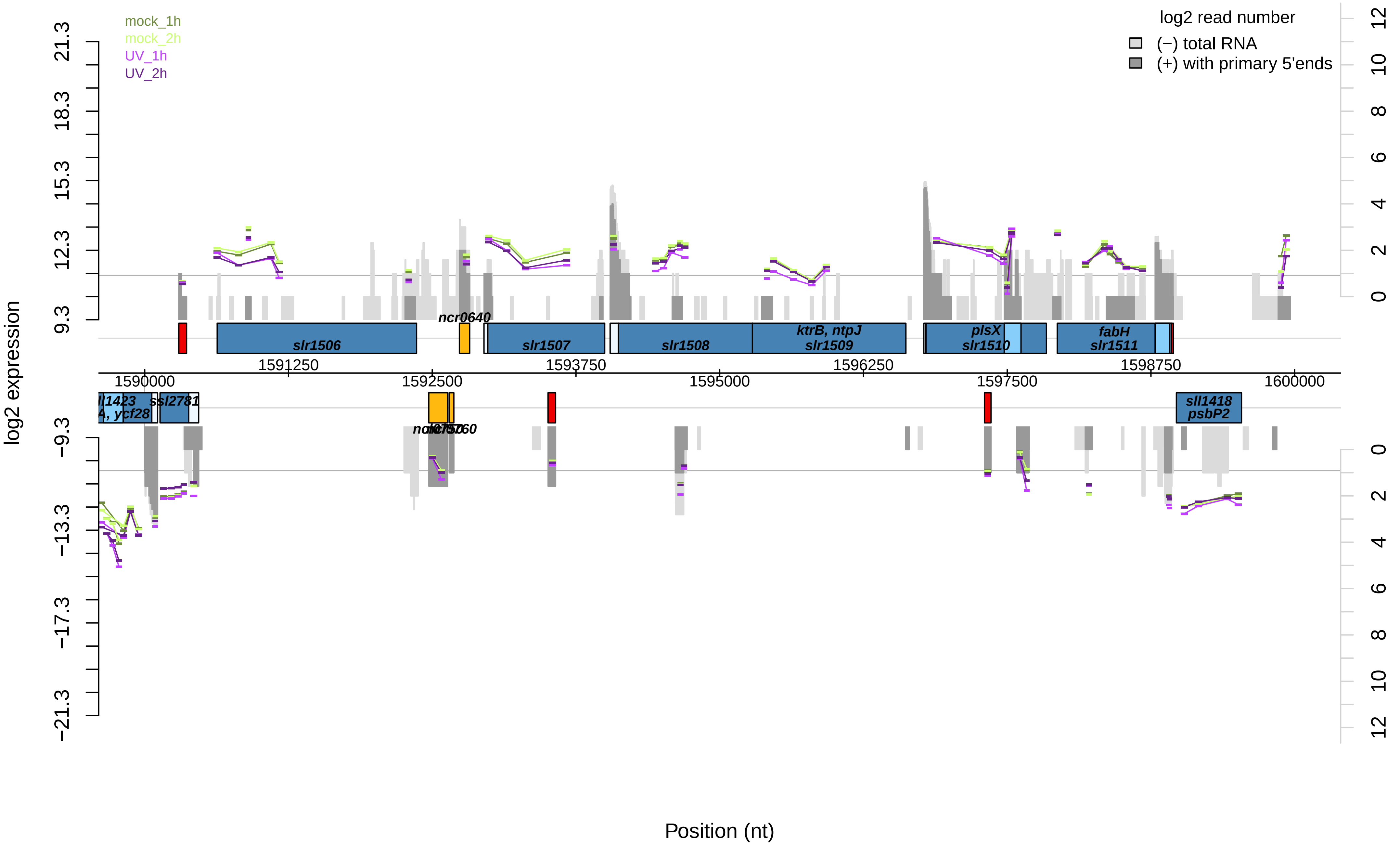

mock_1h
mock_2h
UV_1h
UV_2h
log2 read number
(−) total RNA
(+) with primary 5'ends
log2 expression
Position (nt)
slr1506
ncr0640
slr1507
slr1508
ktrB, ntpJ
slr1509
plsX
slr1510
fabH
slr1511
ssl2781
sll1418
psbP2
1590000
1591250
1592500
1593750
1595000
1596250
1597500
1598750
1600000

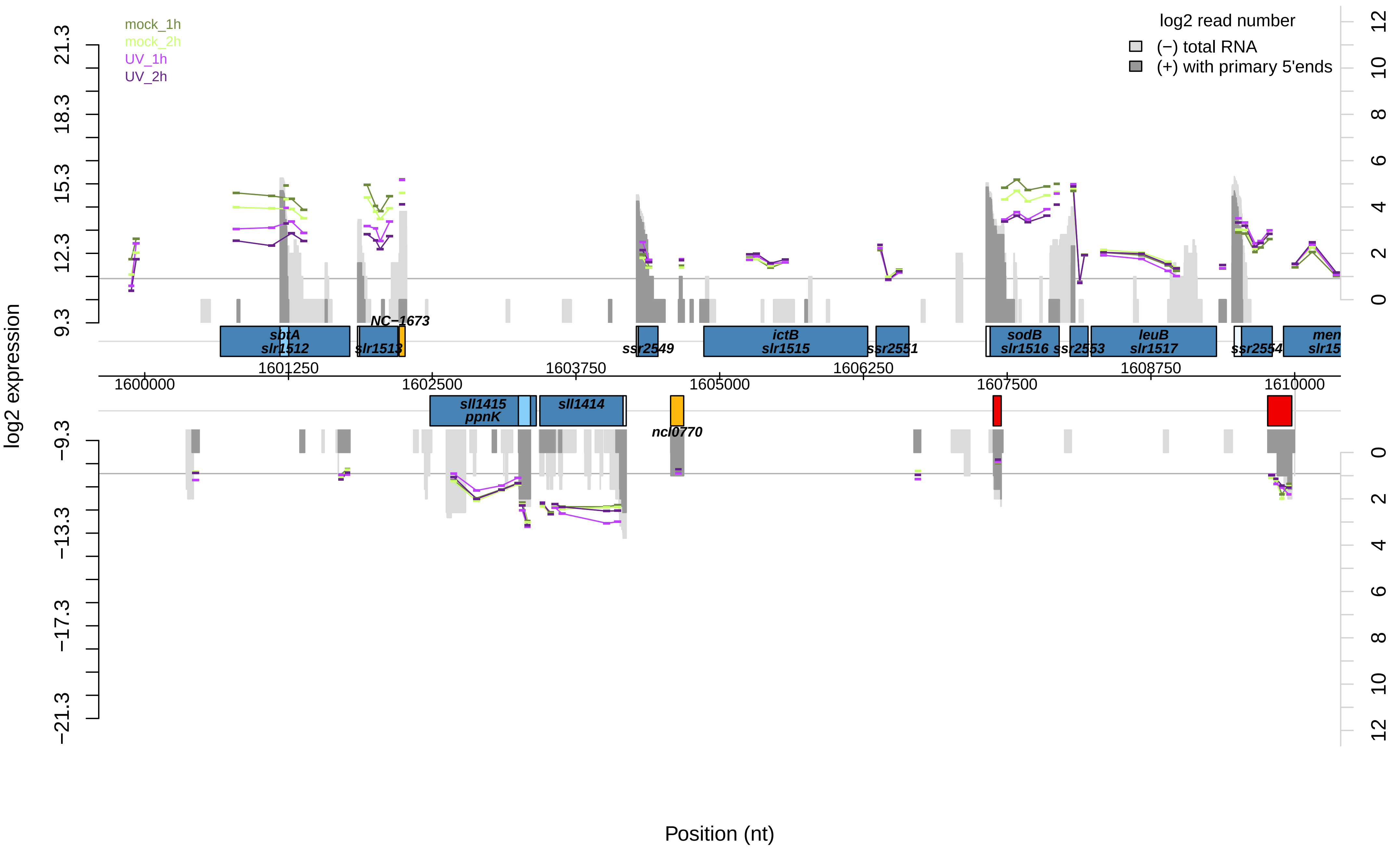

mock_1h
mock_2h
UV_1h
UV_2h
log2 read number
(−) total RNA
(+) with primary 5'ends
log2 expression
Position (nt)
21.3
18.3
15.3
12.3
9.3
−9.3
−13.3
−17.3
−21.3
12
10
8
6
4
2
0
sbtA
slr1512
slr1513
NC−1673
ssr2549
ictB
slr1515
ssr2551
sodB
slr1516
ssr2553
leuB
slr1517
ssr2554
sll1415
ppnK
sll1414
ncl0770
1600000
1601250
1602500
1603750
1605000
1606250
1607500
1608750
1610000

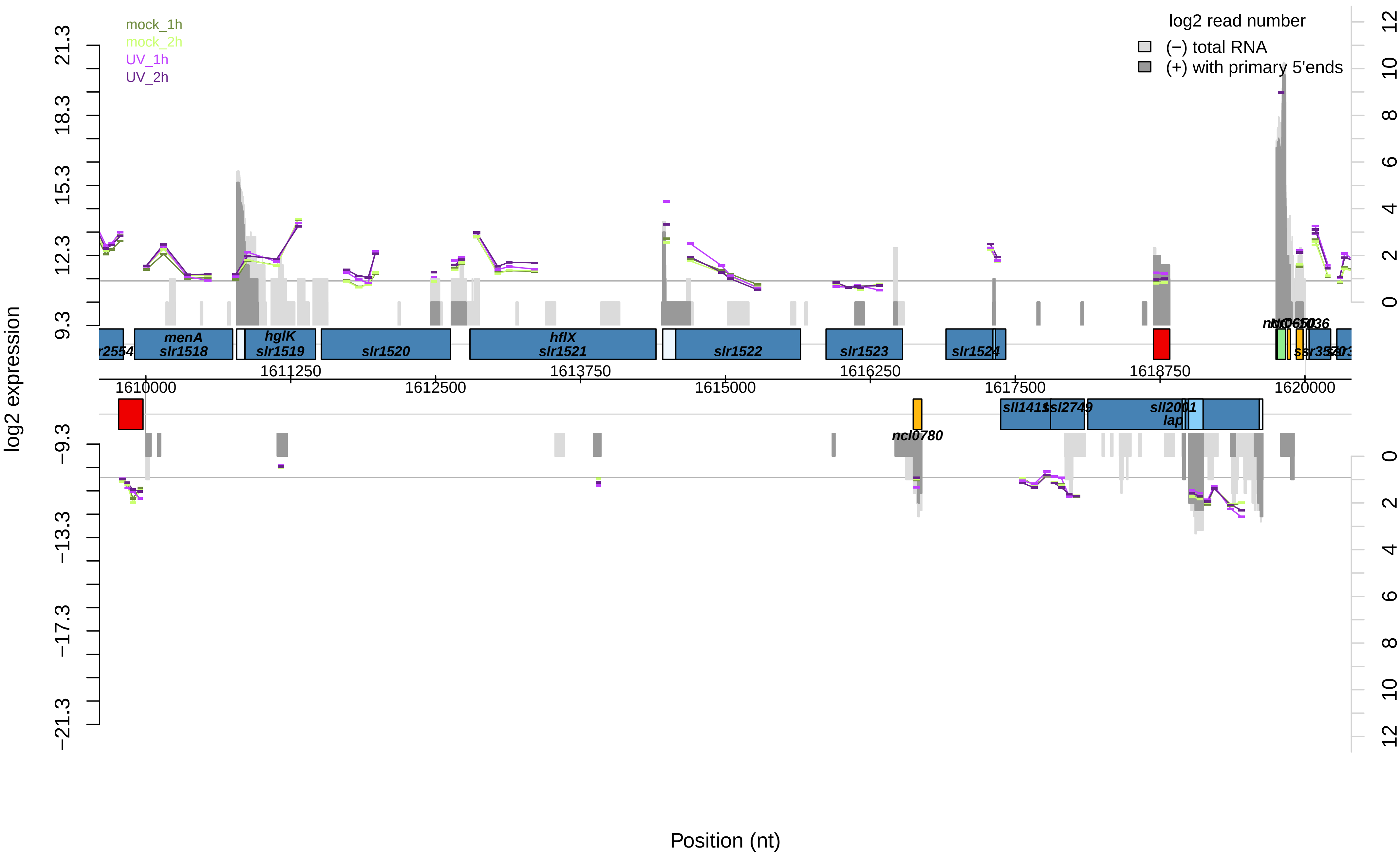

mock_1h
mock_2h
UV_1h
UV_2h
log2 read number
(−) total RNA
(+) with primary 5'ends
log2 expression
21.3
18.3
15.3
12.3
9.3
−9.3
−13.3
−17.3
−21.3
12
10
8
6
4
2
0
menA
slr1518
hglK
slr1519
slr1520
hflX
slr1521
slr1522
slr1523
slr1524
1611250
1613750
1616250
1618750
1610000
1612500
1615000
1617500
1620000
ncl0780
sll2749
sll2001
lap
Position (nt)

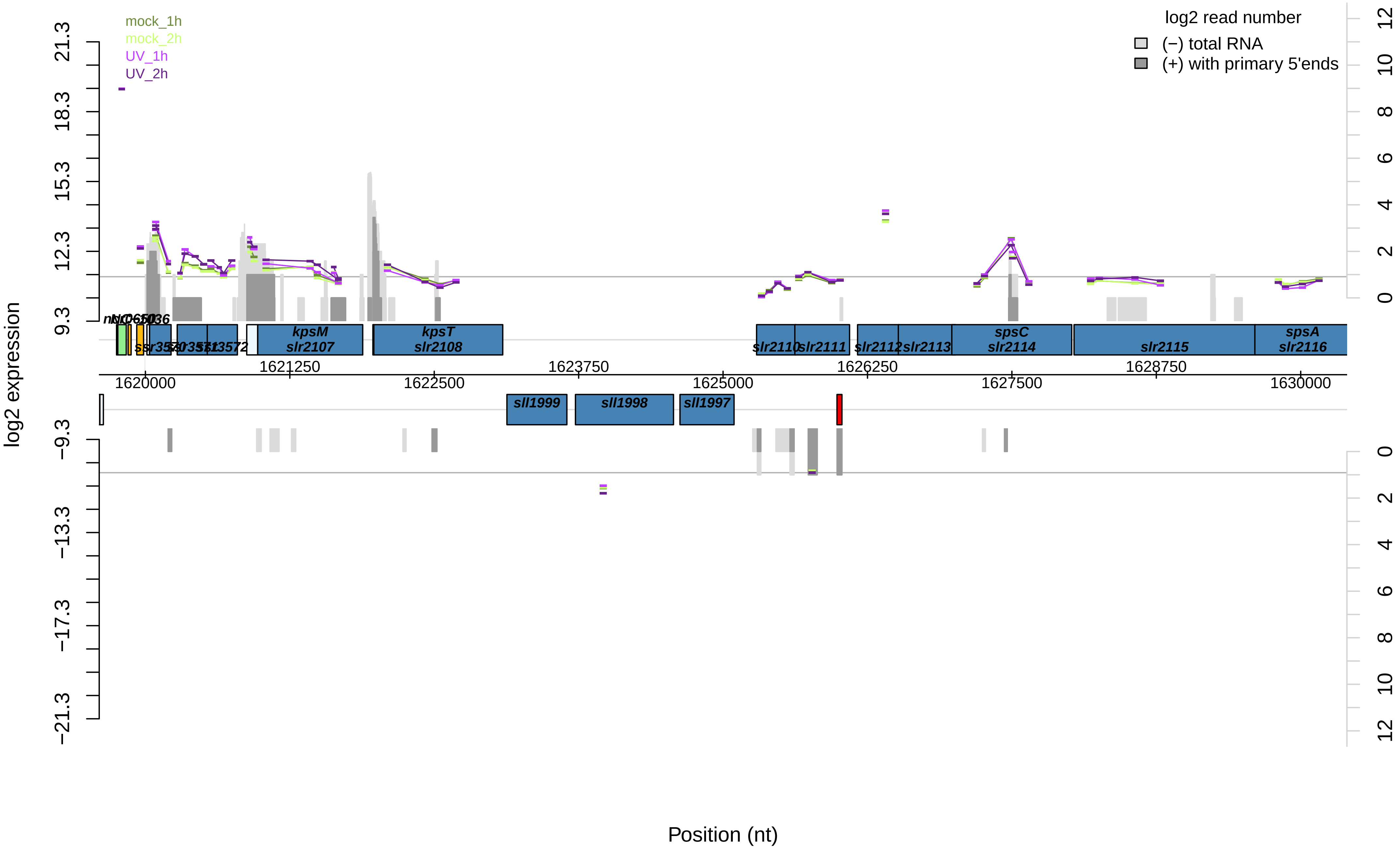

mock_1h
mock_2h
UV_1h
UV_2h
log2 read number
(−) total RNA
(+) with primary 5'ends
log2 expression
Position (nt)
21.3
18.3
15.3
12.3
9.3
−9.3
−13.3
−17.3
−21.3
1620000
1621250
1622500
1623750
1625000
1626250
1627500
1628750
1630000
kpsM
slr2107
kpsT
slr2108
slr2110
slr2111
slr2112
slr2113
spsC
slr2114
slr2115
spsA
slr2116
sll1999
sll1998
sll1997

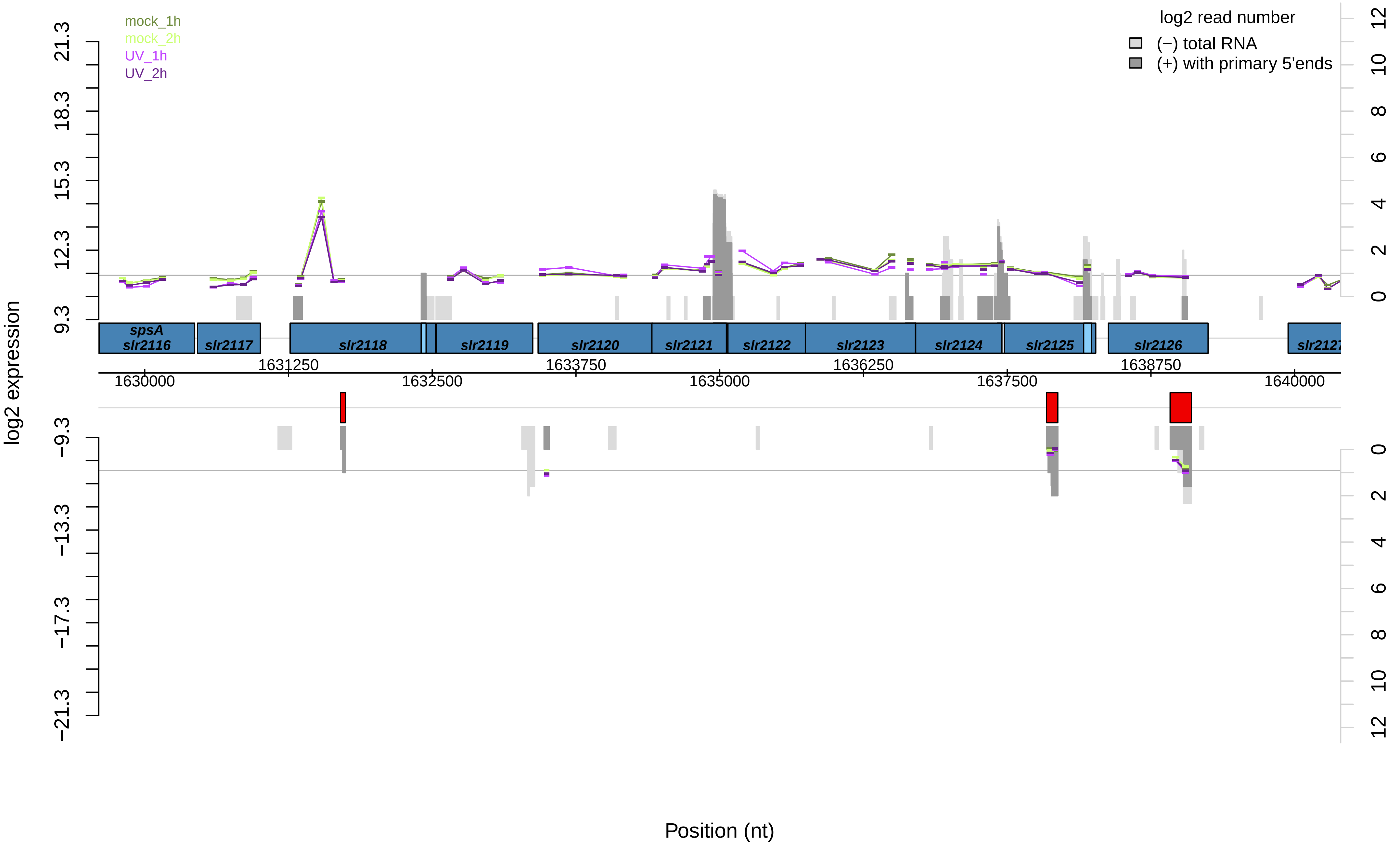

mock_1h
mock_2h
UV_1h
UV_2h
log2 read number
(−) total RNA
(+) with primary 5'ends
log2 expression
21.3
18.3
15.3
12.3
9.3
−9.3
−13.3
−17.3
−21.3
12
10
8
6
4
2
0
spsA
slr2116
slr2117
slr2118
slr2119
slr2120
slr2121
slr2122
slr2123
slr2124
slr2125
slr2126
1631250
1633750
1636250
1638750
1630000
1632500
1635000
1637500
1640000
Position (nt)

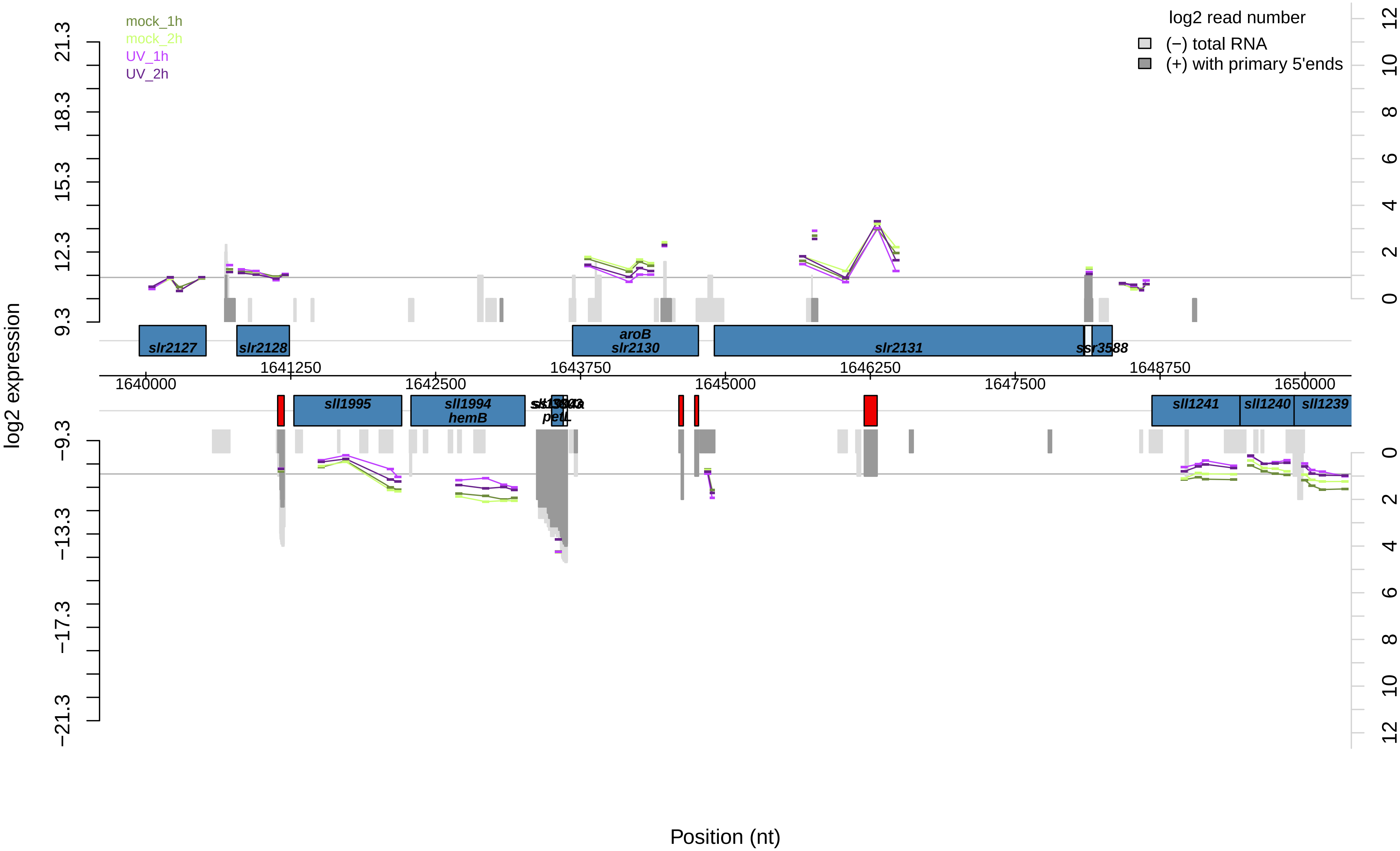

mock_1h
mock_2h
UV_1h
UV_2h
log2 read number
(−) total RNA
(+) with primary 5'ends
log2 expression
21.3
18.3
15.3
12.3
9.3
−9.3
−13.3
−17.3
−21.3
slr2127
slr2128
aroB
slr2130
slr2131
ssr3588
sll1995
sll1994
hemB
petL
sll1241
sll1240
sll1239
1640000
1641250
1642500
1643750
1645000
1646250
1647500
1648750
1650000
Position (nt)

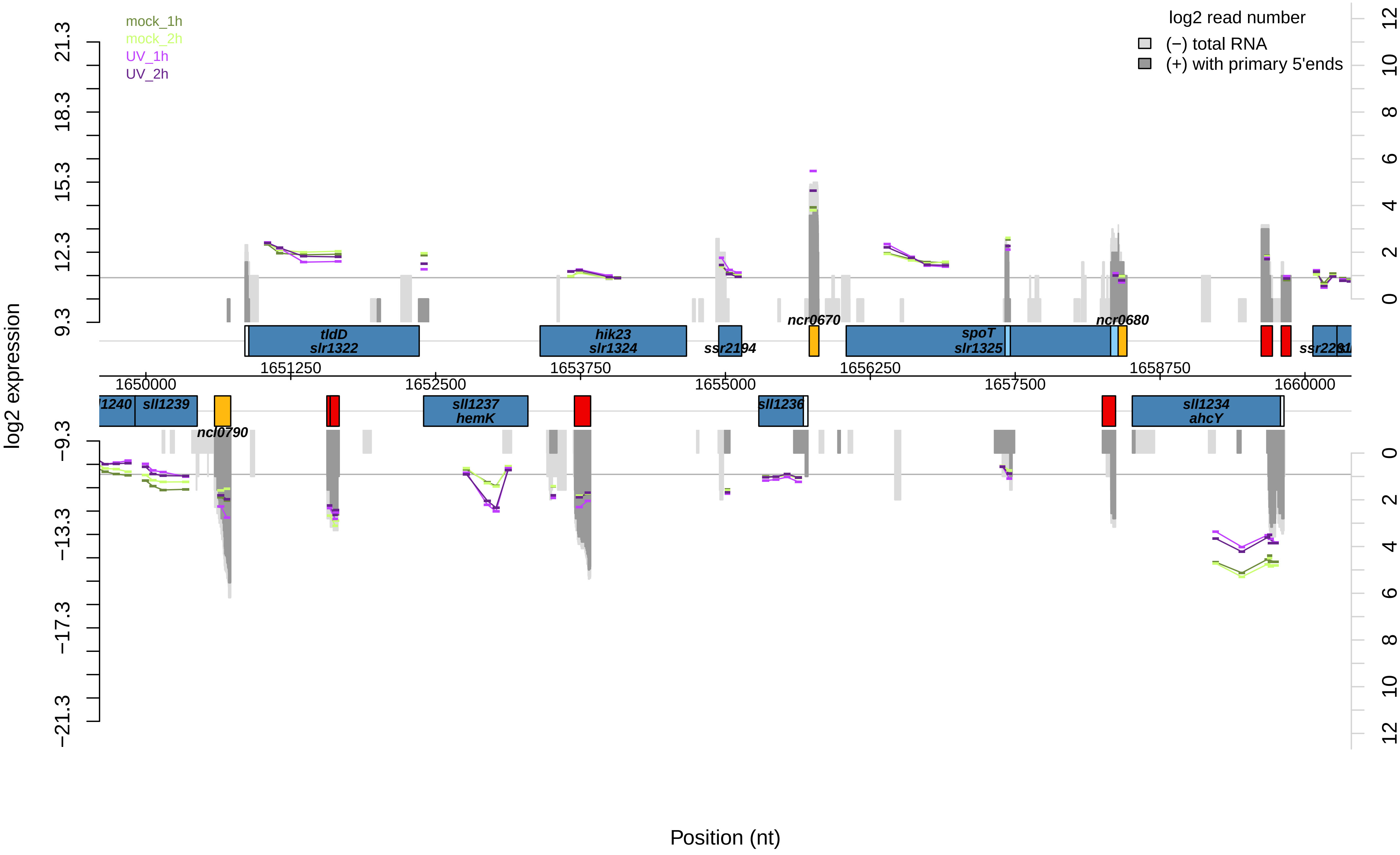
mock_1h
mock_2h
UV_1h
UV_2h
log2 read number
(−) total RNA
(+) with primary 5'ends
log2 expression
Position (nt)
21.3
18.3
15.3
12.3
9.3
−9.3
−13.3
−17.3
−21.3
12
10
8
6
4
2
0
1650000
1651250
1652500
1653750
1655000
1656250
1657500
1658750
1660000
tldD
slr1322
hik23
slr1324
ssr2194
ncr0670
spoT
slr1325
ncr0680
sll1239
ncl0790
sll1237
hemK
sll1236
sll1234
ahcY

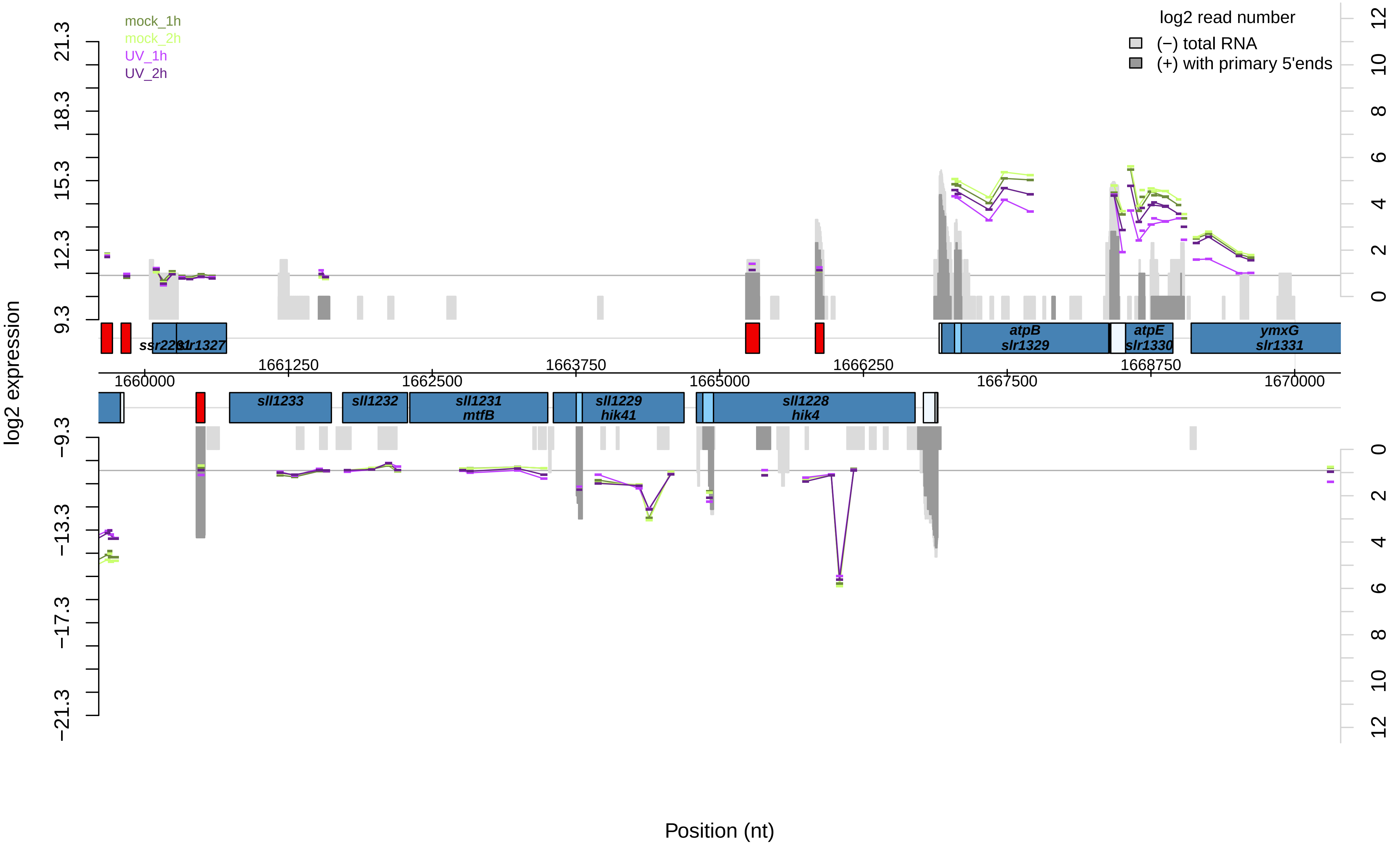
mock_1h
mock_2h
UV_1h
UV_2h
log2 read number
(−) total RNA
(+) with primary 5'ends
log2 expression
21.3
18.3
15.3
12.3
9.3
−9.3
−13.3
−17.3
−21.3
12
10
8
6
4
2
0
ssr2281
slr1327
atpB
slr1329
atpE
slr1330
ymxG
slr1331
1661250
1663750
1666250
1668750
1660000
1662500
1665000
1667500
1670000
sll1233
sll1232
sll1231
mtfB
sll1229
hik41
sll1228
hik4
Position (nt)

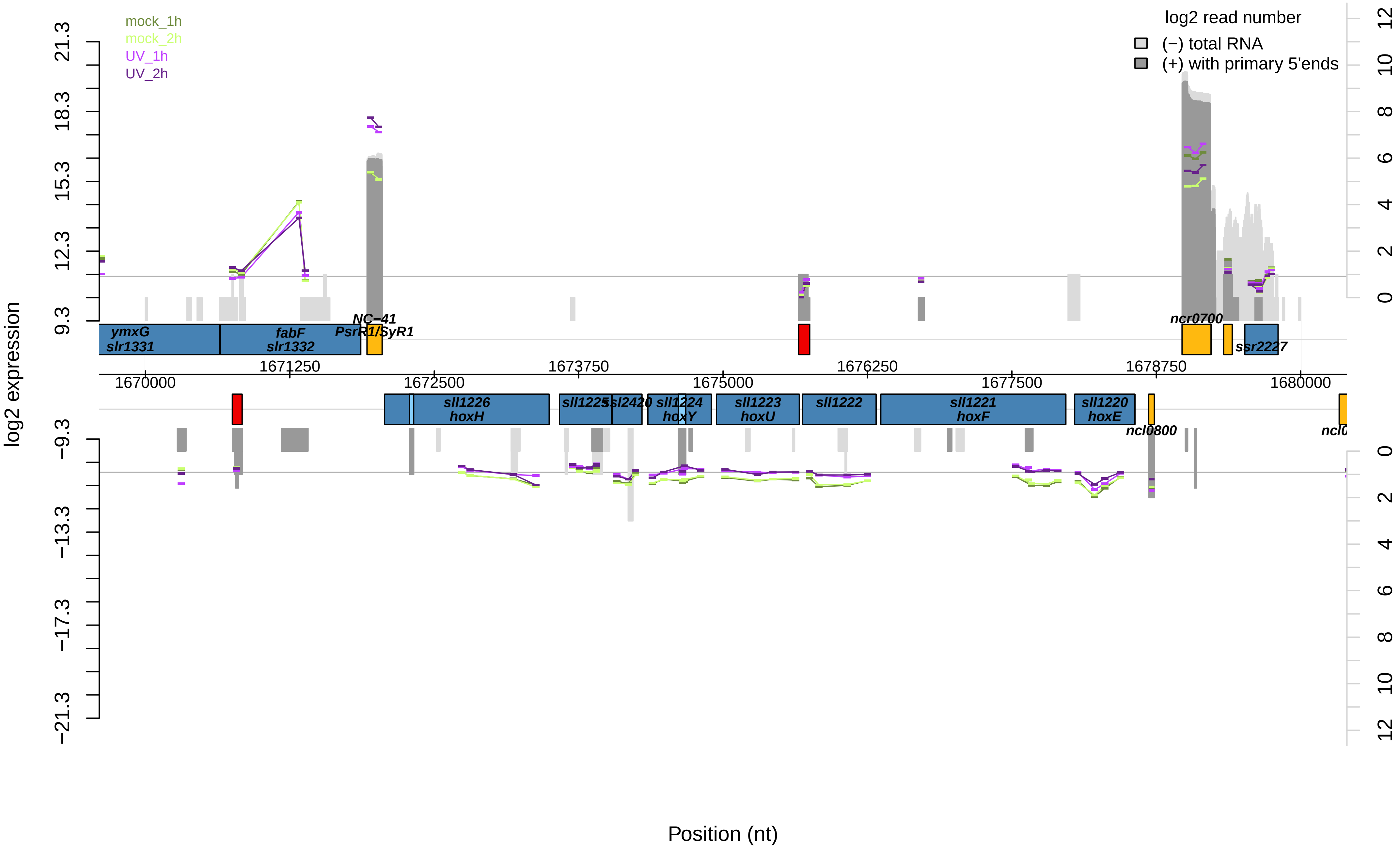

mock_1h
mock_2h
UV_1h
UV_2h
log2 read number
(−) total RNA
(+) with primary 5'ends
log2 expression
21.3
18.3
15.3
12.3
9.3
−9.3
−13.3
−17.3
−21.3
12
10
8
6
4
2
0
ymxG
slr1331
fabF
slr1332
NC−41
PsrR1/SyR1
ncr0700
ssr2227
1670000
1671250
1672500
1673750
1675000
1676250
1677500
1678750
1680000
sll1226
hoxH
sll1225
ssl2420
sll1224
hoxY
sll1223
hoxU
sll1222
sll1221
hoxF
sll1220
hoxE
ncl0800
Position (nt)

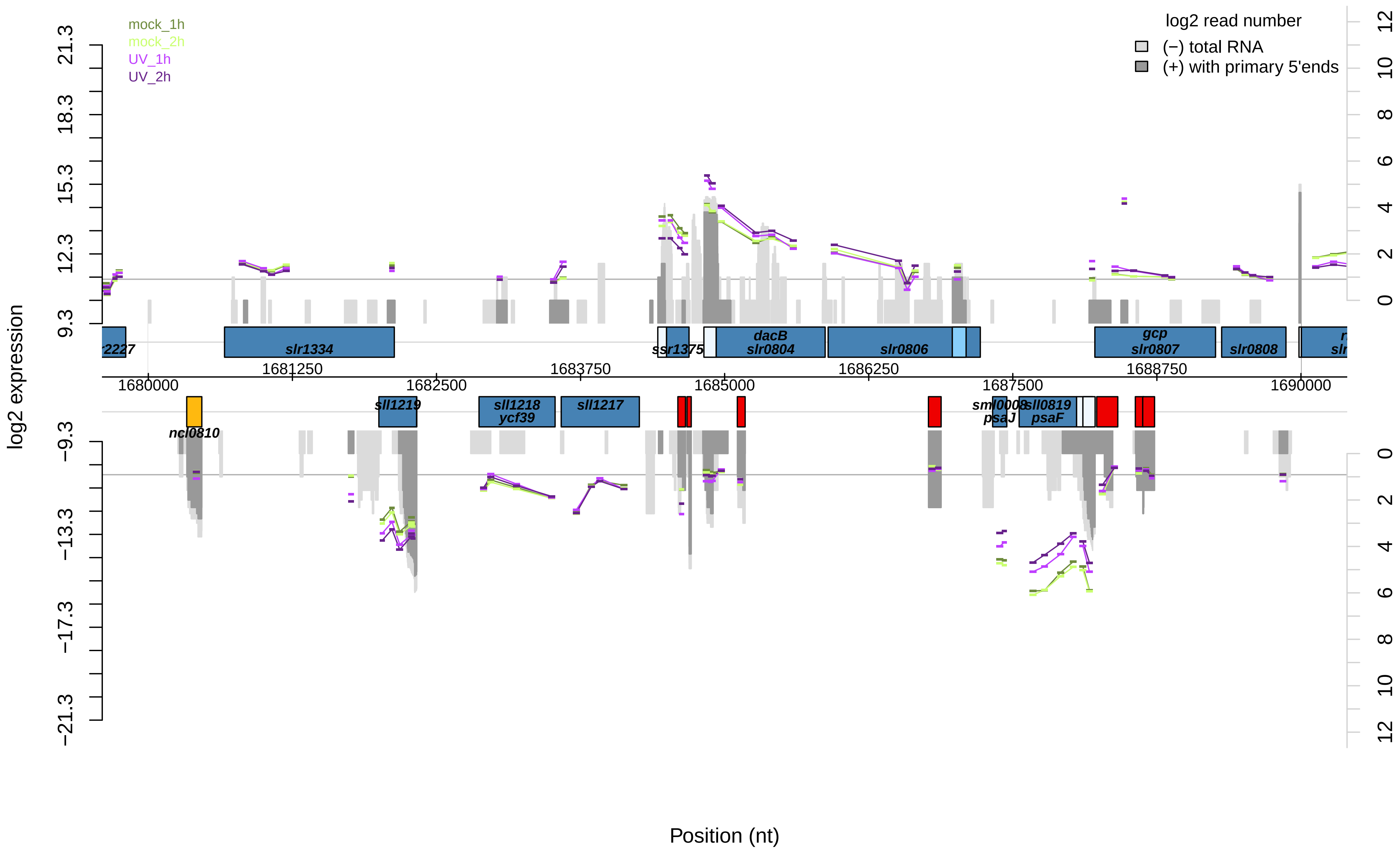

mock_1h
mock_2h
UV_1h
UV_2h
log2 read number
(−) total RNA
(+) with primary 5'ends
log2 expression
21.3
18.3
15.3
12.3
9.3
−9.3
−13.3
−17.3
−21.3
12
10
8
6
4
2
0
r2227
slr1334
ssr1375
dacB
slr0804
slr0806
gcp
slr0807
slr0808
1681250
1683750
1686250
1688750
1680000
1682500
1685000
1687500
1690000
ncl0810
sll1219
sll1218
ycf39
sll1217
sml0008
psaJ
sll0819
psaF
Position (nt)

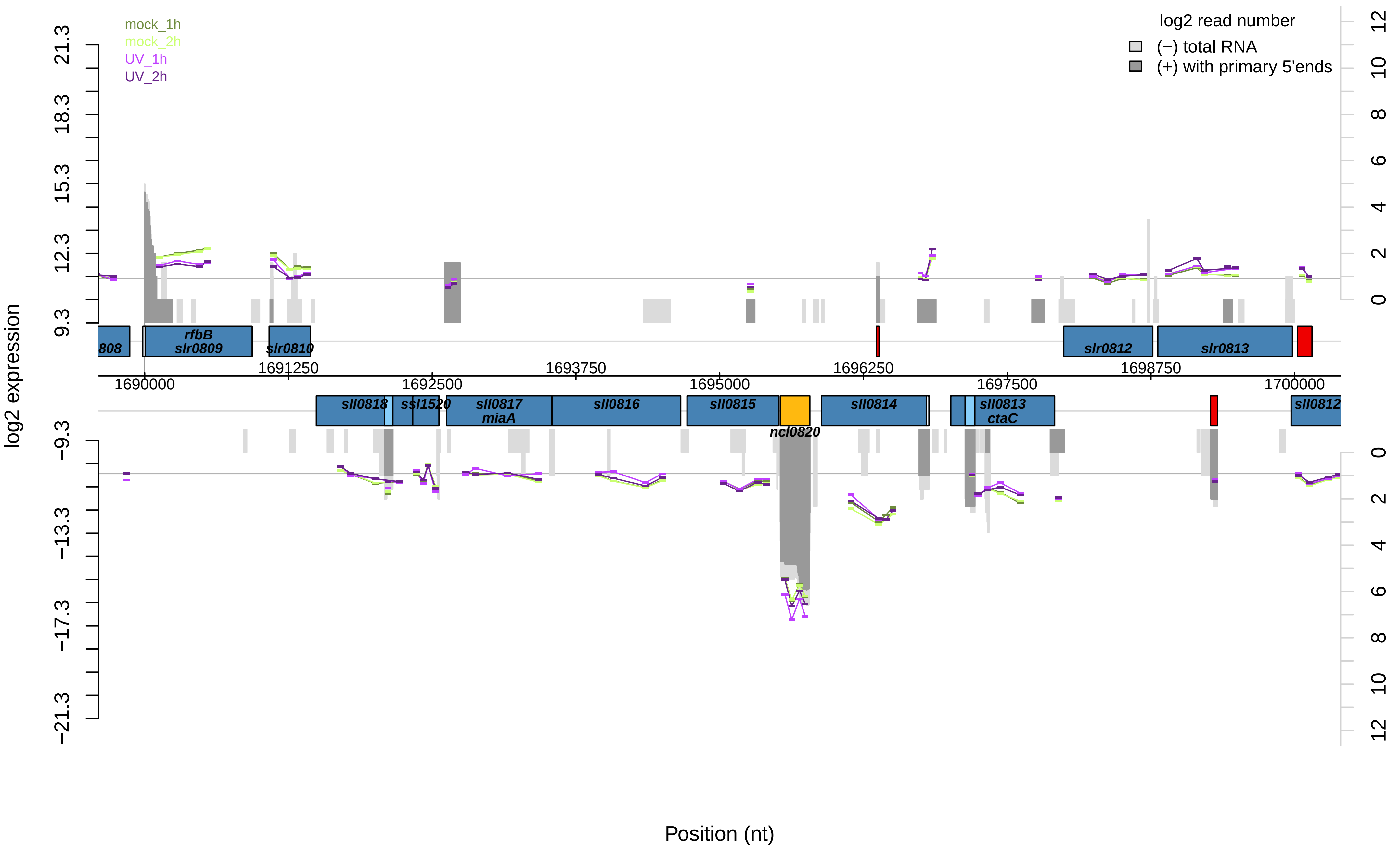

mock_1h
mock_2h
UV_1h
UV_2h
log2 read number
(−) total RNA
(+) with primary 5'ends
log2 expression
21.3
18.3
15.3
12.3
9.3
−9.3
−13.3
−17.3
−21.3
12
10
8
6
4
2
0
808
rfbB
slr0809
slr0810
slr0812
slr0813
1690000
1691250
1692500
1693750
1695000
1696250
1697500
1698750
1700000
sll0818
ssl1520
sll0817
miaA
sll0816
sll0815
ncl0820
sll0814
sll0813
ctaC
sll0812
Position (nt)

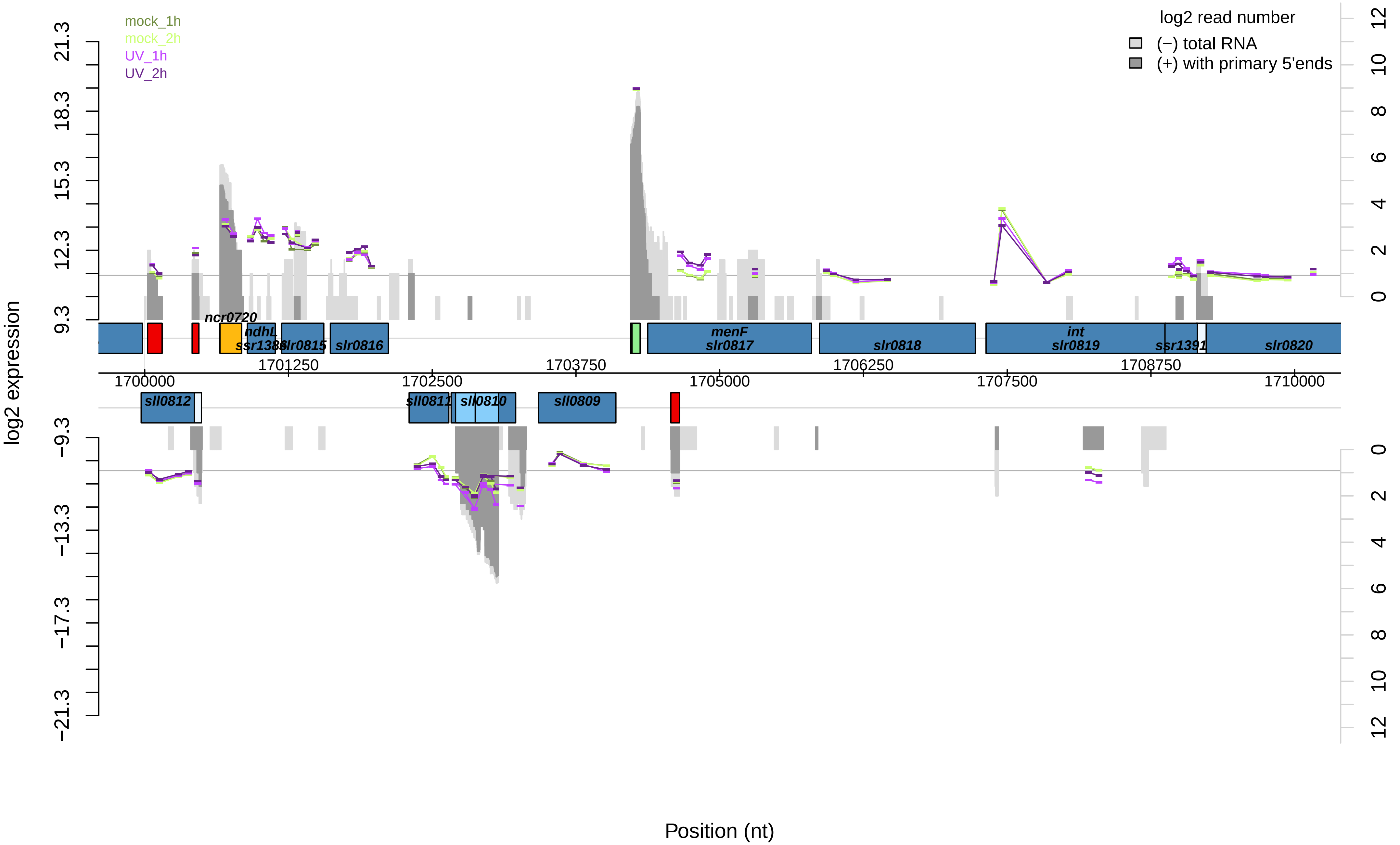

mock_1h
mock_2h
UV_1h
UV_2h
log2 read number
(−) total RNA
(+) with primary 5'ends
log2 expression
21.3
18.3
15.3
12.3
9.3
−9.3
−13.3
−17.3
−21.3
12
10
8
6
4
2
0
ncr0720
ndhL
ssr1386
slr0815
slr0816
menF
slr0817
slr0818
int
slr0819
ssr1391
slr0820
1701250
1703750
1706250
1708750
1700000
1702500
1705000
1707500
1710000
sll0812
sll0811
sll0810
sll0809
Position (nt)

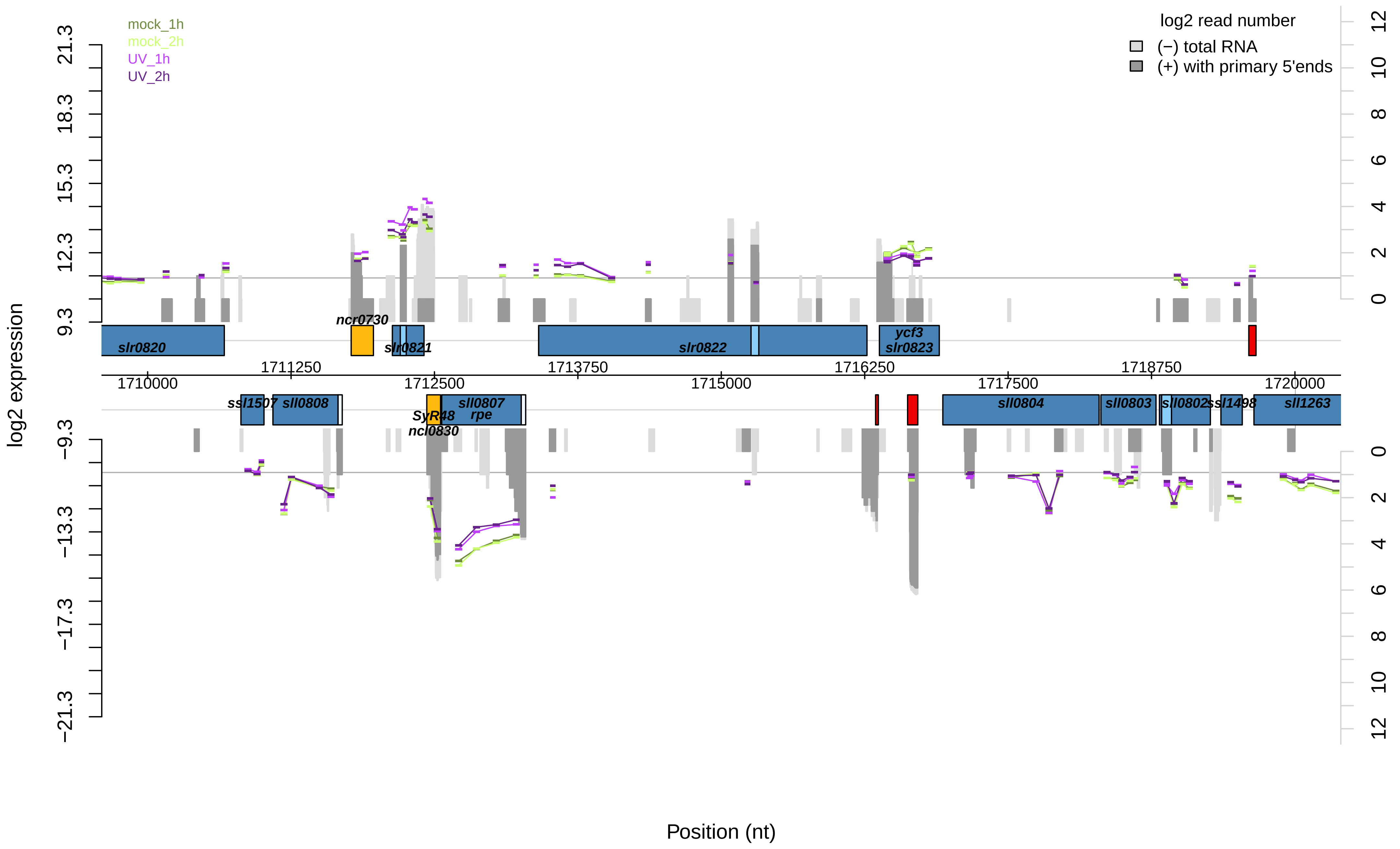

mock_1h
mock_2h
UV_1h
UV_2h
log2 read number
(−) total RNA
(+) with primary 5'ends
log2 expression
21.3
18.3
15.3
12.3
9.3
−9.3
−13.3
−17.3
−21.3
12
10
8
6
4
2
0
slr0820
ncr0730
slr0821
slr0822
ycf3
slr0823
1711250
1712500
1713750
1715000
1716250
1717500
1718750
1710000
1720000
ssl1507
sll0808
SyR48
ncl0830
sll0807
rpe
sll0804
sll0803
sll0802
ssl1498
sll1263
Position (nt)

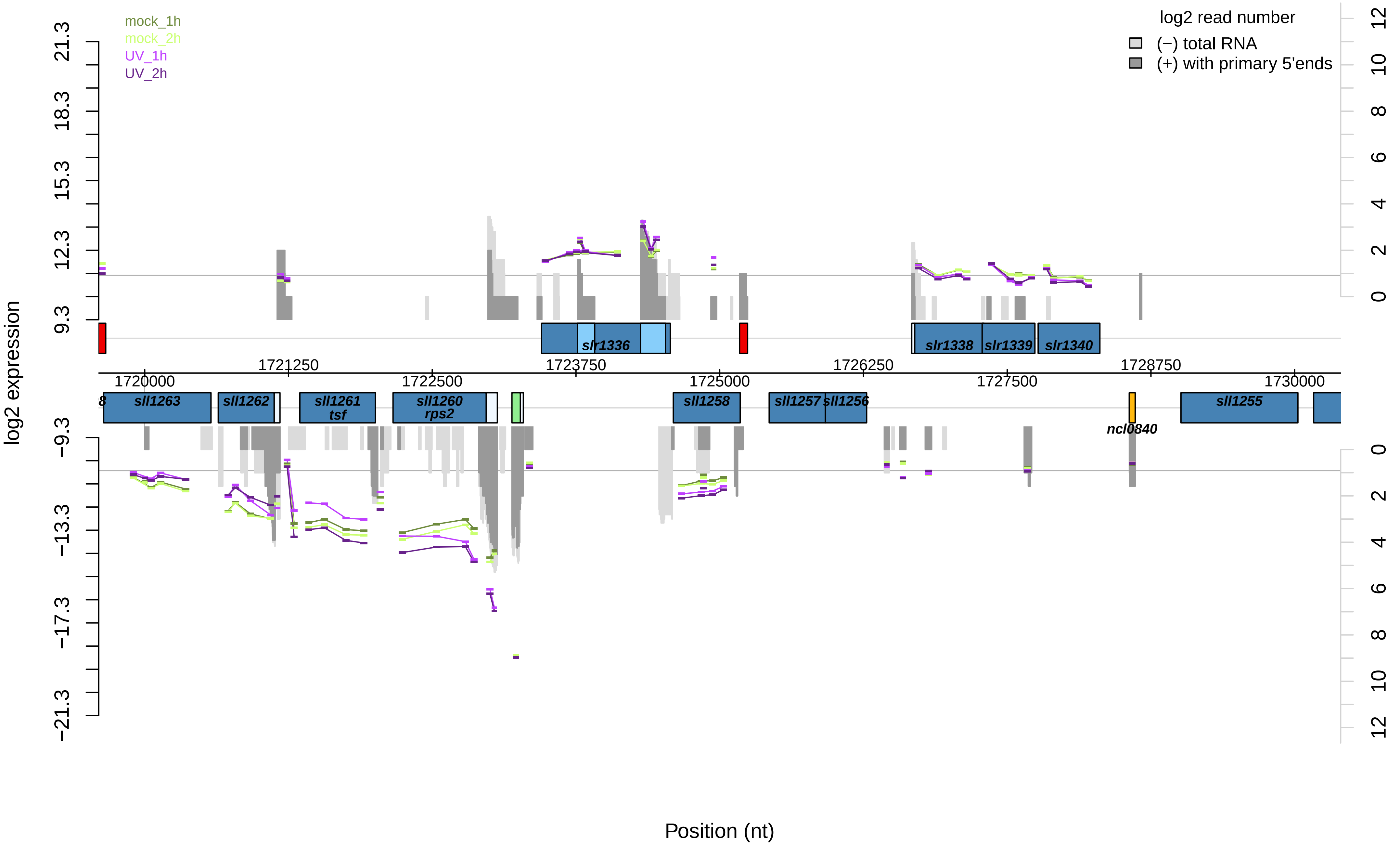

mock_1h
mock_2h
UV_1h
UV_2h
log2 read number
(−) total RNA
(+) with primary 5'ends
log2 expression
21.3
18.3
15.3
12.3
9.3
−9.3
−13.3
−17.3
−21.3
12
10
8
6
4
2
0
slr1336
slr1338
slr1339
slr1340
1721250
1723750
1726250
1728750
1720000
1722500
1725000
1727500
1730000
sll1263
sll1262
sll1261
tsf
sll1260
rps2
sll1258
sll1257
sll1256
sll1255
ncl0840
Position (nt)

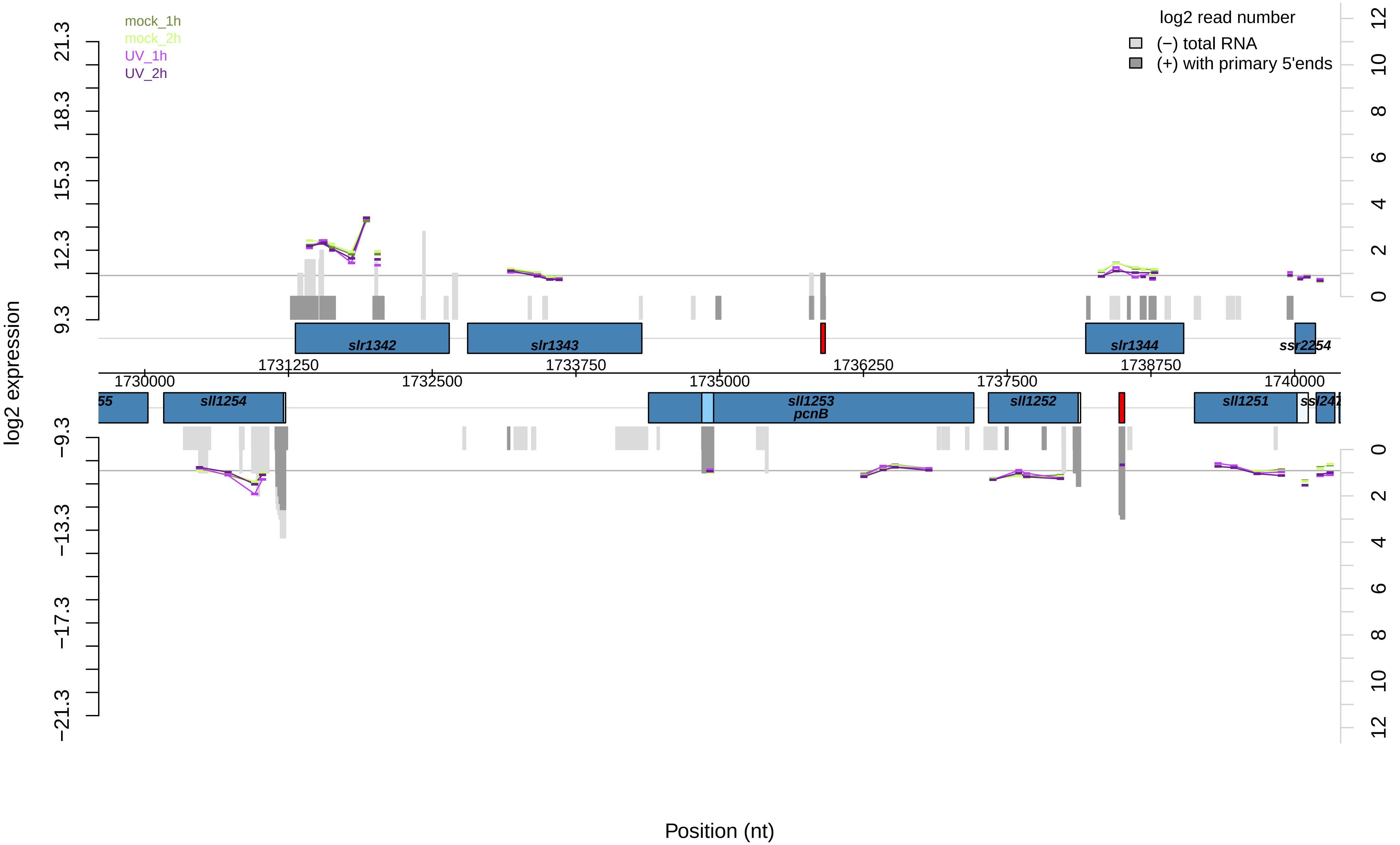

mock_1h
mock_2h
UV_1h
UV_2h
log2 read number
(−) total RNA
(+) with primary 5'ends
log2 expression
21.3
18.3
15.3
12.3
9.3
−9.3
−13.3
−17.3
−21.3
12
10
8
6
4
2
0
slr1342
slr1343
slr1344
ssr2254
1731250
1733750
1736250
1738750
1730000
1732500
1735000
1737500
1740000
sll1254
sll1253
pcnB
sll1252
sll1251
Position (nt)

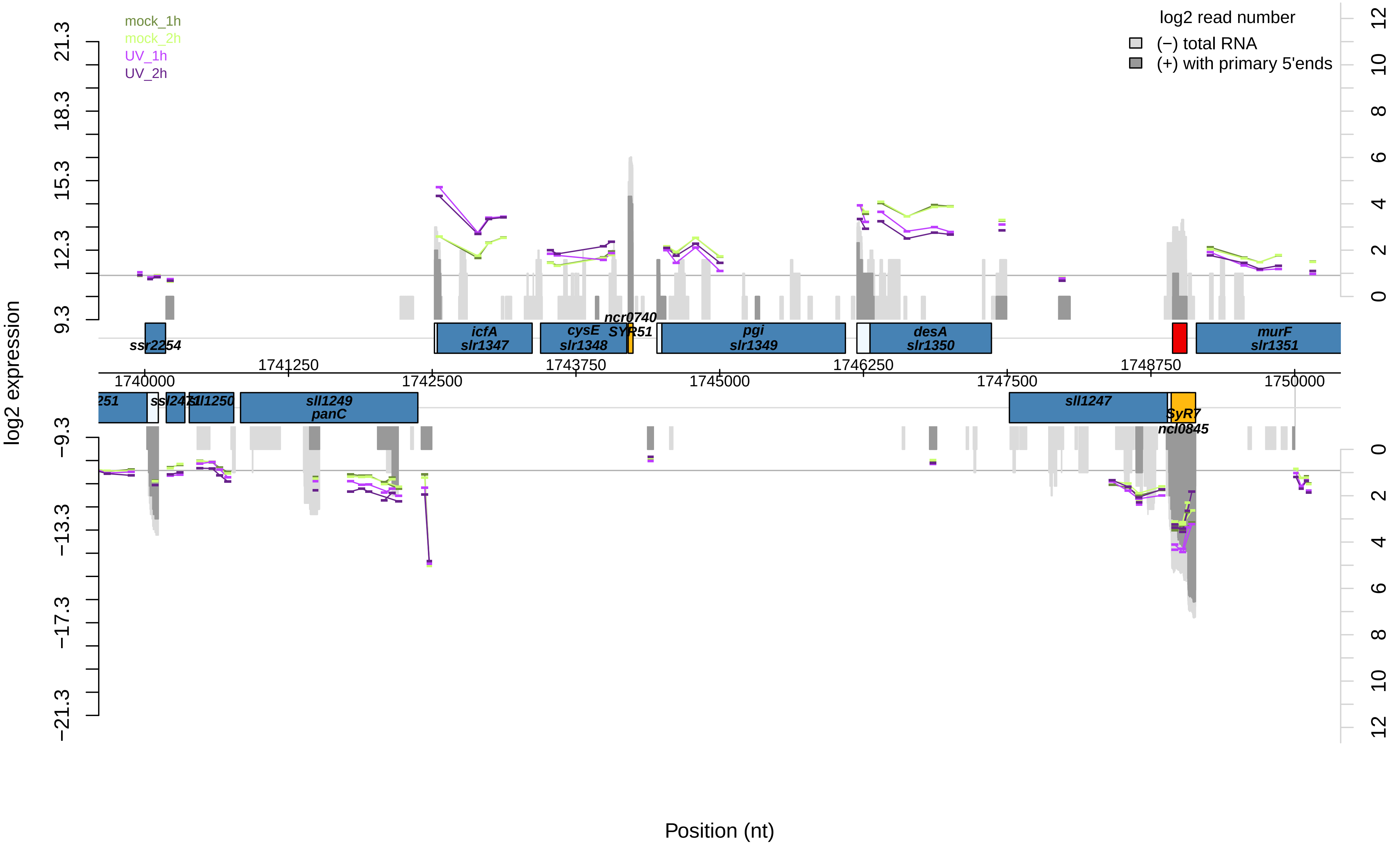

mock_1h
mock_2h
UV_1h
UV_2h
log2 read number
(−) total RNA
(+) with primary 5'ends
log2 expression
21.3
18.3
15.3
12.3
9.3
−9.3
−13.3
−17.3
−21.3
12
10
8
6
4
2
0
ssr2254
icfA
slr1347
cysE
slr1348
ncr0740
SYR51
pgi
slr1349
desA
slr1350
murF
slr1351
1741250
1743750
1746250
1748750
1740000
1742500
1745000
1747500
1750000
251
ssl2471
sll1250
sll1249
panC
sll1247
SyR7
ncl0845
Position (nt)

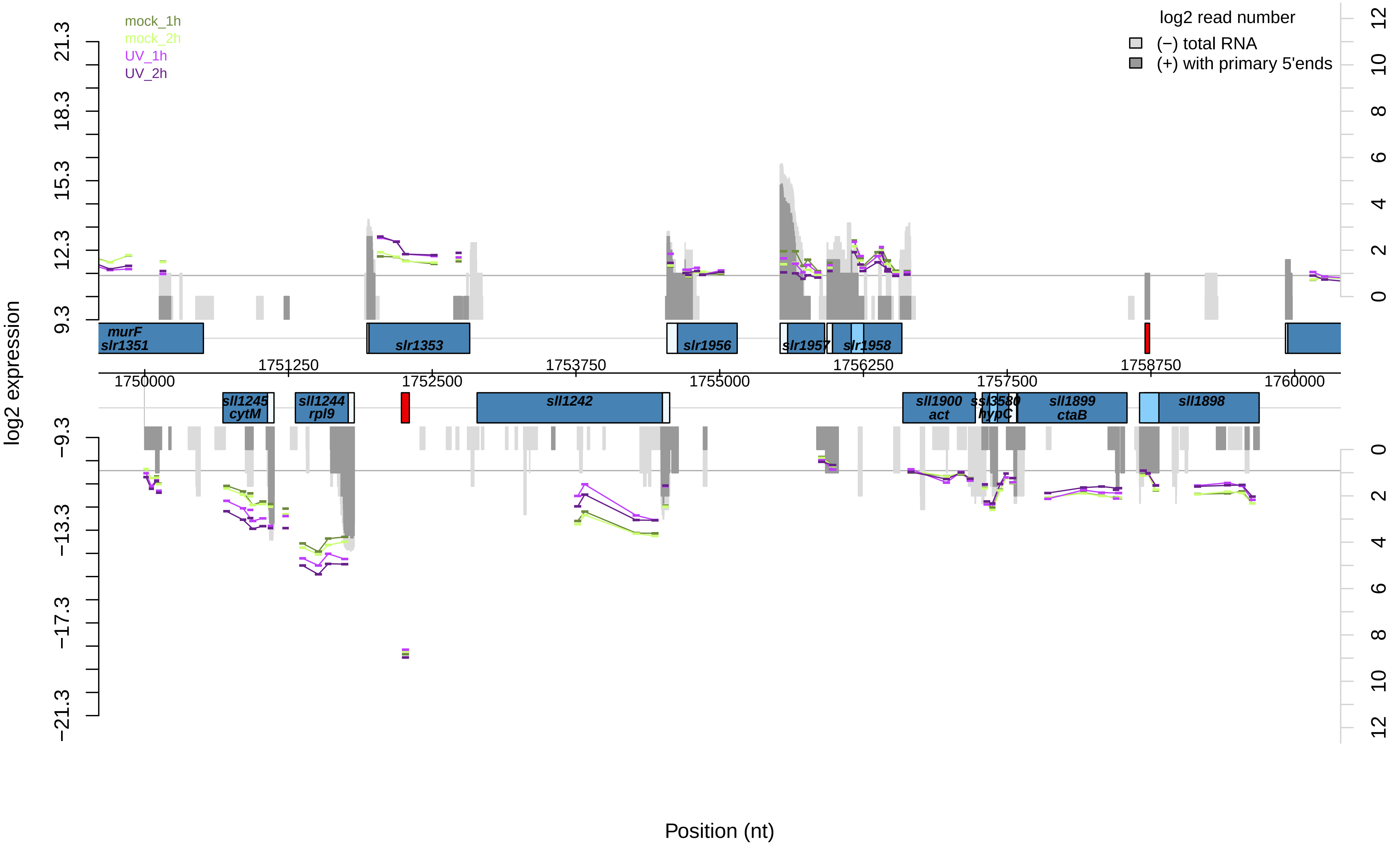
mock_1h
mock_2h
UV_1h
UV_2h
log2 read number
(−) total RNA
(+) with primary 5'ends
log2 expression
21.3
18.3
15.3
12.3
9.3
−9.3
−13.3
−17.3
−21.3
12
10
8
6
4
2
0
murF
slr1351
slr1353
slr1956
slr1957
slr1958
1751250
1753750
1756250
1758750
1750000
1752500
1755000
1757500
1760000
sll1245
cytM
sll1244
rpl9
sll1242
sll1900
act
ssl3580
hypC
sll1899
ctaB
sll1898
Position (nt)

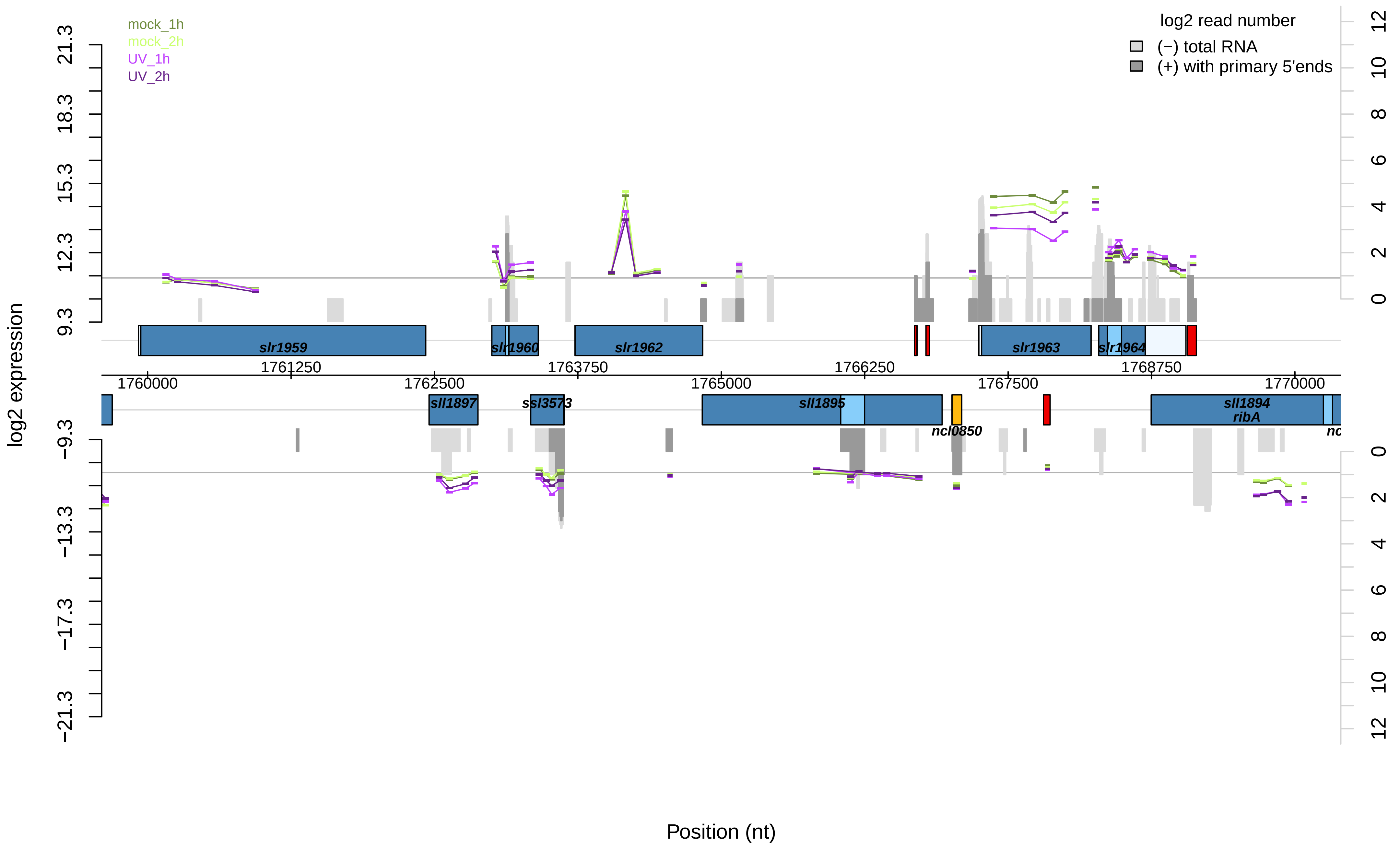
mock_1h
mock_2h
UV_1h
UV_2h
log2 read number
(−) total RNA
(+) with primary 5'ends
log2 expression
21.3
18.3
15.3
12.3
9.3
−9.3
−13.3
−17.3
−21.3
12
10
8
6
4
2
0
slr1959
slr1960
slr1962
slr1963
slr1964
1761250
1763750
1766250
1768750
1760000
1762500
1765000
1767500
1770000
sll1897
ssl3573
sll1895
ncl0850
sll1894
ribA
Position (nt)

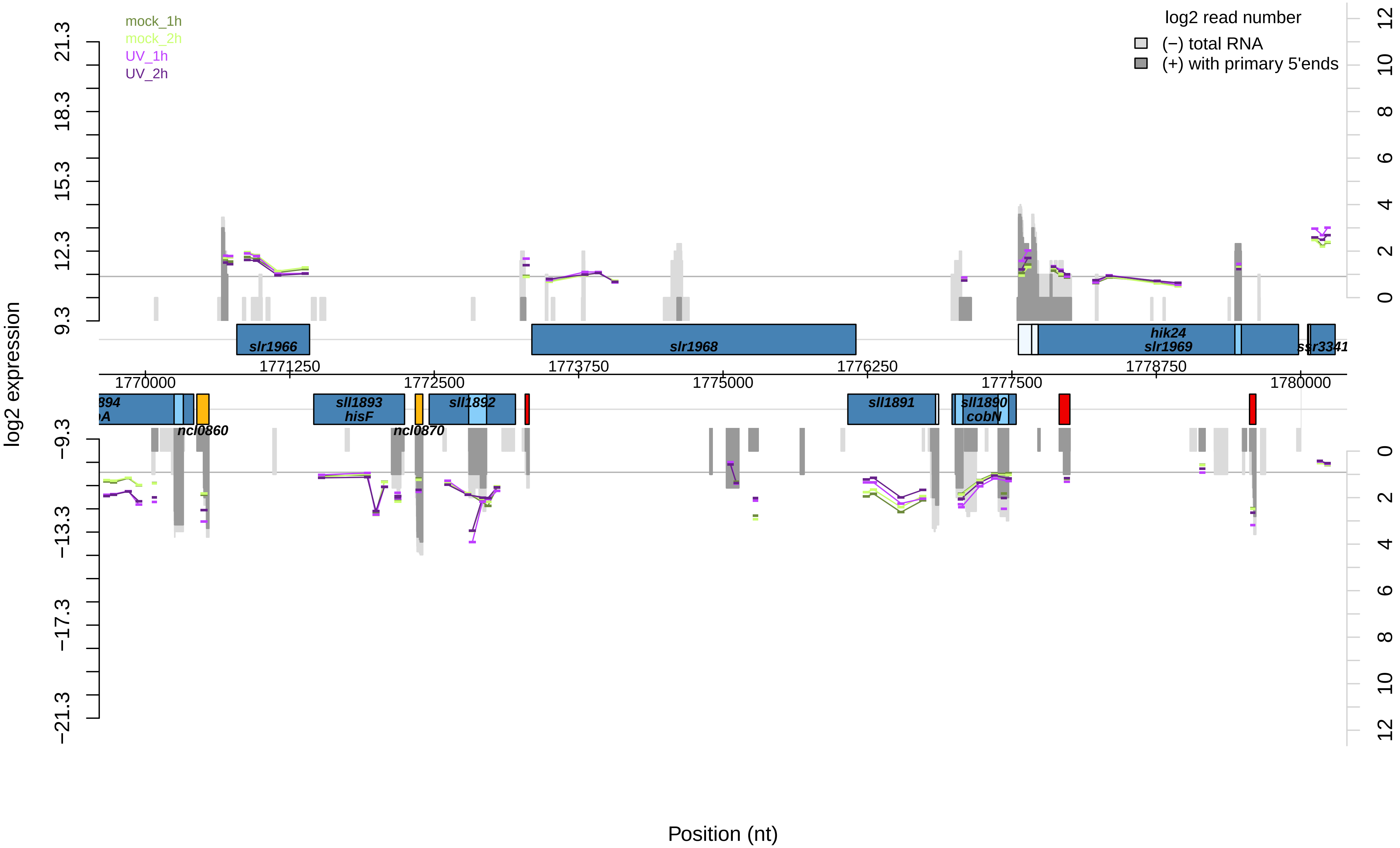

mock_1h
mock_2h
UV_1h
UV_2h
log2 read number
(−) total RNA
(+) with primary 5'ends
log2 expression
21.3
18.3
15.3
12.3
9.3
−9.3
−13.3
−17.3
−21.3
12
10
8
6
4
2
0
slr1966
slr1968
hik24
slr1969
ssr3341
1771250
1773750
1776250
1778750
1770000
1772500
1775000
1777500
1780000
894
sll1893
hisF
sll1892
sll1891
sll1890
cobN
ncl0860
ncl0870
Position (nt)

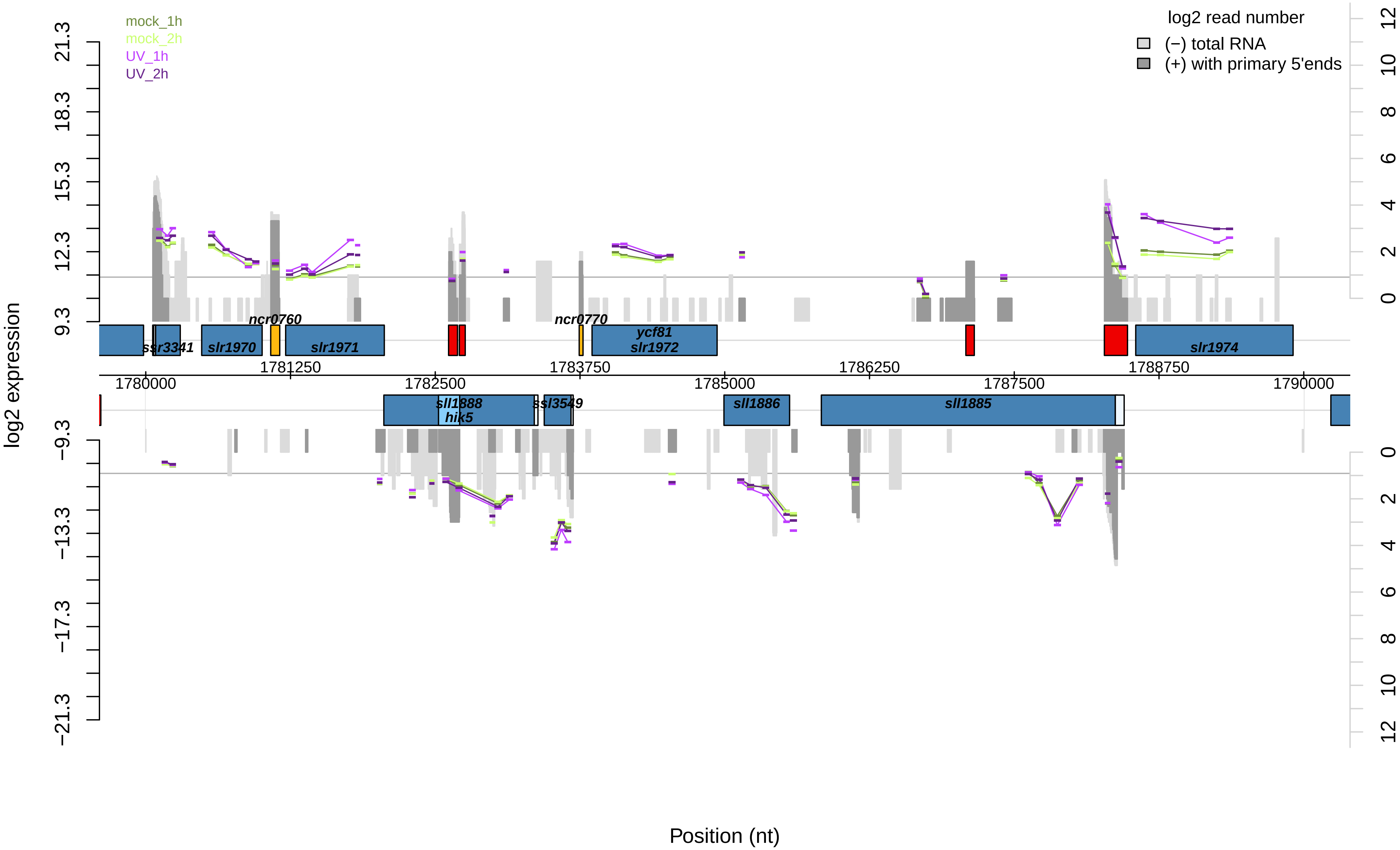

mock_1h
mock_2h
UV_1h
UV_2h
log2 read number
(−) total RNA
(+) with primary 5'ends
log2 expression
21.3
18.3
15.3
12.3
9.3
−9.3
−13.3
−17.3
−21.3
12
10
8
6
4
2
0
ssr3341
slr1970
ncr0760
slr1971
ncr0770
ycf81
slr1972
slr1974
1781250
1782500
1783750
1785000
1786250
1787500
1788750
1790000
1780000
sll1888
hik5
ssl3549
sll1886
sll1885
Position (nt)

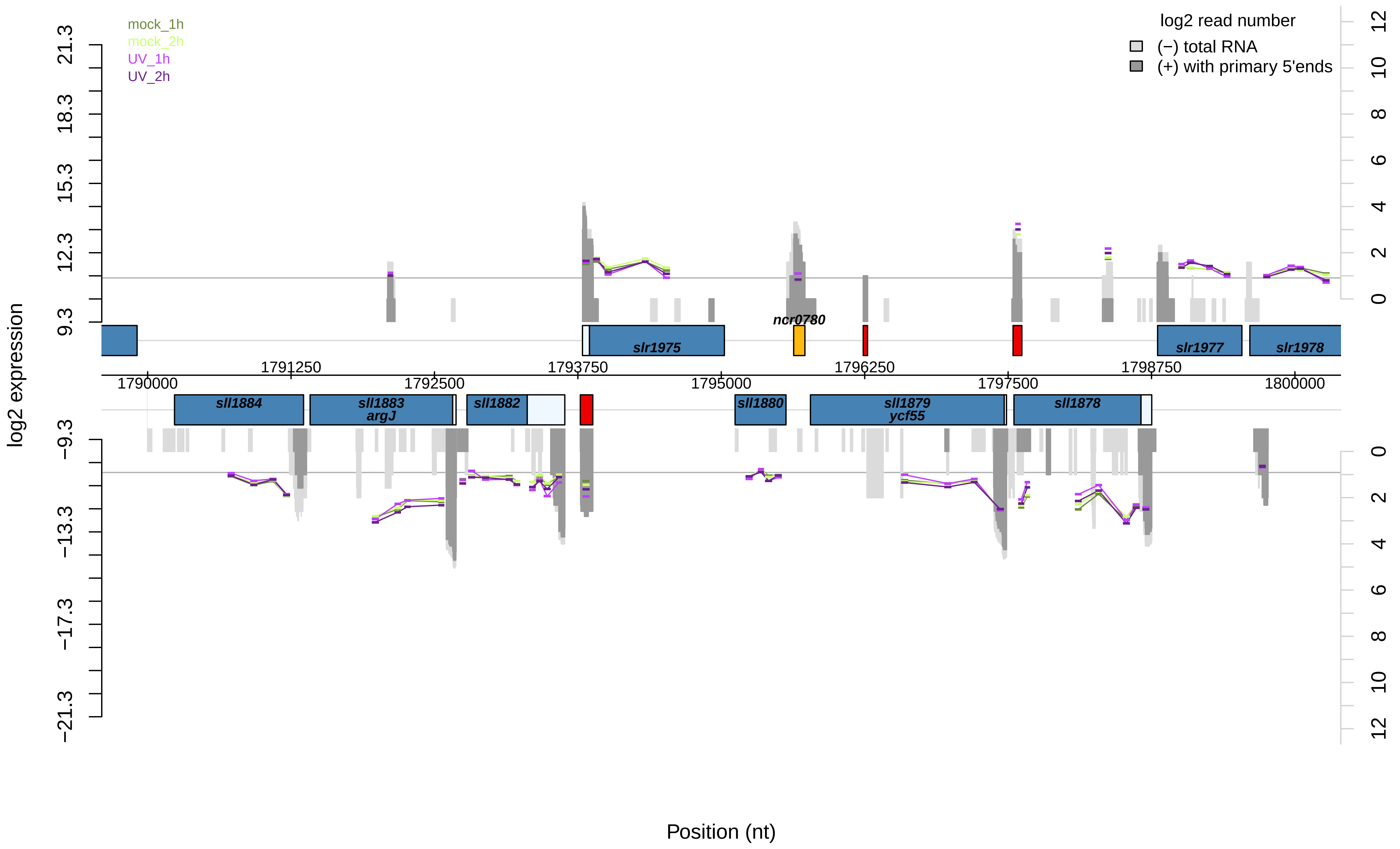

mock_1h
mock_2h
UV_1h
UV_2h
log2 read number
(−) total RNA
(+) with primary 5'ends
log2 expression
21.3
18.3
15.3
12.3
9.3
−9.3
−13.3
−17.3
−21.3
12
10
8
6
4
2
0
ncr0780
slr1975
slr1977
slr1978
1791250
1793750
1796250
1798750
1790000
1792500
1795000
1797500
1800000
sll1884
sll1883
argJ
sll1882
sll1880
sll1879
ycf55
sll1878
Position (nt)

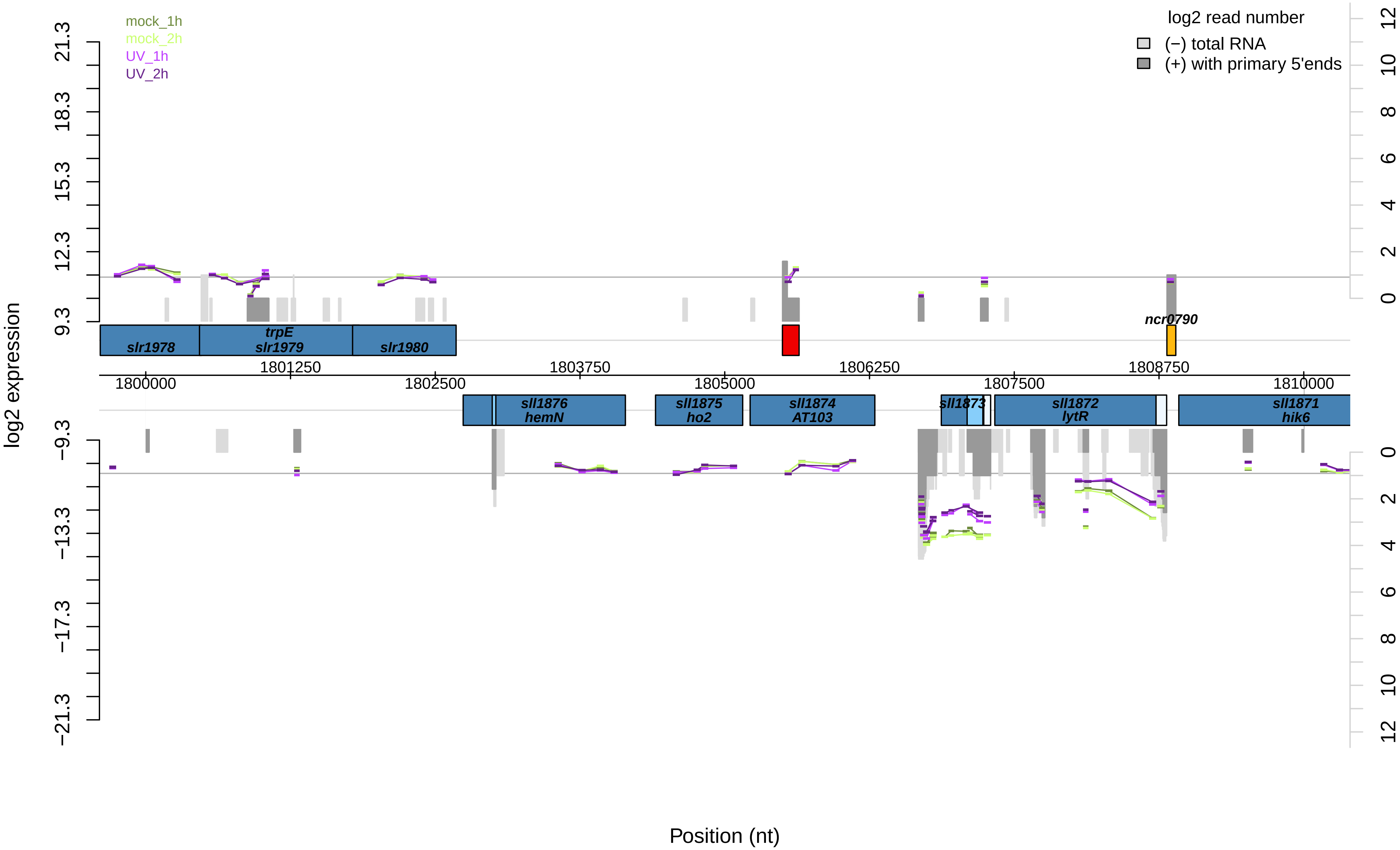
mock_1h
mock_2h
UV_1h
UV_2h
log2 read number
(−) total RNA
(+) with primary 5'ends
log2 expression
21.3
18.3
15.3
12.3
9.3
−9.3
−13.3
−17.3
−21.3
slr1978
trpE
slr1979
slr1980
ncr0790
1801250
1803750
1806250
1808750
1800000
1802500
1805000
1807500
1810000
sll1876
hemN
sll1875
ho2
sll1874
AT103
sll1873
sll1872
lytR
sll1871
hik6
Position (nt)

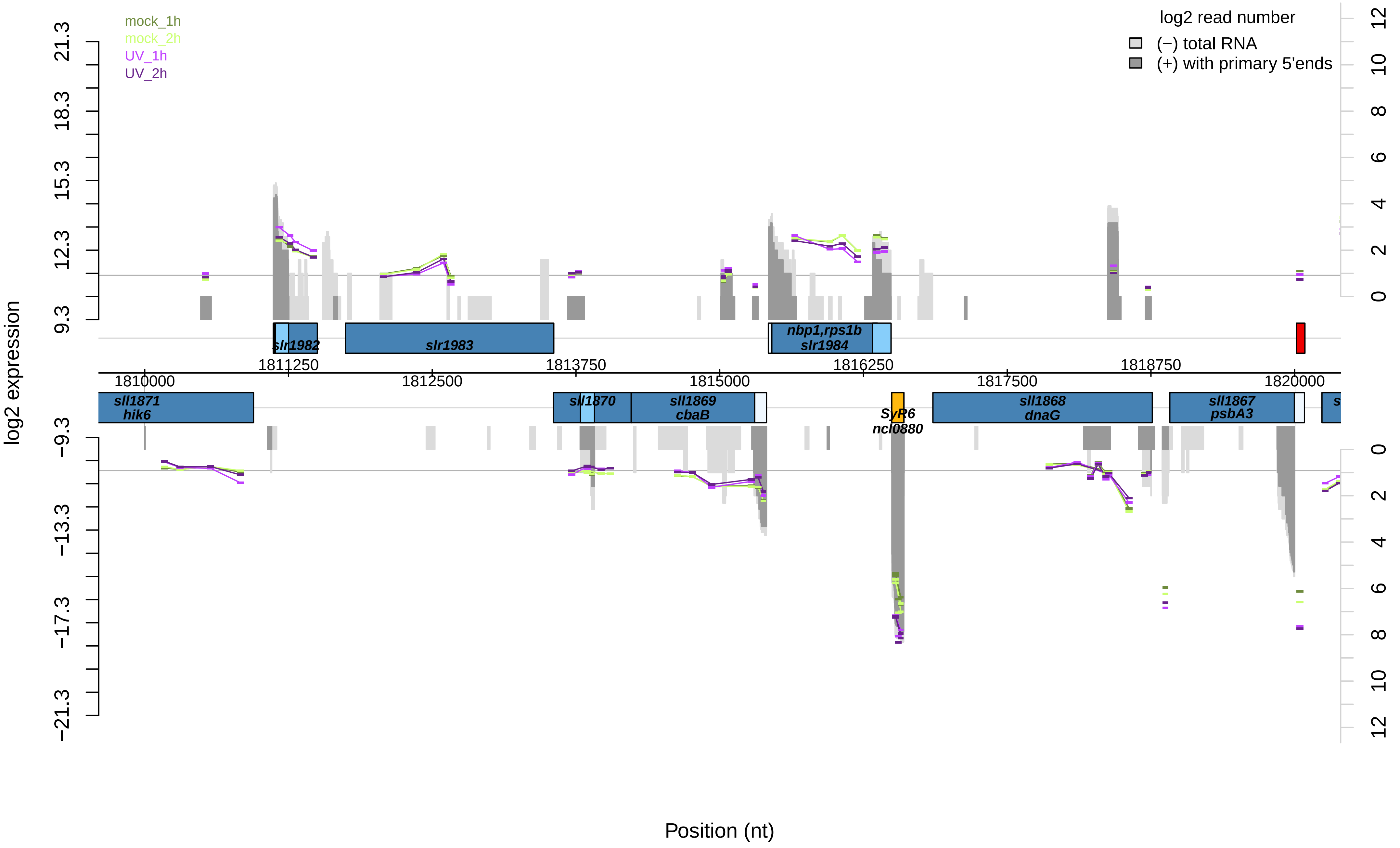

mock_1h
mock_2h
UV_1h
UV_2h
log2 read number
(−) total RNA
(+) with primary 5'ends
log2 expression
21.3
18.3
15.3
12.3
9.3
−9.3
−13.3
−17.3
−21.3
12
10
8
6
4
2
0
slr1982
slr1983
nbp1,rps1b
slr1984
1811250
1813750
1816250
1818750
1810000
1812500
1815000
1817500
1820000
sll1871
hik6
sll1870
sll1869
cbaB
SyR6
ncl0880
sll1868
dnaG
sll1867
psbA3
Position (nt)

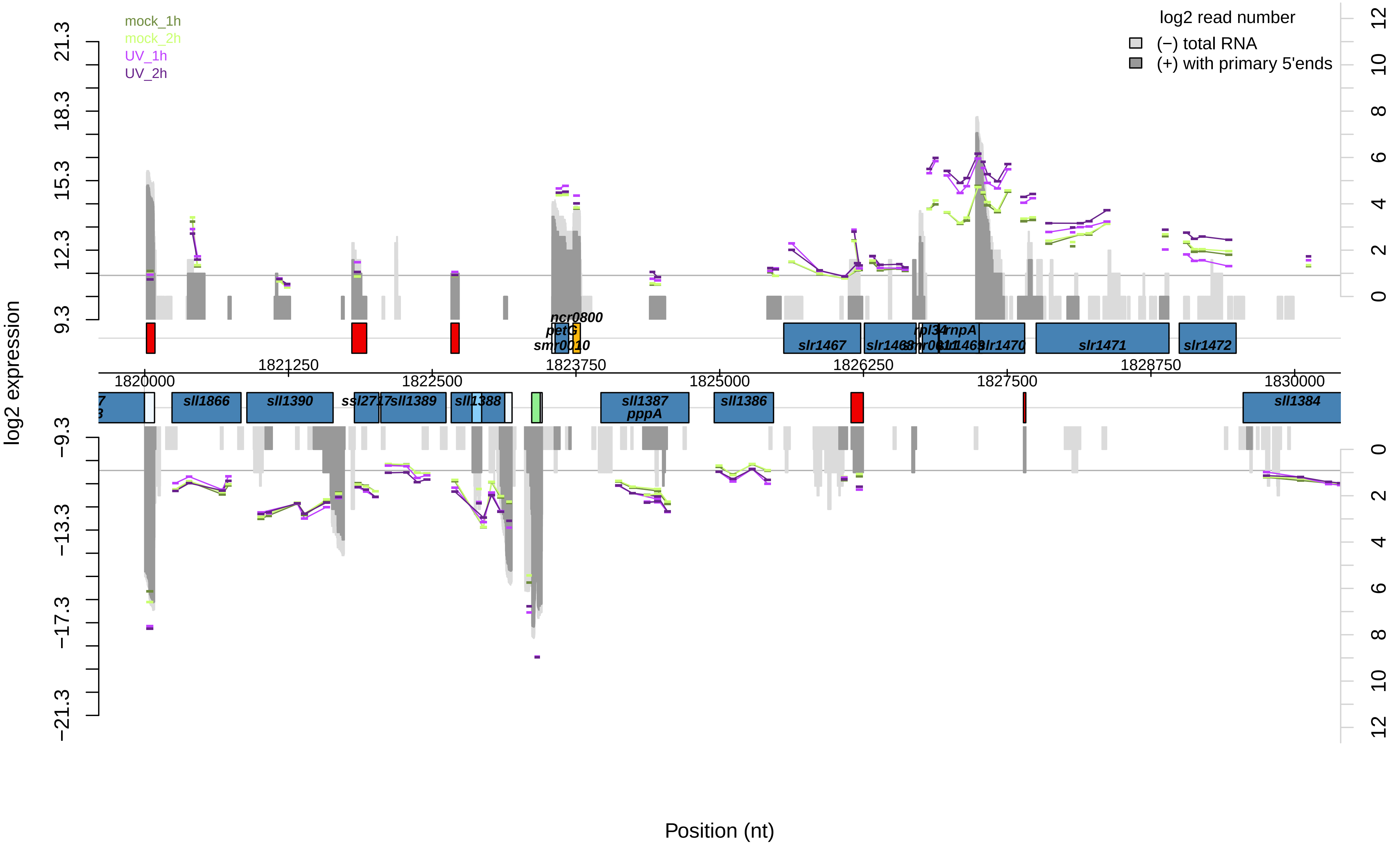
mock_1h
mock_2h
UV_1h
UV_2h
log2 read number
(−) total RNA
(+) with primary 5'ends
log2 expression
Position (nt)
21.3
18.3
15.3
12.3
9.3
−9.3
−13.3
−17.3
−21.3
1820000
1821250
1822500
1823750
1825000
1826250
1827500
1828750
1830000
ncr0800
petG
smr0010
slr1467
slr1468
rpl34
rnpA
smr0011
slr1469
slr1470
slr1471
slr1472
sll1866
sll1390
ssl2717
sll1389
sll1388
sll1387
pppA
sll1386
sll1384

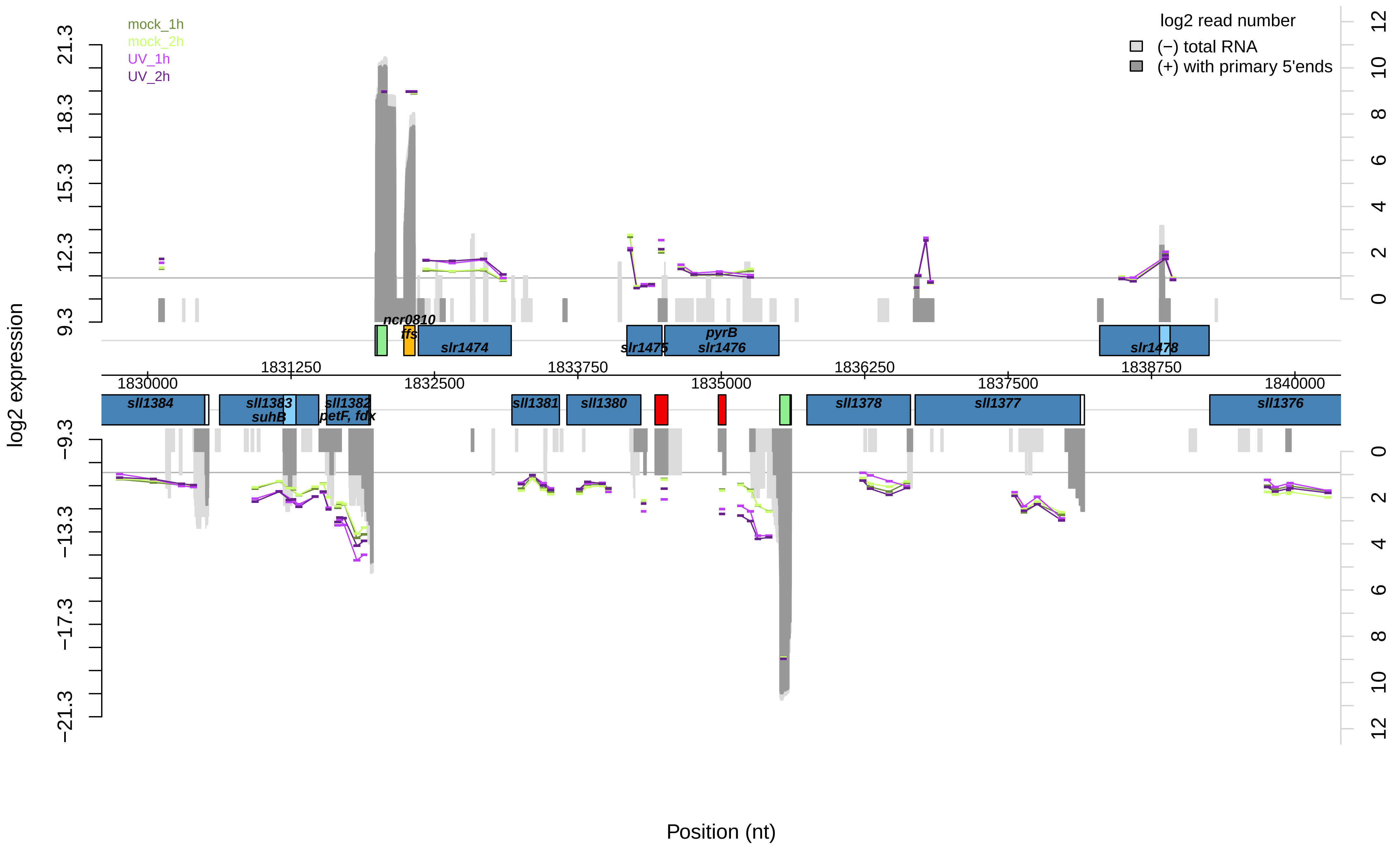

mock_1h
mock_2h
UV_1h
UV_2h
log2 read number
(−) total RNA
(+) with primary 5'ends
log2 expression
21.3
18.3
15.3
12.3
9.3
−9.3
−13.3
−17.3
−21.3
12
10
8
6
4
2
0
ncr0810
ffs
slr1474
slr1475
pyrB
slr1476
slr1478
1831250
1832500
1833750
1835000
1836250
1837500
1838750
1840000
1830000
sll1384
sll1383
suhB
sll1382
petF, fdx
sll1381
sll1380
sll1378
sll1377
sll1376
Position (nt)

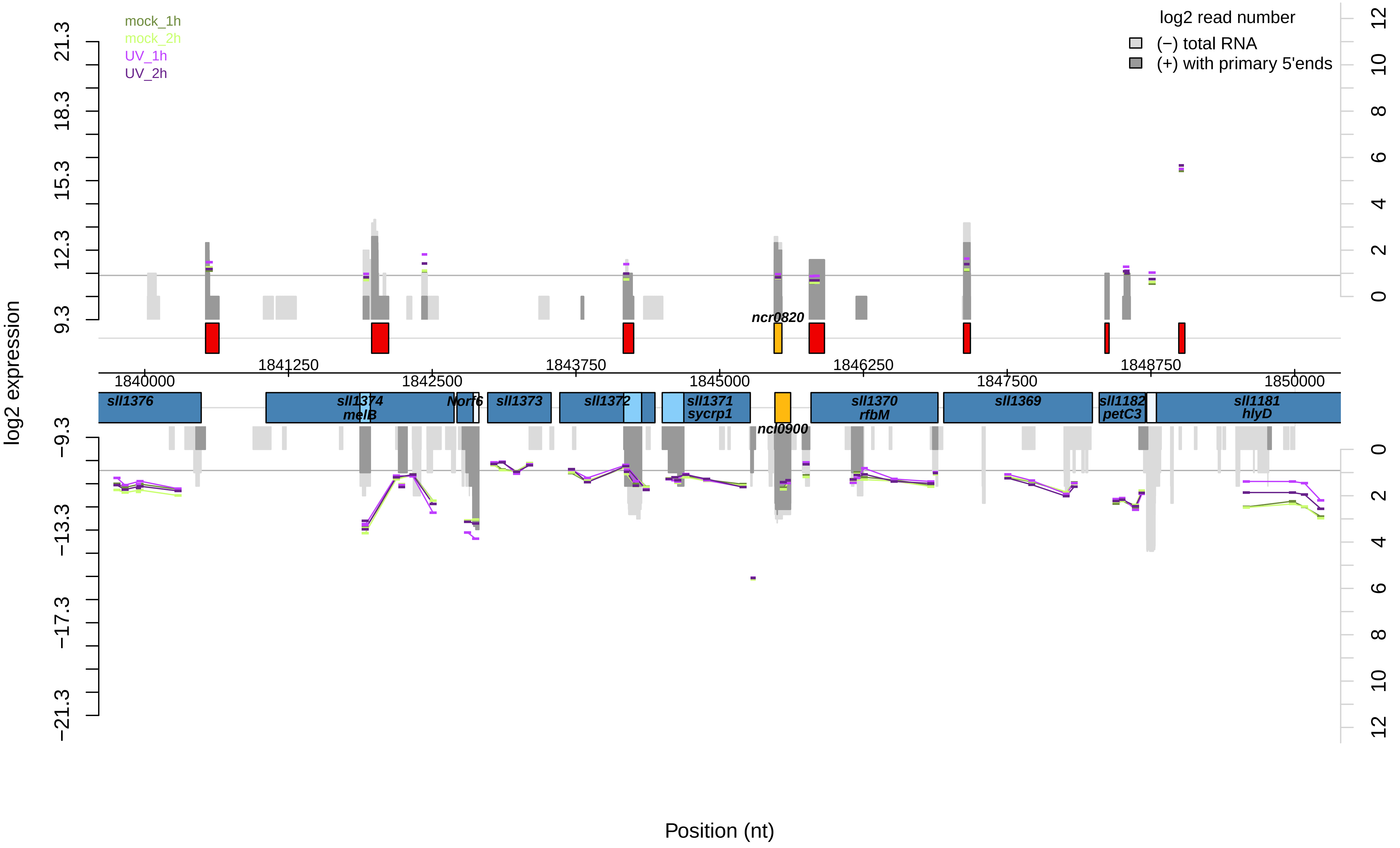

mock_1h
mock_2h
UV_1h
UV_2h
log2 read number
(−) total RNA
(+) with primary 5'ends
log2 expression
21.3
18.3
15.3
12.3
9.3
−9.3
−13.3
−17.3
−21.3
12
10
8
6
4
2
0
ncr0820
ncl0900
1840000
1841250
1842500
1843750
1845000
1846250
1847500
1848750
1850000
sll1376
sll1374
melB
Norf6
sll1373
sll1372
sll1371
sycrp1
sll1370
rfbM
sll1369
sll1182
petC3
sll1181
hlyD
Position (nt)

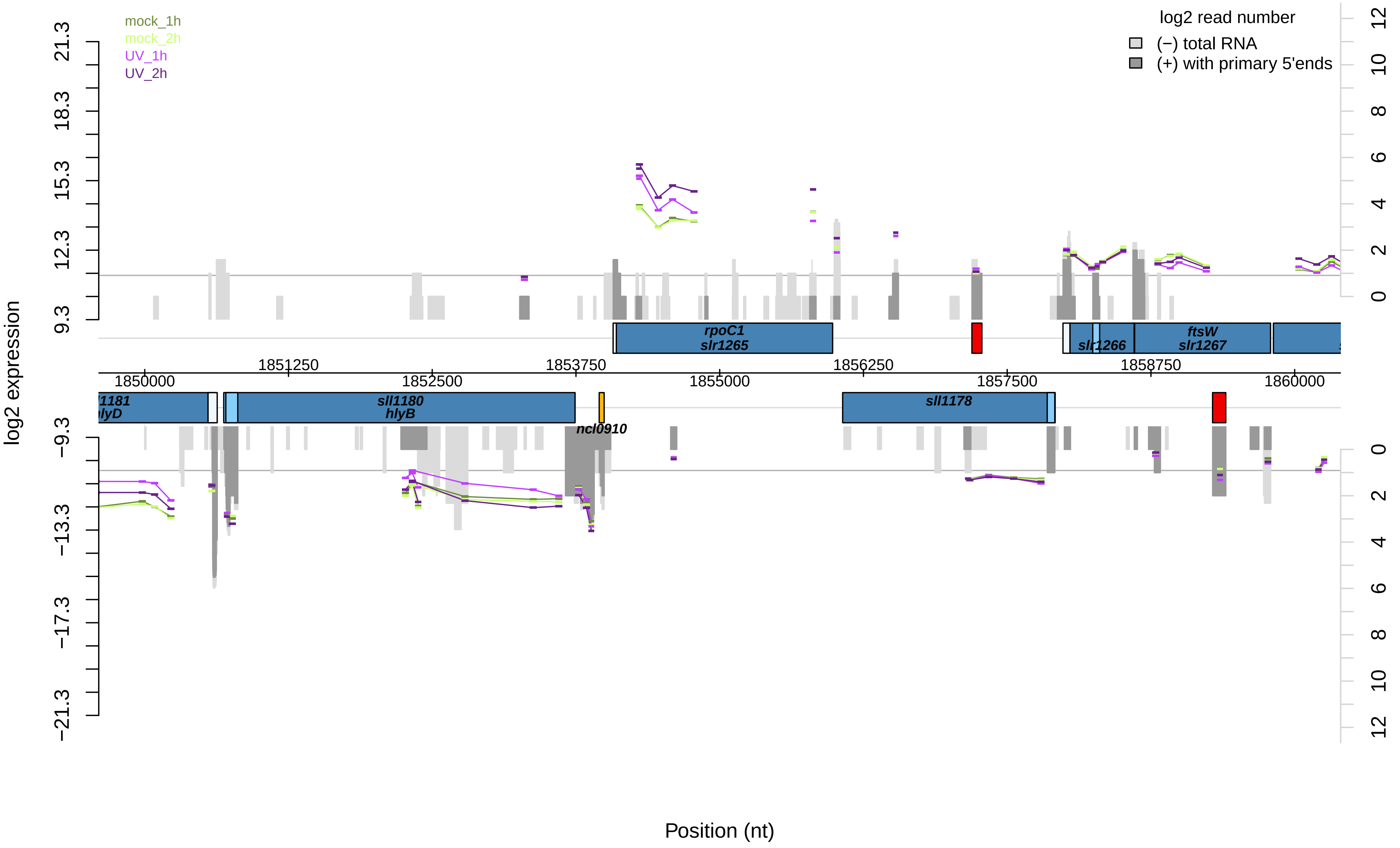

mock_1h
mock_2h
UV_1h
UV_2h
log2 read number
(−) total RNA
(+) with primary 5'ends
log2 expression
Position (nt)
rpoC1
slr1265
slr1266
ftsW
slr1267
sll1180
hlyB
ncl0910
sll1178
1851250
1853750
1856250
1858750
1850000
1852500
1855000
1857500
1860000
21.3
18.3
15.3
12.3
9.3
−9.3
−13.3
−17.3
−21.3

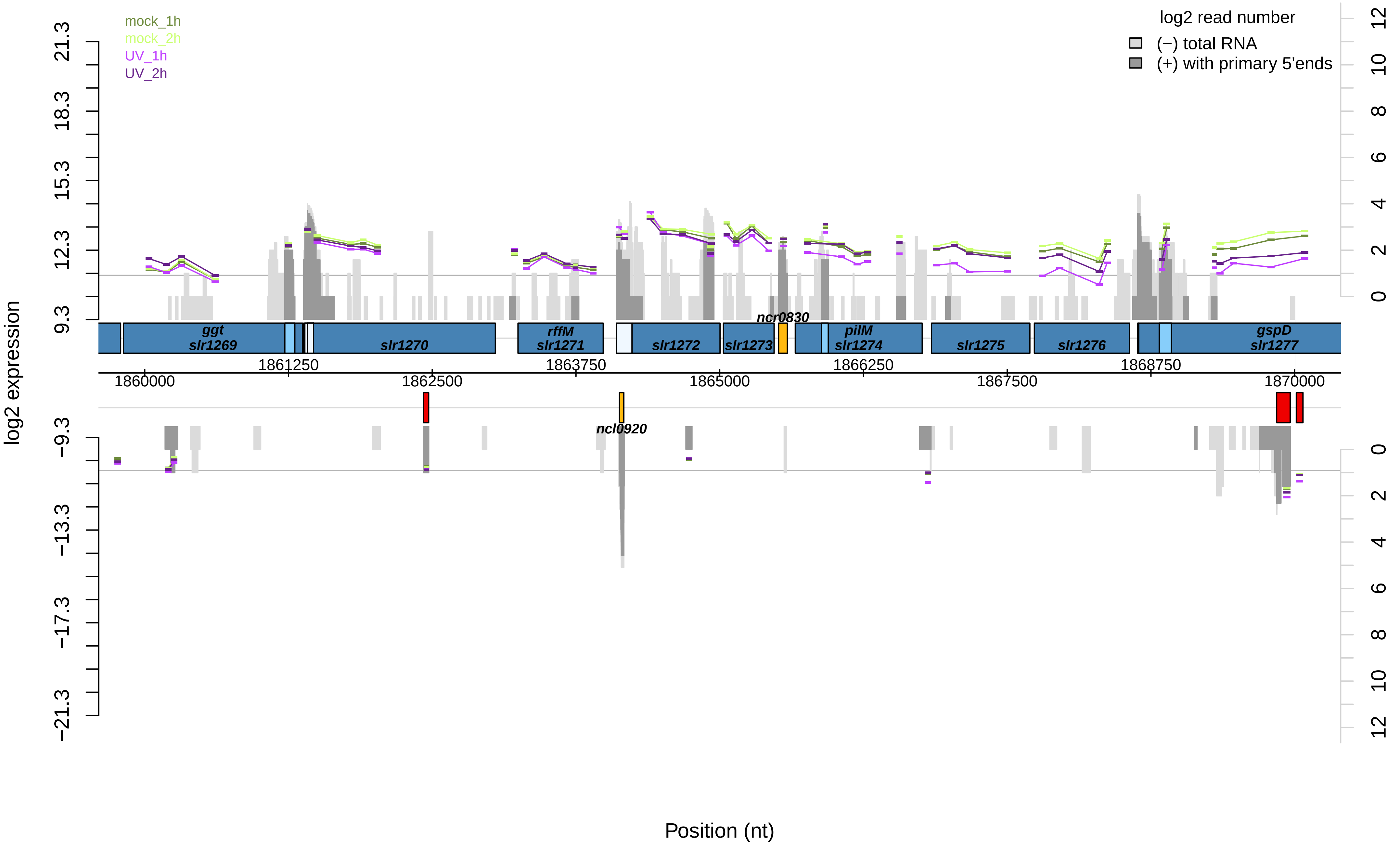

mock_1h
mock_2h
UV_1h
UV_2h
log2 read number
(−) total RNA
(+) with primary 5'ends
log2 expression
21.3
18.3
15.3
12.3
9.3
−9.3
−13.3
−17.3
−21.3
12
10
8
6
4
2
0
ggt
slr1269
slr1270
rffM
slr1271
slr1272
slr1273
ncr0830
pilM
slr1274
slr1275
slr1276
gspD
slr1277
1861250
1863750
1866250
1868750
1860000
1862500
1865000
1867500
1870000
ncl0920
Position (nt)

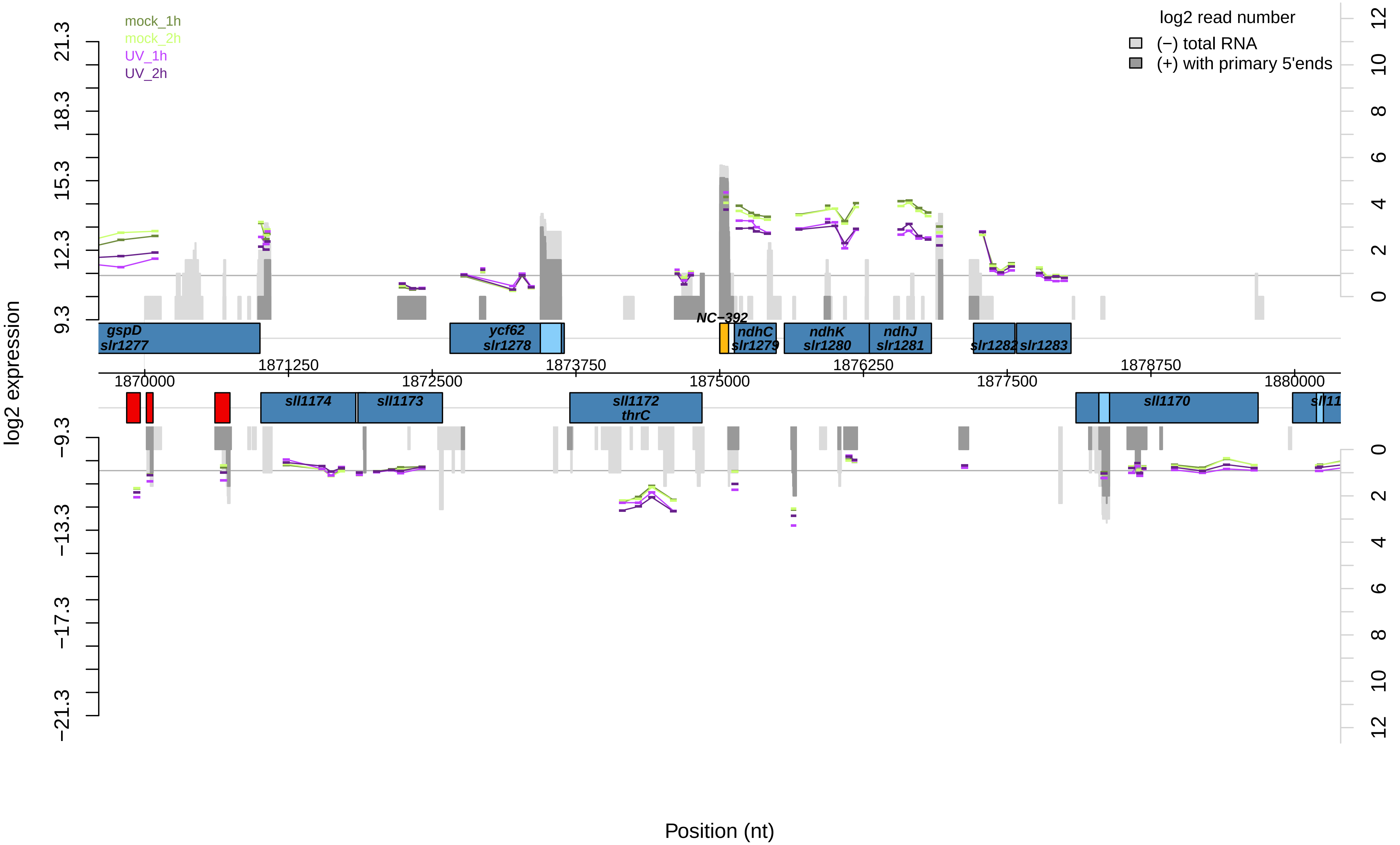

mock_1h
mock_2h
UV_1h
UV_2h
log2 read number
(−) total RNA
(+) with primary 5'ends
log2 expression
21.3
18.3
15.3
12.3
9.3
−9.3
−13.3
−17.3
−21.3
12
10
8
6
4
2
0
gspD
slr1277
ycf62
slr1278
NC−392
ndhC
slr1279
ndhK
slr1280
ndhJ
slr1281
slr1282
slr1283
1871250
1872500
1873750
1875000
1876250
1877500
1878750
1870000
1880000
sll1174
sll1173
sll1172
thrC
sll1170
Position (nt)

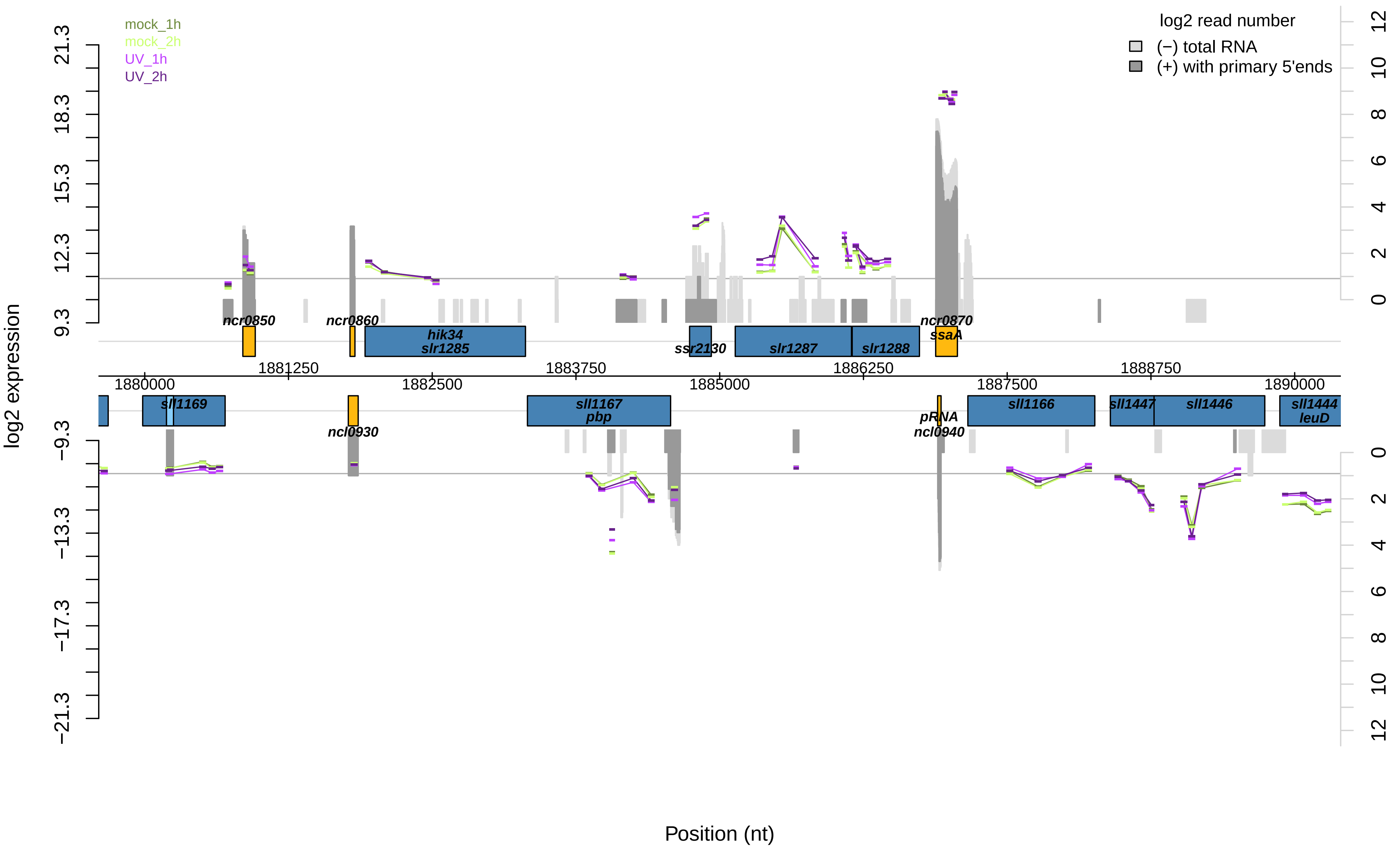

mock_1h
mock_2h
UV_1h
UV_2h
log2 read number
(−) total RNA
(+) with primary 5'ends
log2 expression
21.3
18.3
15.3
12.3
9.3
−9.3
−13.3
−17.3
−21.3
12
10
8
6
4
2
0
ncr0850
ncr0860
hik34
slr1285
ssr2130
slr1287
slr1288
ncr0870
ssaA
1881250
1883750
1886250
1888750
1880000
1882500
1885000
1887500
1890000
sll1169
ncl0930
sll1167
pbp
pRNA
ncl0940
sll1166
sll1447
sll1446
sll1444
leuD
Position (nt)

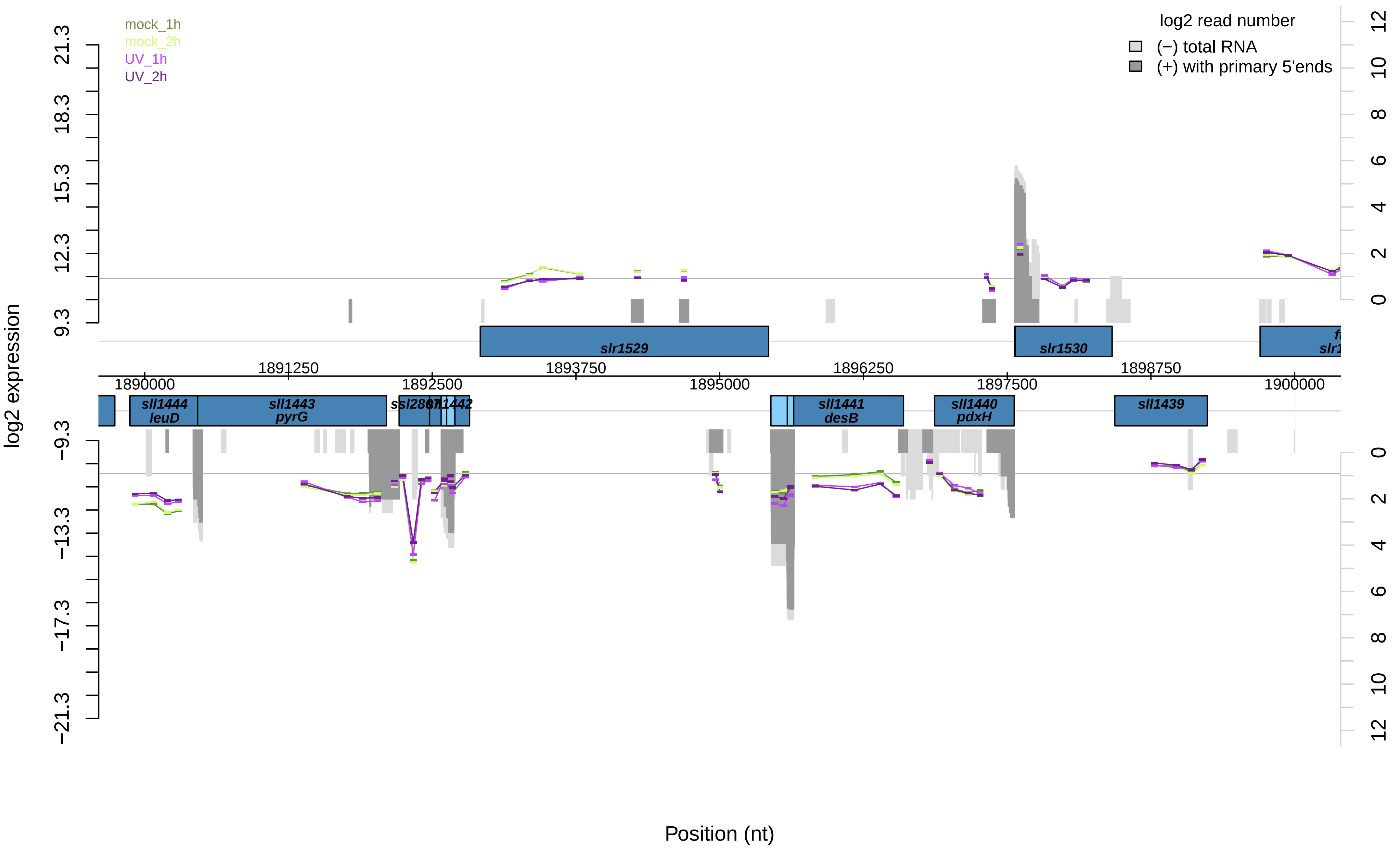

mock_1h
mock_2h
UV_1h
UV_2h
log2 read number
(−) total RNA
(+) with primary 5'ends
log2 expression
21.3
18.3
15.3
12.3
9.3
−9.3
−13.3
−17.3
−21.3
12
10
8
6
4
2
0
slr1529
slr1530
1891250
1893750
1896250
1898750
1890000
1892500
1895000
1897500
1900000
sll1444
leuD
sll1443
pyrG
sll1441
desB
sll1440
pdxH
sll1439
Position (nt)

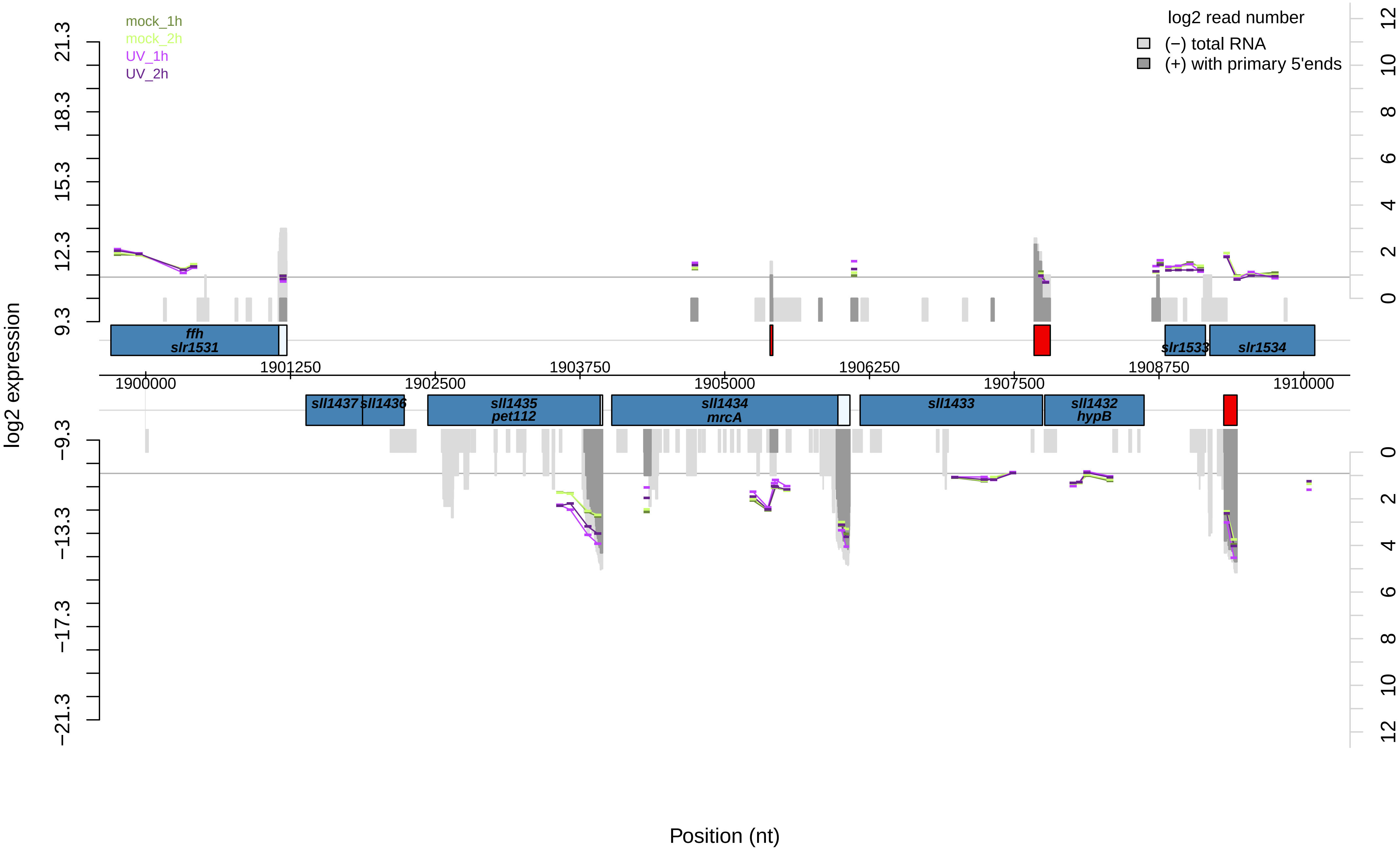

mock_1h
mock_2h
UV_1h
UV_2h
log2 read number
(−) total RNA
(+) with primary 5'ends
log2 expression
21.3
18.3
15.3
12.3
9.3
−9.3
−13.3
−17.3
−21.3
12
10
8
6
4
2
0
ffh
slr1531
slr1533
slr1534
sll1437
sll1436
sll1435
pet112
sll1434
mrcA
sll1433
sll1432
hypB
1900000
1901250
1902500
1903750
1905000
1906250
1907500
1908750
1910000
Position (nt)

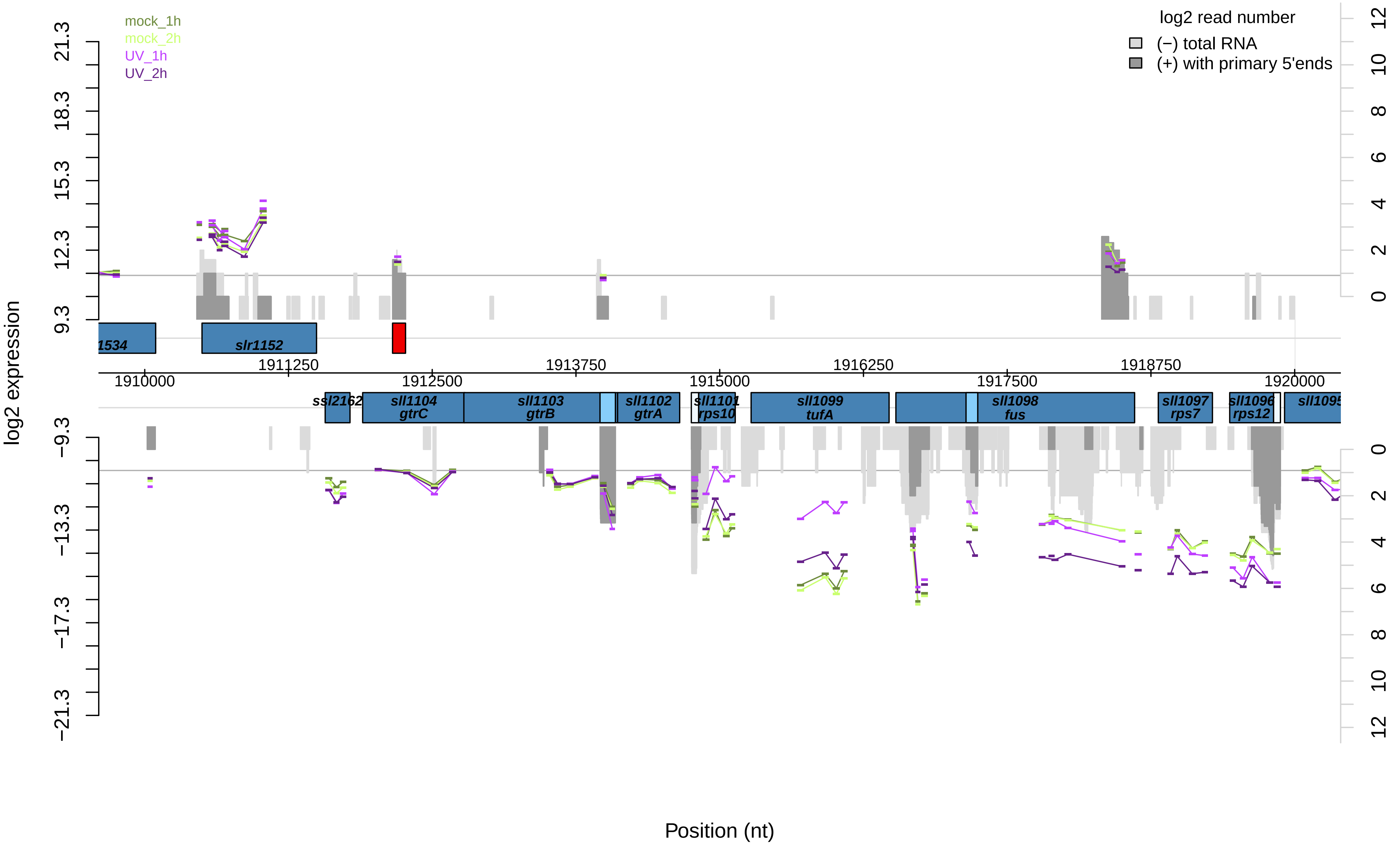

mock_1h
mock_2h
UV_1h
UV_2h
log2 read number
(−) total RNA
(+) with primary 5'ends
log2 expression
21.3
18.3
15.3
12.3
9.3
−9.3
−13.3
−17.3
−21.3
12
10
8
6
4
2
0
1534
slr1152
1911250
1912500
1913750
1915000
1916250
1917500
1918750
1920000
1910000
ssl2162
sll1104
gtrC
sll1103
gtrB
sll1102
gtrA
sll1101
rps10
sll1099
tufA
sll1098
fus
sll1097
rps7
sll1096
rps12
Position (nt)

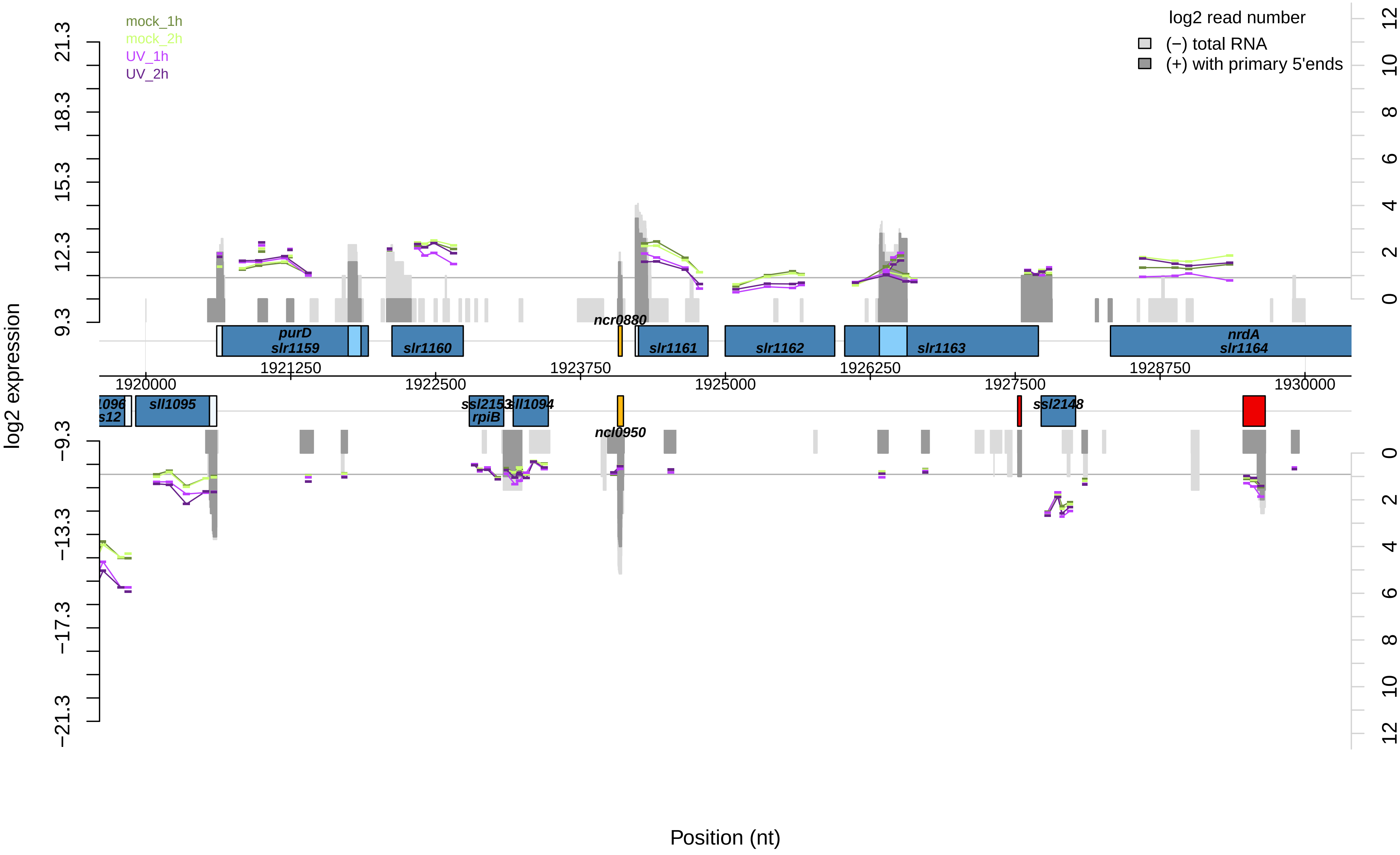

mock_1h
mock_2h
UV_1h
UV_2h
log2 read number
(−) total RNA
(+) with primary 5'ends
log2 expression
21.3
18.3
15.3
12.3
9.3
−9.3
−13.3
−17.3
−21.3
12
10
8
6
4
2
0
purD
slr1159
slr1160
ncr0880
slr1161
slr1162
slr1163
nrdA
slr1164
1921250
1923750
1926250
1928750
1920000
1922500
1925000
1927500
1930000
sll1095
ssl2153
rpiB
sll1094
ncl0950
ssl2148
Position (nt)

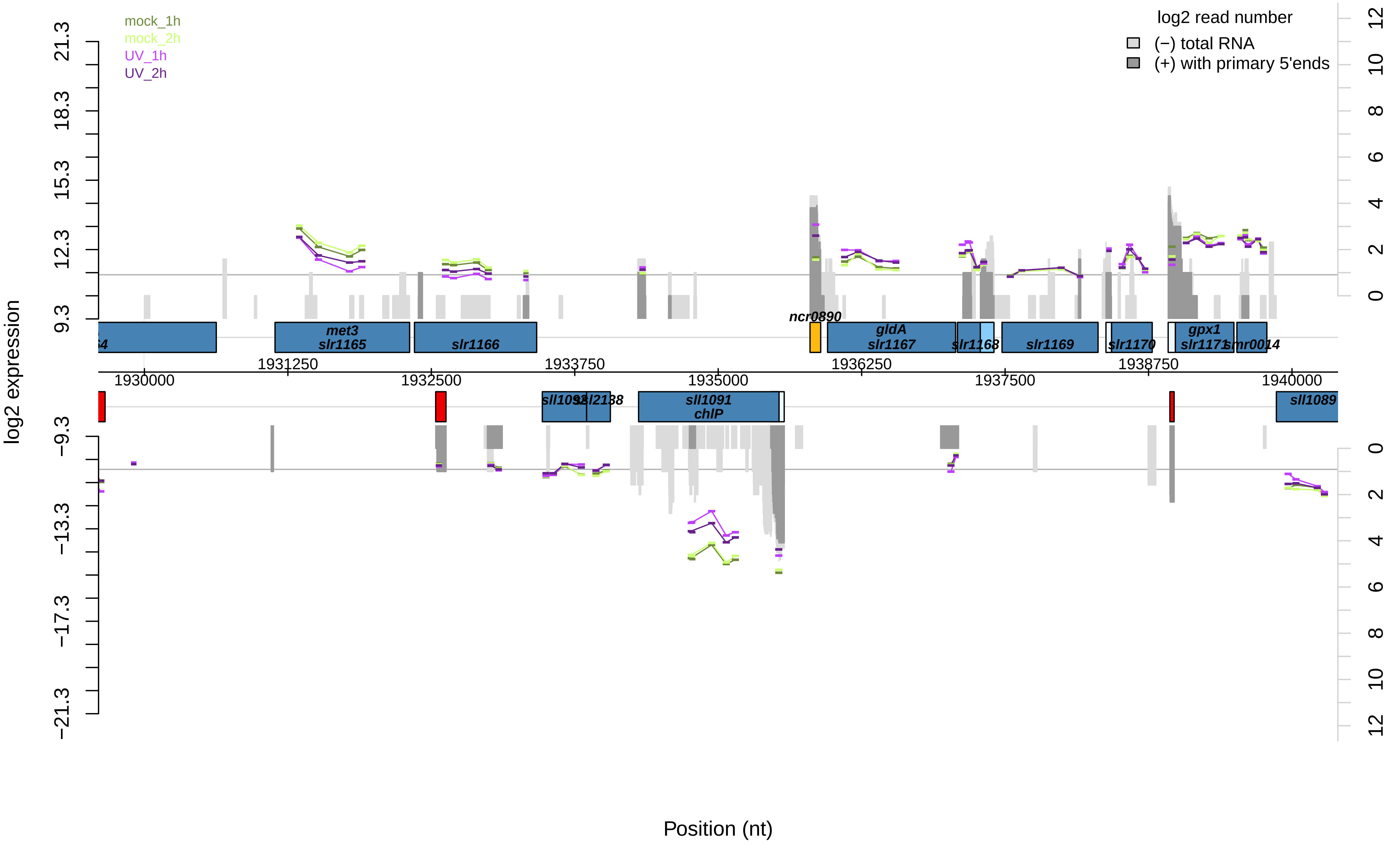

mock_1h
mock_2h
UV_1h
UV_2h
log2 read number
(−) total RNA
(+) with primary 5'ends
log2 expression
Position (nt)
met3
slr1165
slr1166
ncr0890
gldA
slr1167
slr1168
slr1169
slr1170
gpx1
slr1171
smr0014
sll1092
sll2138
sll1091
chlP
sll1089
1930000
1931250
1932500
1933750
1935000
1936250
1937500
1938750
1940000
21.3
18.3
15.3
12.3
9.3
−9.3
−13.3
−17.3
−21.3

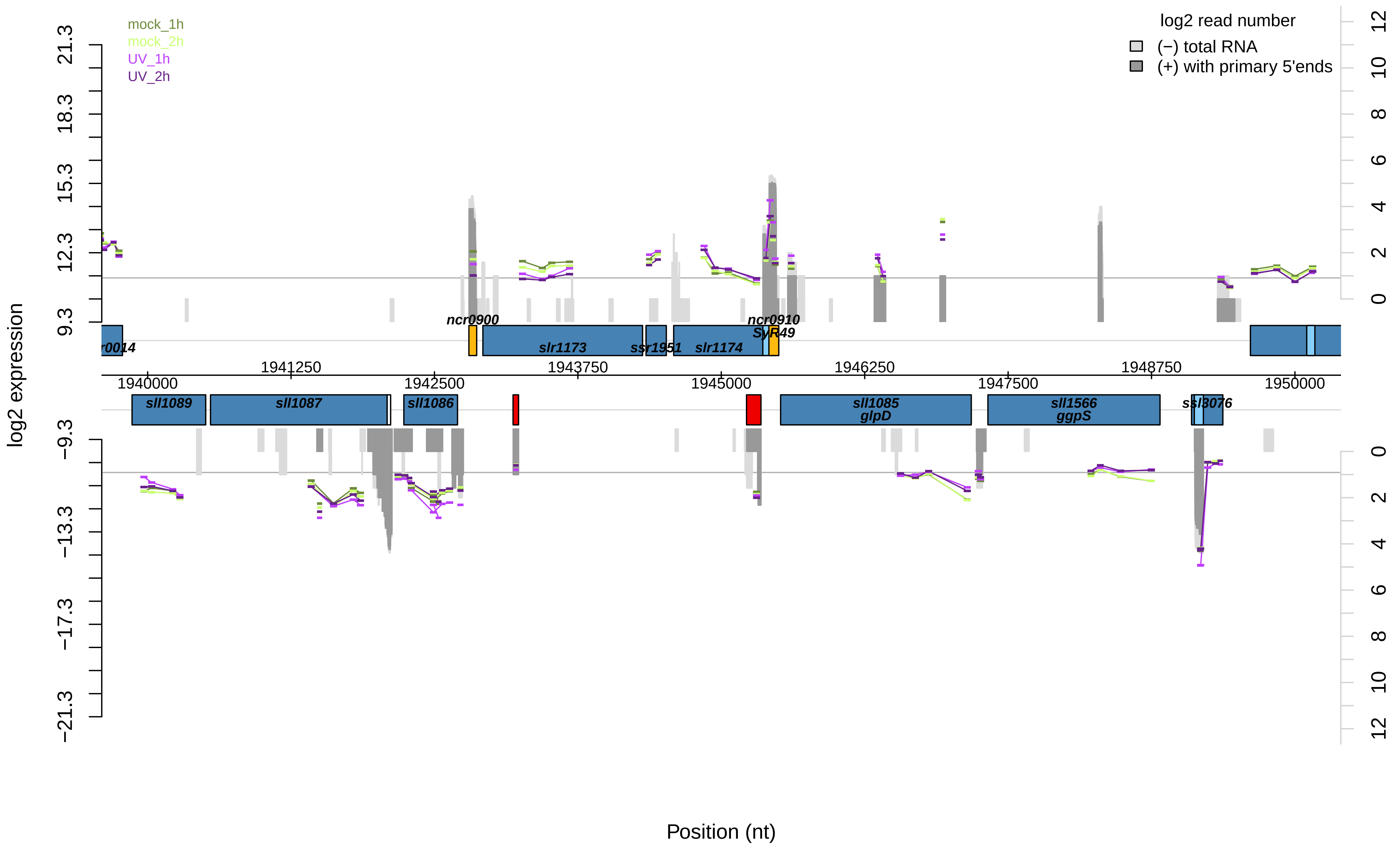

mock_1h
mock_2h
UV_1h
UV_2h
log2 read number
(−) total RNA
(+) with primary 5'ends
log2 expression
21.3
18.3
15.3
12.3
9.3
−9.3
−13.3
−17.3
−21.3
12
10
8
6
4
2
0
ncr0900
ncr0910
SyR49
r0014
slr1173
ssr1951
slr1174
1941250
1943750
1946250
1948750
1940000
1942500
1945000
1947500
1950000
sll1089
sll1087
sll1086
sll1085
glpD
sll1566
ggpS
ssl3076
Position (nt)

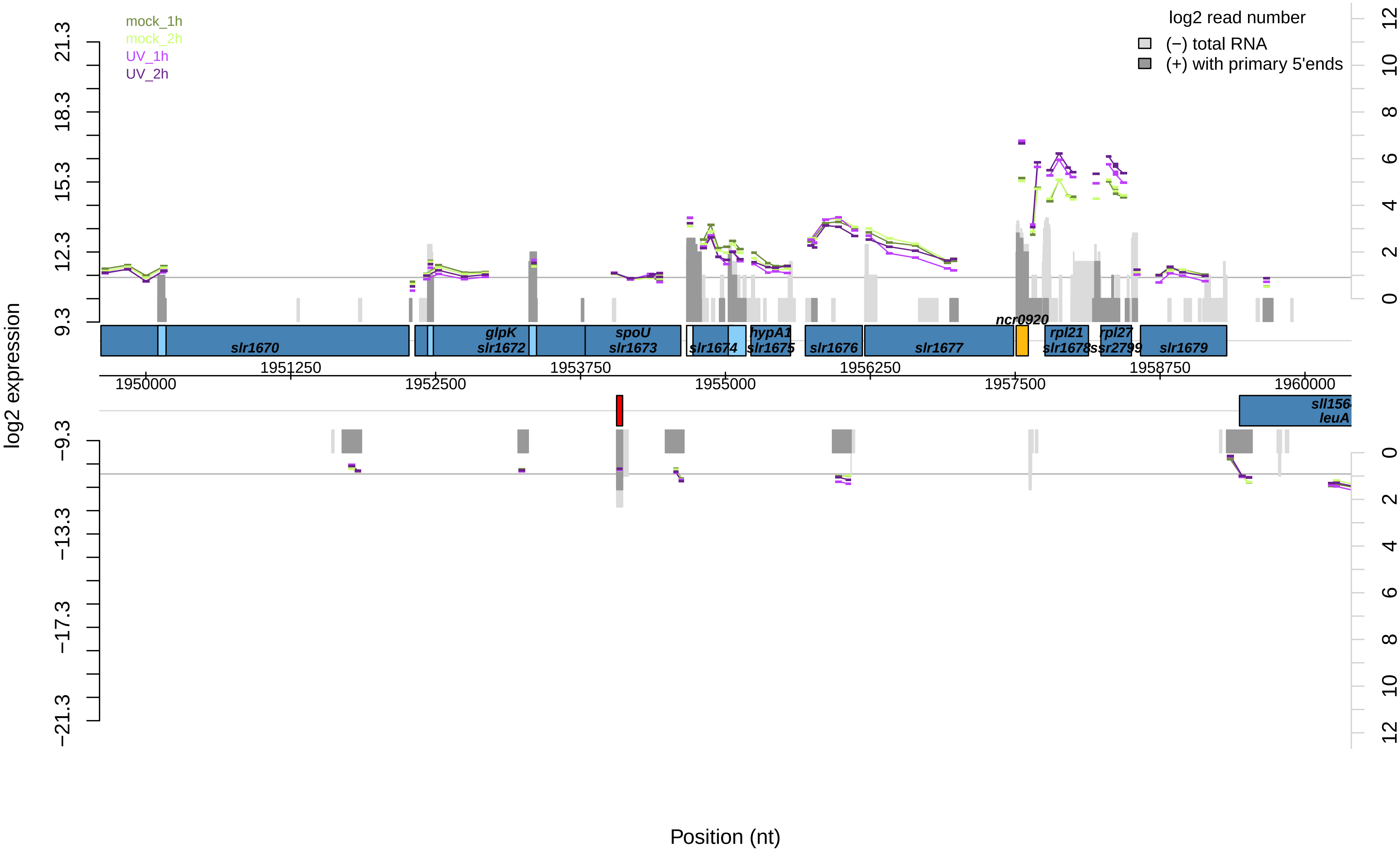

mock_1h
mock_2h
UV_1h
UV_2h
log2 read number
(−) total RNA
(+) with primary 5'ends
log2 expression
21.3
18.3
15.3
12.3
9.3
−9.3
−13.3
−17.3
−21.3
12
10
8
6
4
2
0
slr1670
glpK
slr1672
spoU
slr1673
slr1674
hypA1
slr1675
slr1676
slr1677
ncr0920
rpl21
slr1678
rpl27
ssr2799
slr1679
leuA
1951250
1953750
1956250
1958750
1950000
1952500
1955000
1957500
1960000
Position (nt)

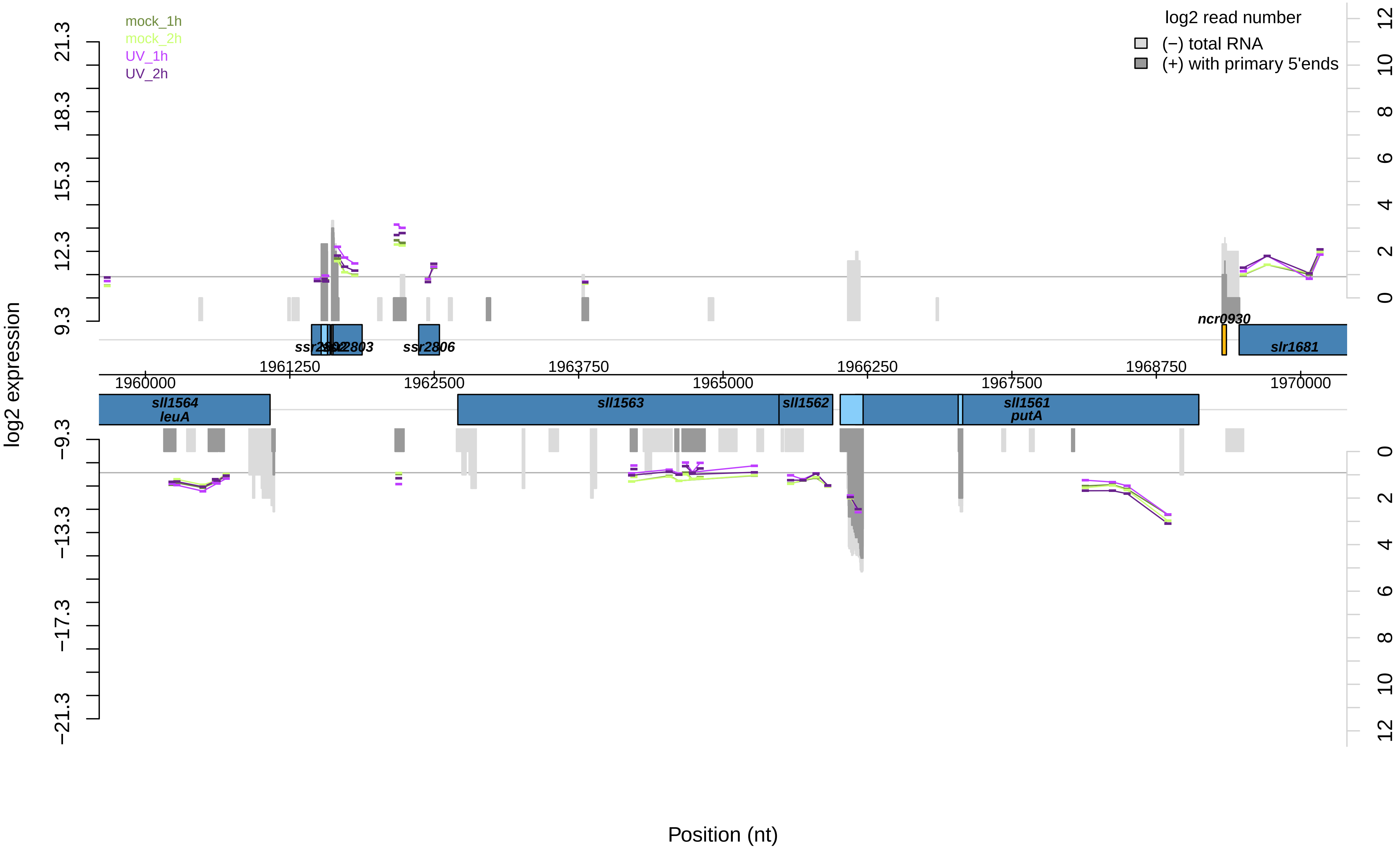

mock_1h
mock_2h
UV_1h
UV_2h
log2 read number
(−) total RNA
(+) with primary 5'ends
log2 expression
21.3
18.3
15.3
12.3
9.3
−9.3
−13.3
−17.3
−21.3
12
10
8
6
4
2
0
ssr2803
ssr2806
ncr0930
slr1681
1961250
1962500
1963750
1965000
1966250
1967500
1968750
1960000
1970000
sll1564
leuA
sll1563
sll1562
sll1561
putA
Position (nt)

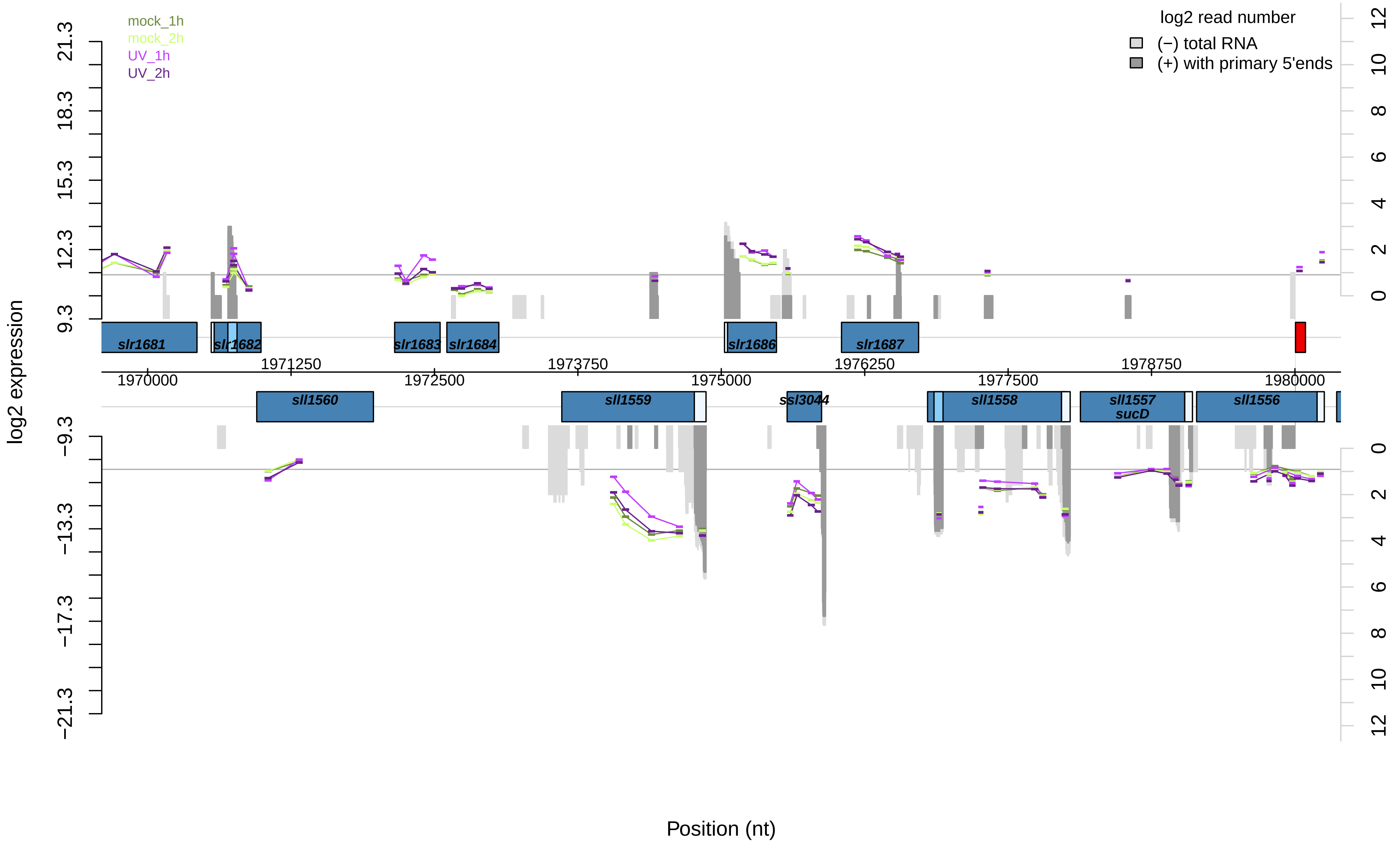

mock_1h
mock_2h
UV_1h
UV_2h
log2 read number
(−) total RNA
(+) with primary 5'ends
log2 expression
21.3
18.3
15.3
12.3
9.3
−9.3
−13.3
−17.3
−21.3
12
10
8
6
4
2
0
slr1681
slr1682
slr1683
slr1684
slr1686
slr1687
1971250
1973750
1976250
1978750
1970000
1972500
1975000
1977500
1980000
sll1560
sll1559
ssl3044
sll1558
sll1557
sucD
sll1556
Position (nt)

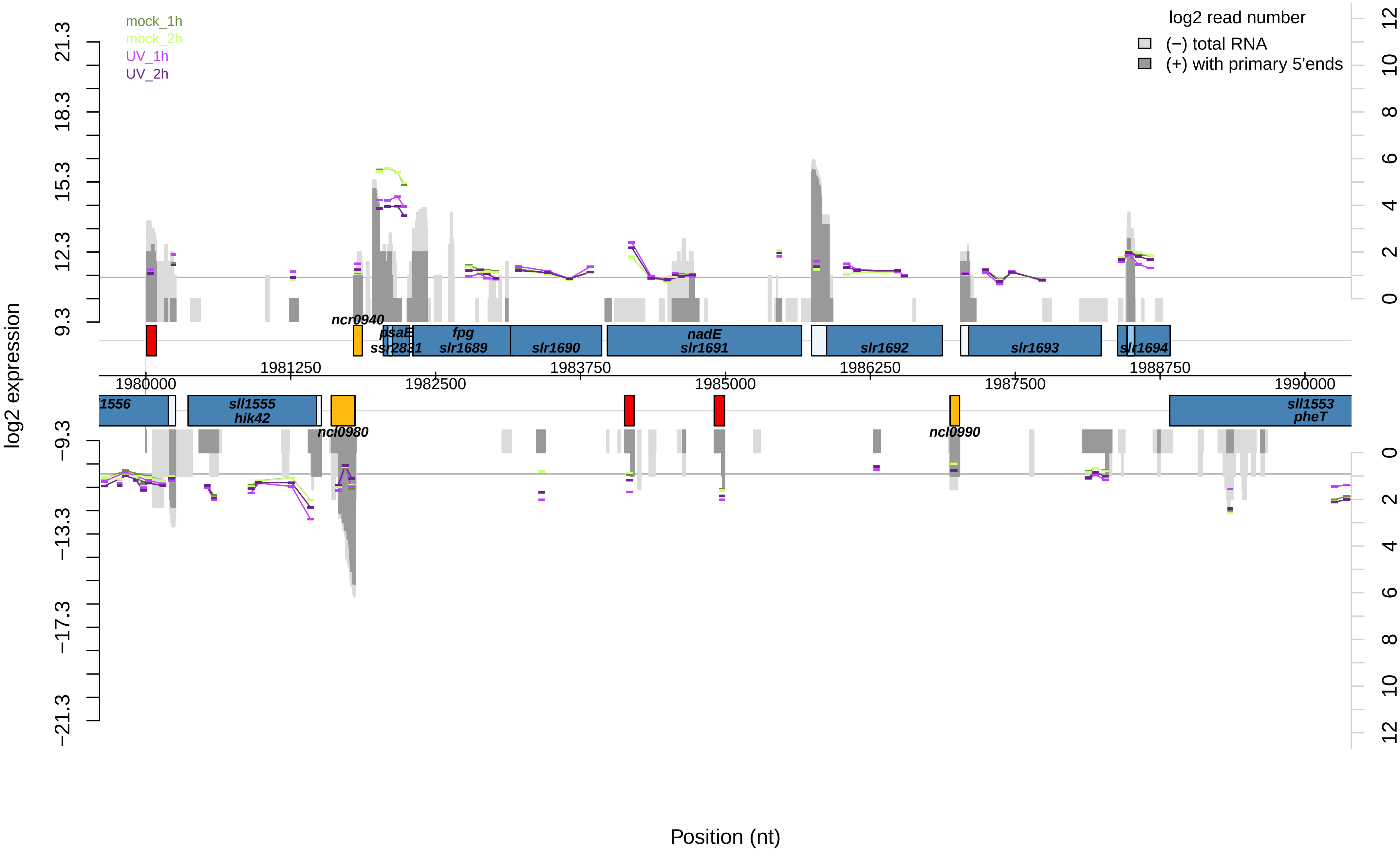
mock_1h
mock_2h
UV_1h
UV_2h
log2 read number
(−) total RNA
(+) with primary 5'ends
log2 expression
Position (nt)
ncr0940
psaE
ssr2831
fpg
slr1689
slr1690
nadE
slr1691
slr1692
slr1693
slr1694
1556
sll1555
hik42
ncl0980
ncl0990
sll1553
pheT
1980000
1981250
1982500
1983750
1985000
1986250
1987500
1988750
1990000

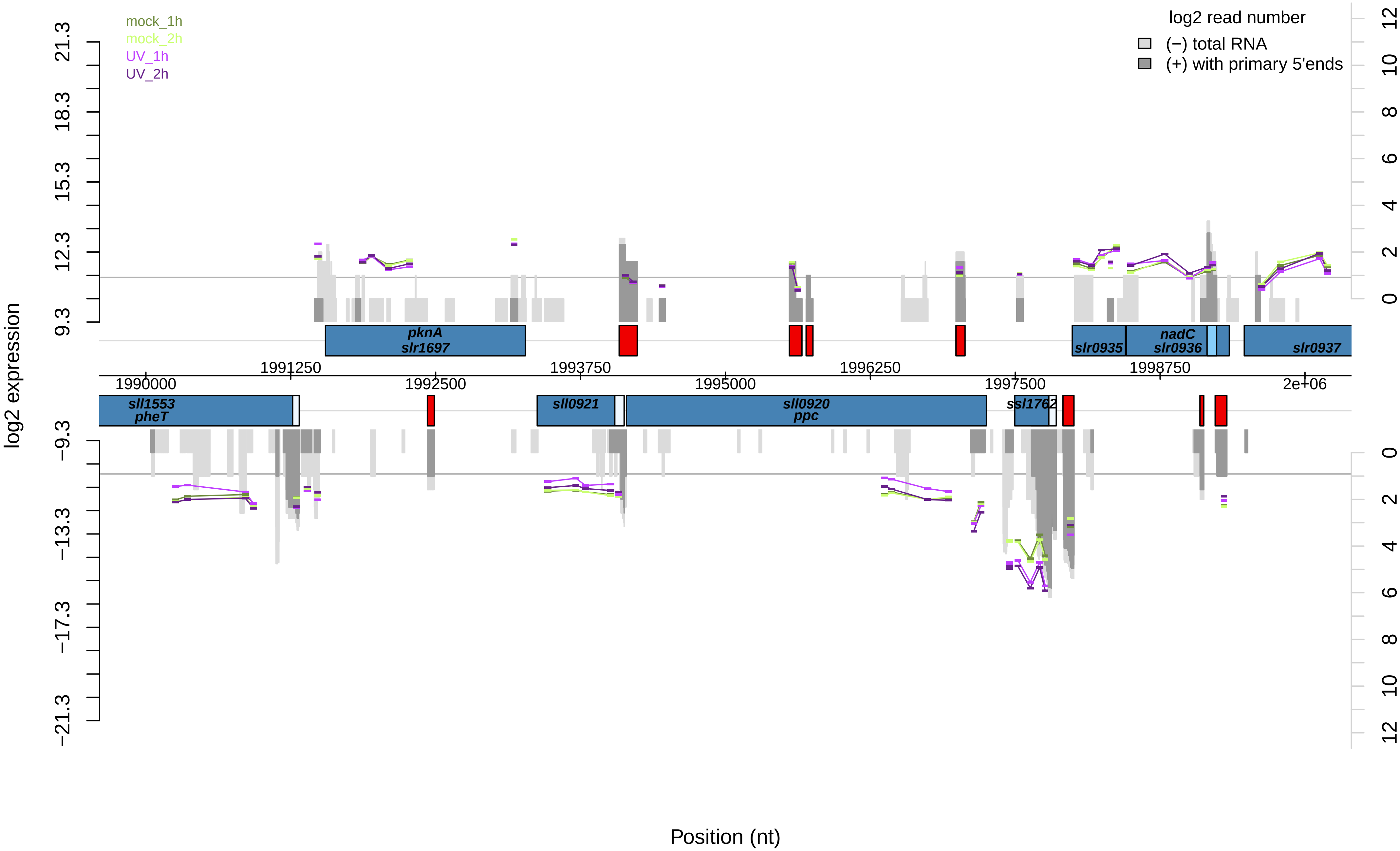

mock_1h
mock_2h
UV_1h
UV_2h
log2 read number
(−) total RNA
(+) with primary 5'ends
log2 expression
21.3
18.3
15.3
12.3
9.3
−9.3
−13.3
−17.3
−21.3
12
10
8
6
4
2
0
pknA
slr1697
slr0935
nadC
slr0936
slr0937
1991250
1993750
1996250
1998750
1990000
1992500
1995000
1997500
2e+06
sll1553
pheT
sll0921
sll0920
ppc
ssl1762
Position (nt)

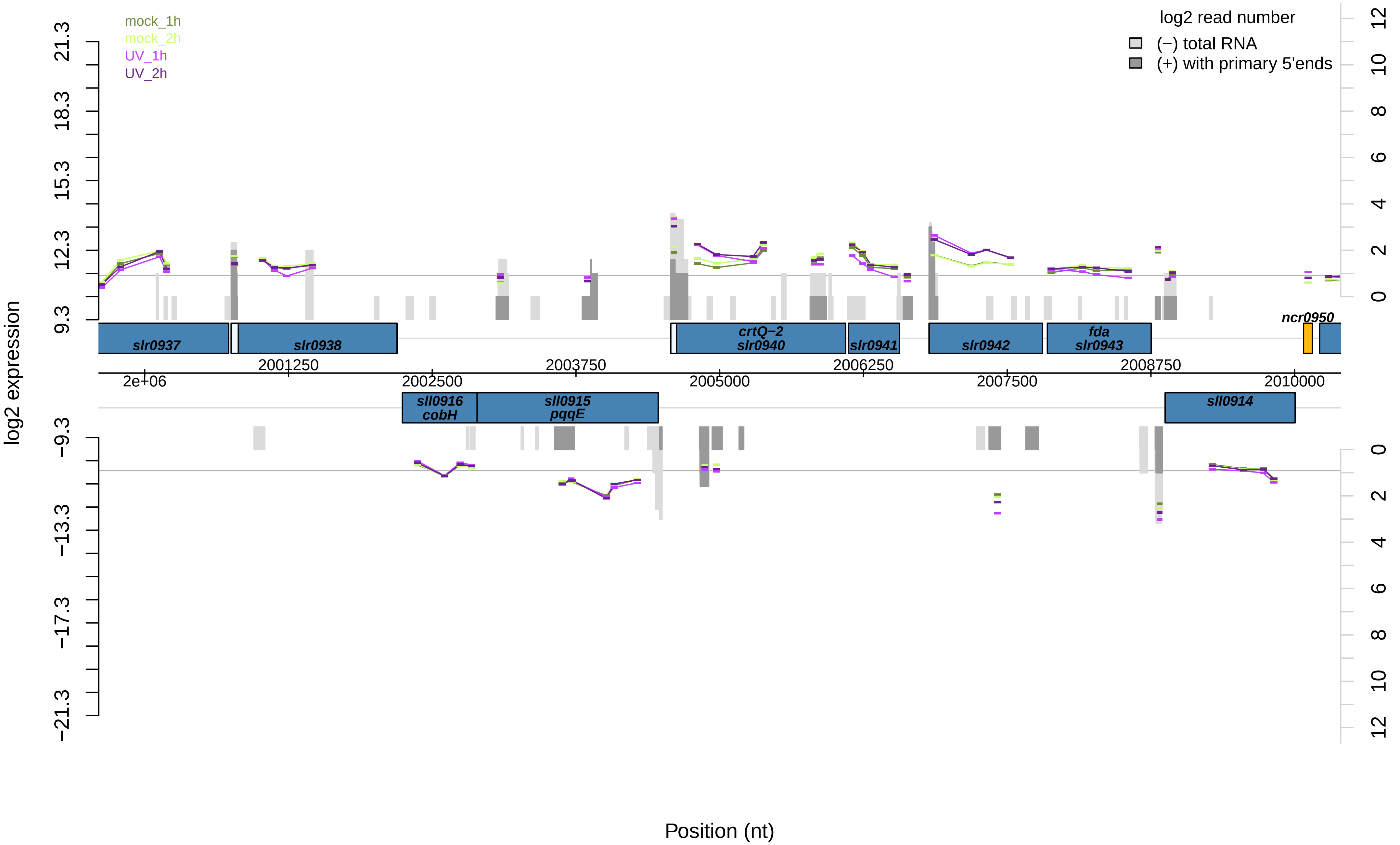
mock_1h
mock_2h
UV_1h
UV_2h
log2 read number
(−) total RNA
(+) with primary 5'ends
log2 expression
21.3
18.3
15.3
12.3
9.3
−9.3
−13.3
−17.3
−21.3
12
10
8
6
4
2
0
slr0937
slr0938
crtQ−2
slr0940
slr0941
slr0942
fda
slr0943
ncr0950
2001250
2003750
2006250
2008750
2e+06
2002500
2005000
2007500
2010000
sll0916
cobH
sll0915
pqqE
sll0914
Position (nt)

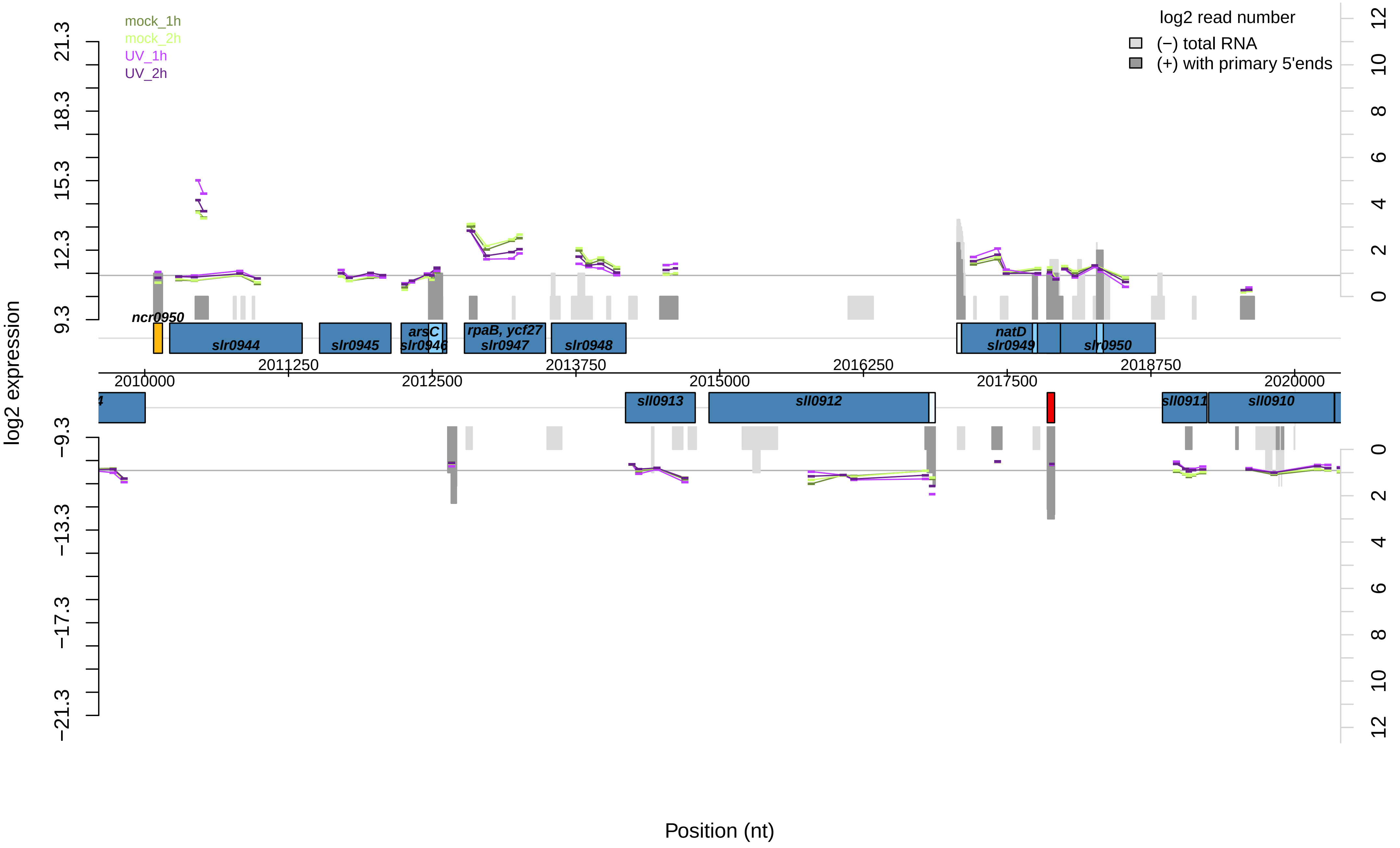
mock_1h
mock_2h
UV_1h
UV_2h
log2 read number
(−) total RNA
(+) with primary 5'ends
log2 expression
21.3
18.3
15.3
12.3
9.3
−9.3
−13.3
−17.3
−21.3
12
10
8
6
4
2
0
ncr0950
slr0944
slr0945
arsC
slr0946
rpaB, ycf27
slr0947
slr0948
natD
slr0949
slr0950
2010000
2011250
2012500
2013750
2015000
2016250
2017500
2018750
2020000
sll0913
sll0912
sll0911
sll0910
Position (nt)

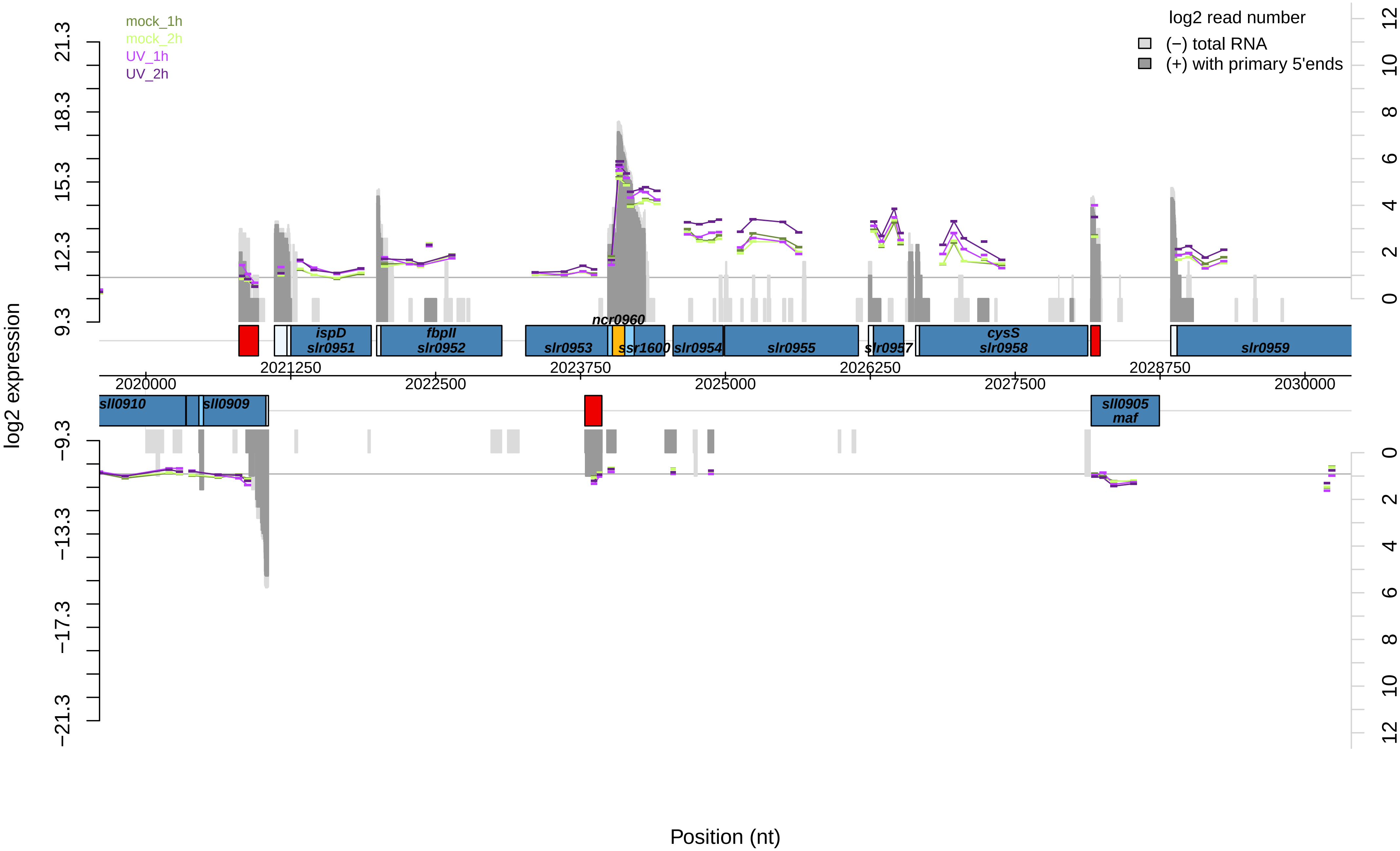

mock_1h
mock_2h
UV_1h
UV_2h
log2 read number
(−) total RNA
(+) with primary 5'ends
log2 expression
21.3
18.3
15.3
12.3
9.3
−9.3
−13.3
−17.3
−21.3
0
2
4
6
8
10
12
ispD
slr0951
fbpII
slr0952
slr0953
ncr0960
ssr1600
slr0954
slr0955
slr0957
cysS
slr0958
slr0959
sll0910
sll0909
sll0905
maf
2020000
2021250
2022500
2023750
2025000
2026250
2027500
2028750
2030000
Position (nt)

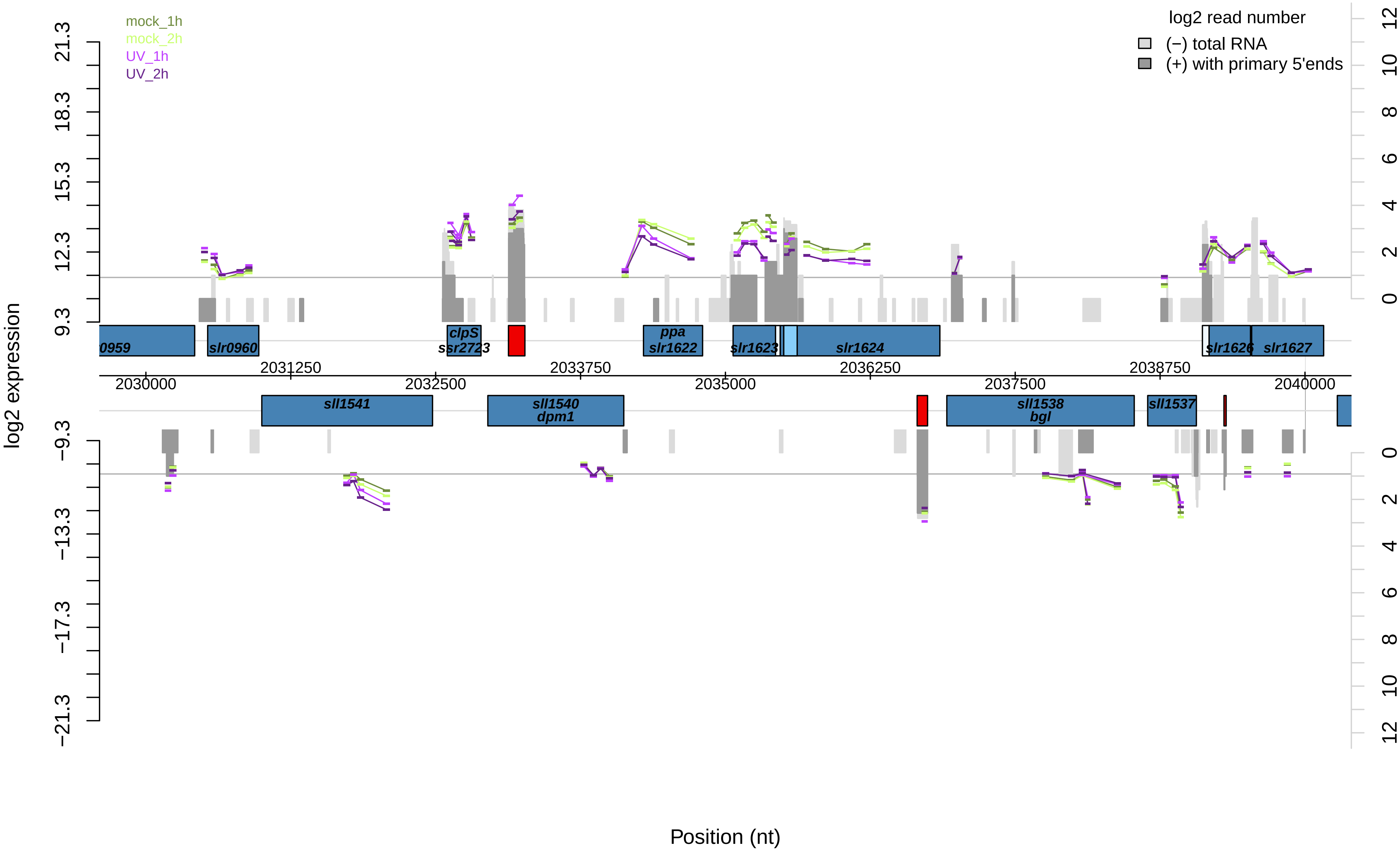

mock_1h
mock_2h
UV_1h
UV_2h
log2 read number
(−) total RNA
(+) with primary 5'ends
log2 expression
21.3
18.3
15.3
12.3
9.3
−9.3
−13.3
−17.3
−21.3
12
10
8
6
4
2
0
0959
slr0960
clpS
ssr2723
ppa
slr1622
slr1623
slr1624
slr1626
slr1627
2031250
2033750
2036250
2038750
2030000
2032500
2035000
2037500
2040000
sll1541
sll1540
dpm1
sll1538
bgl
sll1537
Position (nt)

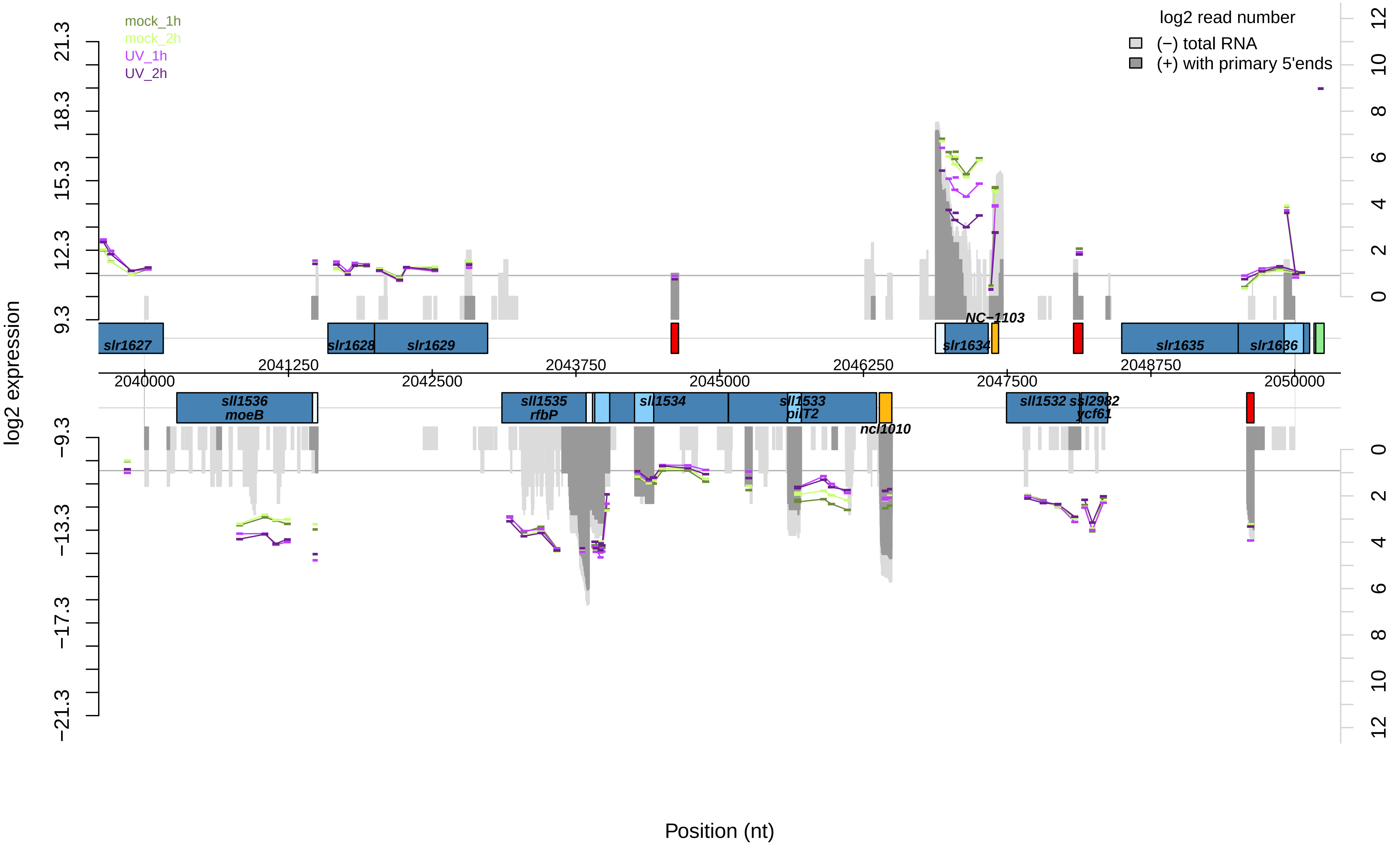

mock_1h
mock_2h
UV_1h
UV_2h
log2 read number
(−) total RNA
(+) with primary 5'ends
log2 expression
21.3
18.3
15.3
12.3
9.3
−9.3
−13.3
−17.3
−21.3
12
10
8
6
4
2
0
slr1627
slr1628
slr1629
slr1634
NC−1103
slr1635
slr1636
2041250
2043750
2046250
2048750
2040000
2042500
2045000
2047500
2050000
sll1536
moeB
sll1535
rfbP
sll1534
sll1533
pilT2
ncl1010
sll1532
ssl2982
ycf61
Position (nt)

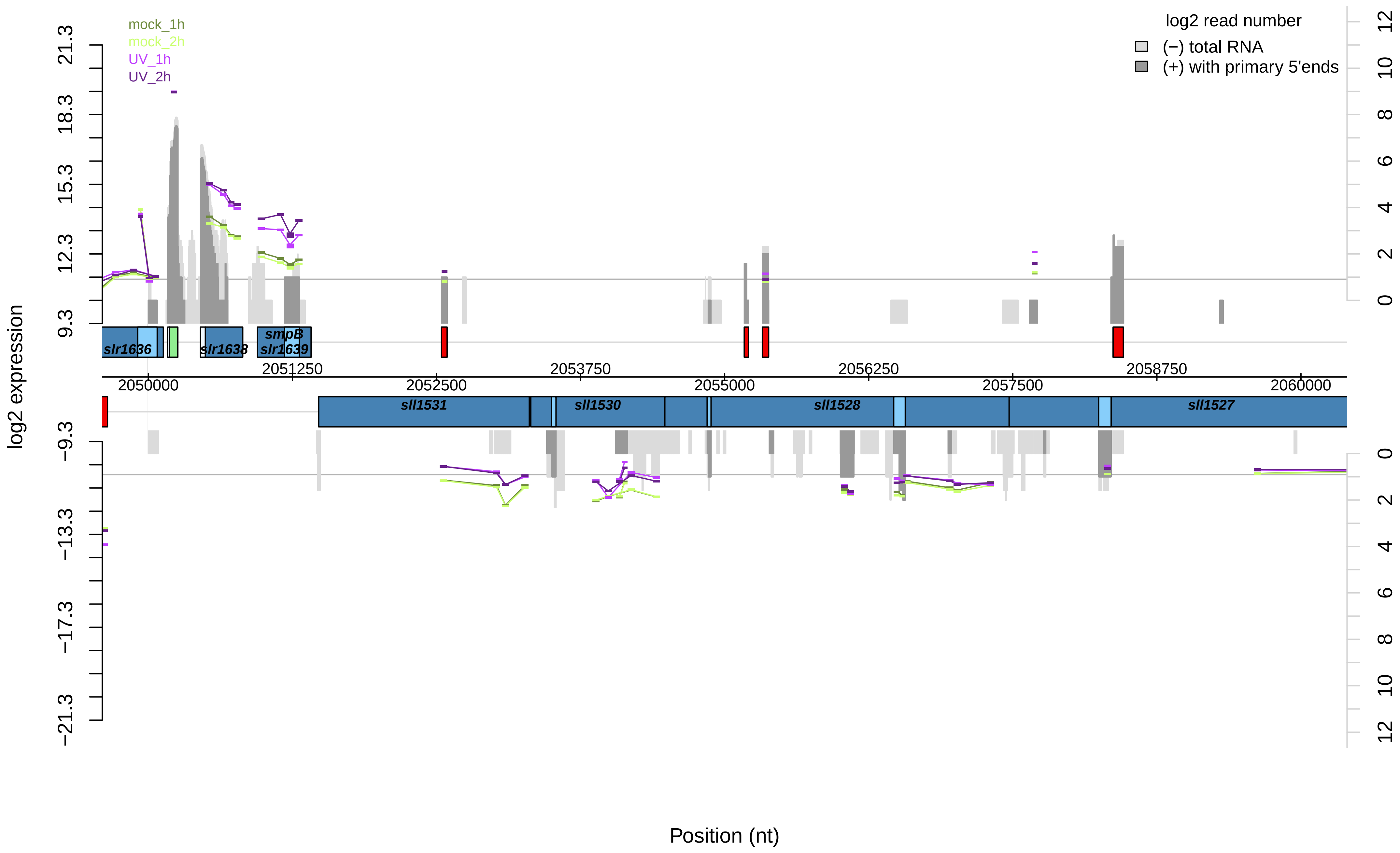

mock_1h
mock_2h
UV_1h
UV_2h
log2 read number
(−) total RNA
(+) with primary 5'ends
log2 expression
21.3
18.3
15.3
12.3
9.3
−9.3
−13.3
−17.3
−21.3
12
10
8
6
4
2
0
slr1636
slr1638
smpB
slr1639
sll1531
sll1530
sll1528
sll1527
2050000
2051250
2052500
2053750
2055000
2056250
2057500
2058750
2060000
Position (nt)

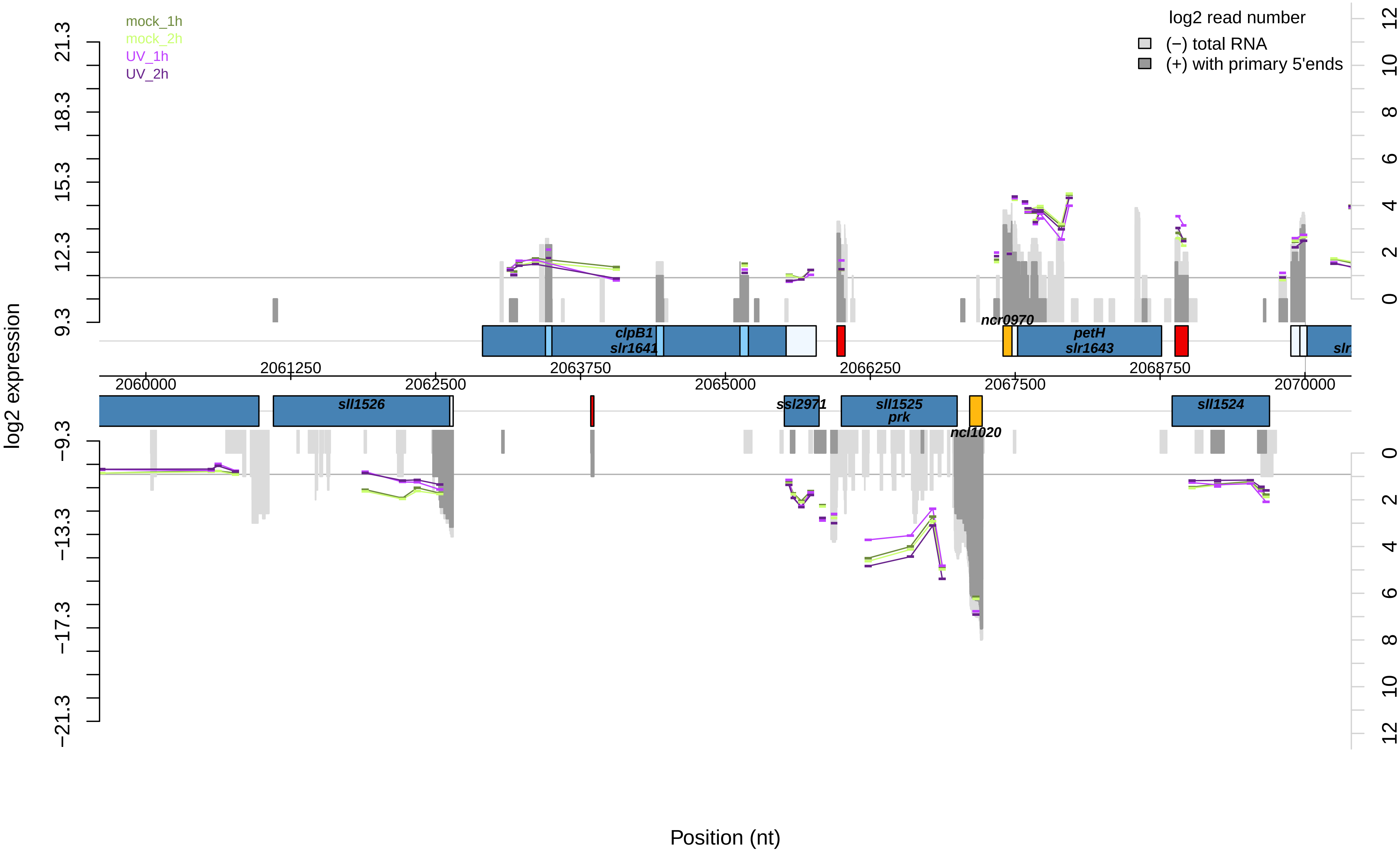
mock_1h
mock_2h
UV_1h
UV_2h
log2 read number
(−) total RNA
(+) with primary 5'ends
log2 expression
21.3
18.3
15.3
12.3
9.3
−9.3
−13.3
−17.3
−21.3
12
10
8
6
4
2
0
clpB1
slr1641
ncr0970
petH
slr1643
2060000
2061250
2062500
2063750
2065000
2066250
2067500
2068750
2070000
sll1526
ssl2971
sll1525
prk
ncl1020
sll1524
Position (nt)

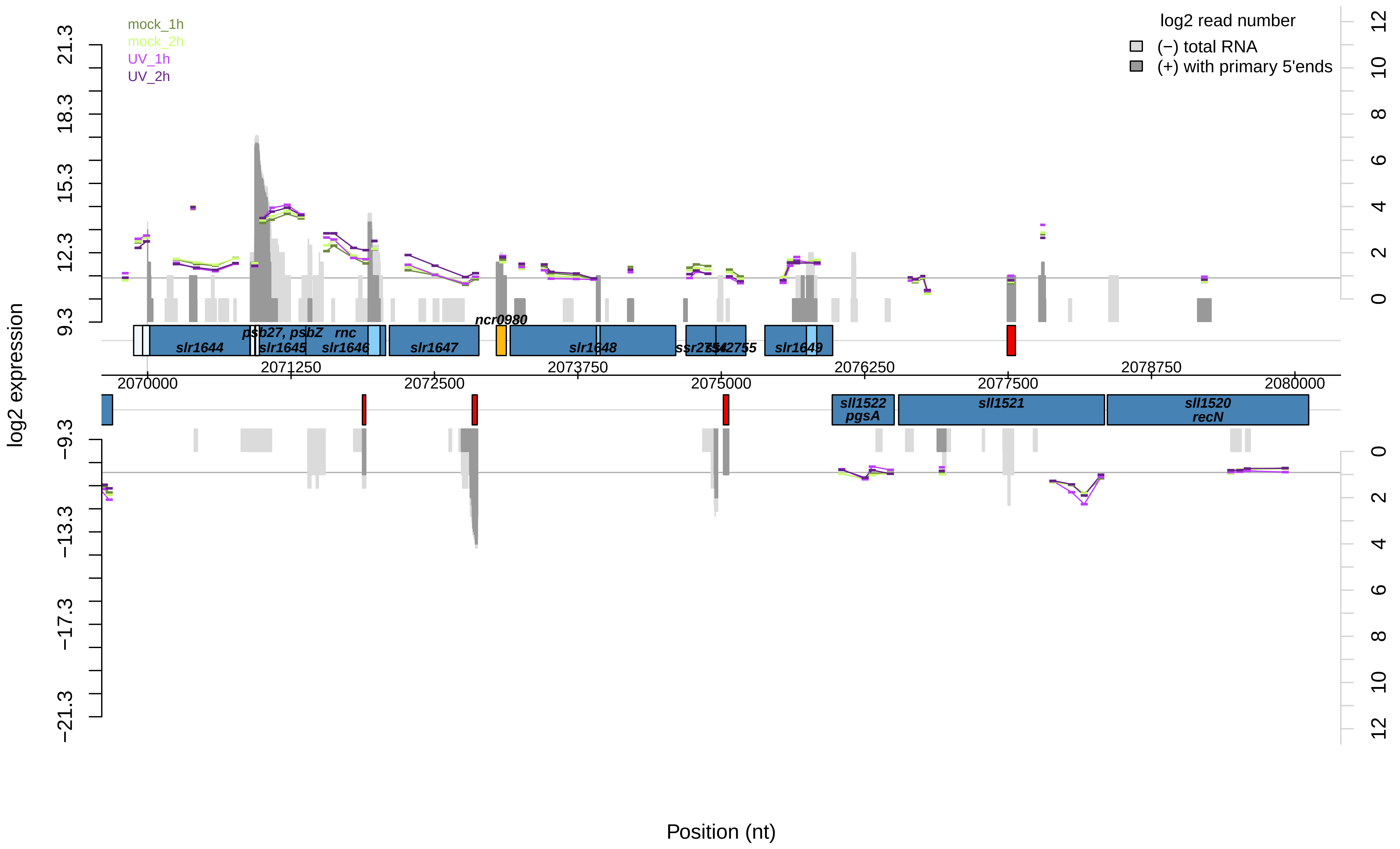

mock_1h
mock_2h
UV_1h
UV_2h
log2 read number
(−) total RNA
(+) with primary 5'ends
log2 expression
21.3
18.3
15.3
12.3
9.3
−9.3
−13.3
−17.3
−21.3
12
10
8
6
4
2
0
psb27, psbZ
rnc
slr1644
slr1645
slr1646
slr1647
ncr0980
slr1648
slr1649
2071250
2073750
2076250
2078750
2070000
2072500
2075000
2077500
2080000
sll1522
pgsA
sll1521
sll1520
recN
Position (nt)

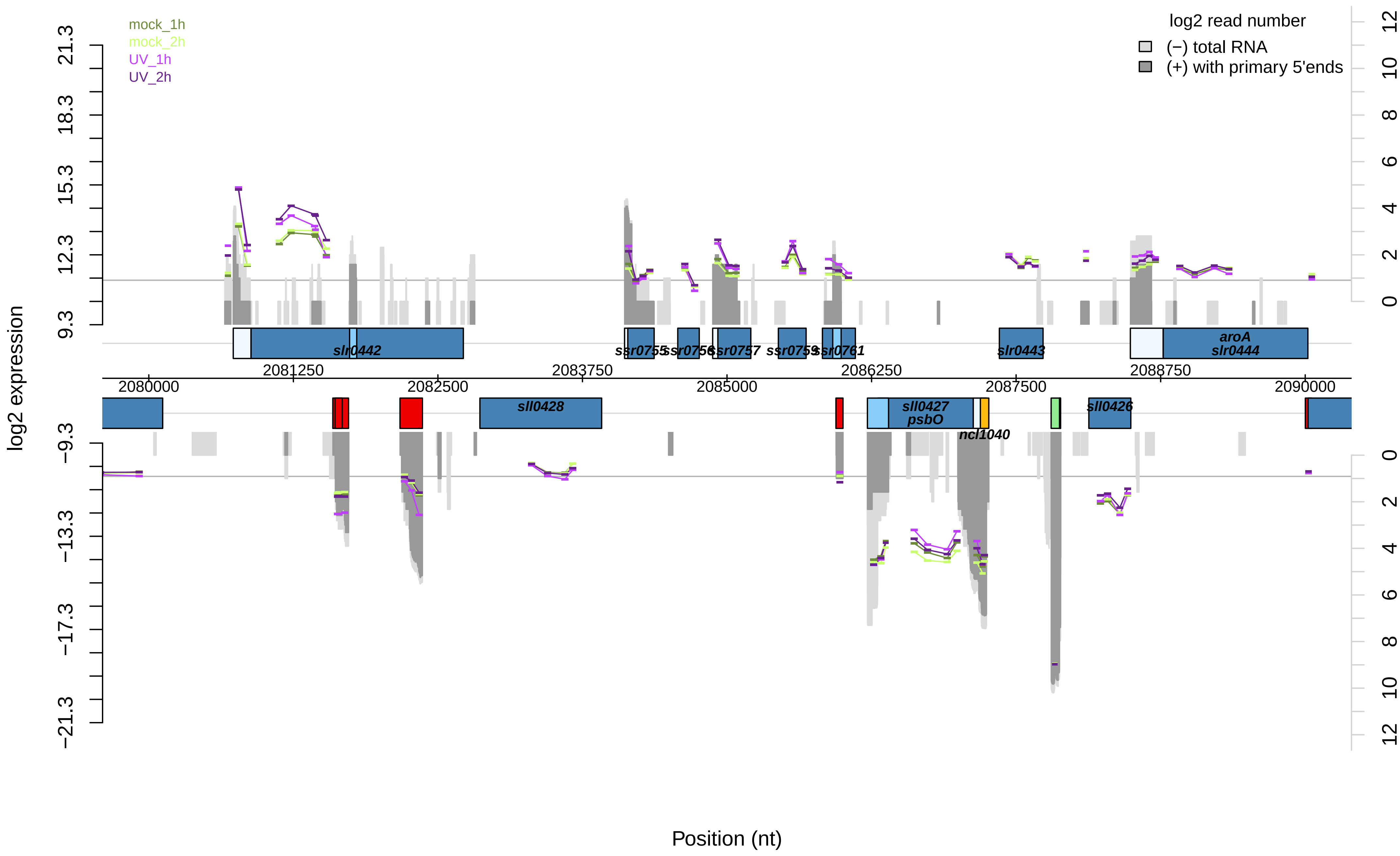

mock_1h
mock_2h
UV_1h
UV_2h
log2 read number
(−) total RNA
(+) with primary 5'ends
log2 expression
21.3
18.3
15.3
12.3
9.3
−9.3
−13.3
−17.3
−21.3
12
10
8
6
4
2
0
slr0442
ssr0755
ssr0756
ssr0757
ssr0759
ssr0761
slr0443
aroA
slr0444
2081250
2083750
2086250
2088750
2080000
2082500
2085000
2087500
2090000
sll0428
sll0427
psbO
ncl1040
sll0426
Position (nt)

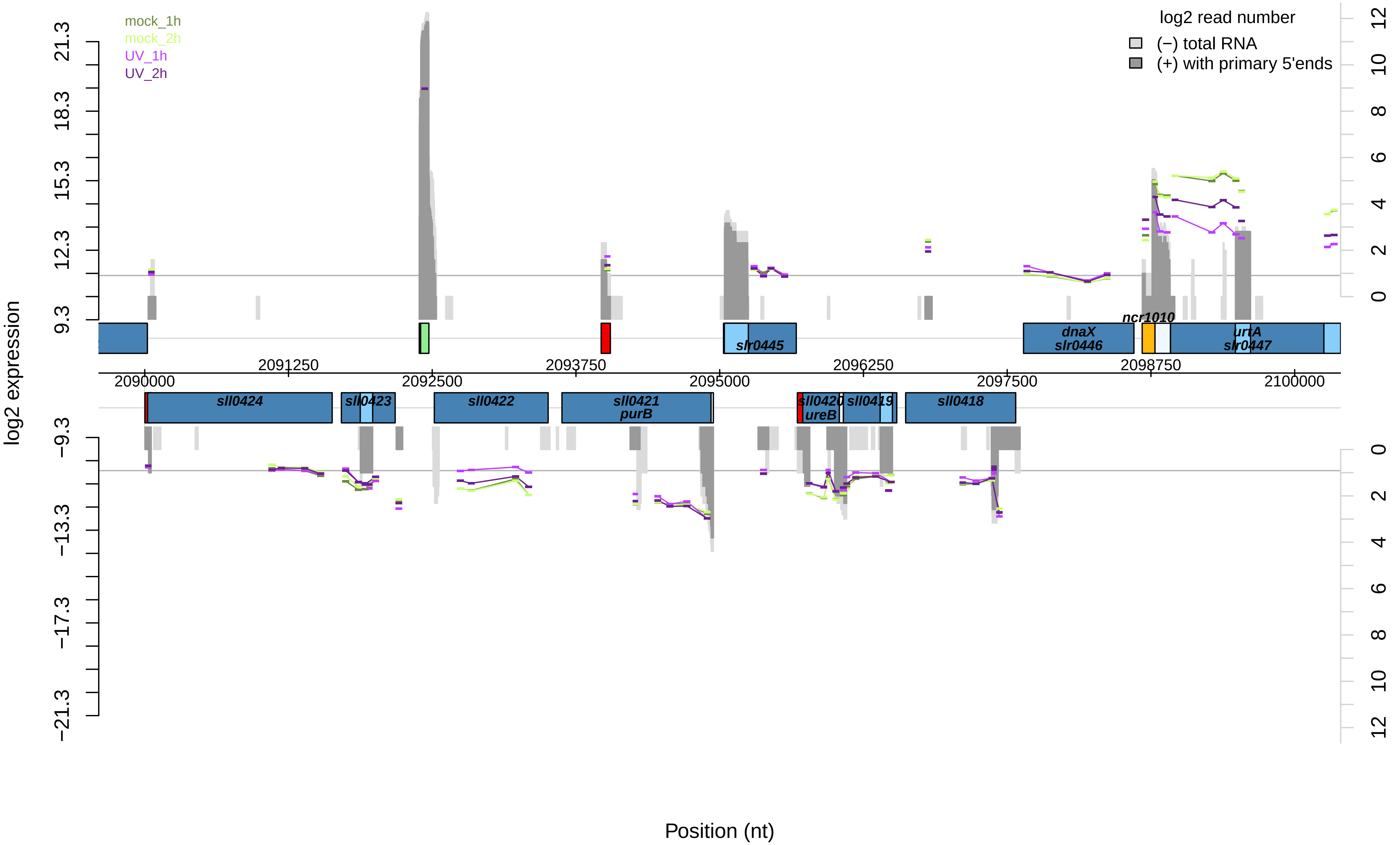

mock_1h
mock_2h
UV_1h
UV_2h
log2 read number
(−) total RNA
(+) with primary 5'ends
log2 expression
21.3
18.3
15.3
12.3
9.3
−9.3
−13.3
−17.3
−21.3
0
2
4
6
8
10
12
dnaX
slr0446
ncr1010
urtA
slr0447
slr0445
2090000
2091250
2092500
2093750
2095000
2096250
2097500
2098750
2100000
sll0424
sll0423
sll0422
sll0421
purB
sll0420
ureB
sll0419
sll0418
Position (nt)

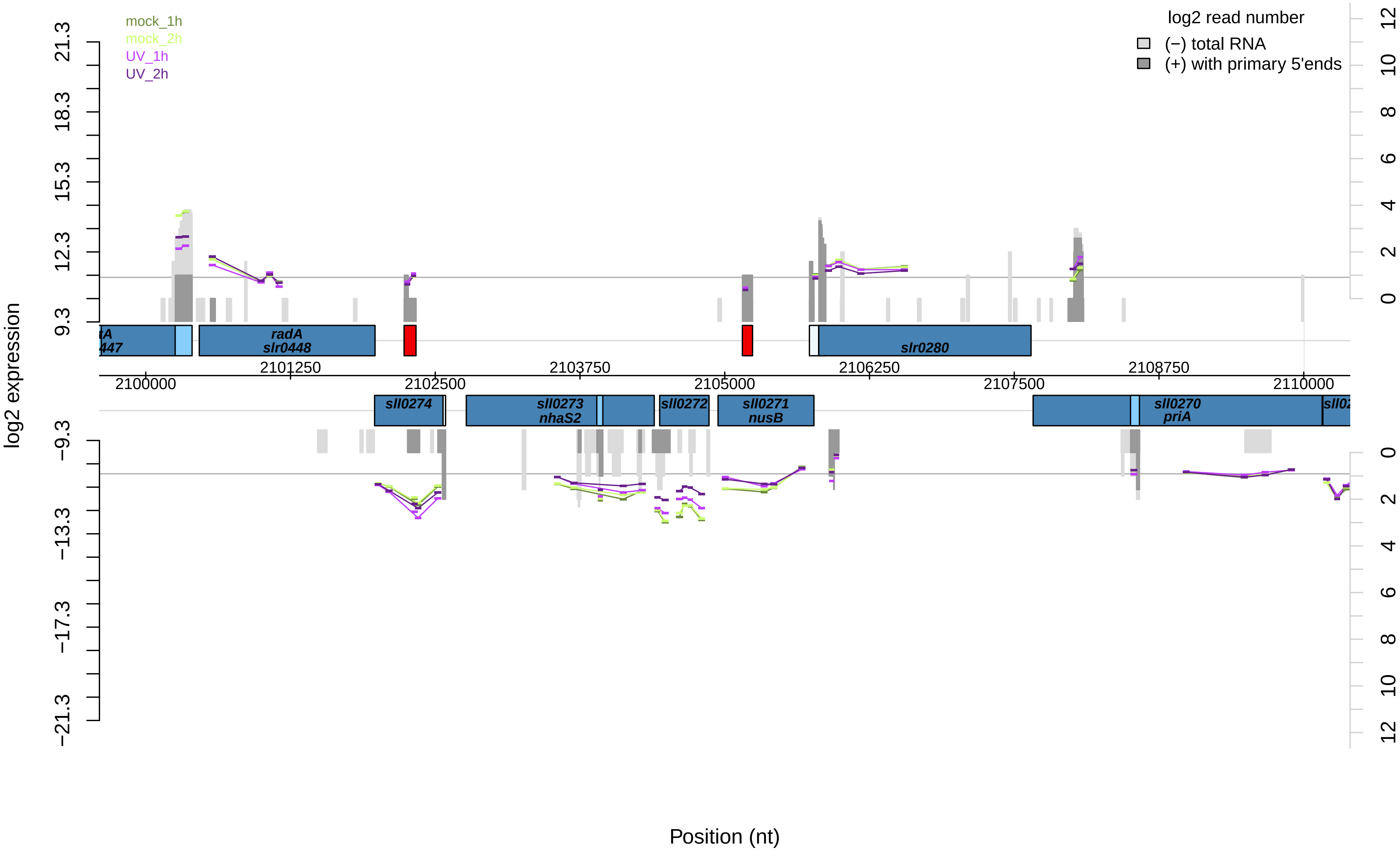

mock_1h
mock_2h
UV_1h
UV_2h
log2 read number
(−) total RNA
(+) with primary 5'ends
log2 expression
Position (nt)
radA
slr0448
slr0280
sll0274
sll0273
nhaS2
sll0272
sll0271
nusB
sll0270
priA
2100000
2101250
2102500
2103750
2105000
2106250
2107500
2108750
2110000
21.3
18.3
15.3
12.3
9.3
−9.3
−13.3
−17.3
−21.3

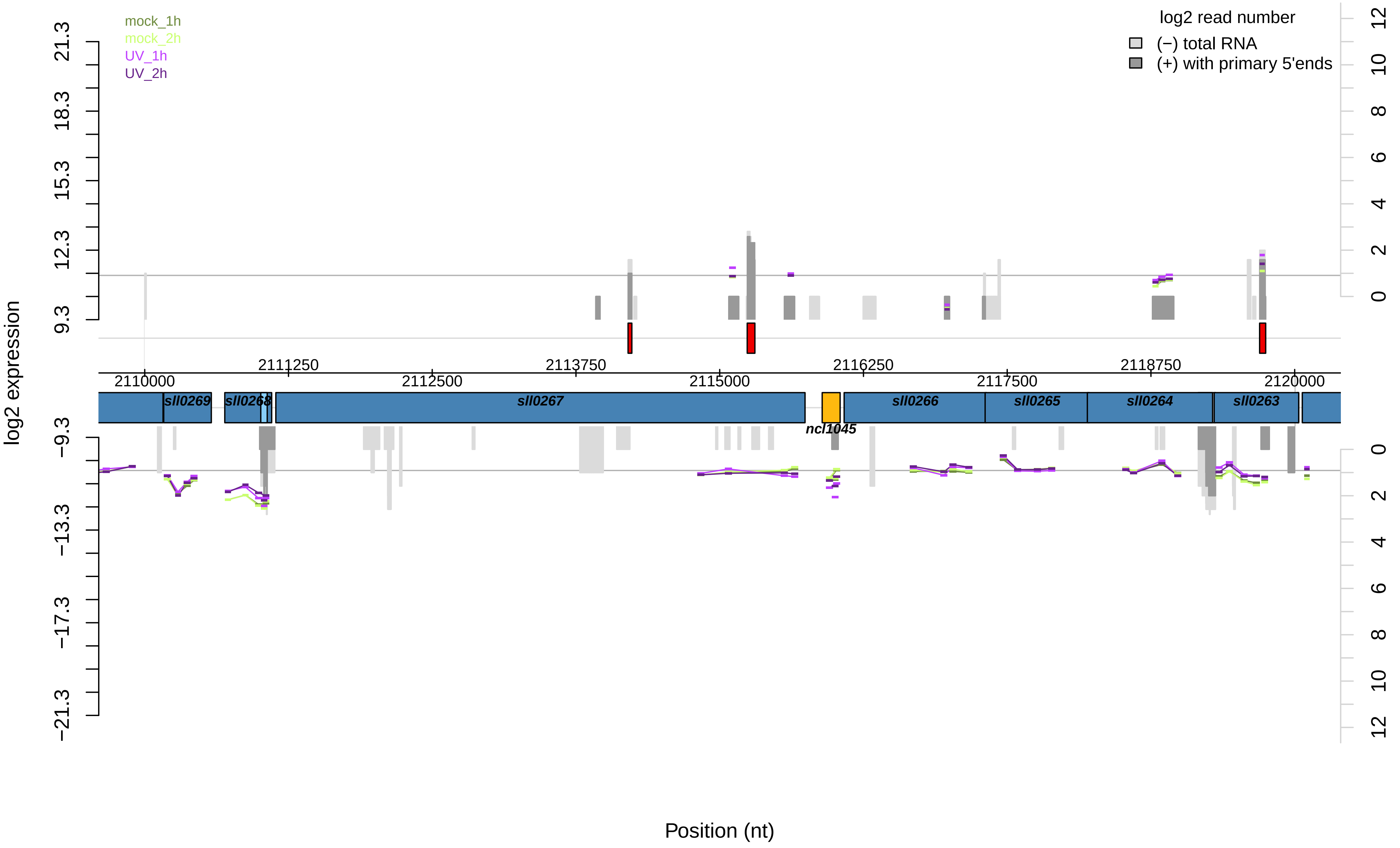

mock_1h
mock_2h
UV_1h
UV_2h
log2 read number
(−) total RNA
(+) with primary 5'ends
log2 expression
21.3
18.3
15.3
12.3
9.3
−9.3
−13.3
−17.3
−21.3
12
10
8
6
4
2
0
2110000
2111250
2112500
2113750
2115000
2116250
2117500
2118750
2120000
sll0269
sll0268
sll0267
ncl1045
sll0266
sll0265
sll0264
sll0263
Position (nt)

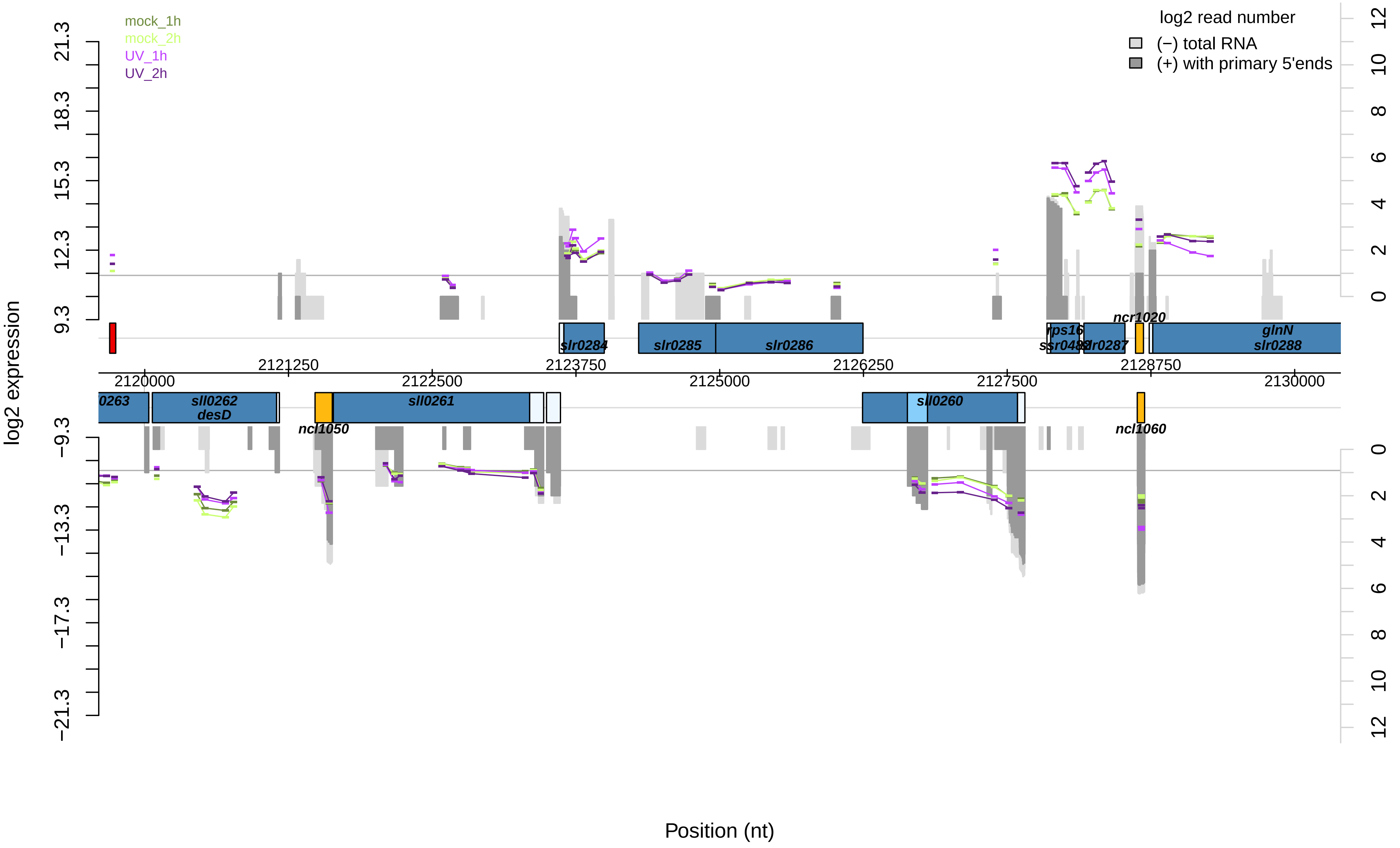

mock_1h
mock_2h
UV_1h
UV_2h
log2 read number
(−) total RNA
(+) with primary 5'ends
log2 expression
21.3
18.3
15.3
12.3
9.3
−9.3
−13.3
−17.3
−21.3
12
10
8
6
4
2
0
slr0284
slr0285
slr0286
rps16
ssr0482
slr0287
ncr1020
glnN
slr0288
2121250
2123750
2126250
2128750
2120000
2122500
2125000
2127500
2130000
0263
sll0262
desD
sll0261
ncl1050
sll0260
ncl1060
Position (nt)

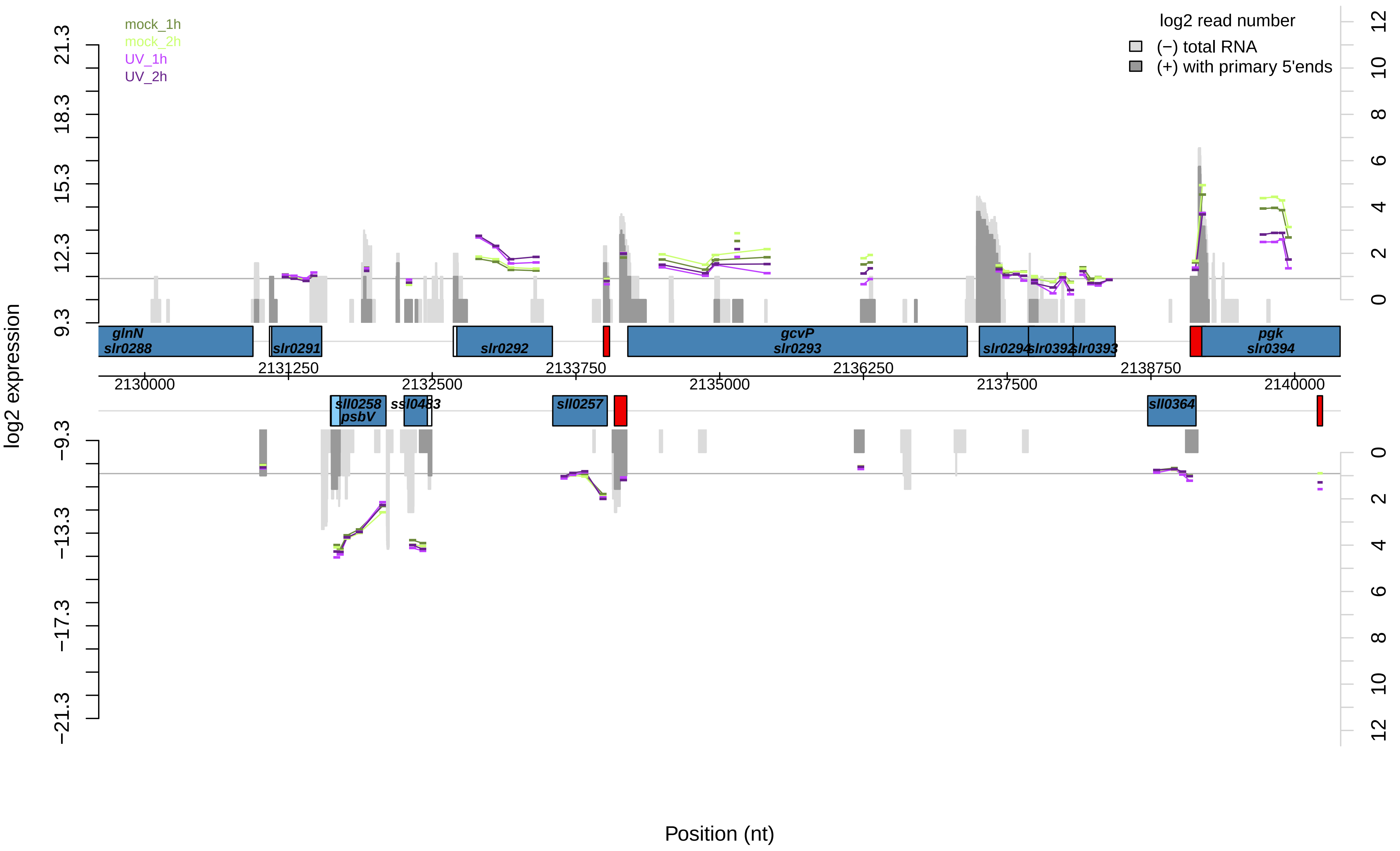
mock_1h
mock_2h
UV_1h
UV_2h
log2 read number
(−) total RNA
(+) with primary 5'ends
log2 expression
21.3
18.3
15.3
12.3
9.3
−9.3
−13.3
−17.3
−21.3
0
2
4
6
8
10
12
glnN
slr0288
slr0291
slr0292
gcvP
slr0293
slr0294
slr0392
slr0393
pgk
slr0394
2131250
2132500
2133750
2136250
2138750
2130000
2135000
2137500
2140000
sll0258
psbV
ssl0453
sll0257
sll0364
Position (nt)

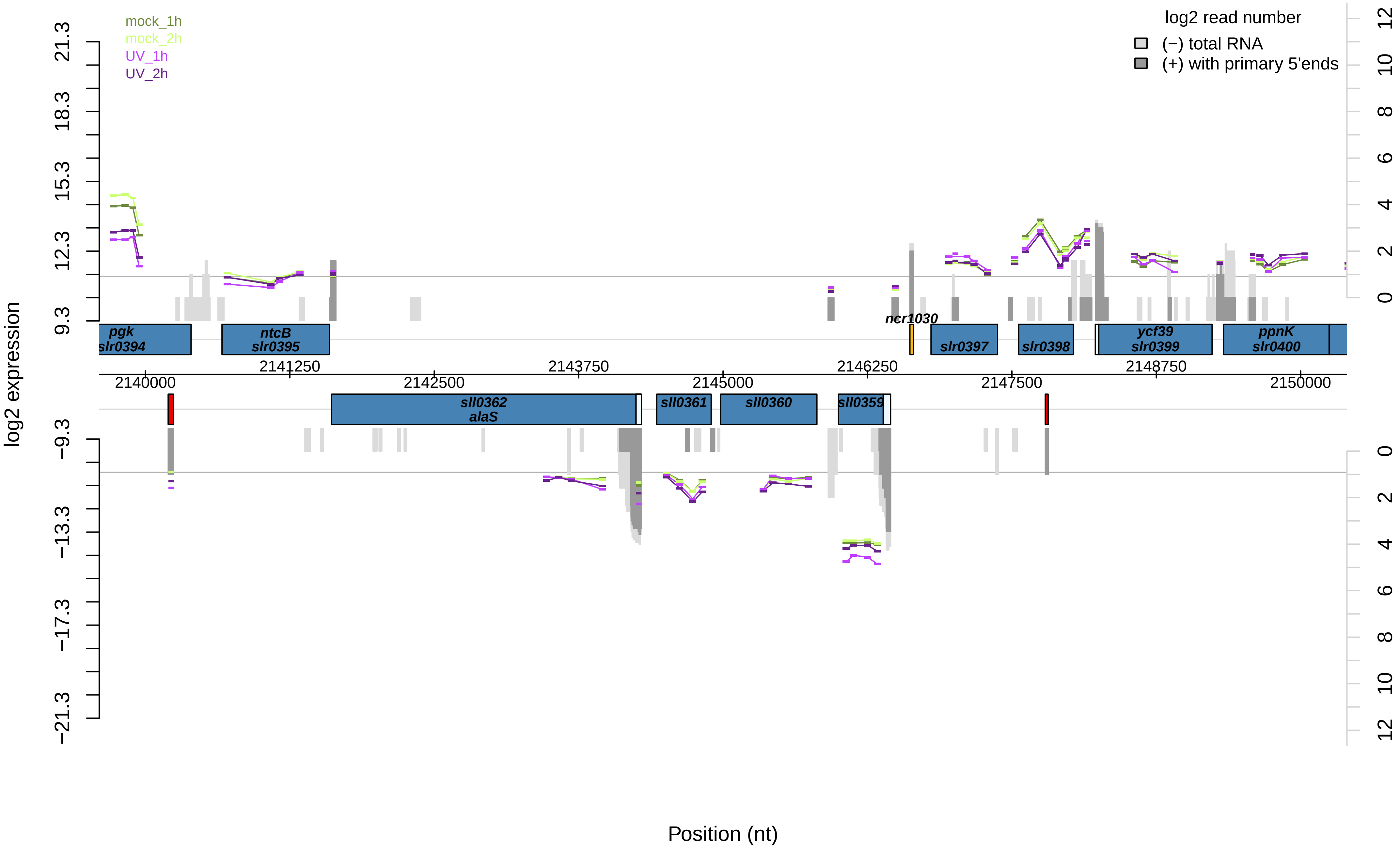

mock_1h
mock_2h
UV_1h
UV_2h
log2 read number
(−) total RNA
(+) with primary 5'ends
log2 expression
Position (nt)
21.3
18.3
15.3
12.3
9.3
−9.3
−13.3
−17.3
−21.3
12
10
8
6
4
2
0
pgk
slr0394
ntcB
slr0395
ncr1030
slr0397
slr0398
ycf39
slr0399
ppnK
slr0400
2141250
2143750
2146250
2148750
2140000
2142500
2145000
2147500
2150000
sll0362
alaS
sll0361
sll0360
sll0359

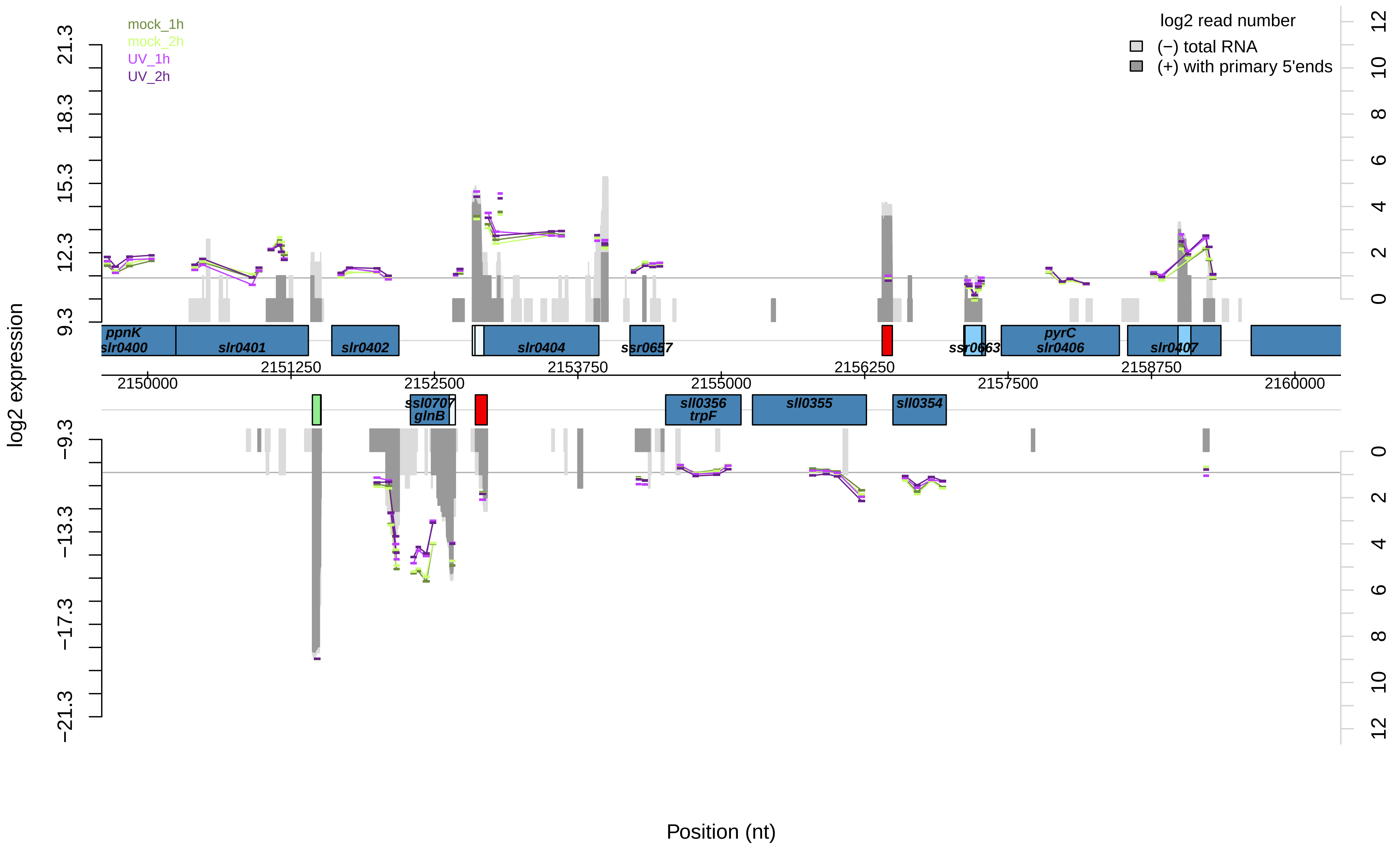

mock_1h
mock_2h
UV_1h
UV_2h
log2 read number
(−) total RNA
(+) with primary 5'ends
log2 expression
Position (nt)
21.3
18.3
15.3
12.3
9.3
−9.3
−13.3
−17.3
−21.3
12
10
8
6
4
2
0
ppnK
slr0400
slr0401
slr0402
slr0404
ssr0657
ssr0663
pyrC
slr0406
slr0407
ssl0707
glnB
sll0356
trpF
sll0355
sll0354
2150000
2151250
2152500
2153750
2155000
2156250
2157500
2158750
2160000

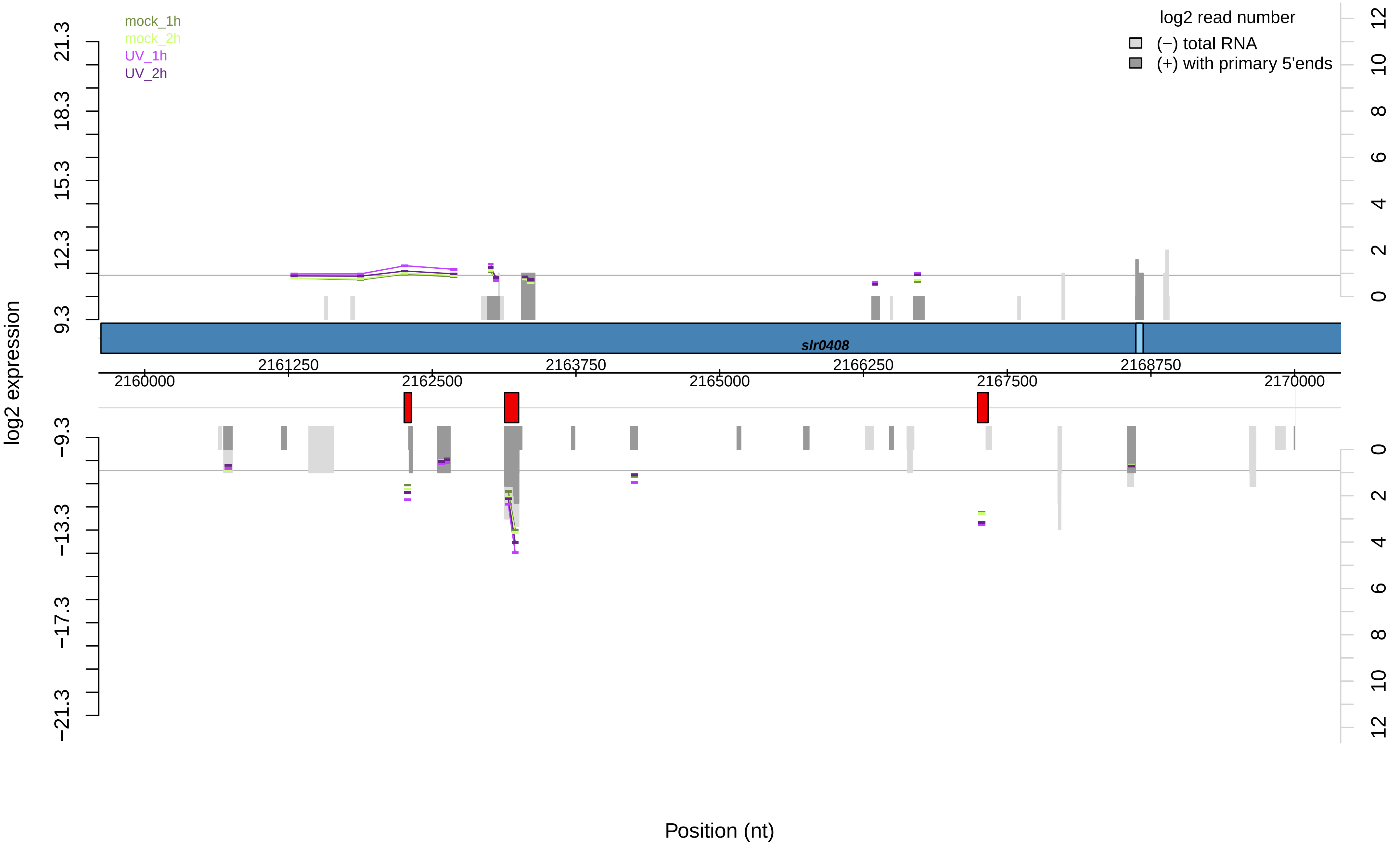

mock_1h
mock_2h
UV_1h
UV_2h
log2 read number
(−) total RNA
(+) with primary 5'ends
slr0408
log2 expression
Position (nt)
21.3
18.3
15.3
12.3
9.3
−9.3
−13.3
−17.3
−21.3
2160000
2161250
2162500
2163750
2165000
2166250
2167500
2168750
2170000

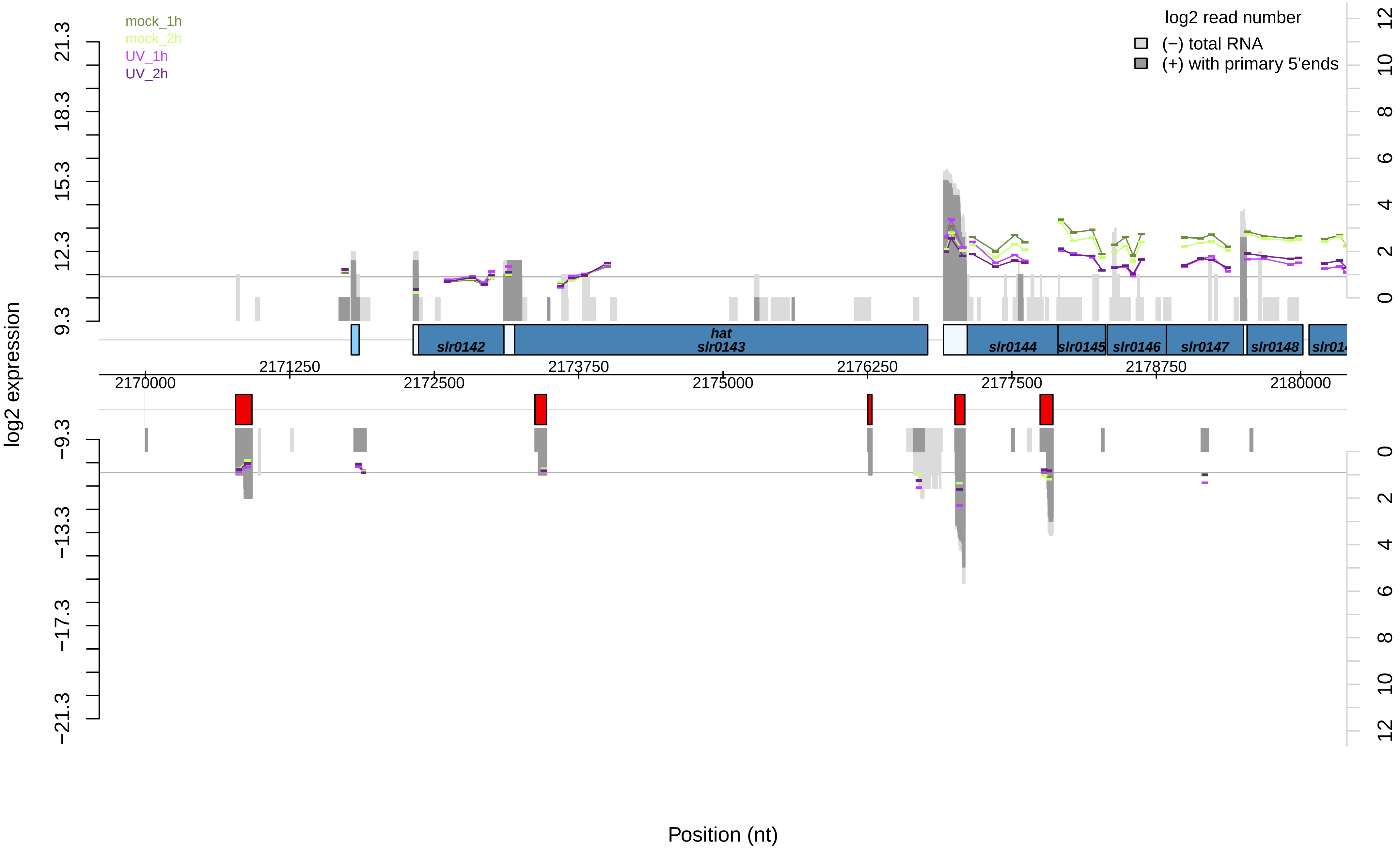

mock_1h
mock_2h
UV_1h
UV_2h
log2 read number
(−) total RNA
(+) with primary 5'ends
log2 expression
21.3
18.3
15.3
12.3
9.3
−9.3
−13.3
−17.3
−21.3
12
10
8
6
4
2
0
hat
slr0142
slr0143
slr0144
slr0145
slr0146
slr0147
slr0148
2170000
2171250
2172500
2173750
2175000
2176250
2177500
2178750
2180000
Position (nt)

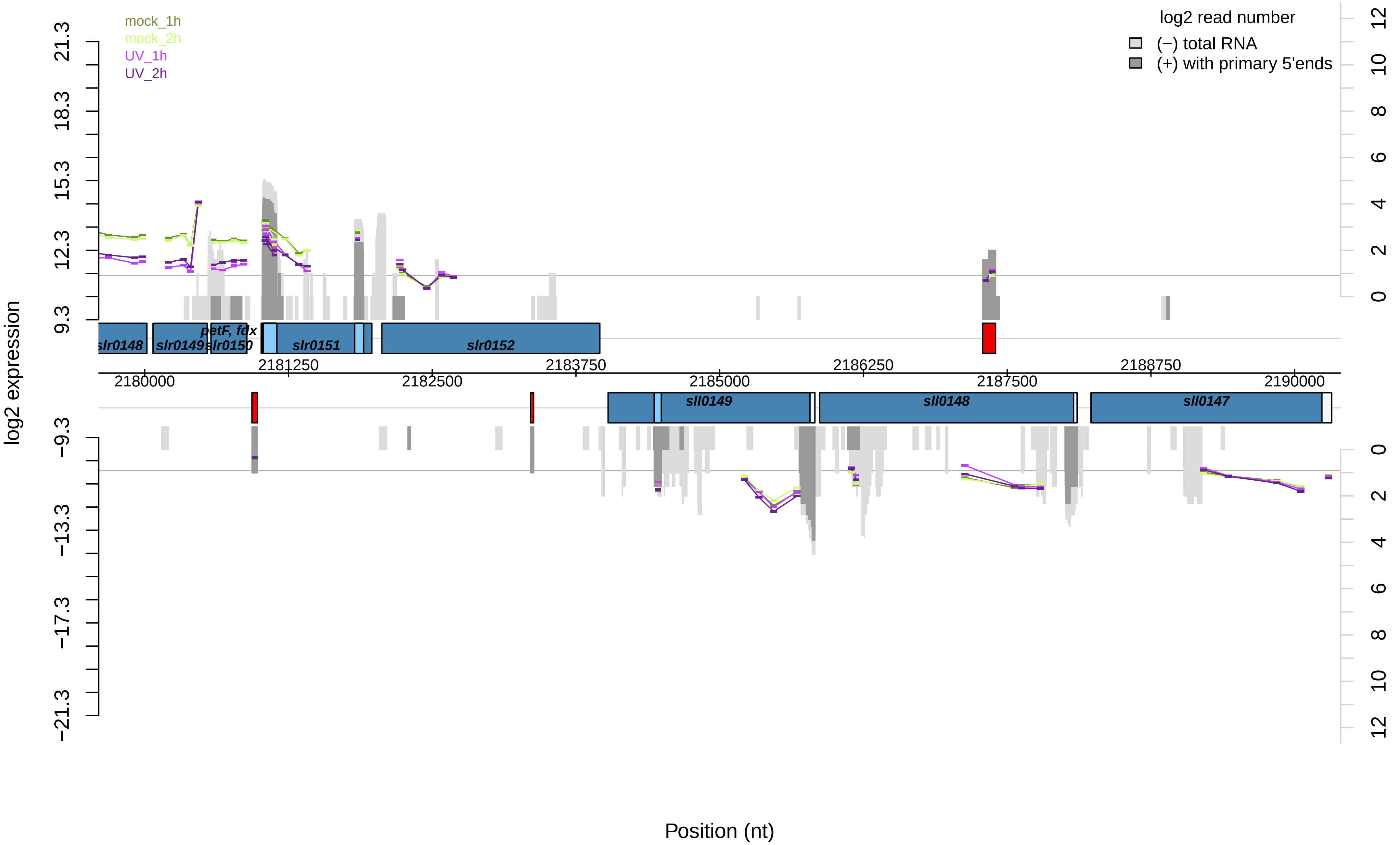

mock_1h
mock_2h
UV_1h
UV_2h
log2 read number
(−) total RNA
(+) with primary 5'ends
log2 expression
21.3
18.3
15.3
12.3
9.3
−9.3
−13.3
−17.3
−21.3
12
10
8
6
4
2
0
slr0148
slr0149
petF, fdx
slr0150
slr0151
slr0152
sll0149
sll0148
sll0147
2180000
2181250
2182500
2183750
2185000
2186250
2187500
2188750
2190000
Position (nt)

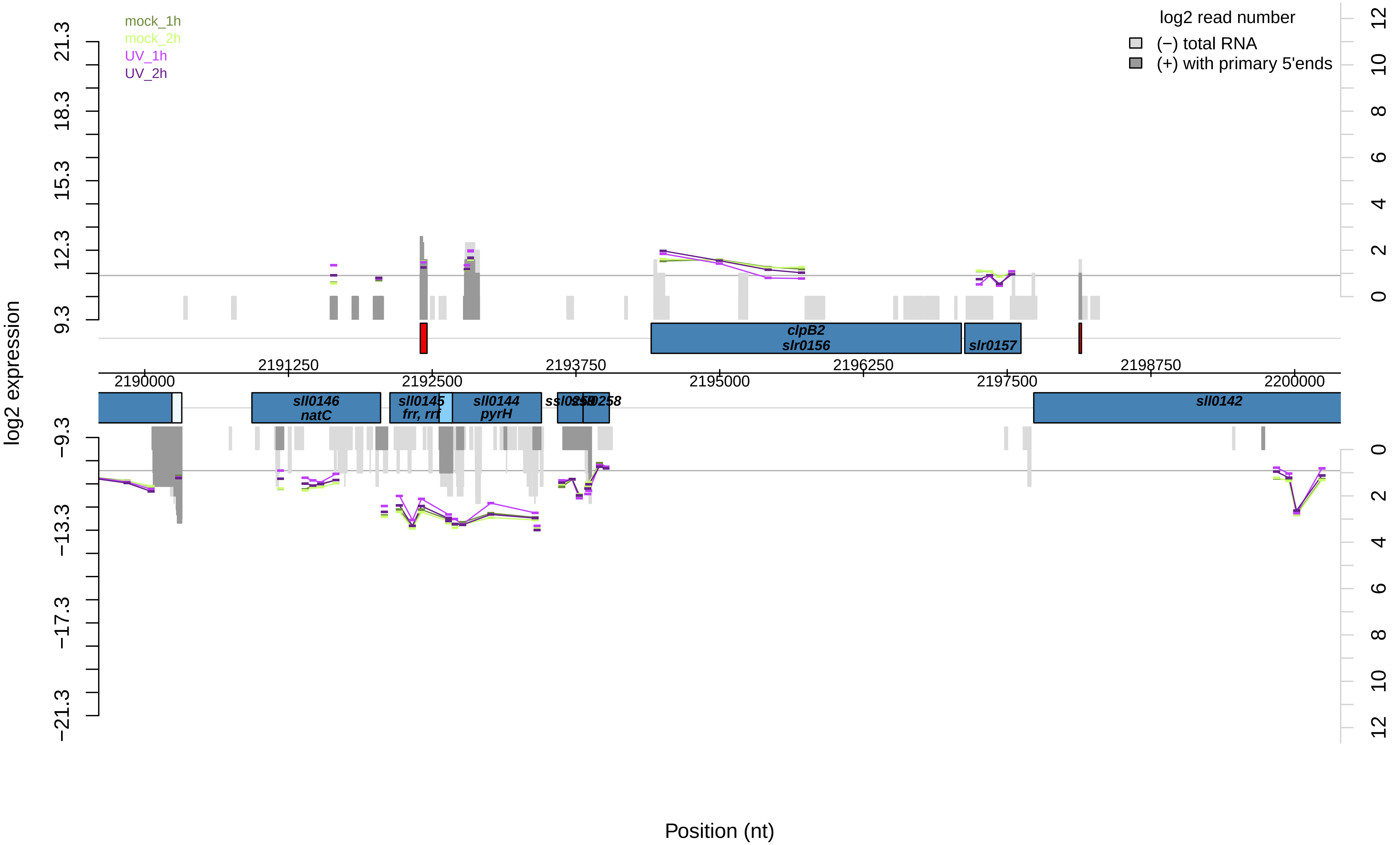

mock_1h
mock_2h
UV_1h
UV_2h
log2 read number
(−) total RNA
(+) with primary 5'ends
log2 expression
21.3
18.3
15.3
12.3
9.3
−9.3
−13.3
−17.3
−21.3
12
10
8
6
4
2
0
clpB2
slr0156
slr0157
2190000
2191250
2192500
2193750
2195000
2196250
2197500
2198750
2200000
sll0146
natC
sll0145
frr, rrf
sll0144
pyrH
sll0142
Position (nt)

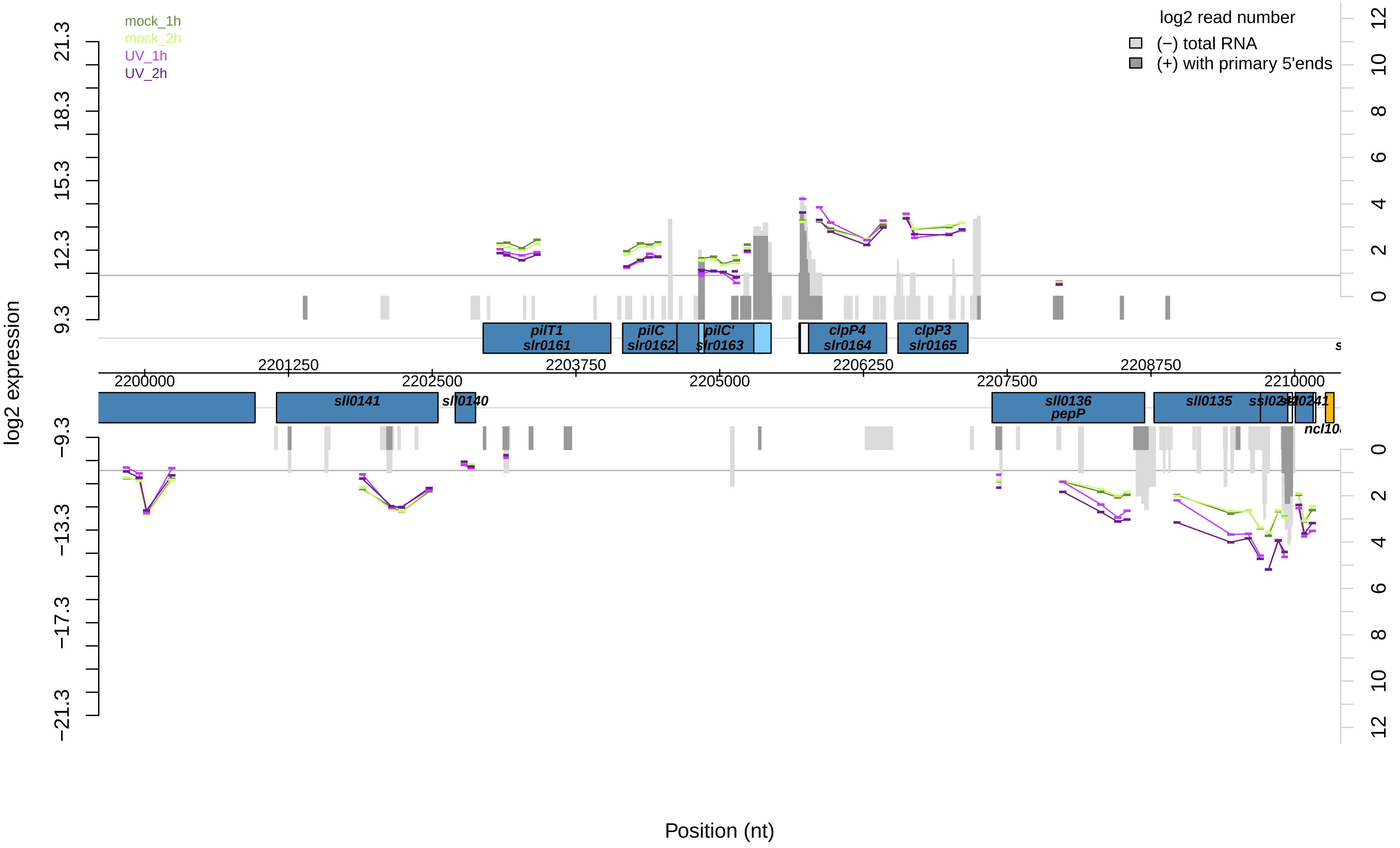
mock_1h
mock_2h
UV_1h
UV_2h
log2 read number
(−) total RNA
(+) with primary 5'ends
log2 expression
21.3
18.3
15.3
12.3
9.3
−9.3
−13.3
−17.3
−21.3
12
10
8
6
4
2
0
pilT1
slr0161
pilC
slr0162
pilC'
slr0163
clpP4
slr0164
clpP3
slr0165
2201250
2203750
2206250
2208750
2200000
2202500
2205000
2207500
2210000
sll0141
sll0140
sll0136
pepP
sll0135
Position (nt)

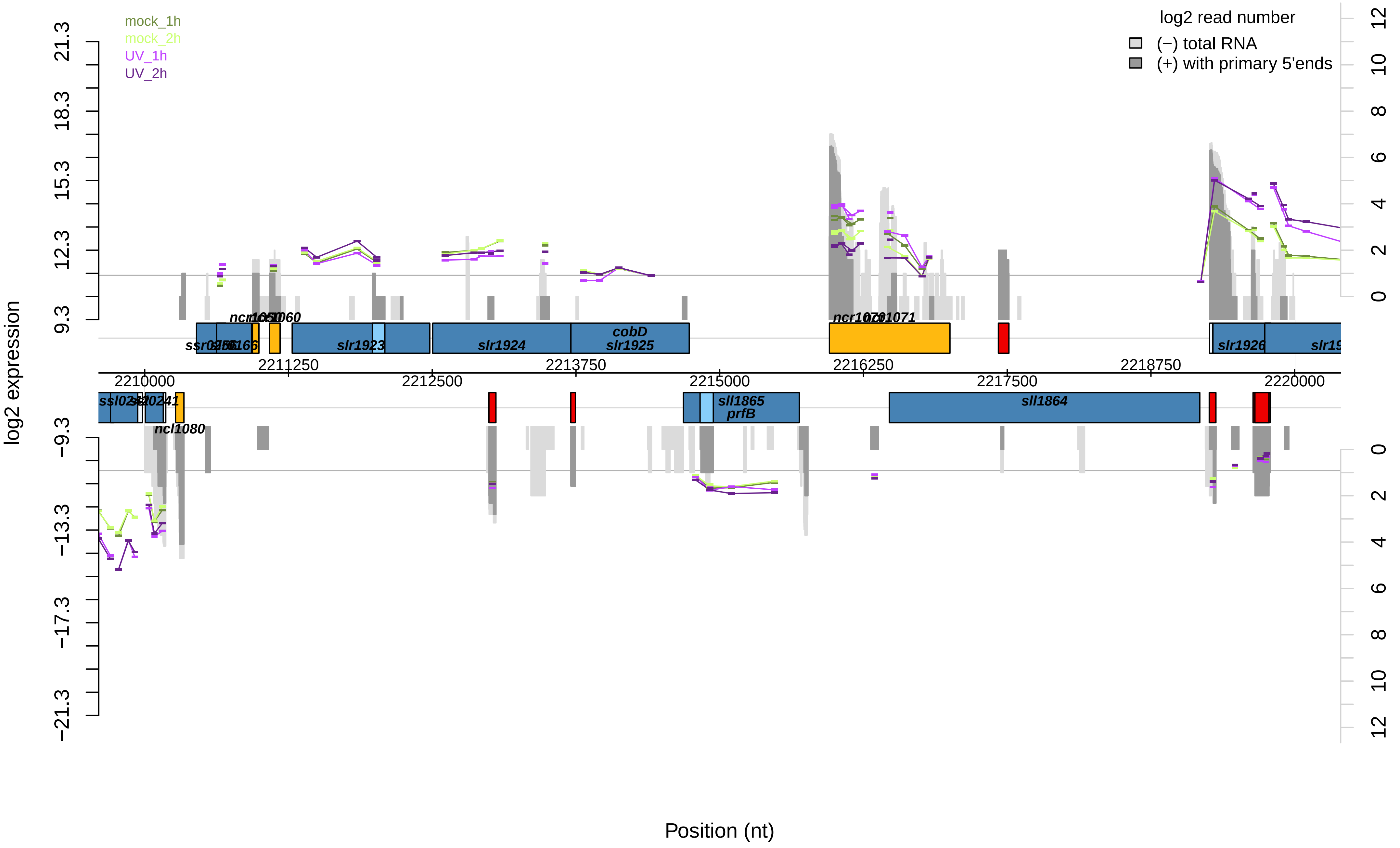

mock_1h
mock_2h
UV_1h
UV_2h
log2 read number
(−) total RNA
(+) with primary 5'ends
log2 expression
Position (nt)
slr1923
slr1924
cobD
slr1925
slr1926
ncr1070
ncl1080
sll1865
prfB
sll1864
2210000
2211250
2212500
2213750
2215000
2216250
2217500
2218750
2220000

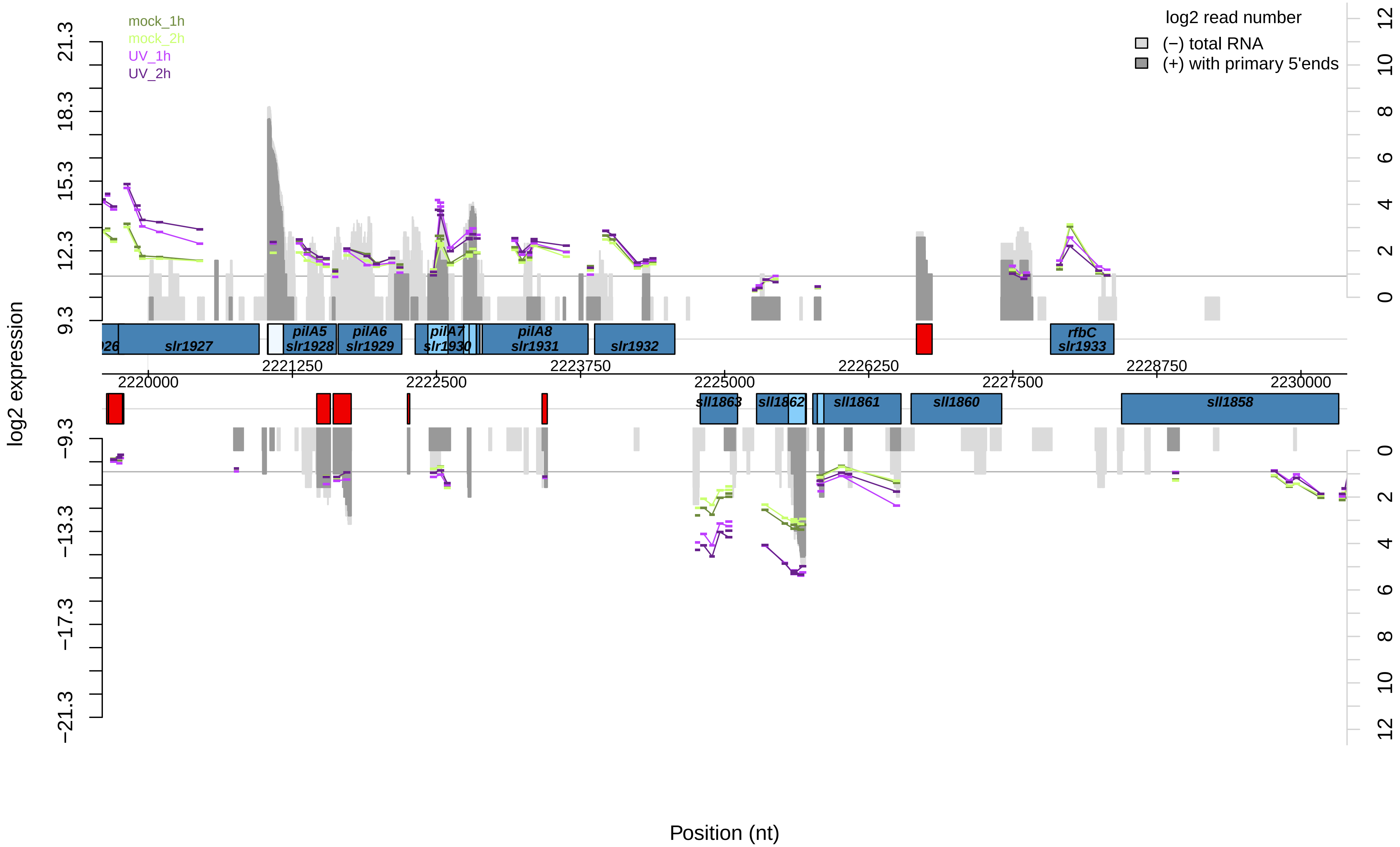

mock_1h
mock_2h
UV_1h
UV_2h
log2 read number
(−) total RNA
(+) with primary 5'ends
log2 expression
Position (nt)
21.3
18.3
15.3
12.3
9.3
−9.3
−13.3
−17.3
−21.3
12
10
8
6
4
2
0
slr1927
pilA5
slr1928
pilA6
slr1929
pilA7
slr1930
pilA8
slr1931
slr1932
rfbC
slr1933
2221250
2222500
2223750
2225000
2226250
2227500
2228750
2230000
2220000
sll1863
sll1862
sll1861
sll1860
sll1858

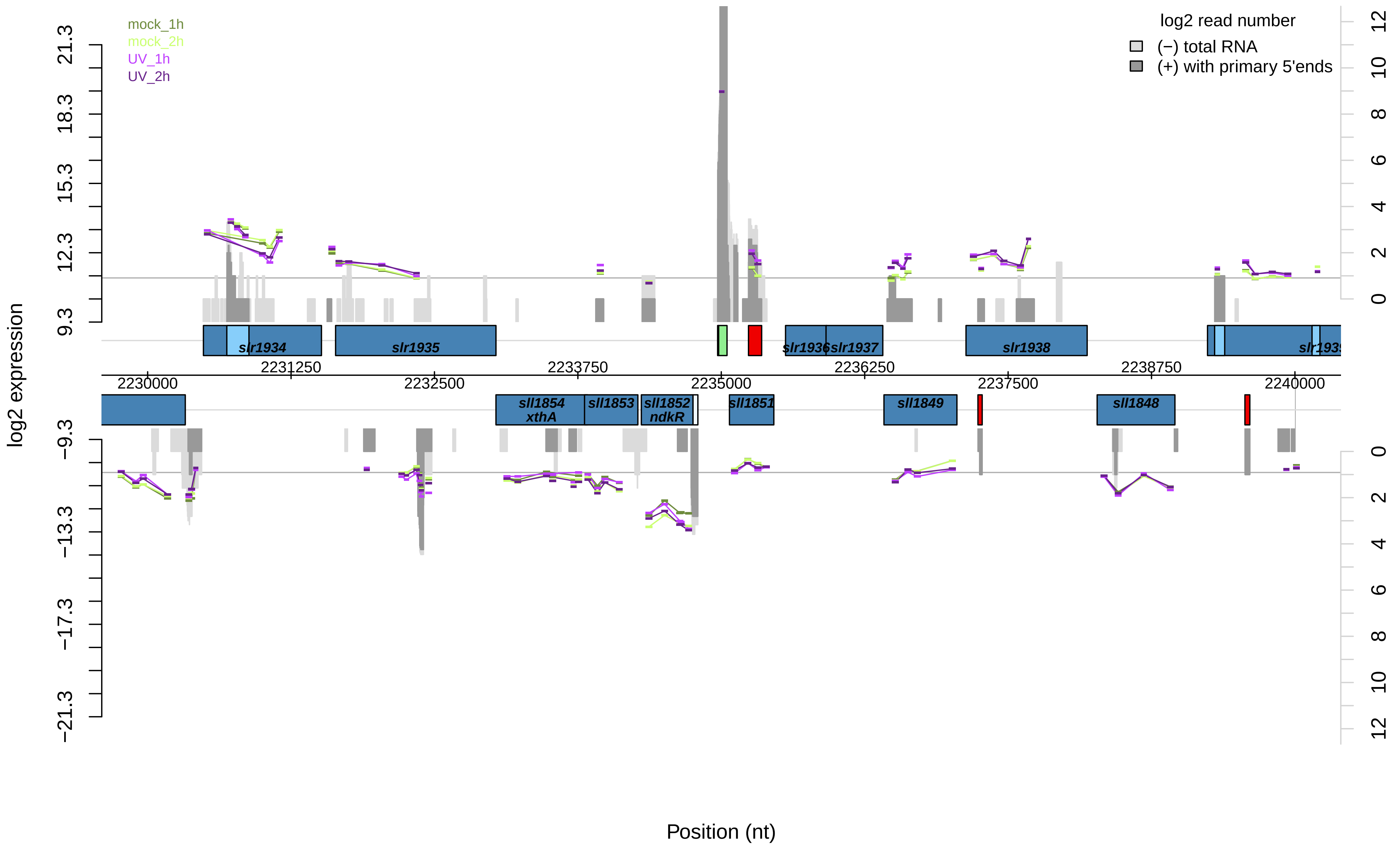
mock_1h
mock_2h
UV_1h
UV_2h
log2 read number
(−) total RNA
(+) with primary 5'ends
log2 expression
21.3
18.3
15.3
12.3
9.3
−9.3
−13.3
−17.3
−21.3
0
2
4
6
8
10
12
slr1934
slr1935
slr1936
slr1937
slr1938
sll1854
xthA
sll1853
sll1852
ndkR
sll1851
sll1849
sll1848
2230000
2231250
2232500
2233750
2235000
2236250
2237500
2238750
2240000
Position (nt)

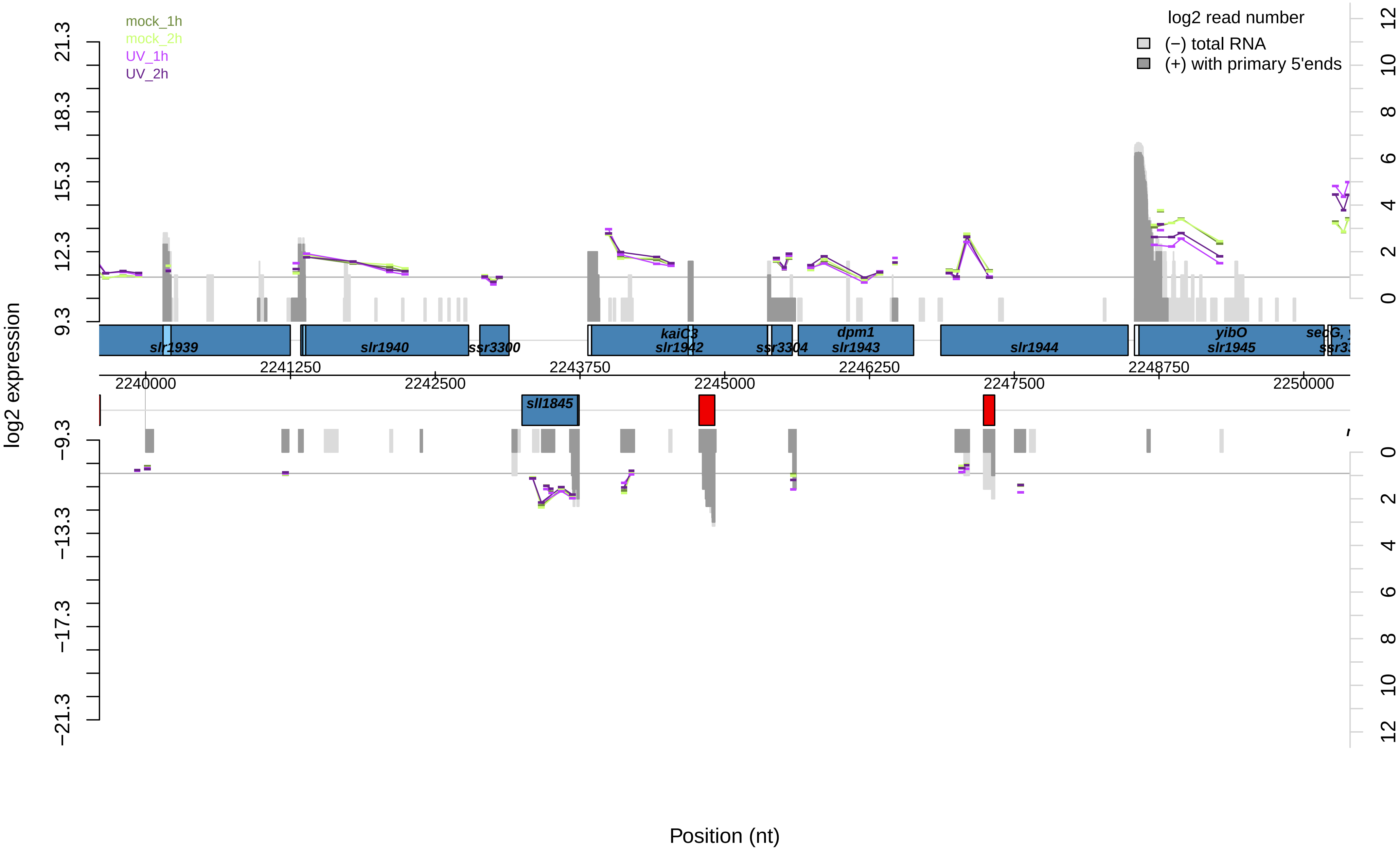

mock_1h
mock_2h
UV_1h
UV_2h
log2 read number
(−) total RNA
(+) with primary 5'ends
log2 expression
21.3
18.3
15.3
12.3
9.3
−9.3
−13.3
−17.3
−21.3
slr1939
slr1940
ssr3300
kaiC3
slr1942
ssr3304
dpm1
slr1943
slr1944
yibO
slr1945
sll1845
2241250
2243750
2246250
2248750
2240000
2242500
2245000
2247500
2250000
Position (nt)

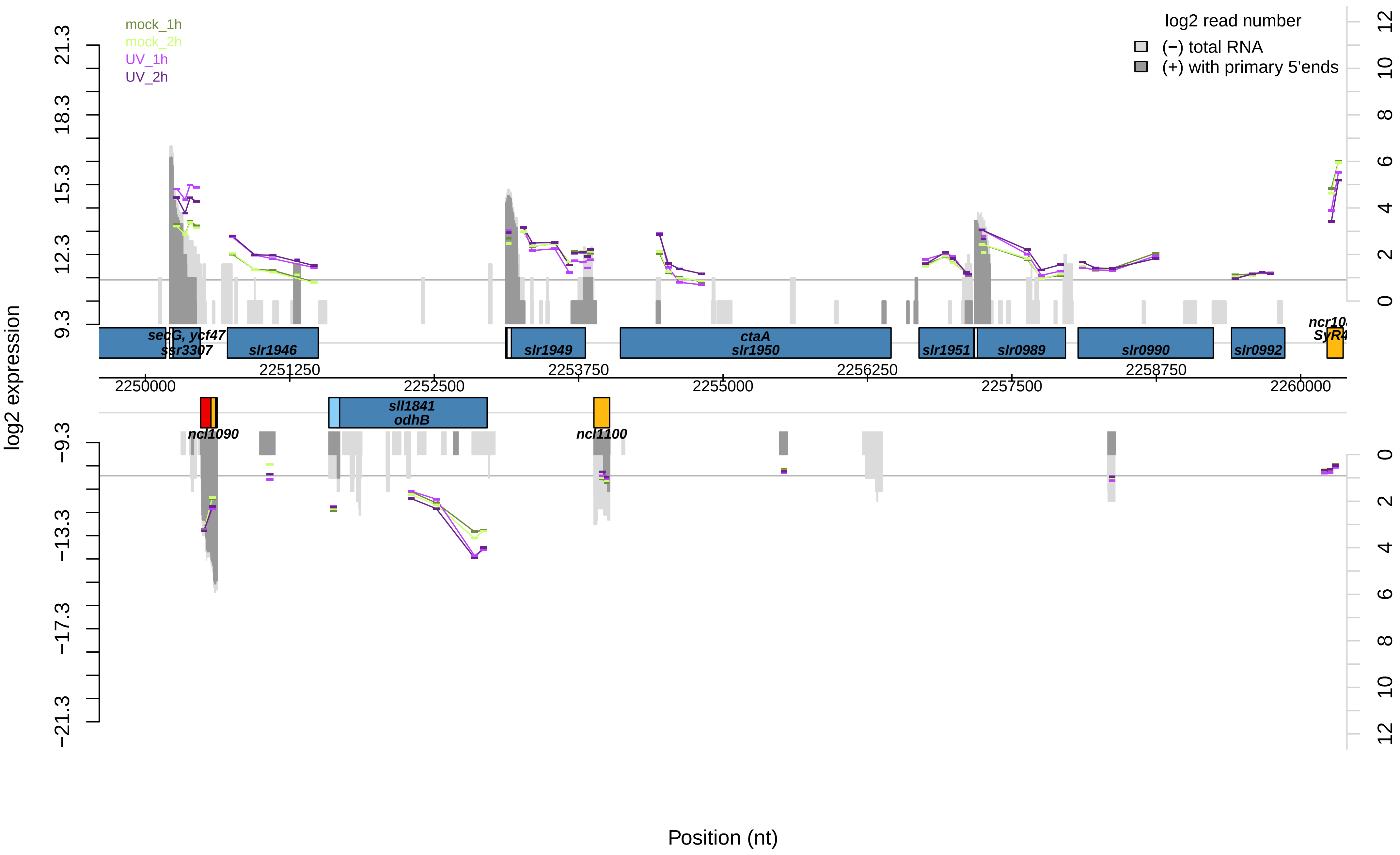

mock_1h
mock_2h
UV_1h
UV_2h
log2 read number
(−) total RNA
(+) with primary 5'ends
log2 expression
secG, ycf47
ssr3307
slr1946
slr1949
ctaA
slr1950
slr1951
slr0989
slr0990
slr0992
SyR4
ncl1090
sll1841
odhB
ncl1100
Position (nt)

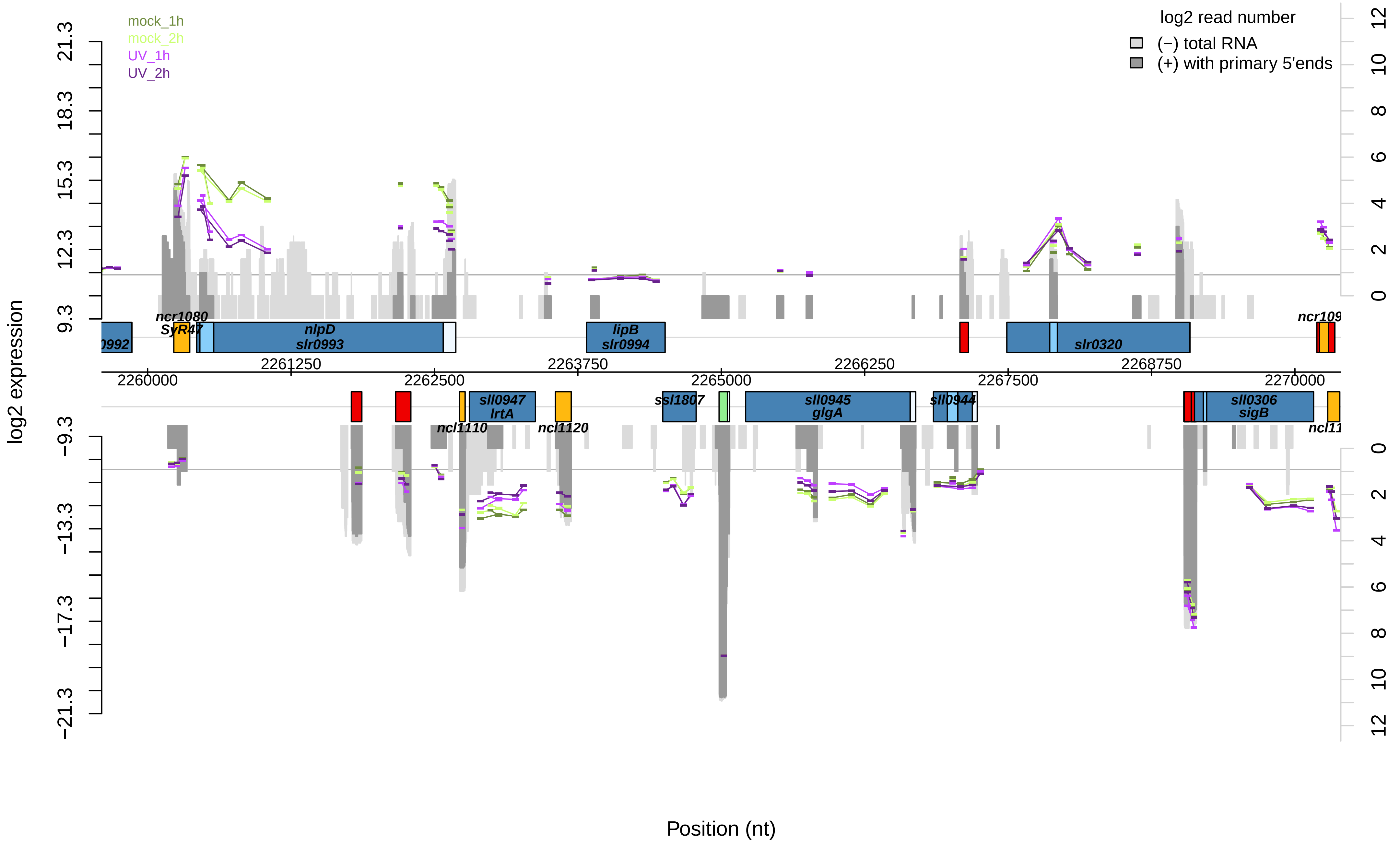

mock_1h
mock_2h
UV_1h
UV_2h
log2 read number
(−) total RNA
(+) with primary 5'ends
log2 expression
21.3
18.3
15.3
12.3
9.3
−9.3
−13.3
−17.3
−21.3
12
10
8
6
4
2
0
ncr1080
SyR47
nlpD
slr0993
lipB
slr0994
slr0320
ncr109
sll0947
lrtA
ncl1110
ncl1120
ssl1807
sll0945
glgA
sll0944
sll0306
sigB
2260000
2261250
2262500
2263750
2265000
2266250
2267500
2268750
2270000
Position (nt)

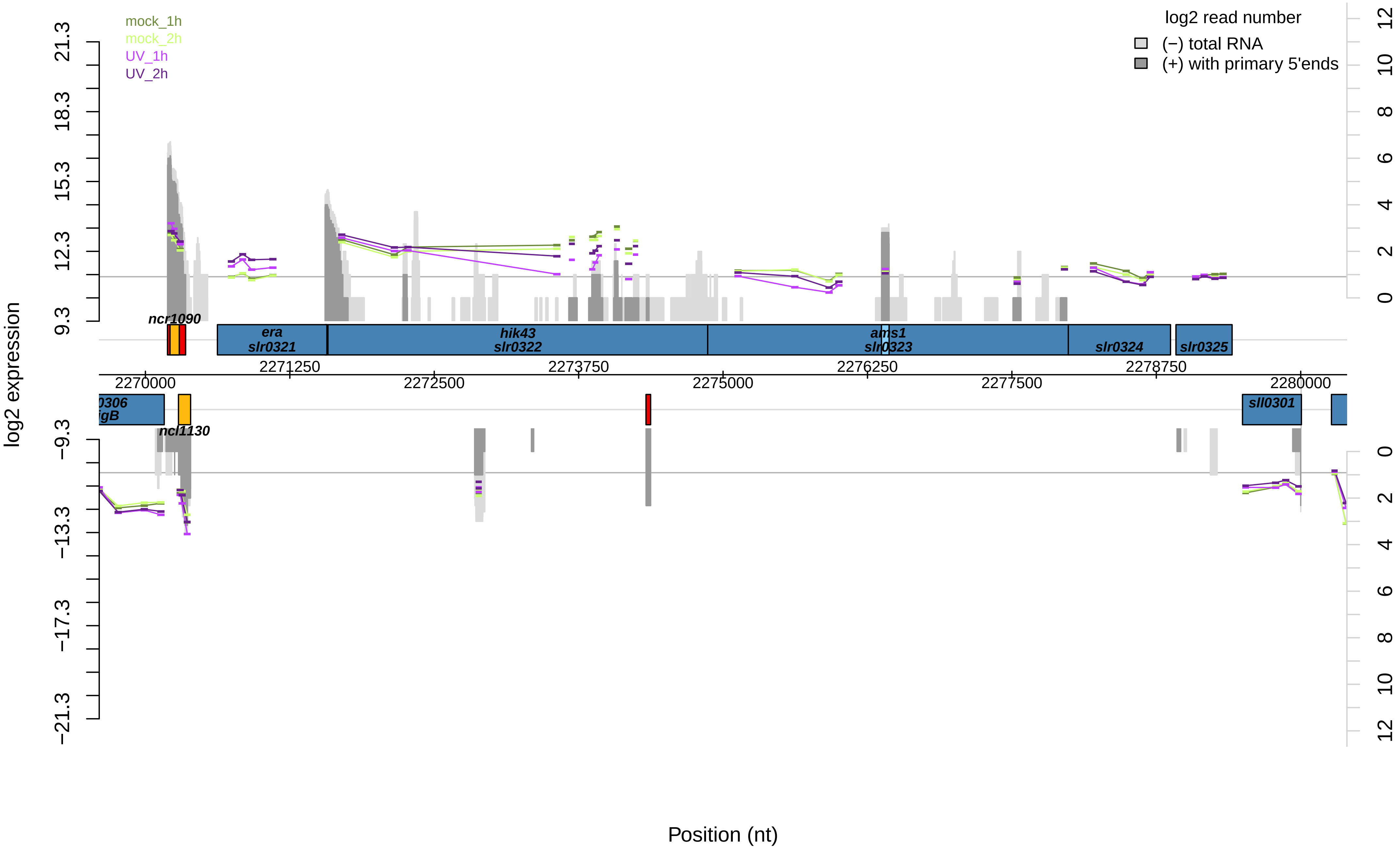
mock_1h
mock_2h
UV_1h
UV_2h
log2 read number
(−) total RNA
(+) with primary 5'ends
log2 expression
21.3
18.3
15.3
12.3
9.3
−9.3
−13.3
−17.3
−21.3
12
10
8
6
4
2
0
ncr1090
era
slr0321
hik43
slr0322
ams1
slr0323
slr0324
slr0325
2271250
2273750
2276250
2278750
2270000
2272500
2275000
2277500
2280000
0306
igB
ncl1130
sll0301
Position (nt)

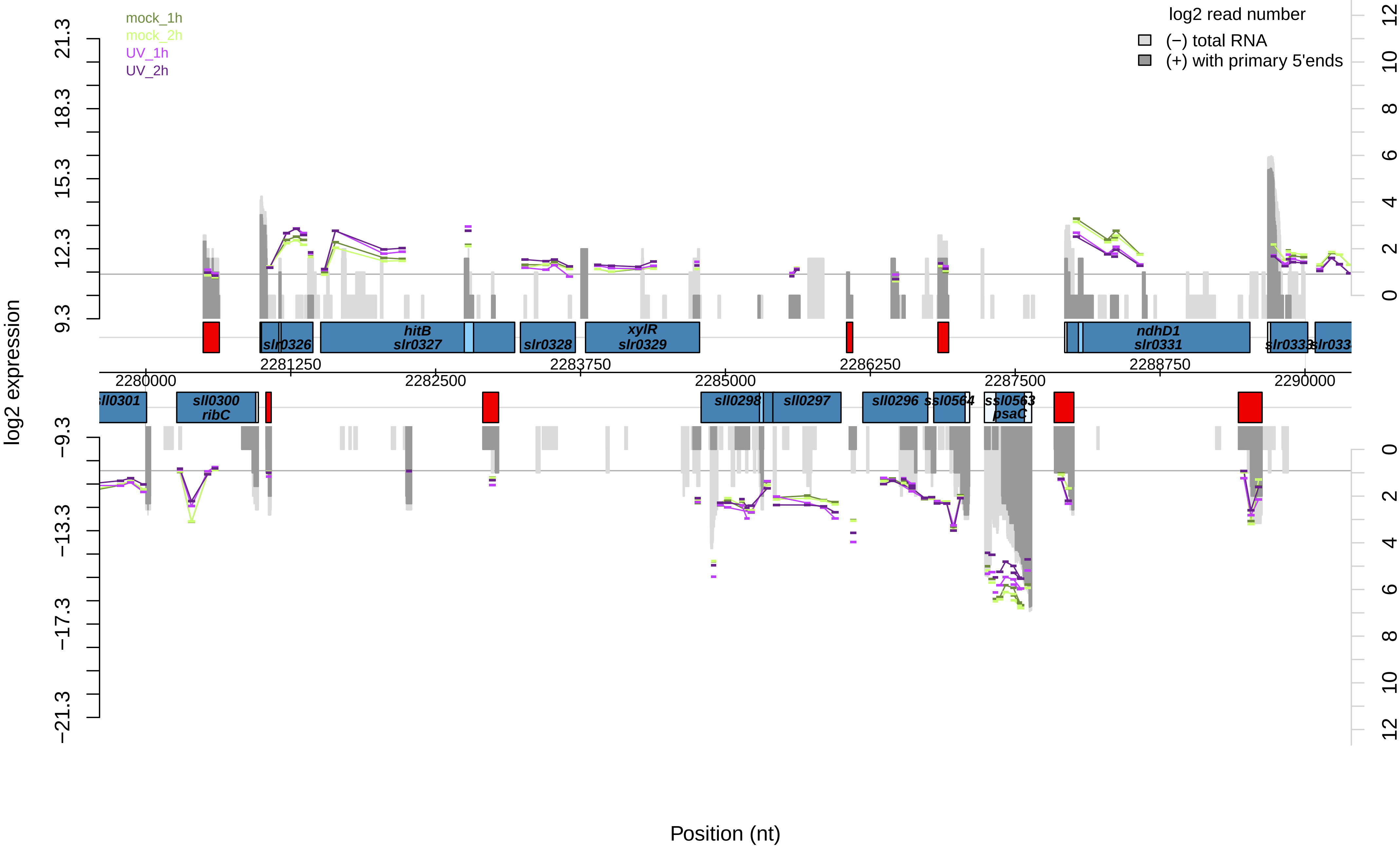

mock_1h
mock_2h
UV_1h
UV_2h
log2 read number
(−) total RNA
(+) with primary 5'ends
log2 expression
21.3
18.3
15.3
12.3
9.3
−9.3
−13.3
−17.3
−21.3
0
2
4
6
8
10
12
slr0326
hitB
slr0327
slr0328
xylR
slr0329
ndhD1
slr0331
slr0333
2281250
2283750
2286250
2288750
2280000
2282500
2285000
2287500
2290000
sll0300
ribC
sll0298
sll0297
sll0296
ssl0564
ssl0563
psaC
Position (nt)

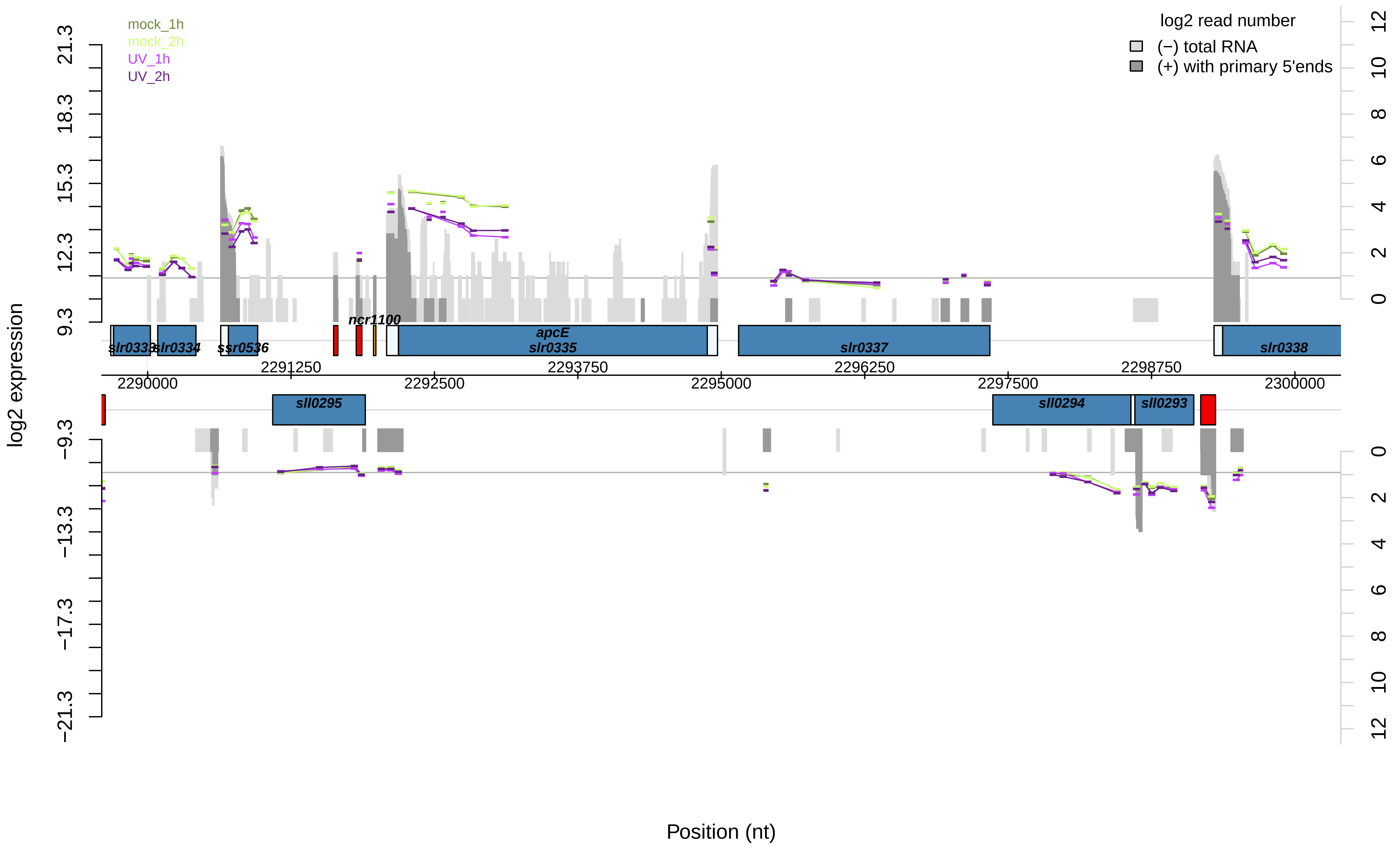
mock_1h
mock_2h
UV_1h
UV_2h
log2 read number
(−) total RNA
(+) with primary 5'ends
log2 expression
21.3
18.3
15.3
12.3
9.3
−9.3
−13.3
−17.3
−21.3
slr0333
slr0334
ssr0536
ncr1100
apcE
slr0335
slr0337
slr0338
sll0295
sll0294
sll0293
2290000
2291250
2292500
2293750
2295000
2296250
2297500
2298750
2300000
Position (nt)

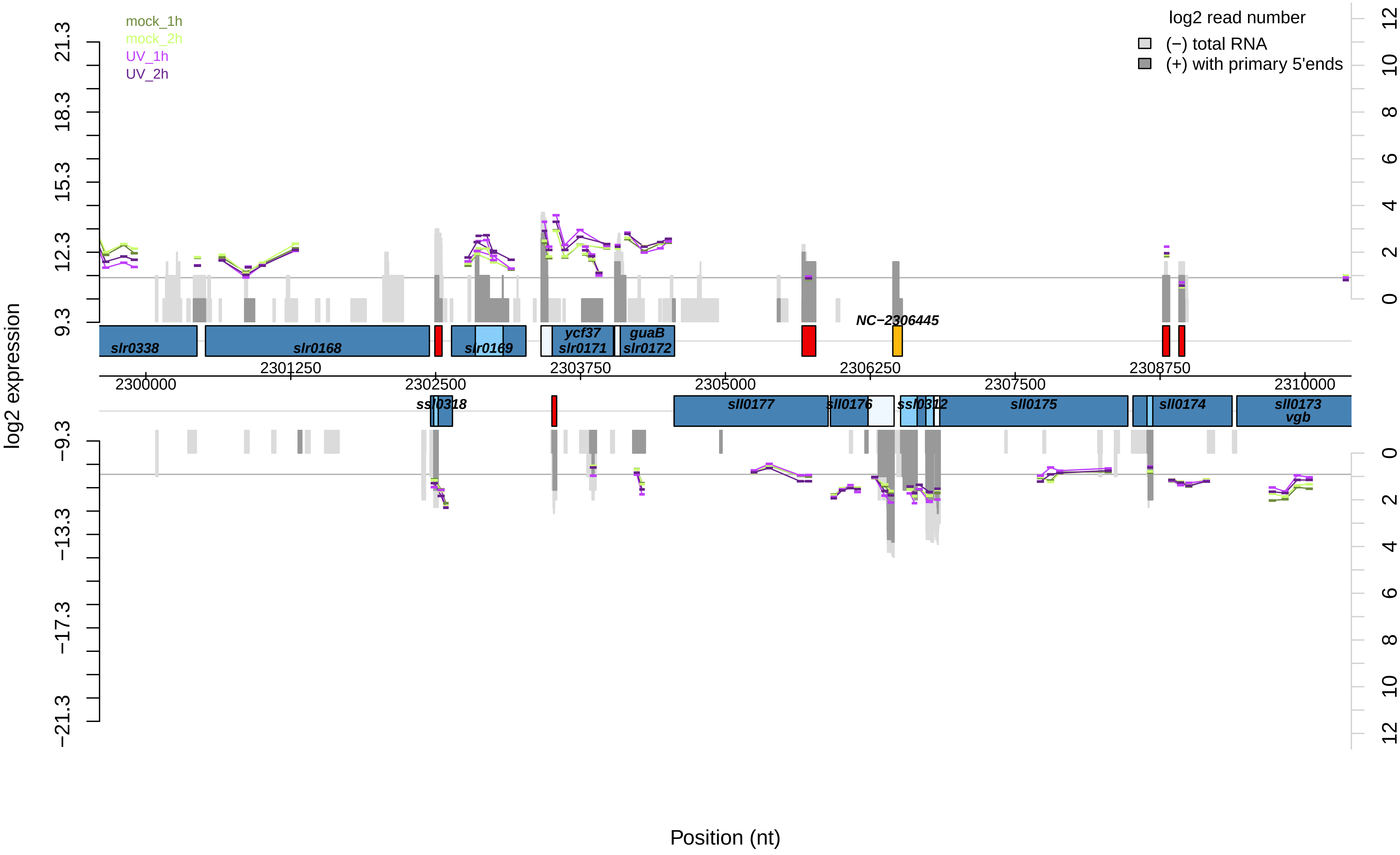

mock_1h
mock_2h
UV_1h
UV_2h
log2 read number
(−) total RNA
(+) with primary 5'ends
log2 expression
21.3
18.3
15.3
12.3
9.3
−9.3
−13.3
−17.3
−21.3
12
10
8
6
4
2
0
slr0338
slr0168
slr0169
ycf37
slr0171
guaB
slr0172
NC−2306445
2301250
2303750
2306250
2308750
2300000
2302500
2305000
2307500
2310000
ssl0318
sll0177
sll0176
ssl0312
sll0175
sll0174
sll0173
vgb
Position (nt)

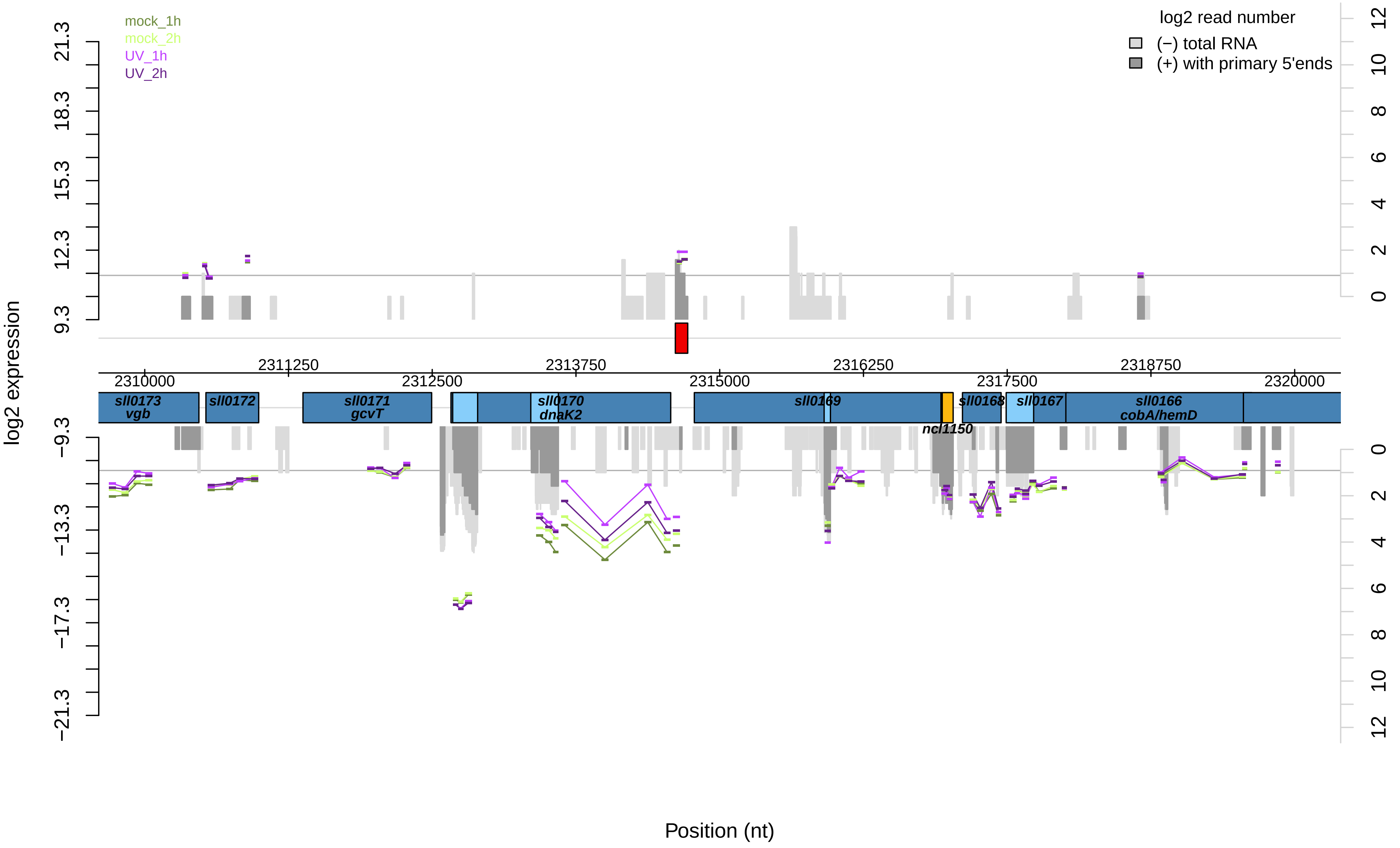
mock_1h
mock_2h
UV_1h
UV_2h
log2 read number
(−) total RNA
(+) with primary 5'ends
log2 expression
21.3
18.3
15.3
12.3
9.3
−9.3
−13.3
−17.3
−21.3
12
10
8
6
4
2
0
2311250
2313750
2316250
2318750
2310000
2312500
2315000
2317500
2320000
sll0173
vgb
sll0172
sll0171
gcvT
sll0170
dnaK2
sll0169
ncl1150
sll0168
sll0167
sll0166
cobA/hemD
Position (nt)

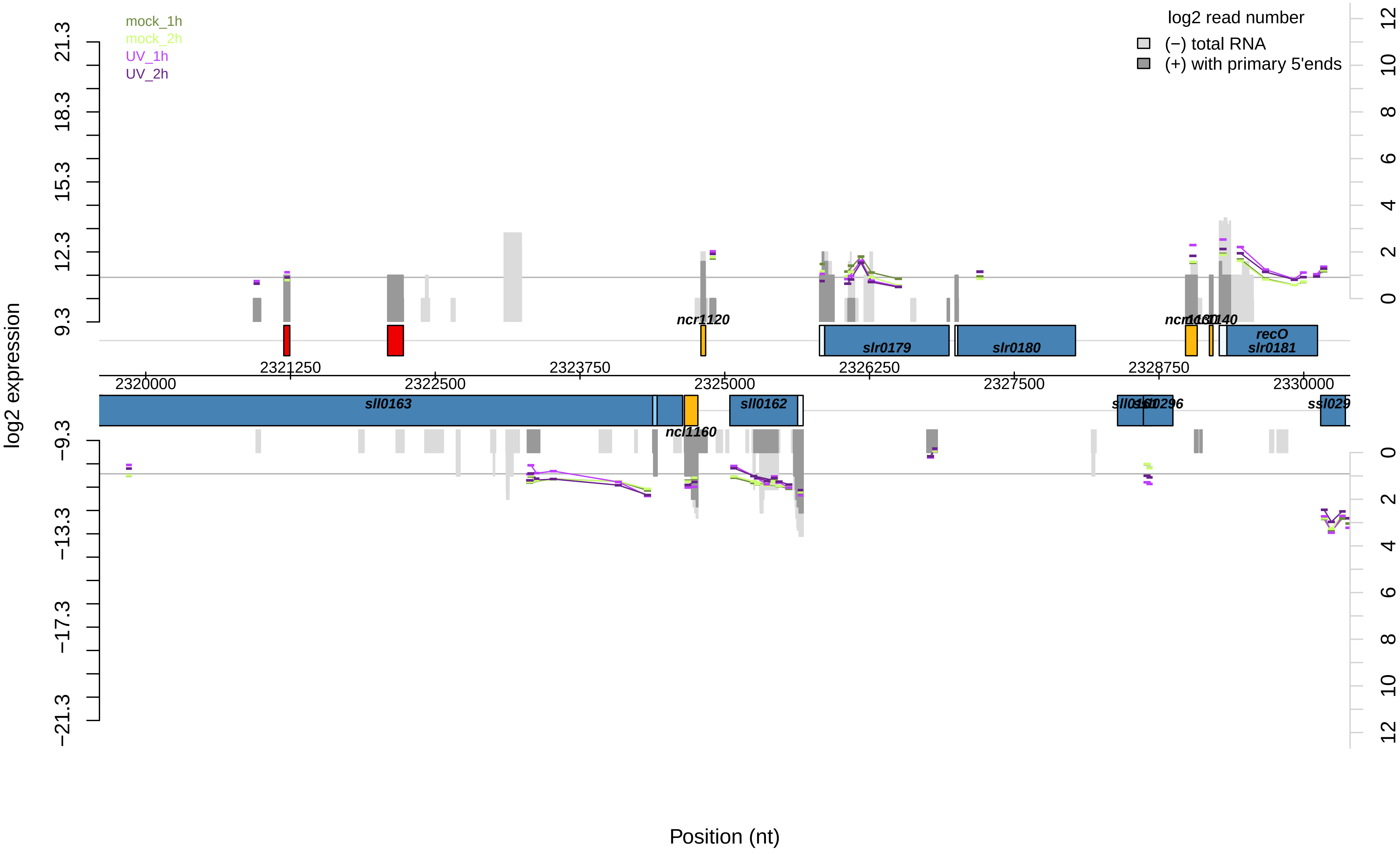

mock_1h
mock_2h
UV_1h
UV_2h
log2 read number
(−) total RNA
(+) with primary 5'ends
log2 expression
21.3
18.3
15.3
12.3
9.3
−9.3
−13.3
−17.3
−21.3
ncr1120
ncr1130
slr0179
slr0180
recO
slr0181
2321250
2323750
2326250
2328750
2320000
2322500
2325000
2327500
2330000
sll0163
ncl1160
sll0162
ssl029
Position (nt)

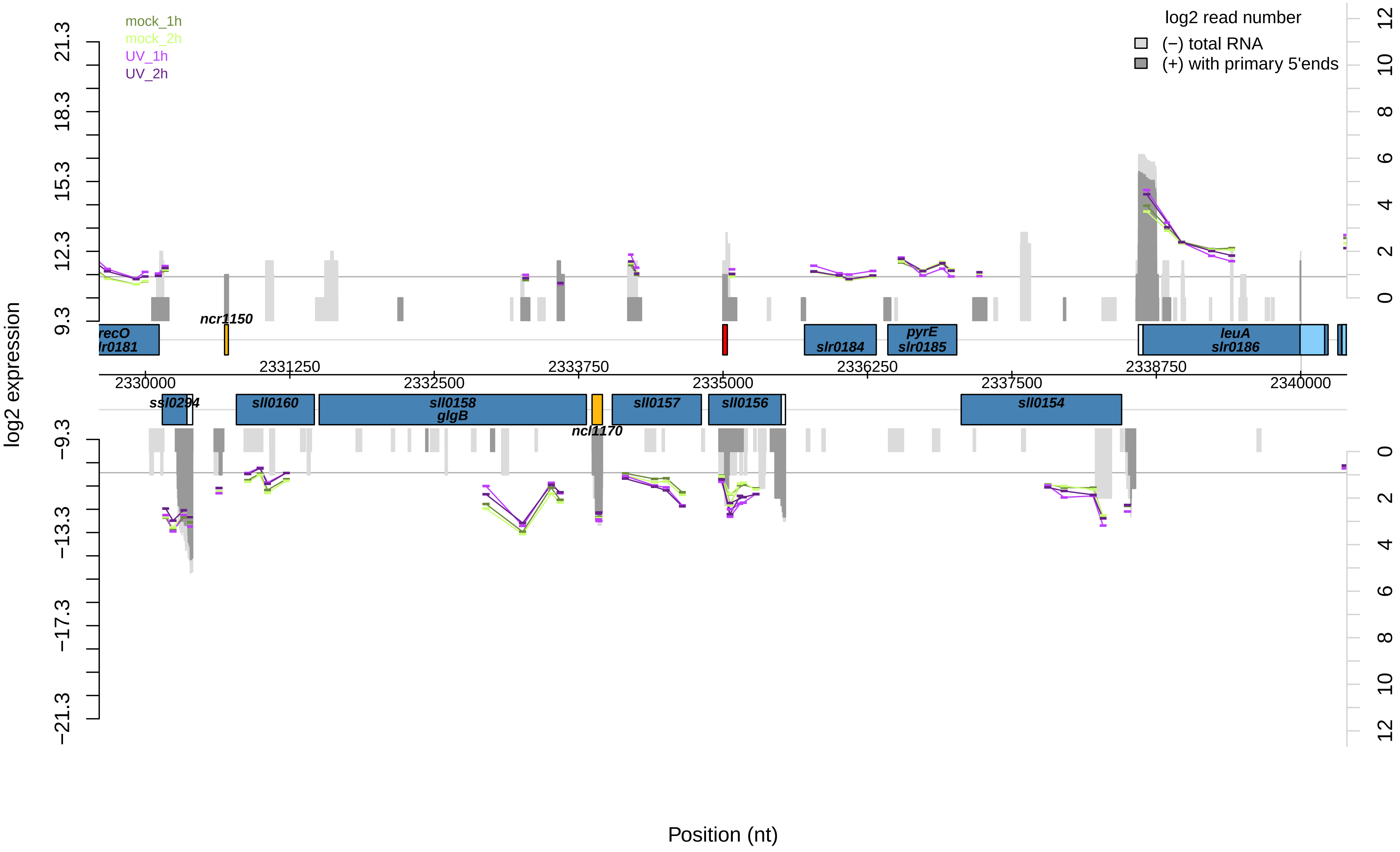

mock_1h
mock_2h
UV_1h
UV_2h
log2 read number
(−) total RNA
(+) with primary 5'ends
21.3
18.3
15.3
12.3
9.3
−9.3
−13.3
−17.3
−21.3
log2 expression
0
2
4
6
8
10
12
recO
lr0181
ncr1150
slr0184
pyrE
slr0185
leuA
slr0186
2331250
2333750
2336250
2338750
2330000
2332500
2335000
2337500
2340000
ssl0294
sll0160
sll0158
glgB
ncl1170
sll0157
sll0156
sll0154
Position (nt)

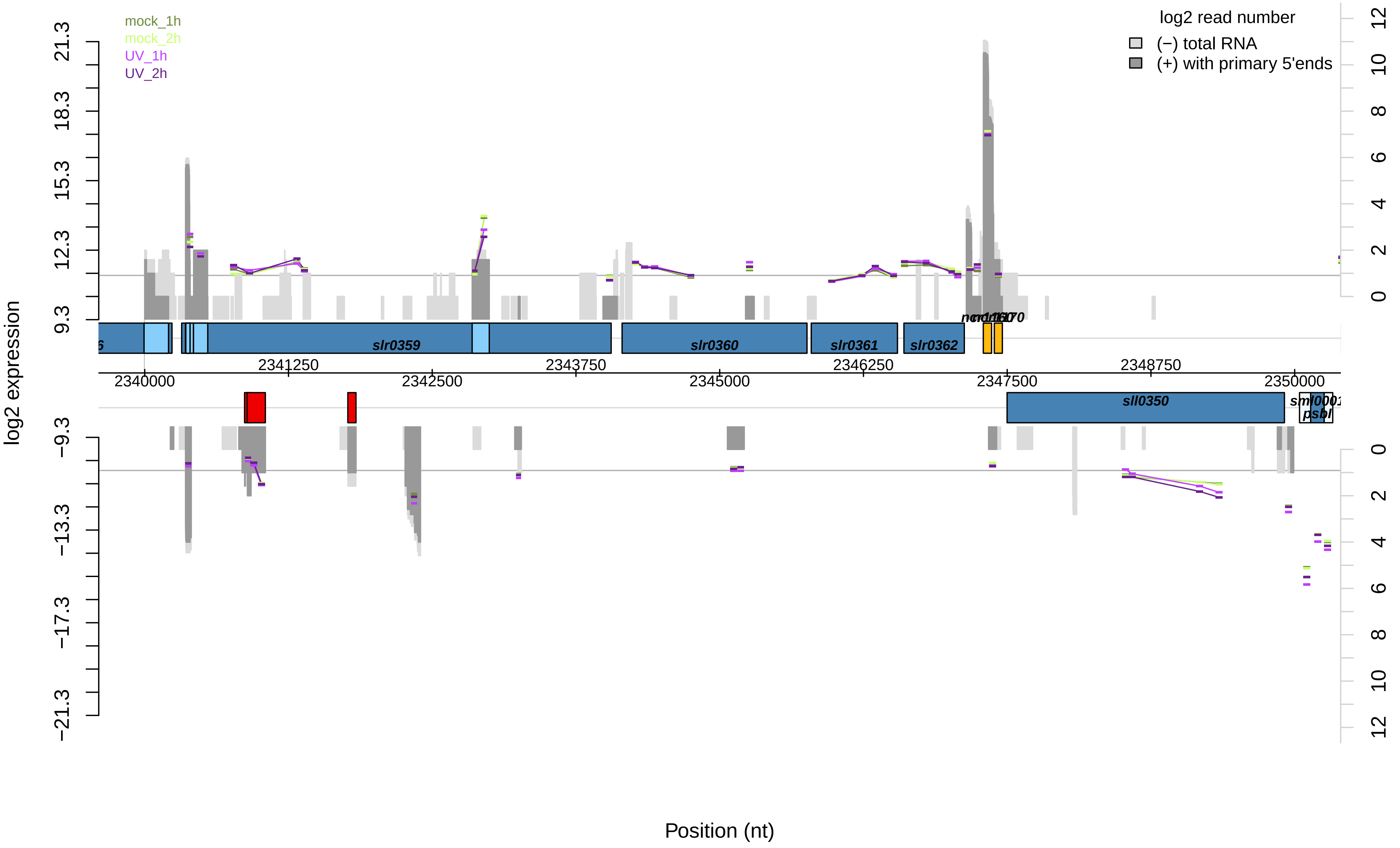

mock_1h
mock_2h
UV_1h
UV_2h
log2 read number
(−) total RNA
(+) with primary 5'ends
log2 expression
21.3
18.3
15.3
12.3
9.3
−9.3
−13.3
−17.3
−21.3
12
10
8
6
4
2
0
slr0359
slr0360
slr0361
slr0362
sll0350
psbI
2341250
2343750
2346250
2348750
2340000
2342500
2345000
2347500
2350000
Position (nt)

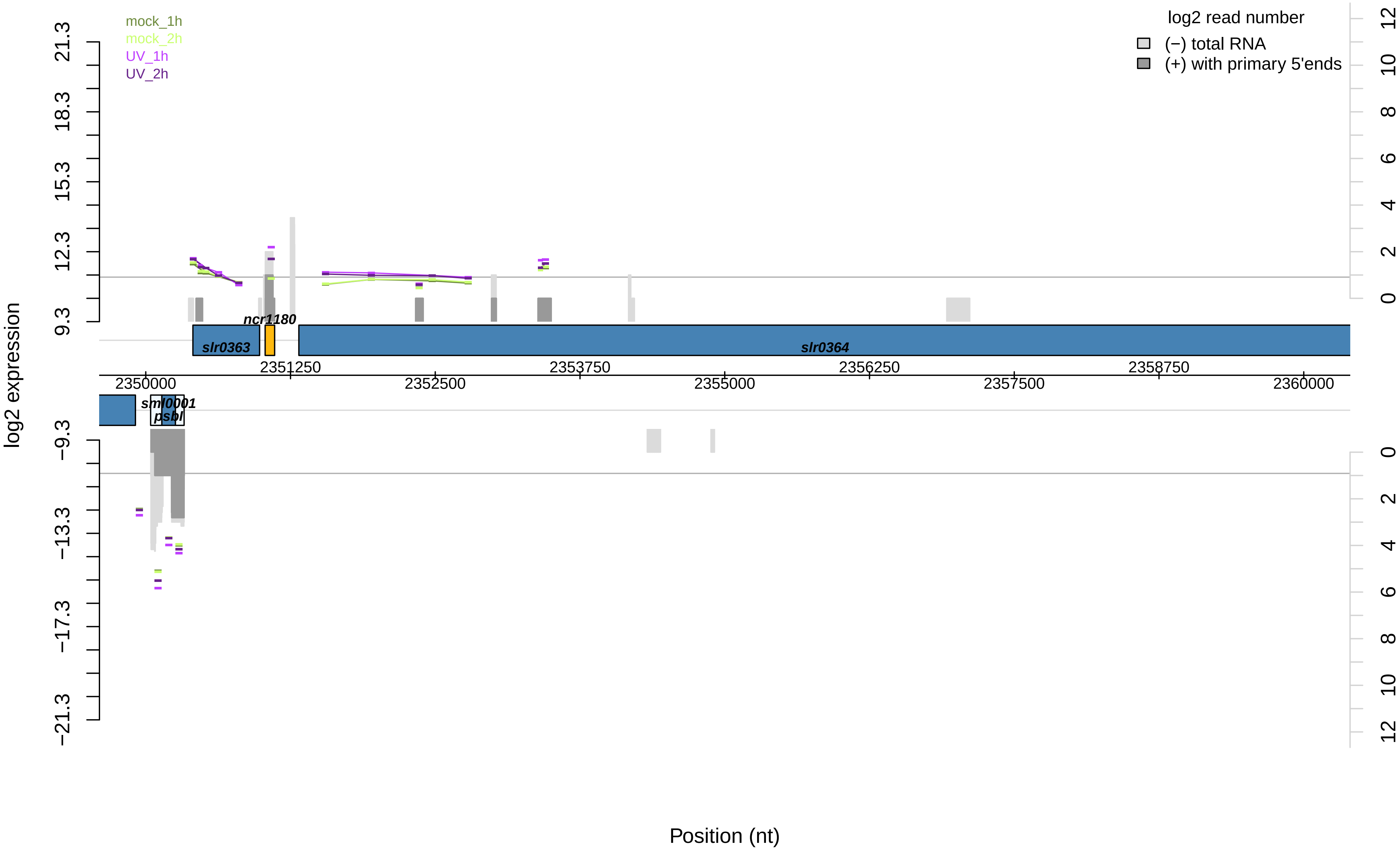

mock_1h
mock_2h
UV_1h
UV_2h
log2 read number
(−) total RNA
(+) with primary 5'ends
log2 expression
21.3
18.3
15.3
12.3
9.3
−9.3
−13.3
−17.3
−21.3
12
10
8
6
4
2
0
ncr1180
slr0363
slr0364
sml0001
psbI
2350000
2351250
2352500
2353750
2355000
2356250
2357500
2358750
2360000
Position (nt)

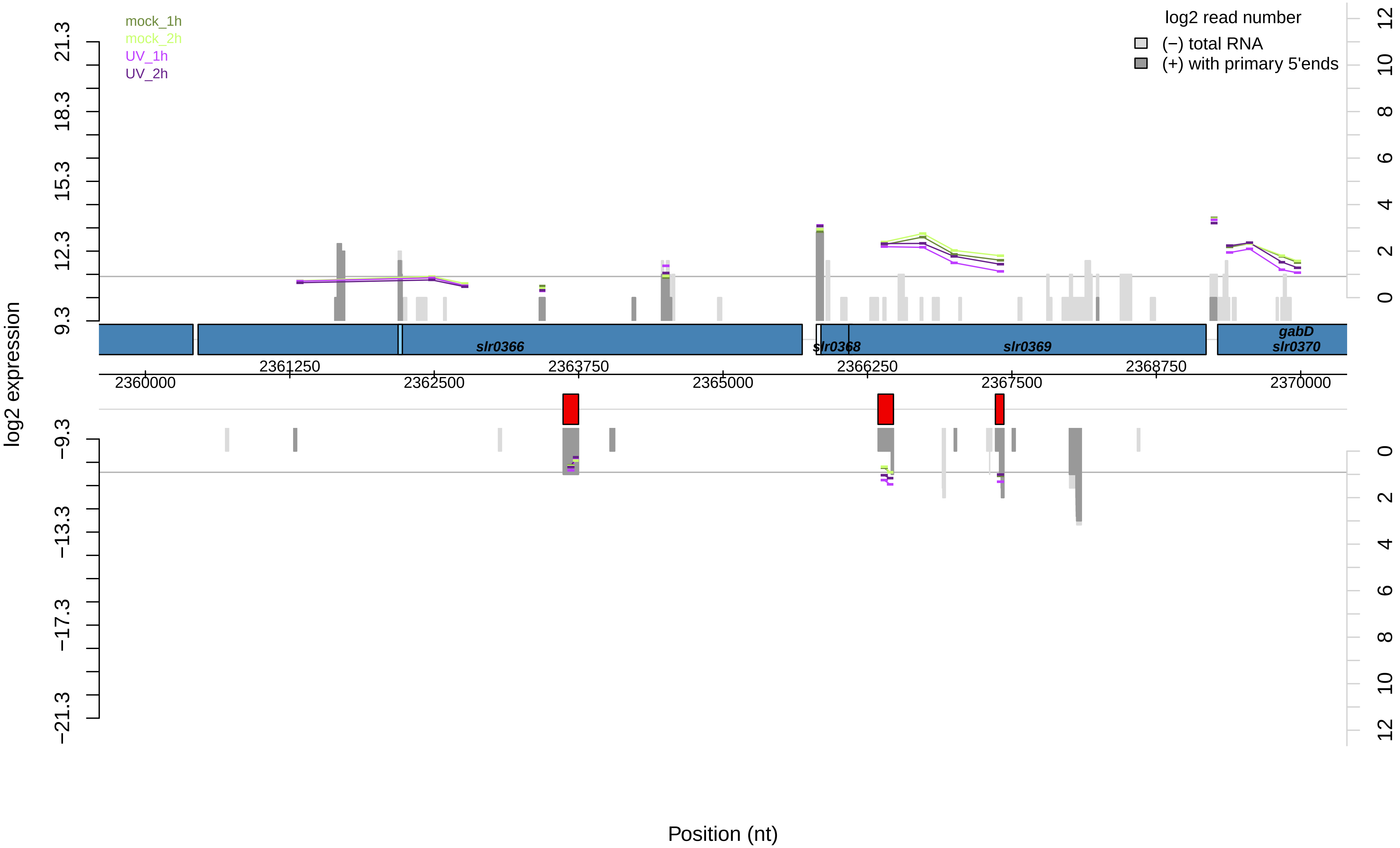

mock_1h
mock_2h
UV_1h
UV_2h
log2 read number
(−) total RNA
(+) with primary 5'ends
log2 expression
21.3
18.3
15.3
12.3
9.3
−9.3
−13.3
−17.3
−21.3
12
10
8
6
4
2
0
slr0366
slr0368
slr0369
gabD
slr0370
2361250
2363750
2366250
2368750
2360000
2362500
2365000
2367500
2370000
Position (nt)

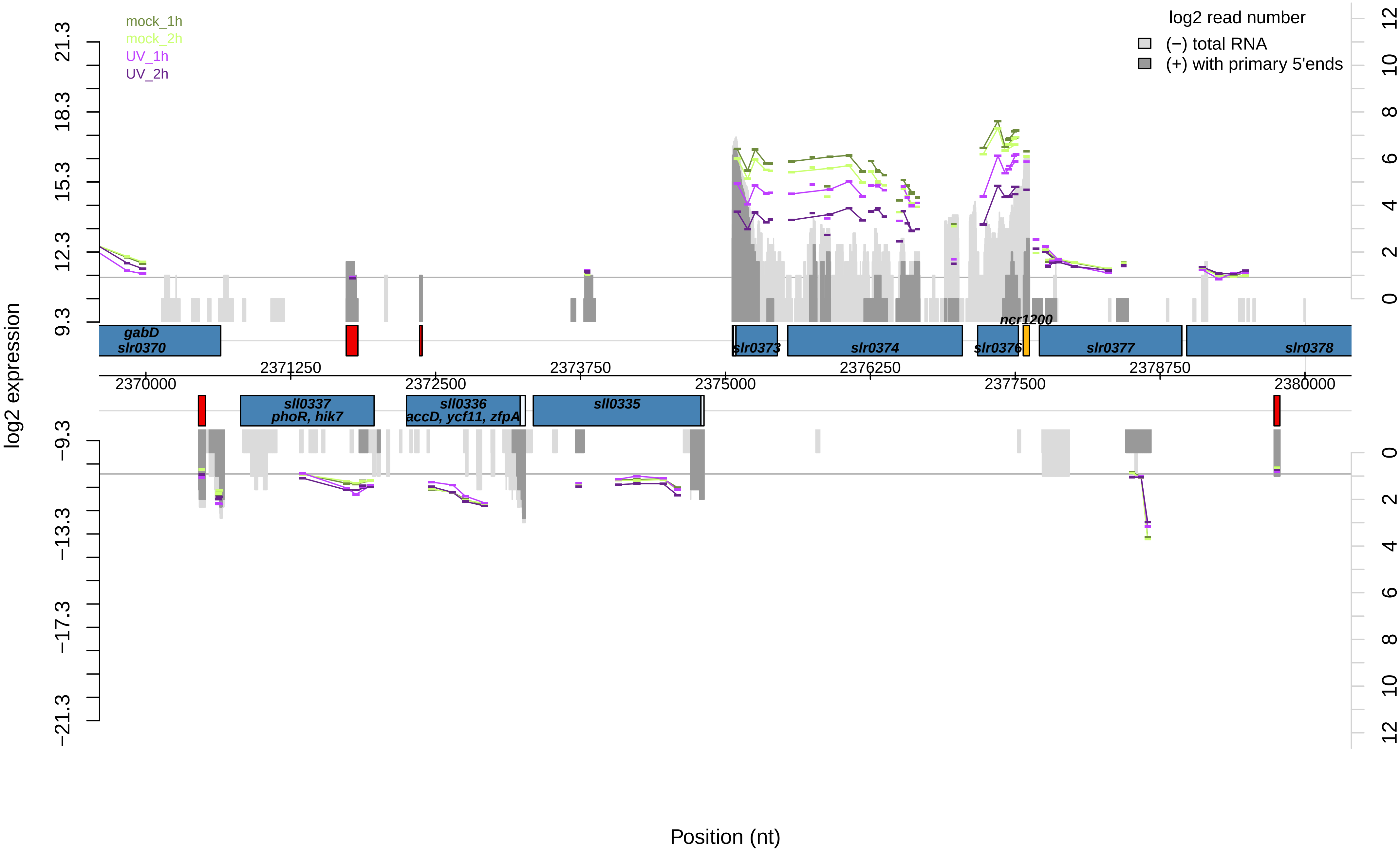

mock_1h
mock_2h
UV_1h
UV_2h
log2 read number
(−) total RNA
(+) with primary 5'ends
log2 expression
21.3
18.3
15.3
12.3
9.3
−9.3
−13.3
−17.3
−21.3
12
10
8
6
4
2
0
gabD
slr0370
slr0373
slr0374
slr0376
ncr1200
slr0377
slr0378
2371250
2373750
2376250
2378750
2370000
2372500
2375000
2377500
2380000
sll0337
phoR, hik7
sll0336
accD, ycf11, zfpA
sll0335
Position (nt)

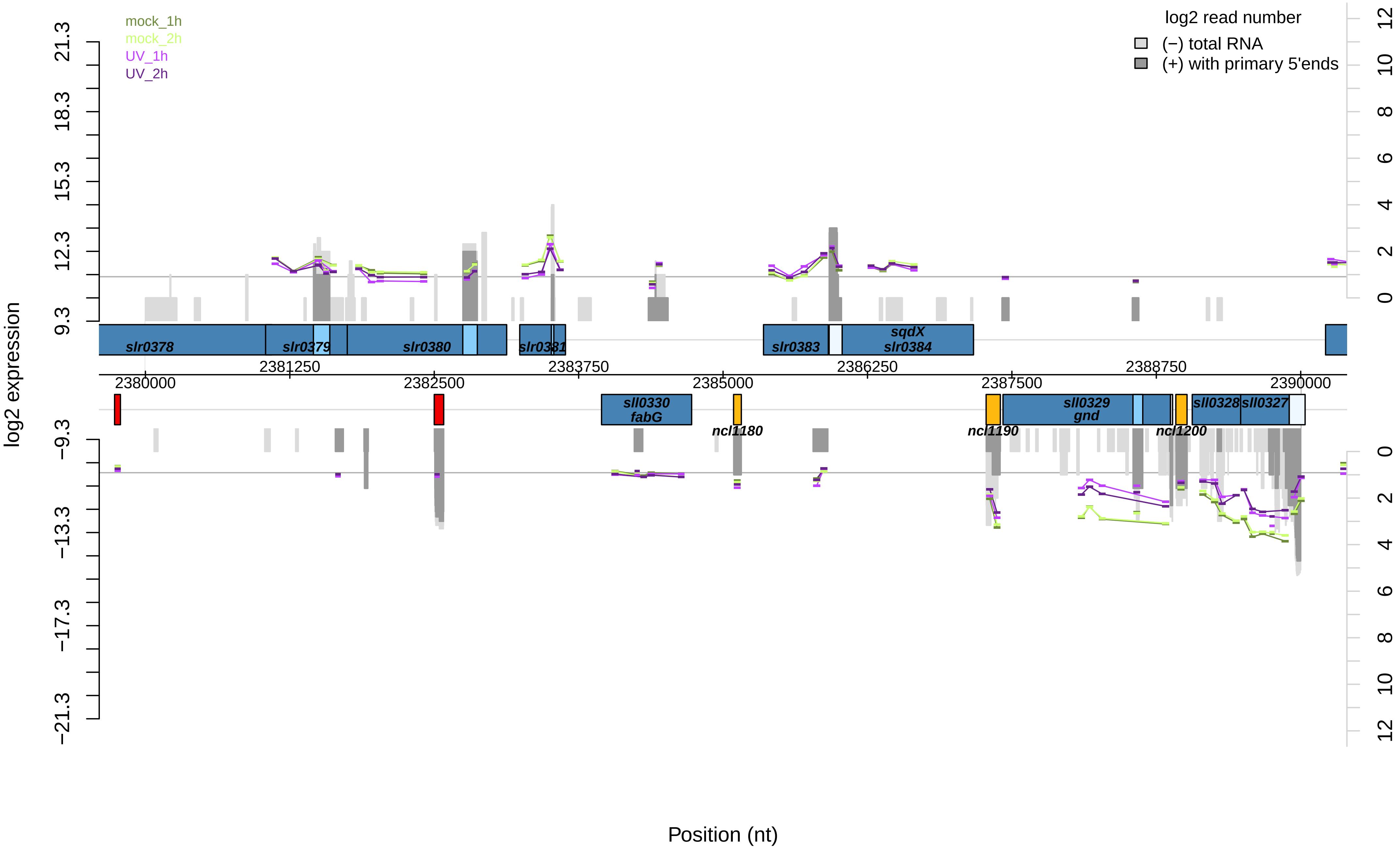

mock_1h
mock_2h
UV_1h
UV_2h
log2 read number
(−) total RNA
(+) with primary 5'ends
log2 expression
21.3
18.3
15.3
12.3
9.3
−9.3
−13.3
−17.3
−21.3
12
10
8
6
4
2
0
slr0378
slr0379
slr0380
slr0381
slr0383
sqdX
slr0384
2381250
2383750
2386250
2388750
2380000
2382500
2385000
2387500
2390000
sll0330
fabG
ncl1180
ncl1190
sll0329
gnd
ncl1200
sll0328
sll0327
Position (nt)

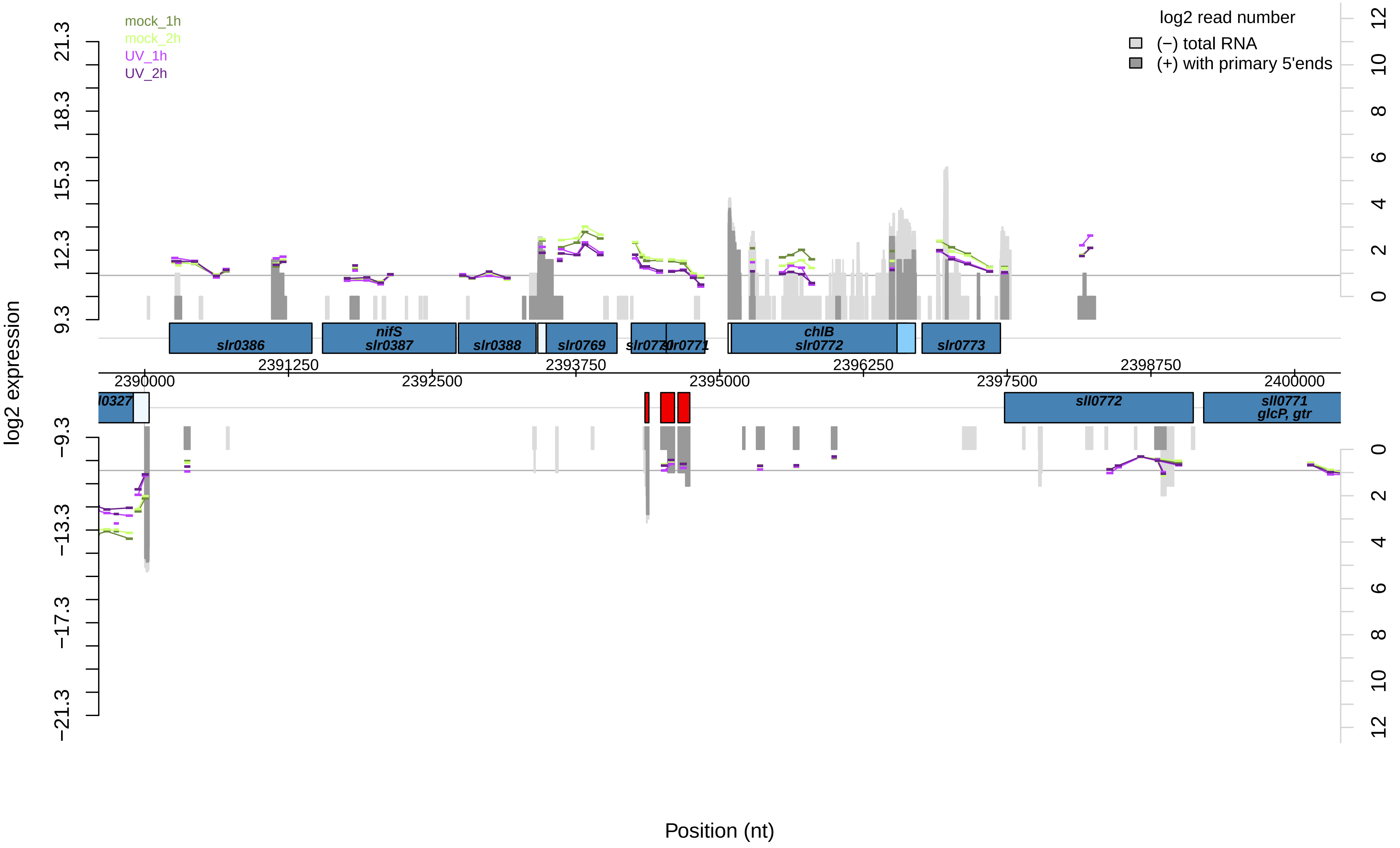

mock_1h
mock_2h
UV_1h
UV_2h
log2 read number
(−) total RNA
(+) with primary 5'ends
log2 expression
21.3
18.3
15.3
12.3
9.3
−9.3
−13.3
−17.3
−21.3
slr0386
nifS
slr0387
slr0388
slr0769
slr0770
slr0771
chlB
slr0772
slr0773
2391250
2393750
2396250
2398750
2390000
2392500
2395000
2397500
2400000
ll0327
sll0772
sll0771
glcP, gtr
Position (nt)

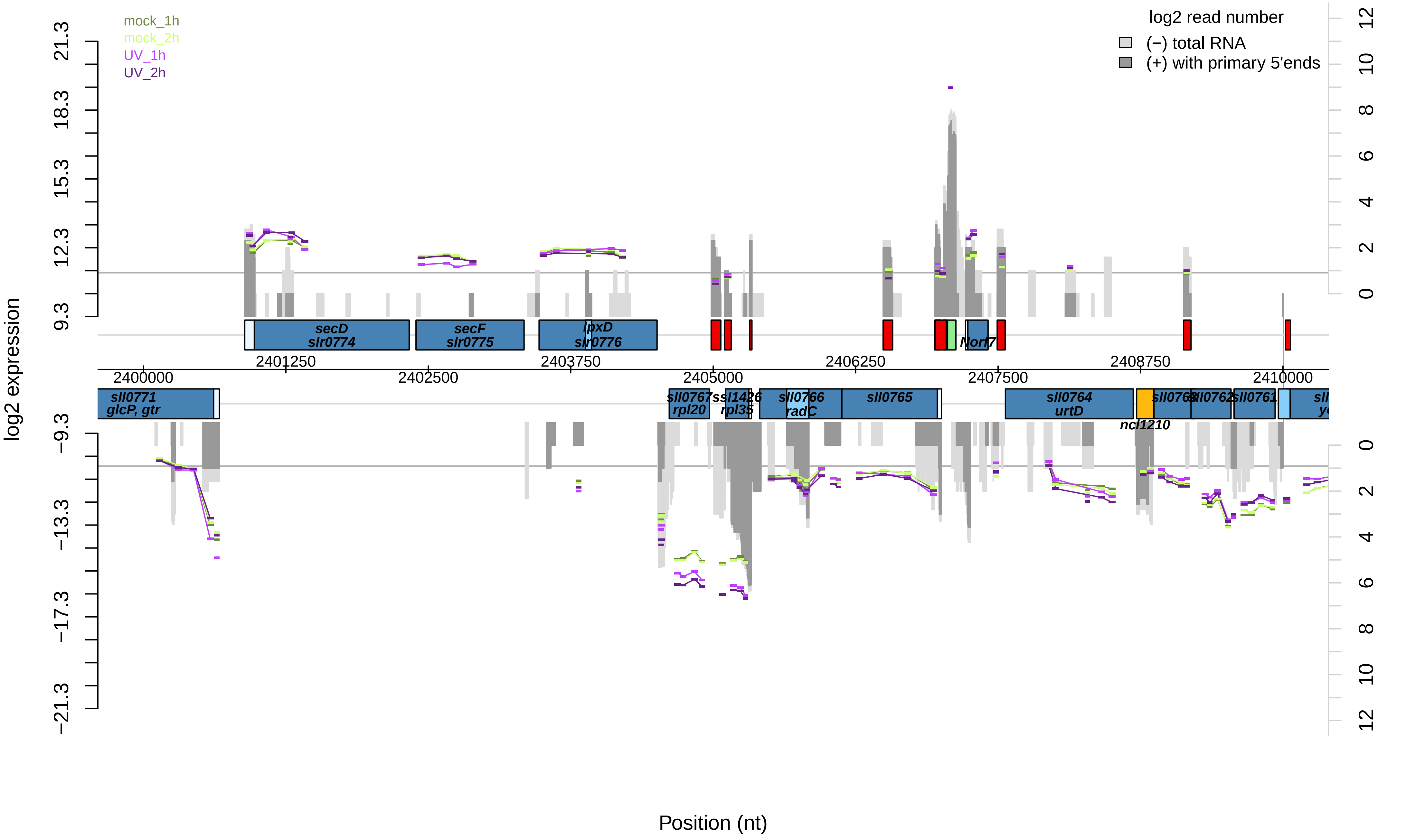

mock_1h
mock_2h
UV_1h
UV_2h
log2 read number
(−) total RNA
(+) with primary 5'ends
log2 expression
21.3
18.3
15.3
12.3
9.3
−9.3
−13.3
−17.3
−21.3
12
10
8
6
4
2
0
secD
slr0774
secF
slr0775
lpxD
slr0776
Norf7
2401250
2403750
2406250
2408750
2400000
2402500
2405000
2407500
2410000
sll0771
glcP, gtr
sll0767
rpl20
ssl1426
rpl35
sll0766
radC
sll0765
sll0764
urtD
ncl1210
sll0763
sll0762
sll0761
Position (nt)

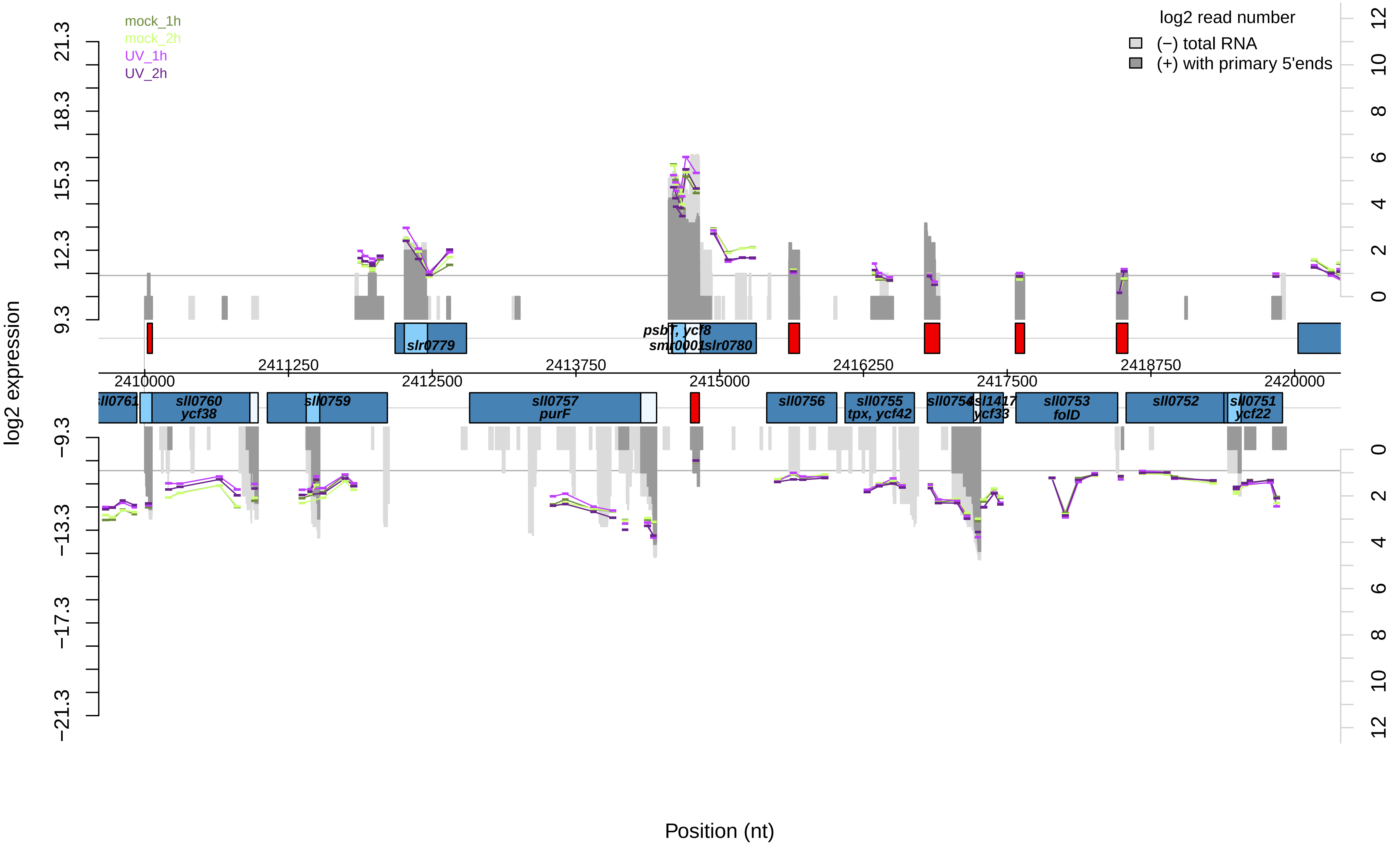

mock_1h
mock_2h
UV_1h
UV_2h
log2 read number
(−) total RNA
(+) with primary 5'ends
log2 expression
21.3
18.3
15.3
12.3
9.3
−9.3
−13.3
−17.3
−21.3
12
10
8
6
4
2
0
slr0779
psbT, ycf8
sml0001
slr0780
2411250
2413750
2416250
2418750
2410000
2412500
2415000
2417500
2420000
sll0761
sll0760
ycf38
sll0759
sll0757
purF
sll0756
sll0755
tpx, ycf42
sll0754
ssl1417
ycf33
sll0753
folD
sll0752
sll0751
ycf22
Position (nt)

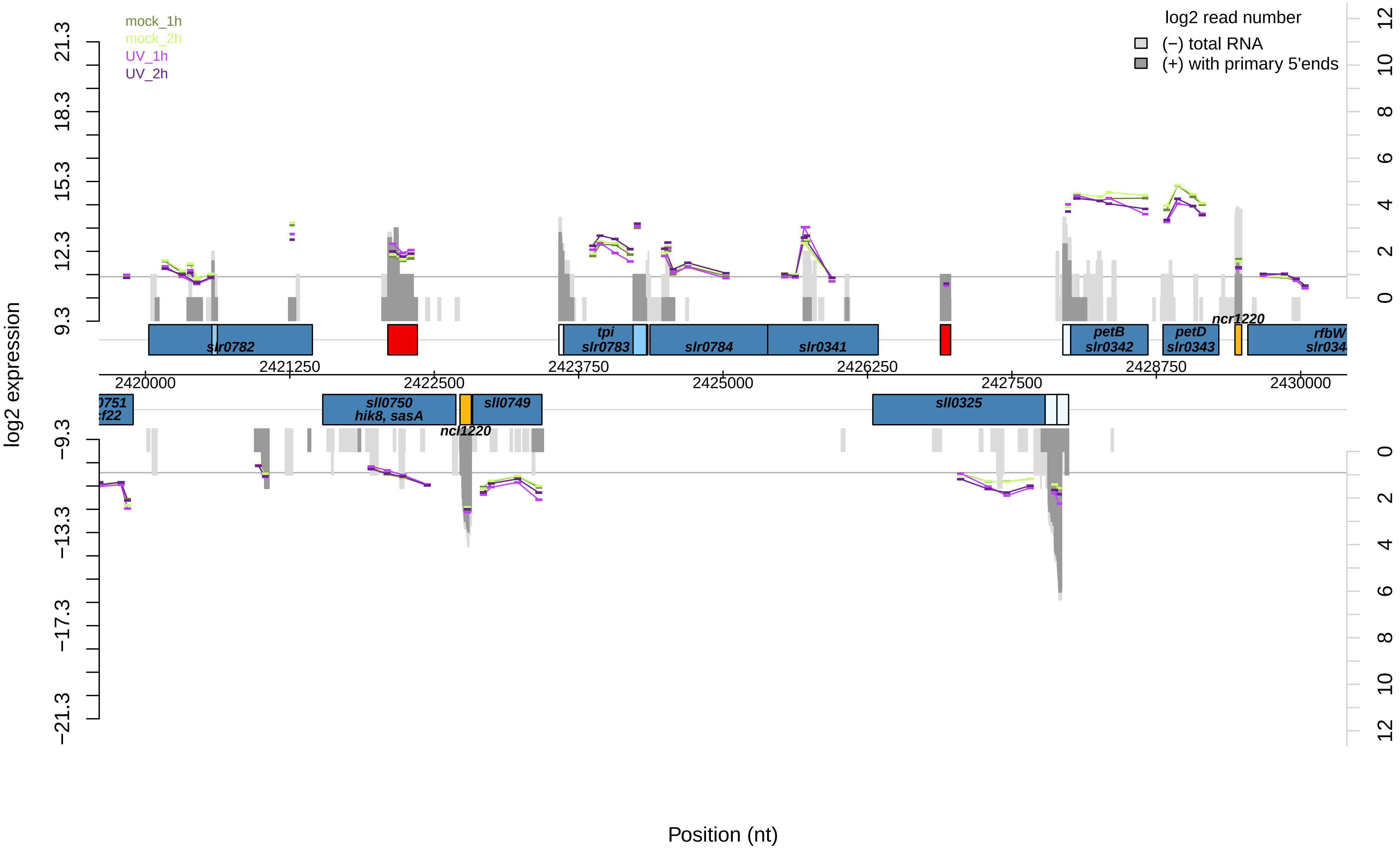
mock_1h
mock_2h
UV_1h
UV_2h
log2 read number
(−) total RNA
(+) with primary 5'ends
log2 expression
Position (nt)
slr0782
tpi
slr0783
slr0784
slr0341
petB
slr0342
petD
slr0343
ncr1220
rfbW
sll0750
hik8, sasA
ncl1220
sll0749
sll0325
2420000
2421250
2422500
2423750
2425000
2426250
2427500
2428750
2430000

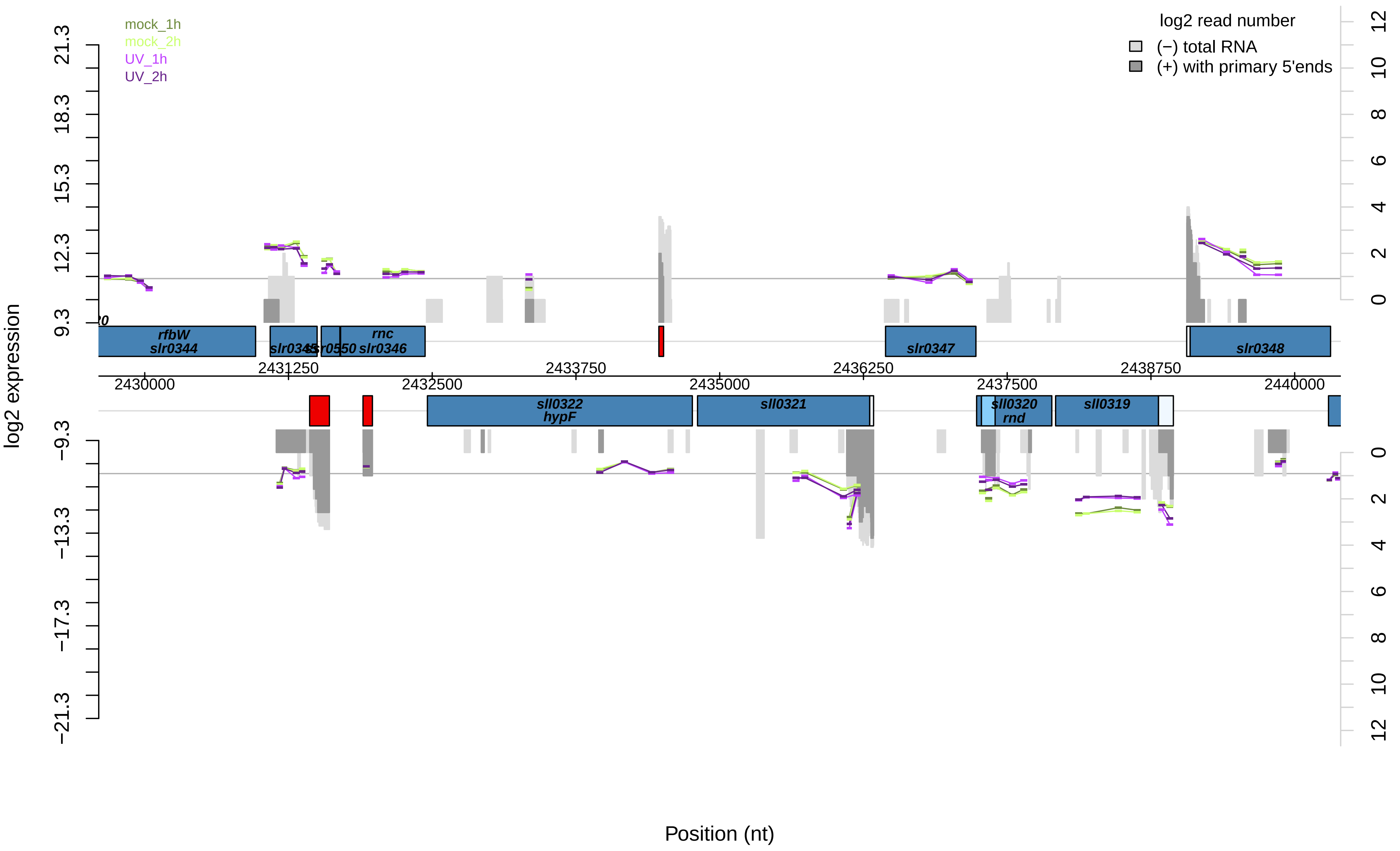

mock_1h
mock_2h
UV_1h
UV_2h
log2 read number
(−) total RNA
(+) with primary 5'ends
log2 expression
rfbW
slr0344
slr0345
ssr0550
rnc
slr0346
slr0347
slr0348
sll0322
hypF
sll0321
sll0320
rnd
sll0319
2430000
2431250
2432500
2433750
2435000
2436250
2437500
2438750
2440000
Position (nt)

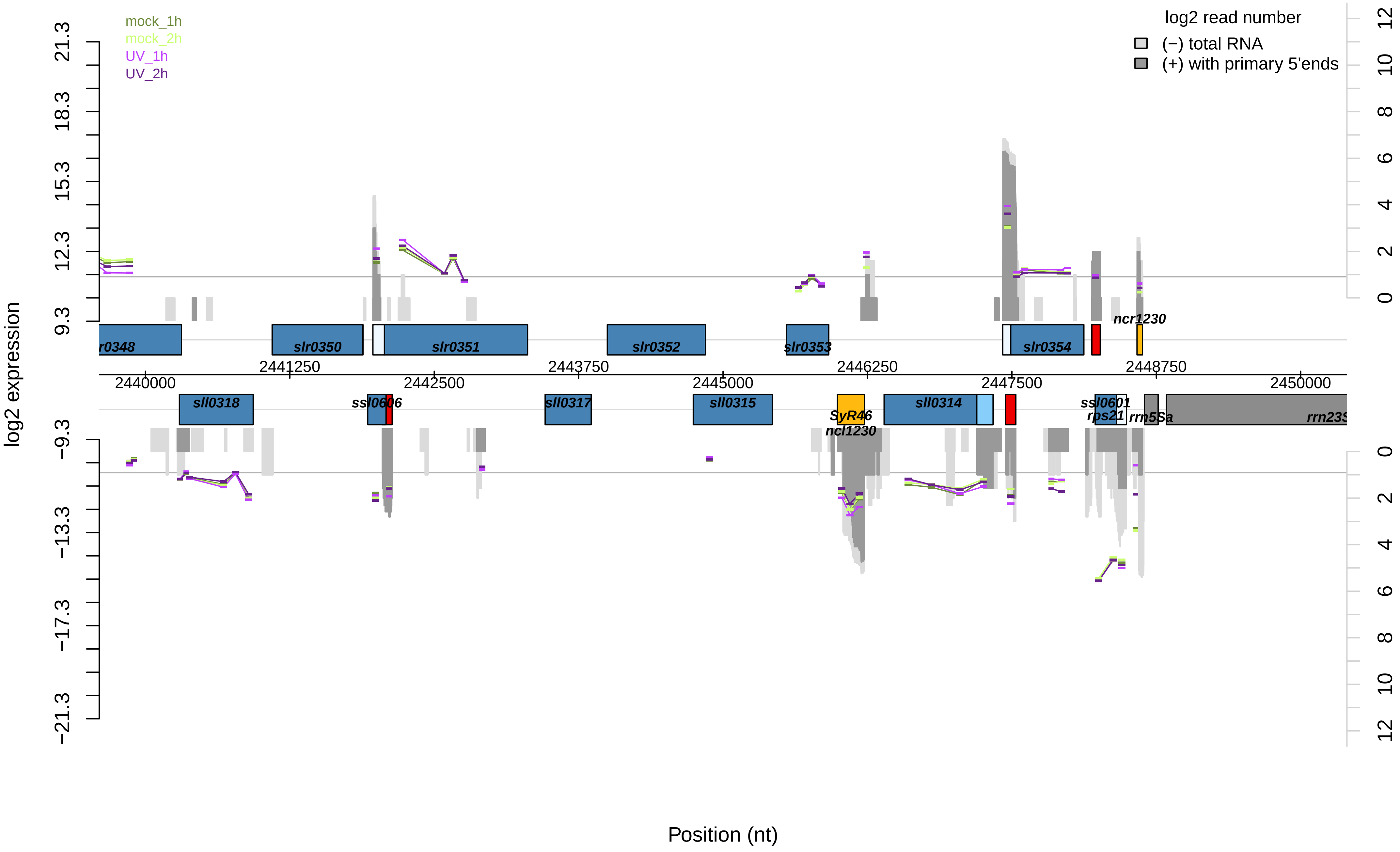

mock_1h
mock_2h
UV_1h
UV_2h
log2 read number
(−) total RNA
(+) with primary 5'ends
log2 expression
21.3
18.3
15.3
12.3
9.3
−9.3
−13.3
−17.3
−21.3
12
10
8
6
4
2
0
r0348
slr0350
slr0351
slr0352
slr0353
slr0354
ncr1230
2440000
2441250
2442500
2443750
2445000
2446250
2447500
2448750
2450000
sll0318
ssl0606
sll0317
sll0315
SyR46
ncl1230
sll0314
ssl0601
rps21
rrn5Sa
rrn23S
Position (nt)

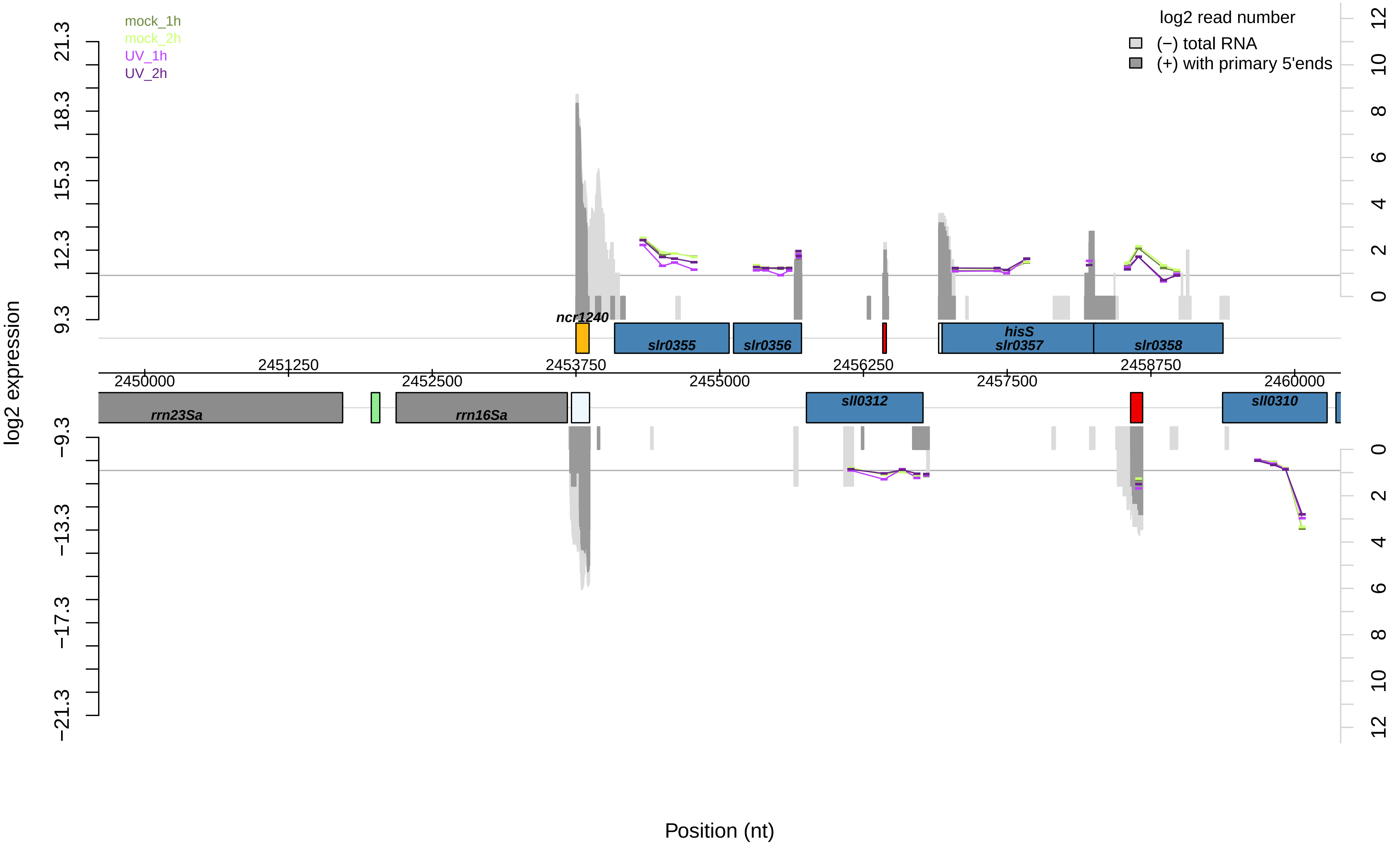

mock_1h
mock_2h
UV_1h
UV_2h
log2 read number
(−) total RNA
(+) with primary 5'ends
log2 expression
21.3
18.3
15.3
12.3
9.3
−9.3
−13.3
−17.3
−21.3
12
10
8
6
4
2
0
ncr1240
slr0355
slr0356
hisS
slr0357
slr0358
2451250
2453750
2456250
2458750
2450000
2452500
2455000
2457500
2460000
rrn23Sa
rrn16Sa
sll0312
sll0310
Position (nt)

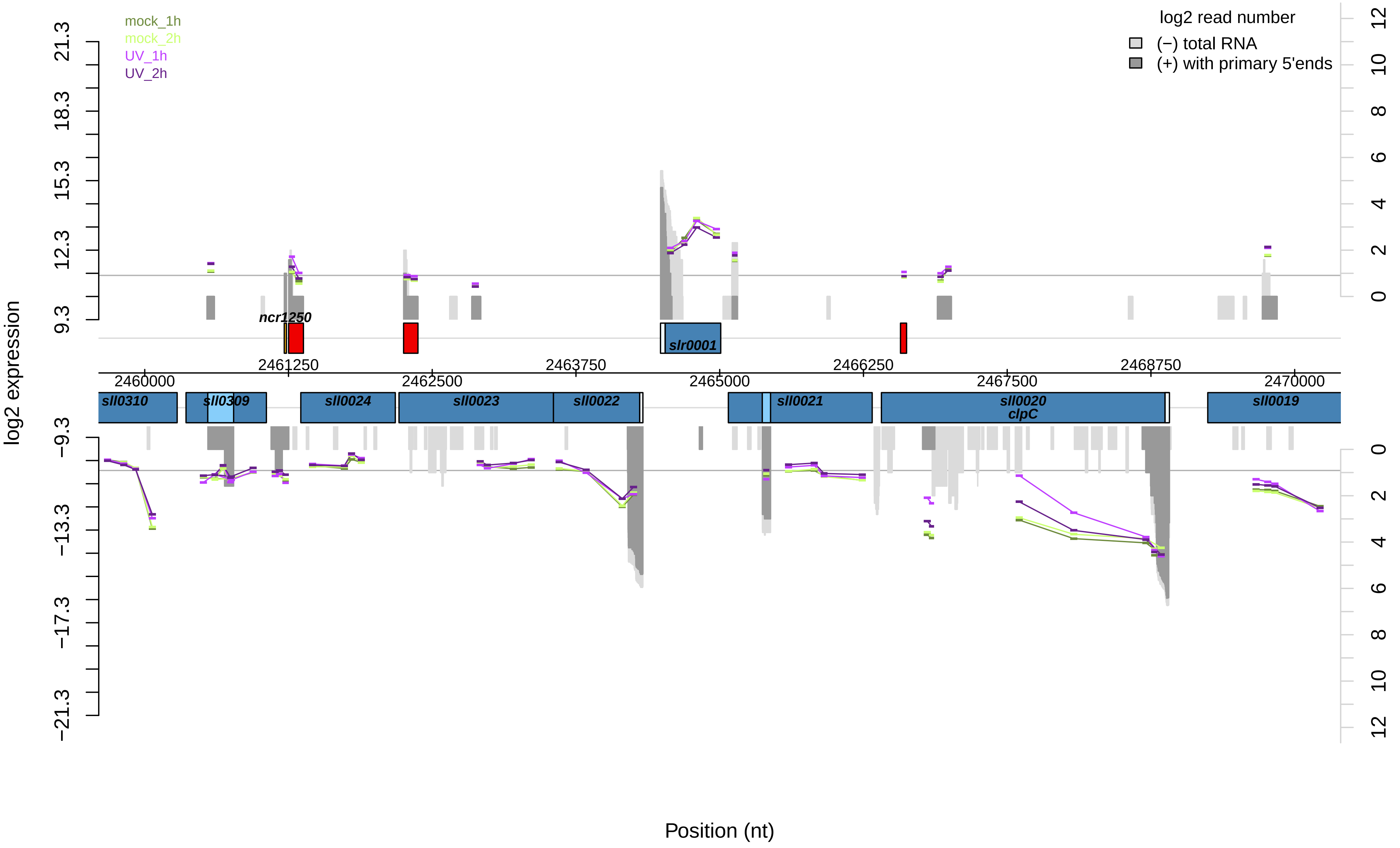

mock_1h
mock_2h
UV_1h
UV_2h
log2 read number
(−) total RNA
(+) with primary 5'ends
log2 expression
21.3
18.3
15.3
12.3
9.3
−9.3
−13.3
−17.3
−21.3
12
10
8
6
4
2
0
ncr1250
slr0001
2461250
2463750
2466250
2468750
2460000
2462500
2465000
2467500
2470000
sll0310
sll0309
sll0024
sll0023
sll0022
sll0021
sll0020
clpC
sll0019
Position (nt)

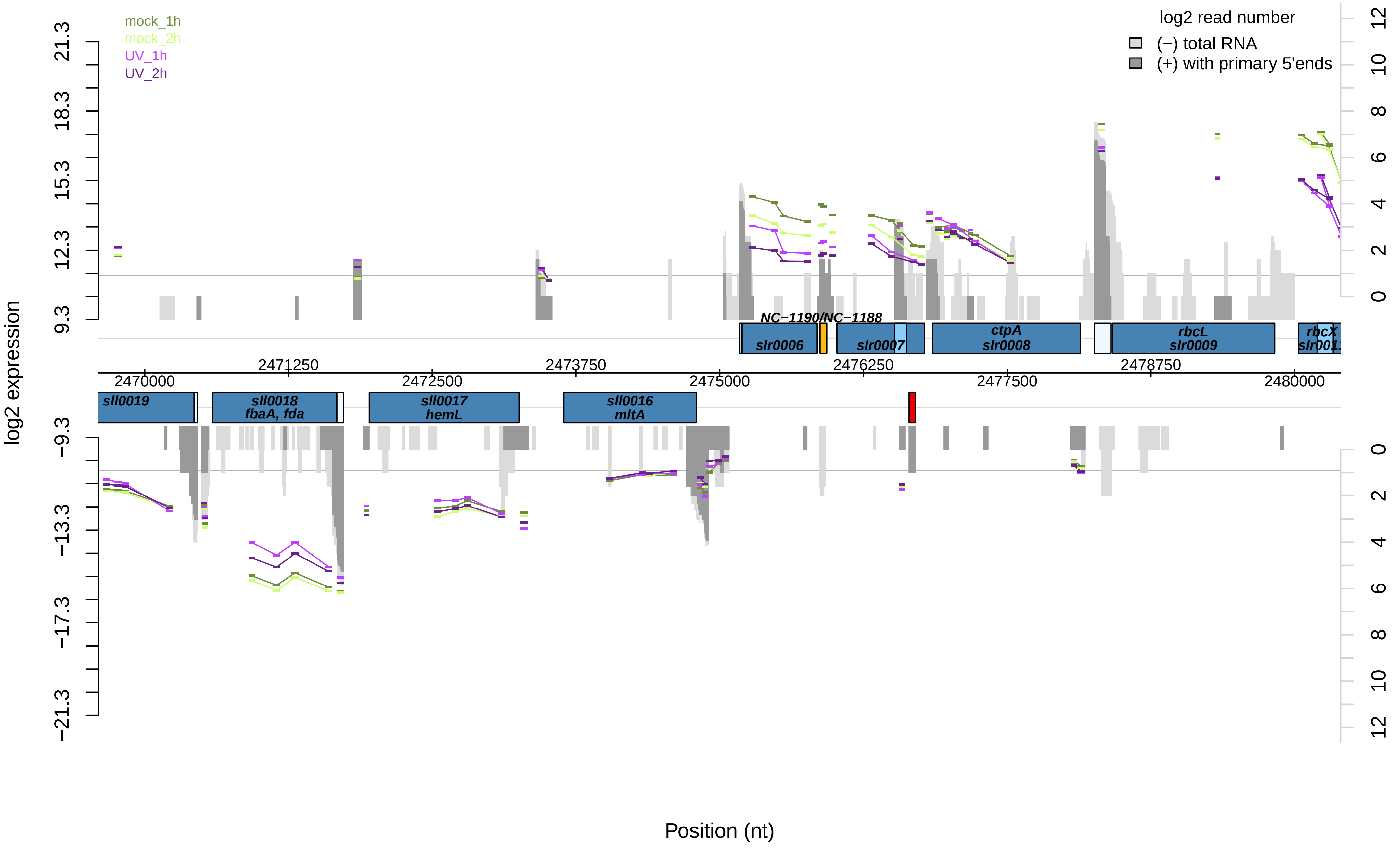
mock_1h
mock_2h
UV_1h
UV_2h
log2 read number
(−) total RNA
(+) with primary 5'ends
21.3
18.3
15.3
12.3
9.3
−9.3
−13.3
−17.3
−21.3
12
10
8
6
4
2
0
log2 expression
NC−1190/NC−1188
slr0006
slr0007
ctpA
slr0008
rbcL
slr0009
rbcX
2471250
2473750
2476250
2478750
2470000
2472500
2475000
2477500
2480000
sll0019
sll0018
fbaA, fda
sll0017
hemL
sll0016
mltA
Position (nt)

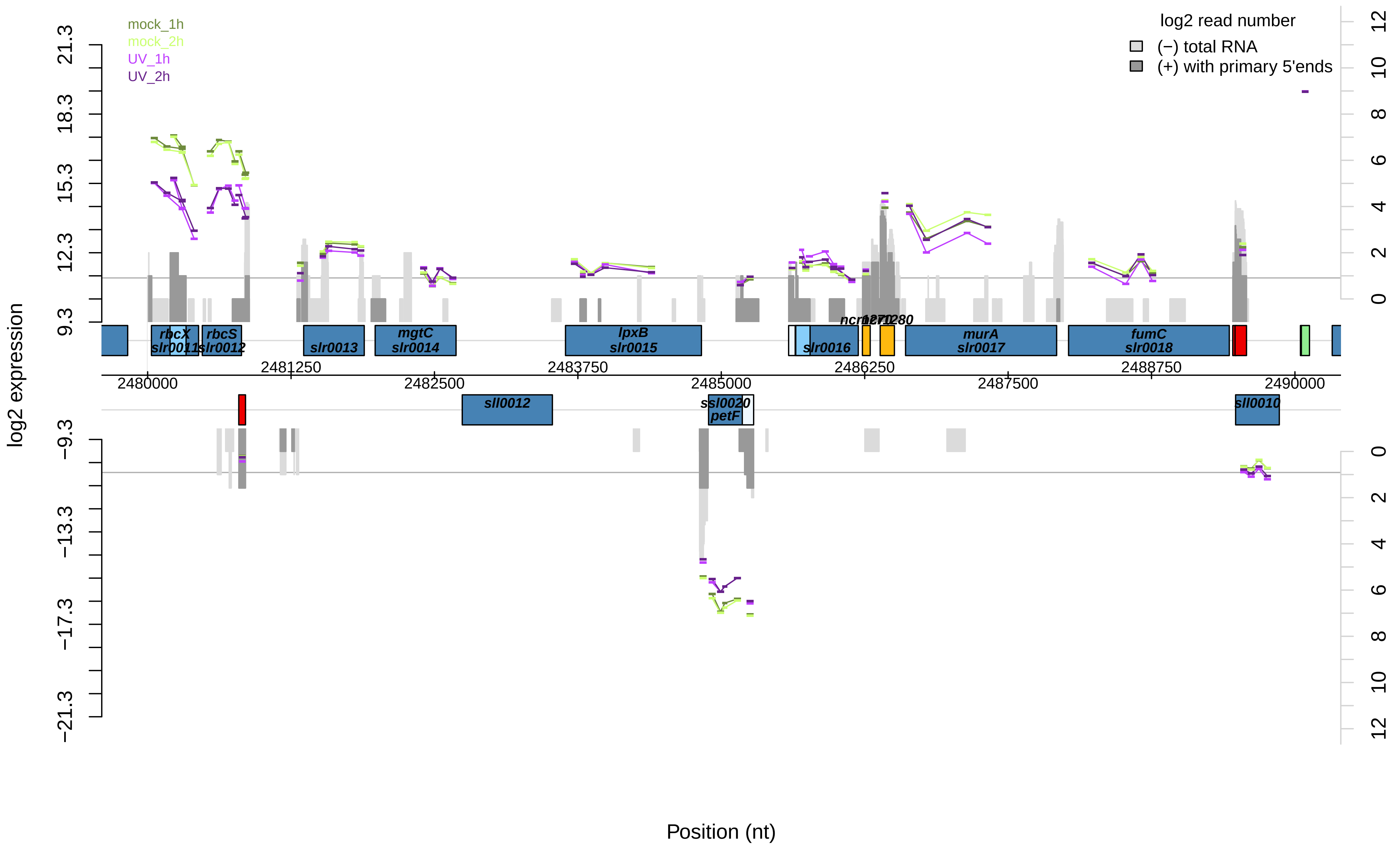

mock_1h
mock_2h
UV_1h
UV_2h
log2 read number
(−) total RNA
(+) with primary 5'ends
log2 expression
Position (nt)
rbcX
slr0011
rbcS
slr0012
slr0013
mgtC
slr0014
lpxB
slr0015
slr0016
ncr1270
1280
murA
slr0017
fumC
slr0018
sll0012
ssl0020
petF
sll0010
2480000
2481250
2482500
2483750
2485000
2486250
2487500
2488750
2490000
21.3
18.3
15.3
12.3
9.3
−9.3
−13.3
−17.3
−21.3

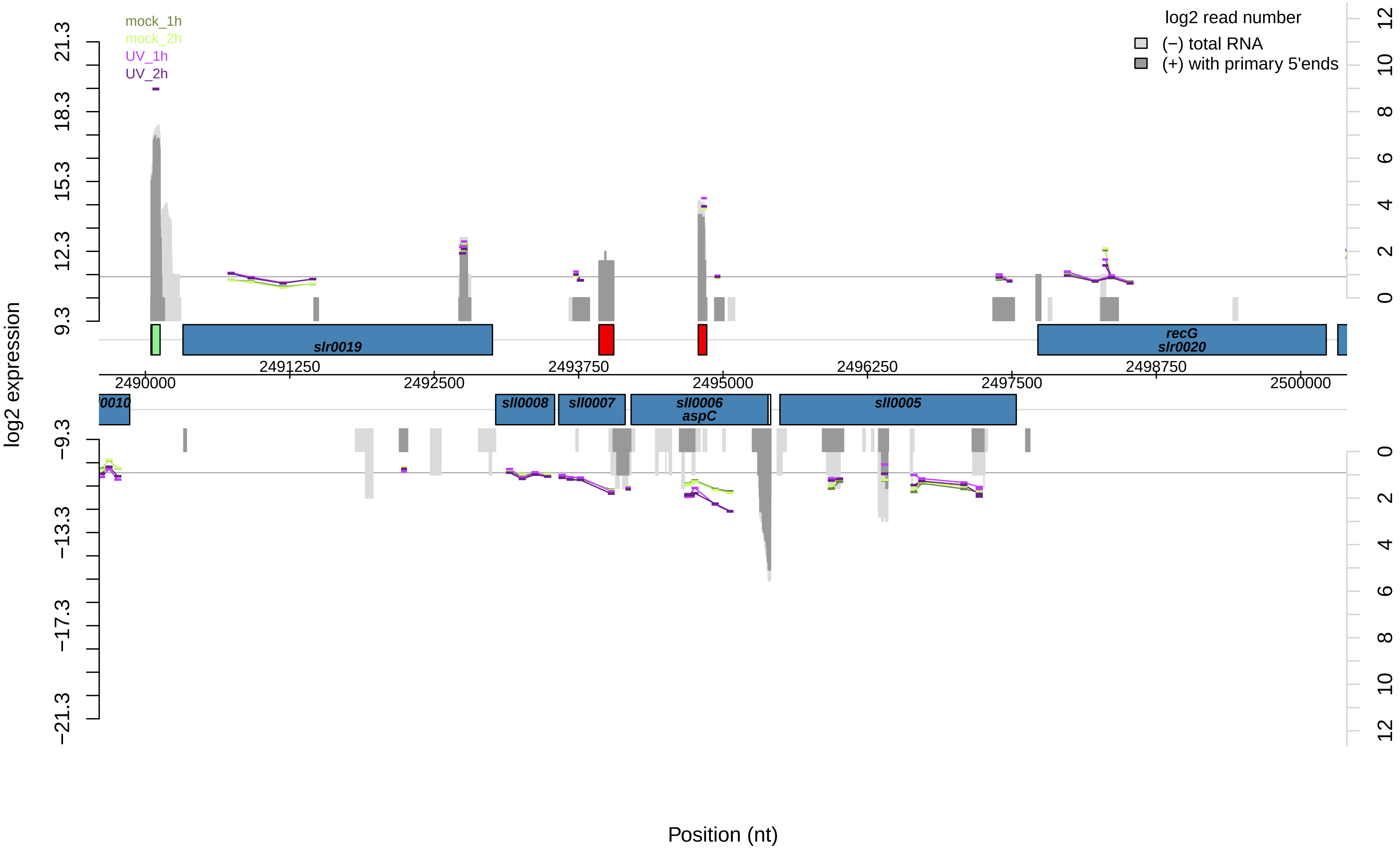

mock_1h
mock_2h
UV_1h
UV_2h
log2 read number
(−) total RNA
(+) with primary 5'ends
log2 expression
21.3
18.3
15.3
12.3
9.3
−9.3
−13.3
−17.3
−21.3
12
10
8
6
4
2
0
slr0019
recG
slr0020
2491250
2493750
2496250
2498750
2490000
2492500
2495000
2497500
2500000
0010
sll0008
sll0007
sll0006
aspC
sll0005
Position (nt)

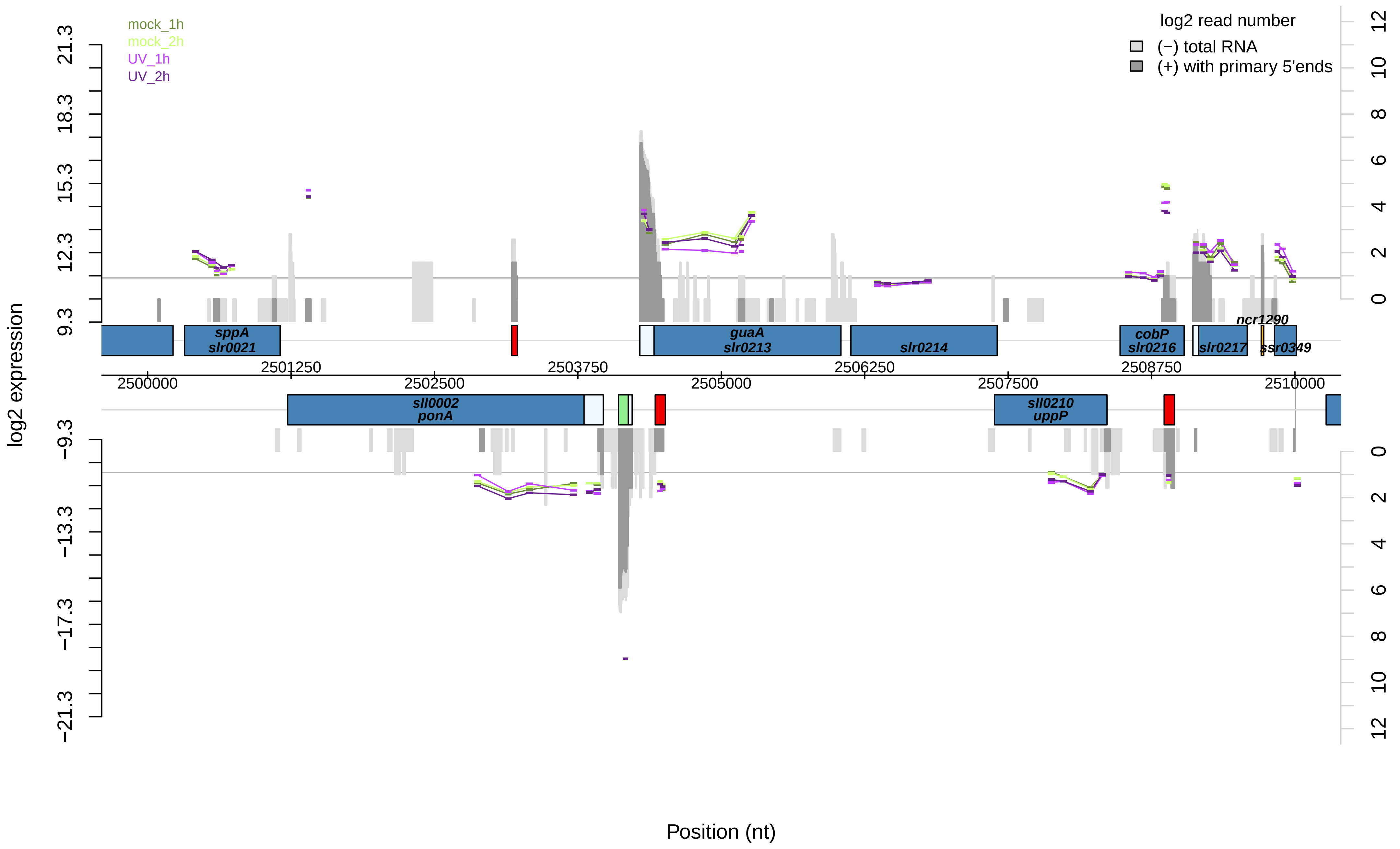

mock_1h
mock_2h
UV_1h
UV_2h
log2 read number
(−) total RNA
(+) with primary 5'ends
log2 expression
21.3
18.3
15.3
12.3
9.3
−9.3
−13.3
−17.3
−21.3
sppA
slr0021
guaA
slr0213
slr0214
cobP
slr0216
slr0217
ncr1290
ssr0349
sll0002
ponA
sll0210
uppP
2500000
2501250
2502500
2503750
2505000
2506250
2507500
2508750
2510000
Position (nt)

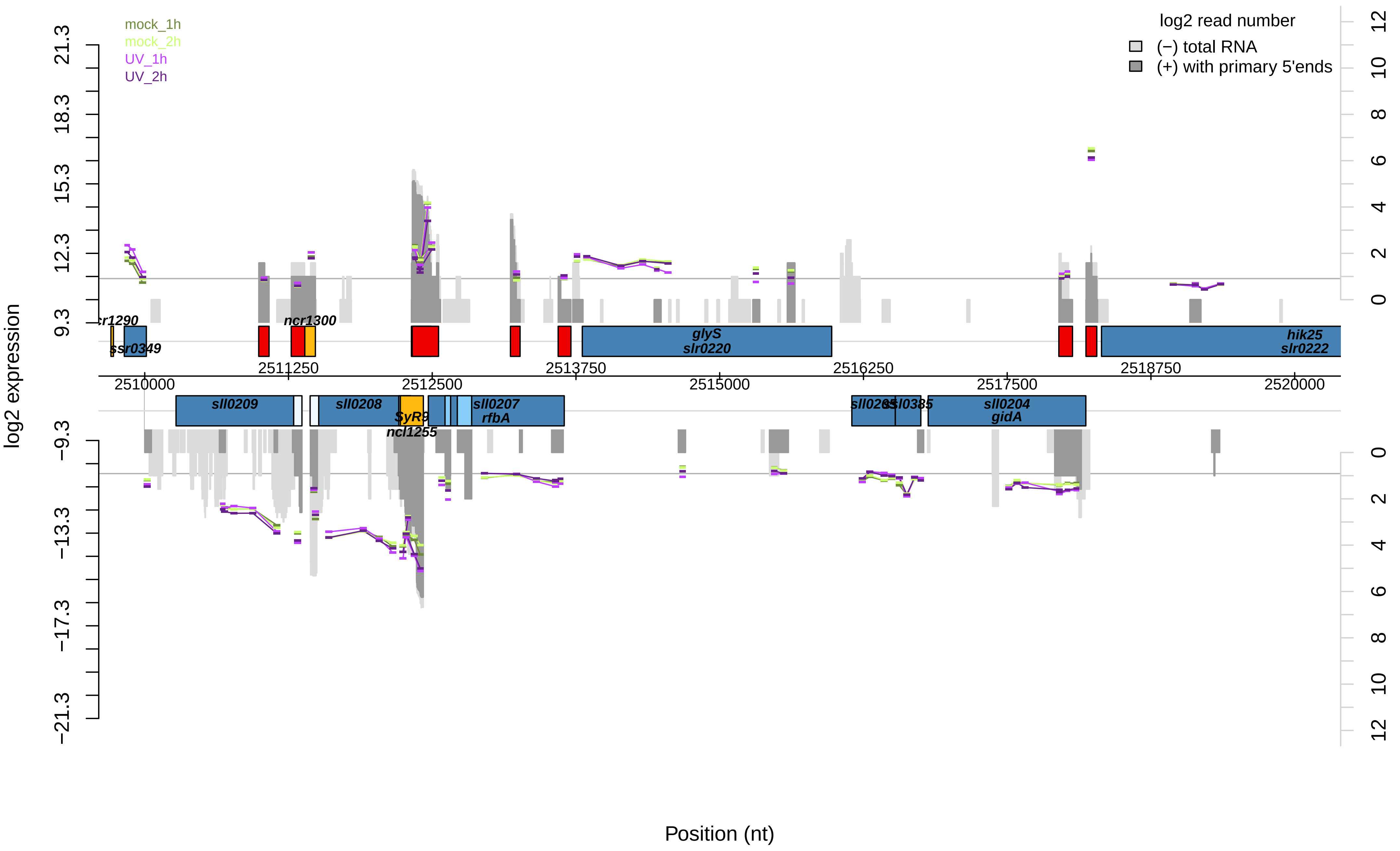

mock_1h
mock_2h
UV_1h
UV_2h
log2 read number
(−) total RNA
(+) with primary 5'ends
log2 expression
21.3
18.3
15.3
12.3
9.3
−9.3
−13.3
−17.3
−21.3
12
10
8
6
4
2
0
ncr1300
ssr0349
glyS
slr0220
hik25
slr0222
2511250
2513750
2516250
2518750
2510000
2512500
2515000
2517500
2520000
sll0209
sll0208
SyR9
ncl1255
sll0207
rfbA
sll0204
gidA
Position (nt)

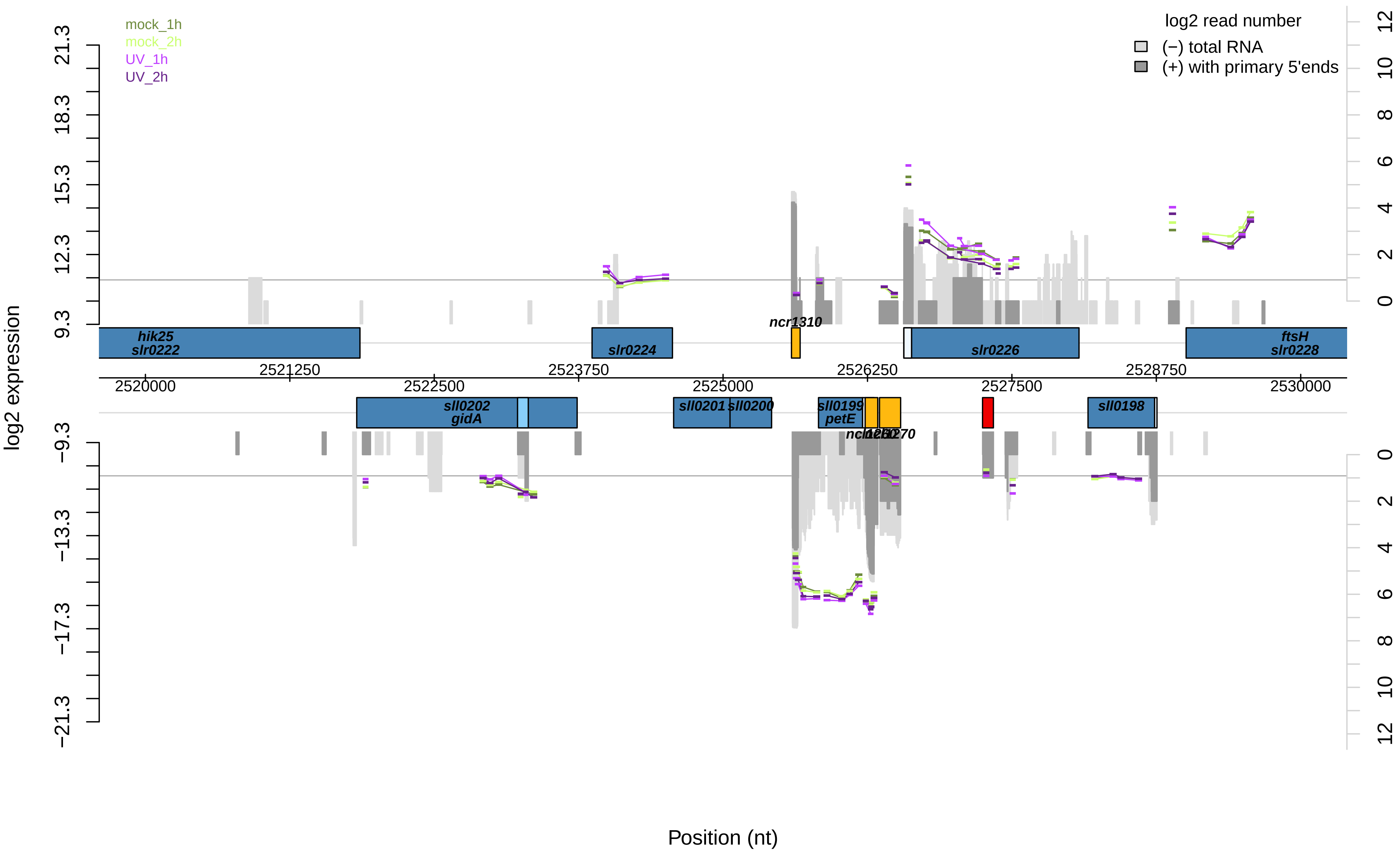
mock_1h
mock_2h
UV_1h
UV_2h
log2 read number
(−) total RNA
(+) with primary 5'ends
log2 expression
21.3
18.3
15.3
12.3
9.3
−9.3
−13.3
−17.3
−21.3
hik25
slr0222
slr0224
ncr1310
slr0226
ftsH
slr0228
2521250
2523750
2526250
2528750
2520000
2522500
2525000
2527500
2530000
sll0202
gidA
sll0201
sll0200
sll0199
petE
sll0198
Position (nt)

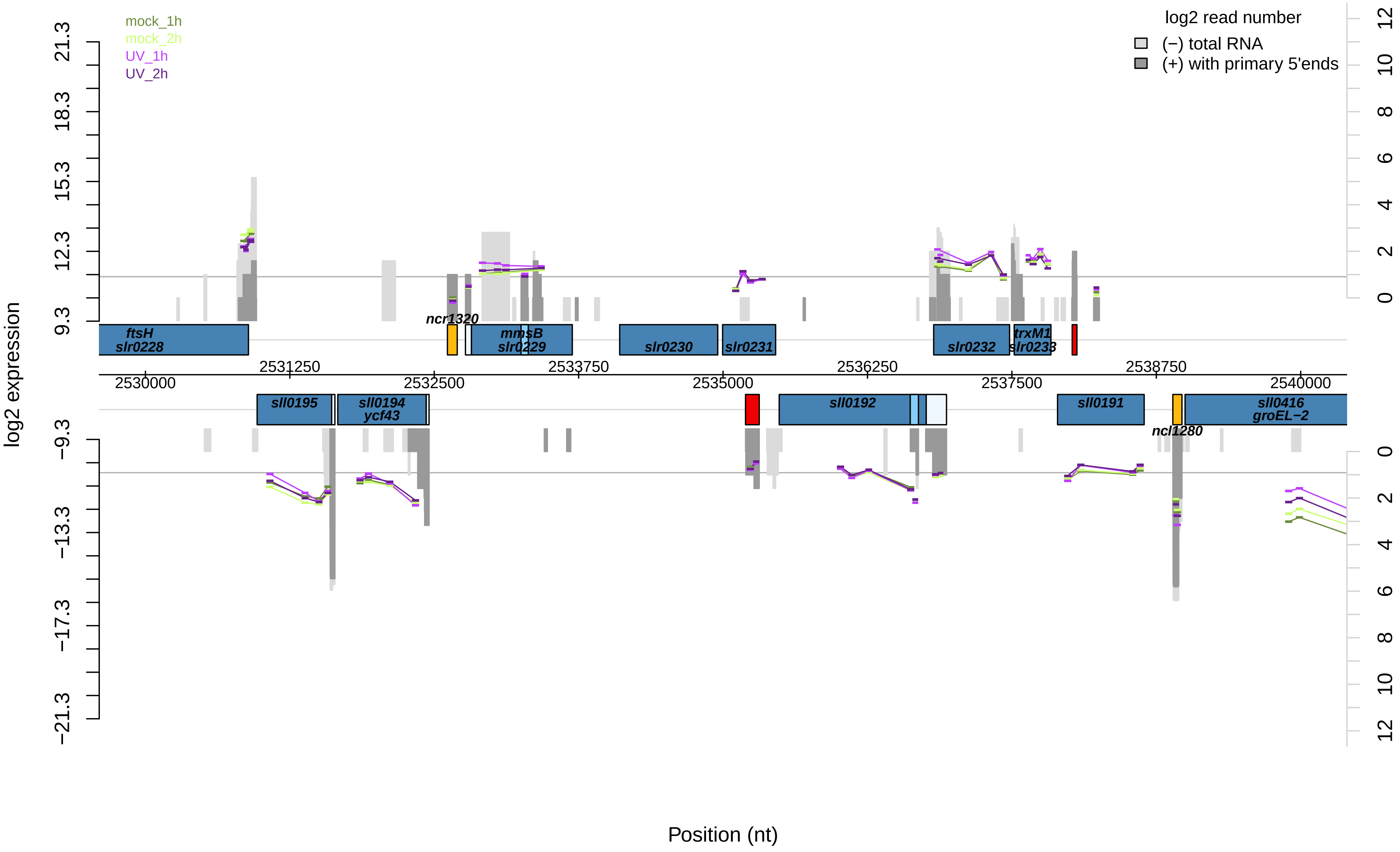

mock_1h
mock_2h
UV_1h
UV_2h
log2 read number
(−) total RNA
(+) with primary 5'ends
log2 expression
21.3
18.3
15.3
12.3
9.3
−9.3
−13.3
−17.3
−21.3
12
10
8
6
4
2
0
ftsH
slr0228
ncr1320
mmsB
slr0229
slr0230
slr0231
slr0232
trxM1
slr0233
2530000
2531250
2532500
2533750
2535000
2536250
2537500
2538750
2540000
sll0195
sll0194
ycf43
sll0192
sll0191
sll0416
groEL−2
ncl1280
Position (nt)

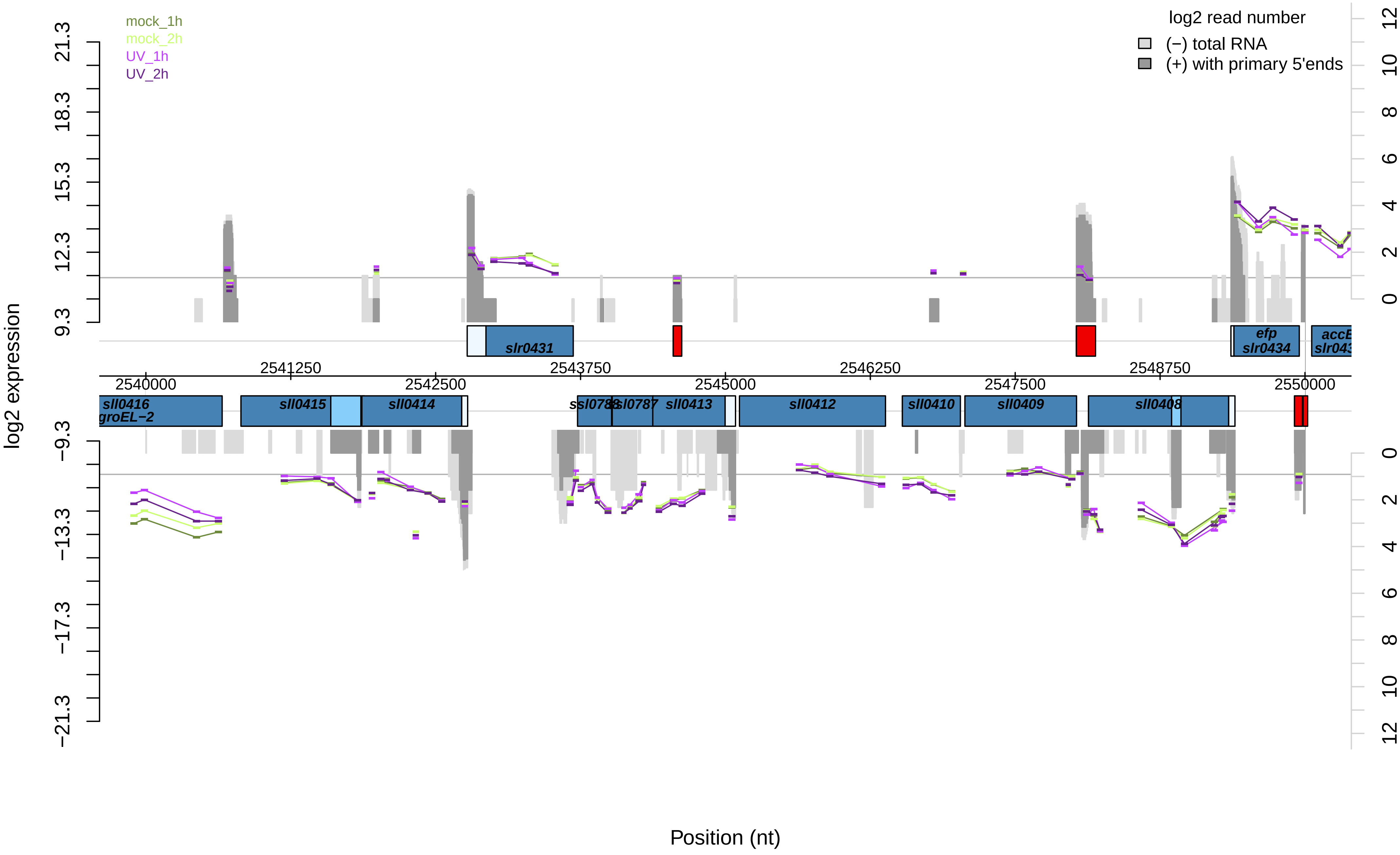

mock_1h
mock_2h
UV_1h
UV_2h
log2 read number
(−) total RNA
(+) with primary 5'ends
log2 expression
21.3
18.3
15.3
12.3
9.3
−9.3
−13.3
−17.3
−21.3
12
10
8
6
4
2
0
slr0431
efp
slr0434
2541250
2543750
2546250
2548750
2540000
2542500
2545000
2547500
2550000
sll0416
groEL−2
sll0415
sll0414
sll0413
sll0412
sll0410
sll0409
sll0408
Position (nt)

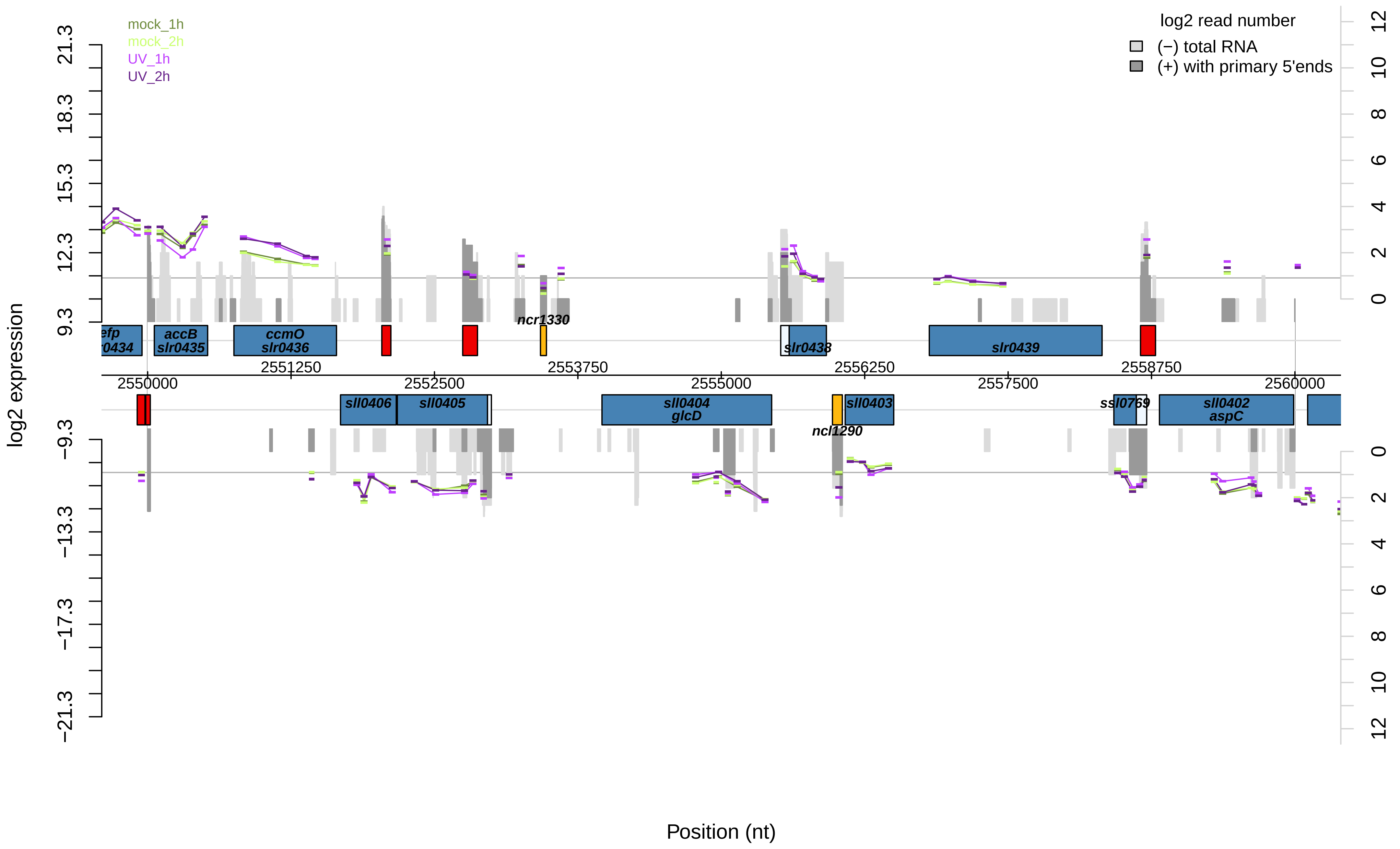

mock_1h
mock_2h
UV_1h
UV_2h
log2 read number
(−) total RNA
(+) with primary 5'ends
log2 expression
21.3
18.3
15.3
12.3
9.3
−9.3
−13.3
−17.3
−21.3
12
10
8
6
4
2
0
efp
accB
slr0435
ccmO
slr0436
ncr1330
slr0438
slr0439
2551250
2553750
2556250
2558750
2550000
2552500
2555000
2557500
2560000
sll0406
sll0405
sll0404
glcD
sll0403
ncl1290
ssl0769
sll0402
aspC
Position (nt)

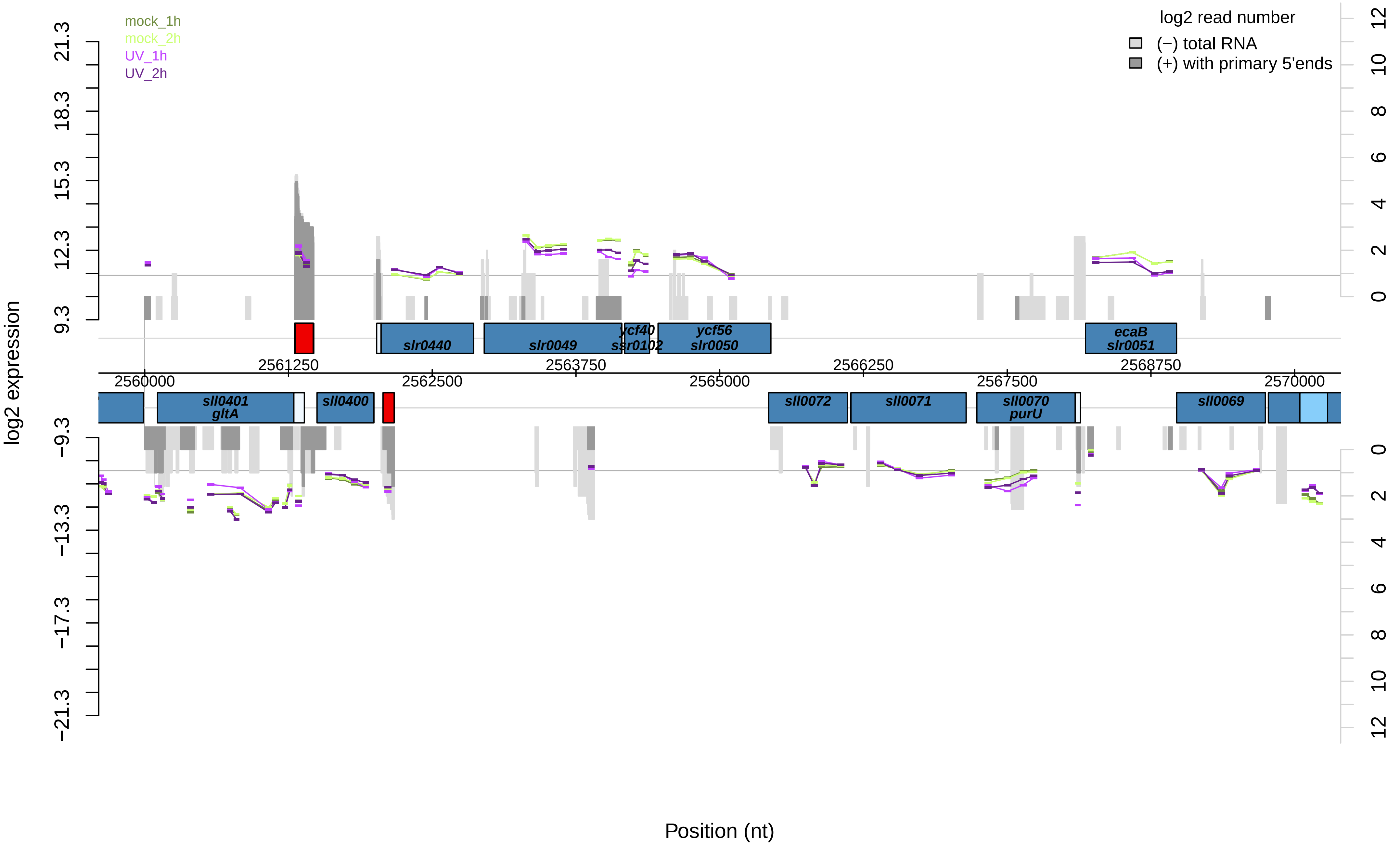

mock_1h
mock_2h
UV_1h
UV_2h
log2 read number
(−) total RNA
(+) with primary 5'ends
log2 expression
21.3
18.3
15.3
12.3
9.3
−9.3
−13.3
−17.3
−21.3
12
10
8
6
4
2
0
slr0440
slr0049
ycf40
ssr0102
ycf56
slr0050
ecaB
slr0051
2561250
2563750
2566250
2568750
2560000
2562500
2565000
2567500
2570000
sll0401
gltA
sll0400
sll0072
sll0071
sll0070
purU
sll0069
Position (nt)

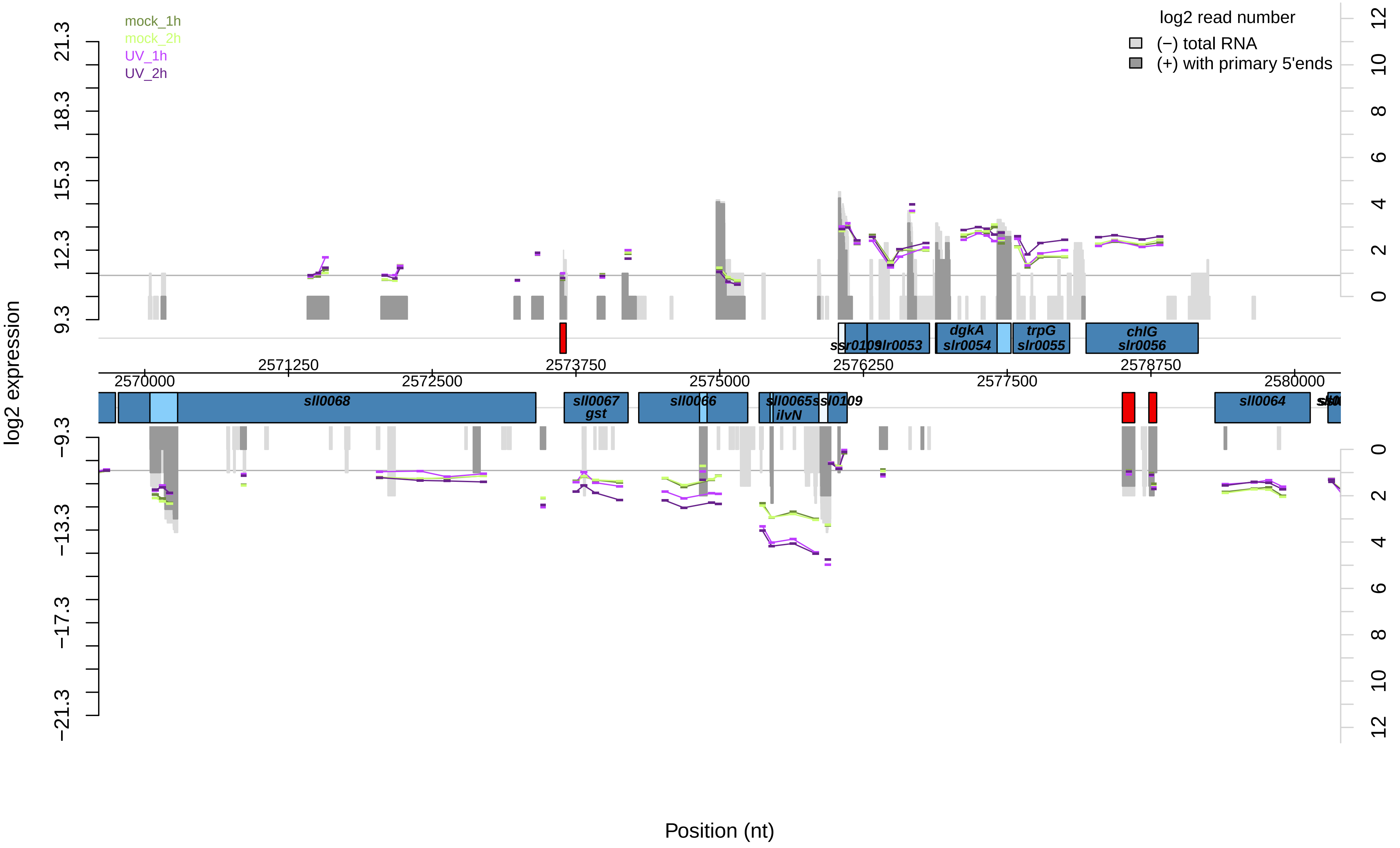

mock_1h
mock_2h
UV_1h
UV_2h
log2 read number
(−) total RNA
(+) with primary 5'ends
log2 expression
Position (nt)
dgkA
slr0054
trpG
slr0055
chlG
slr0056
slr0053
sll0068
sll0067
gst
sll0066
sll0065
ilvN
ssl0109
sll0064

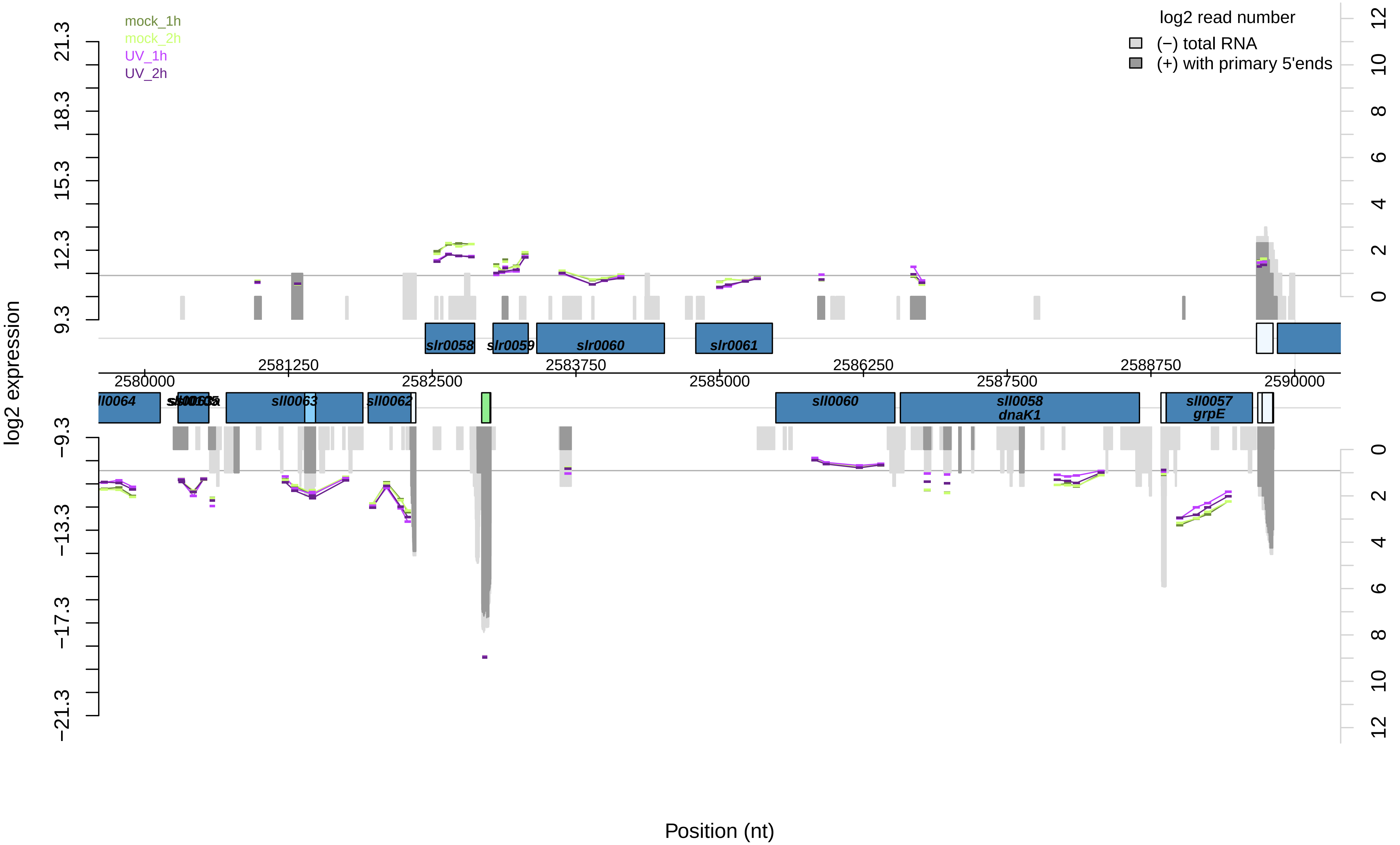

mock_1h
mock_2h
UV_1h
UV_2h
log2 read number
(−) total RNA
(+) with primary 5'ends
log2 expression
21.3
18.3
15.3
12.3
9.3
−9.3
−13.3
−17.3
−21.3
12
10
8
6
4
2
0
slr0058
slr0059
slr0060
slr0061
2580000
2581250
2582500
2583750
2585000
2586250
2587500
2588750
2590000
sll0063
sll0062
sll0060
sll0058
dnaK1
sll0057
grpE
Position (nt)

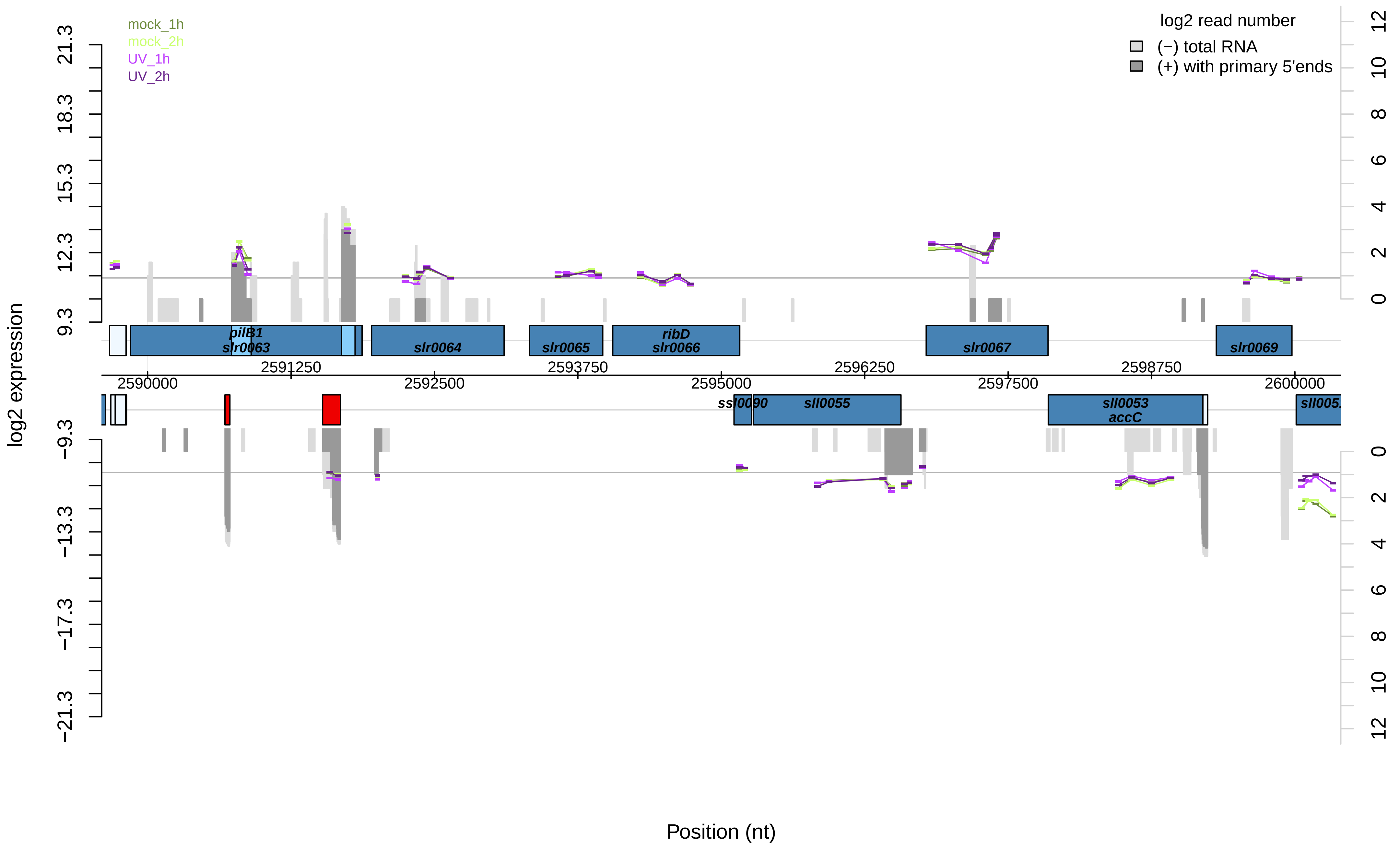

mock_1h
mock_2h
UV_1h
UV_2h
log2 read number
(−) total RNA
(+) with primary 5'ends
log2 expression
21.3
18.3
15.3
12.3
9.3
−9.3
−13.3
−17.3
−21.3
12
10
8
6
4
2
0
pilB1
slr0063
slr0064
slr0065
ribD
slr0066
slr0067
slr0069
2591250
2593750
2596250
2598750
2590000
2592500
2595000
2597500
2600000
ssl0090
sll0055
sll0053
accC
Position (nt)

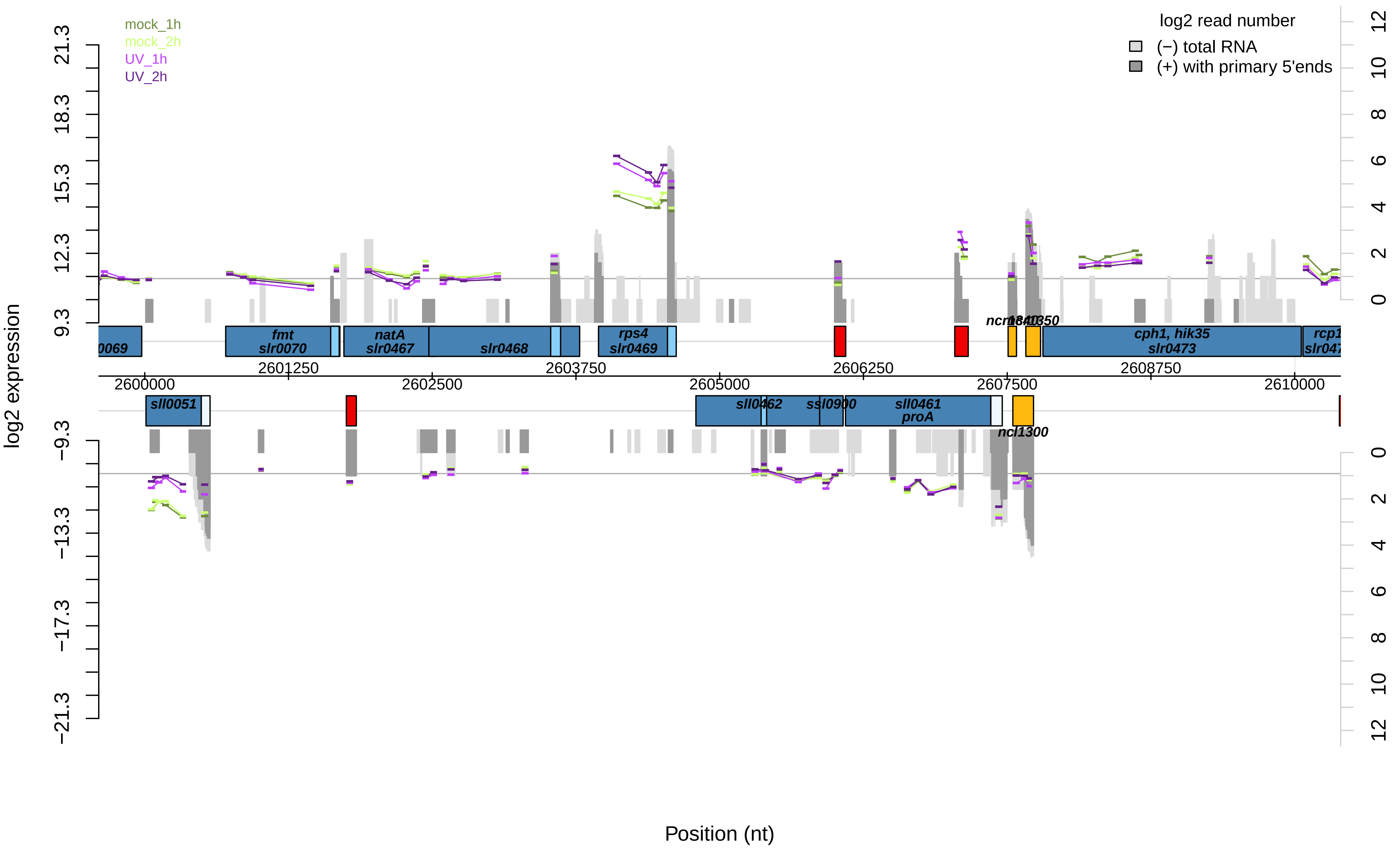

mock_1h
mock_2h
UV_1h
UV_2h
log2 read number
(−) total RNA
(+) with primary 5'ends
log2 expression
21.3
18.3
15.3
12.3
9.3
−9.3
−13.3
−17.3
−21.3
12
10
8
6
4
2
0
0069
fmt
slr0070
natA
slr0467
slr0468
rps4
slr0469
ncr1840
ncl0350
cph1, hik35
slr0473
rcp1
slr047
2601250
2603750
2606250
2608750
2600000
2602500
2605000
2607500
2610000
sll0051
sll0462
ssl0900
sll0461
proA
ncl1300
Position (nt)

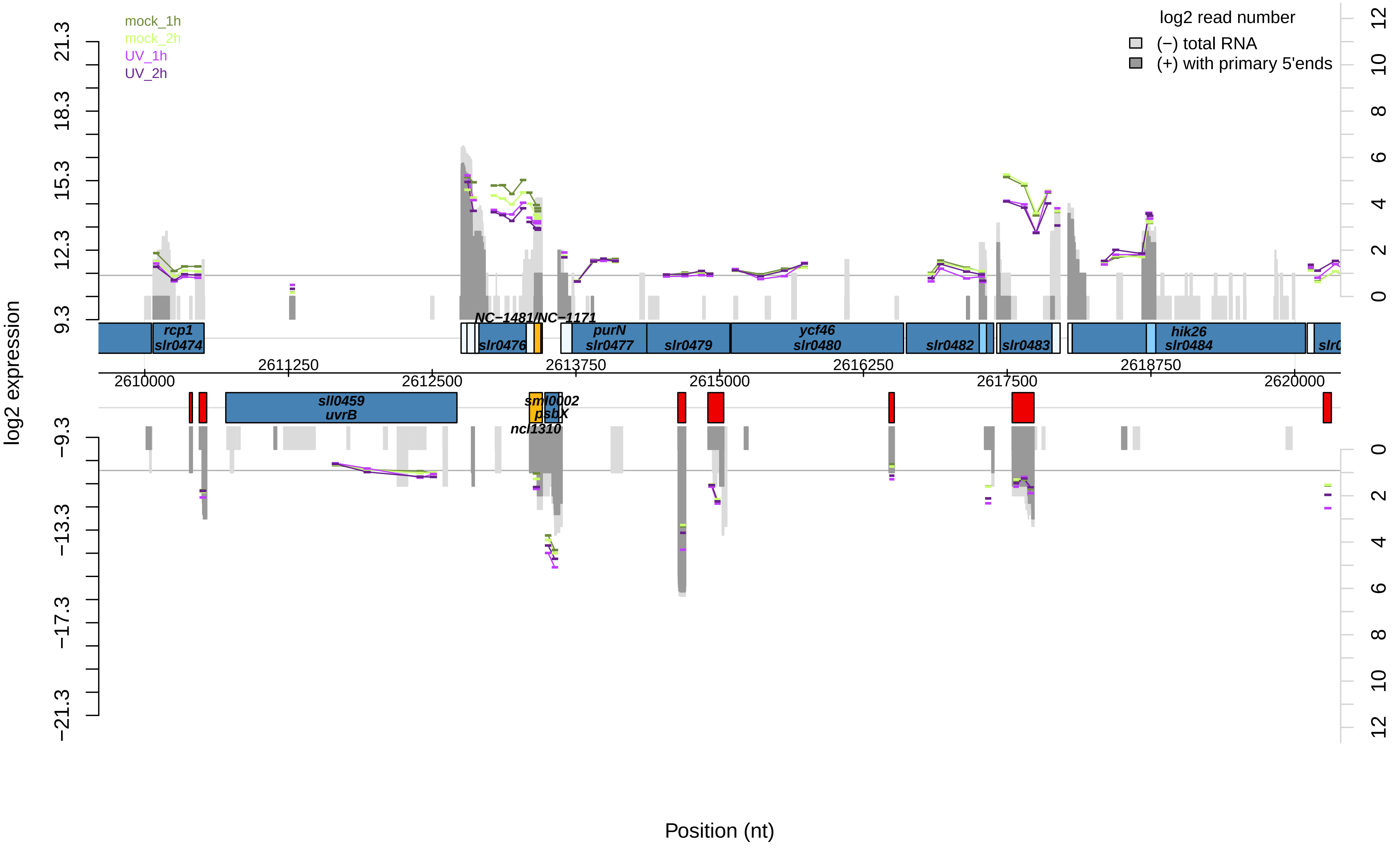

mock_1h
mock_2h
UV_1h
UV_2h
log2 read number
(−) total RNA
(+) with primary 5'ends
log2 expression
Position (nt)
21.3
18.3
15.3
12.3
9.3
−9.3
−13.3
−17.3
−21.3
NC−1481/NC−1171
rcp1
slr0474
slr0476
purN
slr0477
slr0479
ycf46
slr0480
slr0482
slr0483
hik26
slr0484
sll0459
uvrB
sml0002
psbX
ncl1310
2610000
2611250
2612500
2613750
2615000
2616250
2617500
2618750
2620000

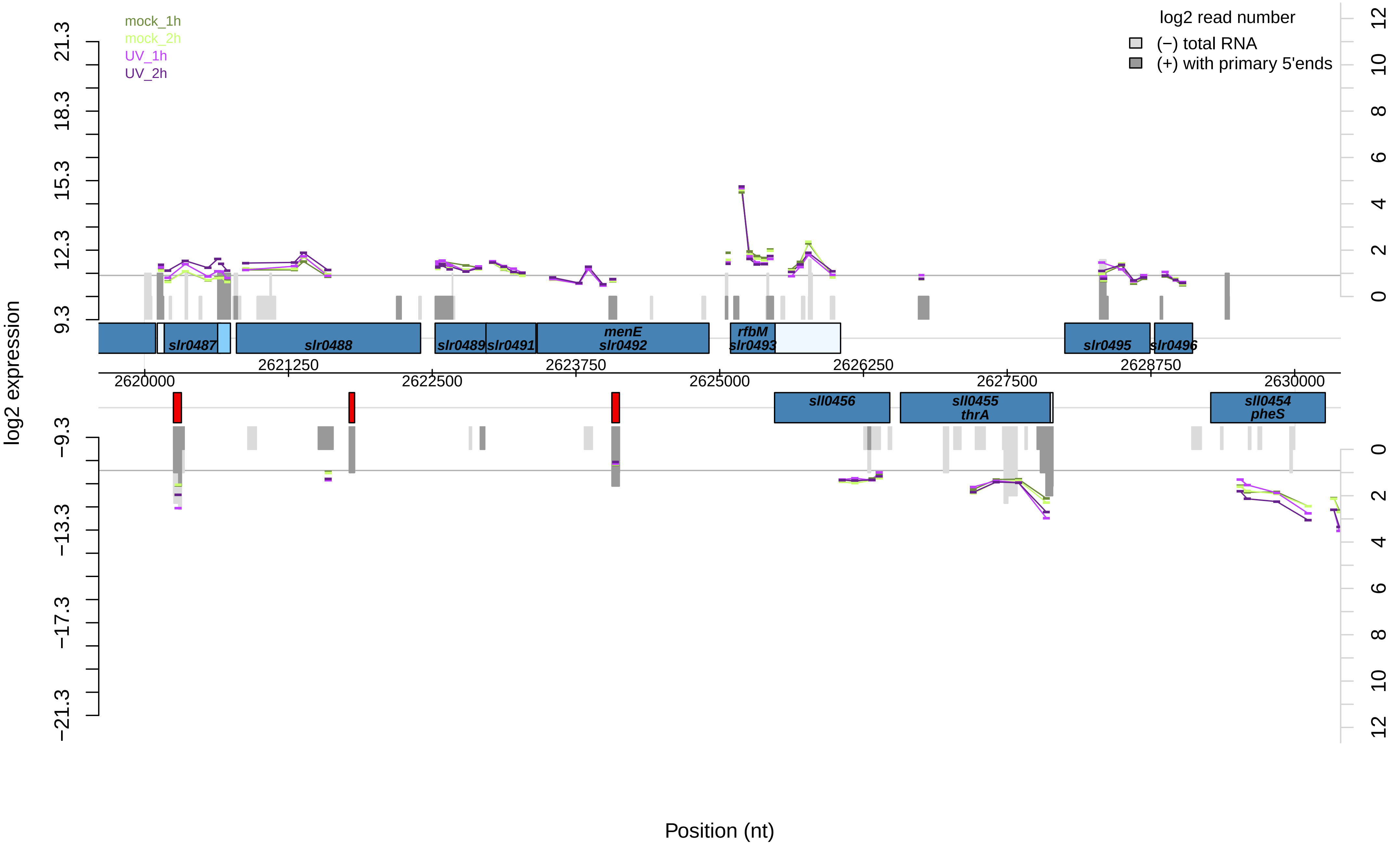

mock_1h
mock_2h
UV_1h
UV_2h
log2 read number
(−) total RNA
(+) with primary 5'ends
log2 expression
21.3
18.3
15.3
12.3
9.3
−9.3
−13.3
−17.3
−21.3
slr0487
slr0488
slr0489
slr0491
menE
slr0492
rfbM
slr0493
slr0495
slr0496
sll0456
sll0455
thrA
sll0454
pheS
2620000
2621250
2622500
2623750
2625000
2626250
2627500
2628750
2630000
Position (nt)

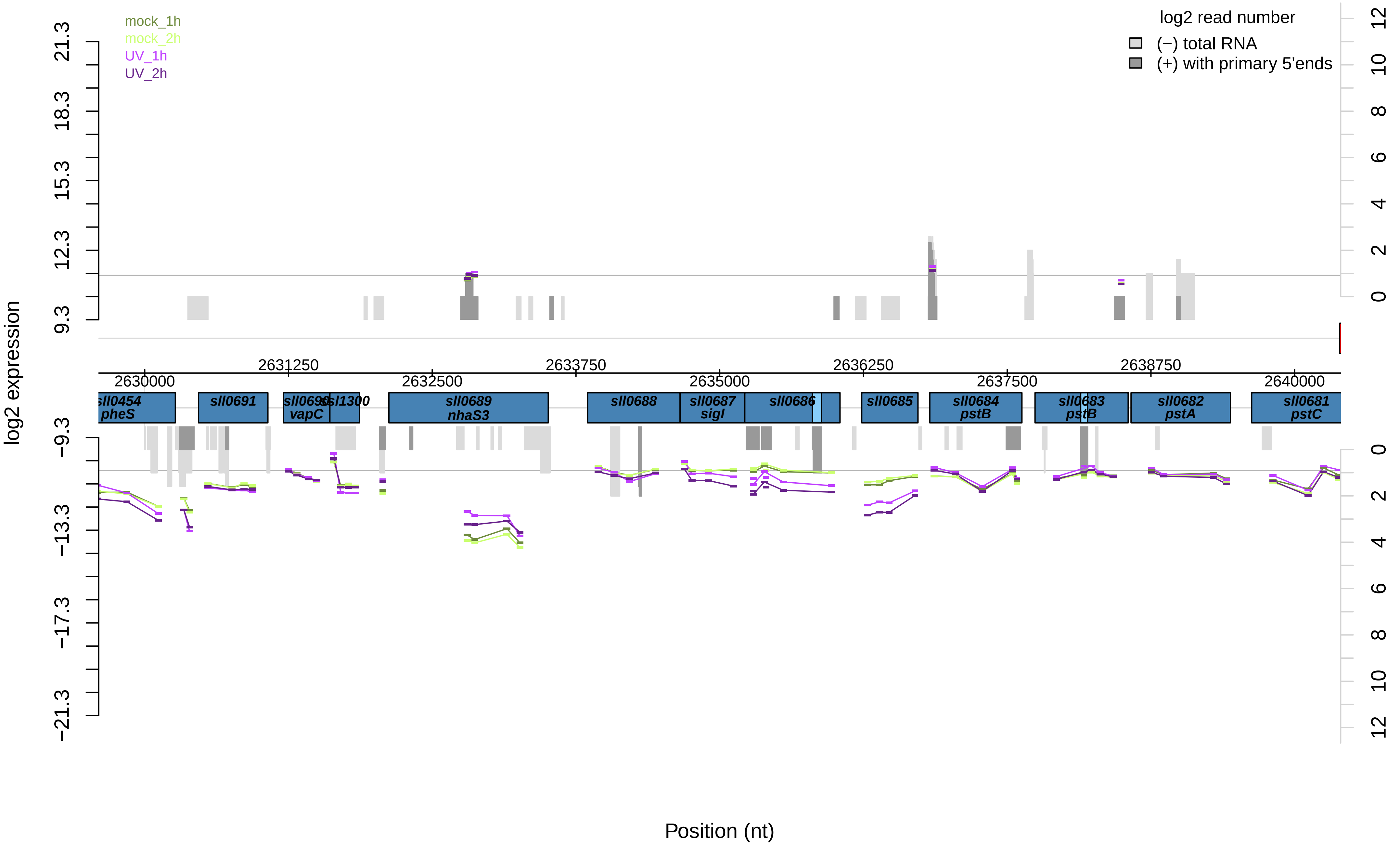

mock_1h
mock_2h
UV_1h
UV_2h
log2 read number
(−) total RNA
(+) with primary 5'ends
log2 expression
21.3
18.3
15.3
12.3
9.3
−9.3
−13.3
−17.3
−21.3
12
10
8
6
4
2
0
2631250
2633750
2636250
2638750
2630000
2632500
2635000
2637500
2640000
sll0454
pheS
sll0691
sll0690
vapC
ssl1300
sll0689
nhaS3
sll0688
sll0687
sigI
sll0686
sll0685
sll0684
pstB
sll0683
pstB
sll0682
pstA
sll0681
pstC
Position (nt)

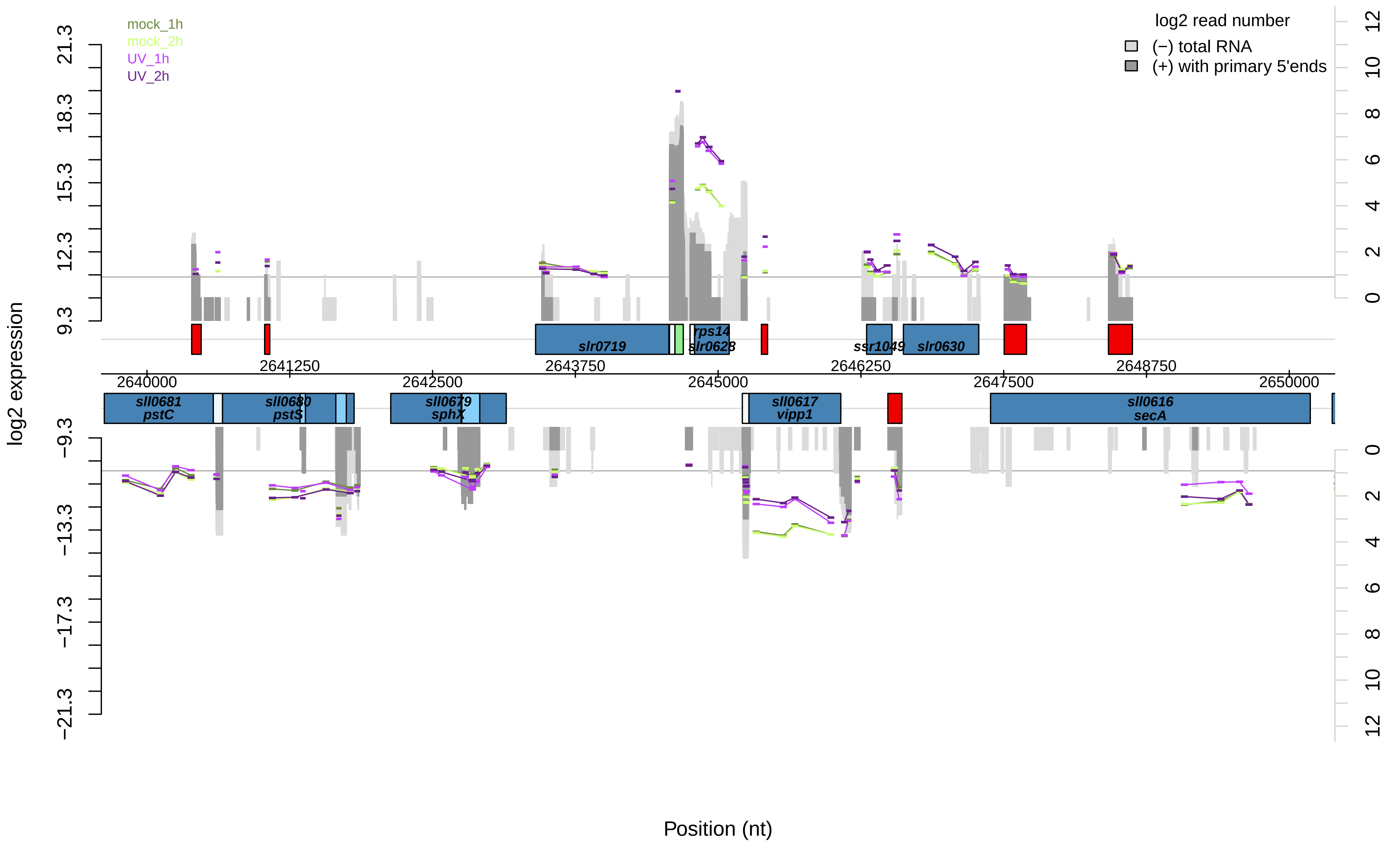

mock_1h
mock_2h
UV_1h
UV_2h
log2 read number
(−) total RNA
(+) with primary 5'ends
log2 expression
21.3
18.3
15.3
12.3
9.3
−9.3
−13.3
−17.3
−21.3
12
10
8
6
4
2
0
slr0719
rps14
slr0628
ssr1049
slr0630
2641250
2643750
2646250
2648750
2640000
2642500
2645000
2647500
2650000
sll0681
pstC
sll0680
pstS
sll0679
sphX
sll0617
vipp1
sll0616
secA
Position (nt)

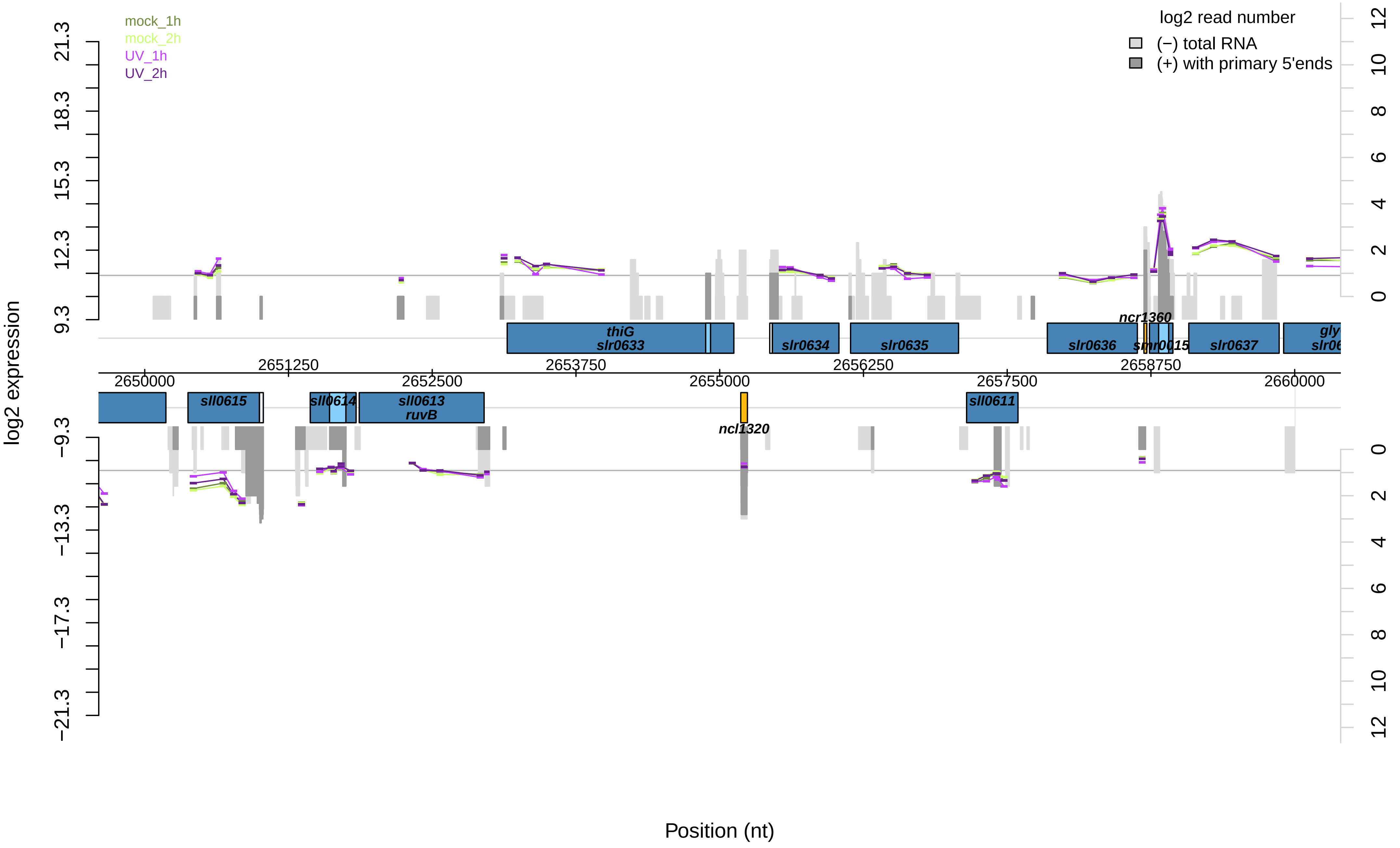
mock_1h
mock_2h
UV_1h
UV_2h
log2 read number
(−) total RNA
(+) with primary 5'ends
log2 expression
21.3
18.3
15.3
12.3
9.3
−9.3
−13.3
−17.3
−21.3
12
10
8
6
4
2
0
thiG
slr0633
slr0634
slr0635
slr0636
ncr1360
smr0015
slr0637
2651250
2653750
2656250
2658750
2650000
2652500
2655000
2657500
2660000
sll0615
sll0614
sll0613
ruvB
ncl1320
sll0611
Position (nt)

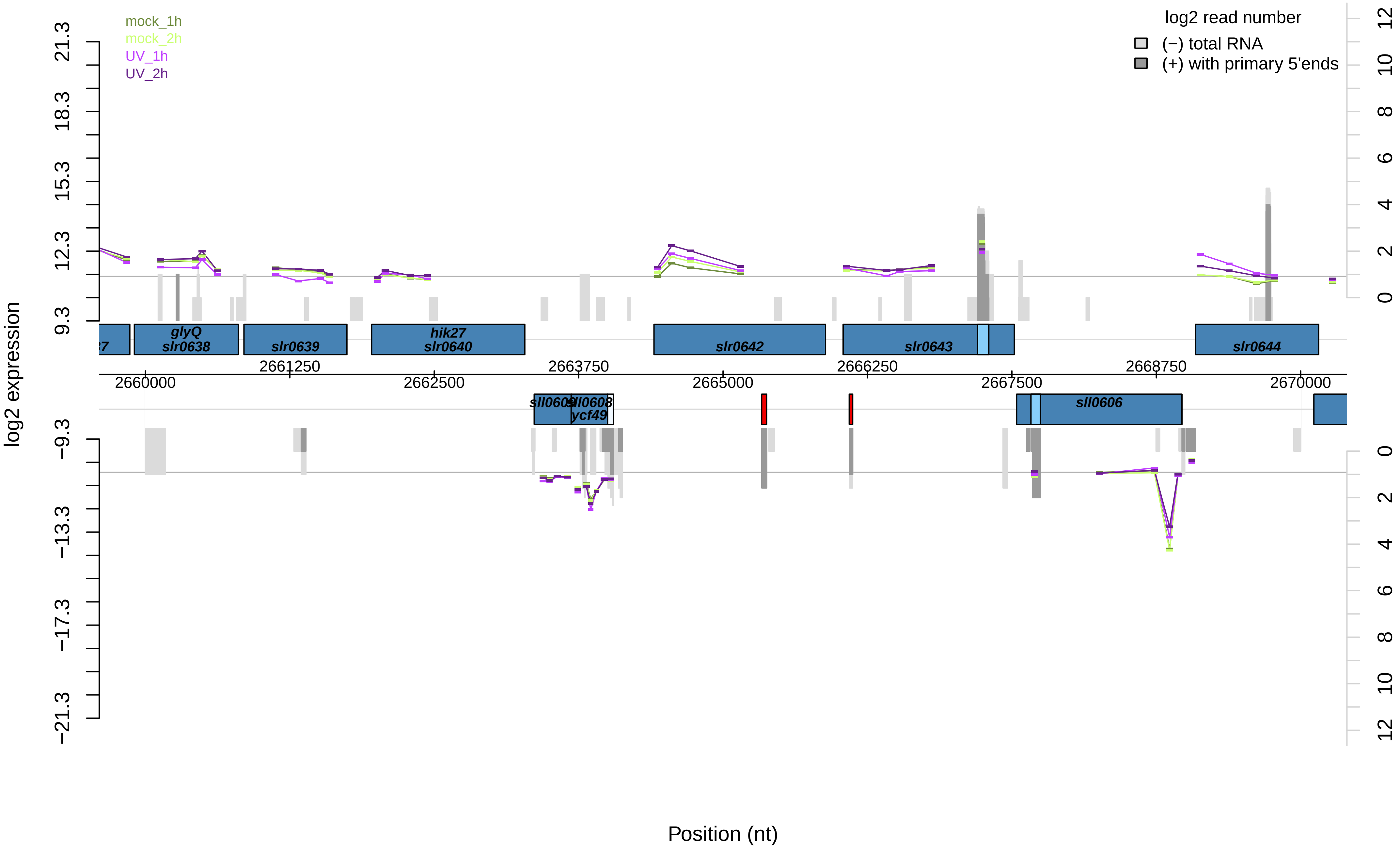

mock_1h
mock_2h
UV_1h
UV_2h
log2 read number
(−) total RNA
(+) with primary 5'ends
log2 expression
21.3
18.3
15.3
12.3
9.3
−9.3
−13.3
−17.3
−21.3
12
10
8
6
4
2
0
glyQ
slr0638
slr0639
hik27
slr0640
slr0642
slr0643
slr0644
2660000
2661250
2662500
2663750
2665000
2666250
2667500
2668750
2670000
sll0608
ycf49
sll0606
Position (nt)

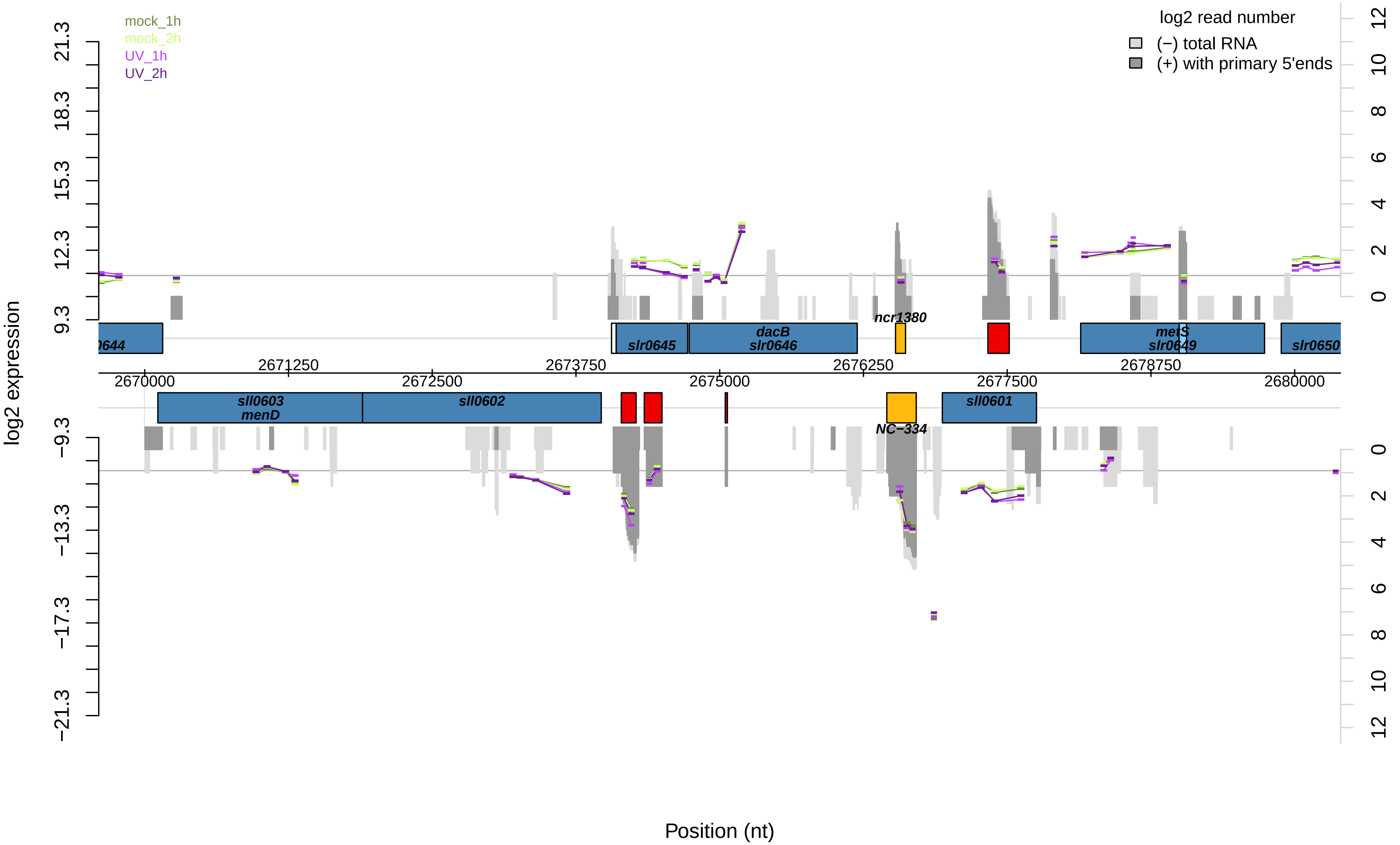

mock_1h
mock_2h
UV_1h
UV_2h
log2 read number
(−) total RNA
(+) with primary 5'ends
log2 expression
21.3
18.3
15.3
12.3
9.3
−9.3
−13.3
−17.3
−21.3
12
10
8
6
4
2
0
0644
slr0645
dacB
slr0646
ncr1380
metS
slr0649
slr0650
2671250
2673750
2676250
2678750
2670000
2672500
2675000
2677500
2680000
sll0603
menD
sll0602
NC−334
sll0601
Position (nt)

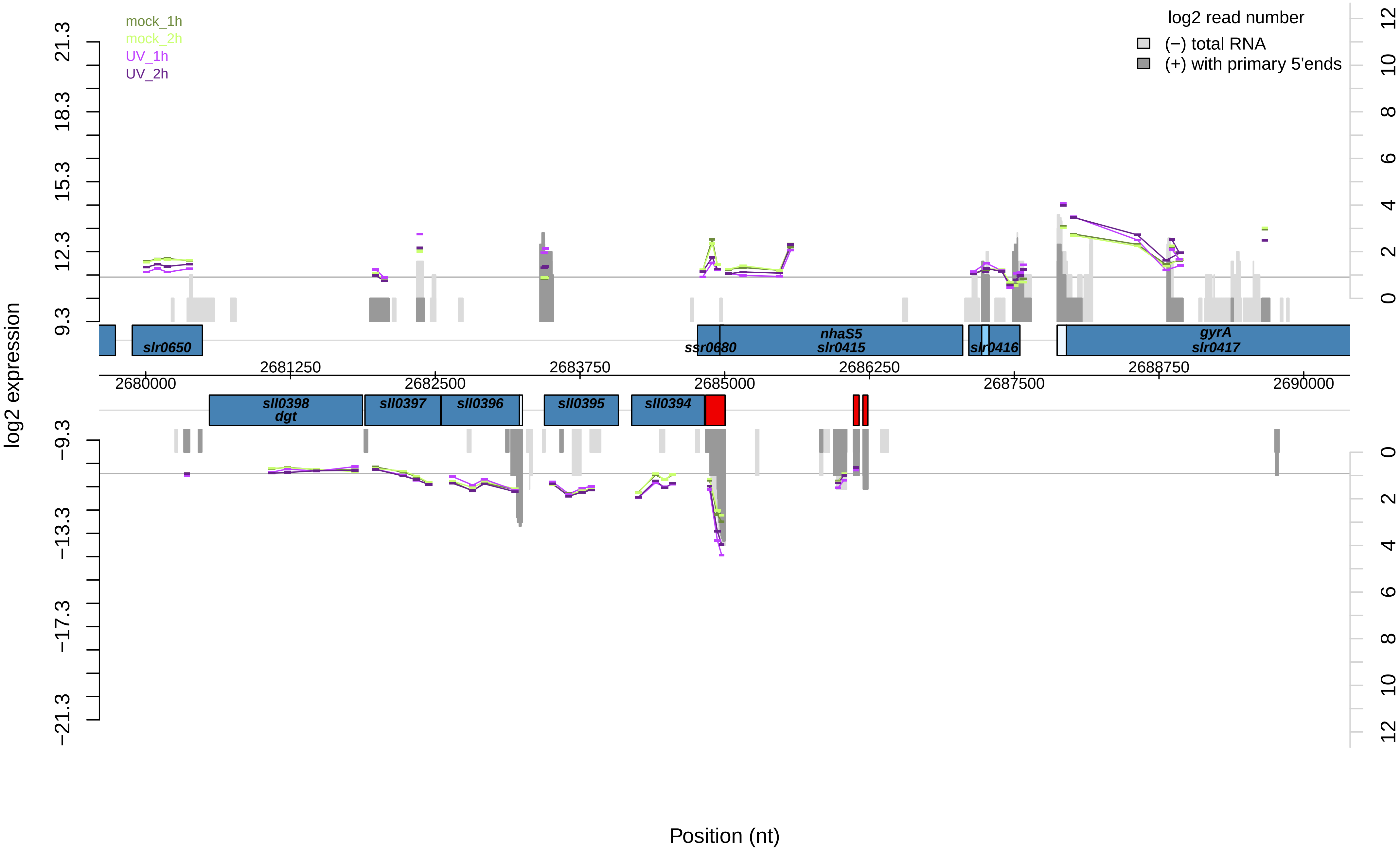

mock_1h
mock_2h
UV_1h
UV_2h
log2 read number
(−) total RNA
(+) with primary 5'ends
log2 expression
21.3
18.3
15.3
12.3
9.3
−9.3
−13.3
−17.3
−21.3
12
10
8
6
4
2
0
slr0650
ssr0680
nhaS5
slr0415
slr0416
gyrA
slr0417
2681250
2683750
2686250
2688750
2680000
2682500
2685000
2687500
2690000
sll0398
dgt
sll0397
sll0396
sll0395
sll0394
Position (nt)

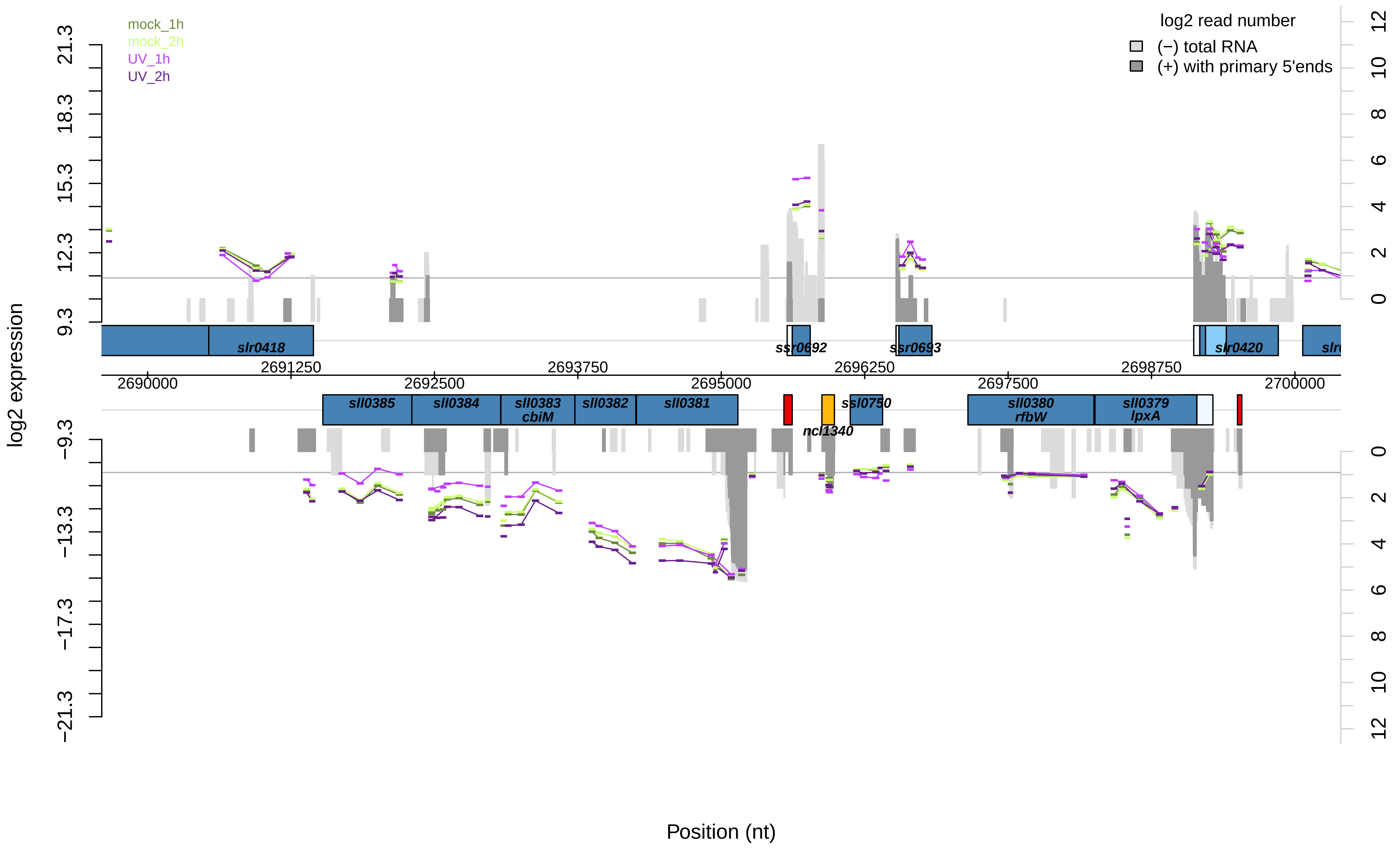

mock_1h
mock_2h
UV_1h
UV_2h
log2 read number
(−) total RNA
(+) with primary 5'ends
log2 expression
21.3
18.3
15.3
12.3
9.3
−9.3
−13.3
−17.3
−21.3
12
10
8
6
4
2
0
slr0418
ssr0692
ssr0693
slr0420
2691250
2693750
2696250
2698750
2690000
2692500
2695000
2697500
2700000
sll0385
sll0384
sll0383
cbiM
sll0382
sll0381
ncl1340
ssl0750
sll0380
rfbW
sll0379
lpxA
Position (nt)

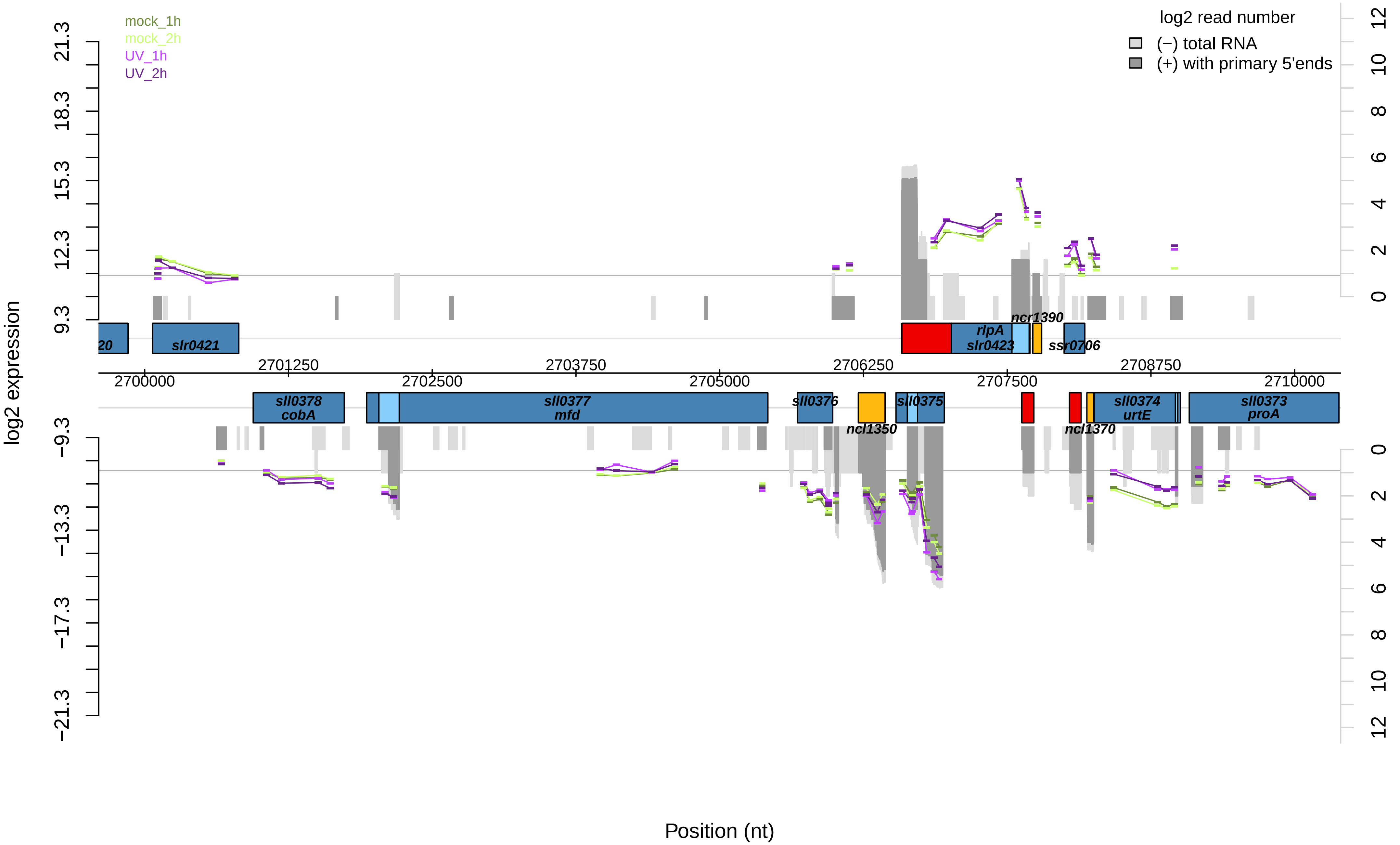

mock_1h
mock_2h
UV_1h
UV_2h
log2 read number
(−) total RNA
(+) with primary 5'ends
log2 expression
Position (nt)
21.3
18.3
15.3
12.3
9.3
−9.3
−13.3
−17.3
−21.3
12
10
8
6
4
2
0
slr0421
rlpA
slr0423
ncr1390
ssr0706
2700000
2701250
2702500
2703750
2705000
2706250
2707500
2708750
2710000
sll0378
cobA
sll0377
mfd
sll0376
ncl1350
sll0375
ncl1370
sll0374
urtE
sll0373
proA

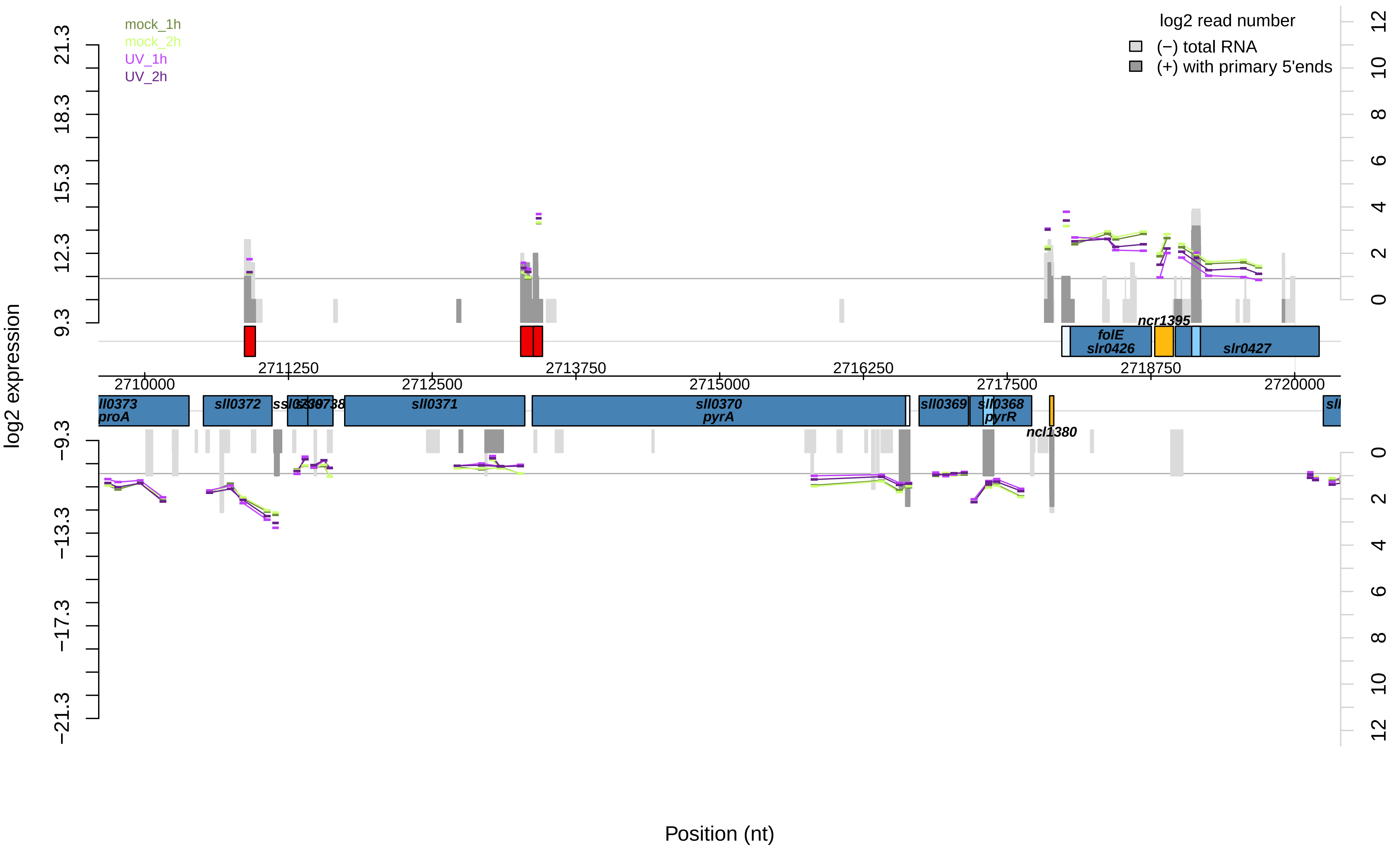
mock_1h
mock_2h
UV_1h
UV_2h
log2 read number
(−) total RNA
(+) with primary 5'ends
log2 expression
21.3
18.3
15.3
12.3
9.3
−9.3
−13.3
−17.3
−21.3
12
10
8
6
4
2
0
ncr1395
folE
slr0426
slr0427
2711250
2713750
2716250
2718750
2710000
2712500
2715000
2717500
2720000
ll0373
proA
sll0372
sll0371
sll0370
pyrA
sll0369
sll0368
pyrR
ncl1380
Position (nt)

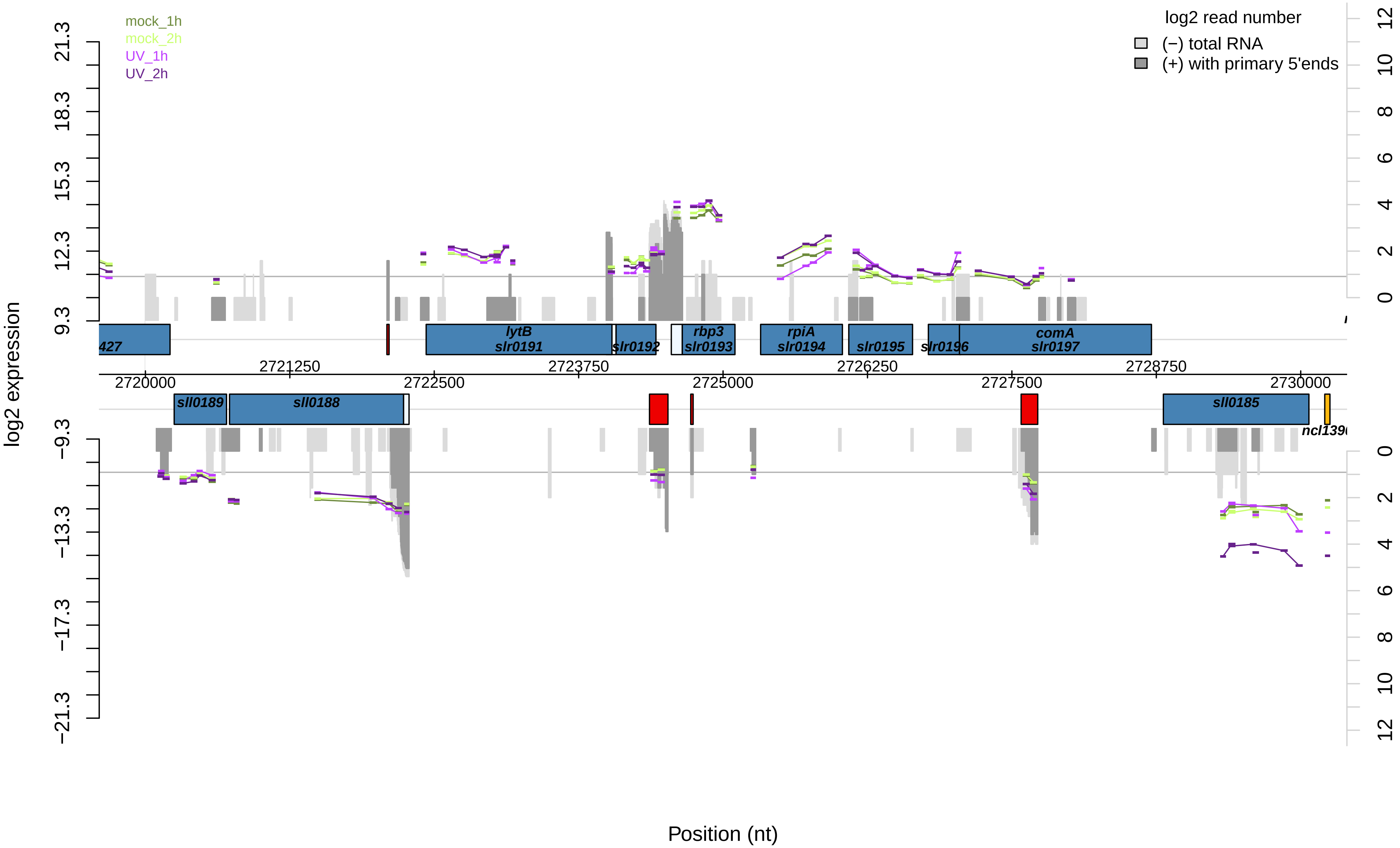

mock_1h
mock_2h
UV_1h
UV_2h
log2 read number
(−) total RNA
(+) with primary 5'ends
log2 expression
21.3
18.3
15.3
12.3
9.3
−9.3
−13.3
−17.3
−21.3
12
10
8
6
4
2
0
427
lytB
slr0191
slr0192
rbp3
slr0193
rpiA
slr0194
slr0195
slr0196
comA
slr0197
2720000
2721250
2722500
2723750
2725000
2726250
2727500
2728750
2730000
sll0189
sll0188
sll0185
Position (nt)

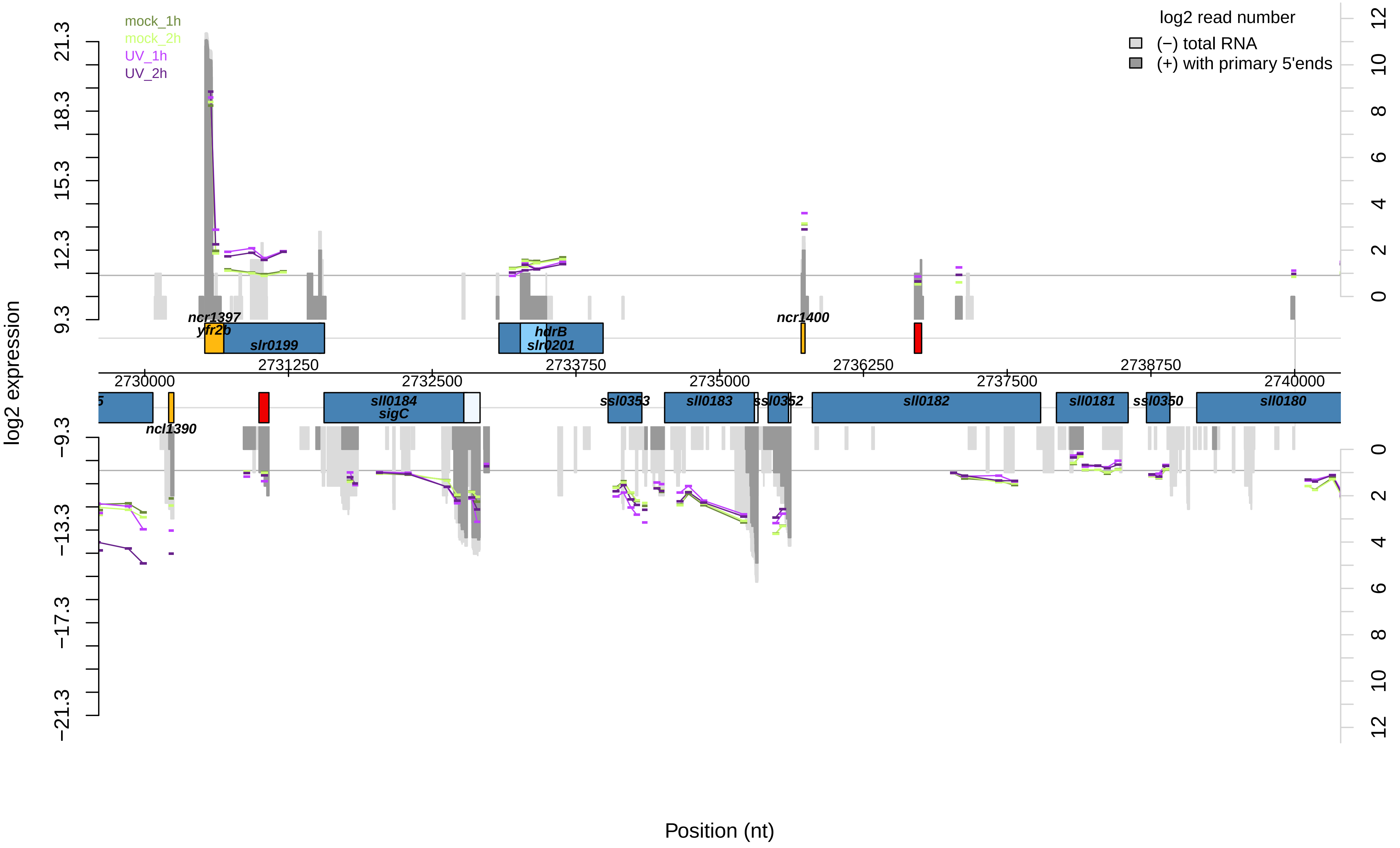

mock_1h
mock_2h
UV_1h
UV_2h
log2 read number
(−) total RNA
(+) with primary 5'ends
log2 expression
21.3
18.3
15.3
12.3
9.3
−9.3
−13.3
−17.3
−21.3
12
10
8
6
4
2
0
ncr1397
yfr2b
slr0199
hdrB
slr0201
ncr1400
2730000
2731250
2732500
2733750
2735000
2736250
2737500
2738750
2740000
ncl1390
sll0184
sigC
ssl0353
sll0183
ssl0352
sll0182
sll0181
ssl0350
sll0180
Position (nt)

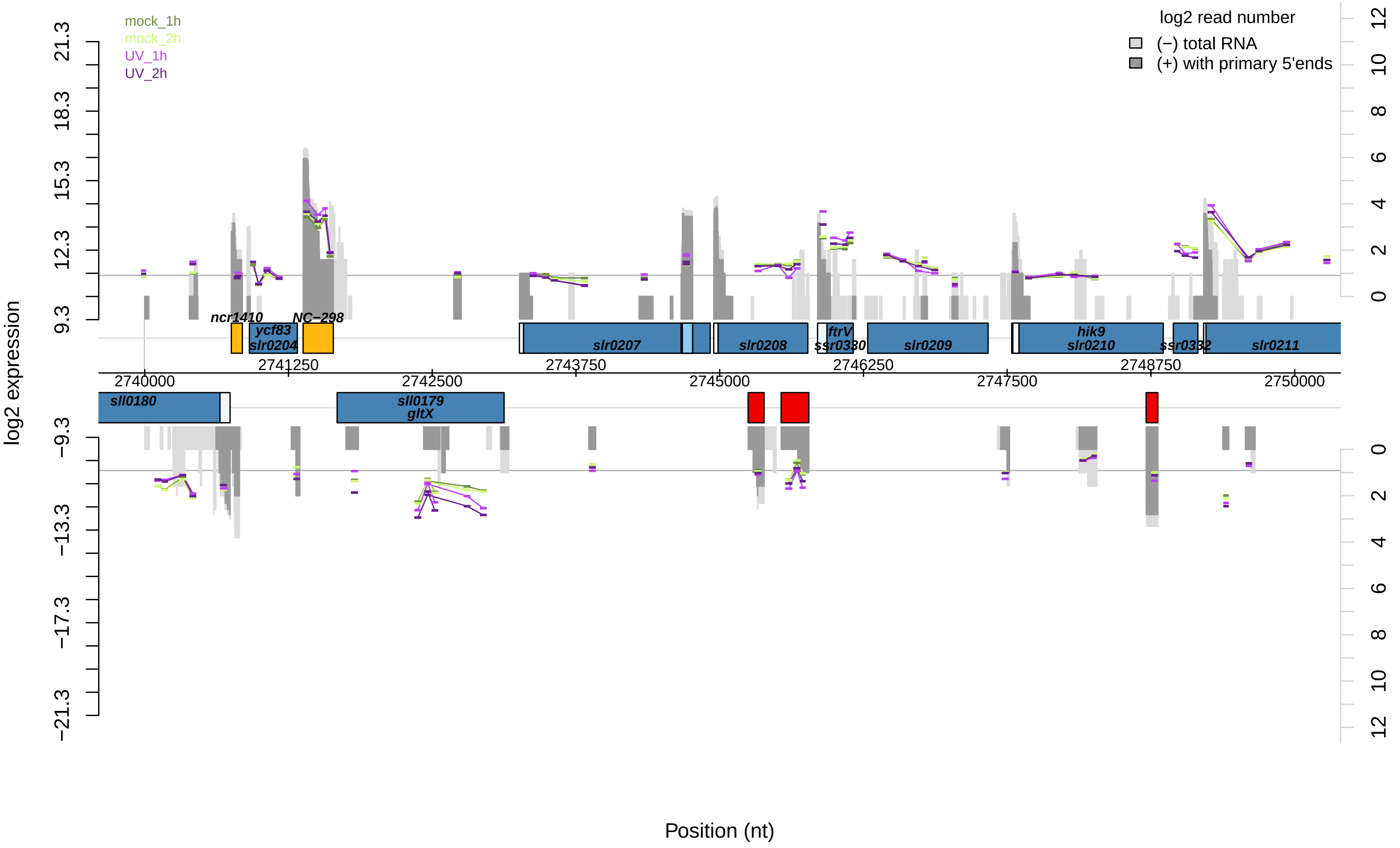

mock_1h
mock_2h
UV_1h
UV_2h
log2 read number
(−) total RNA
(+) with primary 5'ends
log2 expression
21.3
18.3
15.3
12.3
9.3
−9.3
−13.3
−17.3
−21.3
ncr1410
ycf83
slr0204
NC−298
slr0207
slr0208
ftrV
ssr0330
slr0209
hik9
slr0210
ssr0332
slr0211
sll0180
sll0179
gltX
2740000
2741250
2742500
2743750
2745000
2746250
2747500
2748750
2750000
Position (nt)

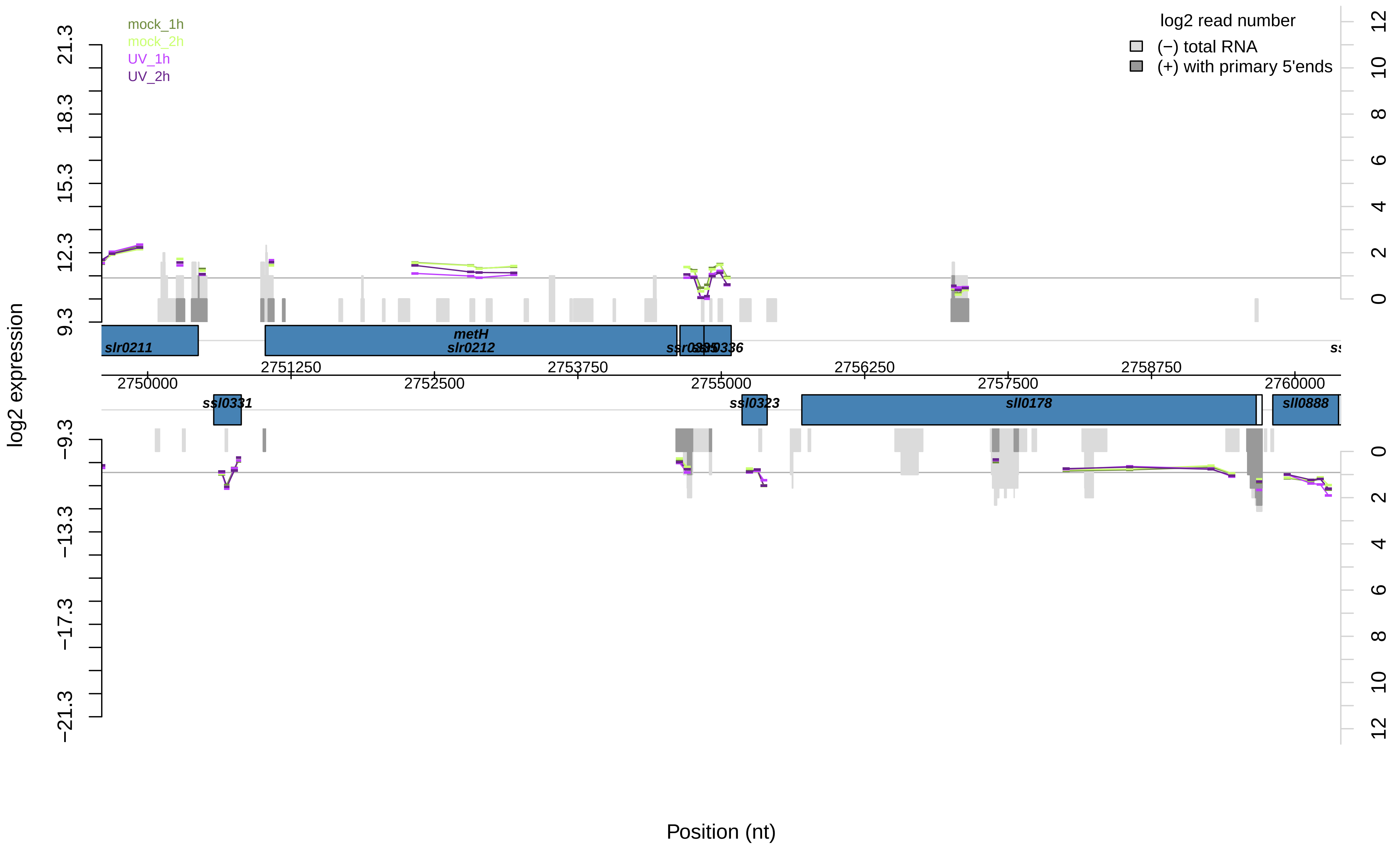

mock_1h
mock_2h
UV_1h
UV_2h
log2 read number
(−) total RNA
(+) with primary 5'ends
log2 expression
21.3
18.3
15.3
12.3
9.3
−9.3
−13.3
−17.3
−21.3
12
10
8
6
4
2
0
slr0211
metH
slr0212
ssl0331
ssl0323
sll0178
sll0888
2750000
2751250
2752500
2753750
2755000
2756250
2757500
2758750
2760000
Position (nt)

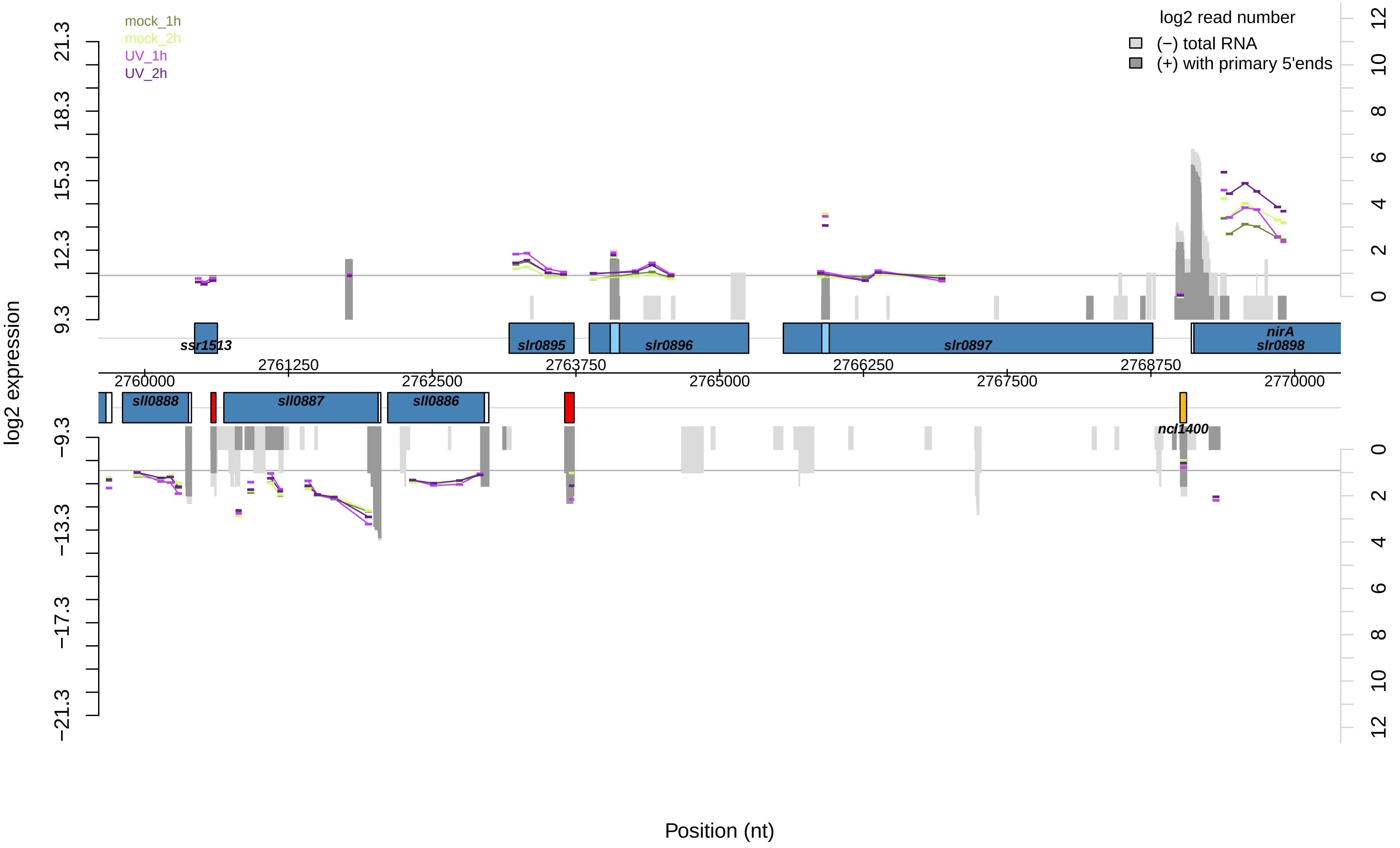

mock_1h
mock_2h
UV_1h
UV_2h
log2 read number
(−) total RNA
(+) with primary 5'ends
log2 expression
21.3
18.3
15.3
12.3
9.3
−9.3
−13.3
−17.3
−21.3
12
10
8
6
4
2
0
ssr1513
slr0895
slr0896
slr0897
nirA
slr0898
sll0888
sll0887
sll0886
ncl1400
2760000
2761250
2762500
2763750
2765000
2766250
2767500
2768750
2770000
Position (nt)

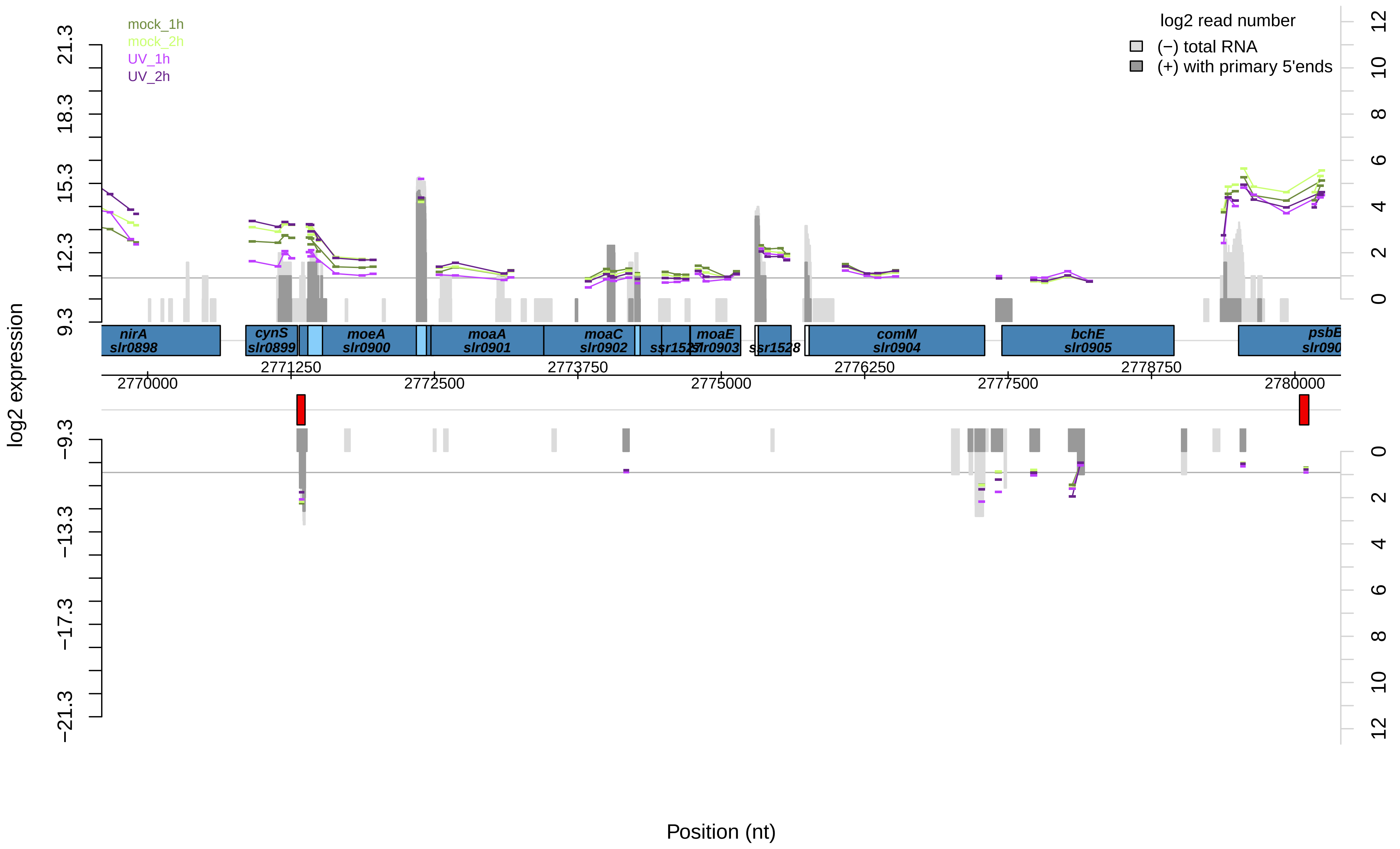

mock_1h
mock_2h
UV_1h
UV_2h
log2 read number
(−) total RNA
(+) with primary 5'ends
log2 expression
21.3
18.3
15.3
12.3
9.3
−9.3
−13.3
−17.3
−21.3
12
10
8
6
4
2
0
nirA
slr0898
cynS
slr0899
moeA
slr0900
moaA
slr0901
moaC
slr0902
ssr1527
moaE
slr0903
ssr1528
comM
slr0904
bchE
slr0905
psbE
slr090
2770000
2771250
2772500
2773750
2775000
2776250
2777500
2778750
2780000
Position (nt)

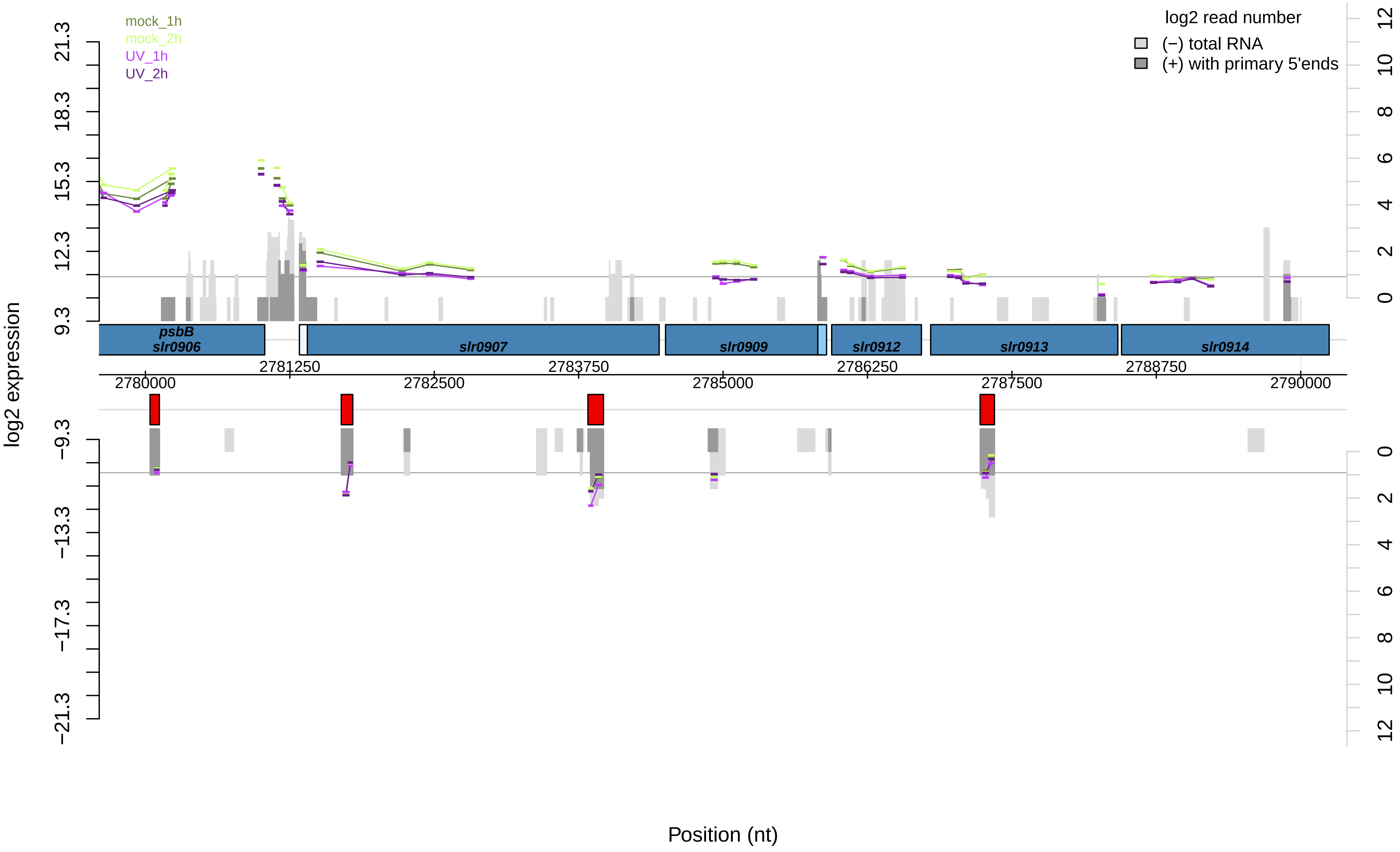

mock_1h
mock_2h
UV_1h
UV_2h
log2 read number
(−) total RNA
(+) with primary 5'ends
log2 expression
21.3
18.3
15.3
12.3
9.3
−9.3
−13.3
−17.3
−21.3
12
10
8
6
4
2
0
psbB
slr0906
slr0907
slr0909
slr0912
slr0913
slr0914
2780000
2781250
2782500
2783750
2785000
2786250
2787500
2788750
2790000
Position (nt)

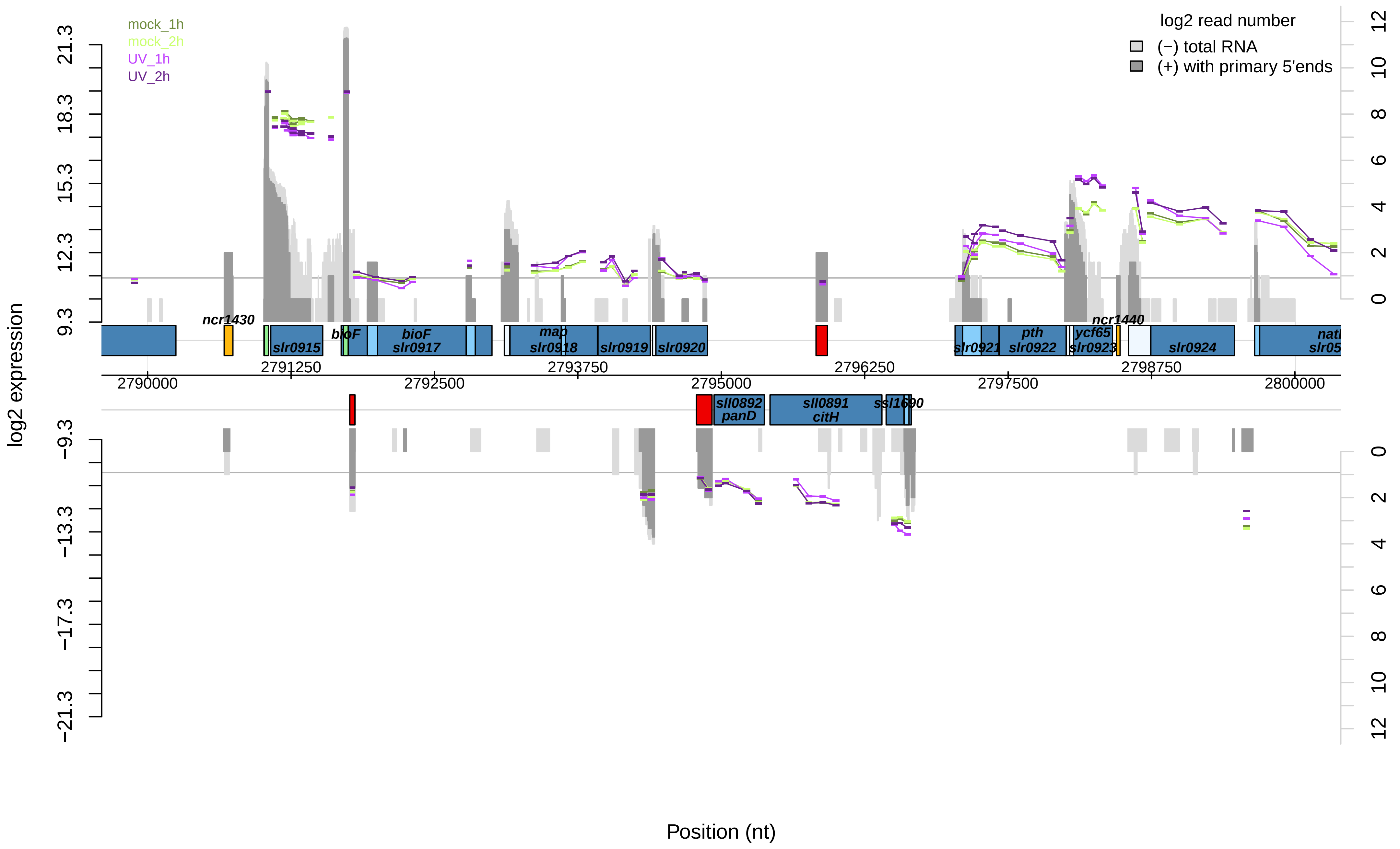

mock_1h
mock_2h
UV_1h
UV_2h
log2 read number
(−) total RNA
(+) with primary 5'ends
log2 expression
Position (nt)
ncr1430
slr0915
bioF
bioF
slr0917
map
slr0918
slr0919
slr0920
slr0921
pth
slr0922
ycf65
slr0923
ncr1440
slr0924
sll0892
panD
sll0891
citH
ssl1690
2790000
2791250
2792500
2793750
2795000
2796250
2797500
2798750
2800000

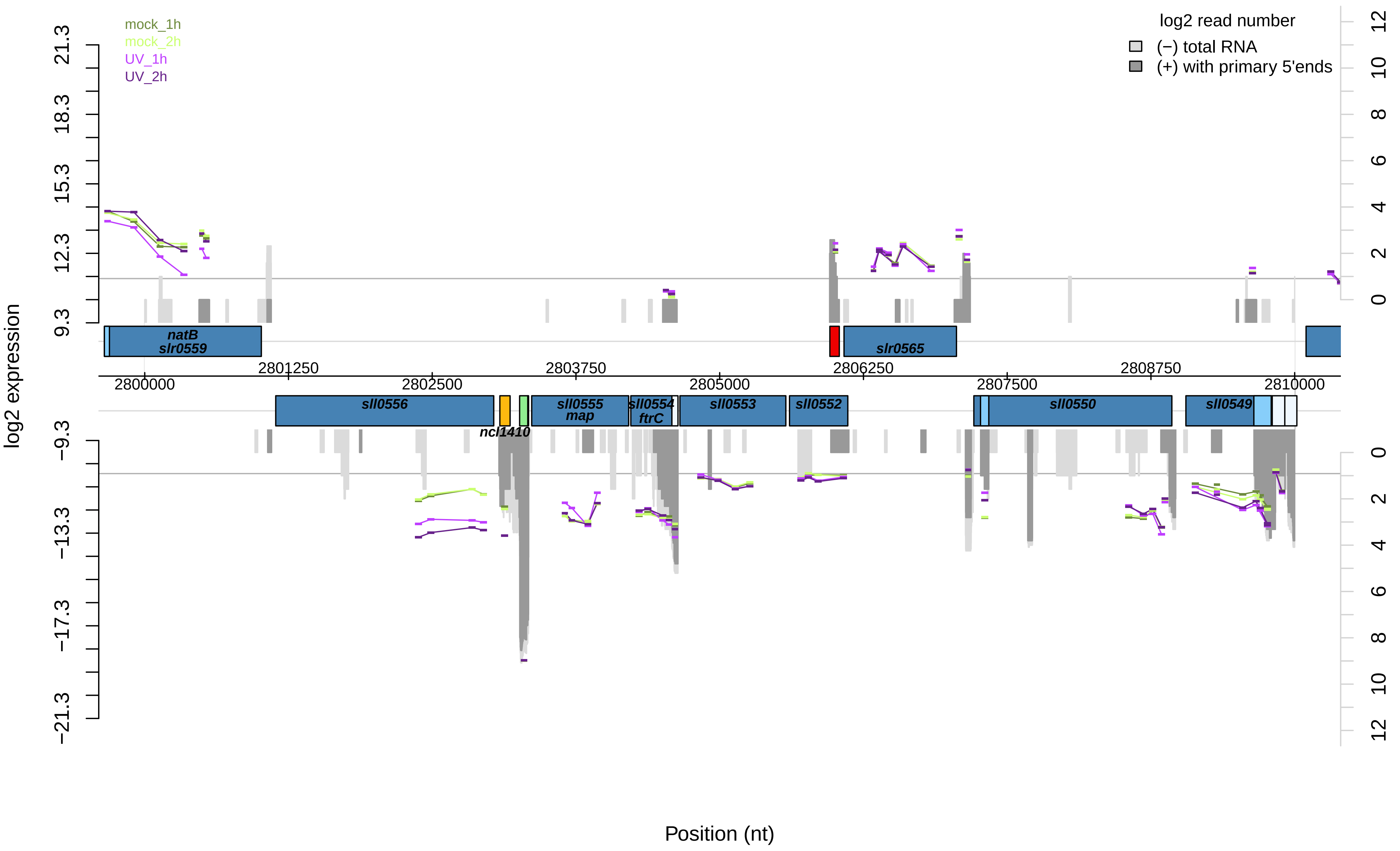

mock_1h
mock_2h
UV_1h
UV_2h
log2 read number
(−) total RNA
(+) with primary 5'ends
log2 expression
21.3
18.3
15.3
12.3
9.3
−9.3
−13.3
−17.3
−21.3
12
10
8
6
4
2
0
natB
slr0559
slr0565
sll0556
ncl1410
sll0555
map
sll0554
ftrC
sll0553
sll0552
sll0550
sll0549
2800000
2801250
2802500
2803750
2805000
2806250
2807500
2808750
2810000
Position (nt)

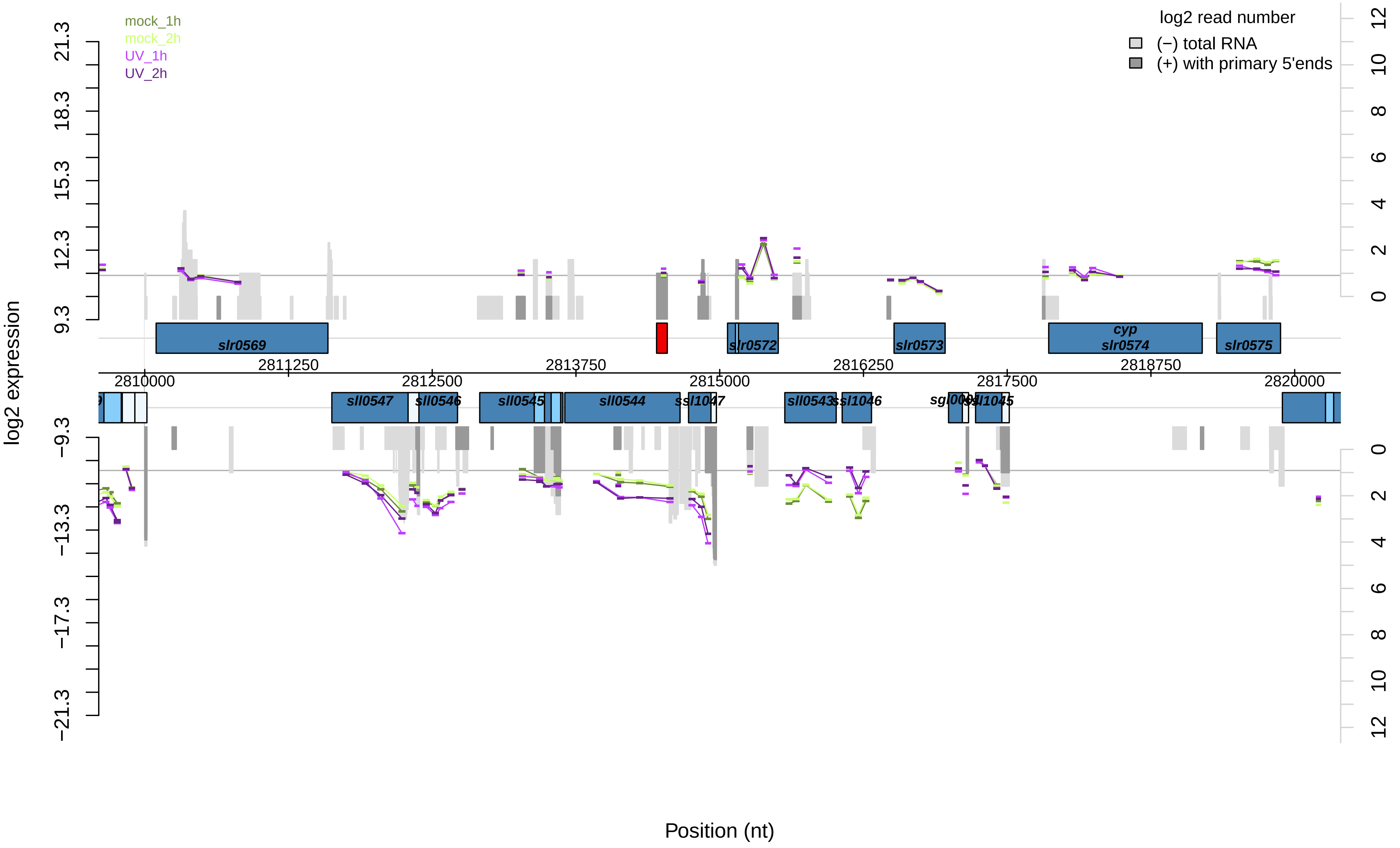

mock_1h
mock_2h
UV_1h
UV_2h
log2 read number
(−) total RNA
(+) with primary 5'ends
log2 expression
21.3
18.3
15.3
12.3
9.3
−9.3
−13.3
−17.3
−21.3
12
10
8
6
4
2
0
slr0569
slr0572
slr0573
cyp
slr0574
slr0575
2811250
2813750
2816250
2818750
2810000
2812500
2815000
2817500
2820000
sll0547
sll0546
sll0545
sll0544
ssl1047
sll0543
ssl1046
ssl1045
Position (nt)

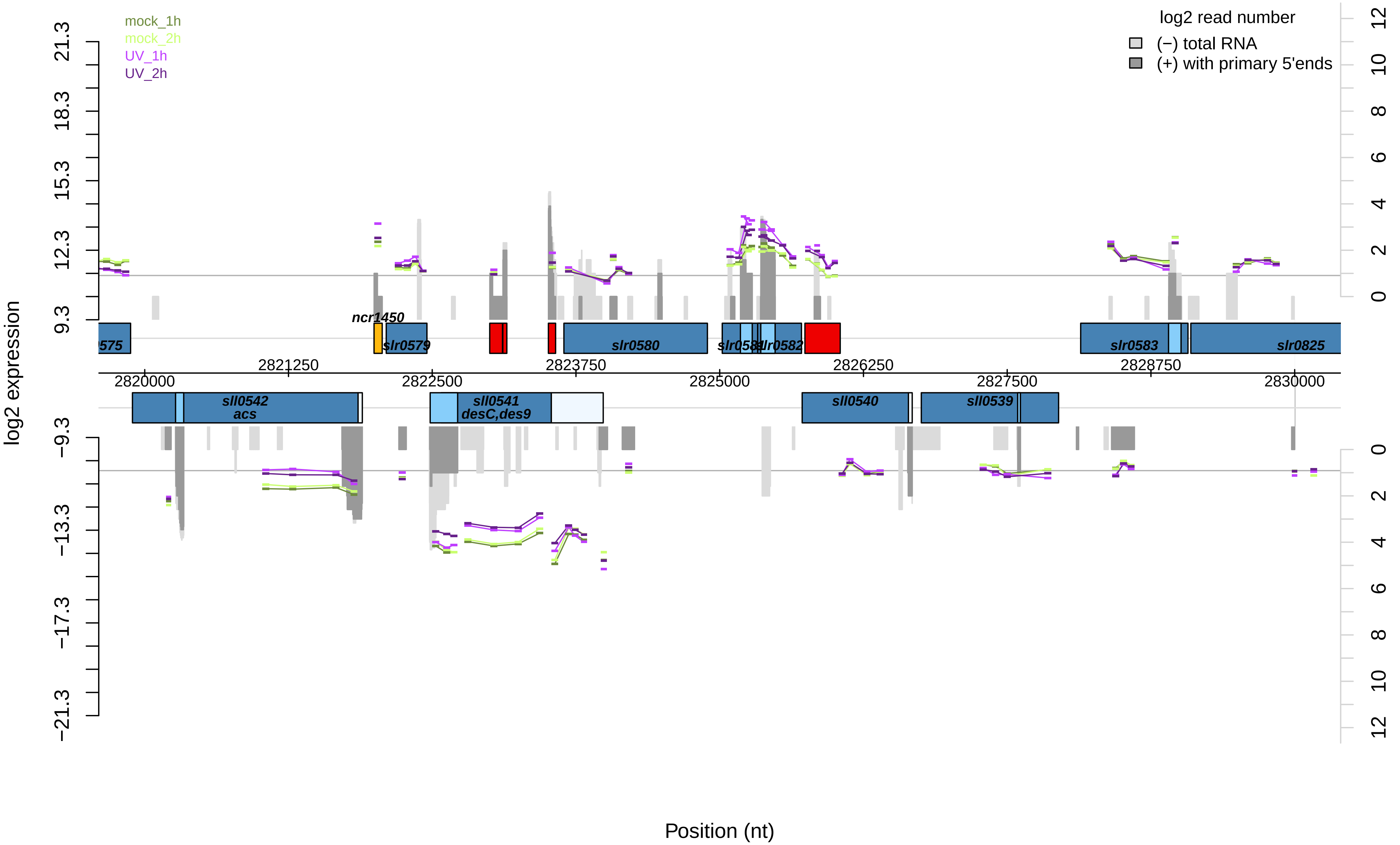

mock_1h
mock_2h
UV_1h
UV_2h
log2 read number
(−) total RNA
(+) with primary 5'ends
log2 expression
21.3
18.3
15.3
12.3
9.3
−9.3
−13.3
−17.3
−21.3
12
10
8
6
4
2
0
ncr1450
575
slr0579
slr0580
slr0581
slr0582
slr0583
slr0825
2821250
2823750
2826250
2828750
2820000
2822500
2825000
2827500
2830000
sll0542
acs
sll0541
desC,des9
sll0540
sll0539
Position (nt)

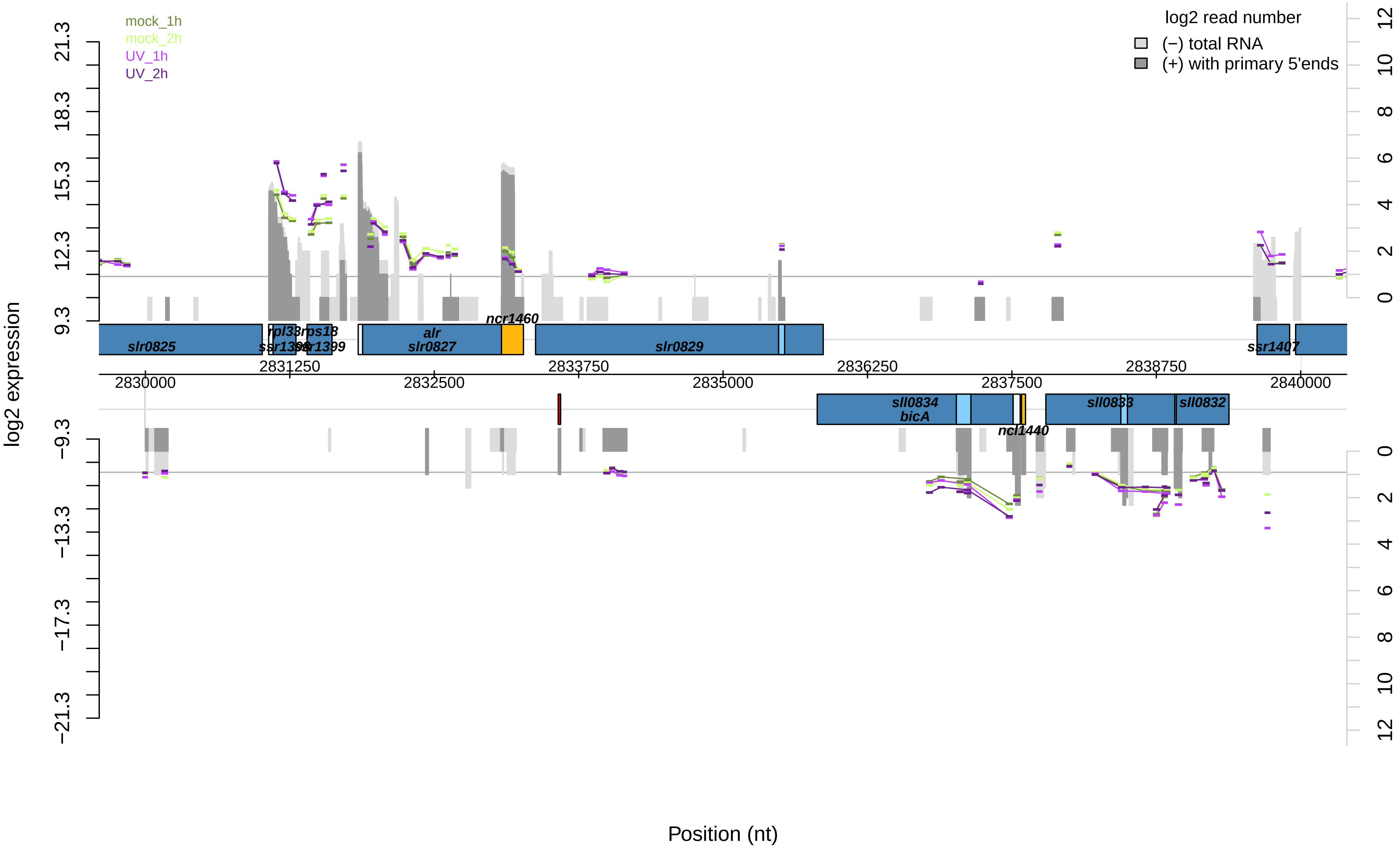

mock_1h
mock_2h
UV_1h
UV_2h
log2 read number
(−) total RNA
(+) with primary 5'ends
log2 expression
Position (nt)
21.3
18.3
15.3
12.3
9.3
−9.3
−13.3
−17.3
−21.3
12
10
8
6
4
2
0
slr0825
rpl33
rps18
ssr1398
ssr1399
alr
slr0827
ncr1460
slr0829
ssr1407
sll0834
bicA
ncl1440
sll0833
sll0832
2830000
2831250
2832500
2833750
2835000
2836250
2837500
2838750
2840000

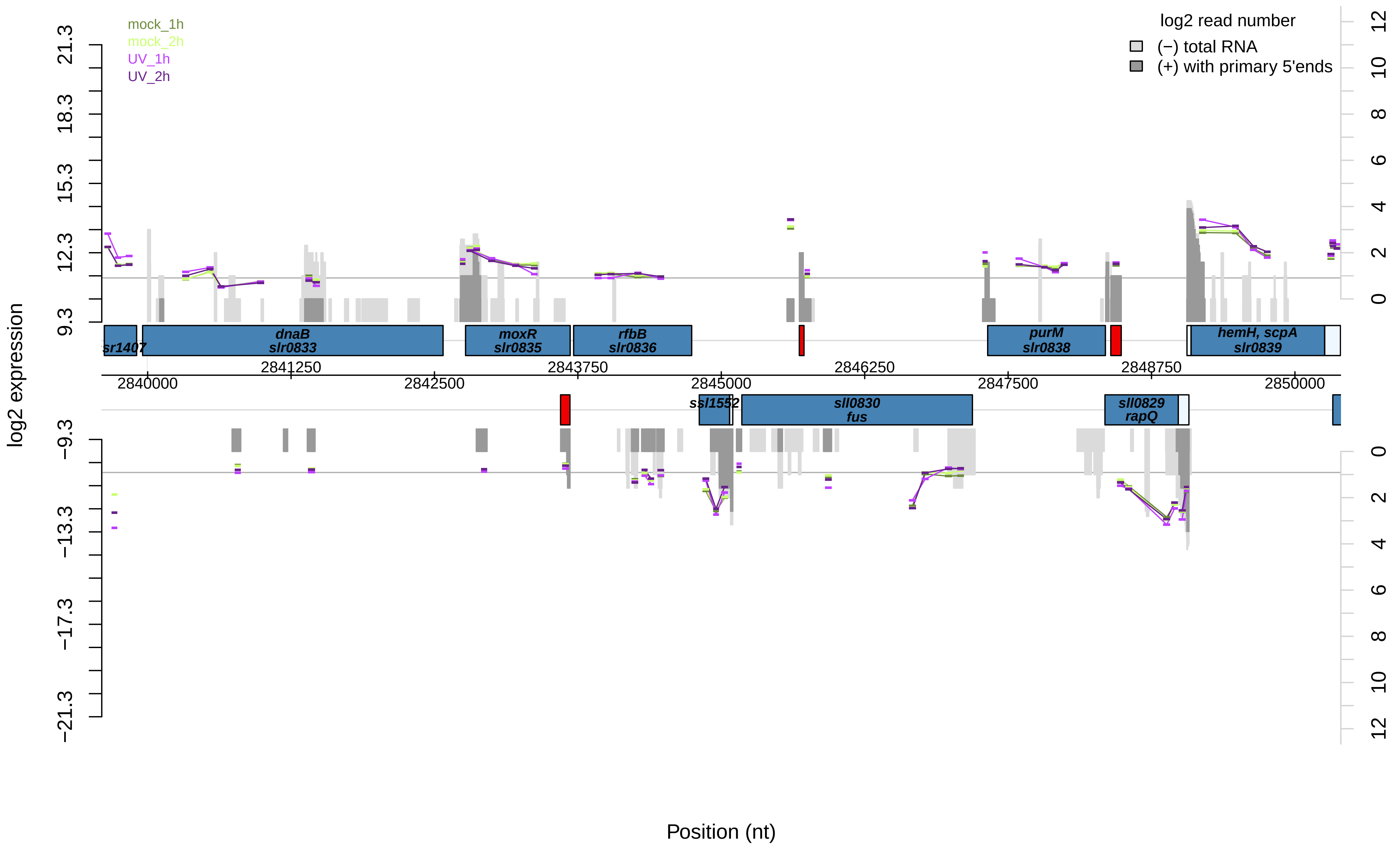

mock_1h
mock_2h
UV_1h
UV_2h
log2 read number
(−) total RNA
(+) with primary 5'ends
log2 expression
Position (nt)
dnaB
slr0833
moxR
slr0835
rfbB
slr0836
purM
slr0838
hemH, scpA
slr0839
ssl1552
sll0830
fus
sll0829
rapQ
2840000
2841250
2842500
2843750
2845000
2846250
2847500
2848750
2850000

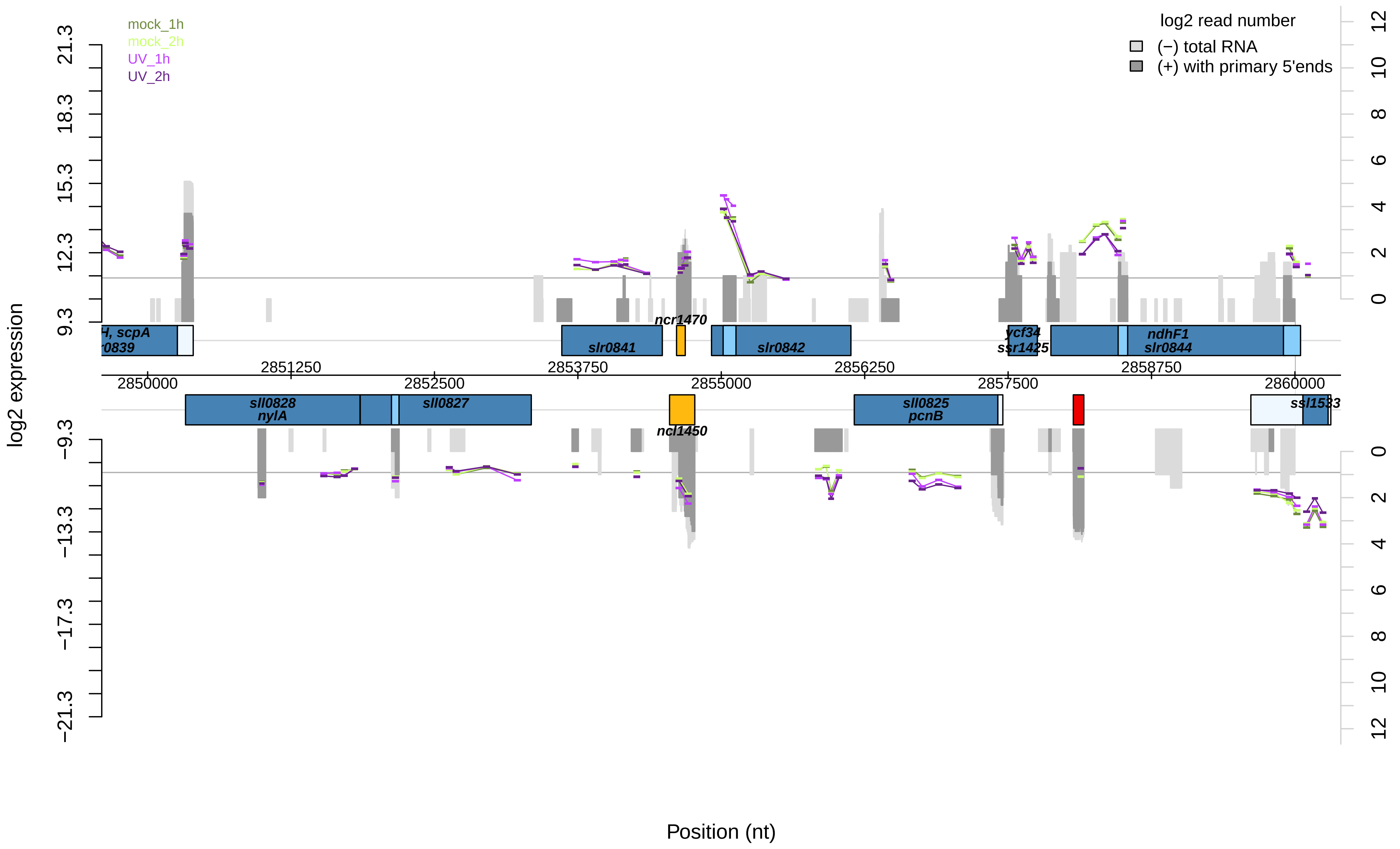

mock_1h
mock_2h
UV_1h
UV_2h
log2 read number
(−) total RNA
(+) with primary 5'ends
log2 expression
21.3
18.3
15.3
12.3
9.3
−9.3
−13.3
−17.3
−21.3
12
10
8
6
4
2
0
H, scpA
r0839
slr0841
ncr1470
slr0842
ycf34
ssr1425
ndhF1
slr0844
2851250
2853750
2856250
2858750
2850000
2852500
2855000
2857500
2860000
sll0828
nylA
sll0827
ncl1450
sll0825
pcnB
ssl1533
Position (nt)

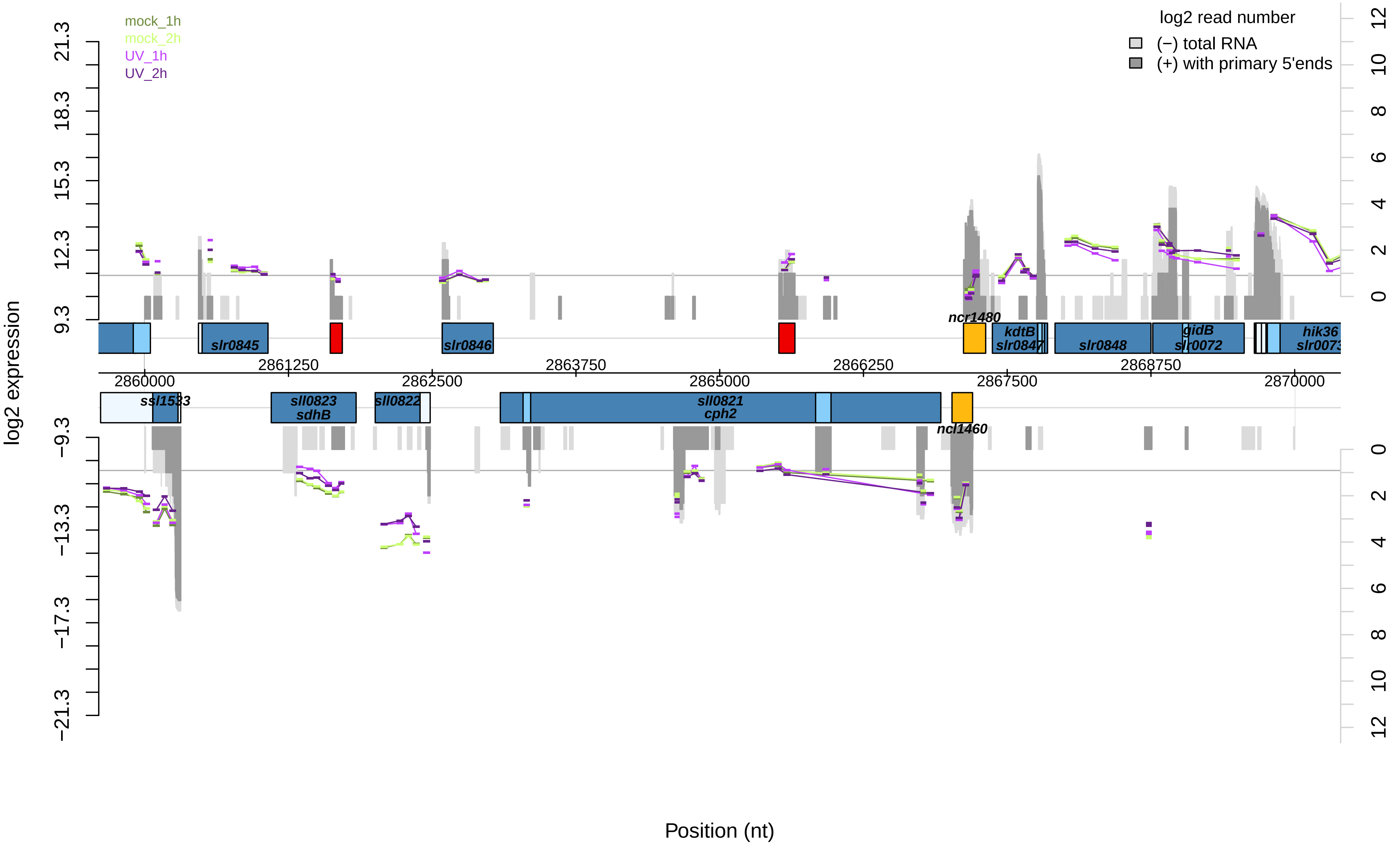

mock_1h
mock_2h
UV_1h
UV_2h
log2 read number
(−) total RNA
(+) with primary 5'ends
log2 expression
Position (nt)
21.3
18.3
15.3
12.3
9.3
−9.3
−13.3
−17.3
−21.3
12
10
8
6
4
2
0
2860000
2861250
2862500
2863750
2865000
2866250
2867500
2868750
2870000
slr0845
slr0846
ncr1480
kdtB
slr0847
slr0848
gidB
slr0072
hik36
ssl1533
sll0823
sdhB
sll0822
sll0821
cph2
ncl1460

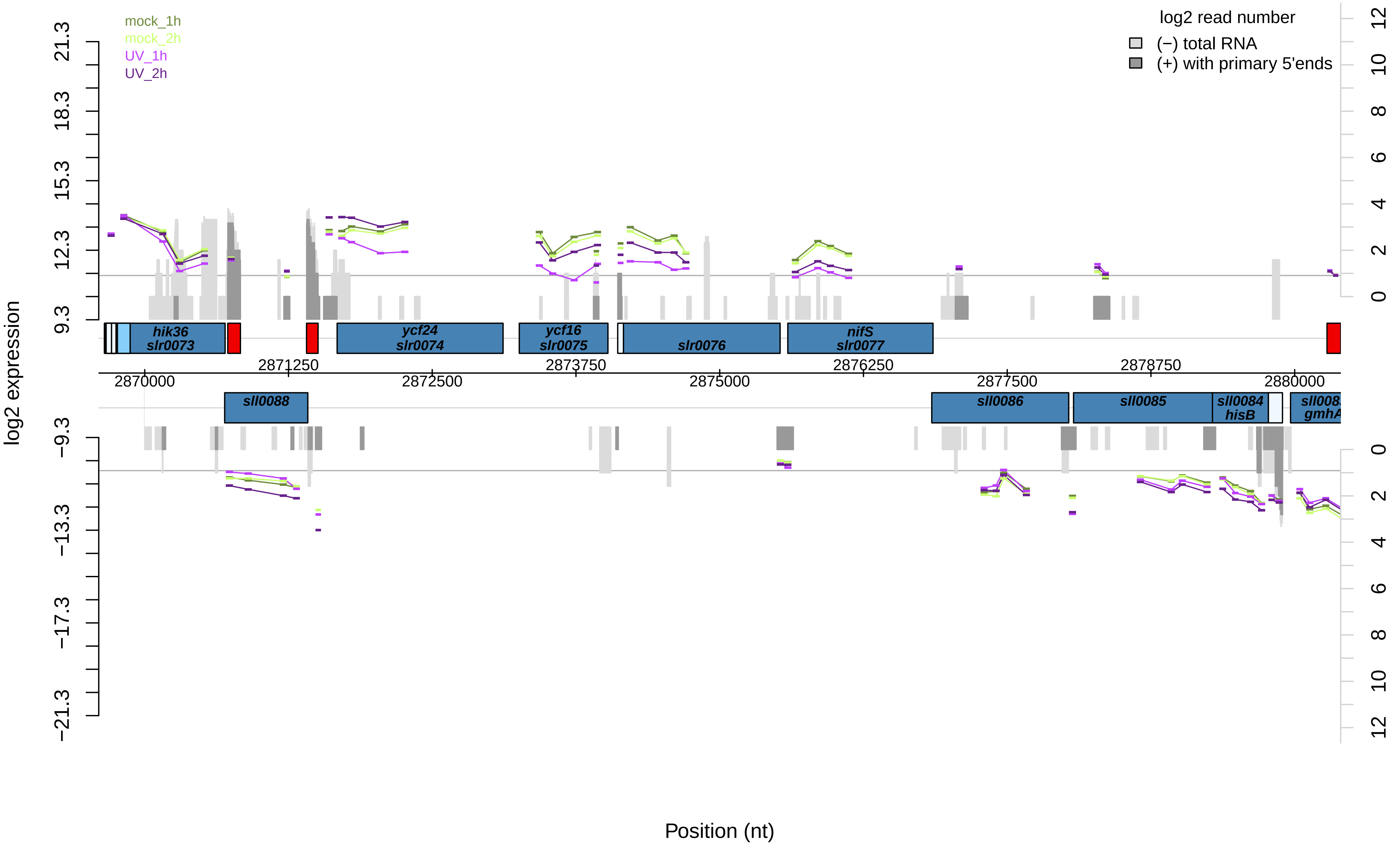

mock_1h
mock_2h
UV_1h
UV_2h
log2 read number
(−) total RNA
(+) with primary 5'ends
log2 expression
21.3
18.3
15.3
12.3
9.3
−9.3
−13.3
−17.3
−21.3
12
10
8
6
4
2
0
hik36
slr0073
ycf24
slr0074
ycf16
slr0075
slr0076
nifS
slr0077
2871250
2873750
2876250
2878750
2870000
2872500
2875000
2877500
2880000
sll0088
sll0086
sll0085
sll0084
hisB
Position (nt)

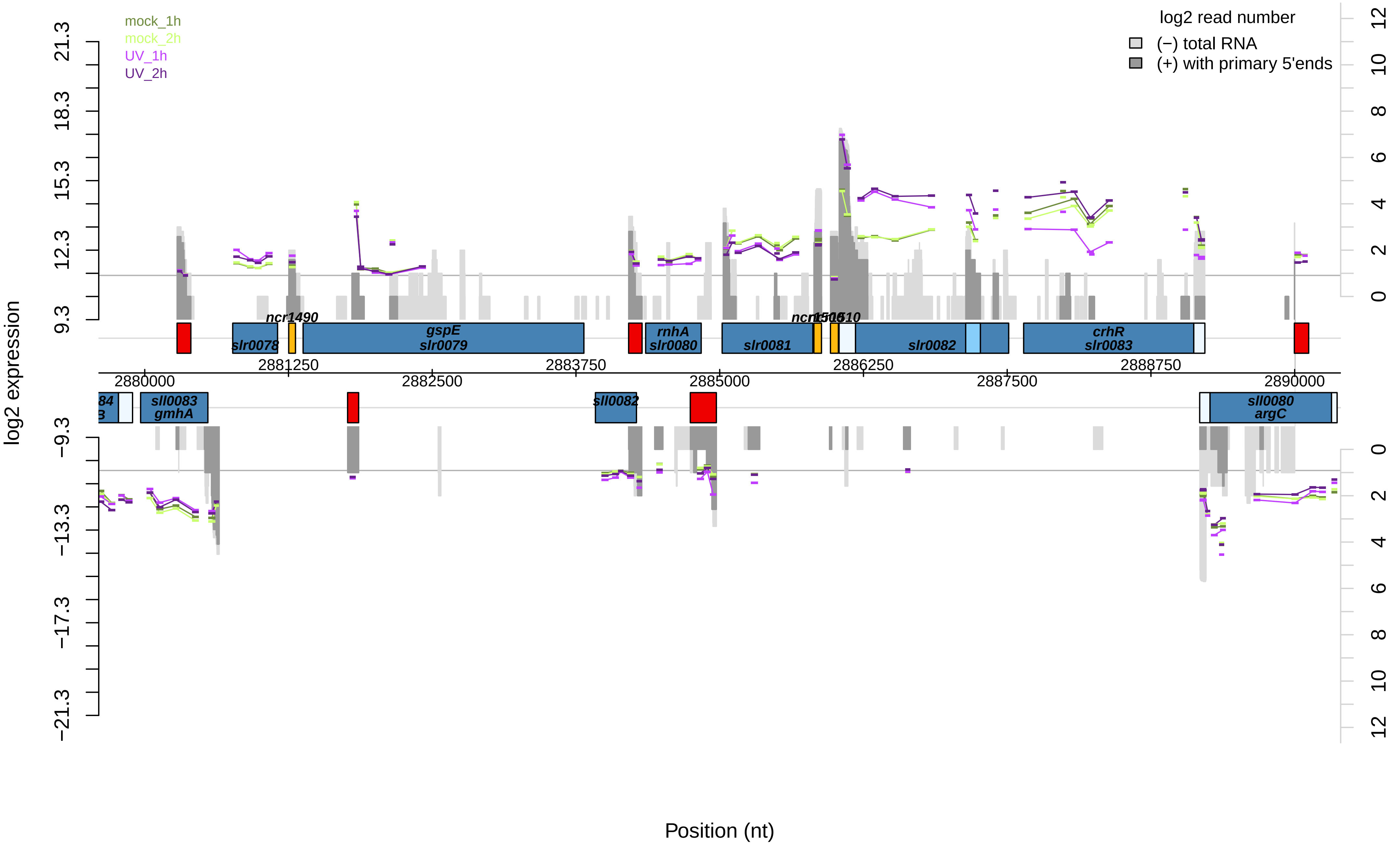
mock_1h
mock_2h
UV_1h
UV_2h
log2 read number
(−) total RNA
(+) with primary 5'ends
log2 expression
Position (nt)
ncr1490
slr0078
gspE
slr0079
rnhA
slr0080
slr0081
slr0082
crhR
slr0083
sll0083
gmhA
sll0082
sll0080
argC
2880000
2881250
2882500
2883750
2885000
2886250
2887500
2888750
2890000

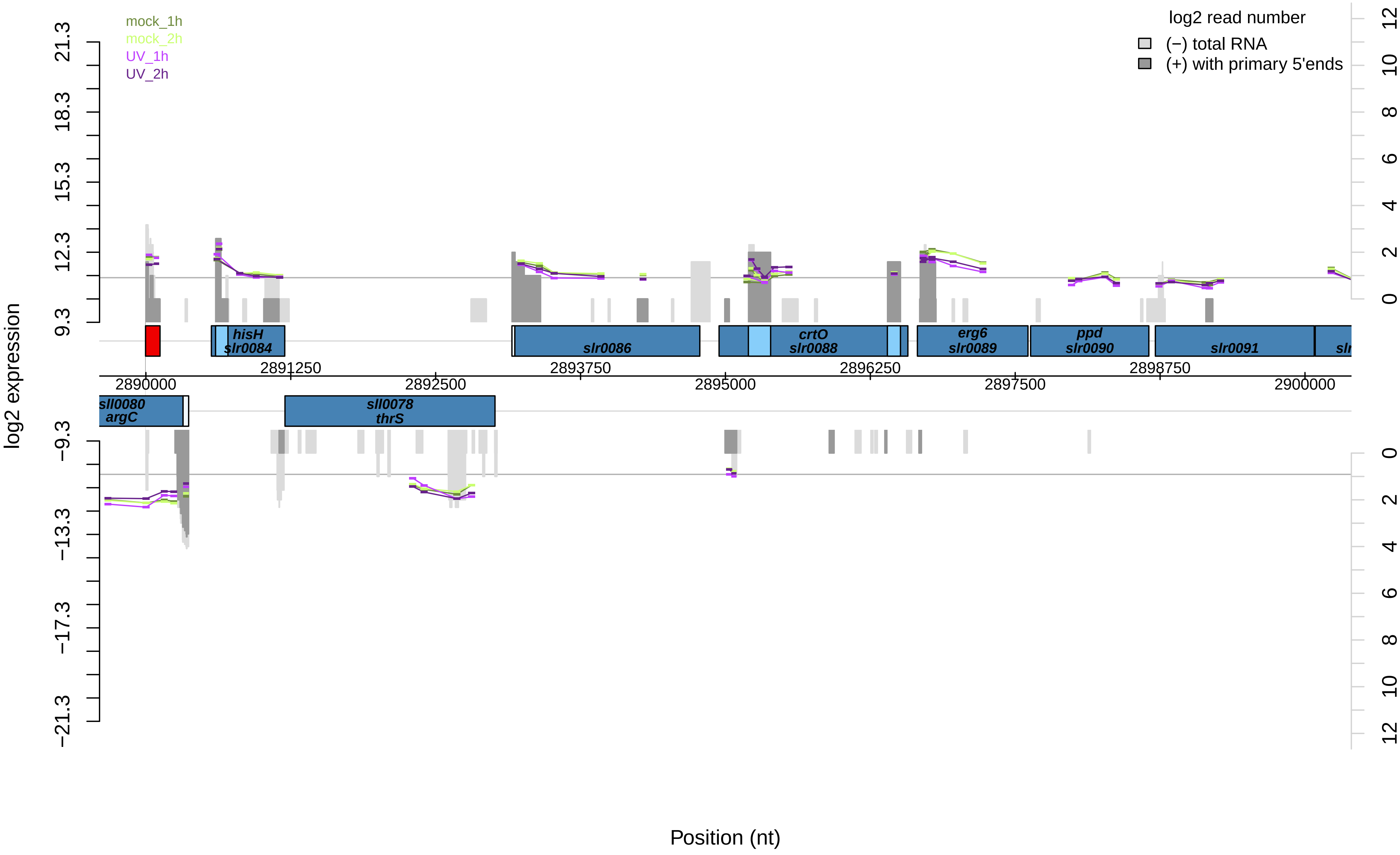

mock_1h
mock_2h
UV_1h
UV_2h
log2 read number
(−) total RNA
(+) with primary 5'ends
log2 expression
21.3
18.3
15.3
12.3
9.3
−9.3
−13.3
−17.3
−21.3
hisH
slr0084
slr0086
crtO
slr0088
erg6
slr0089
ppd
slr0090
slr0091
sll0080
argC
sll0078
thrS
2890000
2891250
2892500
2893750
2895000
2896250
2897500
2898750
2900000
Position (nt)

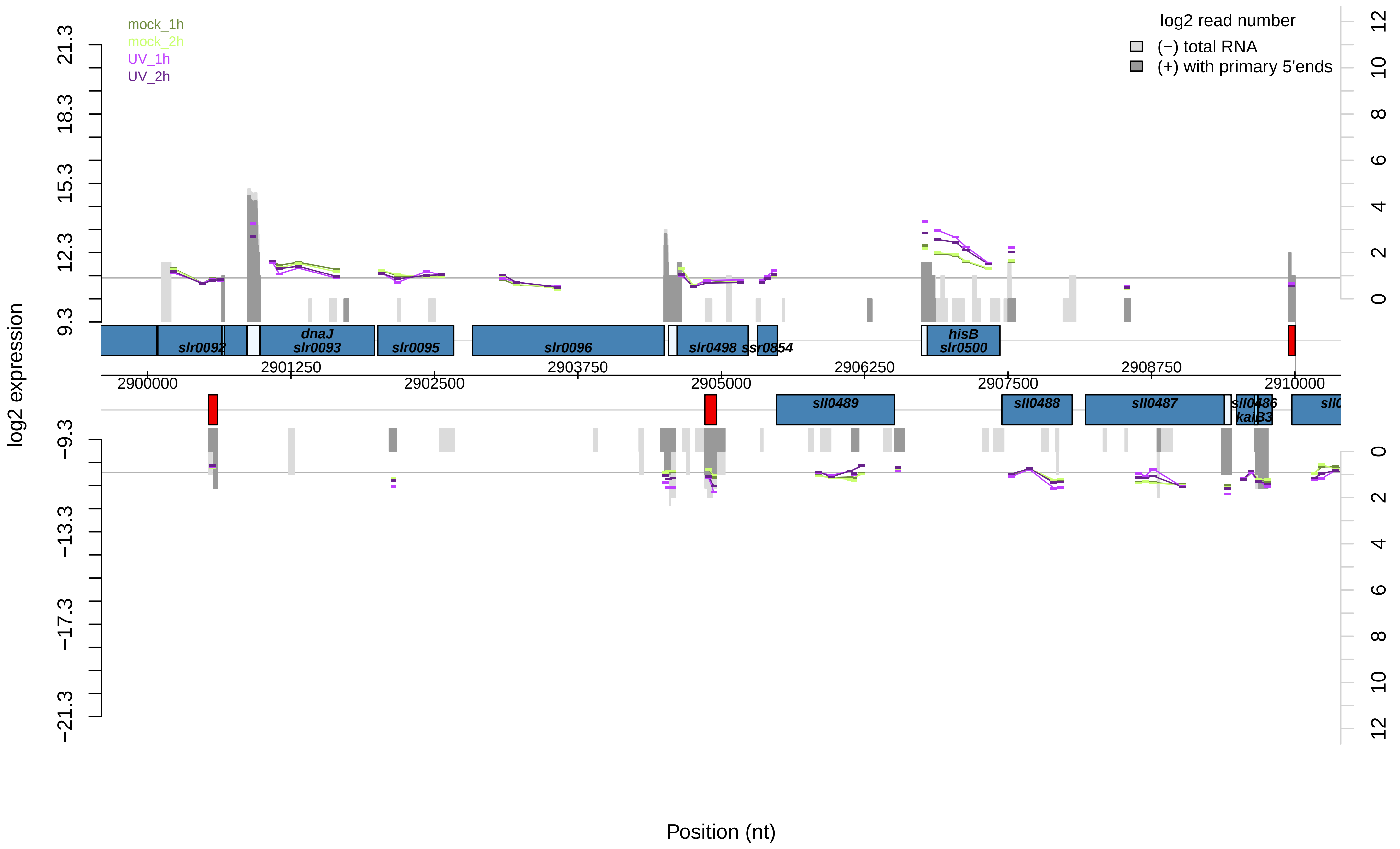

mock_1h
mock_2h
UV_1h
UV_2h
log2 read number
(−) total RNA
(+) with primary 5'ends
log2 expression
21.3
18.3
15.3
12.3
9.3
−9.3
−13.3
−17.3
−21.3
12
10
8
6
4
2
0
slr0092
dnaJ
slr0093
slr0095
slr0096
slr0498
ssr0854
hisB
slr0500
2900000
2901250
2902500
2903750
2905000
2906250
2907500
2908750
2910000
sll0489
sll0488
sll0487
sll0486
kaiB3
Position (nt)

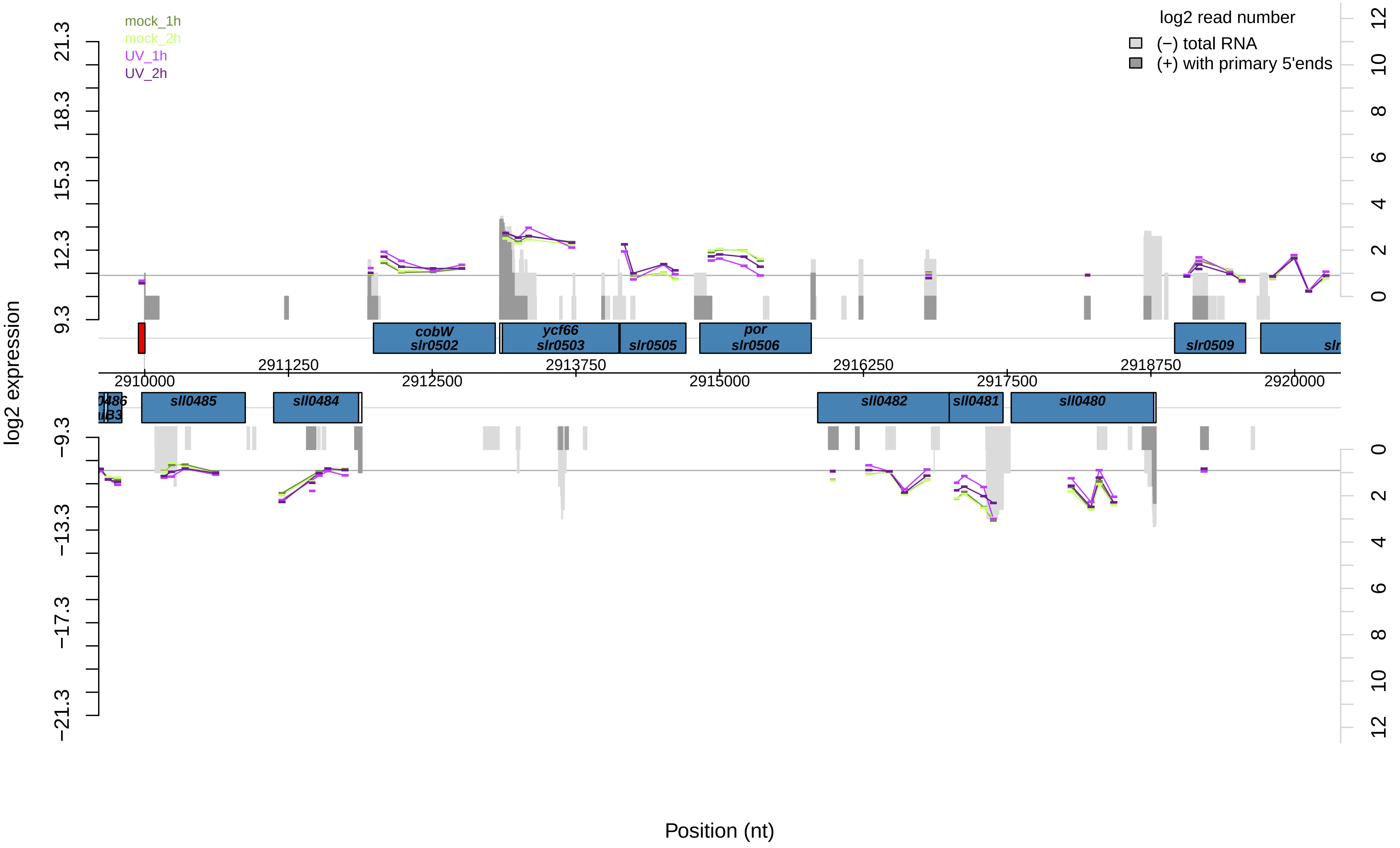

mock_1h
mock_2h
UV_1h
UV_2h
log2 read number
(−) total RNA
(+) with primary 5'ends
log2 expression
21.3
18.3
15.3
12.3
9.3
−9.3
−13.3
−17.3
−21.3
12
10
8
6
4
2
0
cobW
slr0502
ycf66
slr0503
slr0505
por
slr0506
slr0509
2911250
2913750
2916250
2918750
2910000
2912500
2915000
2917500
2920000
sll0485
sll0484
sll0482
sll0481
sll0480
Position (nt)

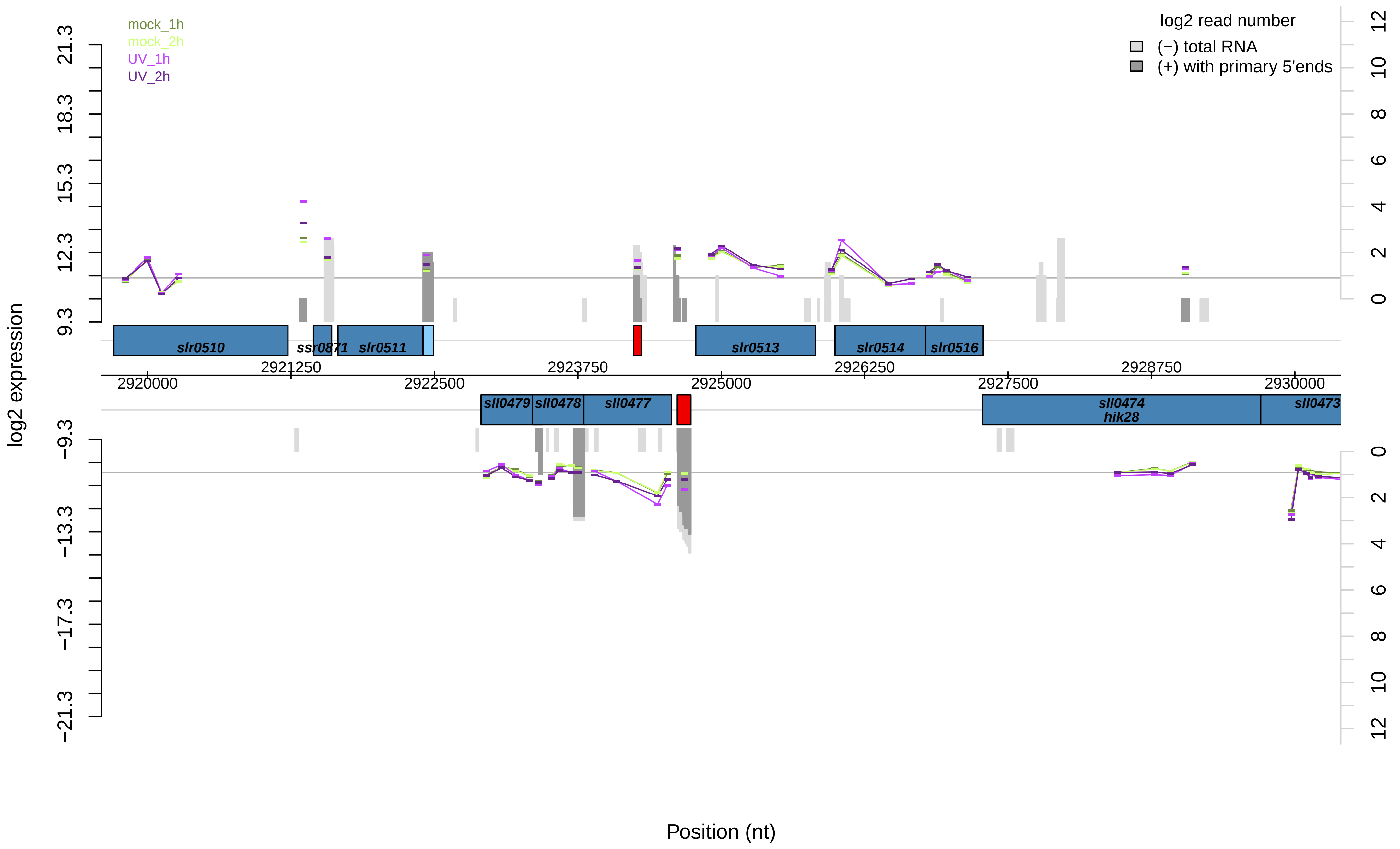

mock_1h
mock_2h
UV_1h
UV_2h
log2 read number
(−) total RNA
(+) with primary 5'ends
log2 expression
21.3
18.3
15.3
12.3
9.3
−9.3
−13.3
−17.3
−21.3
12
10
8
6
4
2
0
slr0510
ssr0871
slr0511
slr0513
slr0514
slr0516
2921250
2923750
2926250
2928750
2920000
2922500
2925000
2927500
2930000
sll0479
sll0478
sll0477
sll0474
hik28
sll0473
Position (nt)

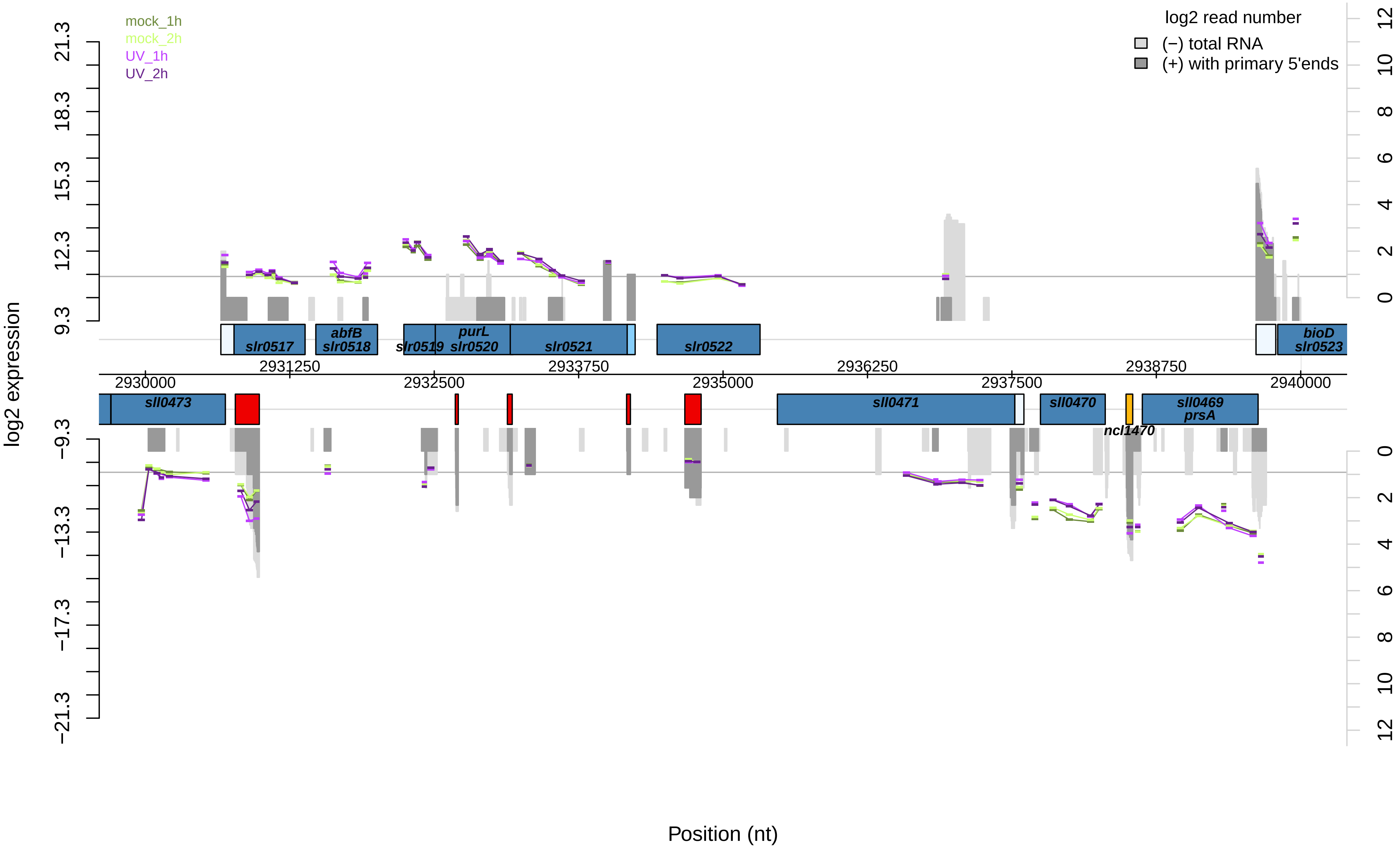

mock_1h
mock_2h
UV_1h
UV_2h
log2 read number
(−) total RNA
(+) with primary 5'ends
log2 expression
21.3
18.3
15.3
12.3
9.3
−9.3
−13.3
−17.3
−21.3
12
10
8
6
4
2
0
slr0517
abfB
slr0518
slr0519
purL
slr0520
slr0521
slr0522
bioD
slr0523
2930000
2931250
2932500
2933750
2935000
2936250
2937500
2938750
2940000
sll0473
sll0471
sll0470
ncl1470
sll0469
prsA
Position (nt)

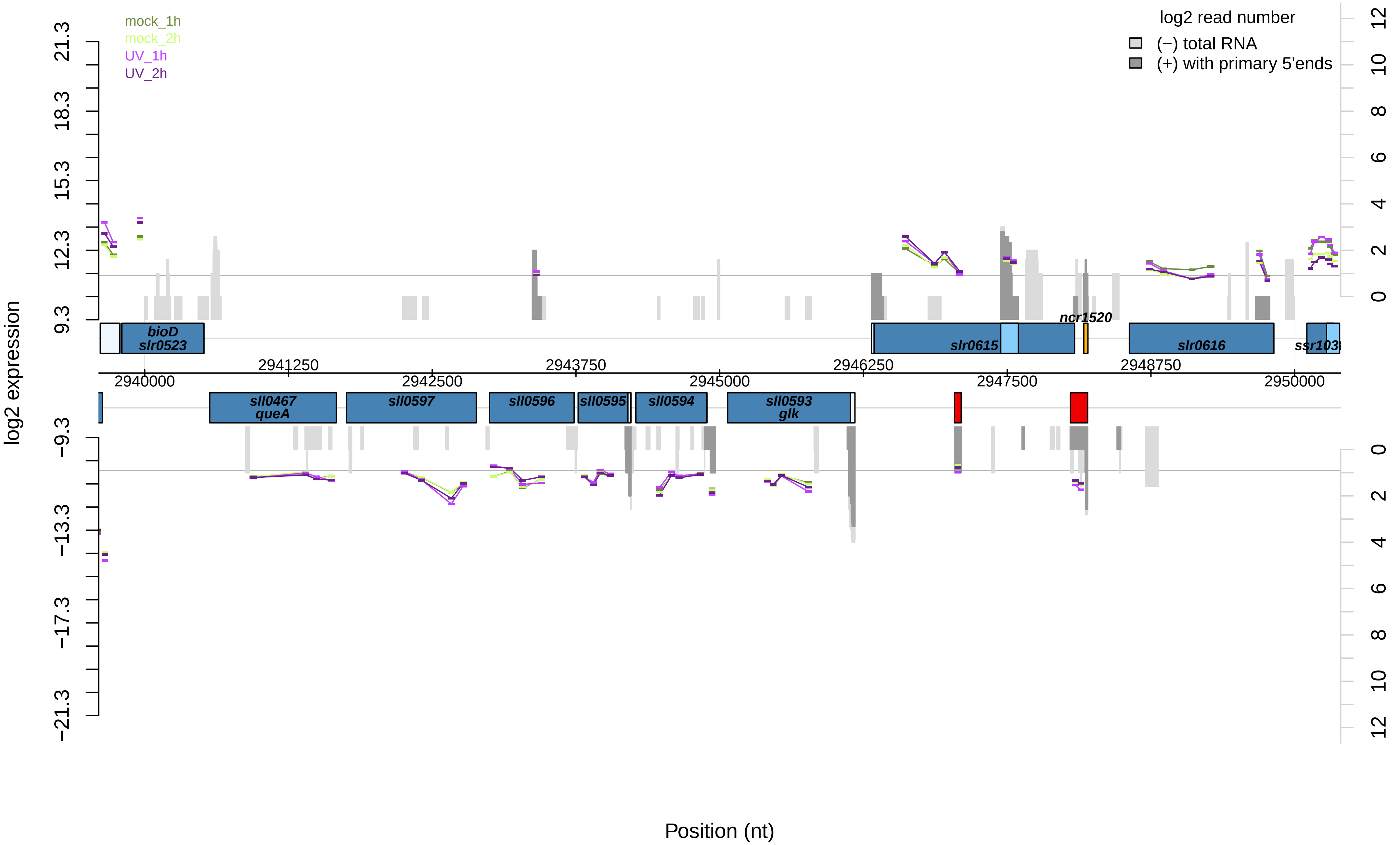

mock_1h
mock_2h
UV_1h
UV_2h
log2 read number
(−) total RNA
(+) with primary 5'ends
log2 expression
21.3
18.3
15.3
12.3
9.3
−9.3
−13.3
−17.3
−21.3
12
10
8
6
4
2
0
bioD
slr0523
slr0615
ncr1520
slr0616
ssr103
2940000
2941250
2942500
2943750
2945000
2946250
2947500
2948750
2950000
sll0467
queA
sll0597
sll0596
sll0595
sll0594
sll0593
glk
Position (nt)

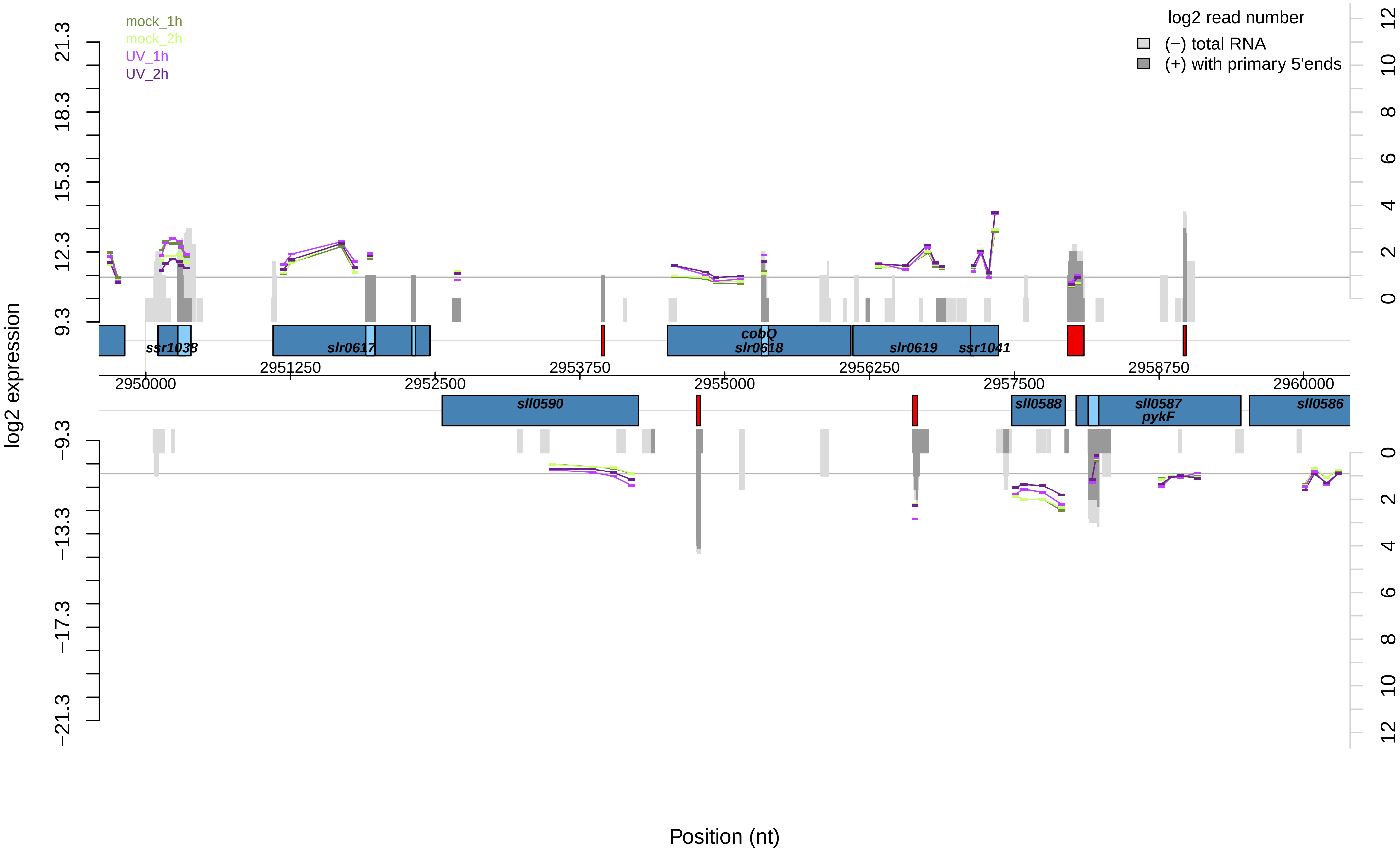
mock_1h
mock_2h
UV_1h
UV_2h
log2 read number
(−) total RNA
(+) with primary 5'ends
log2 expression
Position (nt)
ssr1038
slr0617
cobQ
slr0618
slr0619
ssr1041
sll0590
sll0588
sll0587
pykF
sll0586
2950000
2951250
2952500
2953750
2955000
2956250
2957500
2958750
2960000

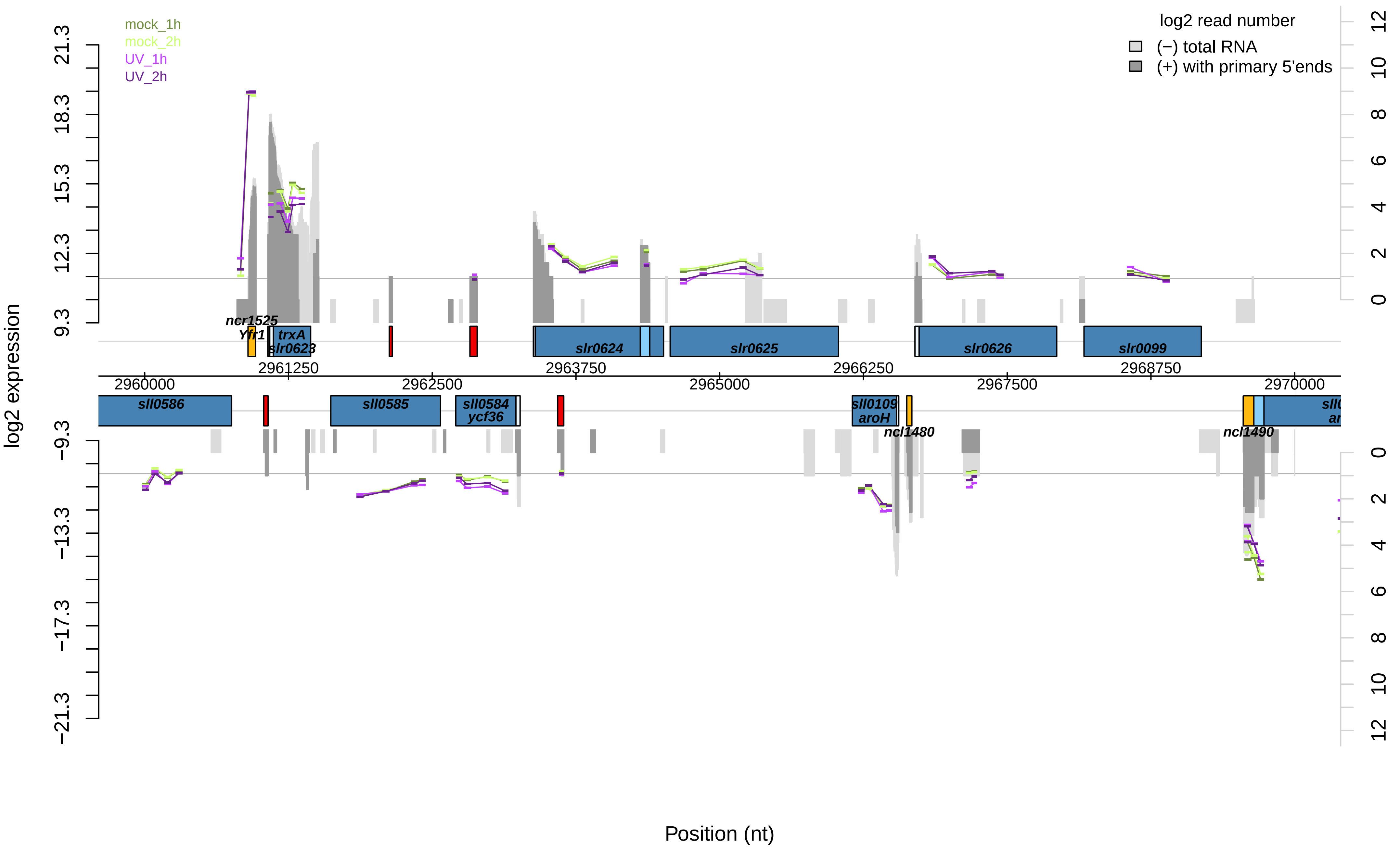

mock_1h
mock_2h
UV_1h
UV_2h
log2 read number
(−) total RNA
(+) with primary 5'ends
log2 expression
21.3
18.3
15.3
12.3
9.3
−9.3
−13.3
−17.3
−21.3
12
10
8
6
4
2
0
ncr1525
Yfr1
trxA
slr0623
slr0624
slr0625
slr0626
slr0099
2961250
2963750
2966250
2968750
2960000
2962500
2965000
2967500
2970000
sll0586
sll0585
sll0584
ycf36
sll0109
aroH
ncl1480
ncl1490
Position (nt)

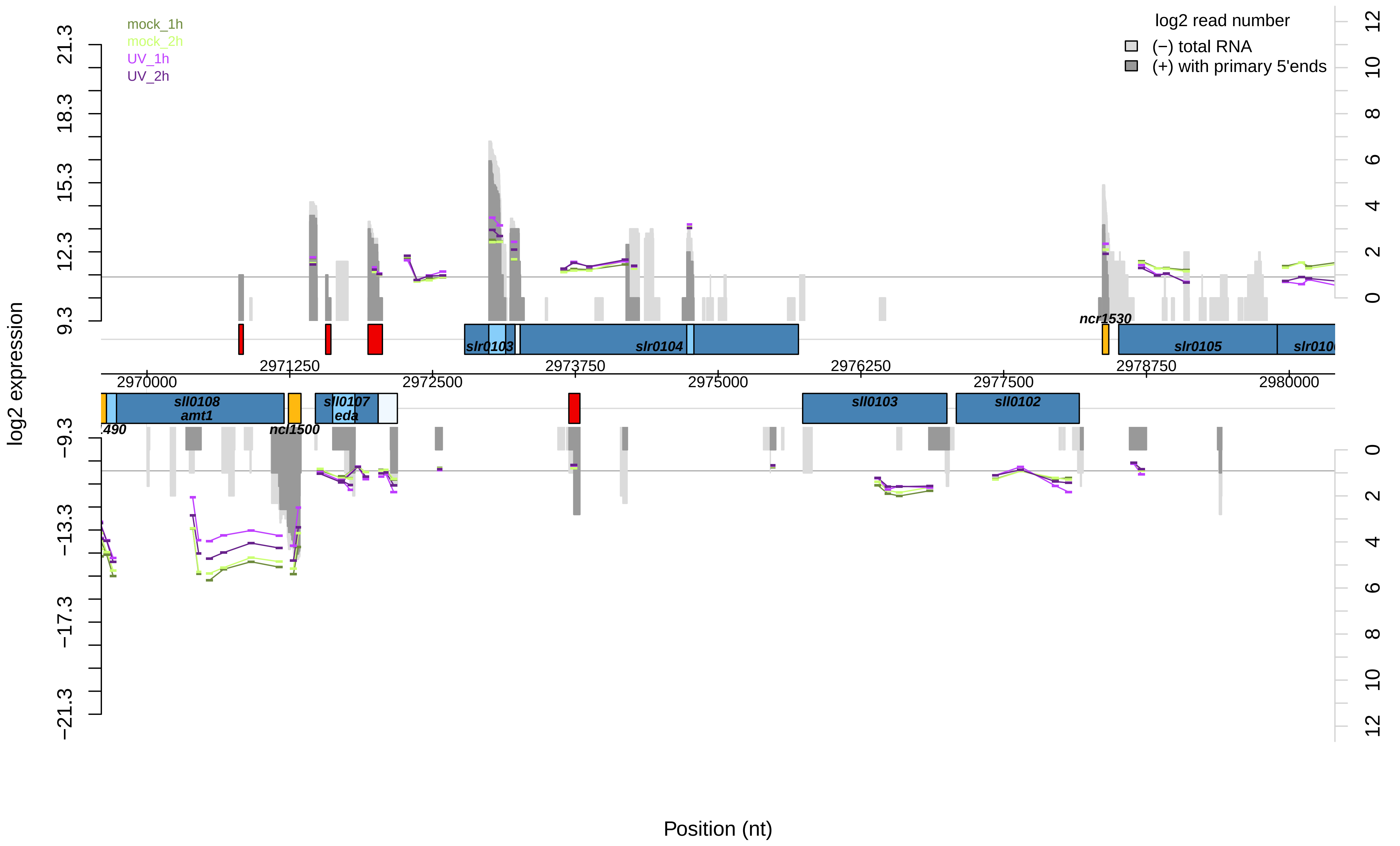

mock_1h
mock_2h
UV_1h
UV_2h
log2 read number
(−) total RNA
(+) with primary 5'ends
log2 expression
21.3
18.3
15.3
12.3
9.3
−9.3
−13.3
−17.3
−21.3
12
10
8
6
4
2
0
ncr1530
slr0103
slr0104
slr0105
2971250
2973750
2976250
2978750
2970000
2972500
2975000
2977500
2980000
sll0108
amt1
sll0107
eda
sll0103
sll0102
ncl1500
Position (nt)

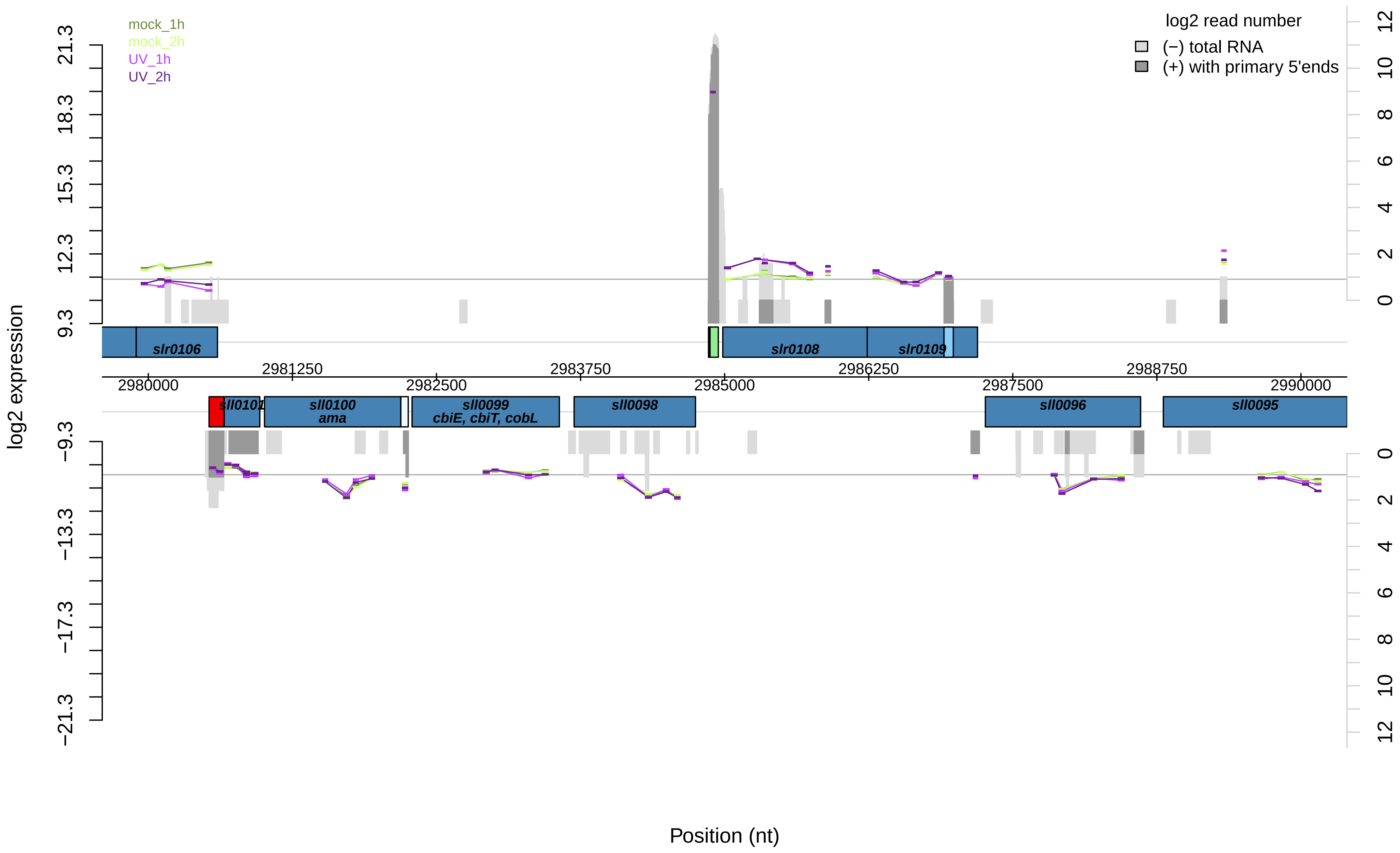
mock_1h
mock_2h
UV_1h
UV_2h
log2 read number
(−) total RNA
(+) with primary 5'ends
log2 expression
21.3
18.3
15.3
12.3
9.3
−9.3
−13.3
−17.3
−21.3
12
10
8
6
4
2
0
slr0106
slr0108
slr0109
2980000
2981250
2982500
2983750
2985000
2986250
2987500
2988750
2990000
sll0101
sll0100
ama
sll0099
cbiE, cbiT, cobL
sll0098
sll0096
sll0095
Position (nt)

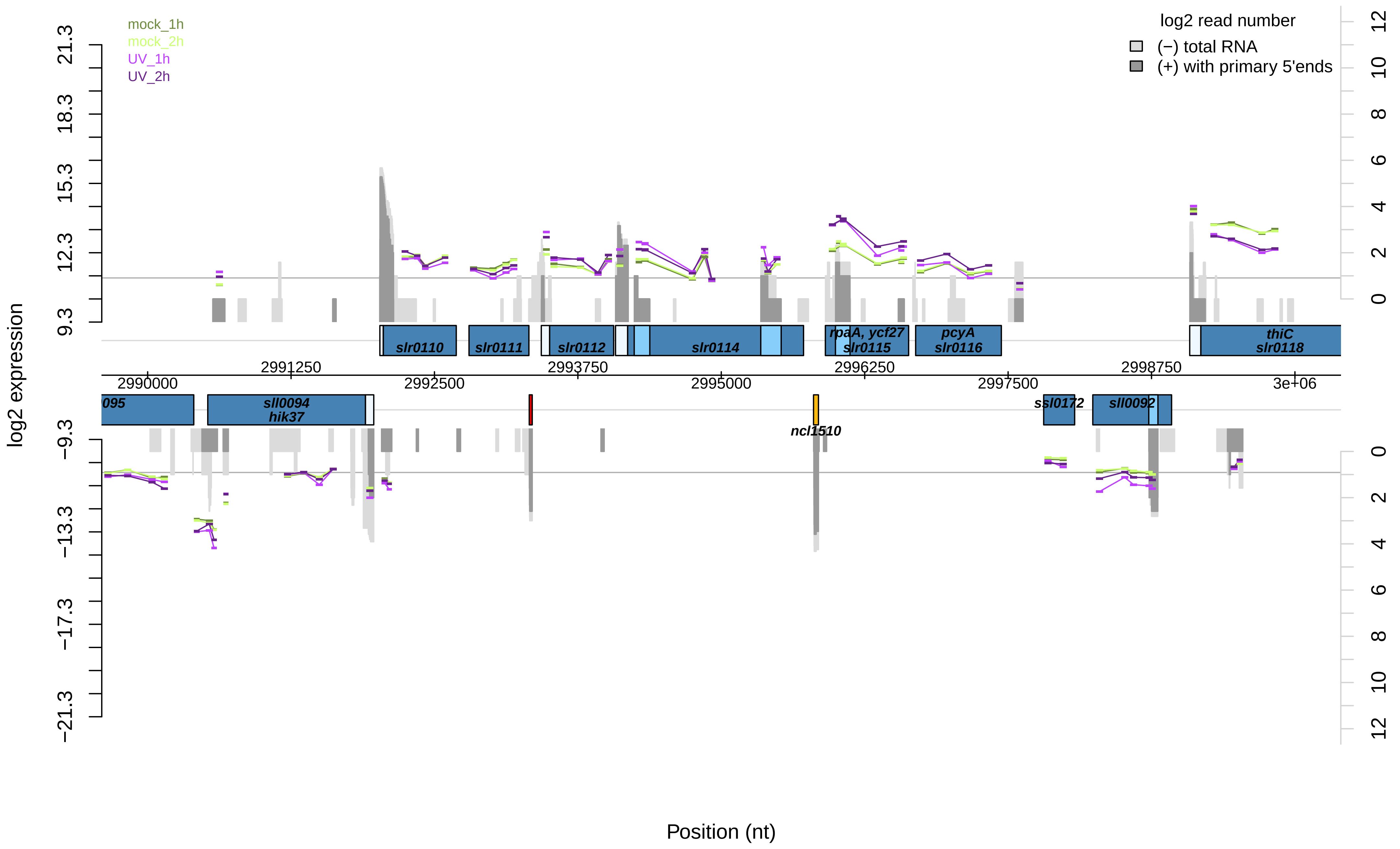
mock_1h
mock_2h
UV_1h
UV_2h
log2 read number
(−) total RNA
(+) with primary 5'ends
log2 expression
21.3
18.3
15.3
12.3
9.3
−9.3
−13.3
−17.3
−21.3
12
10
8
6
4
2
0
slr0110
slr0111
slr0112
slr0114
rpaA, ycf27
slr0115
pcyA
slr0116
thiC
slr0118
2991250
2993750
2996250
2998750
2990000
2992500
2995000
2997500
3e+06
095
sll0094
hik37
ncl1510
ssl0172
sll0092
Position (nt)

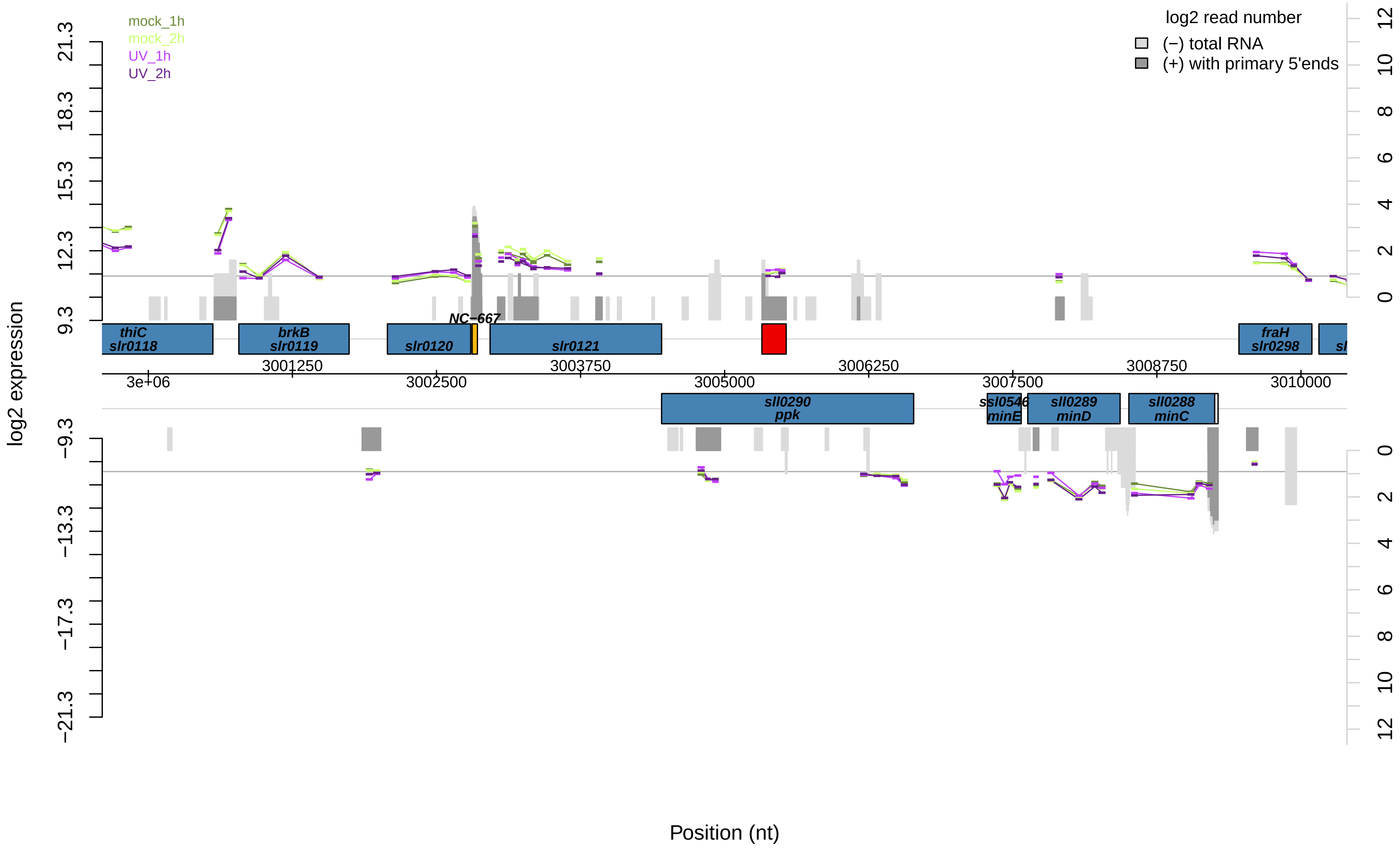

mock_1h
mock_2h
UV_1h
UV_2h
log2 read number
(−) total RNA
(+) with primary 5'ends
log2 expression
Position (nt)
thiC
slr0118
brkB
slr0119
slr0120
NC−667
slr0121
fraH
slr0298
sll0290
ppk
ssl0546
minE
sll0289
minD
sll0288
minC
3e+06
3001250
3002500
3003750
3005000
3006250
3007500
3008750
3010000
21.3
18.3
15.3
12.3
9.3
−9.3
−13.3
−17.3
−21.3

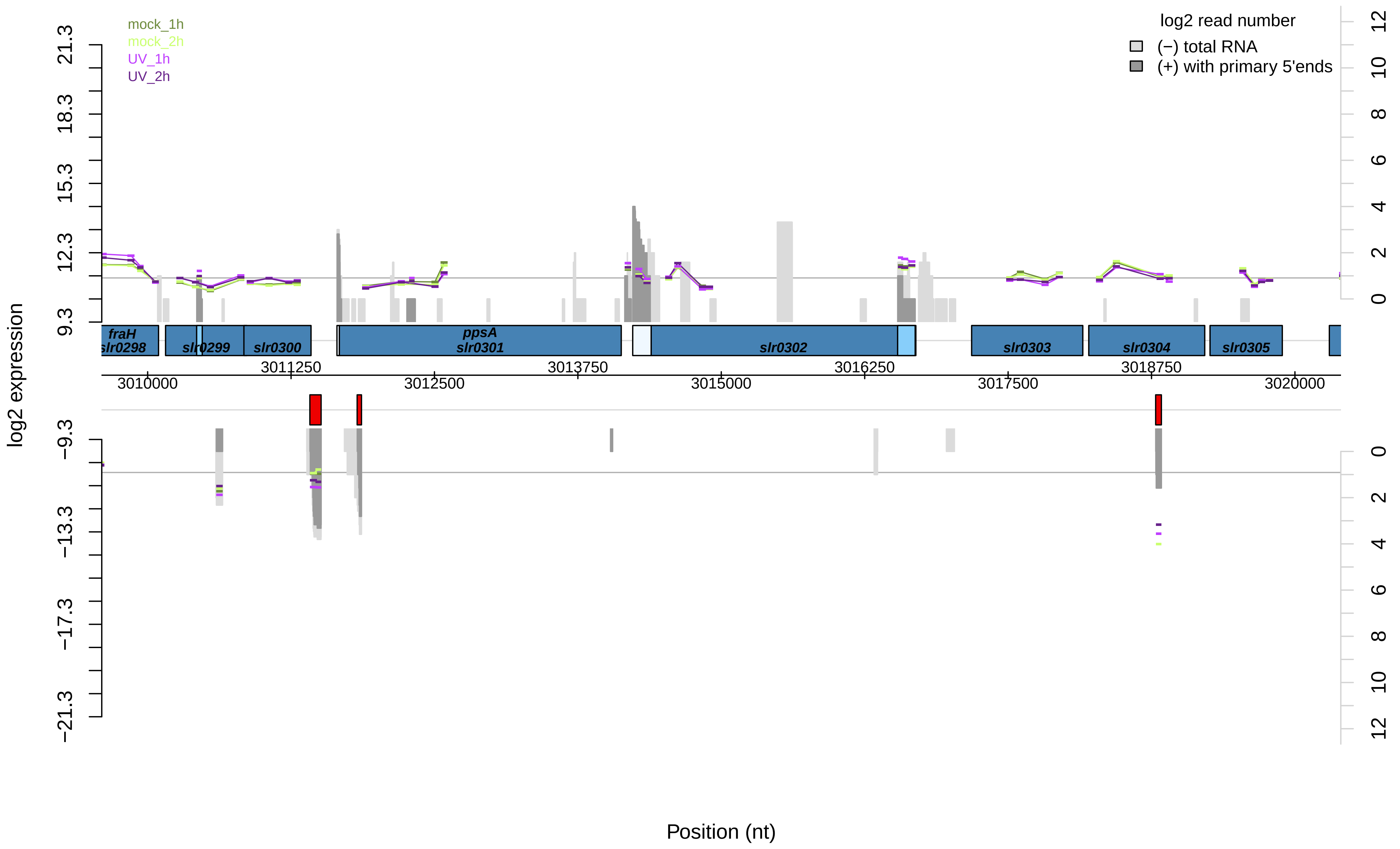

mock_1h
mock_2h
UV_1h
UV_2h
log2 read number
(−) total RNA
(+) with primary 5'ends
log2 expression
21.3
18.3
15.3
12.3
9.3
−9.3
−13.3
−17.3
−21.3
12
10
8
6
4
2
0
fraH
slr0298
slr0299
slr0300
ppsA
slr0301
slr0302
slr0303
slr0304
slr0305
3010000
3011250
3012500
3013750
3015000
3016250
3017500
3018750
3020000
Position (nt)

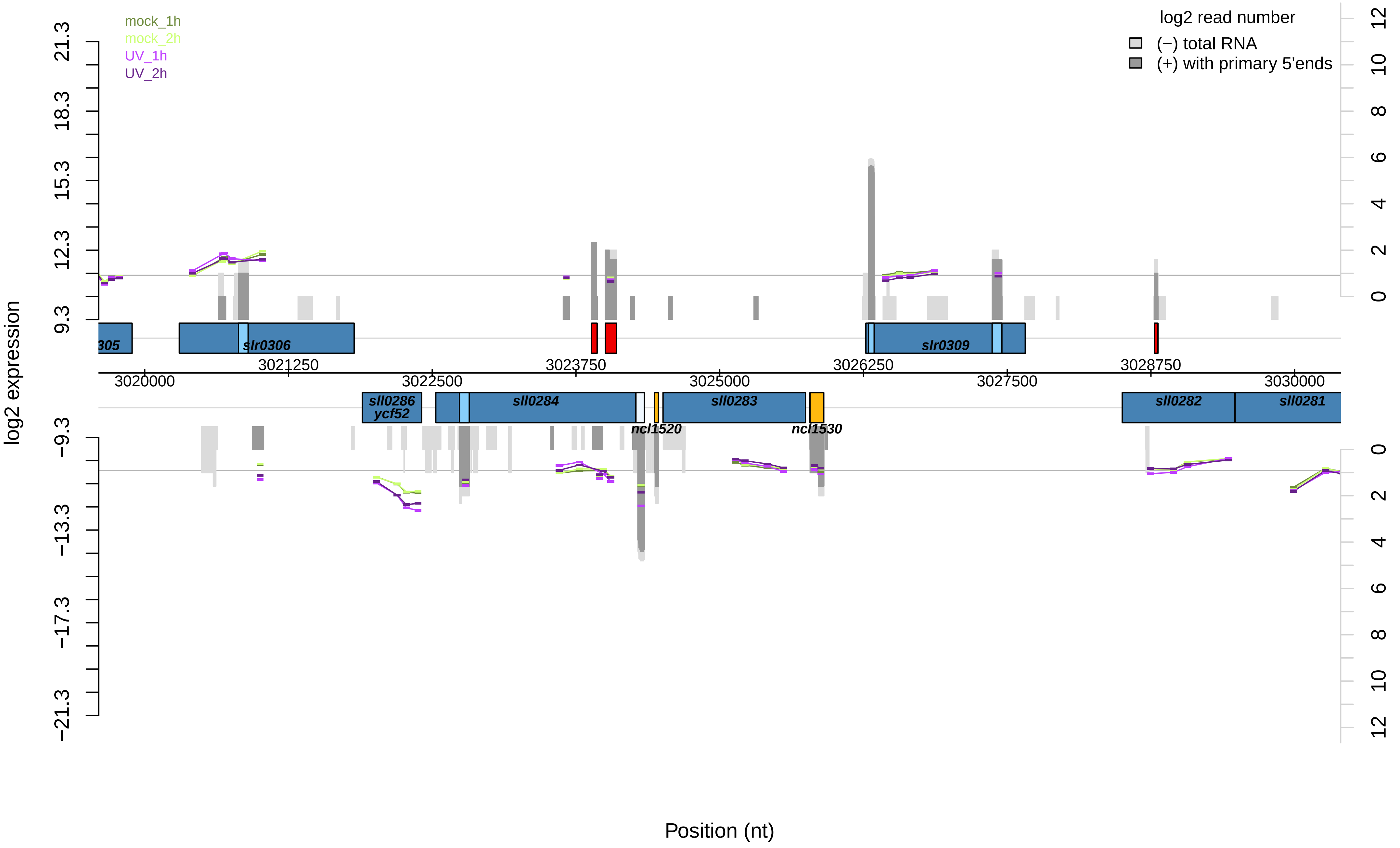

mock_1h
mock_2h
UV_1h
UV_2h
log2 read number
(−) total RNA
(+) with primary 5'ends
log2 expression
21.3
18.3
15.3
12.3
9.3
−9.3
−13.3
−17.3
−21.3
12
10
8
6
4
2
0
305
slr0306
slr0309
3021250
3023750
3026250
3028750
3020000
3022500
3025000
3027500
3030000
sll0286
ycf52
sll0284
sll0283
sll0282
sll0281
ncl1520
ncl1530
Position (nt)

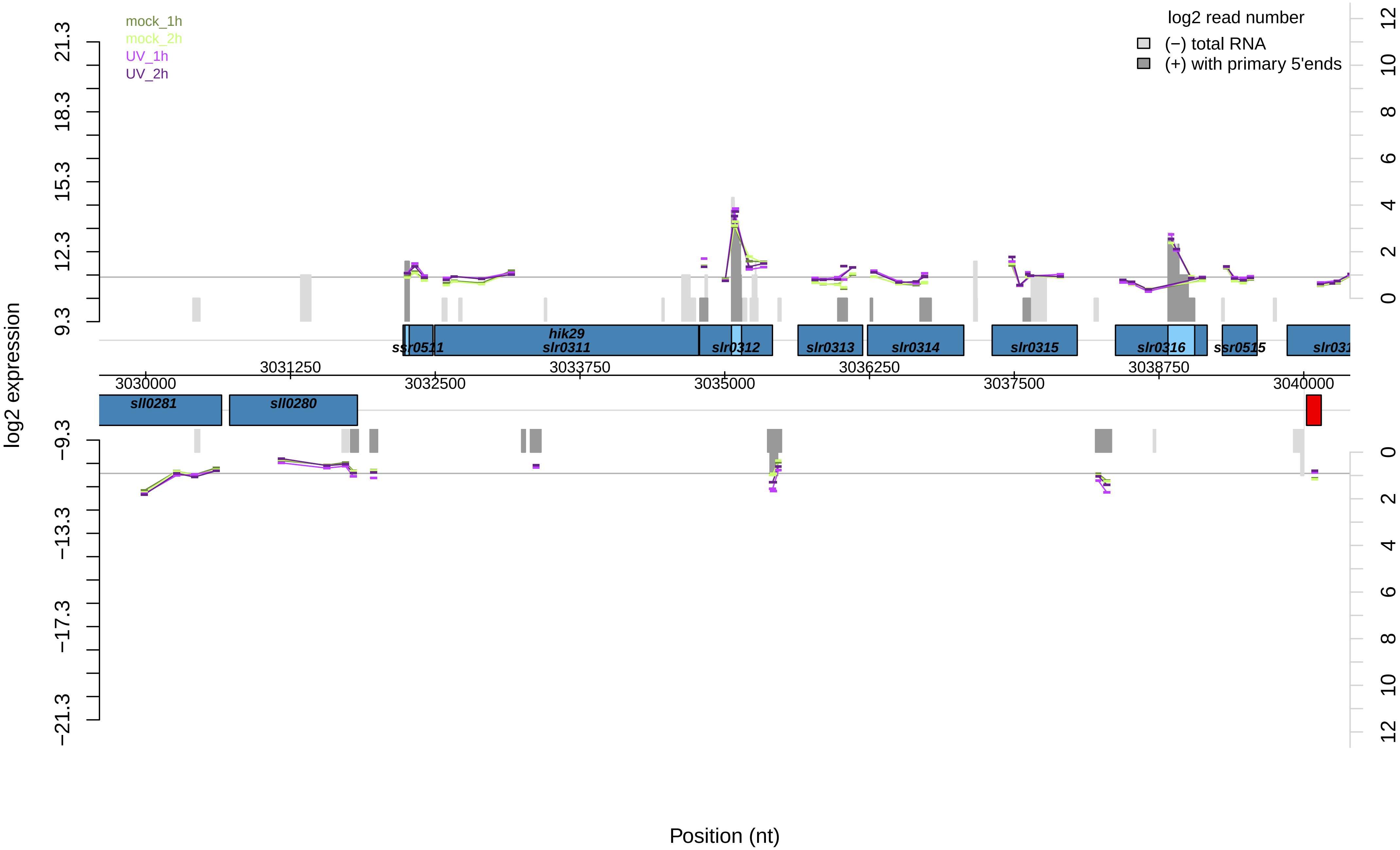

mock_1h
mock_2h
UV_1h
UV_2h
log2 read number
(−) total RNA
(+) with primary 5'ends
log2 expression
21.3
18.3
15.3
12.3
9.3
−9.3
−13.3
−17.3
−21.3
12
10
8
6
4
2
0
hik29
ssr0511
slr0311
slr0312
slr0313
slr0314
slr0315
slr0316
ssr0515
3031250
3033750
3036250
3038750
3030000
3032500
3035000
3037500
3040000
sll0281
sll0280
Position (nt)

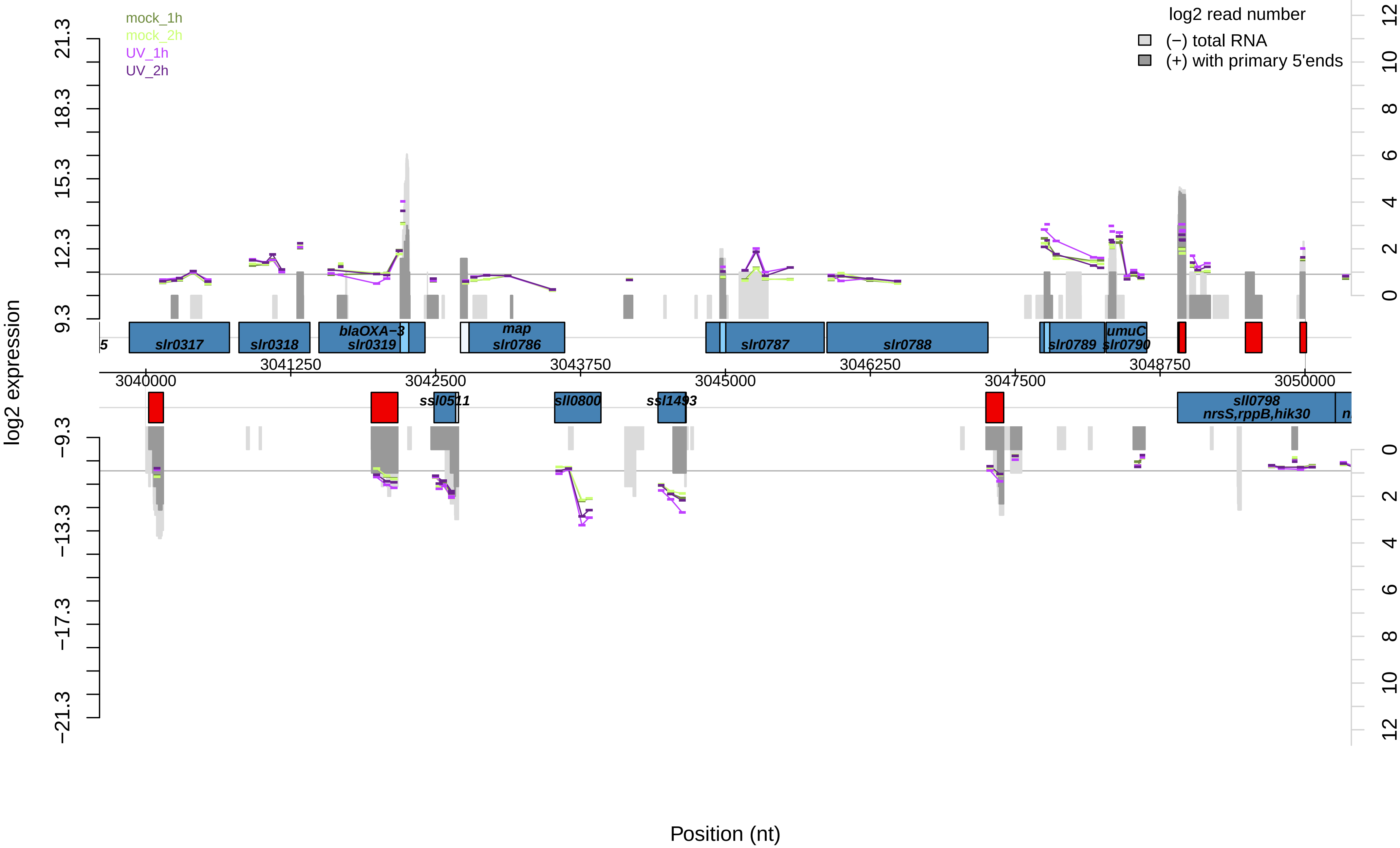

mock_1h
mock_2h
UV_1h
UV_2h
log2 read number
(−) total RNA
(+) with primary 5'ends
log2 expression
Position (nt)
21.3
18.3
15.3
12.3
9.3
−9.3
−13.3
−17.3
−21.3
12
10
8
6
4
2
0
3040000
3041250
3042500
3043750
3045000
3046250
3047500
3048750
3050000
slr0317
slr0318
blaOXA−3
slr0319
map
slr0786
slr0787
slr0788
slr0789
umuC
slr0790
ssl0511
sll0800
ssl1493
sll0798
nrsS,rppB,hik30

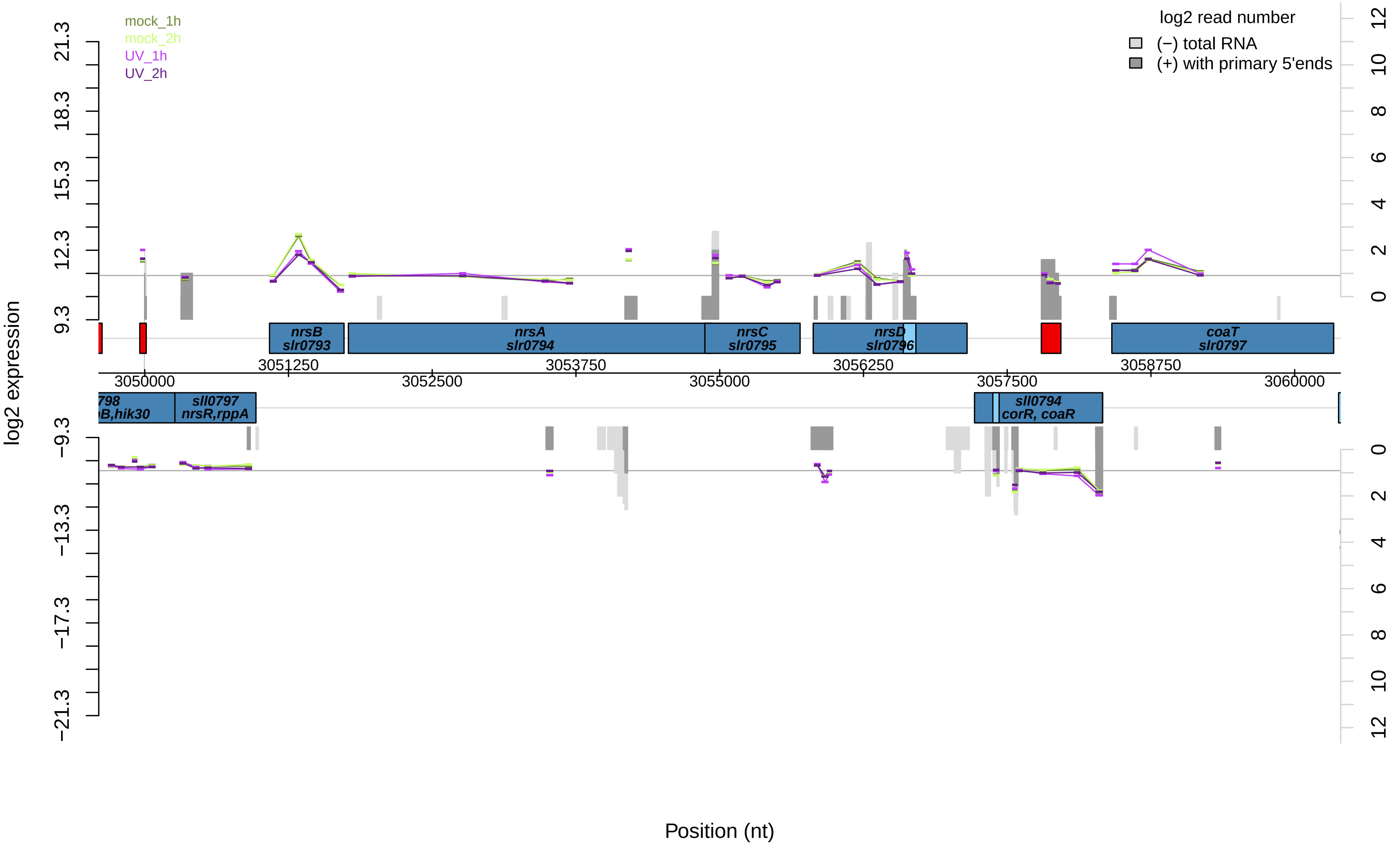
mock_1h
mock_2h
UV_1h
UV_2h
log2 read number
(−) total RNA
(+) with primary 5'ends
log2 expression
21.3
18.3
15.3
12.3
9.3
−9.3
−13.3
−17.3
−21.3
12
10
8
6
4
2
0
nrsB
slr0793
nrsA
slr0794
nrsC
slr0795
nrsD
slr0796
coaT
slr0797
3051250
3053750
3056250
3058750
3050000
3052500
3055000
3057500
3060000
798
B,hik30
sll0797
nrsR,rppA
sll0794
corR, coaR
Position (nt)

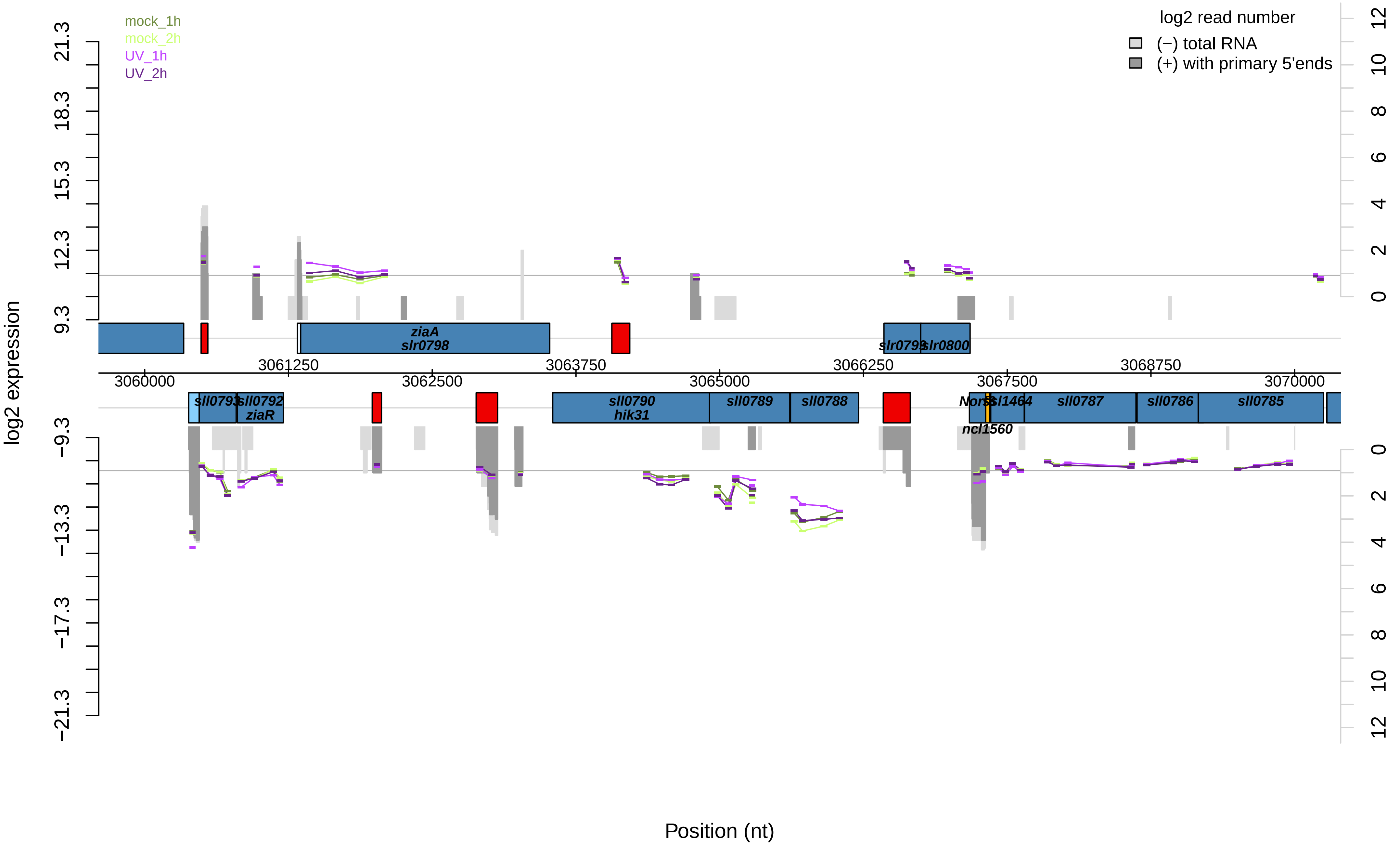

mock_1h
mock_2h
UV_1h
UV_2h
log2 read number
(−) total RNA
(+) with primary 5'ends
log2 expression
21.3
18.3
15.3
12.3
9.3
−9.3
−13.3
−17.3
−21.3
12
10
8
6
4
2
0
ziaA
slr0798
slr0799
slr0800
3061250
3062500
3063750
3065000
3066250
3067500
3068750
3060000
3070000
sll0793
sll0792
ziaR
sll0790
hik31
sll0789
sll0788
ssl1464
sll0787
sll0786
sll0785
ncl1560
Position (nt)

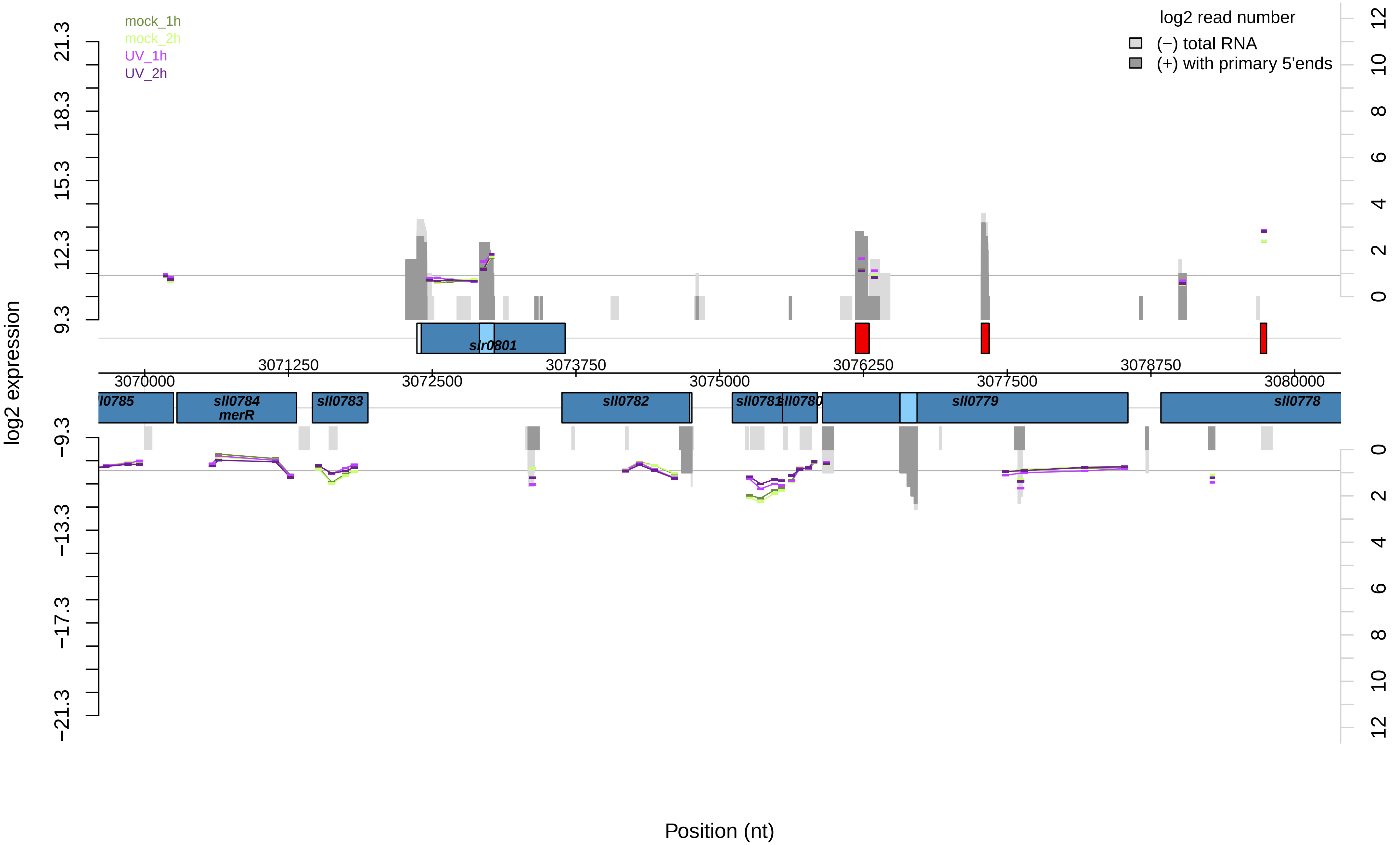
mock_1h
mock_2h
UV_1h
UV_2h
log2 read number
(−) total RNA
(+) with primary 5'ends
log2 expression
21.3
18.3
15.3
12.3
9.3
−9.3
−13.3
−17.3
−21.3
12
10
8
6
4
2
0
slr0801
3071250
3073750
3076250
3078750
3070000
3072500
3075000
3077500
3080000
ll0785
sll0784
merR
sll0783
sll0782
sll0781
sll0780
sll0779
sll0778
Position (nt)

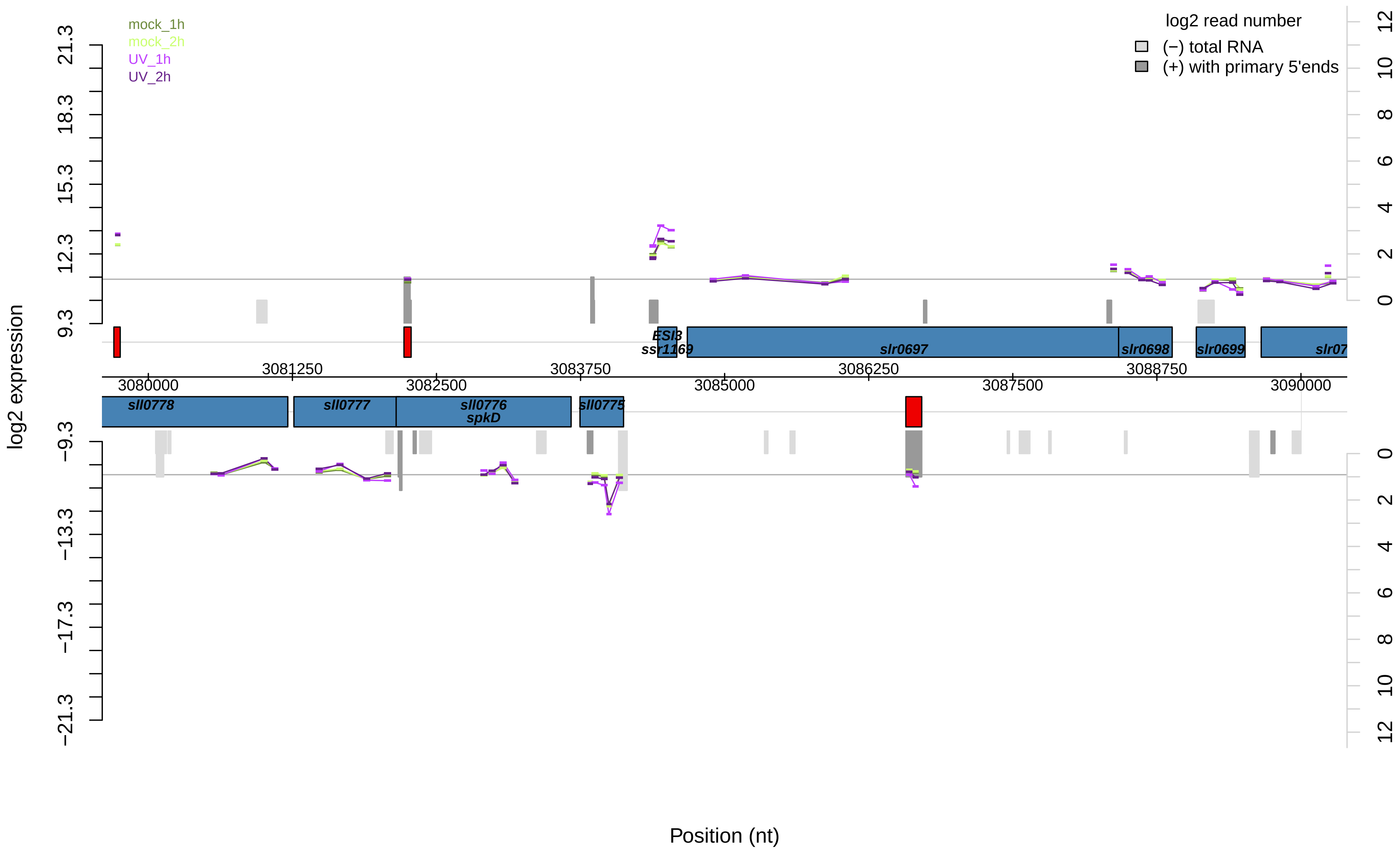

mock_1h
mock_2h
UV_1h
UV_2h
log2 read number
(−) total RNA
(+) with primary 5'ends
log2 expression
21.3
18.3
15.3
12.3
9.3
−9.3
−13.3
−17.3
−21.3
12
10
8
6
4
2
0
ESl3
ssr1169
slr0697
slr0698
slr0699
slr07
sll0778
sll0777
sll0776
spkD
sll0775
3080000
3081250
3082500
3083750
3085000
3086250
3087500
3088750
3090000
Position (nt)

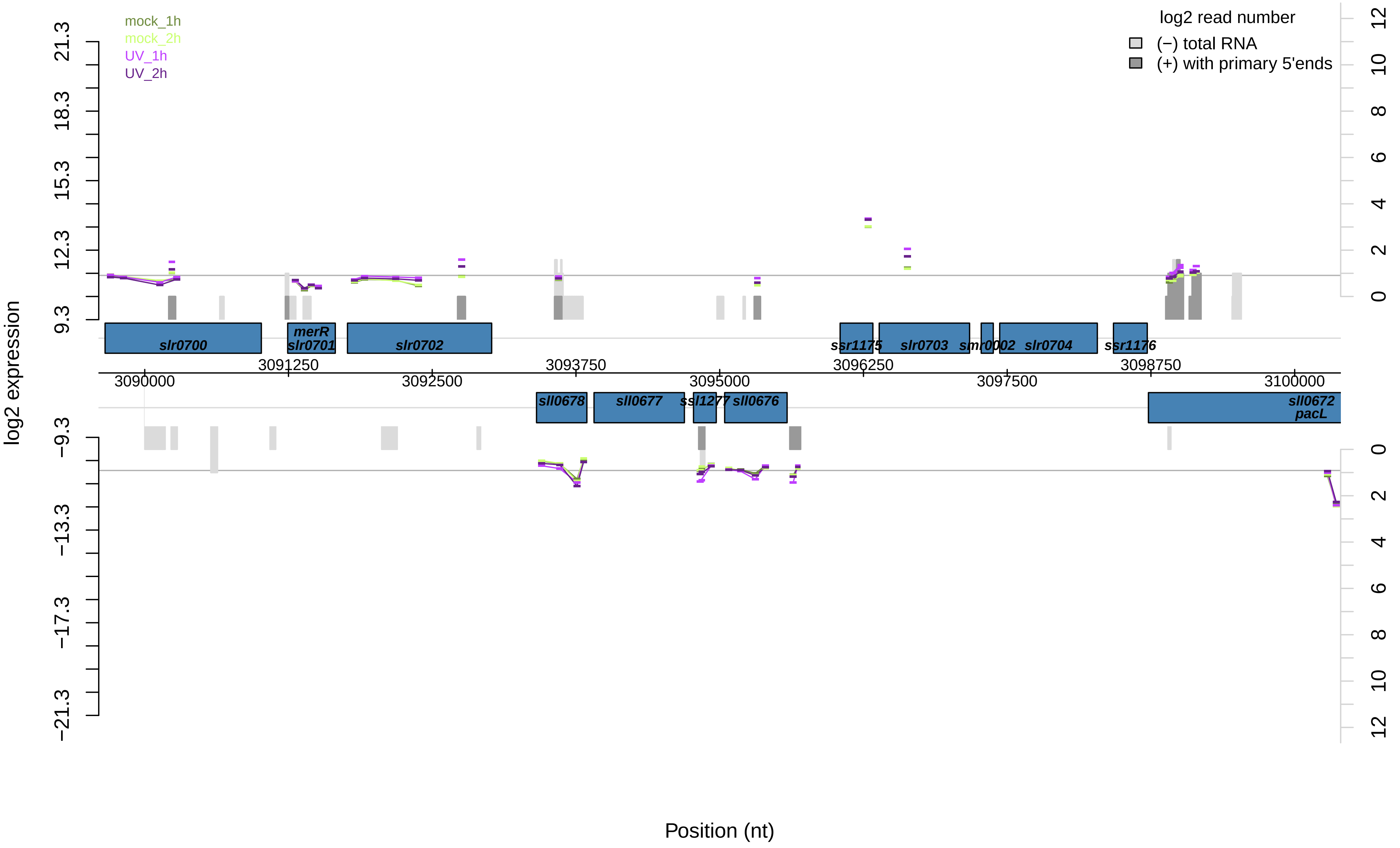

mock_1h
mock_2h
UV_1h
UV_2h
log2 read number
(−) total RNA
(+) with primary 5'ends
log2 expression
21.3
18.3
15.3
12.3
9.3
−9.3
−13.3
−17.3
−21.3
12
10
8
6
4
2
0
slr0700
merR
slr0701
slr0702
ssr1175
slr0703
smr0002
slr0704
ssr1176
3091250
3093750
3096250
3098750
3090000
3092500
3095000
3097500
3100000
sll0678
sll0677
ssl1277
sll0676
sll0672
pacL
Position (nt)

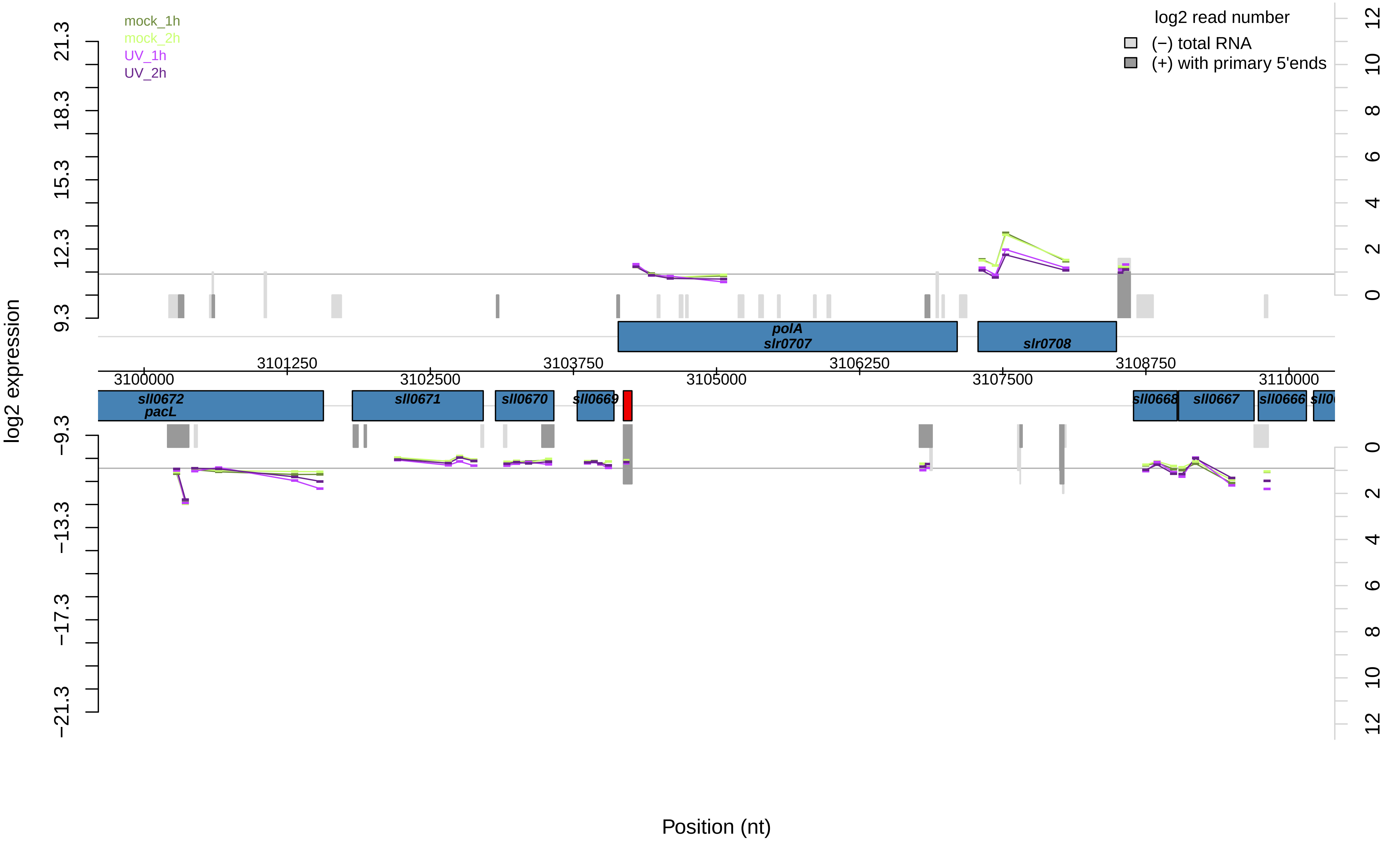

mock_1h
mock_2h
UV_1h
UV_2h
log2 read number
(−) total RNA
(+) with primary 5'ends
log2 expression
21.3
18.3
15.3
12.3
9.3
−9.3
−13.3
−17.3
−21.3
12
10
8
6
4
2
0
polA
slr0707
slr0708
3100000
3101250
3102500
3103750
3105000
3106250
3107500
3108750
3110000
sll0672
pacL
sll0671
sll0670
sll0669
sll0668
sll0667
sll0666
Position (nt)

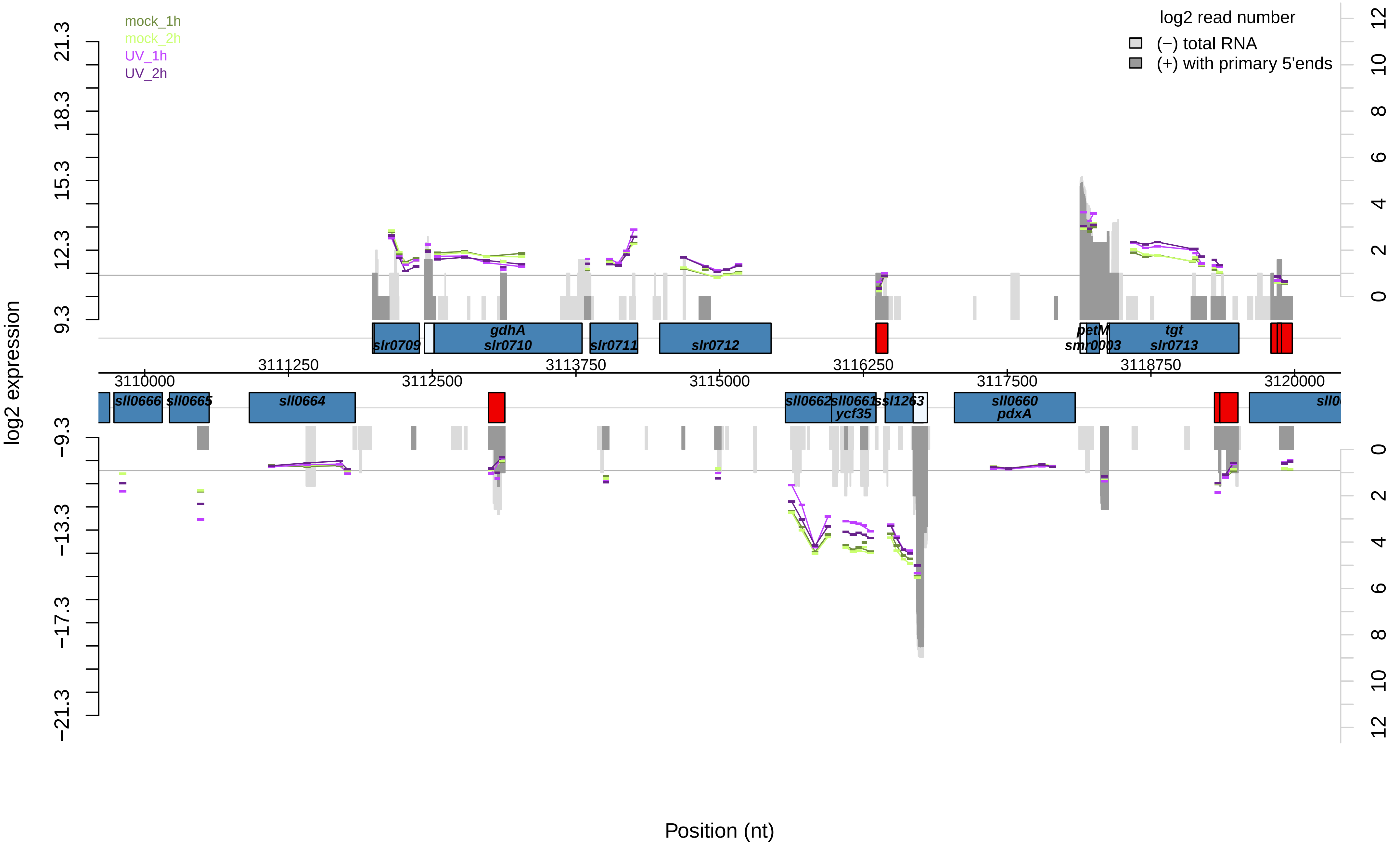

mock_1h
mock_2h
UV_1h
UV_2h
log2 read number
(−) total RNA
(+) with primary 5'ends
log2 expression
21.3
18.3
15.3
12.3
9.3
−9.3
−13.3
−17.3
−21.3
12
10
8
6
4
2
0
slr0709
gdhA
slr0710
slr0711
slr0712
petM
smr0003
tgt
slr0713
3111250
3113750
3116250
3118750
3110000
3112500
3115000
3117500
3120000
sll0666
sll0665
sll0664
sll0662
sll0661
ycf35
ssl1263
sll0660
pdxA
Position (nt)

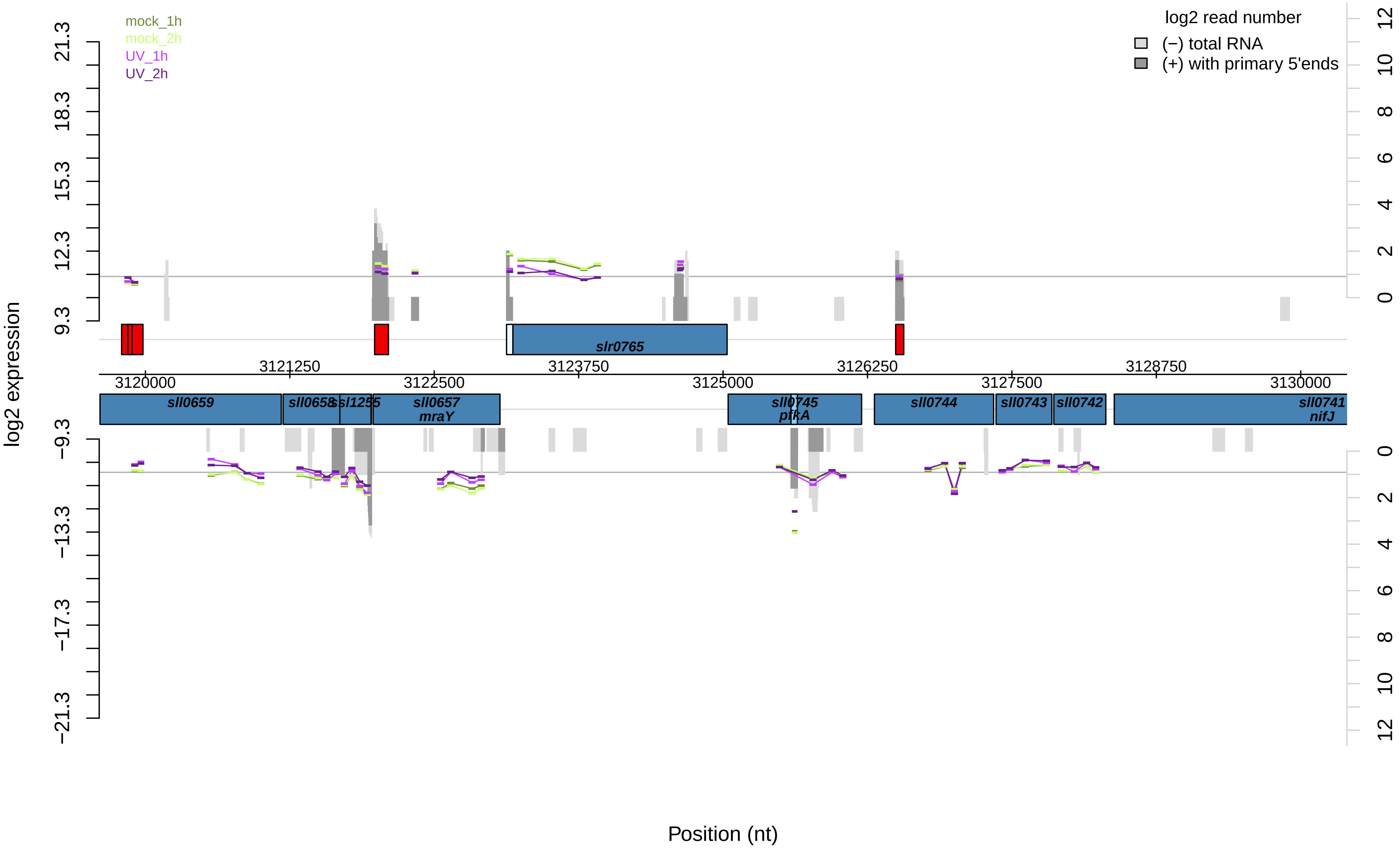
mock_1h
mock_2h
UV_1h
UV_2h
log2 read number
(−) total RNA
(+) with primary 5'ends
log2 expression
21.3
18.3
15.3
12.3
9.3
−9.3
−13.3
−17.3
−21.3
12
10
8
6
4
2
0
slr0765
3121250
3122500
3123750
3125000
3126250
3127500
3128750
3120000
3130000
sll0659
sll0658
ssl1255
sll0657
mraY
sll0745
pfkA
sll0744
sll0743
sll0742
sll0741
nifJ
Position (nt)

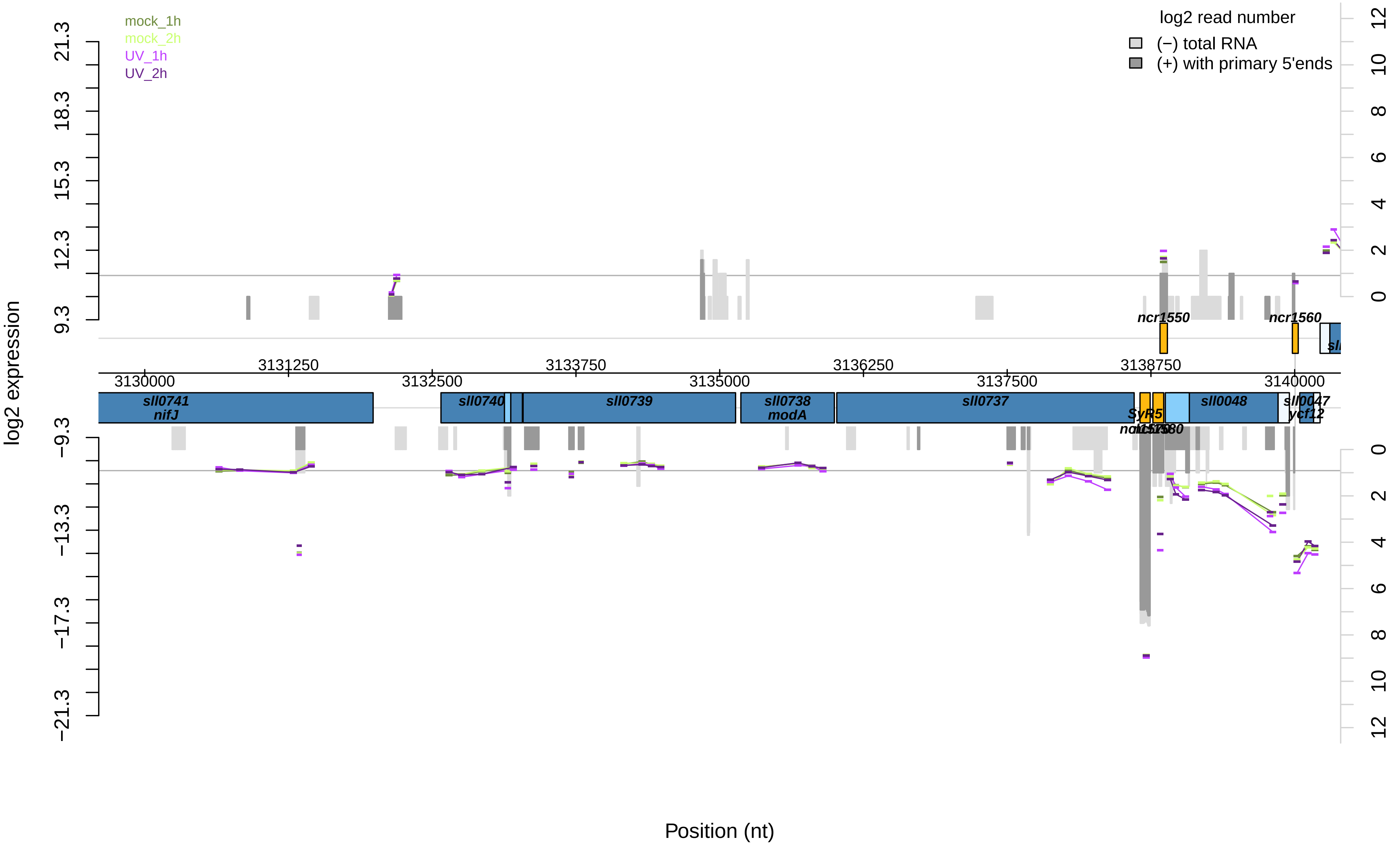

mock_1h
mock_2h
UV_1h
UV_2h
log2 read number
(−) total RNA
(+) with primary 5'ends
log2 expression
21.3
18.3
15.3
12.3
9.3
−9.3
−13.3
−17.3
−21.3
12
10
8
6
4
2
0
ncr1550
ncr1560
3130000
3131250
3132500
3133750
3135000
3136250
3137500
3138750
3140000
sll0741
nifJ
sll0740
sll0739
sll0738
modA
sll0737
SyR5
sll0048
sll0047
ycf12
Position (nt)

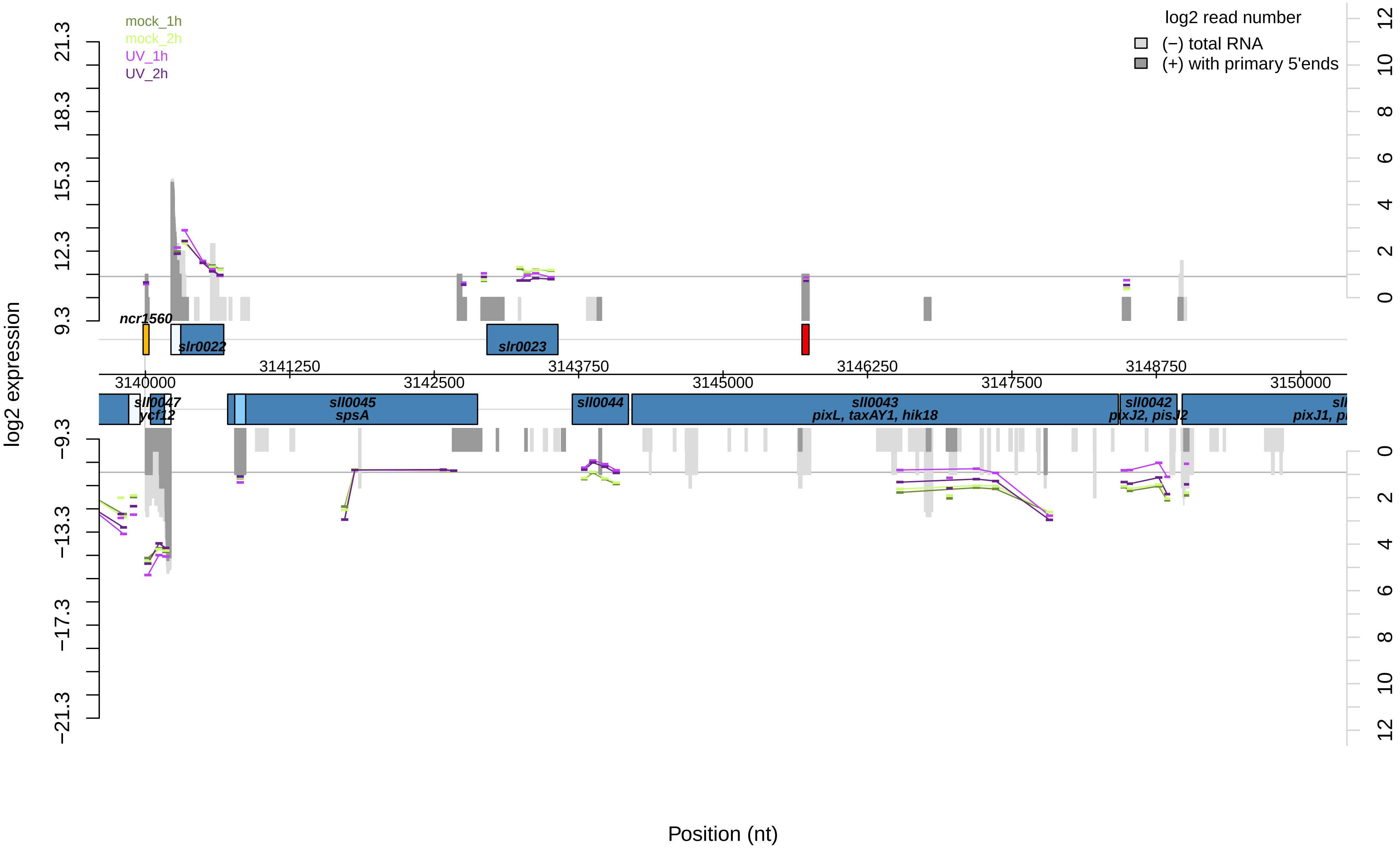

mock_1h
mock_2h
UV_1h
UV_2h
log2 read number
(−) total RNA
(+) with primary 5'ends
log2 expression
21.3
18.3
15.3
12.3
9.3
−9.3
−13.3
−17.3
−21.3
ncr1560
slr0022
slr0023
3141250
3143750
3146250
3148750
3140000
3142500
3145000
3147500
3150000
sll0047
ycf12
sll0045
spsA
sll0044
sll0043
pixL, taxAY1, hik18
sll0042
pixJ2, pisJ2
Position (nt)

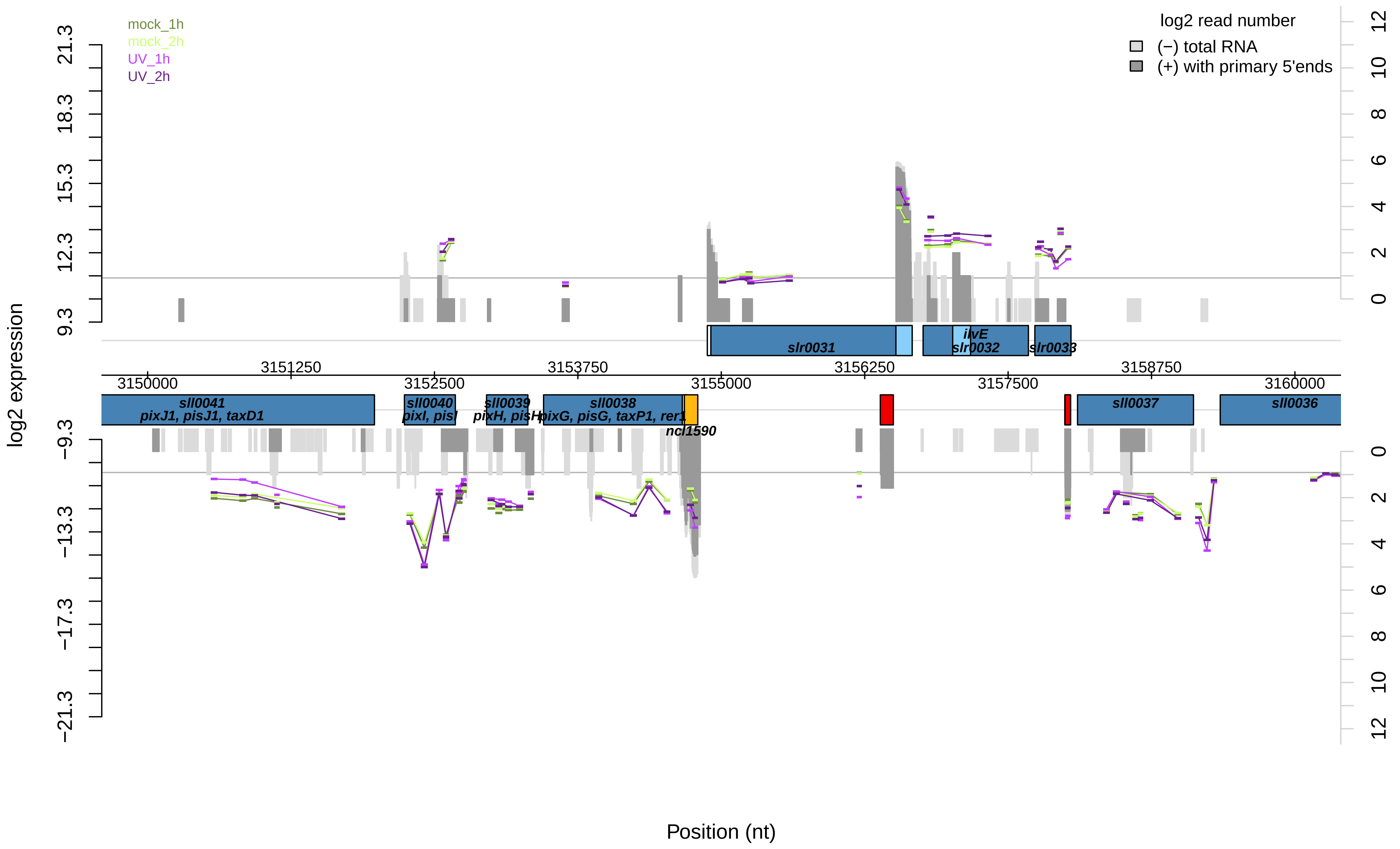

mock_1h
mock_2h
UV_1h
UV_2h
log2 read number
(−) total RNA
(+) with primary 5'ends
log2 expression
21.3
18.3
15.3
12.3
9.3
−9.3
−13.3
−17.3
−21.3
slr0031
ilvE
slr0032
slr0033
3150000
3151250
3152500
3153750
3155000
3156250
3157500
3158750
3160000
sll0041
pixJ1, pisJ1, taxD1
sll0040
pixI, pisI
sll0039
pixH, pisH
sll0038
pixG, pisG, taxP1, rer1
ncl1590
sll0037
sll0036
Position (nt)

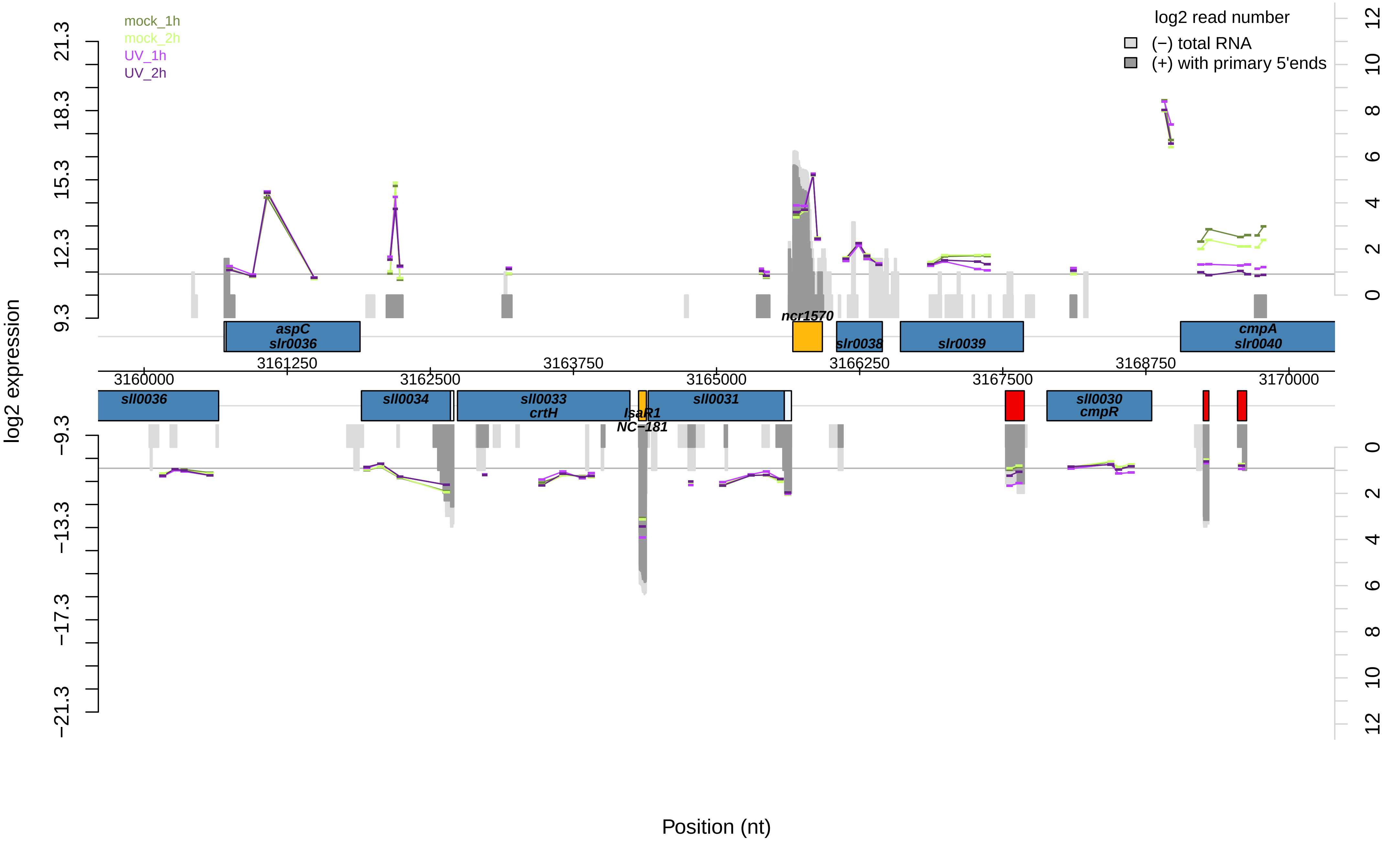
mock_1h
mock_2h
UV_1h
UV_2h
log2 read number
(−) total RNA
(+) with primary 5'ends
log2 expression
21.3
18.3
15.3
12.3
9.3
−9.3
−13.3
−17.3
−21.3
12
10
8
6
4
2
0
ncr1570
aspC
slr0036
slr0038
slr0039
cmpA
slr0040
3161250
3163750
3166250
3168750
3160000
3162500
3165000
3167500
3170000
sll0036
sll0034
sll0033
crtH
IsaR1
NC−181
sll0031
sll0030
cmpR
Position (nt)

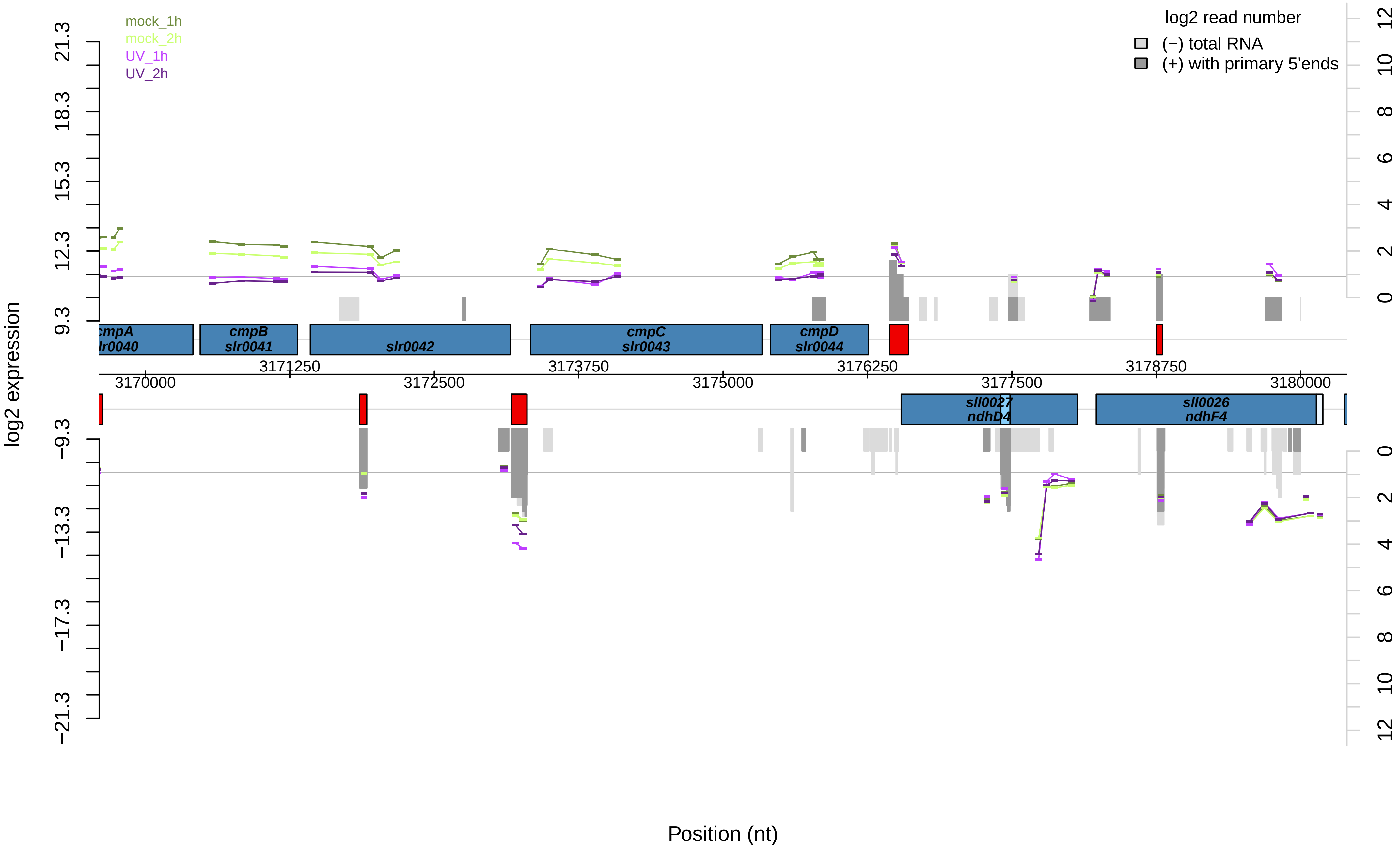

mock_1h
mock_2h
UV_1h
UV_2h
log2 read number
(−) total RNA
(+) with primary 5'ends
log2 expression
21.3
18.3
15.3
12.3
9.3
−9.3
−13.3
−17.3
−21.3
12
10
8
6
4
2
0
cmpA
slr0040
cmpB
slr0041
slr0042
cmpC
slr0043
cmpD
slr0044
3171250
3173750
3176250
3178750
3170000
3172500
3175000
3177500
3180000
sll0027
ndhD4
sll0026
ndhF4
Position (nt)

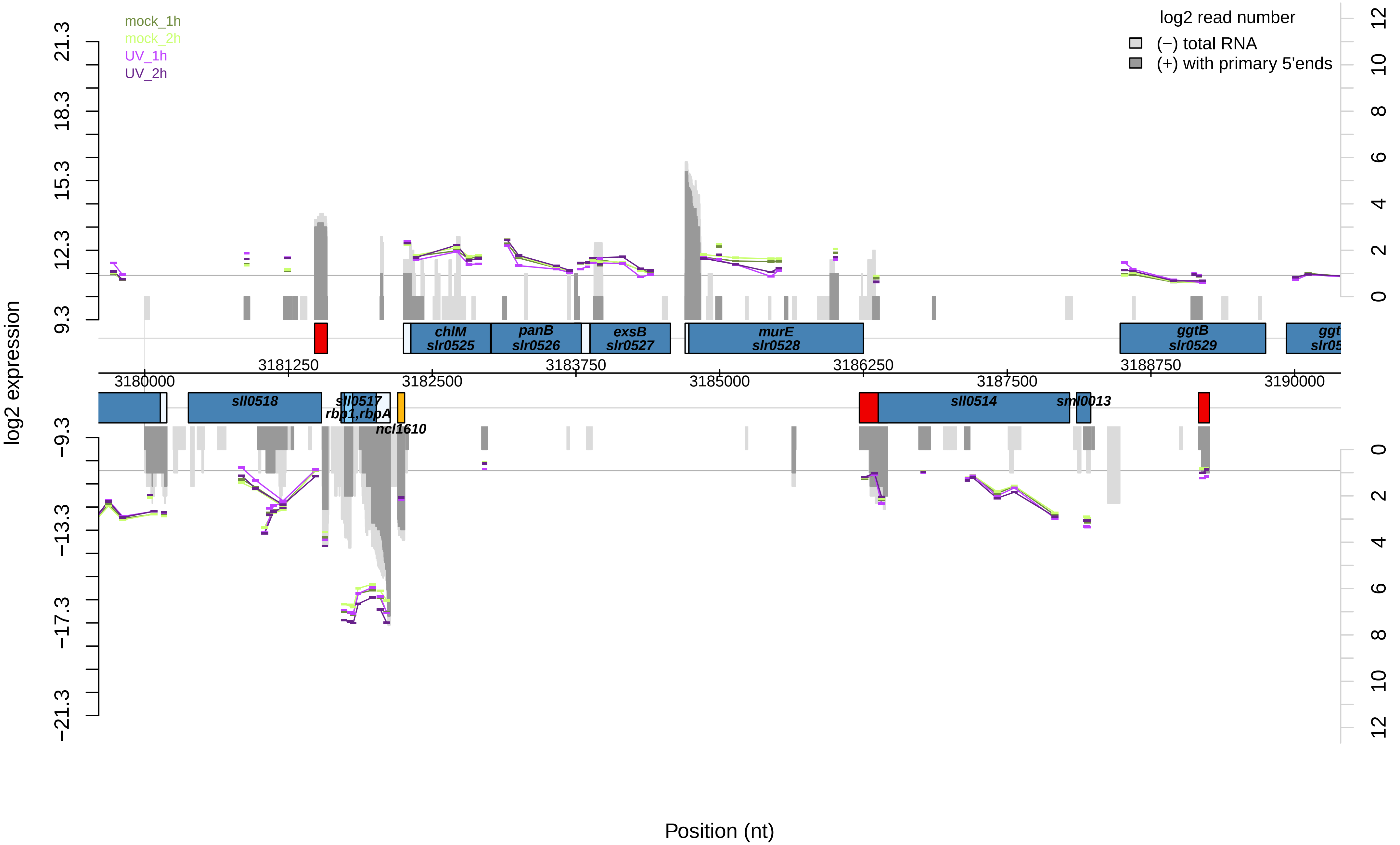

mock_1h
mock_2h
UV_1h
UV_2h
log2 read number
(−) total RNA
(+) with primary 5'ends
log2 expression
21.3
18.3
15.3
12.3
9.3
−9.3
−13.3
−17.3
−21.3
12
10
8
6
4
2
0
chlM
slr0525
panB
slr0526
exsB
slr0527
murE
slr0528
ggtB
slr0529
3181250
3183750
3186250
3188750
3180000
3182500
3185000
3187500
3190000
sll0518
sll0517
rbp1,rbpA
ncl1610
sll0514
sml0013
Position (nt)

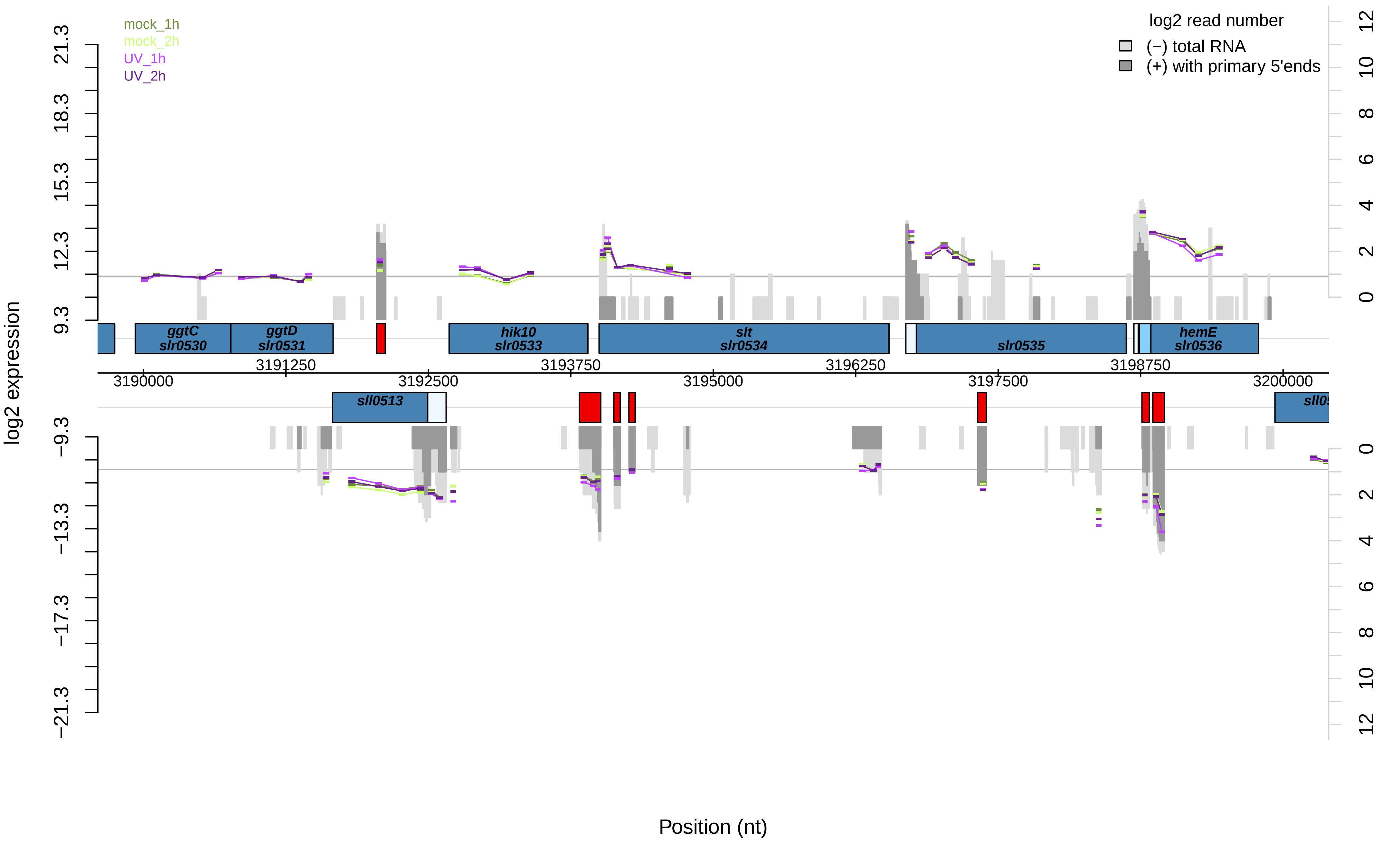
mock_1h
mock_2h
UV_1h
UV_2h
log2 read number
(−) total RNA
(+) with primary 5'ends
log2 expression
21.3
18.3
15.3
12.3
9.3
−9.3
−13.3
−17.3
−21.3
ggtC
slr0530
ggtD
slr0531
hik10
slr0533
slt
slr0534
slr0535
hemE
slr0536
sll0513
3191250
3193750
3196250
3198750
3190000
3192500
3195000
3197500
3200000
Position (nt)

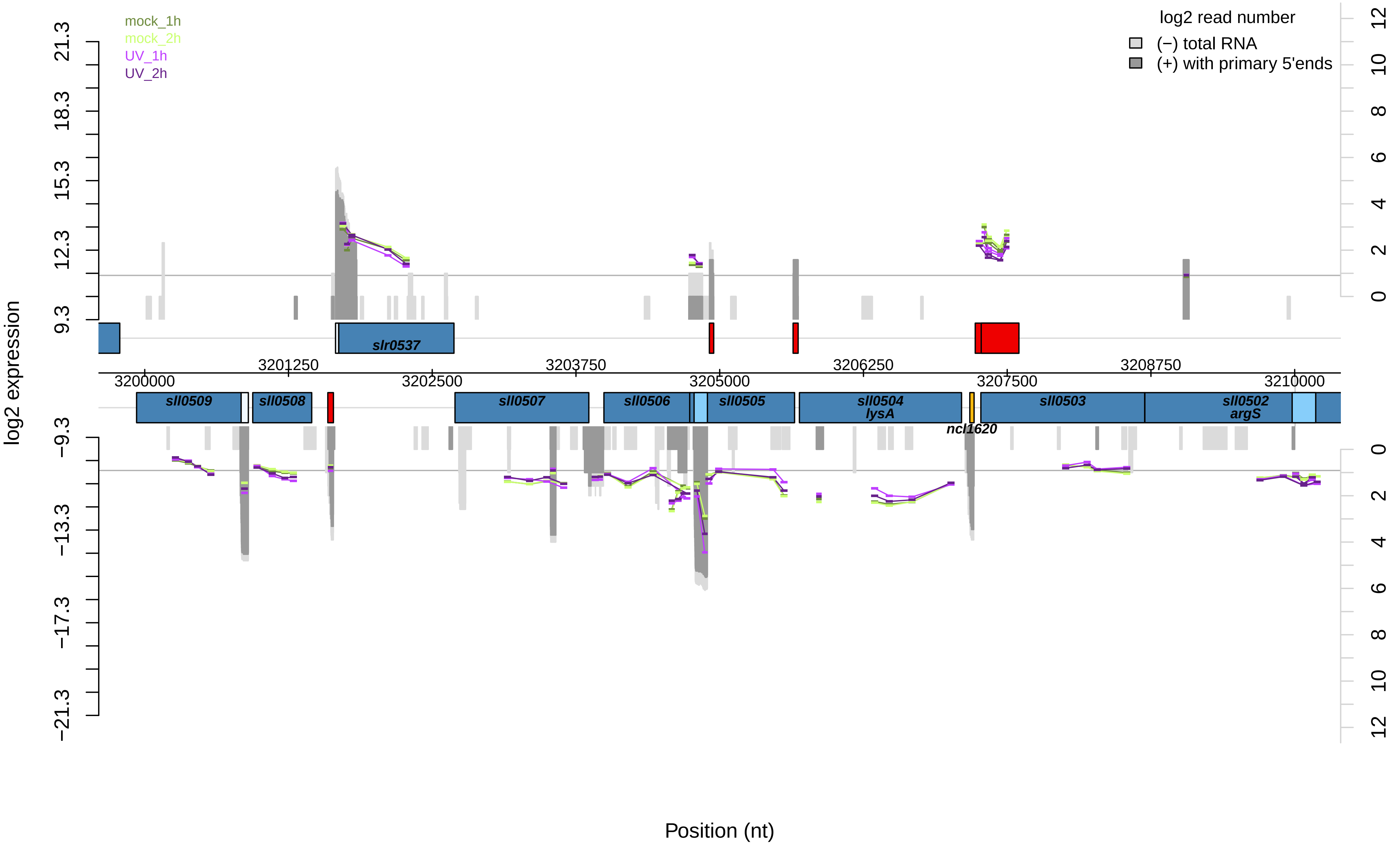

mock_1h
mock_2h
UV_1h
UV_2h
log2 read number
(−) total RNA
(+) with primary 5'ends
log2 expression
21.3
18.3
15.3
12.3
9.3
−9.3
−13.3
−17.3
−21.3
12
10
8
6
4
2
0
slr0537
3201250
3203750
3206250
3208750
3200000
3202500
3205000
3207500
3210000
sll0509
sll0508
sll0507
sll0506
sll0505
sll0504
lysA
ncl1620
sll0503
sll0502
argS
Position (nt)

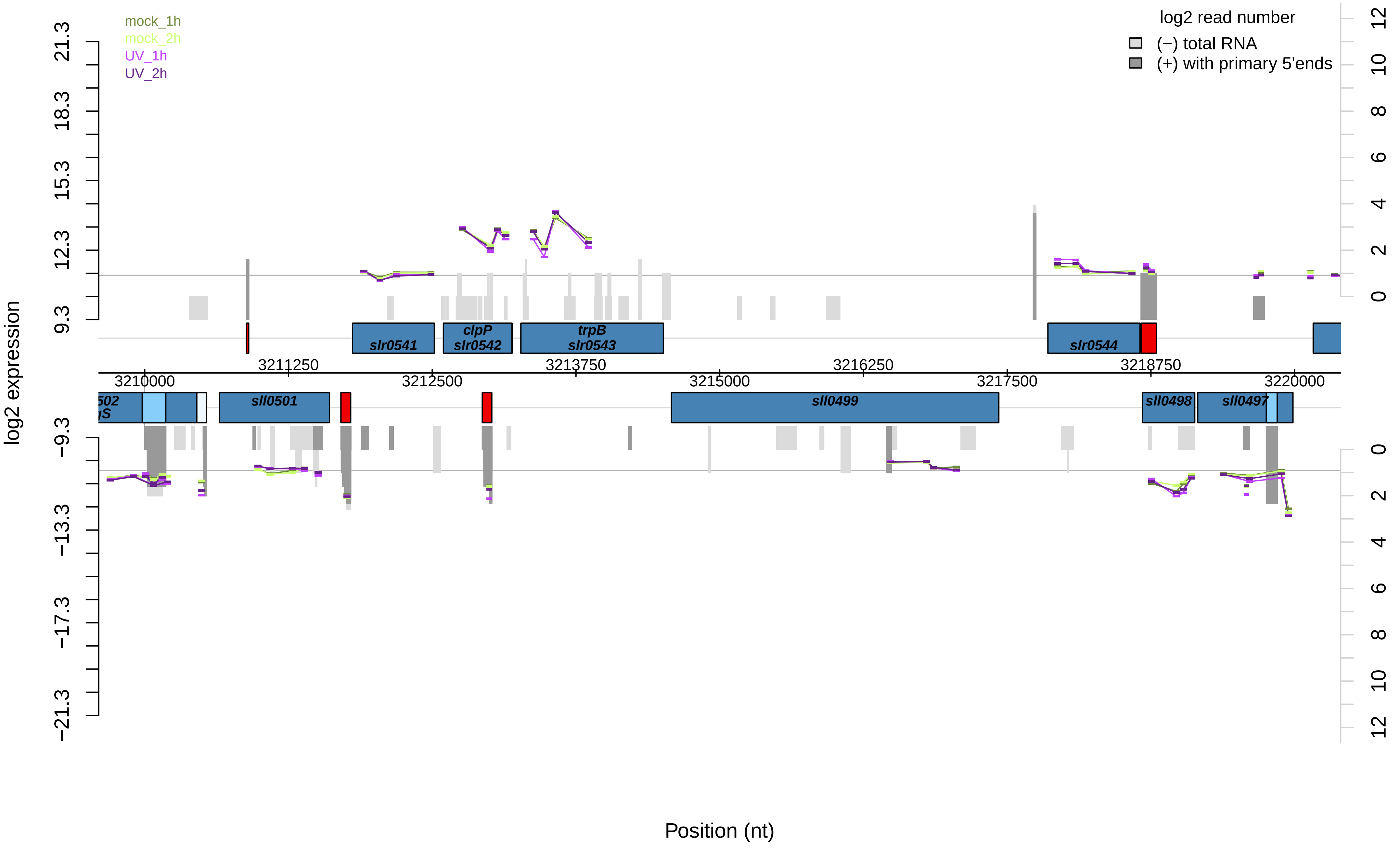

mock_1h
mock_2h
UV_1h
UV_2h
log2 read number
(−) total RNA
(+) with primary 5'ends
log2 expression
21.3
18.3
15.3
12.3
9.3
−9.3
−13.3
−17.3
−21.3
12
10
8
6
4
2
0
slr0541
clpP
slr0542
trpB
slr0543
slr0544
3211250
3212500
3213750
3216250
3218750
3210000
3215000
3217500
3220000
502
gS
sll0501
sll0499
sll0498
sll0497
Position (nt)

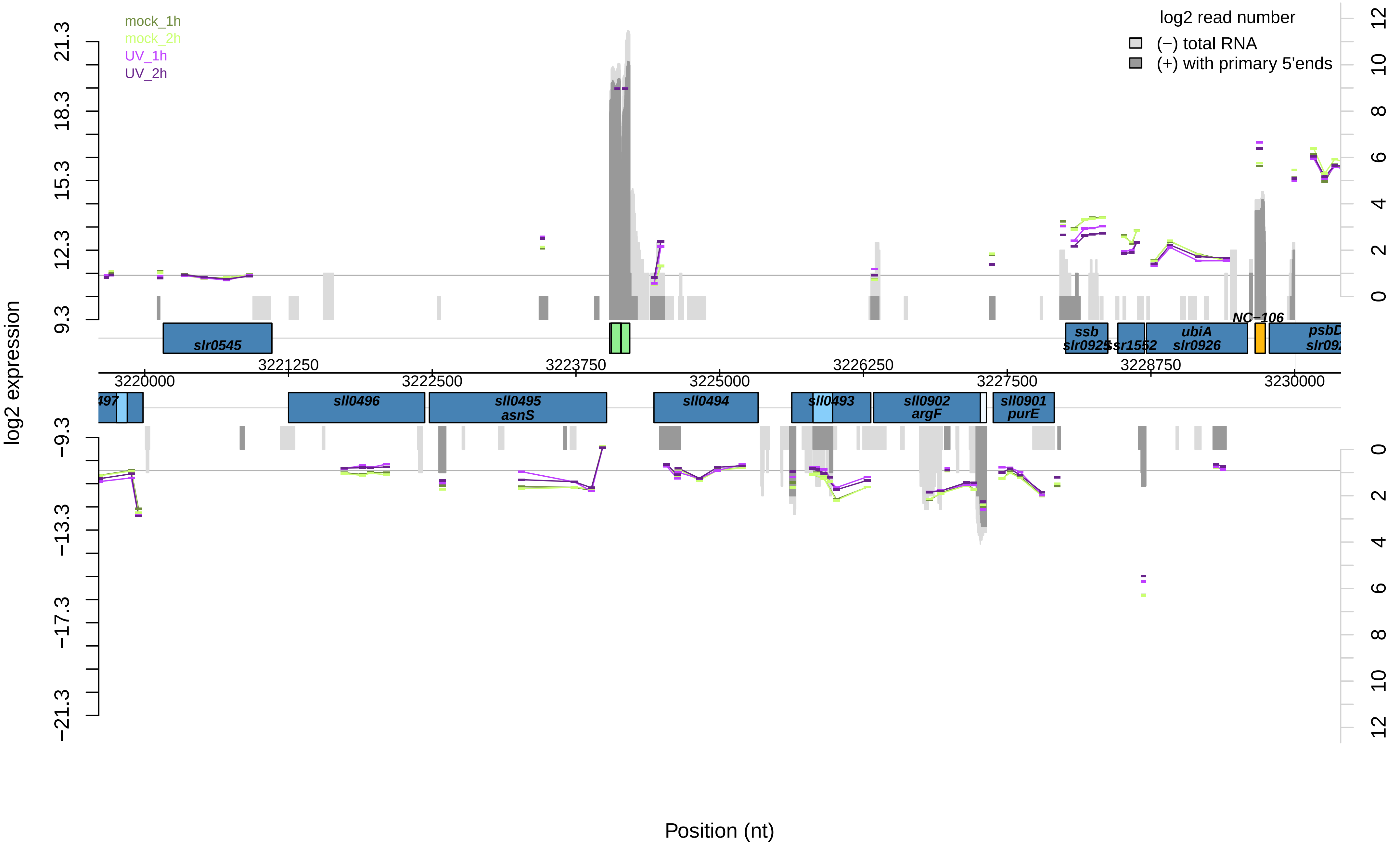

mock_1h
mock_2h
UV_1h
UV_2h
log2 read number
(−) total RNA
(+) with primary 5'ends
log2 expression
Position (nt)
slr0545
ssb
slr0925
ssr1552
ubiA
slr0926
NC−106
sll0496
sll0495
asnS
sll0494
sll0493
sll0902
argF
sll0901
purE
3220000
3221250
3222500
3223750
3225000
3226250
3227500
3228750
3230000
21.3
18.3
15.3
12.3
9.3
−9.3
−13.3
−17.3
−21.3

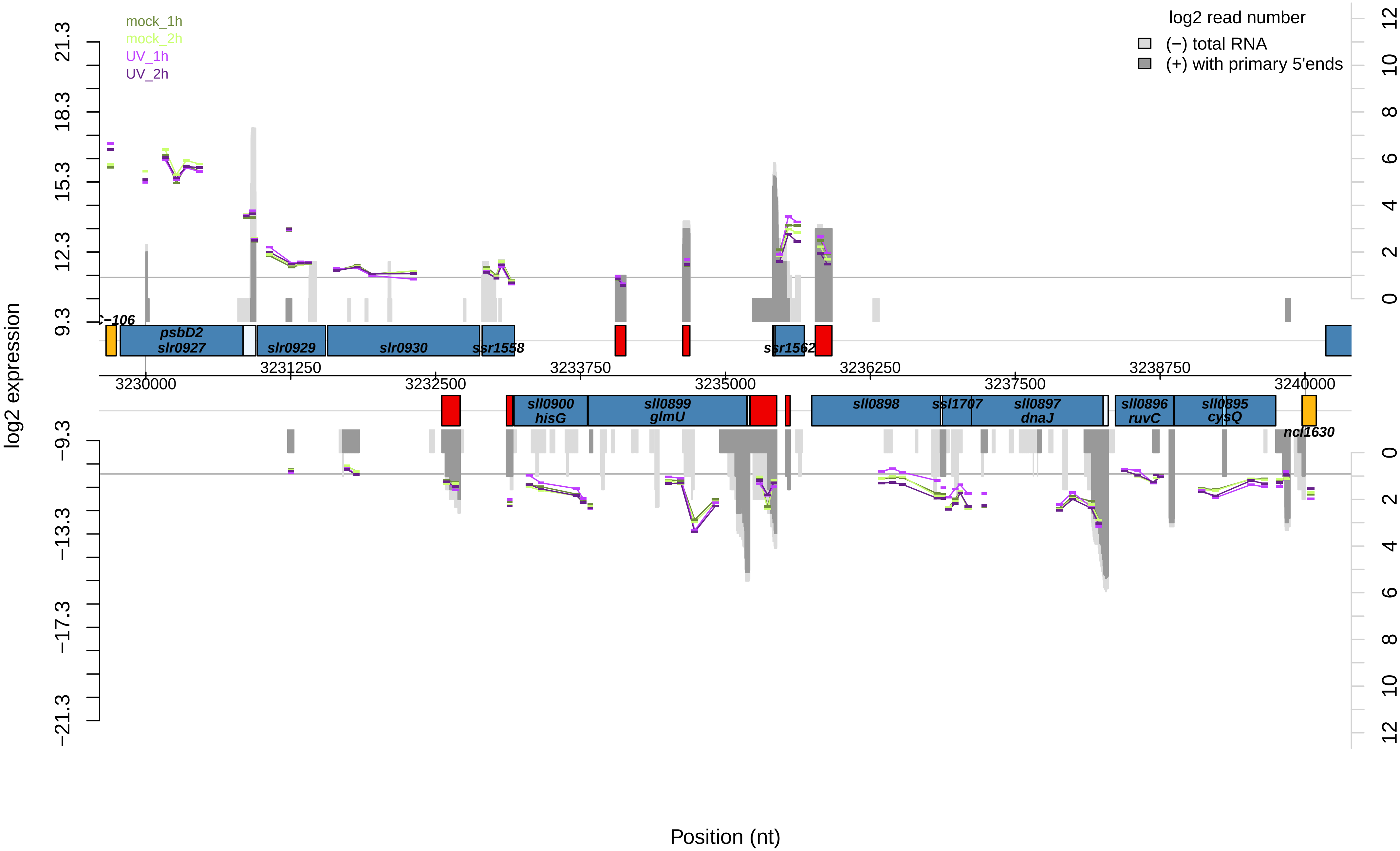
mock_1h
mock_2h
UV_1h
UV_2h
log2 read number
(−) total RNA
(+) with primary 5'ends
log2 expression
Position (nt)
psbD2
slr0927
slr0929
slr0930
ssr1558
ssr1562
sll0900
hisG
sll0899
glmU
sll0898
ssl1707
sll0897
dnaJ
sll0896
ruvC
sll0895
cysQ
ncl1630
3230000
3231250
3232500
3233750
3235000
3236250
3237500
3238750
3240000

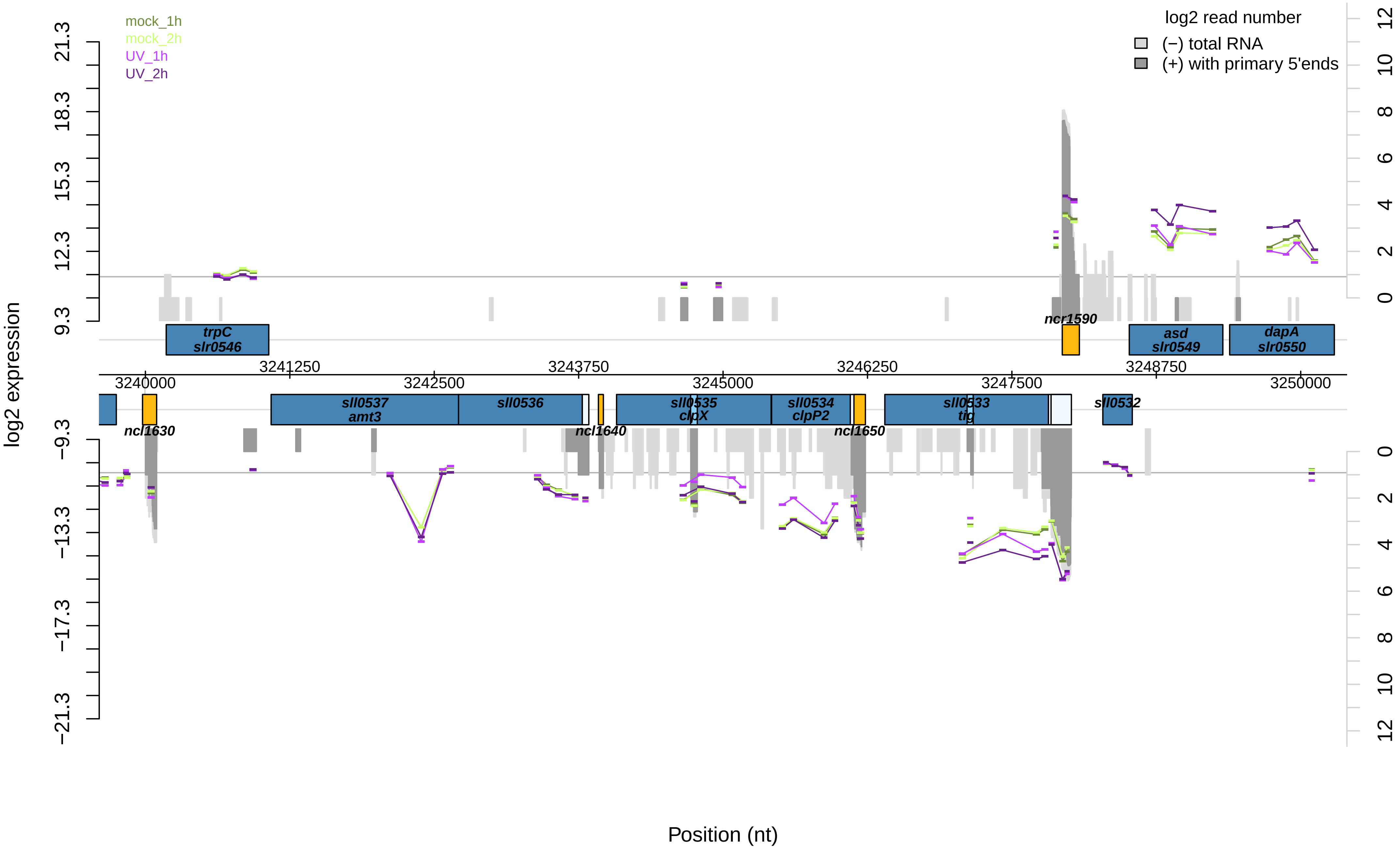

mock_1h
mock_2h
UV_1h
UV_2h
log2 read number
(−) total RNA
(+) with primary 5'ends
log2 expression
21.3
18.3
15.3
12.3
9.3
−9.3
−13.3
−17.3
−21.3
trpC
slr0546
ncr1590
asd
slr0549
dapA
slr0550
3240000
3241250
3242500
3243750
3245000
3246250
3247500
3248750
3250000
ncl1630
sll0537
amt3
sll0536
ncl1640
sll0535
clpX
sll0534
clpP2
ncl1650
sll0533
tig
sll0532
Position (nt)

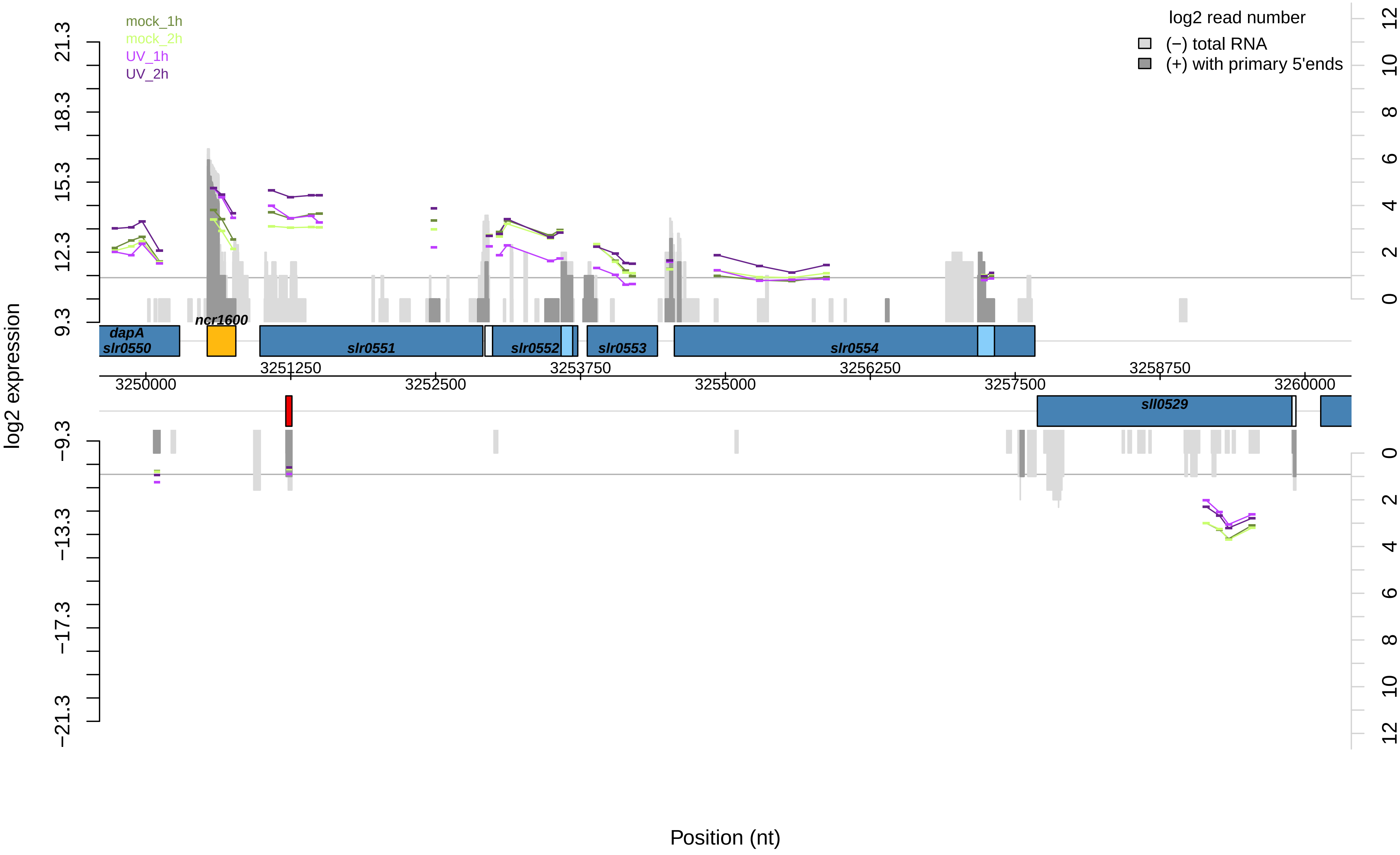

mock_1h
mock_2h
UV_1h
UV_2h
log2 read number
(−) total RNA
(+) with primary 5'ends
log2 expression
21.3
18.3
15.3
12.3
9.3
−9.3
−13.3
−17.3
−21.3
12
10
8
6
4
2
0
dapA
slr0550
ncr1600
slr0551
slr0552
slr0553
slr0554
sll0529
3250000
3251250
3252500
3253750
3255000
3256250
3257500
3258750
3260000
Position (nt)

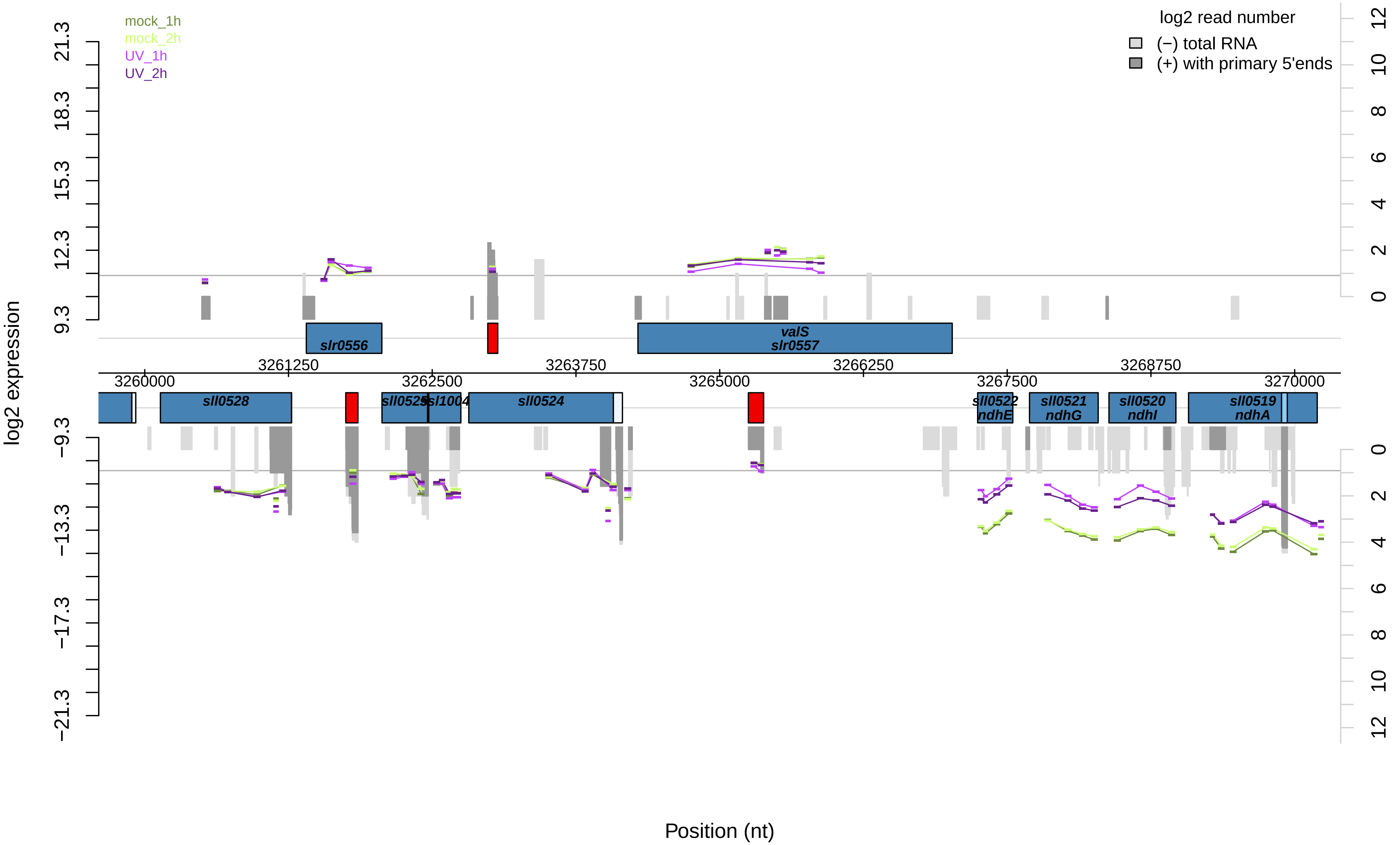
mock_1h
mock_2h
UV_1h
UV_2h
log2 read number
(−) total RNA
(+) with primary 5'ends
log2 expression
21.3
18.3
15.3
12.3
9.3
−9.3
−13.3
−17.3
−21.3
12
10
8
6
4
2
0
slr0556
valS
slr0557
3261250
3262500
3263750
3265000
3266250
3267500
3268750
3260000
3270000
sll0528
sll0525
sll1004
sll0524
sll0522
ndhE
sll0521
ndhG
sll0520
ndhI
sll0519
ndhA
Position (nt)

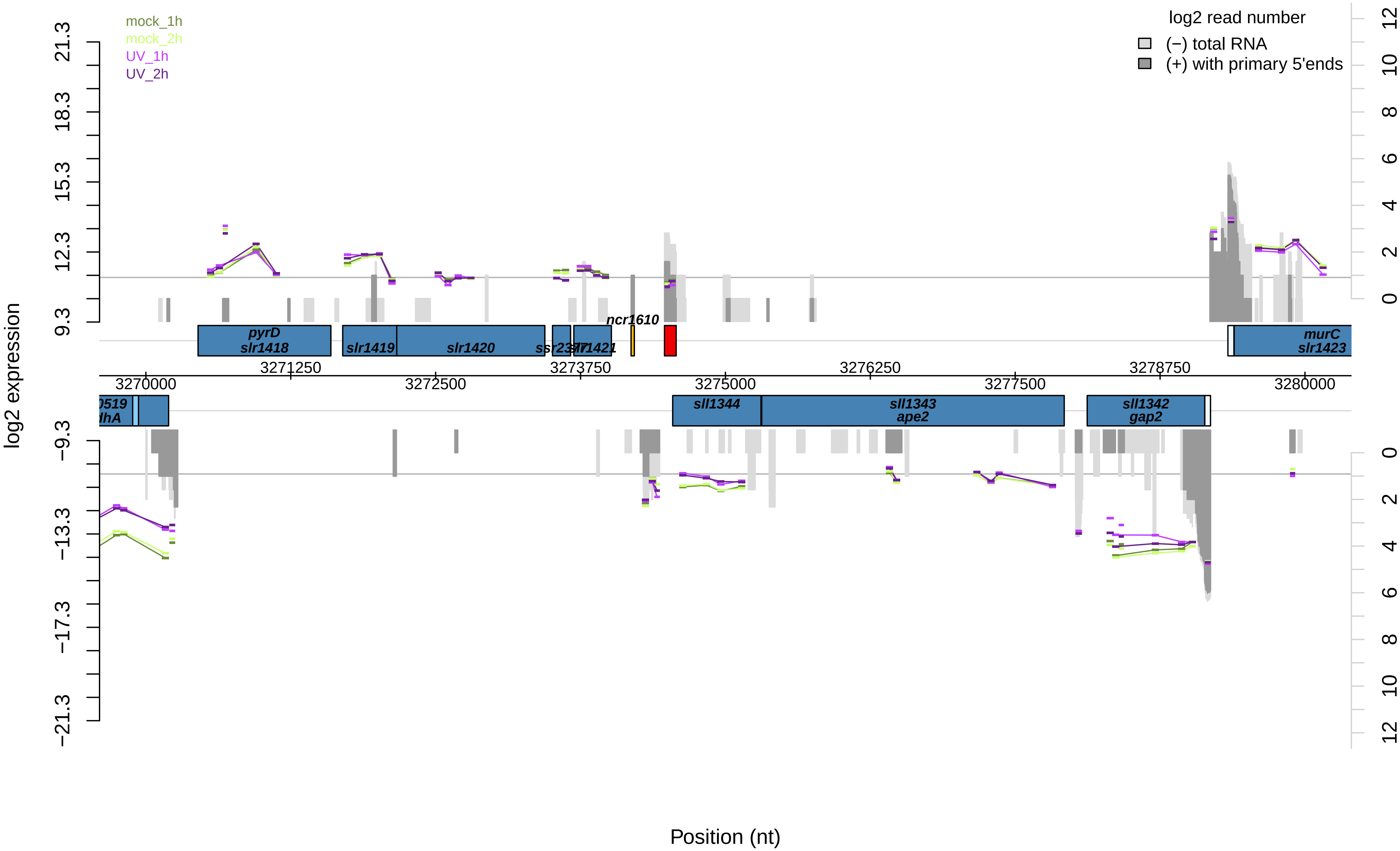
mock_1h
mock_2h
UV_1h
UV_2h
log2 read number
(−) total RNA
(+) with primary 5'ends
log2 expression
21.3
18.3
15.3
12.3
9.3
−9.3
−13.3
−17.3
−21.3
12
10
8
6
4
2
0
pyrD
slr1418
slr1419
slr1420
slr1421
ncr1610
murC
slr1423
3271250
3273750
3276250
3278750
3270000
3272500
3275000
3277500
3280000
sll1344
sll1343
ape2
sll1342
gap2
Position (nt)

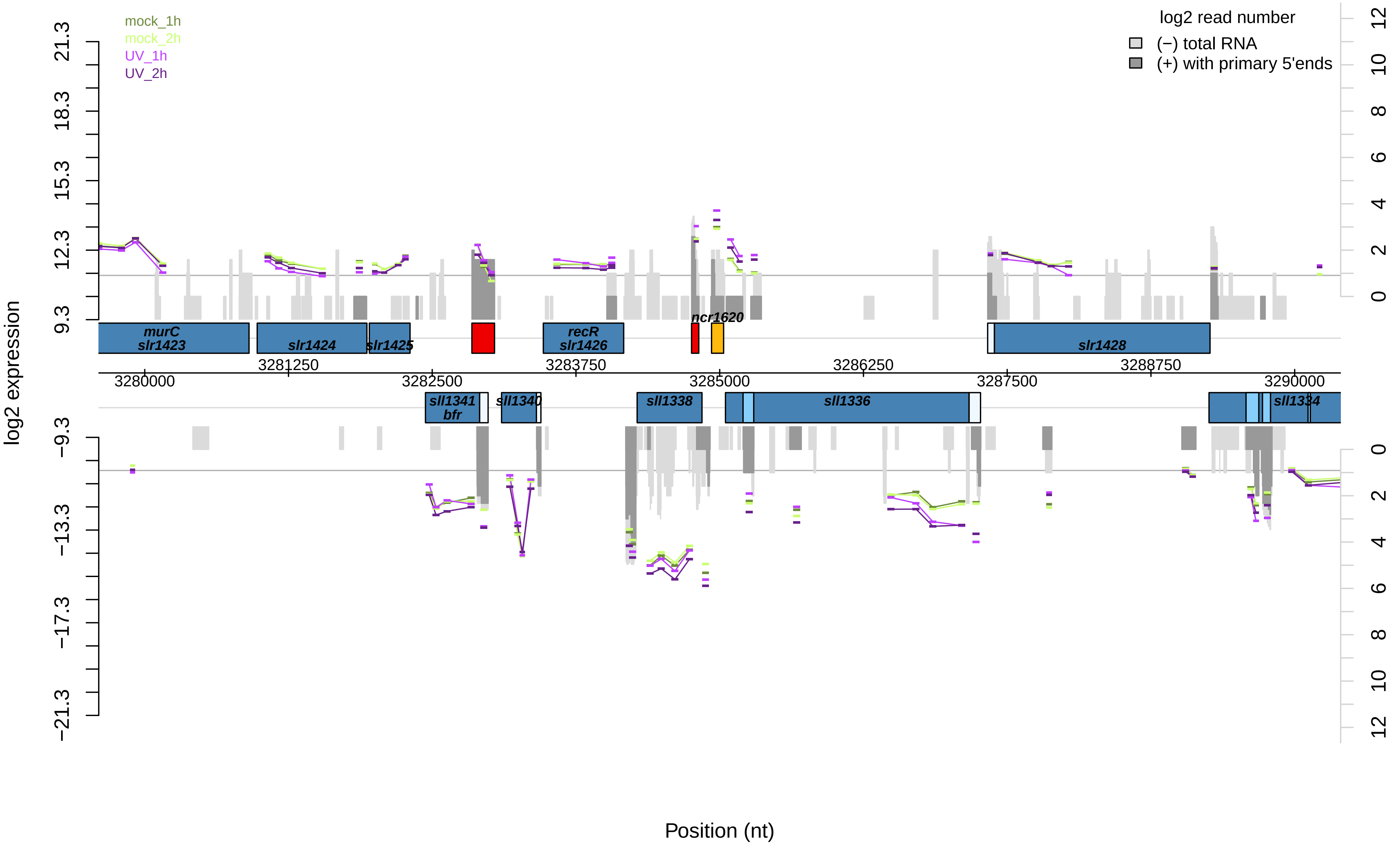

mock_1h
mock_2h
UV_1h
UV_2h
log2 read number
(−) total RNA
(+) with primary 5'ends
log2 expression
Position (nt)
21.3
18.3
15.3
12.3
9.3
−9.3
−13.3
−17.3
−21.3
0
2
4
6
8
10
12
murC
slr1423
slr1424
slr1425
recR
slr1426
ncr1620
slr1428
sll1341
bfr
sll1340
sll1338
sll1336
sll1334
3280000
3281250
3282500
3283750
3285000
3286250
3287500
3288750
3290000

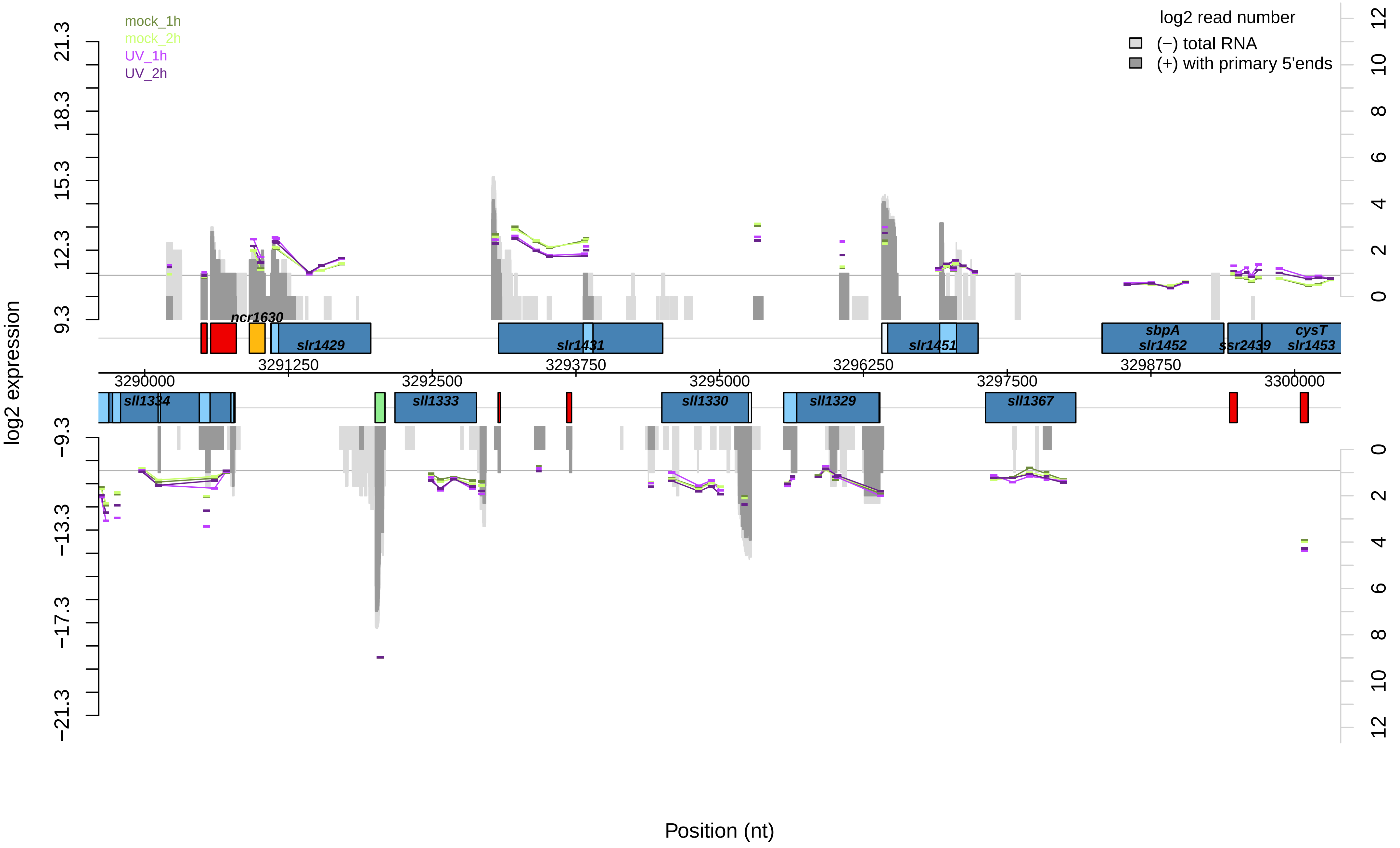
mock_1h
mock_2h
UV_1h
UV_2h
log2 read number
(−) total RNA
(+) with primary 5'ends
log2 expression
Position (nt)
21.3
18.3
15.3
12.3
9.3
−9.3
−13.3
−17.3
−21.3
12
10
8
6
4
2
0
ncr1630
slr1429
slr1431
slr1451
sbpA
slr1452
ssr2439
cysT
slr1453
3291250
3293750
3296250
3298750
3290000
3292500
3295000
3297500
3300000
sll1334
sll1333
sll1330
sll1329
sll1367

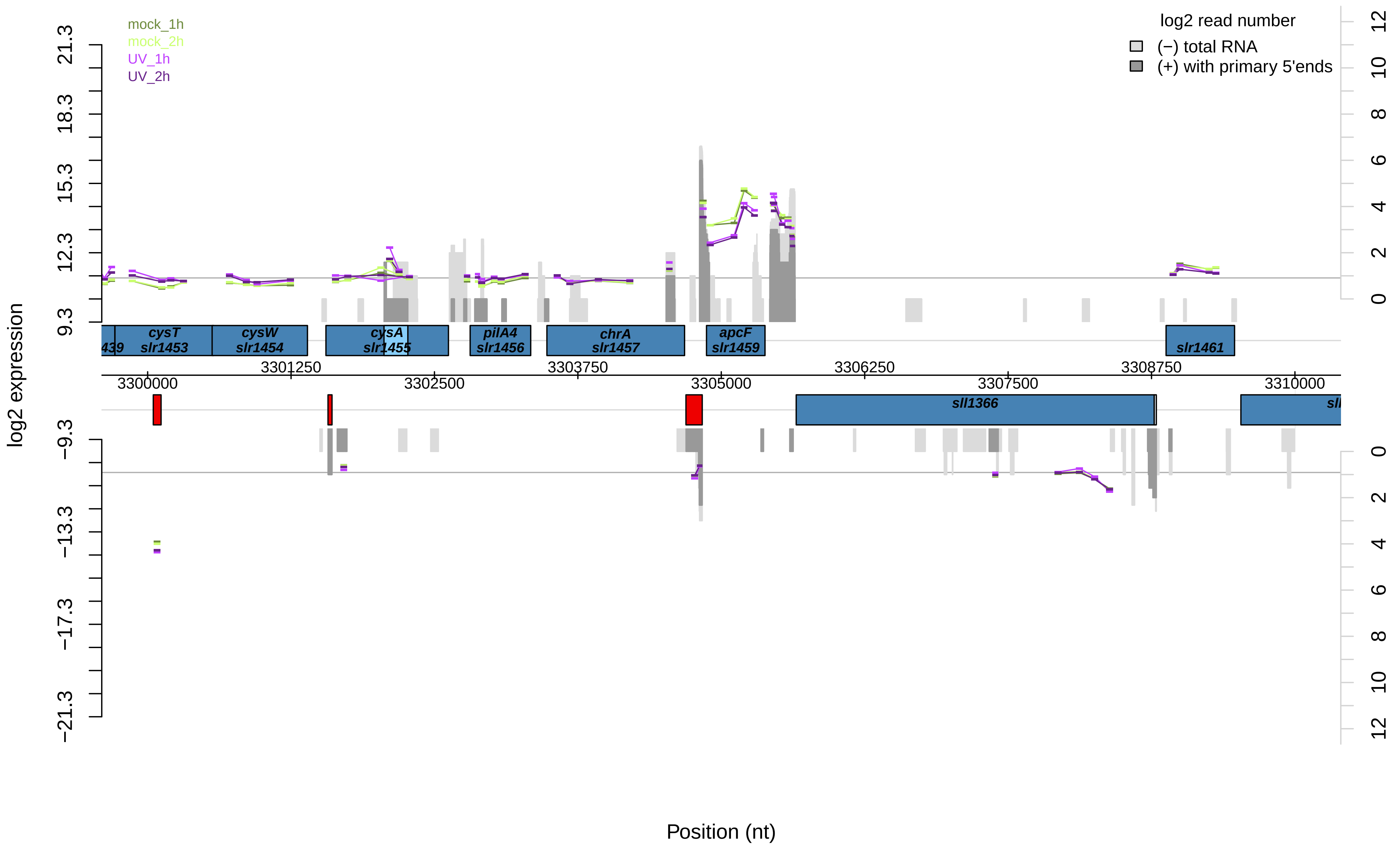

mock_1h
mock_2h
UV_1h
UV_2h
log2 read number
(−) total RNA
(+) with primary 5'ends
log2 expression
21.3
18.3
15.3
12.3
9.3
−9.3
−13.3
−17.3
−21.3
12
10
8
6
4
2
0
439
cysT
slr1453
cysW
slr1454
cysA
slr1455
pilA4
slr1456
chrA
slr1457
apcF
slr1459
slr1461
sll1366
3300000
3301250
3302500
3303750
3305000
3306250
3307500
3308750
3310000
Position (nt)

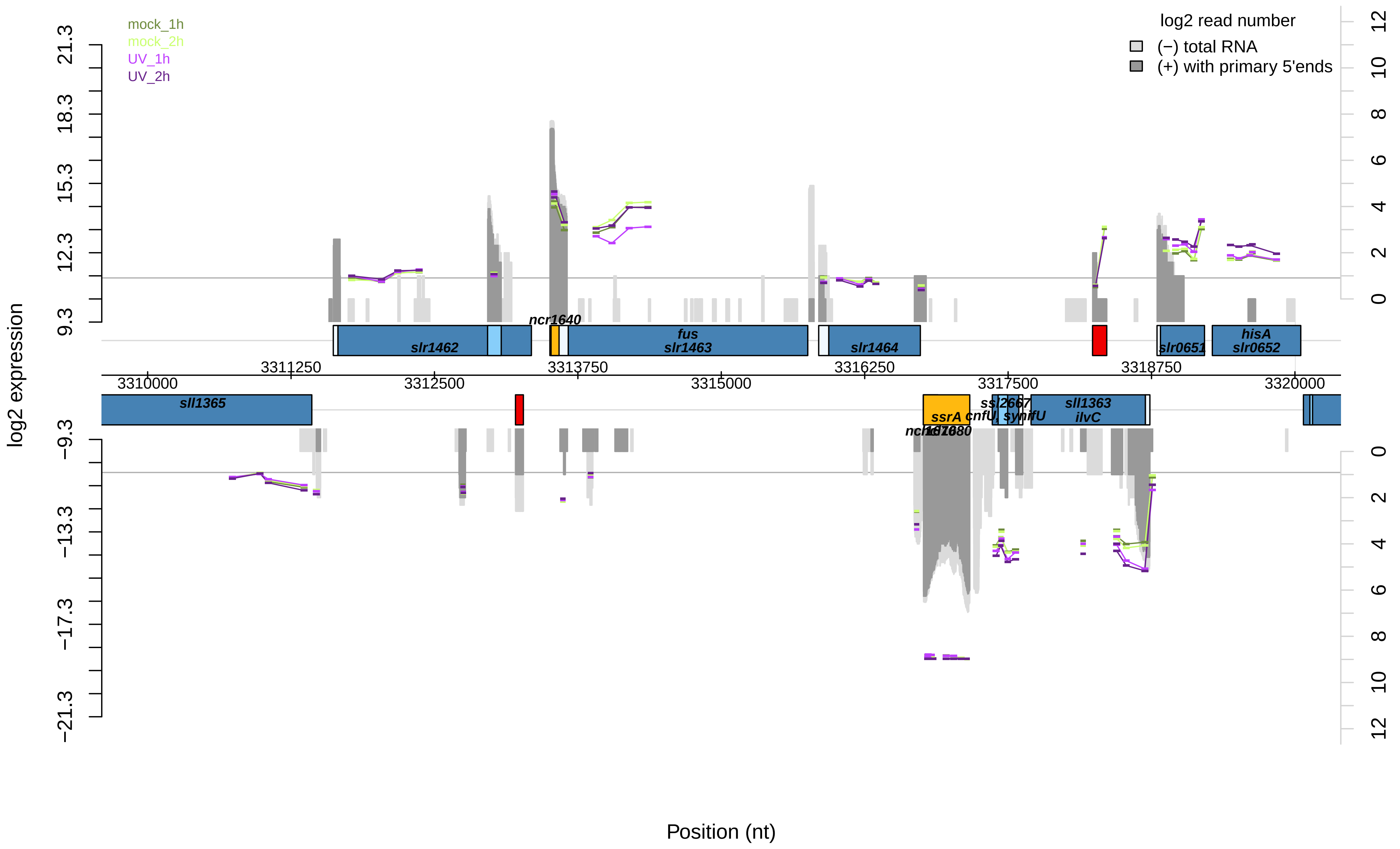

mock_1h
mock_2h
UV_1h
UV_2h
log2 read number
(−) total RNA
(+) with primary 5'ends
log2 expression
21.3
18.3
15.3
12.3
9.3
−9.3
−13.3
−17.3
−21.3
12
10
8
6
4
2
0
ncr1640
slr1462
fus
slr1463
slr1464
slr0651
hisA
slr0652
3311250
3313750
3316250
3317500
3318750
3310000
3312500
3315000
3320000
sll1365
ssrA
ssl2667
cnfU
synifU
sll1363
ilvC
Position (nt)

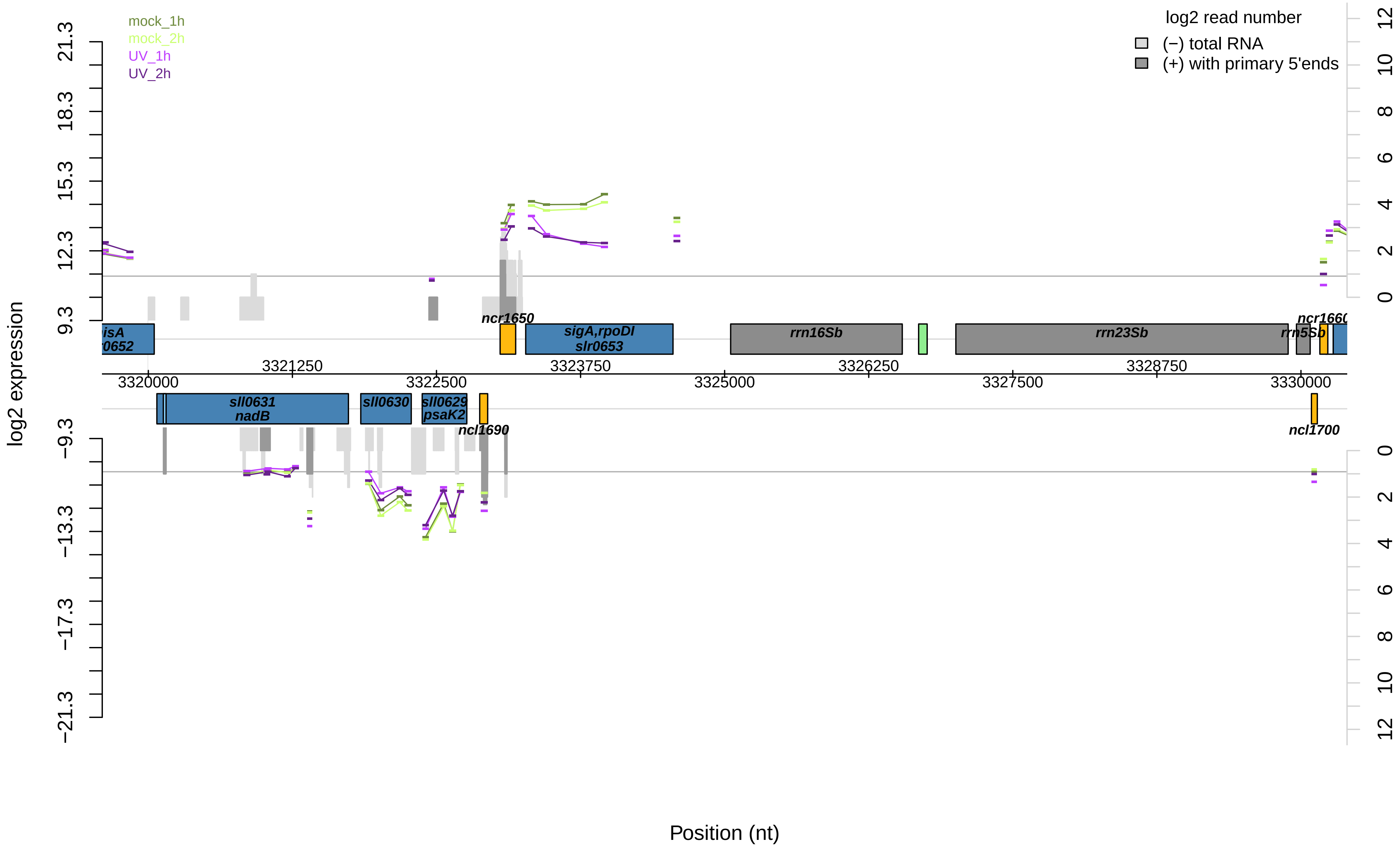

mock_1h
mock_2h
UV_1h
UV_2h
log2 read number
(−) total RNA
(+) with primary 5'ends
log2 expression
21.3
18.3
15.3
12.3
9.3
−9.3
−13.3
−17.3
−21.3
12
10
8
6
4
2
0
isA
r0652
ncr1650
sigA,rpoDI
slr0653
rrn16Sb
rrn23Sb
rrn5Sb
ncr1660
3321250
3323750
3326250
3328750
3320000
3322500
3325000
3327500
3330000
sll0631
nadB
sll0630
sll0629
psaK2
ncl1690
ncl1700
Position (nt)

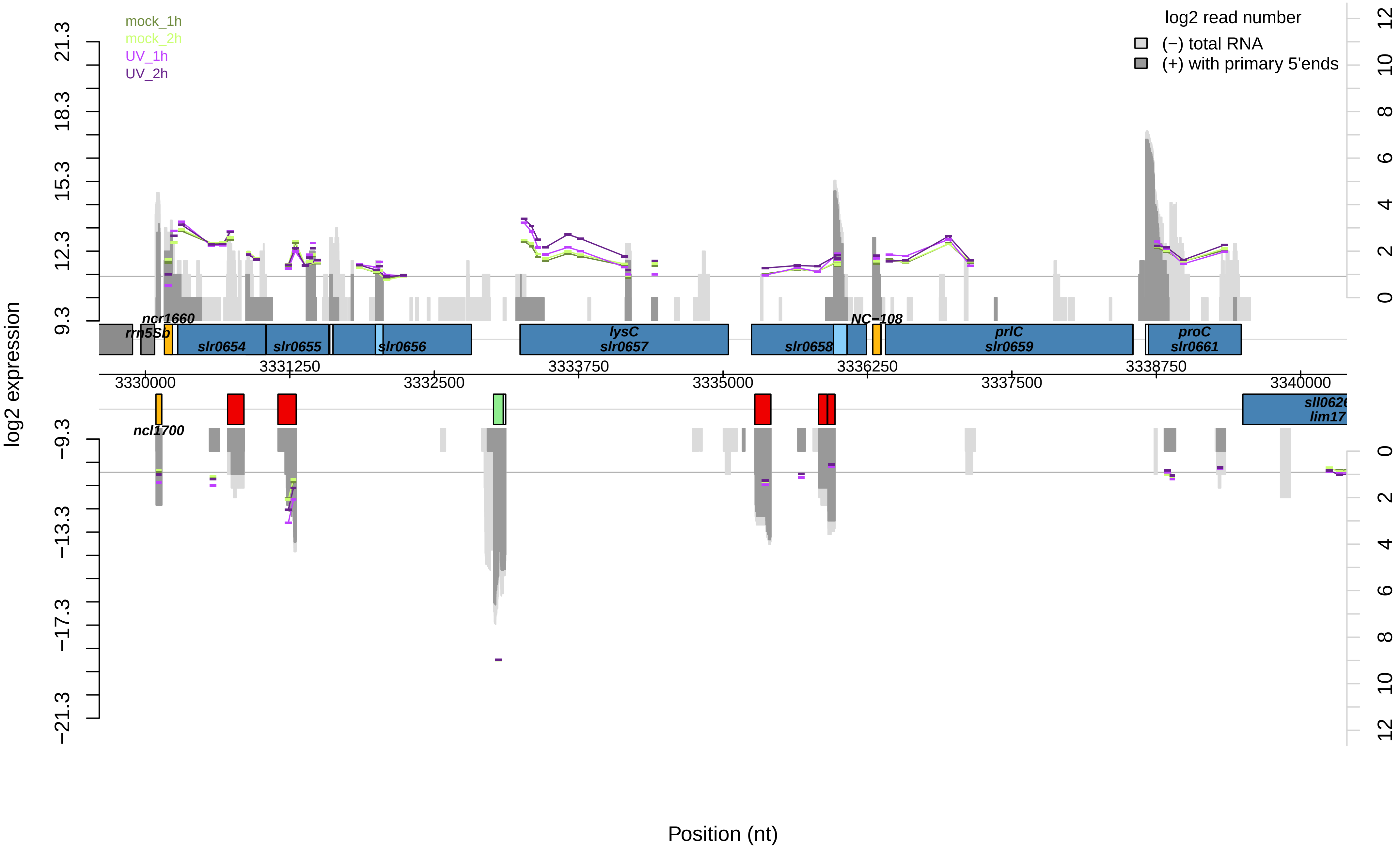

mock_1h
mock_2h
UV_1h
UV_2h
log2 read number
(−) total RNA
(+) with primary 5'ends
log2 expression
21.3
18.3
15.3
12.3
9.3
−9.3
−13.3
−17.3
−21.3
12
10
8
6
4
2
0
ncr1660
rrn5Sb
slr0654
slr0655
slr0656
lysC
slr0657
slr0658
NC−108
prlC
slr0659
proC
slr0661
sll0626
lim17
ncl1700
3330000
3331250
3332500
3333750
3335000
3336250
3337500
3338750
3340000
Position (nt)

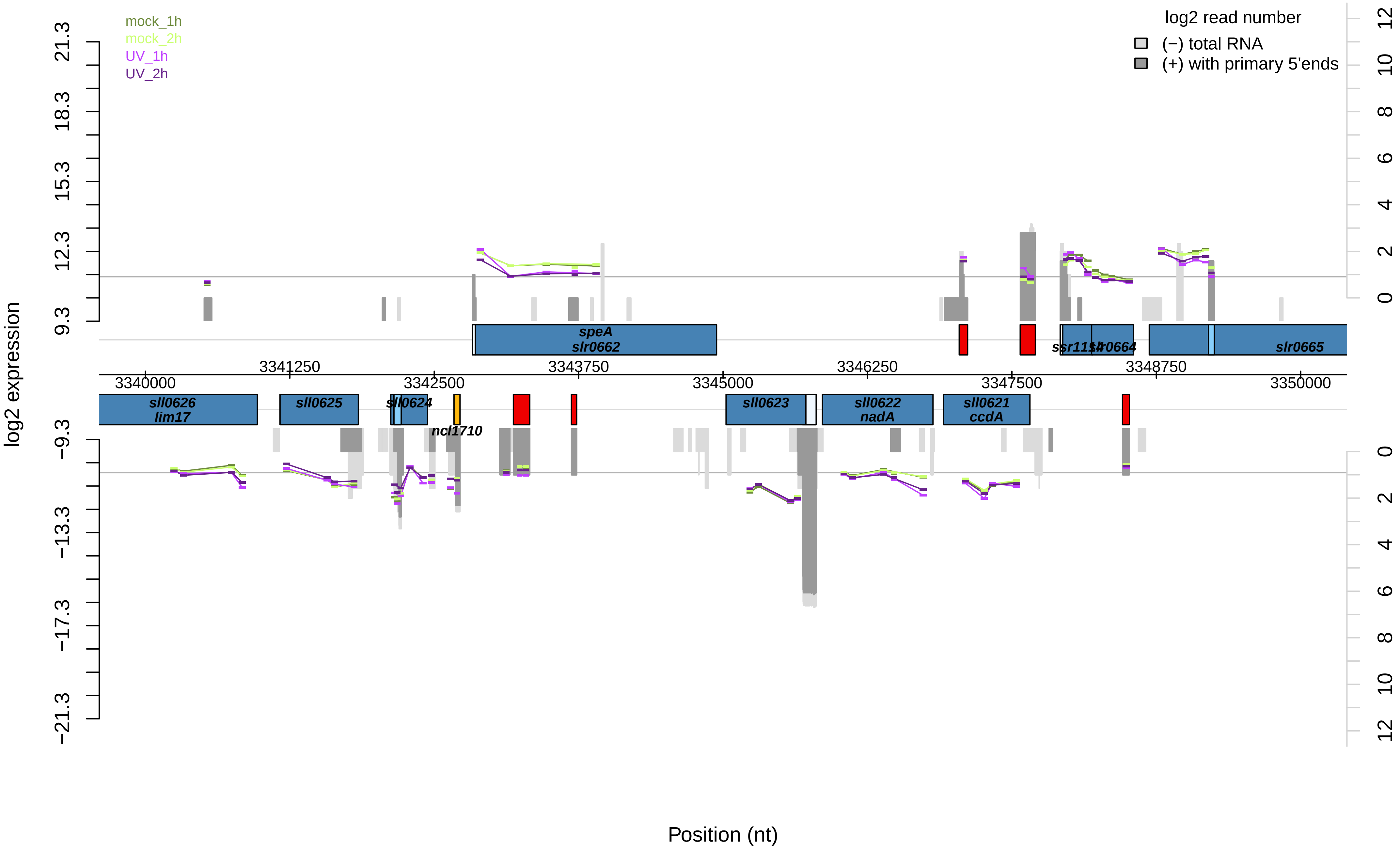

mock_1h
mock_2h
UV_1h
UV_2h
log2 read number
(−) total RNA
(+) with primary 5'ends
log2 expression
21.3
18.3
15.3
12.3
9.3
−9.3
−13.3
−17.3
−21.3
12
10
8
6
4
2
0
speA
slr0662
ssr1154
slr0664
slr0665
3341250
3343750
3346250
3348750
3340000
3342500
3345000
3347500
3350000
sll0626
lim17
sll0625
sll0624
ncl1710
sll0623
sll0622
nadA
sll0621
ccdA
Position (nt)

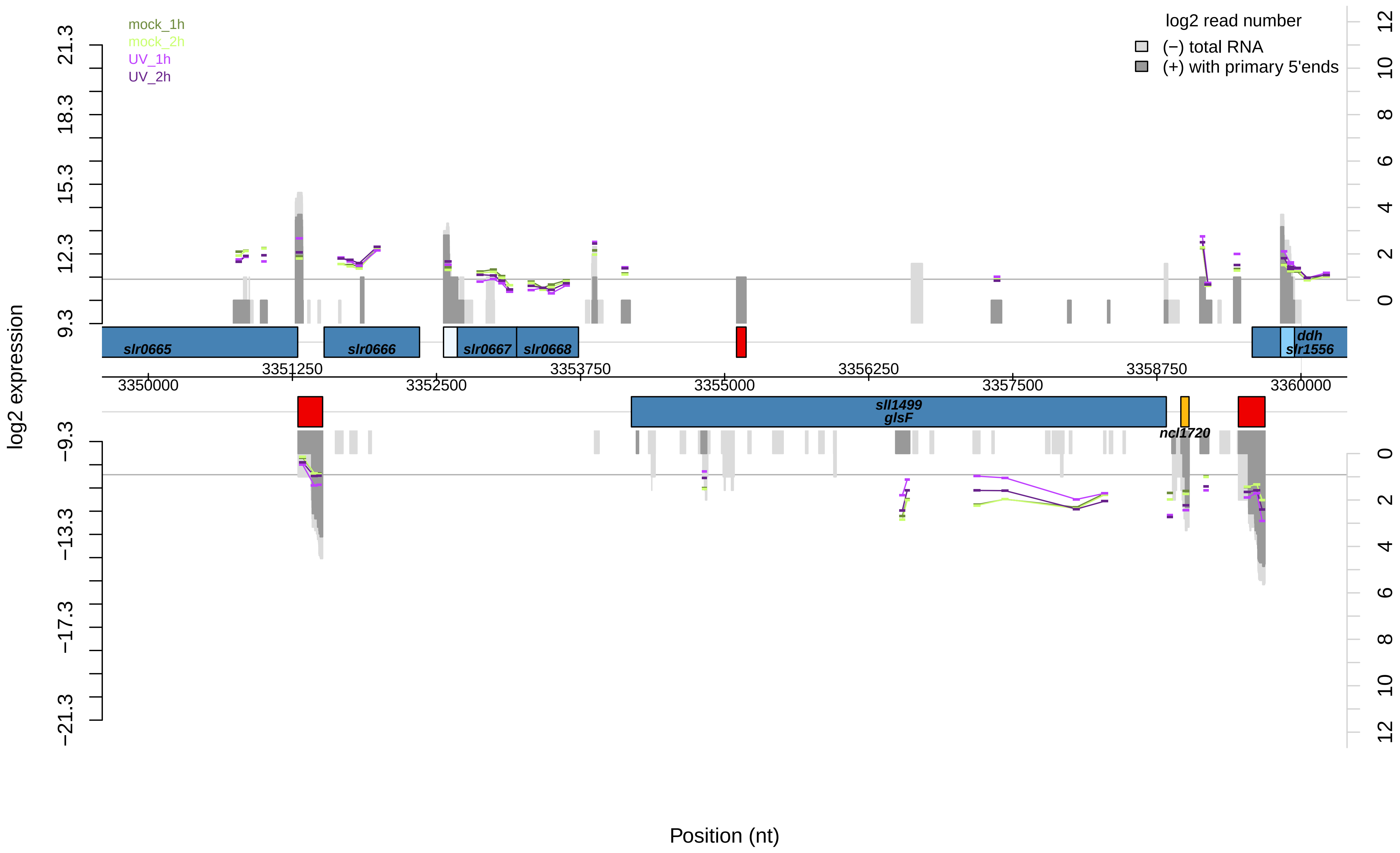

mock_1h
mock_2h
UV_1h
UV_2h
log2 read number
(−) total RNA
(+) with primary 5'ends
log2 expression
21.3
18.3
15.3
12.3
9.3
−9.3
−13.3
−17.3
−21.3
12
10
8
6
4
2
0
slr0665
slr0666
slr0667
slr0668
ddh
slr1556
sll1499
glsF
ncl1720
3350000
3351250
3352500
3353750
3355000
3356250
3357500
3358750
3360000
Position (nt)

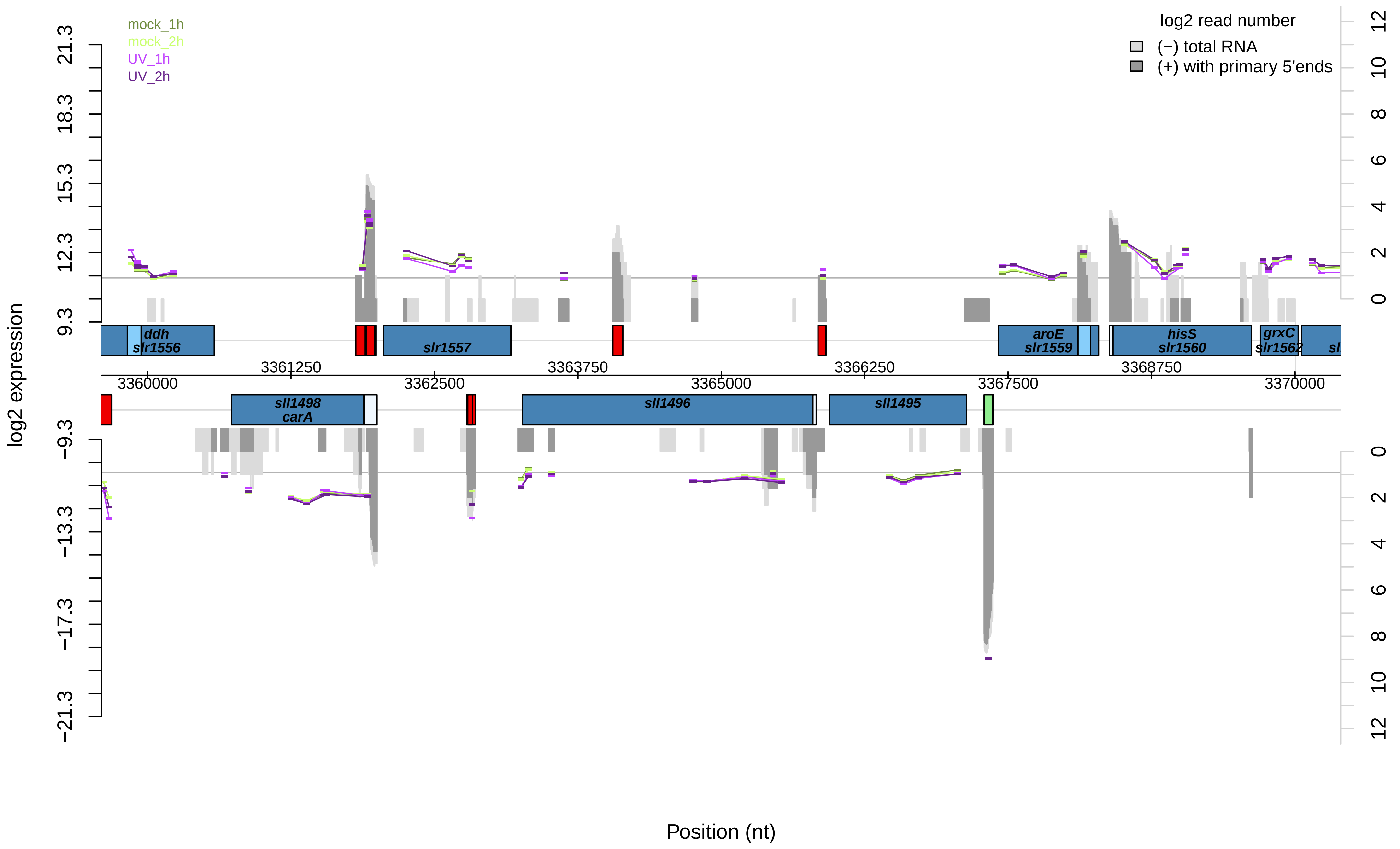

mock_1h
mock_2h
UV_1h
UV_2h
log2 read number
(−) total RNA
(+) with primary 5'ends
log2 expression
21.3
18.3
15.3
12.3
9.3
−9.3
−13.3
−17.3
−21.3
12
10
8
6
4
2
0
ddh
slr1556
slr1557
aroE
slr1559
hisS
slr1560
grxC
slr1562
sll1498
carA
sll1496
sll1495
3360000
3361250
3362500
3363750
3365000
3366250
3367500
3368750
3370000
Position (nt)

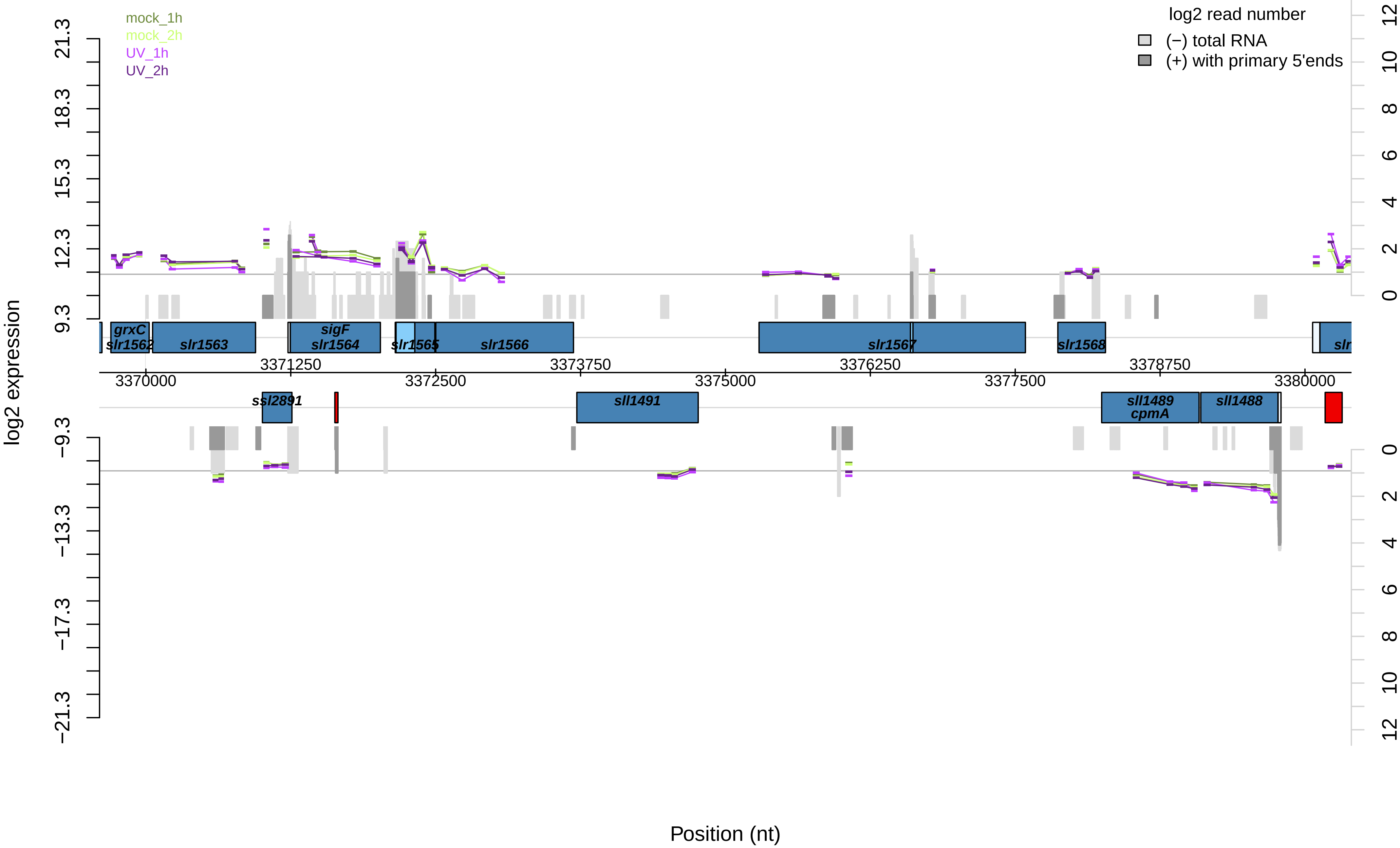

mock_1h
mock_2h
UV_1h
UV_2h
log2 read number
(−) total RNA
(+) with primary 5'ends
log2 expression
21.3
18.3
15.3
12.3
9.3
−9.3
−13.3
−17.3
−21.3
12
10
8
6
4
2
0
grxC
slr1562
slr1563
sigF
slr1564
slr1565
slr1566
slr1567
slr1568
3371250
3373750
3376250
3378750
3370000
3372500
3375000
3377500
3380000
ssl2891
sll1491
sll1489
cpmA
sll1488
Position (nt)

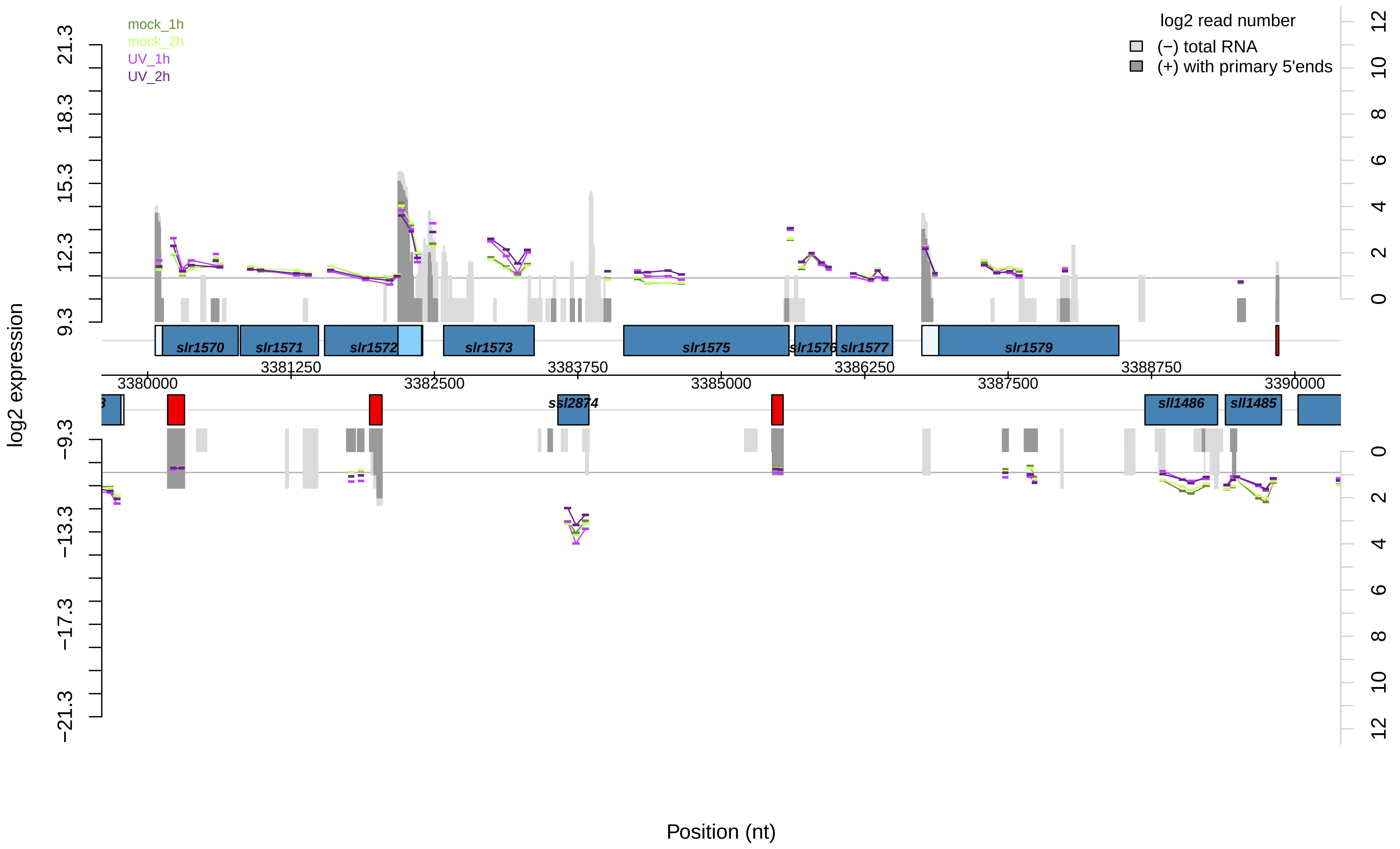

mock_1h
mock_2h
UV_1h
UV_2h
log2 read number
(−) total RNA
(+) with primary 5'ends
log2 expression
21.3
18.3
15.3
12.3
9.3
−9.3
−13.3
−17.3
−21.3
12
10
8
6
4
2
0
slr1570
slr1571
slr1572
slr1573
slr1575
slr1576
slr1577
slr1579
3381250
3383750
3386250
3388750
3380000
3382500
3385000
3387500
3390000
ssl2874
sll1486
sll1485
Position (nt)

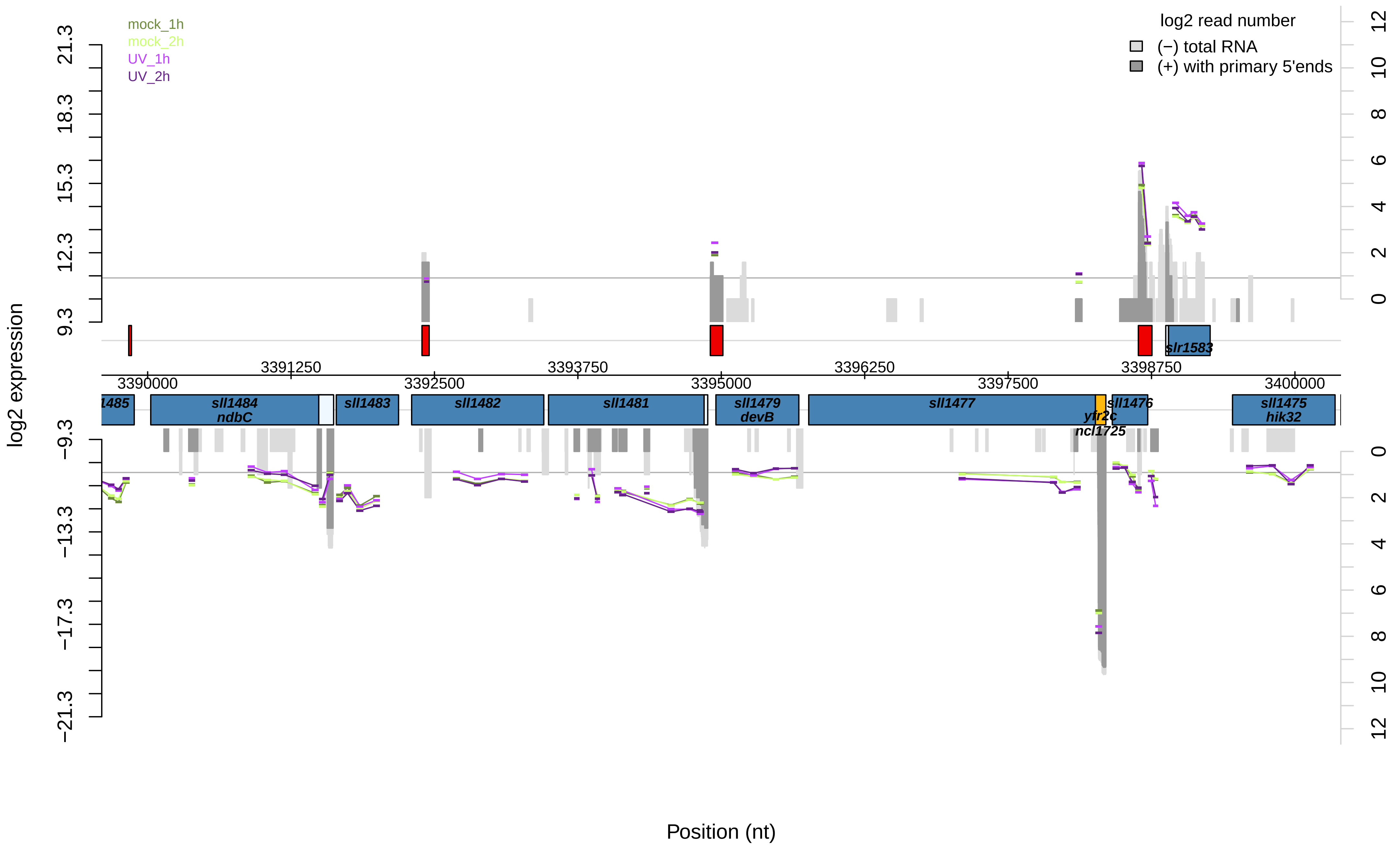

mock_1h
mock_2h
UV_1h
UV_2h
log2 read number
(−) total RNA
(+) with primary 5'ends
log2 expression
21.3
18.3
15.3
12.3
9.3
−9.3
−13.3
−17.3
−21.3
12
10
8
6
4
2
0
slr1583
3391250
3393750
3396250
3398750
3390000
3392500
3395000
3397500
3400000
1485
sll1484
ndbC
sll1483
sll1482
sll1481
sll1479
devB
sll1477
yfr2c
ncl1725
sll1476
sll1475
hik32
Position (nt)

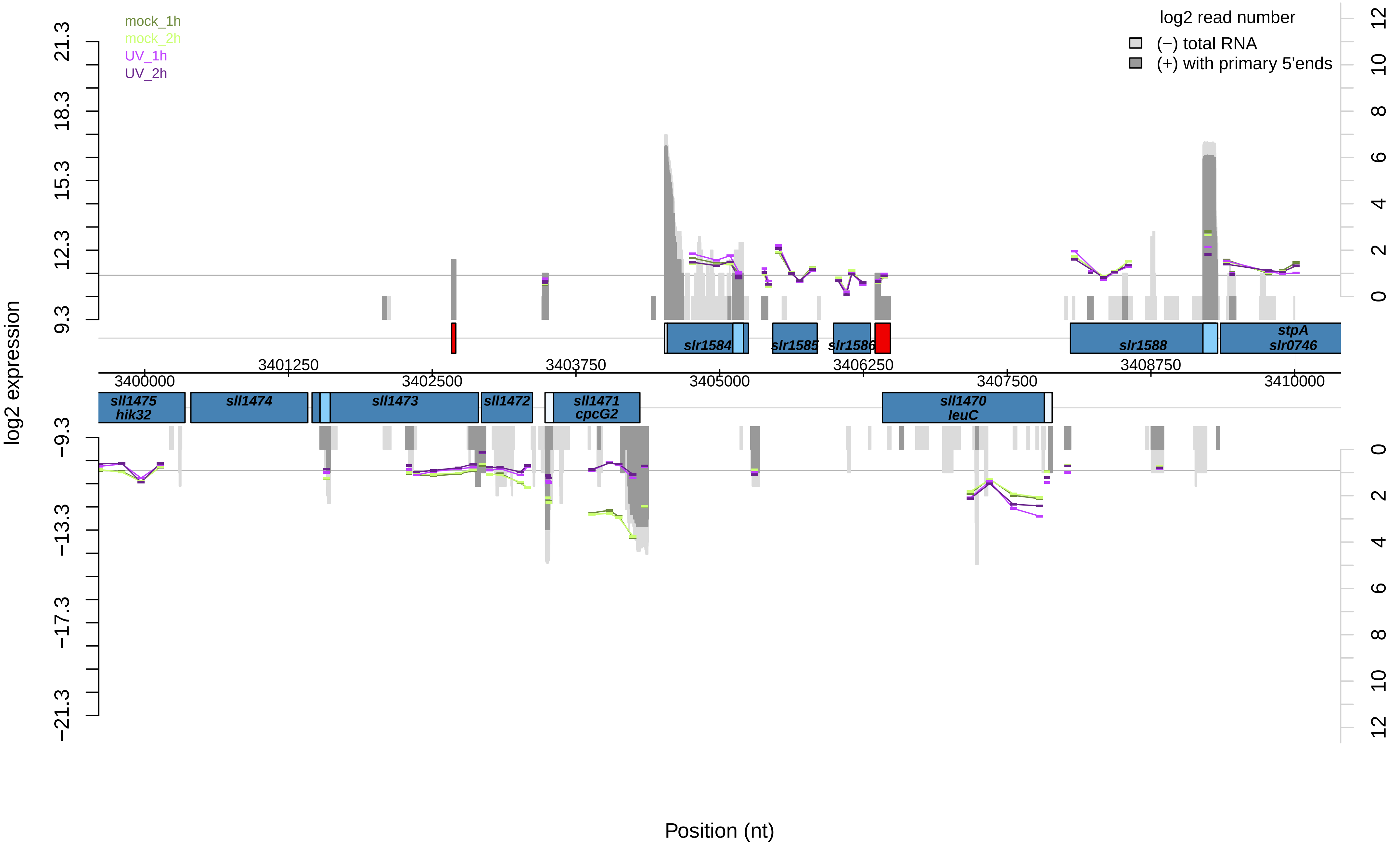

mock_1h
mock_2h
UV_1h
UV_2h
log2 read number
(−) total RNA
(+) with primary 5'ends
log2 expression
21.3
18.3
15.3
12.3
9.3
−9.3
−13.3
−17.3
−21.3
12
10
8
6
4
2
0
slr1584
slr1585
slr1586
slr1588
stpA
slr0746
3401250
3403750
3406250
3408750
3400000
3402500
3405000
3407500
3410000
sll1475
hik32
sll1474
sll1473
sll1472
sll1471
cpcG2
sll1470
leuC
Position (nt)

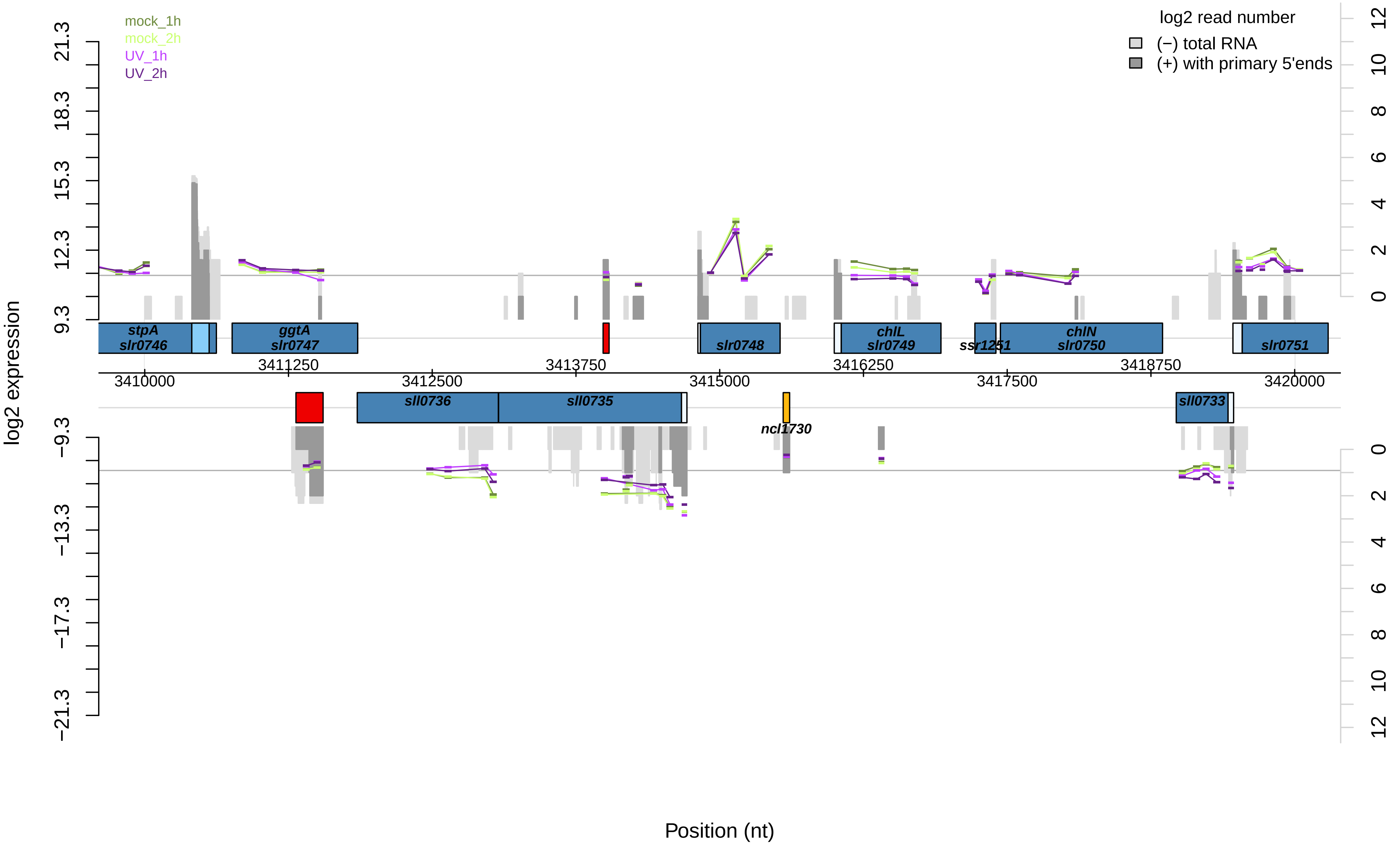

mock_1h
mock_2h
UV_1h
UV_2h
log2 read number
(−) total RNA
(+) with primary 5'ends
log2 expression
21.3
18.3
15.3
12.3
9.3
−9.3
−13.3
−17.3
−21.3
12
10
8
6
4
2
0
stpA
slr0746
ggtA
slr0747
slr0748
chlL
slr0749
ssr1251
chlN
slr0750
slr0751
3410000
3411250
3412500
3413750
3415000
3416250
3417500
3418750
3420000
sll0736
sll0735
ncl1730
sll0733
Position (nt)

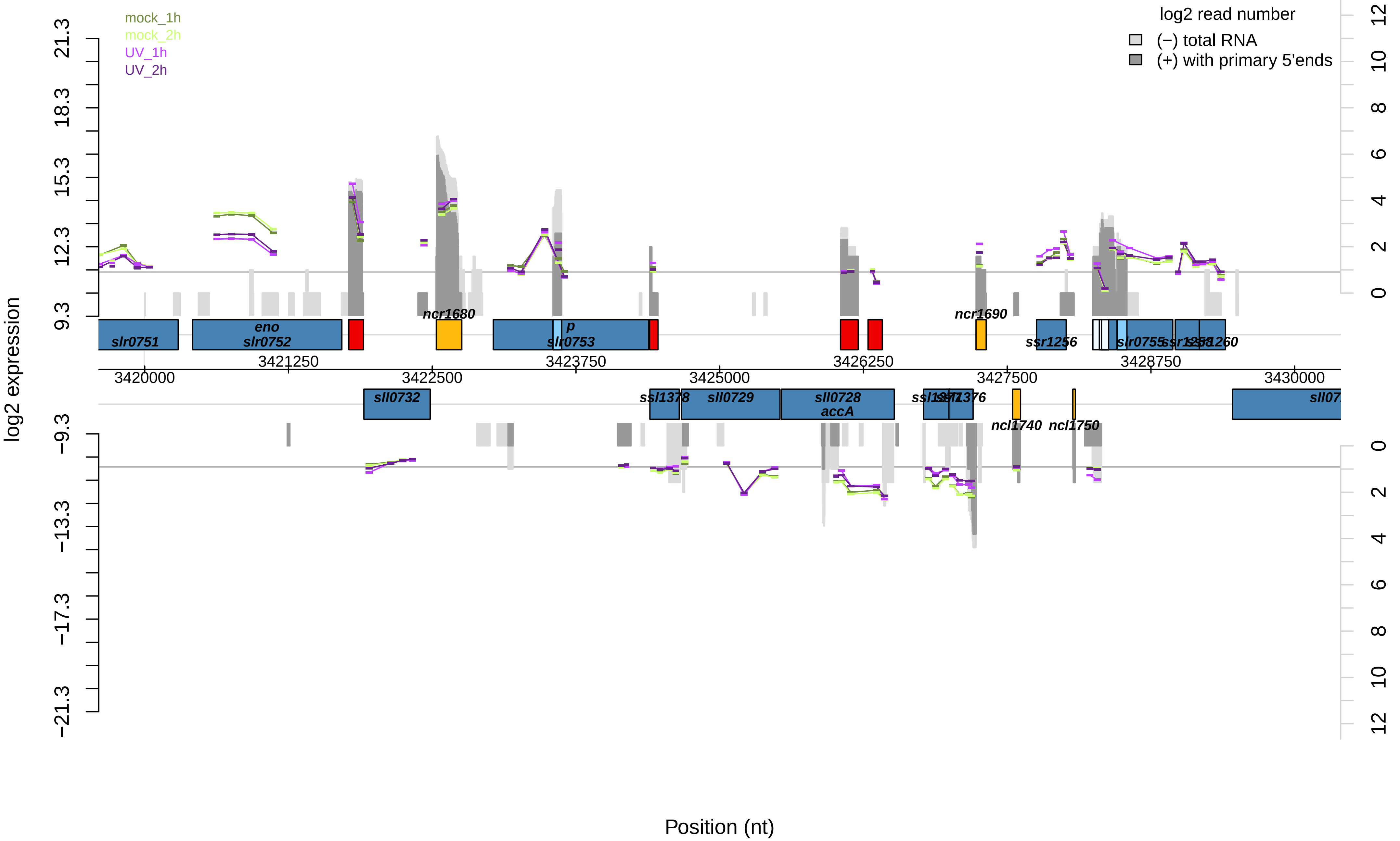

mock_1h
mock_2h
UV_1h
UV_2h
log2 read number
(−) total RNA
(+) with primary 5'ends
log2 expression
21.3
18.3
15.3
12.3
9.3
−9.3
−13.3
−17.3
−21.3
12
10
8
6
4
2
0
ncr1680
ncr1690
eno
slr0752
slr0751
p
slr0753
ssr1256
slr0755
3421250
3423750
3426250
3428750
3420000
3422500
3425000
3427500
3430000
sll0732
ssl1378
sll0729
sll0728
accA
ncl1740
ncl1750
Position (nt)

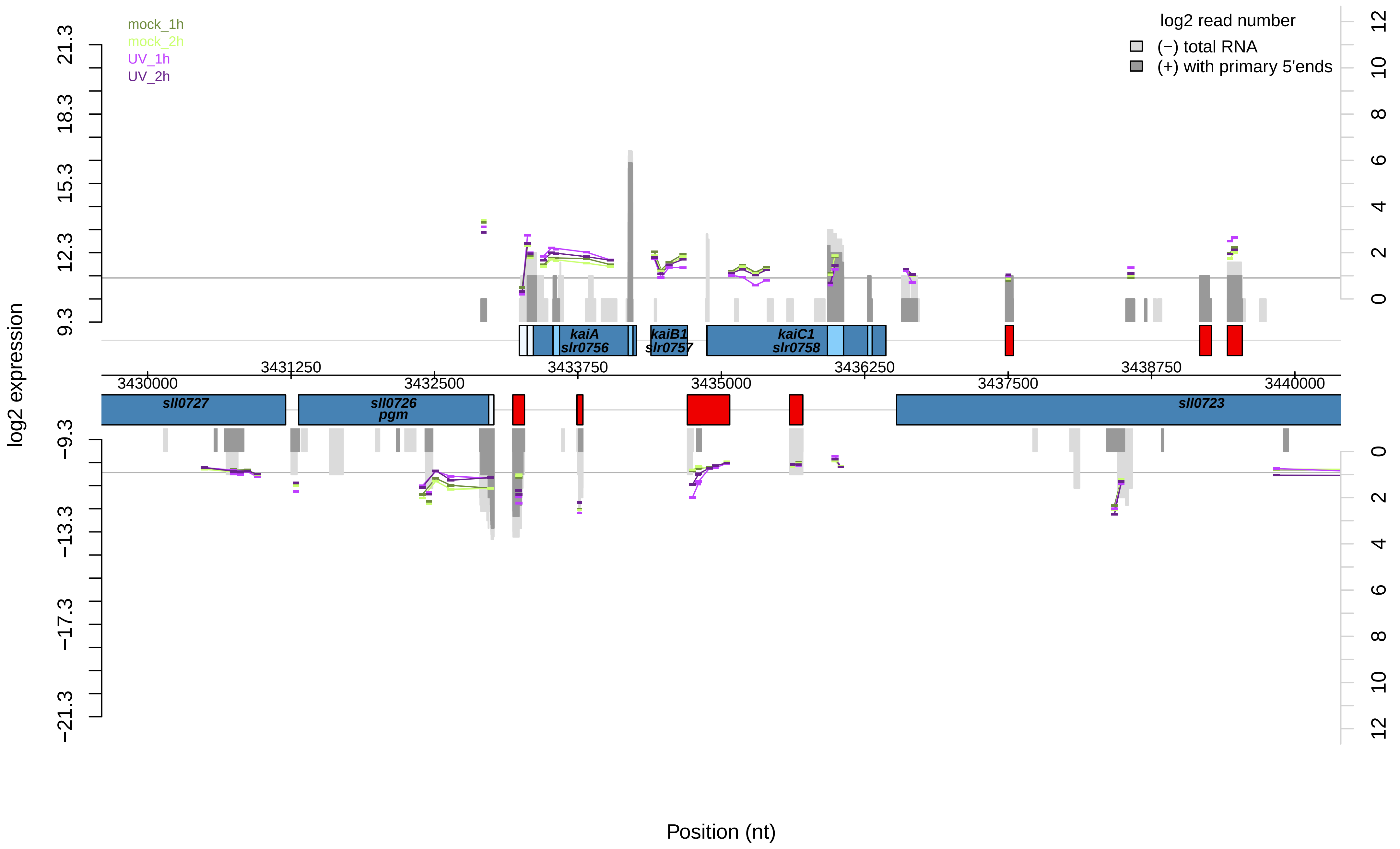

mock_1h
mock_2h
UV_1h
UV_2h
log2 read number
(−) total RNA
(+) with primary 5'ends
log2 expression
21.3
18.3
15.3
12.3
9.3
−9.3
−13.3
−17.3
−21.3
12
10
8
6
4
2
0
kaiA
slr0756
kaiB1
slr0757
kaiC1
slr0758
3431250
3433750
3436250
3438750
3430000
3432500
3435000
3437500
3440000
sll0727
sll0726
pgm
sll0723
Position (nt)

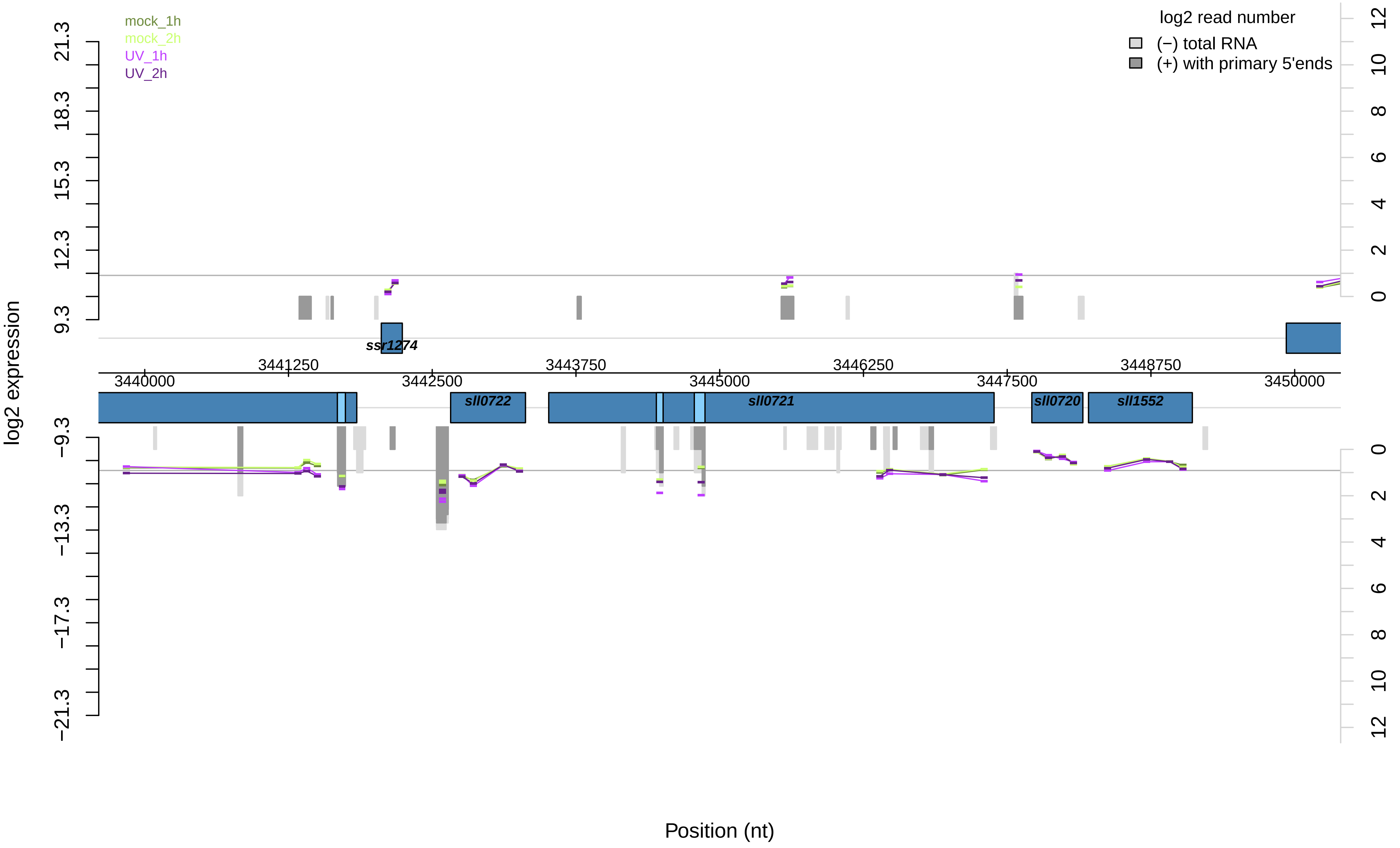

mock_1h
mock_2h
UV_1h
UV_2h
log2 read number
(−) total RNA
(+) with primary 5'ends
log2 expression
21.3
18.3
15.3
12.3
9.3
−9.3
−13.3
−17.3
−21.3
ssr1274
sll0722
sll0721
sll0720
sll1552
3440000
3441250
3442500
3443750
3445000
3446250
3447500
3448750
3450000
Position (nt)

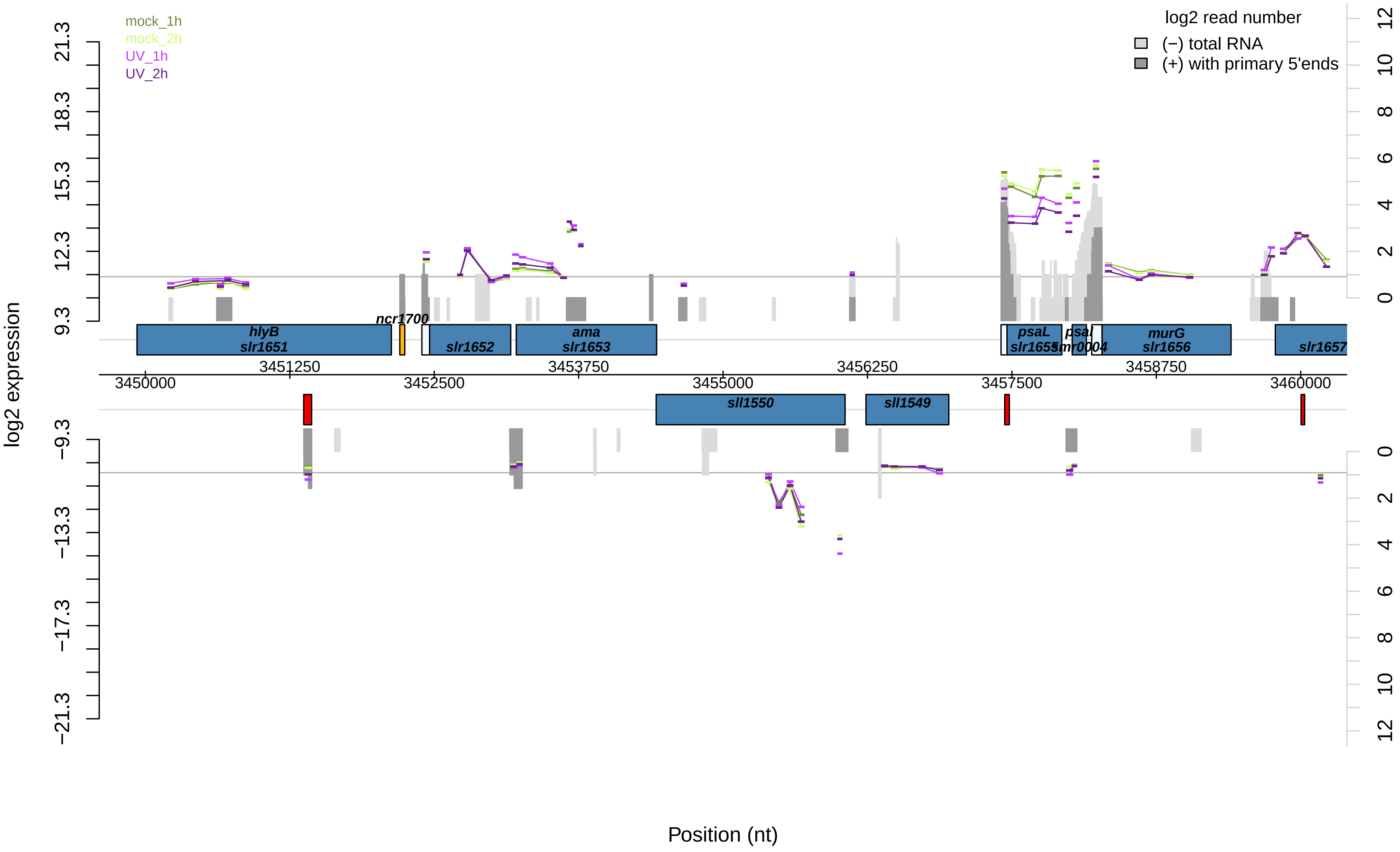
mock_1h
mock_2h
UV_1h
UV_2h
log2 read number
(−) total RNA
(+) with primary 5'ends
log2 expression
hlyB
slr1651
ncr1700
slr1652
ama
slr1653
psaL
slr1655
psaI
smr0004
murG
slr1656
slr1657
sll1550
sll1549
3450000
3451250
3452500
3453750
3455000
3456250
3457500
3458750
3460000
Position (nt)

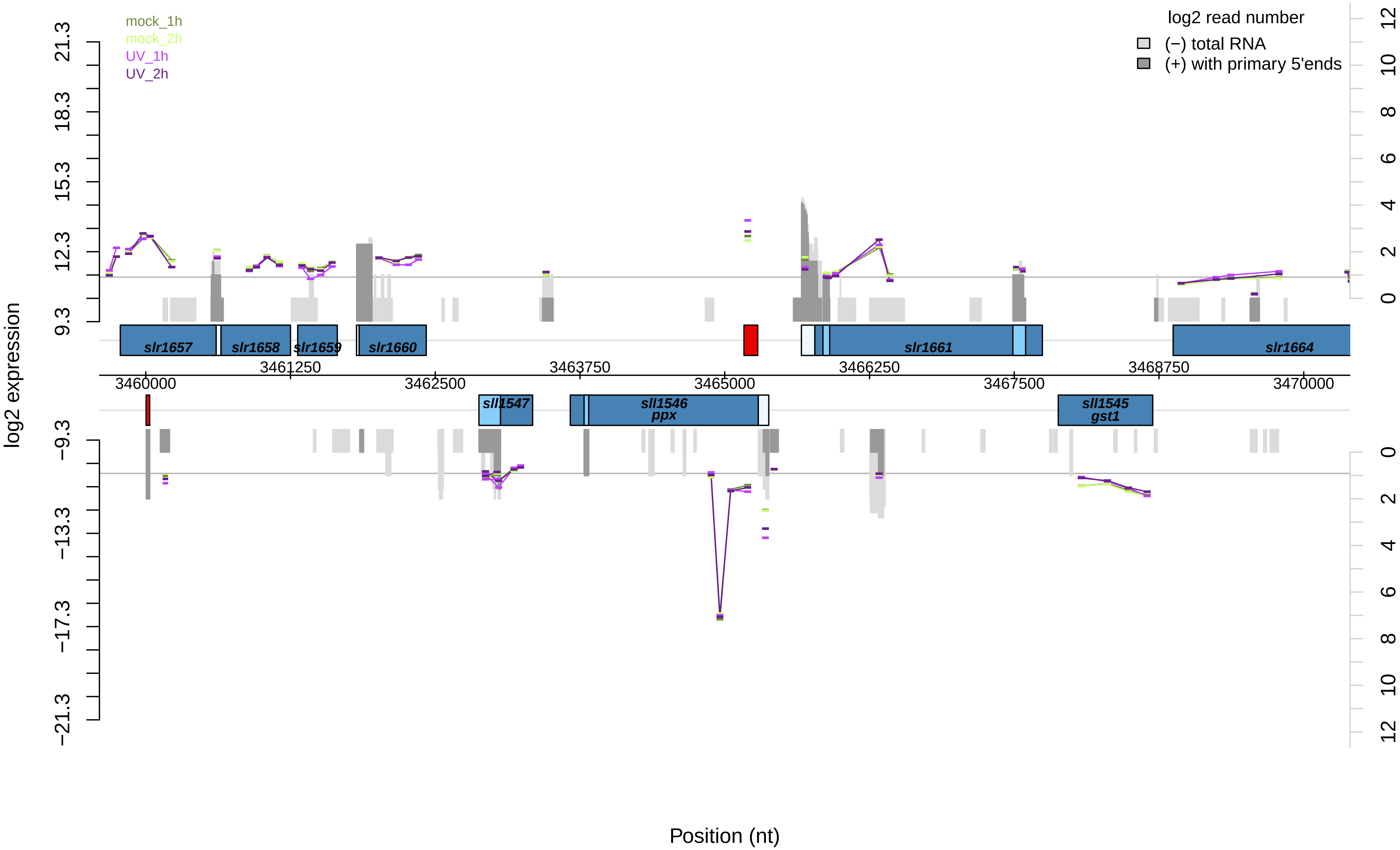

mock_1h
mock_2h
UV_1h
UV_2h
log2 read number
(−) total RNA
(+) with primary 5'ends
log2 expression
21.3
18.3
15.3
12.3
9.3
−9.3
−13.3
−17.3
−21.3
slr1657
slr1658
slr1659
slr1660
slr1661
slr1664
sll1547
sll1546
ppx
sll1545
gst1
3460000
3461250
3462500
3463750
3465000
3466250
3467500
3468750
3470000
Position (nt)

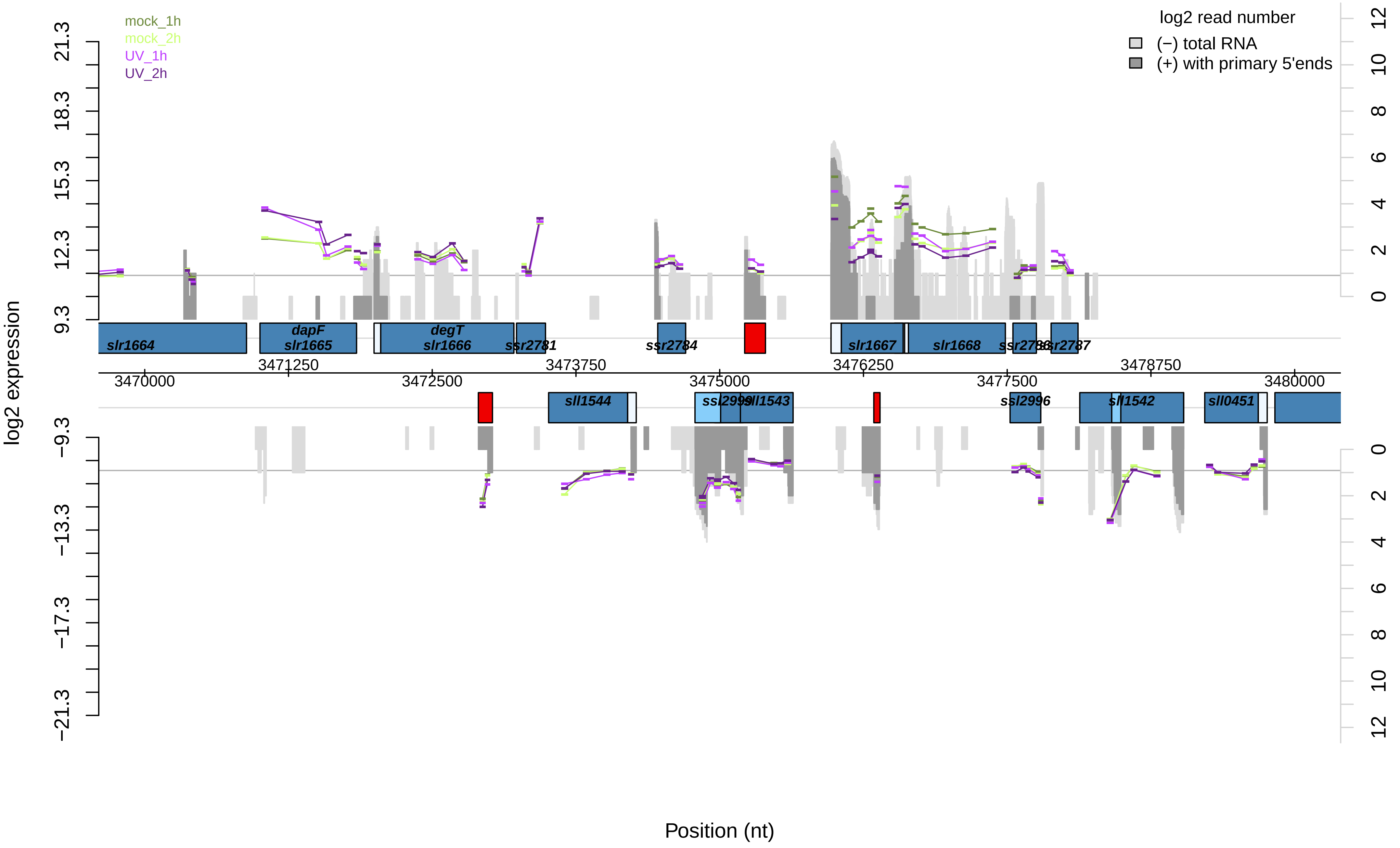

mock_1h
mock_2h
UV_1h
UV_2h
log2 read number
(−) total RNA
(+) with primary 5'ends
log2 expression
21.3
18.3
15.3
12.3
9.3
−9.3
−13.3
−17.3
−21.3
12
10
8
6
4
2
0
slr1664
dapF
slr1665
degT
slr1666
ssr2781
ssr2784
slr1667
slr1668
ssr2787
3470000
3471250
3472500
3473750
3475000
3476250
3477500
3478750
3480000
sll1544
sll1543
ssl2996
sll1542
sll0451
Position (nt)

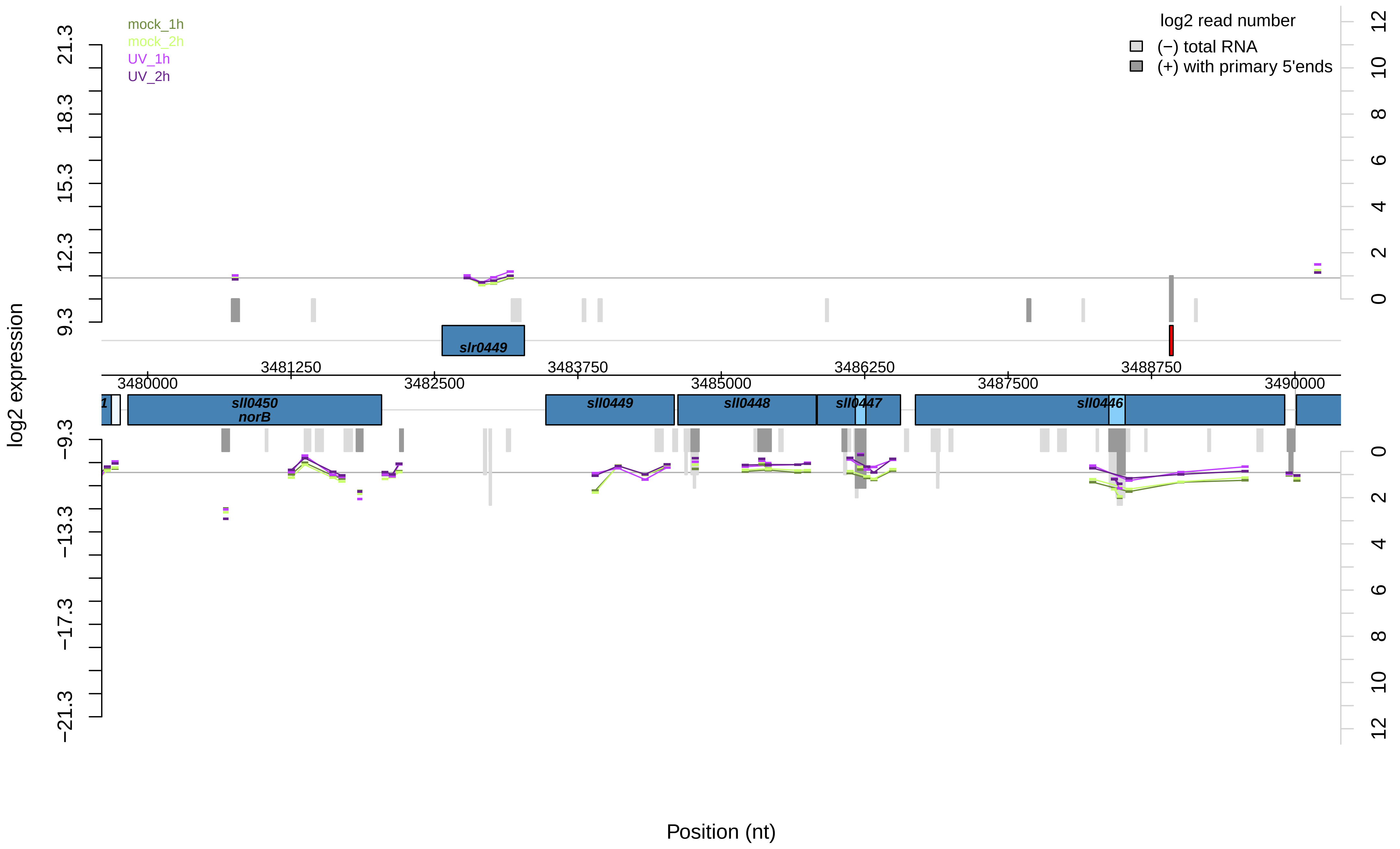

mock_1h
mock_2h
UV_1h
UV_2h
log2 read number
(−) total RNA
(+) with primary 5'ends
log2 expression
21.3
18.3
15.3
12.3
9.3
−9.3
−13.3
−17.3
−21.3
12
10
8
6
4
2
0
slr0449
3480000
3481250
3482500
3483750
3485000
3486250
3487500
3488750
3490000
sll0450
norB
sll0449
sll0448
sll0447
sll0446
Position (nt)

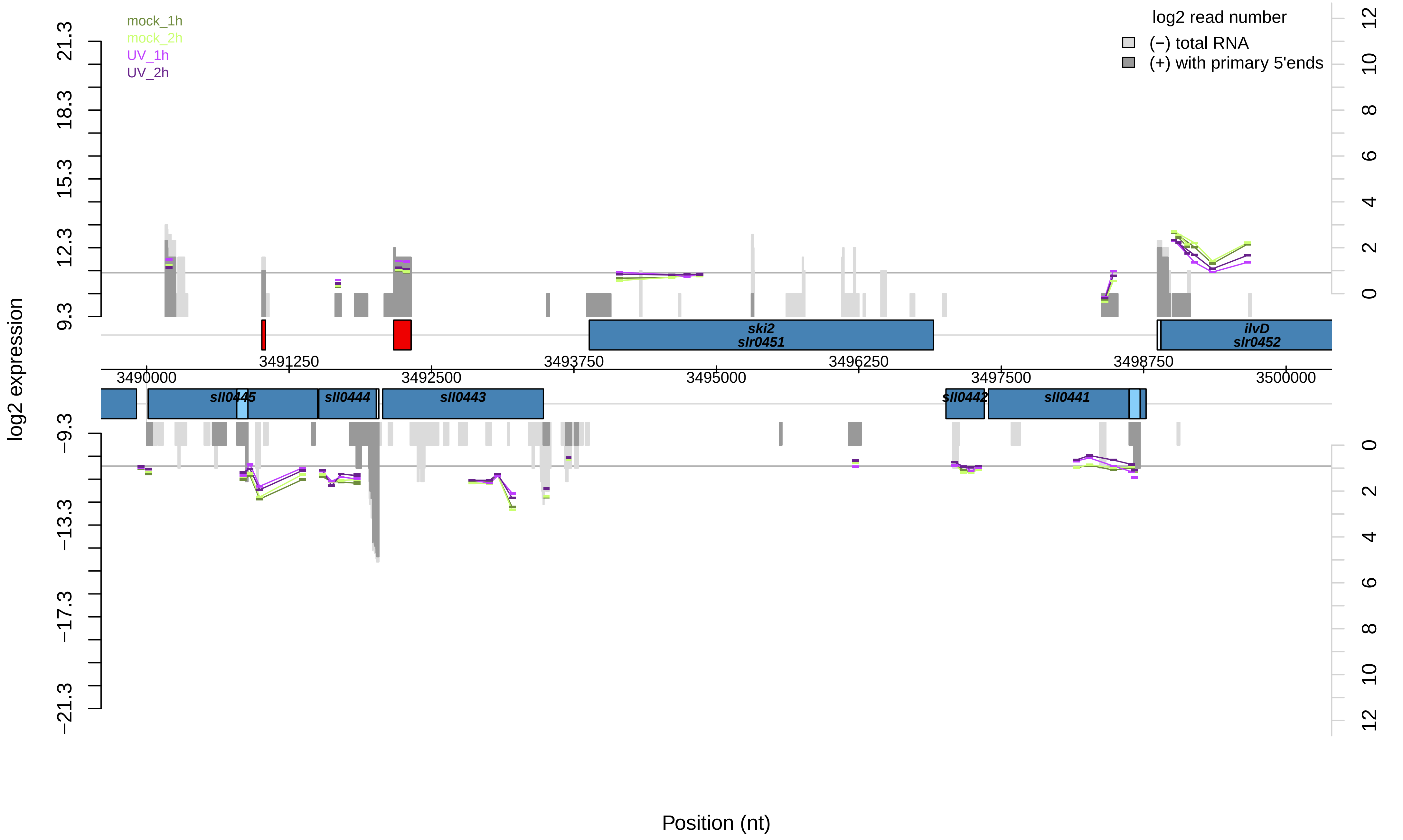

mock_1h
mock_2h
UV_1h
UV_2h
log2 read number
(−) total RNA
(+) with primary 5'ends
log2 expression
21.3
18.3
15.3
12.3
9.3
−9.3
−13.3
−17.3
−21.3
12
10
8
6
4
2
0
ski2
slr0451
ilvD
slr0452
3491250
3492500
3493750
3496250
3497500
3498750
3490000
3495000
3500000
sll0445
sll0444
sll0443
sll0442
sll0441
Position (nt)

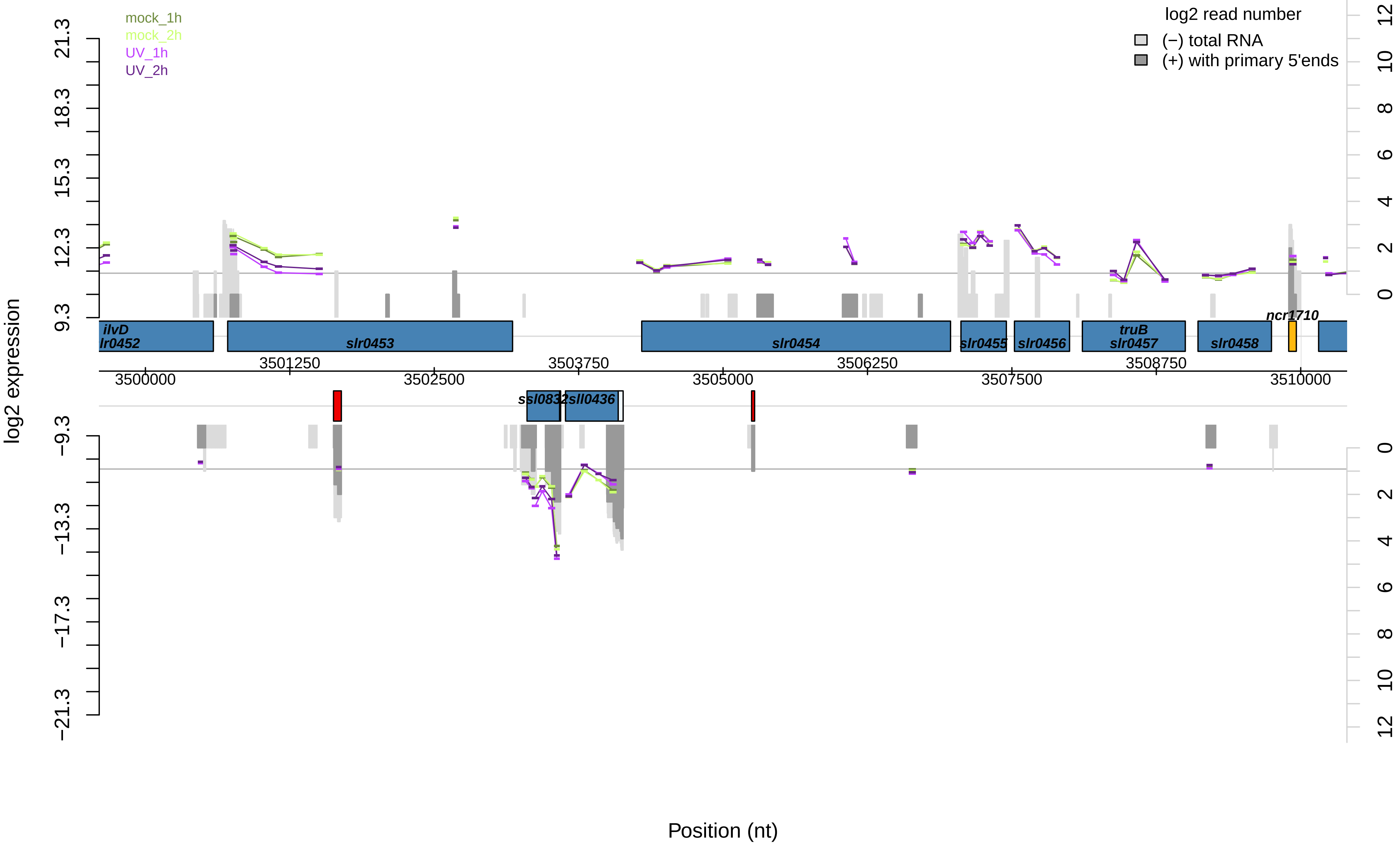

mock_1h
mock_2h
UV_1h
UV_2h
log2 read number
(−) total RNA
(+) with primary 5'ends
log2 expression
21.3
18.3
15.3
12.3
9.3
−9.3
−13.3
−17.3
−21.3
12
10
8
6
4
2
0
ilvD
lr0452
slr0453
slr0454
slr0455
slr0456
truB
slr0457
slr0458
ncr1710
ssl0832
sll0436
3500000
3501250
3502500
3503750
3505000
3506250
3507500
3508750
3510000
Position (nt)

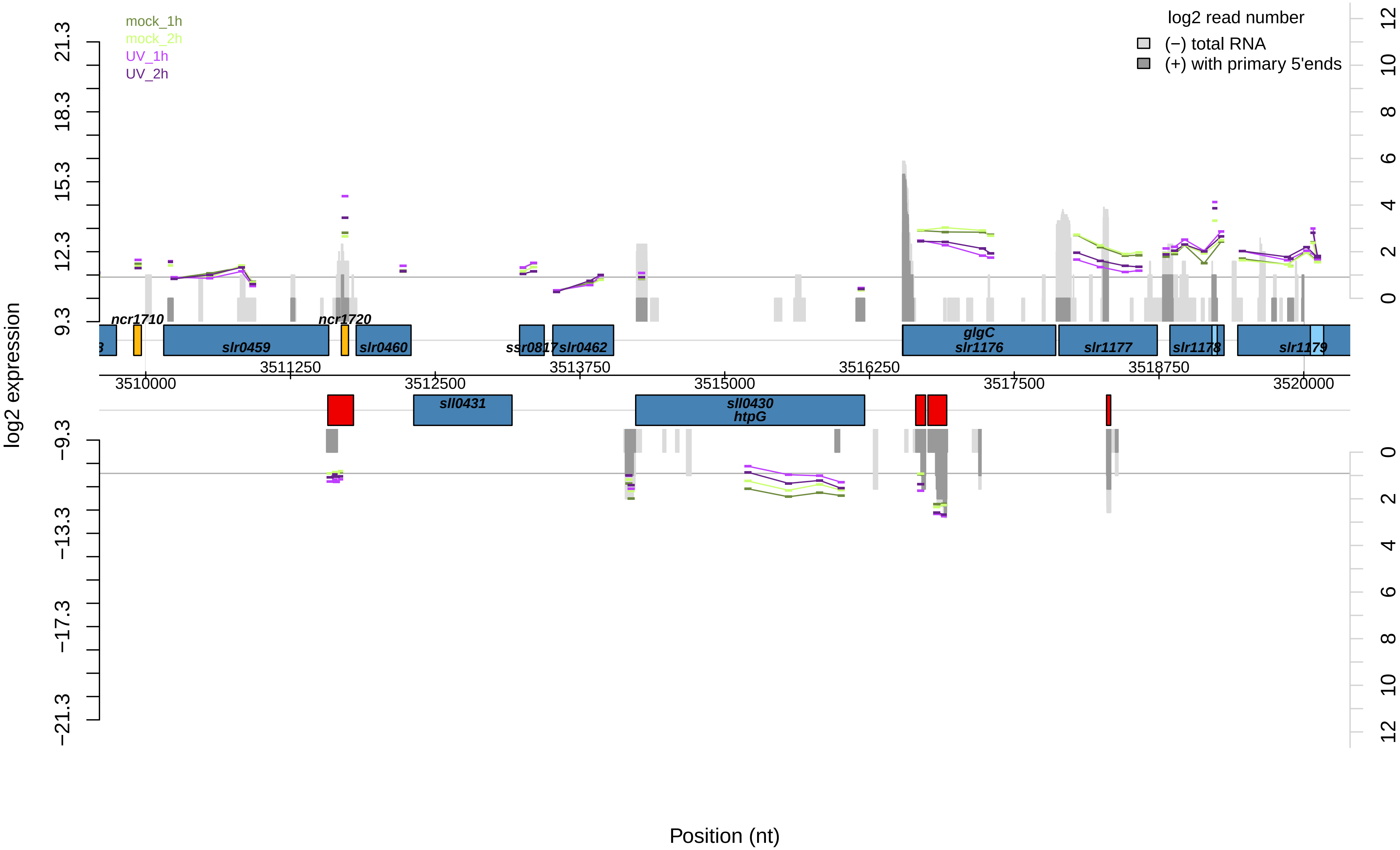

mock_1h
mock_2h
UV_1h
UV_2h
log2 read number
(−) total RNA
(+) with primary 5'ends
log2 expression
21.3
18.3
15.3
12.3
9.3
−9.3
−13.3
−17.3
−21.3
ncr1710
ncr1720
slr0459
slr0460
ssr0817
slr0462
glgC
slr1176
slr1177
slr1178
slr1179
sll0431
sll0430
htpG
3510000
3511250
3512500
3513750
3515000
3516250
3517500
3518750
3520000
Position (nt)

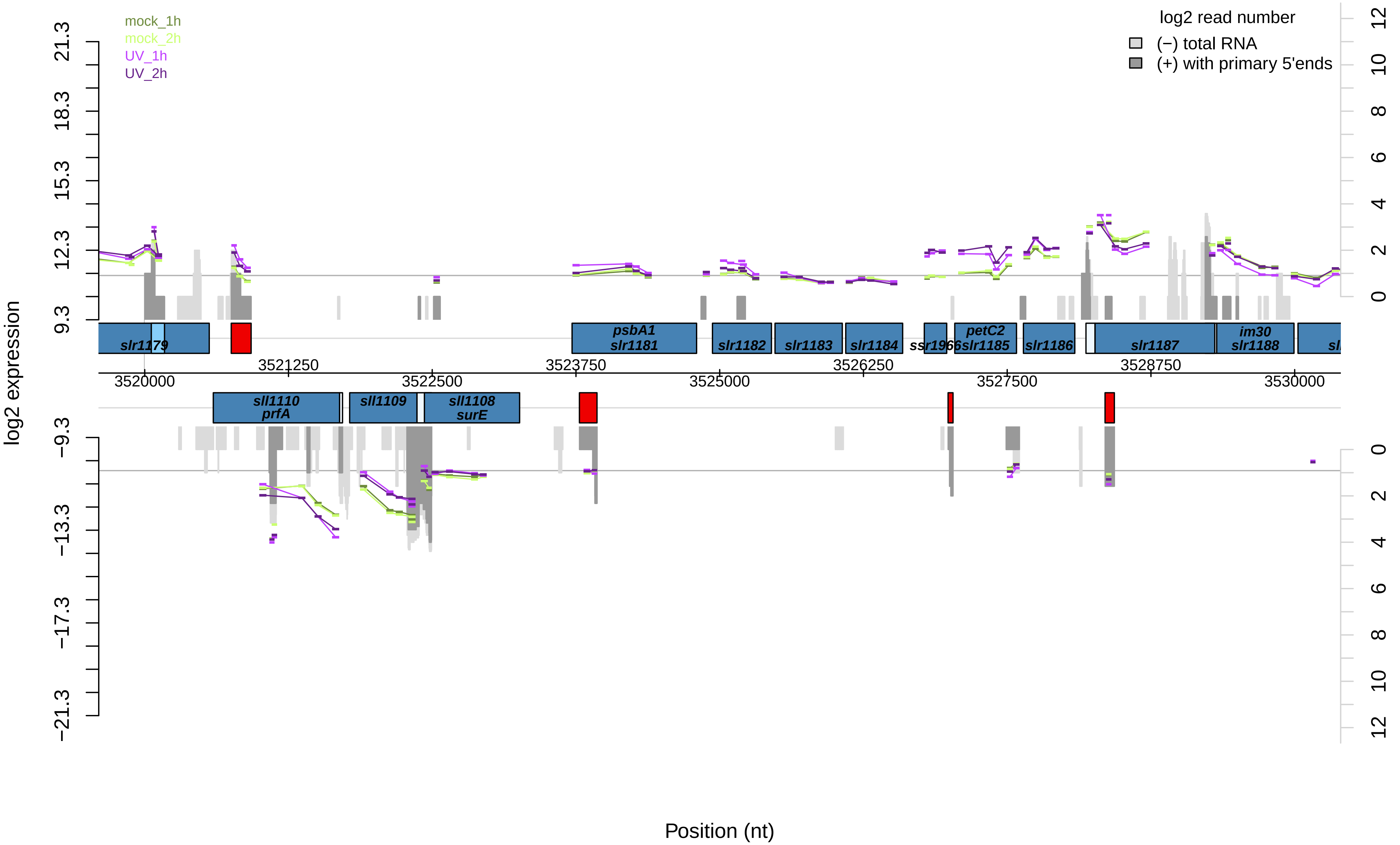
mock_1h
mock_2h
UV_1h
UV_2h
log2 read number
(−) total RNA
(+) with primary 5'ends
log2 expression
slr1179
psbA1
slr1181
slr1182
slr1183
slr1184
ssr1966
petC2
slr1185
slr1186
slr1187
im30
slr1188
sll1110
prfA
sll1109
sll1108
surE
3520000
3521250
3522500
3523750
3525000
3526250
3527500
3528750
3530000
Position (nt)

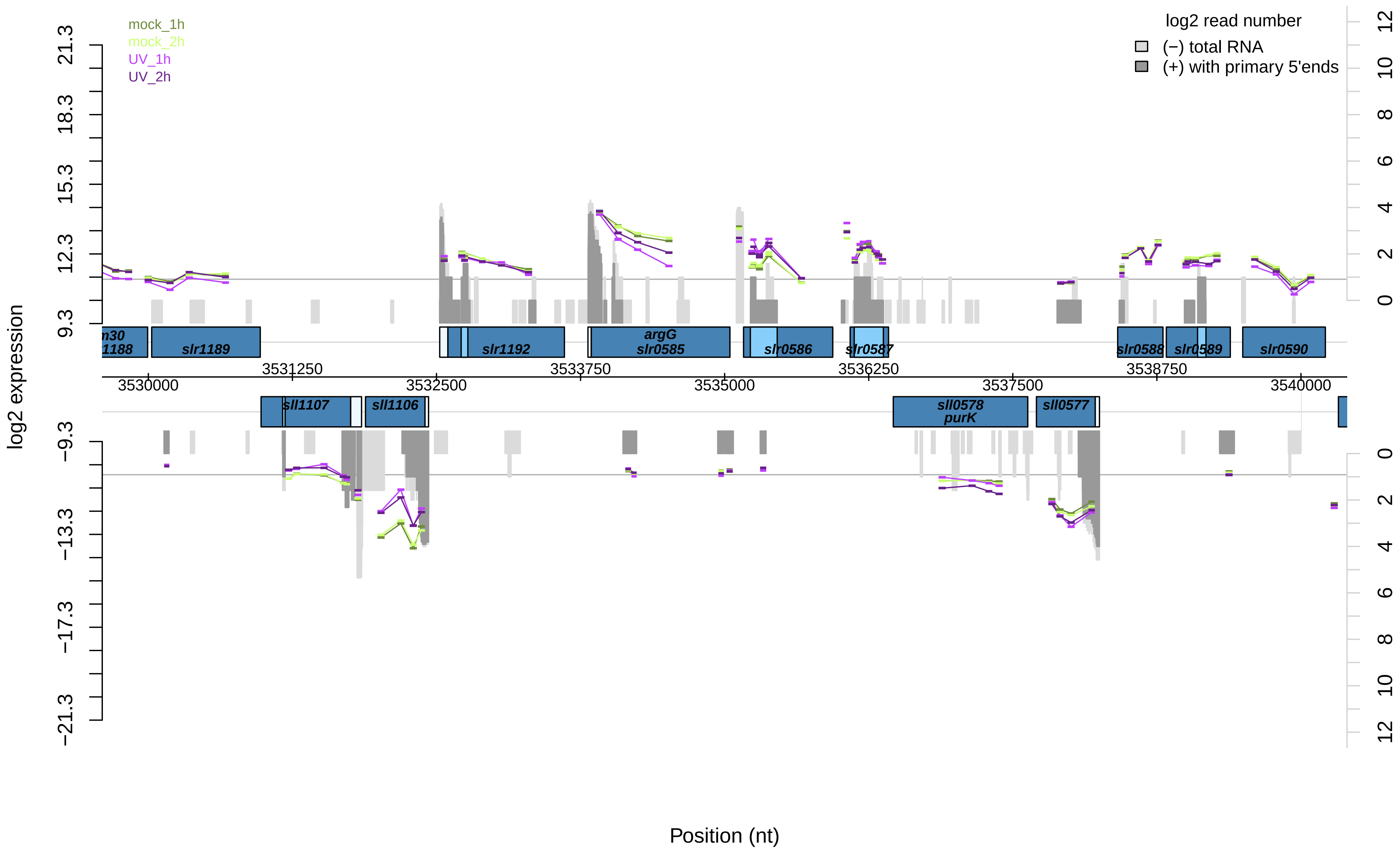

mock_1h
mock_2h
UV_1h
UV_2h
log2 read number
(−) total RNA
(+) with primary 5'ends
log2 expression
21.3
18.3
15.3
12.3
9.3
−9.3
−13.3
−17.3
−21.3
12
10
8
6
4
2
0
n30
1188
slr1189
slr1192
argG
slr0585
slr0586
slr0587
slr0588
slr0589
slr0590
3531250
3532500
3533750
3535000
3536250
3537500
3538750
3540000
3530000
sll1107
sll1106
sll0578
purK
sll0577
Position (nt)

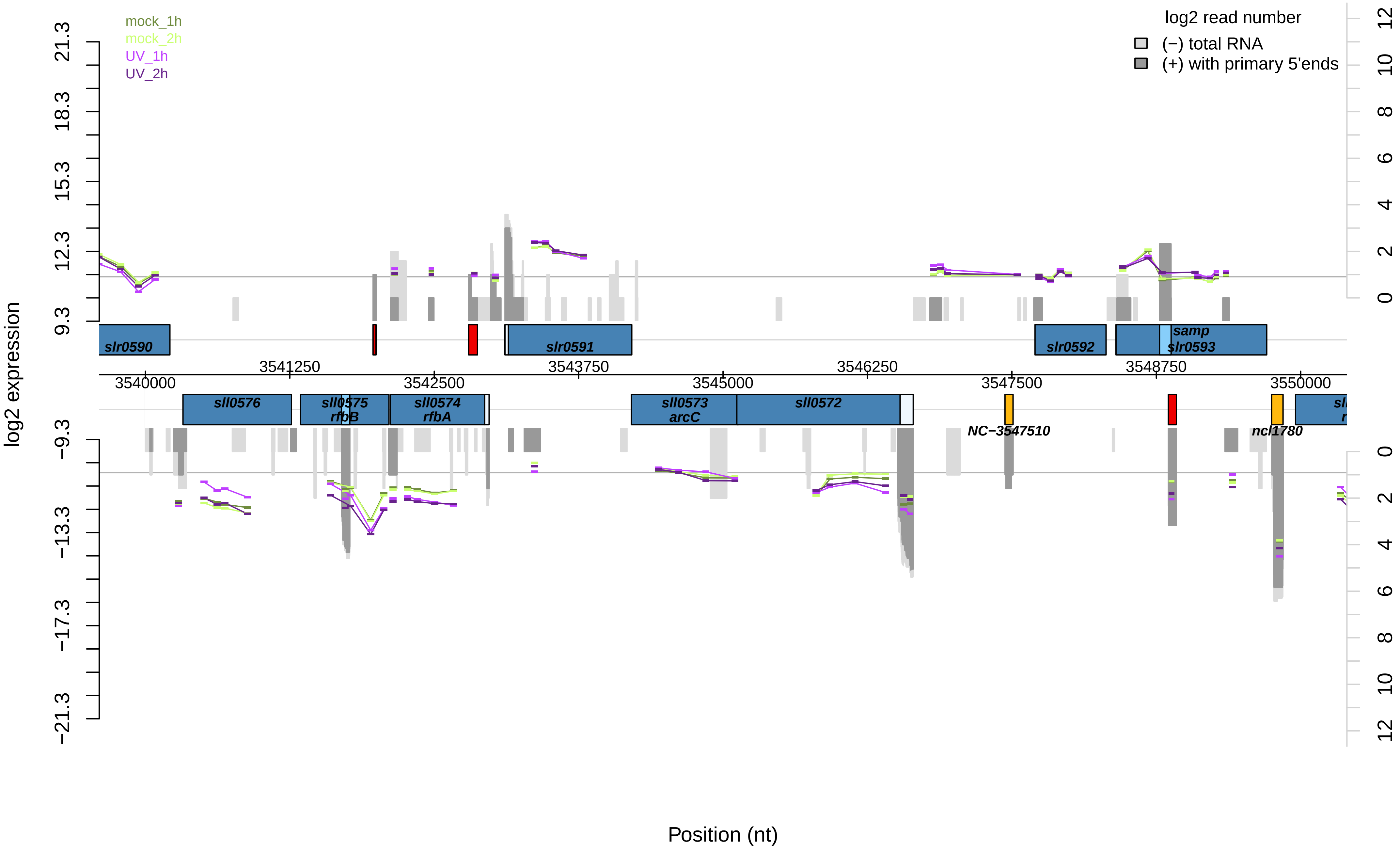

mock_1h
mock_2h
UV_1h
UV_2h
log2 read number
(−) total RNA
(+) with primary 5'ends
log2 expression
21.3
18.3
15.3
12.3
9.3
−9.3
−13.3
−17.3
−21.3
12
10
8
6
4
2
0
slr0590
slr0591
slr0592
samp
slr0593
3541250
3543750
3546250
3548750
3540000
3542500
3545000
3547500
3550000
sll0576
sll0575
rfbB
sll0574
rfbA
sll0573
arcC
sll0572
NC−3547510
ncl1780
Position (nt)

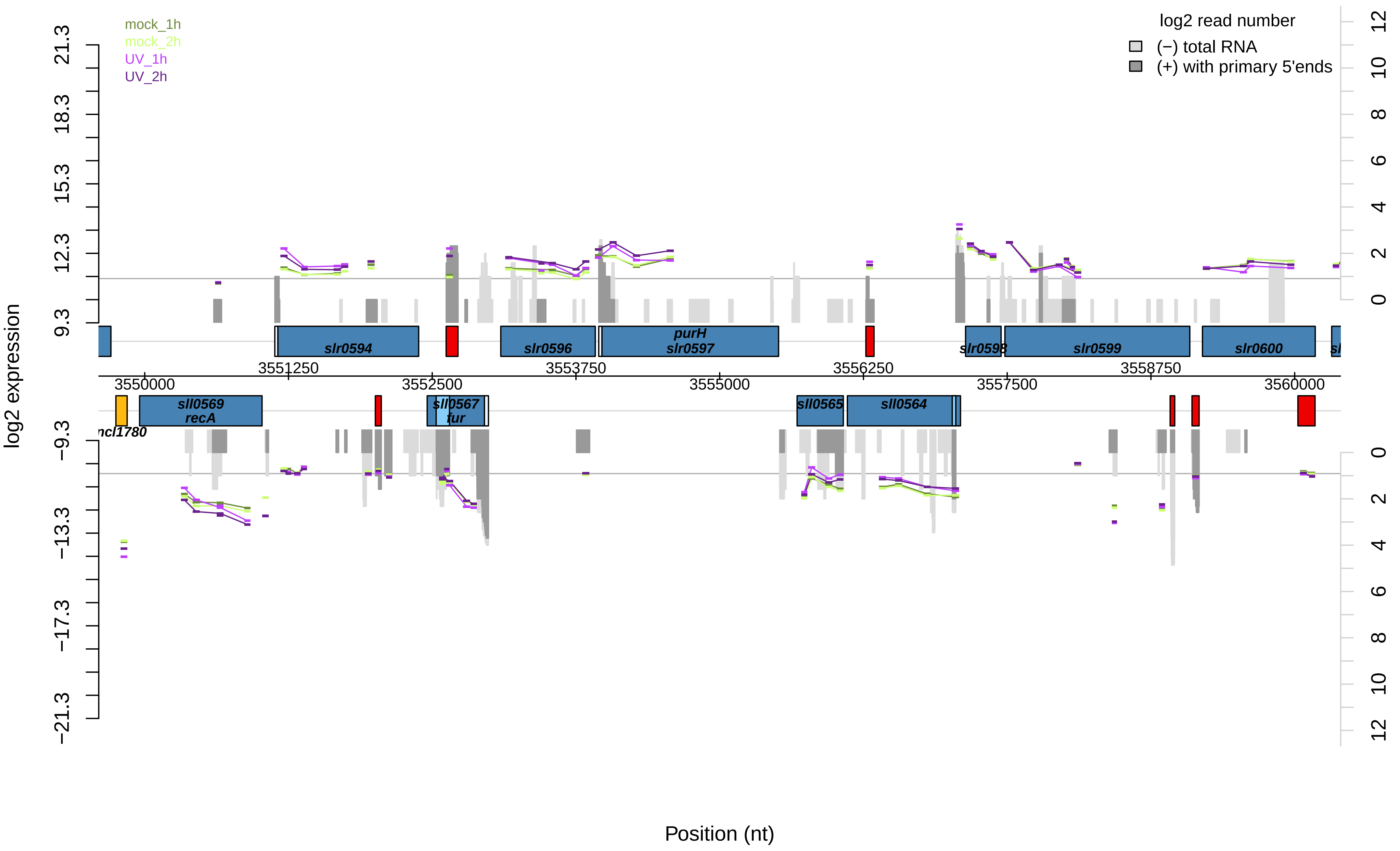

mock_1h
mock_2h
UV_1h
UV_2h
log2 read number
(−) total RNA
(+) with primary 5'ends
log2 expression
Position (nt)
slr0594
slr0596
purH
slr0597
slr0598
slr0599
slr0600
sll0569
recA
sll0567
fur
sll0565
sll0564
ncl1780
3550000
3551250
3552500
3553750
3555000
3556250
3557500
3558750
3560000

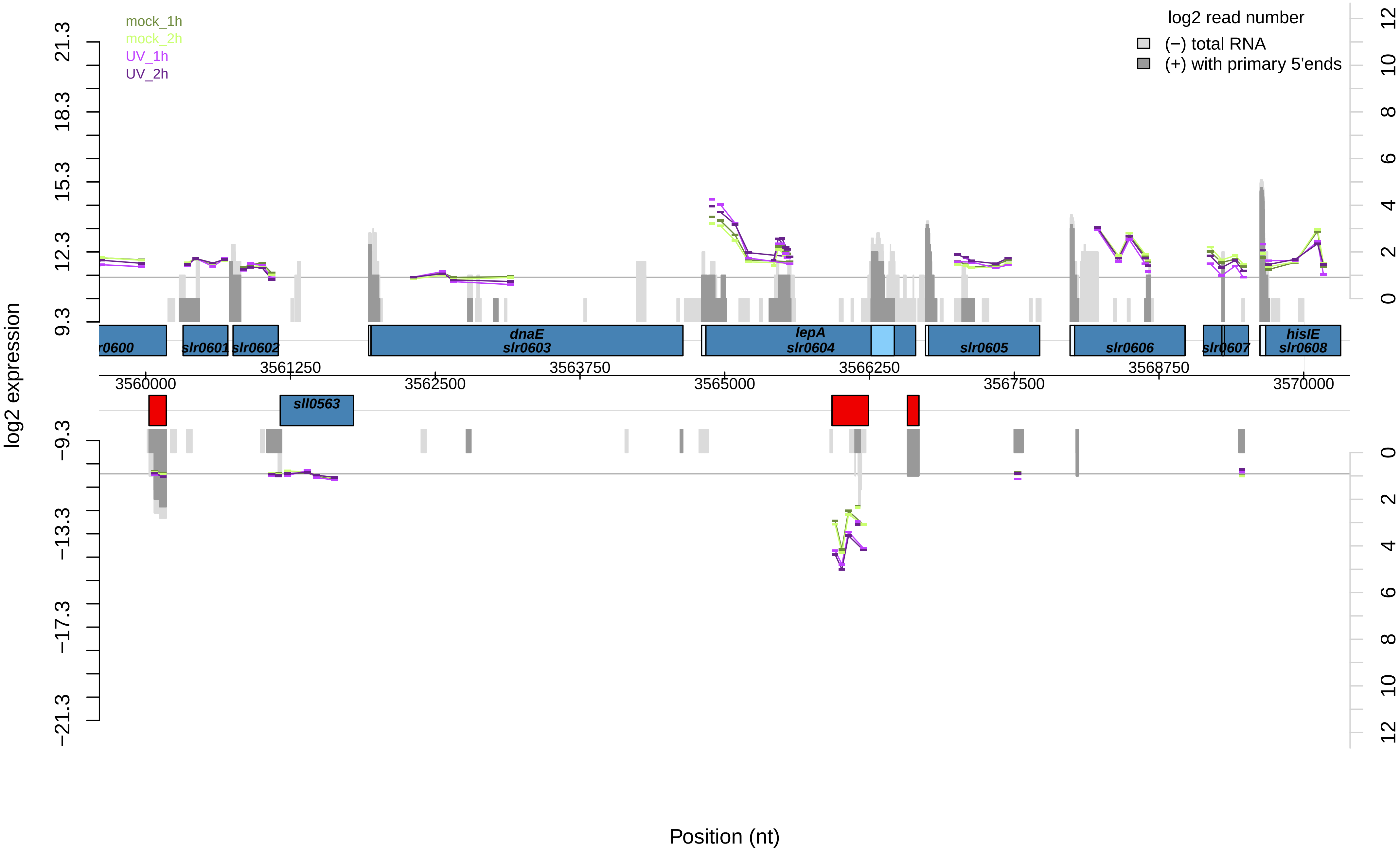

mock_1h
mock_2h
UV_1h
UV_2h
log2 read number
(−) total RNA
(+) with primary 5'ends
log2 expression
21.3
18.3
15.3
12.3
9.3
−9.3
−13.3
−17.3
−21.3
12
10
8
6
4
2
0
r0600
slr0601
slr0602
dnaE
slr0603
lepA
slr0604
slr0605
slr0606
slr0607
hisIE
slr0608
sll0563
3560000
3561250
3562500
3563750
3565000
3566250
3567500
3568750
3570000
Position (nt)

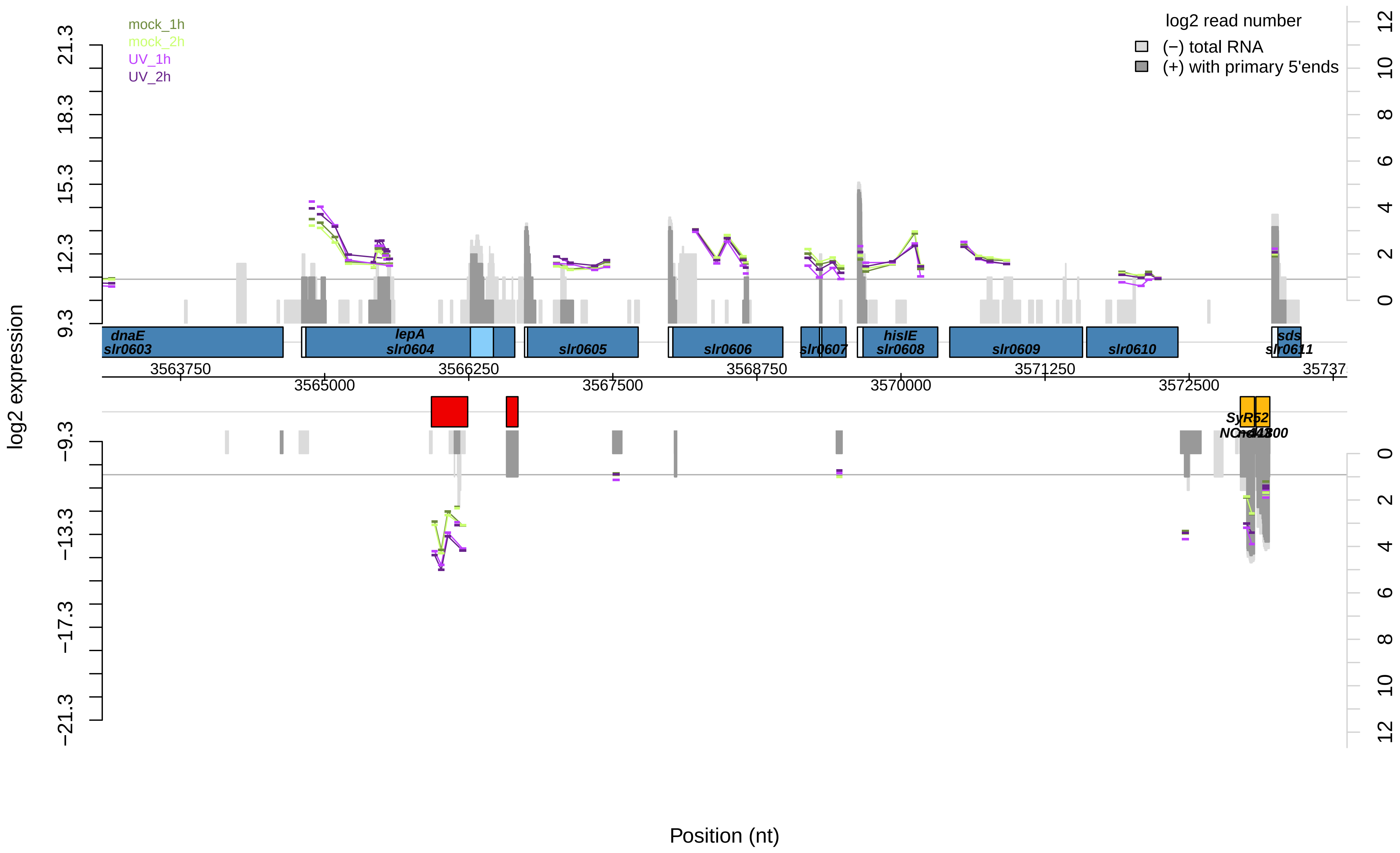

mock_1h
mock_2h
UV_1h
UV_2h
log2 read number
(−) total RNA
(+) with primary 5'ends
log2 expression
21.3
18.3
15.3
12.3
9.3
−9.3
−13.3
−17.3
−21.3
12
10
8
6
4
2
0
dnaE
slr0603
lepA
slr0604
slr0605
slr0606
slr0607
hisIE
slr0608
slr0609
slr0610
sds
slr0611
3563750
3565000
3566250
3567500
3568750
3570000
3571250
3572500
SyR52
Position (nt)
